# *Azotobacter vinelandii* AmrZ is a global regulator linking alginate production and c-di-GMP homeostasis

**DOI:** 10.1101/2025.10.20.683574

**Authors:** Miriam Citlalli Gonzaga-Pérez, Carlos Leonel Ahumada-Manuel, Ana Isabel Chávez-Martínez, Josefina Guzmán, Karel Estrada, Guadalupe Espín, Cinthia Núñez

## Abstract

*Azotobacter vinelandii*, a member of the Pseudomonadaceae, produces the exopolysaccharide alginate during vegetative growth; however, the circuitry linking alginate biosynthesis to lifestyle transitions remains poorly defined. Here we show that the Ribbon–Helix–Helix (RHH) transcription factor AmrZ coordinates alginate production, intracellular c-di-GMP levels, and motility. Deletion of *amrZ* abolished alginate synthesis, whereas chromosomal complementation restored it. A P*algD-gusA* fusion and RT-qPCR demonstrated that *algD*, the first gene in the alginate biosynthetic cluster, depends on AmrZ for expression. Motif analysis identified multiple AmrZ sites upstream of *algD*, and electrophoretic mobility-shift assays (EMSAs) confirmed specific binding to these regions. AmrZ also positively autoregulates: PamrZ-gusA activity decreased in Δ*amrZ*, and purified AmrZ bound the *amrZ* promoter in EMSA. Moreover, P*amrZ* activity required the sigma factor AlgU, consistent with the presence of an AlgU promoter; this positive, AlgU-dependent feedback may stabilize AmrZ under alginate-inducing conditions. To probe AmrZ control of c-di-GMP, we implemented a riboswitch-based biosensor in *A. vinelandii*. The Δ*amrZ* strain showed a markedly reduced signal, similar to a diguanylate cyclase (DGC) mutant; whereas a phosphodiesterase (PDE) mutant displayed elevated output, validating the assay. RNA-seq and RT-qPCR identified two DGC genes, AVAEIV_RS11610 and AVAEIV_RS18795, as AmrZ-activated targets; EMSA verified direct binding at the RS11610 regulatory region. By contrast, transcription of the principal vegetative DGC AvGreg was not AmrZ-regulated. Lower c-di-GMP in Δ*amrZ* correlated with larger swimming halos. Collectively, these genetic, biochemical, and transcriptomic data support a model in which AmrZ directly activates *algD* and elevates c-di-GMP via selected DGCs, thereby promoting alginate synthesis while reducing motility. RNA-seq data also indicate that AmrZ influences broader cellular programs, including metabolism and ion homeostasis, positioning AmrZ as a central regulator that links c-di-GMP homeostasis to coordinated exopolysaccharide production in *A. vinelandii*.

## INTRODUCTION

The free-living bacterium *Azotobacter vinelandii,* a member of the Pseudomonadaceae family, is strictly aerobic and motile by peritrichous flagella (Kennedy 2006). Vegetative cells undergo morphological and physiological differentiation to form cysts that are resistant to desiccation (Segura *et al*., 2020). Alginate, a linear exopolysaccharide composed of β-D-mannuronic and α-L-guluronic acid residues, is the major structural component of the cyst layers (Sadoff, 1975; Segura *et al*., 2020). This polymer is also synthesized in vegetative non-encysting cells of *A. vinelandii* (Galindo *et al*., 2007; Núñez *et al*., 2022). Because alginate has distinctive physicochemical properties that can be exploited as a bio-based material (Dhamecha *et al*., 2019; Moradali and Rehm, 2020), alginate production in *A. vinelandii* has been studied to better understand its genetic regulation, with the ultimate goal of optimizing microbial production (Galindo *et al*., 2007; Urtuvia *et al*., 2017; Núñez *et al*., 2022).

Except for *algC*, the genes required for alginate biosynthesis in *A. vinelandii* are organized in a 12-gene chromosomal cluster (*algD-alg8-alg44-K-J-G-X-L-I-V-F-A*) that encodes the proteins forming the periplasm-spanning biosynthetic machinery (Moradali *et al*., 2015; Urtuvia *et al*., 2017; Núñez *et al*., 2022). This cluster is headed by *algD*, whose transcription is finely tuned by several regulators, including the sigma factors AlgU and RpoS (Martínez-Salazar *et al*., 1996; Castañeda *et al*., 2000; Núñez *et al*., 2000; Castañeda *et al*., 2001), and by the c-di-GMP-responsive regulator FleQ, which binds the *algD* regulatory region to inhibit transcription (Barrios-Rafael *et al*., 2025). The GacS/GacA-Rsm system controls *algD* expression post-transcriptionally (Manzo *et al*., 2011). Alginate production is also post-translationally regulated: c-di-GMP binds the PilZ domain of the co-polymerase Alg44, thereby activating the alginate polymerase Alg8 (Merighi *et al*., 2007; Whitney *et al*., 2015; Moradali *et al*., 2017). Moreover, in *A. vinelandii*, c-di-GMP strongly influences alginate’s physicochemical properties by modulating both molecular mass and monomer composition (Ahumada-Manuel *et al*., 2017; Ahumada-Manuel *et al*., 2020; Martínez-Ortiz *et al*., 2020). Although alginate biosynthesis is largely conserved between *A. vinelandii* and *Pseudomonas aeruginosa*, there are notable regulatory differences that likely reflect the polymer’s distinct functions in these organisms. In *P. aeruginosa*, AlgR, AlgB, and AmrZ are required for transcriptional activation of the *algD* promoter (Ma *et al*., 1998; Leech *et al*., 2008; Jones *et al*., 2014; Kong *et al*., 2015). In *A. vinelandii*, however, AlgR and AlgB are not required for *algD* expression or alginate production (Núñez *et al*., 1999; Mærk *et al*., 2020). In a previous study, a screen of a transposon insertion library in *A. vinelandii* revealed that AmrZ is required for alginate production(Mærk *et al*., 2020).

AmrZ is a 108-amino-acid protein with a flexible N terminus, a central Ribbon–Helix–Helix (RHH) DNA-binding domain, and a C-terminal domain required for oligomerization (Waligora *et al*., 2010; Jr *et al*., 2012; Xu *et al*., 2016a). Initially described in *Pseudomonas* as a regulator of alginate and twitching motility (Baynham and Wozniak, 1996; Baynham *et al*., 2006; Tart *et al*., 2006), AmrZ is now recognized as a global regulator implicated in production of the Psl and Pel polysaccharides, iron acquisition, flagellar structure, type III and type VI secretion systems, swarming motility, and biofilm formation, among other processes (Jones *et al*., 2013; Jones *et al*., 2014; Allsopp *et al*., 2017; Falcone *et al*., 2018; Hou *et al*., 2019). Many of these pathways are also regulated by the second messenger c-di-GMP (Ha and O’Toole, 2015; Jenal *et al*., 2017). Indeed, AmrZ modulates c-di-GMP levels by regulating genes encoding diguanylate cyclases (DGCs) and phosphodiesterases (PDEs), which synthesize and degrade c-di-GMP, respectively (Jones *et al*., 2014; Muriel *et al*., 2018). In *P. aeruginosa*, *algD* transcription is activated by AmrZ (Baynham and Wozniak, 1996; Baynham *et al*., 1999); four AmrZ-binding sites have been identified, and their occupancy promotes formation of a higher-order DNA–AmrZ complex that activates *algD* (Xu *et al*., 2016a).

The aim of this study is to define the role of AmrZ in *A. vinelandii* physiology using genetic, biochemical, and transcriptomic analyses. We show that AmrZ is essential for alginate production because it is required for transcriptional activation of *algD*. Our results further indicate that AmrZ positively influences c-di-GMP levels by regulating specific DGCs and this control also affects swimming motility.

## METHODS

### Bacterial strains and growth conditions

Bacterial strains and plasmids used in this study are listed in S1 Table. The *A. vinelandii* AEIV wild-type strain and derivatives were grown diazotrophically in minimal Burk’s medium supplemented with sucrose (20 g L⁻¹). The composition of Burk’s medium has been described previously (Ahumada-Manuel *et al*., 2017). Cultures were grown at 30 °C with shaking at 200 rpm in 250-ml Erlenmeyer flasks containing 50 ml medium, for the times indicated. Cultures were inoculated with 400 µg of cells from 20-24 h liquid Burk’s-sucrose pre-cultures. The final antibiotic concentrations (µg ml⁻¹) used for *A. vinelandii* and *Escherichia coli*, respectively, were: tetracycline (Tc), 30 and 15; gentamicin (Gm), 1 and 10; spectinomycin (Sp), 100 and 100; and ampicillin (Ap), not used and 200.

### AEIV genome sequencing

Three library types were prepared. Short-read paired-end libraries (2 × 75 bp) were sequenced on an Illumina NextSeq 500, while long-read libraries were sequenced using a PacBio RS II and an Oxford Nanopore MinION. Raw Illumina reads were adapter- and quality-trimmed with Atropos v1.1.32 (Didion *et al*., 2017). PacBio reads together with the processed Illumina reads were used for de novo hybrid assembly with Unicycler v0.5.1(Wick *et al*., 2017) under default settings.

Following the initial assembly, *loci* showing ambiguities or breaks were manually curated. Nanopore reads were mapped to the draft assembly and examined using the visualization tool Tablet v1.21.02.08; problematic regions were then edited to restore contiguity and ensure accurate sequence representation. Assembly integrity was subsequently evaluated by remapping all three datasets (Illumina, PacBio, and Nanopore). The curated assembly was submitted to NCBI, where gene prediction and feature annotation were carried out using the Prokaryotic Genome Annotation Pipeline (PGAP v6.9) (Tatusova *et al*., 2016). Project metadata are available under BioProject accession number PRJNA809843 and BioSample sccession SAMN26203234. The complete assembled and annotated genome of *A. vinelandii* strain AEIV is available in GenBank under assembly accession GCA_030506185.2.

### Standard techniques

Oligonucleotides used for PCR (S2 Table) were designed from the *A. vinelandii* AEIV genome sequence (GenBank: CP092752.2). PCRs were performed with Phusion High-Fidelity DNA Polymerase (Thermo Fisher Scientific). Amplicons were verified by DNA (Sanger) sequencing.

### Analytical methods

β-Glucuronidase activity in *A. vinelandii* was measured as described (Chowdhury-Paul *et al*., 2023). Protein concentration was determined by the Lowry assay (Lowry *et al*., 1951). Alginate was quantified using the carbazole-based spectrophotometric assay for uronic acids (Knutson and Jeanes, 1968). All experiments were performed in biological triplicate (n=3), each with three technical replicates; results are reported as the mean of the biological replicates. Statistical significance was assessed with Student’s t-test (two-tailed; p = 0.05). Swimming motility was assayed as described previously (Ahumada-Manuel *et al*., 2020).

### Construction of *A. vinelandii* mutants

Details of mutant construction and plasmids are listed in S1 Table. Competent *A. vinelandii* cells were prepared as described, exploiting the naturally competent state under iron-limited conditions (Page and Sadoff, 1976; Ahumada-Manuel *et al*., 2017). Competent cells were transformed with 5 µg of linearized plasmid DNA carrying the desired mutation to promote double-crossover allelic exchange, and transformants were selected on the appropriate antibiotic.

### Identification of potential AmrZ binding sites

AmrZ recognition sequences from *P. aeruginosa* (Jones *et al*., 2014) were used to build a position-specific scoring matrix and motif logo with MEME (Multiple EM for Motif Elicitation) in “one occurrence per sequence” mode (Bailey and Elkan, 1994). The matrix was queried against *A. vinelandii* AEIV 5′-UTR sequences using FIMO (Find Individual Motif Occurrences) (Grant *et al*., 2011). 5′-UTR sequences were extracted with bedtools getfasta from the BEDTools toolkit (Quinlan and Hall, 2010), comprising 400 nt upstream and 50 nt downstream of the annotated start coordinate of each gene in the AEIV annotation.

### RNA isolation and RT-qPCR

Total RNA from *A. vinelandii* was extracted with TRIzol reagent (Invitrogen) following the manufacturer’s instructions. RNA integrity was verified by agarose gel electrophoresis, and concentrations were measured with a NanoDrop spectrophotometer. Residual genomic DNA was removed by treatment with RNase-free DNase I (Invitrogen) at 37 °C for 30 min, followed by heat inactivation at 65 °C for 10 min. First-strand cDNA was synthesized with the Maxima Reverse Transcriptase kit (Thermo Fisher Scientific) from 2 µg of total RNA using gene-specific reverse primers (S2 Table). cDNA was quantified and diluted to a working concentration of 50 ng µl⁻¹ with nuclease-free water. Quantitative real-time PCR was performed with Maxima SYBR Green/ROX qPCR Master Mix (Thermo Fisher Scientific) on a QuantStudio^TM^ 5 Real-Time PCR System (Applied Biosystems) in 96-well plates. Each 10 µl reaction contained 50 ng cDNA template (5 ng µl⁻¹), 1× Master Mix, and gene-specific primers (S2 Table). Cycling conditions were an initial denaturation, followed by 35 cycles of 95 °C for 15 s, 58 °C for 30 s, and 72 °C for 30 s. Primers specificity was assessed by agarose-gel analysis of conventional PCR products and by melt-curve analysis. The *recA* (AVAEIV_RS18860) or the *gyrA* (AVAEIV_RS08140) gene served as the internal reference for normalization. Relative expression was calculated using the 2^−ΔΔCt method with the wild-type strain as the calibrator (expression level = 1) (Livak and Schmittgen, 2001). Experiments were performed with three biological replicates, each measured in triplicate.

### Expression and purification of His-AmrZ

AmrZ was produced as an N-terminal His₆-tagged protein (His-AmrZ). A single colony of *E. coli* BL21(DE3) carrying pET-AmrZ was inoculated into 5 ml LB with kanamycin (Km) and grown overnight at 37 °C, 200 rpm. One milliliter of this culture was used to inoculate 250 ml LB (Km); cells were grown ∼3 h to OD₆₀₀ ≈ 0.6, induced with 1 mM IPTG, and incubated for an additional 3 h. Cells were harvested (4,000 × g, 10 min, 4 °C), resuspended in lysis buffer (50 mM NaH₂PO₄, 300 mM NaCl, 10 mM imidazole, pH 7.0), and sonicated on ice (3 × 15 s). The lysate was clarified (4,000 × g, 20 min, 4 °C) and applied to Ni-NTA resin (Thermo Fisher Scientific) pre-equilibrated in lysis buffer; binding was carried out for 30 min at 4 °C with gentle mixing. The column was washed with wash I (50 mM NaH₂PO₄, 300 mM NaCl, 50 mM imidazole, pH 7.0) and wash II (same salts, 250 mM imidazole, pH 7.0), and protein was eluted with elution buffer (same salts, 500 mM imidazole, pH 7.0). The eluate was concentrated using Amicon Ultra-0.5, 10-kDa MWCO devices. Protein concentration was determined by the Lowry assay (Lowry *et al*., 1951). Purity and apparent molecular mass (∼15 kDa) were assessed by 10% SDS–PAGE.

### EMSA

Electrophoretic mobility shift assays were performed using a non-radioactive protocol. Four PCR fragments spanning the *algD* regulatory region were used: (a) 679 bp (AmrZ sites S1–S5), primers palgD-EMSA-F1.21 / palgD-R; (b) 204 bp (S1, S2), palgD-EMSA-F1.21 / palgD-EMSA-R1; (c) 194 bp (S3, S4), palgD-EMSA-F2 / palgD-EMSA-R2; (d) 206 bp (S5), palgD-EMSA-F3 / palgD-R. For testing binding to the *amrZ* regulatory region, pAmrZ-XbaF /pAmrZ-EcoR-Rv amplified P*amrZ*.

Binding reactions (20 µl) contained 100 ng DNA, increasing concentrations of His-AmrZ, and binding buffer (10 mM Tris-HCl, pH 8.0; 50 mM KCl; 1 mM DTT; 0.5 mM EDTA; 5% glycerol; 10 µg ml⁻¹ BSA). After 20 min at room temperature, samples were resolved on 6% nondenaturing polyacrylamide gels in TBE (45.5 mM Tris base, 45.5 mM boric acid, 1 mM EDTA, pH 8.3) and visualized by ethidium bromide staining under UV light.

### Construction of a c-di-GMP biosensor suitable for *A. vinelandii*

A 2,228-bp fragment was PCR-amplified from plasmid pFY4357 using primers pMMB-Rv and pMMBdual-secF. The amplicon carries a c-di-GMP biosensor consisting of a tandem dual riboswitch (Bc4–5) from *Bacillus thuringiensis* and the TurboRFP reporter gene, whose expression is repressed in the absence of c-di-GMP (Zhou *et al*., 2016). The PCR product was cloned into pJET1.2/Blunt, yielding pJB2. A BamHI fragment (1,103 bp) was then excised and ligated into pLA65 (BamHI-digested), downstream of a constitutive σ^70^ promoter, to generate pLA66. The entire cassette, including the promoter, was subsequently excised with BglII and cloned into pUMA-Km(5′–3′), producing pLA68. This vector, pre-linearized with ScaI, enables chromosomal integration of the biosensor into a neutral *locus* (melA) of *A. vinelandii* via double homologous recombination.

### c-di-GMP quantification

Biosensor strains. The AEIV wild type, Δ*mucG,* Δ*avGreg, and* Δ*amrZ* strains were transformed with plasmid pLA68, yielding strains CLAM36, CLAM37, MG07, and MG08, respectively. These strains carry a chromosomally integrated biosensor for intracellular c-di-GMP. CLAM37 (Δ*mucG*) and MG07 (Δ*avGreg*) served as controls because they exhibit increased and decreased c-di-GMP levels, respectively, when compared to the wild-type background (CLAM36).

Fluorescence readout. For c-di-GMP quantification, cultures were sampled over a time course (12, 18, 24, and 48 h). At each time point, 1 ml of culture was centrifuged, the pellet was resuspended in 1 ml water, and cells were washed twice with 10 mM MgSO₄ to remove alginate. Aliquots (200 µl) were transferred to black 96-well plates, and fluorescence was recorded (Ex 550 nm/Em 580 nm) on an Agilent BioTek Synergy H1 multimode microplate reader. Fluorescence values were normalized to total protein content.

### RNA-seq

The *A. vinelandii* AEIV wild-type strain and its Δ*amrZ* derivative were grown diazotrophically in Burk’s-sucrose (BS) medium for 18 h. Aliquots of these cultures (corresponding to 400 µg of protein) were used to inoculate 50 ml fresh BS medium and incubated for 24 h at 30 °C. Cells were harvested by centrifugation (4,000 × g, 10 min, 4 °C). Total RNA was extracted with the RiboPure RNA Purification Kit (Thermo Fisher Scientific) according to the manufacturer’s instructions. For each strain (wild type and Δ*amrZ*), RNA was isolated from three independent cultures (biological replicates). RNA quality was assessed by capillary electrophoresis on a 2100 Bioanalyzer (Agilent Technologies). Ribosomal RNA was depleted using the RiboMinus Bacteria 2.0 Transcriptome Isolation Kit (Thermo Fisher Scientific).

### Library preparation and sequencing

Libraries were prepared following the TruSeq Stranded mRNA Sample Preparation Guide (Illumina). cDNA libraries were sequenced as 2 × 75 bp paired-end reads on an Illumina NextSeq 500 using the NextSeq 500/550 High Output Kit v2.5 (150 cycles). A total yield of 43,253,228 paired-end reads was obtained for the AEIV wild-type strain and 30,955,718 paired-end reads for the Δ*amrZ* derivative.

### Processing of RNA-seq data

Quality control and read mapping. Read quality was assessed with FastQC v0.10.0 (Babraham Bioinformatics; https://www.bioinformatics.babraham.ac.uk/projects/fastqc/). All libraries showed high per-base Phred scores (>Q30), with no evidence of adapter contamination. Reads were aligned to the *A. vinelandii* AEIV reference genome (GenBank assembly GCA_030506185.2) using SMALT v0.7.6 (https://www.sanger.ac.uk/science/tools/smalt-0). Feature-level coverages/counts were generated with BEDTools v2.4.0 (coverageBed), yielding a count table for downstream analysis (Quinlan & Hall, 2010; Barnett *et al*., 2011).

### Differential gene expression analysis

Counts per million (CPM) were calculated using the cpm function in edgeR v3.12.1 (Robinson *et al*., 2010). Genes were retained if they had ≥1 count in at least three samples. Differential expression was assessed with DESeq2 v1.10.1 using the filtered count matrix and default parameters. *P* values were adjusted by the Benjamini-Hochberg procedure. Genes with log₂ fold change > 2 or < −2 and adjusted *P* (Padj) < 0.01 were considered differentially expressed and classified as AmrZ-responsive.

## RESULTS

### The transcriptional regulator AmrZ is required for alginate production

To assess the role of AmrZ (AVAEIV_RS17290) in *A. vinelandii* alginate production, we quantified polysaccharide levels in the Δ*amrZ* mutant and compared them with the wild-type strain. After 48 h in Burk’s-sucrose medium, alginate production was abolished in Δ*amrZ*: values were at or below the limit of detection and comparable to those of the alginate-negative Δ*algU* mutant (Moreno *et al*., 1998) (Fig 1A). Genetic complementation with a chromosomally integrated wild-type *amrZ* allele restored alginate synthesis, indicating that the AmrZ DNA-binding protein is required for alginate production.

**Fig 1.**
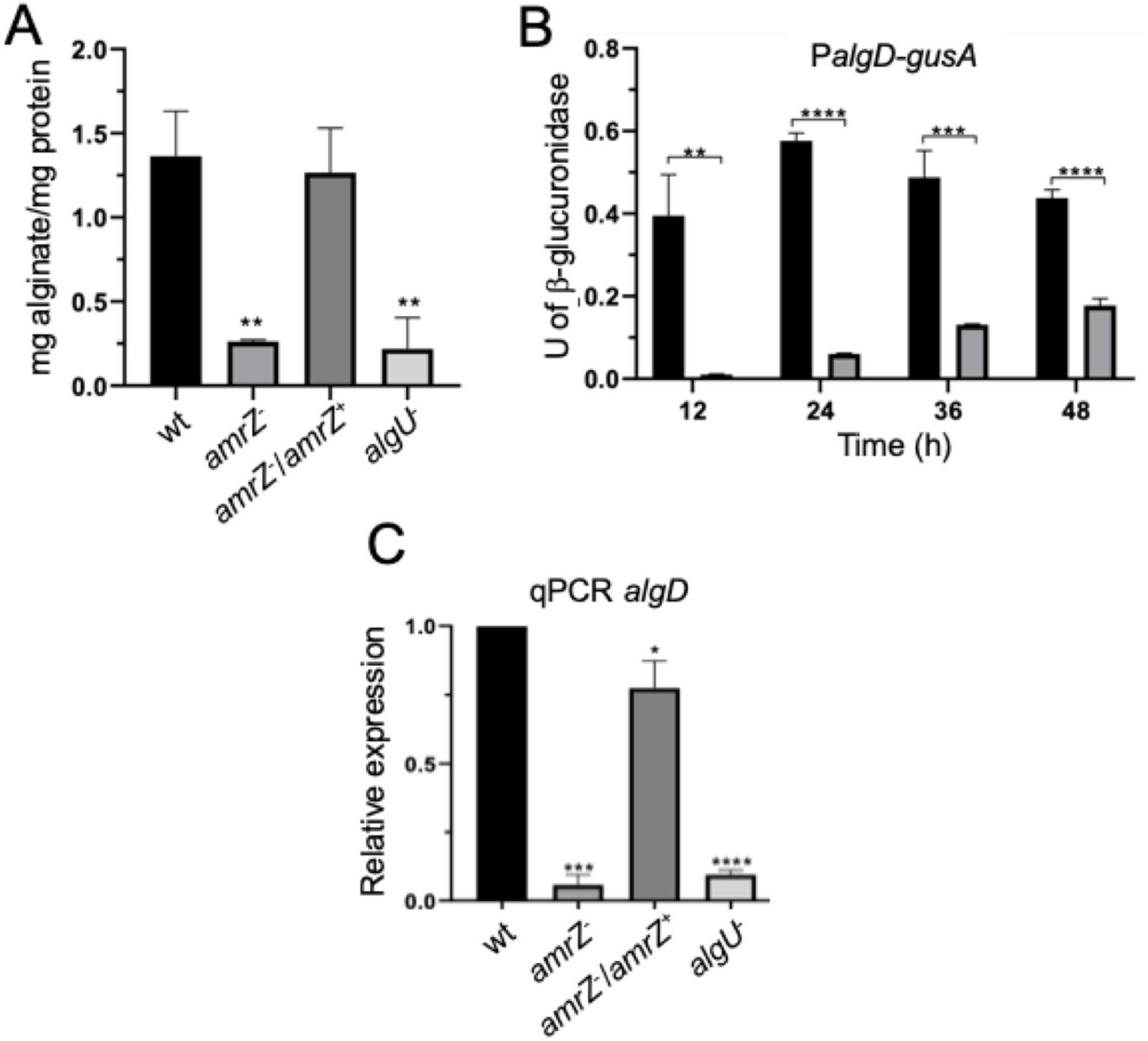
AmrZ is necessary for alginate production and *algD* expression. (A) Alginate quantification for cultures grown 48 h in Burk’s–sucrose medium. (B) β-Glucuronidase activity from an *algD-gusA* transcriptional fusion in the wild type (black) and Δ*amrZ* (gray). Specific activity is shown in U mg⁻¹ protein. (C) Relative *algD* mRNA levels in cultures grown 24 h in Burk’s-sucrose medium (RT-qPCR). Bars denote mean ± SD from three independent experiments. Statistical significance (two-tailed Student’s t-test) is indicated: **, *P* < 0.01; ***, *P* < 0.001; ****, *P* < 0.0001.

### Effect of AmrZ on *algD* expression

To assess the role of AmrZ in *algD* transcription, we introduced the Δ*amrZ* mutation into strain AED-gusA, which carries an *algD-gusA* transcriptional fusion, generating strain JG522. In the absence of AmrZ, *algD* transcription was strongly reduced relative to the wild type at early time points (12 and 24 h). By 48 h, reporter activity in Δ*amrZ* increased slightly, reaching ∼30% of the wild-type level (Fig 1B). Consistent with these data, *algD* mRNA measured by RT-qPCR was nearly undetectable in Δ*amrZ* at 24 h and comparable to the Δ*algU* mutant. As expected, *algD* mRNA levels were restored in the complemented strain (Δ*amrZ*/*amrZ*⁺), confirming that AmrZ is essential for *algD* transcriptional activation (Fig 1C).

### AmrZ regulates *algD* by directly binding its regulatory region

A MEME/FIMO search (Materials and Methods) was performed to identify potential AmrZ targets in the *A. vinelandii* AEIV genome (S3 Table). In *P. aeruginosa*, AmrZ directly binds the *algD* regulatory region to activate transcription (Xu *et al*., 2016b). Consistent with a conserved mechanism in *A. vinelandii*, five candidate AmrZ-binding sites (BS1– BS5) were identified at −565, −548, −426, −388, and −62 relative to the *algD* start codon (Fig 2A; S1 Fig).

**Fig 2.**
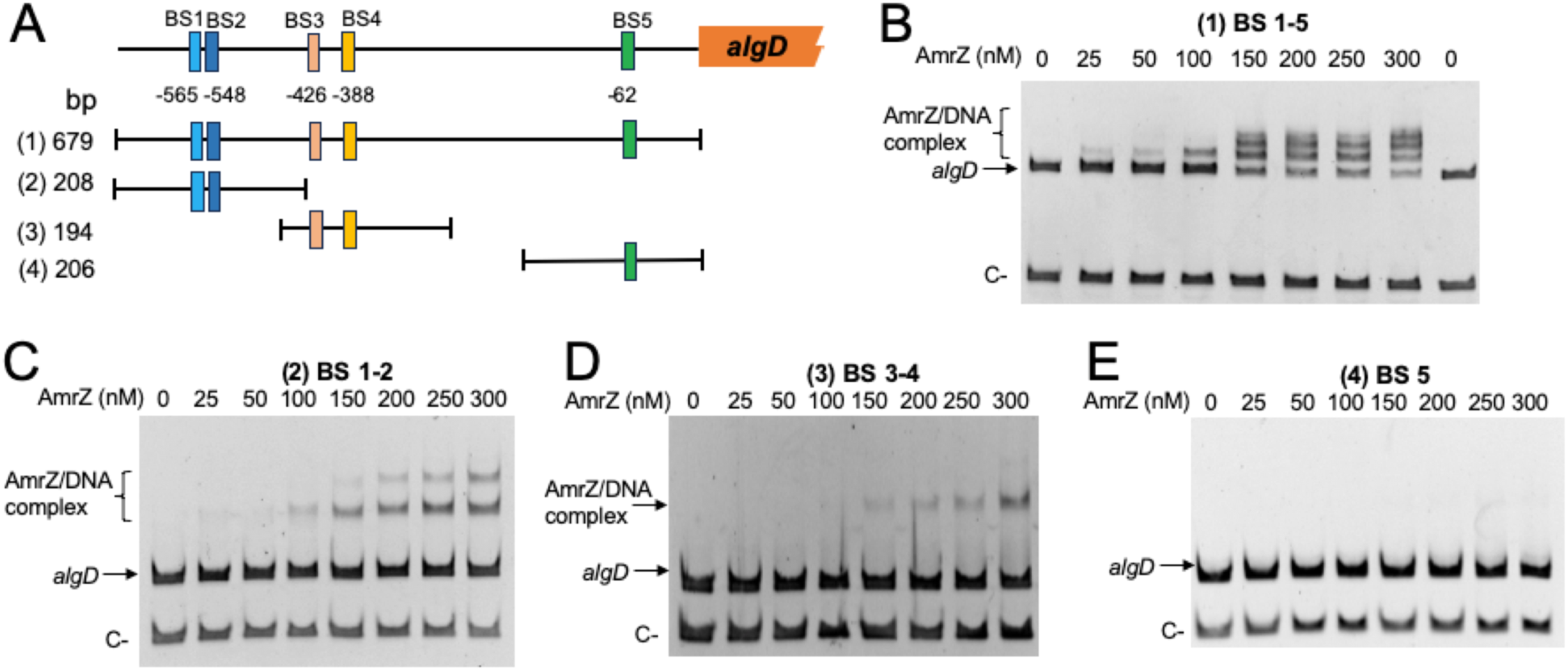
AmrZ directly binds the *algD* regulatory region. (A) Schematic of putative AmrZ- binding sites (BS1-BS5) within the algD regulatory region. The promoter fragments tested in panels B-E are indicated: 679 bp (BS1–BS5), 208 bp (BS1–BS2), 194 bp (BS3–BS4), and 206 bp (BS5). (B-E) EMSAs with increasing concentrations of AmrZ and the indicated *algD* promoter fragments. Shifted bands indicate AmrZ-DNA complex formation. Negative controls (C-) were a 251 bp fragment from ORF AVAEIV_RS16850 (panel B) and a 110 bp fragment from *gyrA* (AVAEIV_RS08140) (panels C-E). DNA was visualized by ethidium bromide staining.

Electrophoretic mobility shift assays (EMSAs) were used to test these predictions. AmrZ caused a mobility shift of a 679-bp P*algD* fragment, indicating DNA-protein complex formation (Fig 2B). With increasing AmrZ concentration, at least three discrete retarded bands were observed, consistent with occupancy of multiple sites and formation of higher-order complexes. Binding was specific, as no shift was detected with a negative-control DNA fragment.

To delimit the sites required for binding, three shorter PCR fragments were tested: 208 bp (BS1-BS2), 194 bp (BS3-BS4), and 206 bp (BS5). AmrZ bound the BS1-BS2 and BS3-BS4 fragments (Fig 2C, D) but not the BS5 fragment (Fig 2E). Together, these data indicate that AmrZ directly recognizes the *algD* regulatory region and acts as a transcriptional activator in *A. vinelandii*.

### Transcriptional regulation of *amrZ*

AmrZ is a RHH transcription factor reported to function as both an activator and a repressor; in several *Pseudomonas* species it autoregulates its own transcription (Ramsey *et al*., 2005). We therefore examined whether this occurs in *A. vinelandii*, noting a putative AmrZ-binding motif in the *amrZ* regulatory region (P*amrZ*) (S3 Table; S2 Fig). *amrZ* transcription was monitored using a PamrZ-gusA fusion. In the wild type, P*amrZ* activity increased gradually over the growth curve, reaching a maximum at ∼36 h. In the Δ*amrZ* mutant, P*amrZ* activity was clearly reduced from 12 h onward (Fig 3A), indicating that, contrary to what has been described in some *Pseudomonas* species, AmrZ positively regulates its own transcription in *A. vinelandii*.

**Fig 3.**
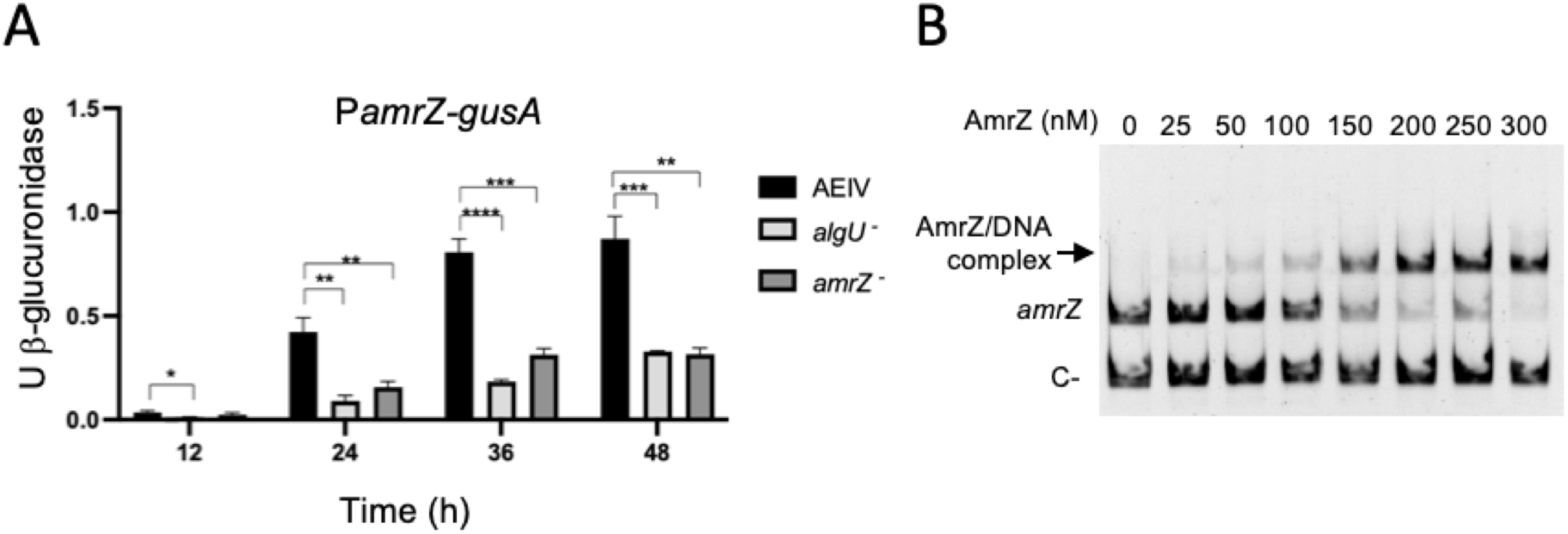
*amrZ* transcription is controlled by AlgU and AmrZ. (A) Activity of a P*amrZ-gusA* transcriptional fusion in the wild type and the Δ*amrZ* and Δ*algU* backgrounds during growth in liquid Burk’s-sucrose medium. (B) EMSA showing AmrZ binding to the *amrZ* regulatory region. A promoter fragment was incubated with increasing concentrations of purified His-AmrZ. A 251-bp fragment from ORF AVAEIV_RS16855 served as a negative control. DNA was visualized by ethidium bromide staining. Bars denote mean ± SD from three independent experiments. Statistical significance (two-tailed Student’s t-test) is indicated: *, *P<0.05;* **, *P* < 0.01; ***, *P* < 0.001; ****, *P* < 0.0001.

To test whether this regulation is direct, EMSAs were performed with a DNA fragment encompassing the *amrZ* promoter region and purified *A. vinelandii* His-AmrZ. A concentration-dependent mobility shift was observed, one or more retarded bands increased with added protein, indicating formation of a DNA-protein complex (Fig 3B). Together with the reporter data, these results support a model in which AmrZ positively autoregulates transcription by directly binding its own regulatory region.

### AlgU is required for *amrZ* transcription

Previous work showed that AmrZ is under positive control of the sigma factor AlgU in *A. vinelandii*: the AmrZ protein was undetectable in the proteome of an AlgU-deficient strain, and an AlgU-dependent promoter motif was identified upstream of *amrZ* (Chowdhury-Paul *et al*., 2023). To test this directly, as also reported for *P. aeruginosa* (Wozniak *et al*., 2003), we introduced a P*amrZ-gusA* transcriptional fusion into a Δ*algU* mutant. P*amrZ* activity was markedly reduced in Δ*algU* relative to wild type, confirming positive regulation of *amrZ* by AlgU (Fig 3A). Notably, loss of either AlgU or AmrZ produced a similar temporal profile, near-complete loss of promoter activity at early time points followed by a gradual increase, suggesting that AmrZ activates transcription from the AlgU-dependent promoter and that an additional promoter contributes later in growth. Consistent with this, a σ⁷⁰-type promoter is predicted within P*amrZ*, ∼205 bp upstream of the AlgU-dependent promoter (Fig. S2).

### AmrZ affects the pool of c-di-GMP in *A. vinelandii*

In *P. aeruginosa*, AmrZ modulates cellular c-di-GMP by regulating the expression of enzymes that synthesize or degrade this second messenger (Jones *et al*., 2014; Muriel *et al*., 2018). Because c-di-GMP is required for alginate production in *A. vinelandii* (Ahumada-Manuel *et al*., 2020; Barrios-Rafael *et al*., 2025) we determined whether AmrZ likewise influences the c-di-GMP pool. We used a previously characterized biosensor in which TurboRFP expression is controlled by three tandem c-di-GMP riboswitches (Bc3–Bc5) (Zhou *et al*., 2016).

To validate the sensor in *A. vinelandii*, we integrated it into Δ*mucG* and Δ*avGreg* mutants, which exhibit ∼4-fold higher and ∼3-fold lower c-di-GMP levels, respectively, relative to wild type AEIV (Ahumada-Manuel *et al*., 2020). TurboRFP fluorescence was measured and normalized to total protein content. As shown in Fig. 4, c-di-GMP levels in AEIV increased over the growth curve, peaking at 48 h. In the Δ*mucG* background (PDE-deficient), a similar trajectory was observed but with consistently higher values than wild type, whereas Δ*avGreg* (DGC-deficient) showed a sustained reduction throughout growth. These results validate the biosensor in *A. vinelandii*, enabling detection of both increased and decreased c-di-GMP relative to wild type (Fig. 4A).

**Fig. 4.**
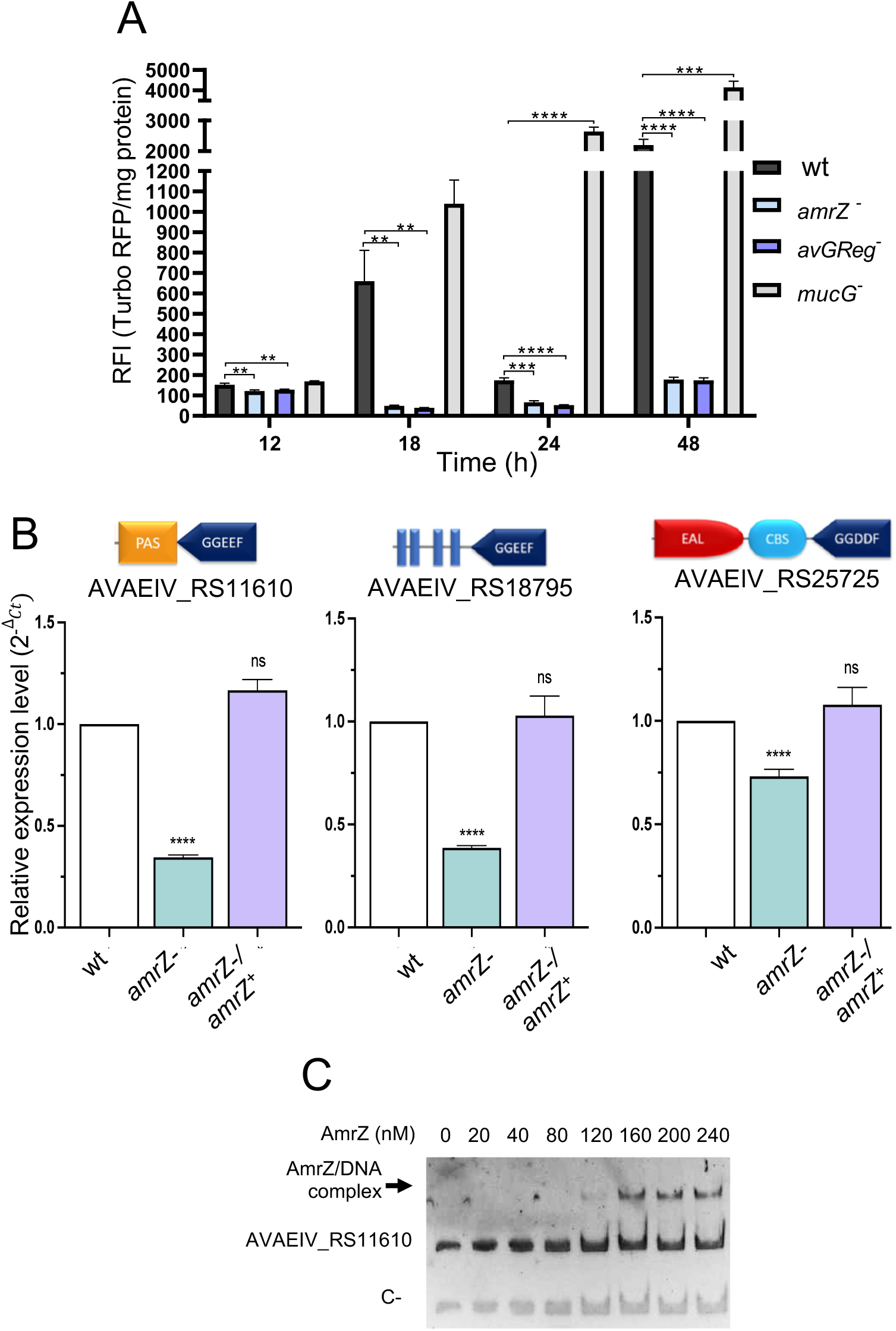
AmrZ regulates intracellular c-di-GMP levels. (A) c-di-GMP biosensor readout in wild type, Δ*amrZ*, Δ*avGreg*, and Δ*mucG* strains. Accumulation was monitored with the TurboRFP riboswitch biosensor (Bc3-Bc5). Relative fluorescence intensity (RFI) = TurboRFP fluorescence/total protein. Bars show mean ± SD from three independent experiments. Statistical significance (two-tailed Student’s t-test): **, *P* < 0.01; ***, *P* < 0.001; ****, *P* < 0.0001. (B) RT-qPCR of AVAEIV_RS11610, AVAEIV_RS18795, and AVAEIV_RS25725 mRNA in the indicated strains grown 24 h in Burk’s-sucrose medium. Domain organizations of the encoded proteins are shown. (C) EMSA demonstrating AmrZ binding to the AVAEIV_RS11610 promoter region. DNA was incubated with increasing concentrations of His-AmrZ. A 251-bp fragment from AVAEIV_RS16855 served as a negative control. DNA was visualized by ethidium bromide staining.

We then introduced the biosensor into the Δ*amrZ* mutant. TurboRFP fluorescence was markedly reduced across the growth curve, approaching the Δ*avGreg* levels, indicating that AmrZ is a key positive determinant of the intracellular c-di-GMP pool in *A. vinelandii*.

### AmrZ modulates c-di-GMP via targets other than *avGreg*

Because Δ*avGreg* and Δ*amrZ* displayed comparably low c-di-GMP levels across the growth curve, we first tested whether AmrZ regulates *avGreg*. This was plausible given that AvGReg is the principal diguanylate cyclase during vegetative growth in *A. vinelandii* (Ahumada-Manuel *et al*., 2020). Expression of *avGreg* was monitored using a P*avGreg-gusA* transcriptional fusion. At 24 h, no significant difference was observed between the wild type and the Δ*amrZ* mutant (S3 Fig), and a slight increase in expression was detected in Δ*amrZ* after 24 h. These results indicate that the reduced basal c-di-GMP in the absence of AmrZ is unlikely to be explained by a transcriptional control of AvGreg and instead suggest that AmrZ modulates c-di-GMP homeostasis through additional DGCs and/or PDEs.

### The AmrZ regulon in *A. vinelandii*

To identify genes directly or indirectly controlled by this transcriptional regulator, we performed RNA-seq–based transcriptional profiling of the wild-type strain AEIV and its Δ*amrZ* derivative (JG521) grown for 24 h in Burk’s-sucrose medium. To accurately define the AmrZ regulon, the AEIV genome was first sequenced (Methods); reads were then mapped to this reference (GenBank GCA_030506185.2).

Principal component analysis revealed a clear separation between wild type and Δ*amrZ* transcriptomes (S4 Fig). Differential expression analysis with DESeq2 (threshold log₂ fold change ≥ 2 and Padj < 0.01) identified 835 differentially expressed genes (DEGs): 498 upregulated and 337 downregulated in Δ*amrZ* relative to wild type (S4 Table). The larger number of genes increased in the mutant suggests a prominent role for AmrZ as a transcriptional repressor, a trend visible in the volcano plot (Fig 5A).

**Fig 5.**
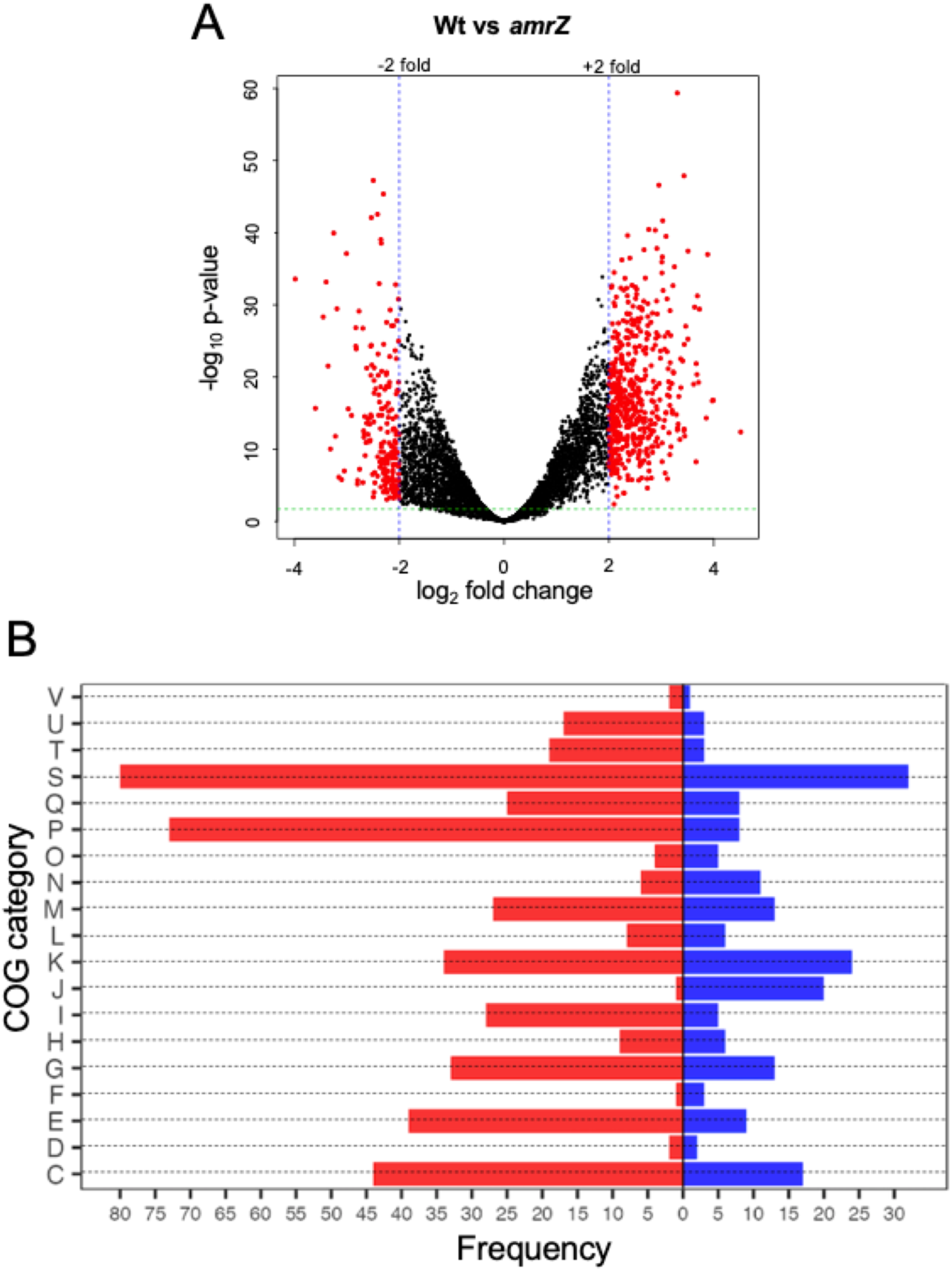
RNA-seq analysis of the Δ*amrZ* mutant versus wild type. (A) Volcano plot of differential expression. Vertical blue lines mark |log₂ fold change| ≥ 2 and the horizontal line marks Padj < 0.01; red points denote genes meeting both criteria. (B) COG functional classification of differentially expressed genes. Bars indicate the number of genes per category; red = upregulated in Δ*amrZ* (negatively controlled by AmrZ), blue = downregulated in Δ*amrZ* (positively controlled by AmrZ). COG codes: C, energy production and conversion; D, cell cycle control, cell division, and chromosome partitioning; E, amino acid transport and metabolism; F, nucleotide transport and metabolism; G, carbohydrate transport and metabolism; H, coenzyme transport and metabolism; I, lipid transport and metabolism; J, translation, ribosomal structure, and biogenesis; K, transcription; L, replication, recombination, and repair; M, cell wall/membrane/envelope biogenesis; N, cell motility; O, posttranslational modification, protein turnover, and chaperones; P, inorganic ion transport and metabolism; Q, secondary metabolite biosynthesis, transport, and catabolism; S, function unknown; T, signal transduction mechanisms; U, intracellular trafficking, secretion, and vesicular transport; V, defense mechanisms. The number of genes in each category is indicated.

Functional categorization of DEGs (Fig 5B) highlighted major effects in energy production and conversion, carbohydrate transport and metabolism, transcription, inorganic ion transport and metabolism, and uncharacterized functions. Inspection of affected genes pointed to specific processes including alginate biosynthesis, ribosome biogenesis/modification, iron homeostasis, aliphatic sulfonate metabolism, nitrogen fixation, and aromatic compound degradation, among others.

Transcriptomic analysis confirmed that AmrZ positively regulates *algD* (log₂ fold change, Δ*amrZ* vs WT = −3.79) and other alginate-biosynthetic genes, including *algL* (log₂ fold change, Δ*amrZ* vs WT = −2.33; S4 Table). The dataset also revealed differentially expressed genes involved in c-di-GMP metabolism, consistent with the altered levels of this second messenger in the Δ*amrZ* mutant.

### AmrZ controls genes for the metabolism of c-di-GMP

AVAEIV_RS11610 and AVAEIV_RS18795, both encoding proteins with conserved c-di-GMP synthesis (GGDEF) domains, showed reduced mRNA accumulation (log₂ fold change = −2.04 and −2.87, respectively) in the Δ*amrZ* mutant; whereas AVAEIV_RS25725, predicted to encode a hybrid DGC/PDE protein, was upregulated (log₂ fold change = 2.38) according to the transcriptome analysis.

RT-qPCR using RNA from the wild type, Δ*amrZ*, and the complemented strain (Δ*amrZ/amrZ*⁺) confirmed downregulation of the DGC-encoding genes in the mutant and restoration upon complementation, indicating positive regulation by AmrZ. In contrast, the RNA-seq upregulation of AVAEIV_RS25725 was not validated by RT-qPCR: its transcript levels were slightly diminished in Δ*amrZ* (Fig 4B).

MEME/FIMO analysis identified a putative AmrZ-binding site upstream of AVAEIV_RS11610 (S3 Table S3; S5 Fig). Consistent with direct regulation, EMSA showed specific binding of AmrZ to this promoter region (Fig 4C), suggesting that AVAEIV_RS11610 is directly activated by AmrZ, whereas regulation of AVAEIV_RS18795 is likely indirect. This regulatory data provides a plausible explanation for the reduced c-di-GMP pool observed in the Δ*amrZ* mutant but does not rule out an AmrZ post-translational control of AvGReg activity.

### AmrZ impacts swimming motility via c-di-GMP

In *Pseudomonas* species, AmrZ regulates both swimming and swarming motility, largely through c-di-GMP–mediated pathways (Ha and O’Toole, 2015; Muriel *et al*., 2018; Hou *et al*., 2019). We therefore examined whether loss of *amrZ* affects swimming motility in *A. vinelandii*. As shown in Fig 6A, deletion of *amrZ* produced a larger swimming halo than the parental strain, and complementation (Δ*amrZ/amrZ*⁺) restored the wild-type phenotype. This behavior aligns with the reduced c-di-GMP levels in Δ*amrZ* and indicates a negative influence of AmrZ on swimming. The enhanced motility is not attributable to alginate deficiency: in an alginate-deficient background (Δ*algA*), the Δ*amrZ* strain still swam farther than the reference strain (data not shown).

**Fig. 6.**
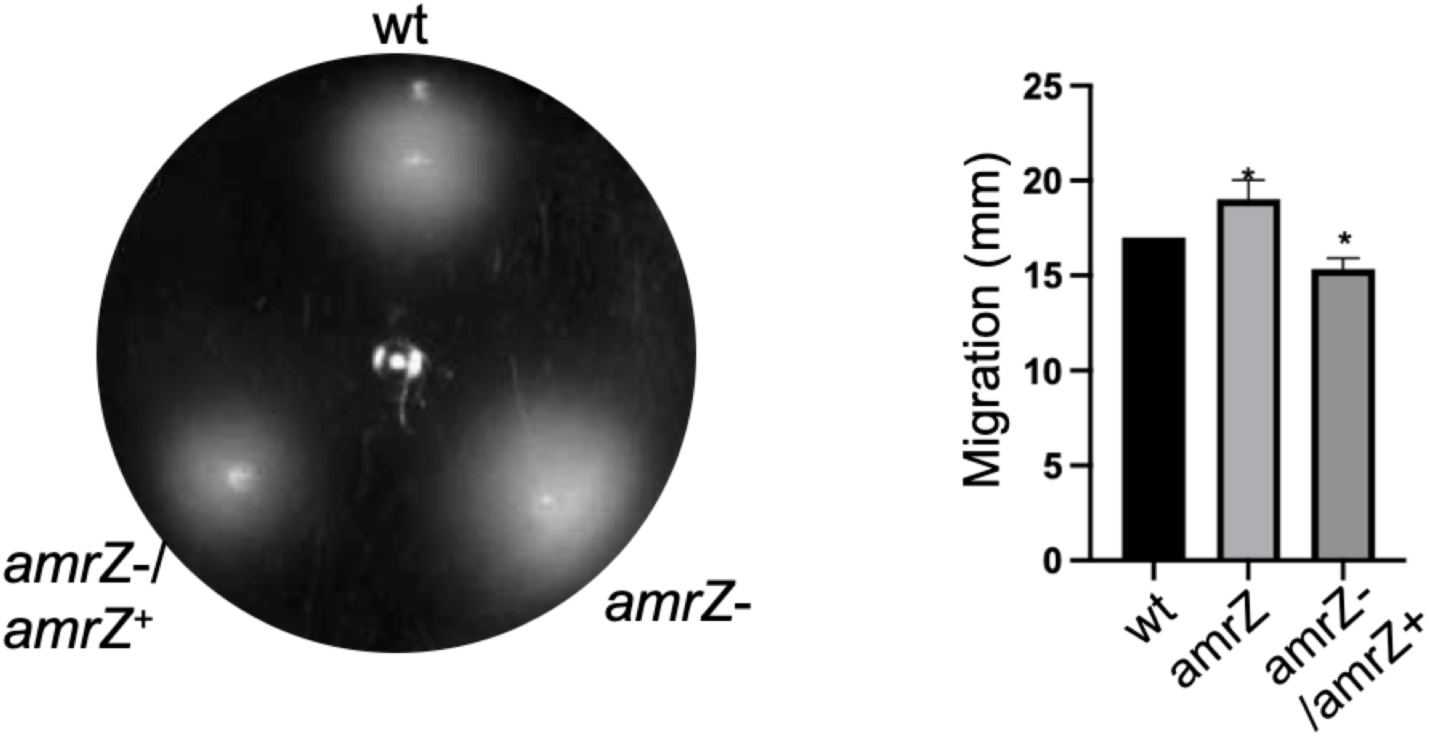
AmrZ negatively regulates swimming motility. Representative swimming motility of the indicated strains on soft agar after 24 h of incubation. The graph shows the quantification of swimming halo diameters.

This effect most likely reflects post-translational consequences of lower c-di-GMP rather than changes in flagellar gene transcription: RNA-seq revealed concerted downregulation of some genes encoding the flagellar basal-body and hook-assembly components (S4 Table).

## DISCUSSION

Our findings demonstrate that the transcriptional regulator AmrZ is essential for alginate biosynthesis in *A. vinelandii*, acting mainly through direct activation of *algD*, encoding the key enzyme in this pathway. Loss of AmrZ completely abolished alginate production, a phenotype restored by genetic complementation, underscoring its central role. This function parallels that of AmrZ in *P. aeruginosa* (Xu *et al*., 2016b), suggesting a conserved regulatory mechanism among different genera of the Pseudomonadaceae.

EMSA results confirmed that AmrZ binds multiple sites in the *algD* promoter, consistent with transcriptional activation. These binding sites are located near or upstream of the AlgU-dependent promoter, indicating that, as in *P. aeruginosa*, AmrZ stimulates transcription from this promoter. Recent evidence also shows that FleQ represses *algD* via two sites but near the RpoS-dependent promoter (Barrios-Rafael *et al*., 2025), suggesting that *algD* expression integrates positive regulation by AmrZ and negative regulation by FleQ. Given that *algD* transcription ultimately requires c-di-GMP, these opposing activities are likely coordinated by intracellular levels of this second messenger.

Interestingly, AmrZ also positively regulates its own expression in *A. vinelandii*, in contrast to the autorepression observed in *Pseudomonas* spp. (Ramsey *et al*., 2005). This positive feedback, dependent on AlgU, may stabilize AmrZ expression under conditions that favor alginate production.

Beyond alginate regulation, our transcriptomic and biochemical data demonstrate that AmrZ exerts broad control over c-di-GMP metabolism. Notably, the principal vegetative DGC, AvGreg, was not transcriptionally affected by AmrZ: PavGreg activity was not diminished, yet cellular c-di-GMP was markedly reduced in Δ*amrZ*. This suggests that, in the Δ*amrZ* background, AvGreg activity may be attenuated post-transcriptionally, potentially via signals sensed by its globin domain (Ahumada-Manuel et al 2020), although this remains to be tested. We identified two additional diguanylate cyclase genes that are positively regulated by AmrZ; their decreased expression in Δ*amrZ* likely contributes to the diminished c-di-GMP pool. These observations are consistent with findings in *P. aeruginosa*, where AmrZ modulates intracellular c-di-GMP to coordinate biofilm formation and motility (Jones *et al*., 2014; Muriel *et al*., 2018). In *A. vinelandii,* low c-di-GMP in the Δ*amrZ* strain also contributes to explain the complete absence of alginate production, given the well-established requirement of this messenger for polysaccharide synthesis (Ahumada-Manuel *et al*., 2020).

The consequences of AmrZ loss extended beyond polysaccharide regulation. RNA-seq analysis revealed over 800 differentially expressed genes, highlighting AmrZ as a global regulator. Categories impacted included central metabolism, iron homeostasis, nitrogen fixation, and ribosome synthesis, underscoring the pleiotropic role of this regulator. Interestingly, the predominance of upregulated genes suggests that AmrZ functions largely as a transcriptional activator in *A. vinelandii*, contrasting with its dual activator/repressor roles in *Pseudomonas* (Hou *et al*., 2019).

Phenotypically, the absence of AmrZ resulted in increased swimming motility, a hallmark of reduced c-di-GMP signaling. This is consistent with the established antagonism between motility and sessile lifestyles across bacteria (Ha and O’Toole, 2015). Thus, AmrZ integrates multiple regulatory layers to coordinate polysaccharide synthesis, motility, and global cellular functions through modulation of c-di-GMP homeostasis.

In conclusion, our results establish AmrZ as a master regulator in *A. vinelandii*, directly activating *algD*, sustaining high intracellular c-di-GMP levels, and broadly influencing gene expression programs beyond alginate production.

## ACKNOWLEDGMENTS

We thank Dr. D. Zamorano-Sánchez for kindly providing plasmid pFY4357 and for his valuable advice. We also thank V. Jiménez for technical assistance, and A. Becerra and R. Bahena for oligonucleotide synthesis, DNA sequencing, and computational support. M.C. Gonzaga-Pérez was a recipient of a PAPIIT-UNAM fellowship. This work was supported by a grant from the Programa de Apoyo a Proyectos de Investigación e Innovación Tecnológica, (PAPIIT IN215924), UNAM to CN.

## SUPPORTING INFORMATION

**S1 Fig.** Putative AmrZ binding sites identified in the regulatory region of *algD*. *algD* regulatory region. The *algD* structural region is in red. The potential AmrZ binding sites are indicated (green) along with the regions used to design the oligonucleotides for PCR amplification of the fragments for EMSAs.

**S2 Fig.** Regulatory region of *amrZ*. The structural region of *amrZ* (AVAEIV_RS17290) is shown along with the predicted AlgU and σ^70^ promoters, and the putative AmrZ binding site (green). The DNA sequence to design oligonucleotides pAPAmrZXbaF and pAPAmrZEcoRIRv, used for PCR amplification of the *amrZ* regulatory region used for EMSA is shown.

**S3 Fig.** Influence of the *amrZ* mutation on the transcription of *avGReg*. The transcriptional fusion P*avGReg-gusA* was introduced into the wt strain and its derivative mutant *amrZ*. The graphic shows a B-glucuronidase kinetics along the growth curve. Strains were cultured in liquid Burks-sucrose medium and samples were taken at the indicated times.

**S4 Fig.** Principal component analysis conducted with the RNA-seq expression data from the wild type strain AEIV (WT) and its derivative mutant *amrZ*.

**S5 Fig**. Regulatory region of *AVAEIV_11610*. The structural region of *AVAEIV_RS11610* is shown in red, along with the detected AmrZ binding site (green). The DNA sequence to design oligonucleotides EMSARS11610Fw and EMSARS11610Rv, used for PCR amplification of the *amrZ* regulatory region is shown. Part of the structural region of ORF AVAEIV_RS11615 located upstream is also shown in red.

**S1 Table.** Strains and plasmids used in this work.

**S2 Table.** Oligonucleotides used in the present study

**S3 Table.** Predicted AmrZ binding sites by MEME/FIMO analysis 400 nt upstream and 50 nt downstream each gene of the *A. vinelandii* AEIV genome.

**S4 Table**. Genes differentially expressed in the τ1*amrZ* mutant when compared to the wt strain AEIV, based on the RNAseq data.

