## Supplemental Material FigS1-S5 and Table S1-S4 for "*Azotobacter vinelandii* AmrZ is a global regulator linking alginate production and c-di-GMP homeostasis"

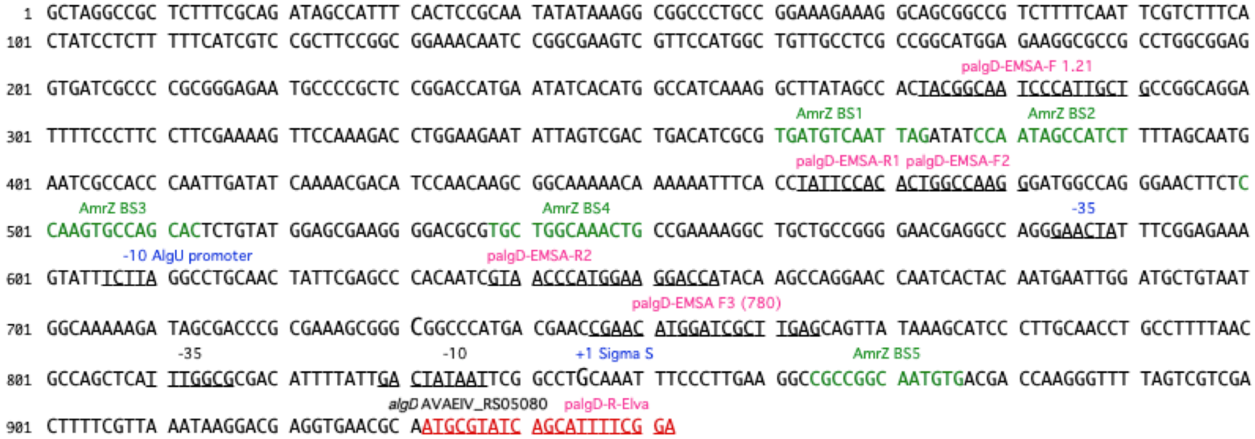


**S1 Fig. Putative AmrZ binding sites identified in the regulatory region of *algD*.** algD regulatory region. The *algD* structural region is in red. The potential AmrZ binding sites are indicated (green) along with the regions used to design the oligonucleotides for PCR amplification of the fragments for EMSAs.

```
                      ← AVAEIV_RS17280
  1 GGCAAGCAGT TGCTGAGGCT TGAGCATGGC GTGCCTCTCC TTTCCACGGC GCTCCGCACA
 61 CAGTACGCAG CCGCCAGCGG GAAGTTCAAG CAGTCCAAAC CGCGTCGAAA GTCAAACCAC
121 TCCCCTGCAG CTGCACAGGC TCAGGAATGC ACGAGGCACT ACGACGAAGA CCGCAGCGCG
181 ACATGCCTGC CATCGAAGCT GCGTATTAAA TGAAGGCAGC GGCATCTTCT GAAATGCCCG
241 GCTCCATTCA GCCATCGCGG CAGTCGGCAT CGTCAGGAAT GCGAACGGGA TCATCTGTCG
301 GGACAGGCGC CACGGCAGGC GACCGGGAGG TGGTGGCGCC CCCTGCTGGA GTCGAACCAG
361 CATCTAGCCC TTAGGAGGGG CTTGTTCTAT CCATTGAACT AAGGGGACTT GCCGAAAAAG
                          σ70 promoter
              -35                         -10      pAPAmrZXbalFw
421 CCATCGATTT TCACACCCCG CCTACGCGGG GATACTCTCG CCAAGCTTCC GATCAGTCGA
481 CCGACCAATG GCGATGGGGC GGCATTCTAG CCATGAAAGA GCCCTTTGTC ATGCCTACAA
541 AAACAGGAAG CGACAACGGA ACAACAGGGT GTCCGCGCAG TCGATTCTTG GGCTCGTCTC
601 ACCGGAAATC CCGGCGTCAG TGGAGCATGC AAATCAACTG CATGCGATTA ATCGATTCAG
                                 AlgU promoter
                       -35                       -10
661 ATATTGAAAA GAGTGAACCA CTTCAGCAAG ACGAGGTCAT ACCGAGAACA CCCTTGCAAG
         AmrZ BS                               amrZ AVAEIV_RS17290 →
721 CTAATTGCCA GCACTCTATA ACAACTCACC GGTCGAAACT ATGCAACCTA TGAAACAGGC
                                                   pAPAmrZEcoRIRv
781 AATCTACTCT AGCCGTACAG CTGACAAATT CGTCGTTCGT CTACCCGATG GCATGCGTGA
```

**S2 Fig. Regulatory region of *amrZ*.** The structural region of *amrZ* (AVAEIV_RS17290) is shown along with the predicted AlgU and $\sigma^{70}$ promoters, and the putative AmrZ binding site (green). The DNA sequence to design oligonucleotides pAPAmrZXbaF and pAPAmrZEcoRIRv, used for PCR amplification of the *amrZ* regulatory region used for EMSA is shown.

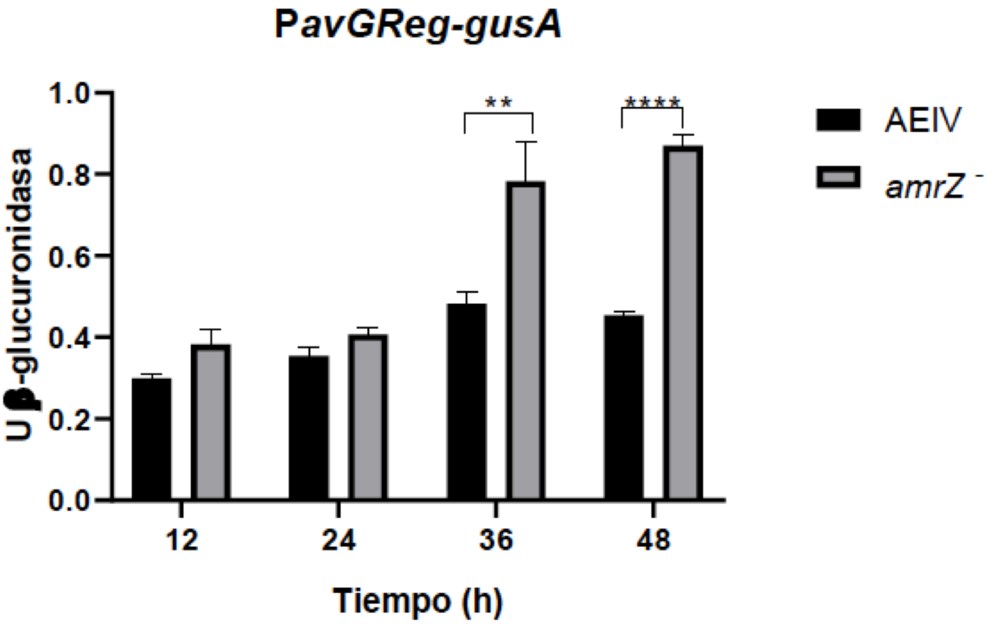


**S3 Fig. Influence of the *amrZ* mutation on the transcription of *avGReg*.** The transcriptional fusion P*avGReg-gusA* was introduced into the wt strain and its derivative mutant *amrZ*. The graphic shows a B-glucuronidase kinetics along the growth curve. Strains were cultured in liquid Burks-sucrose medium and samples were taken at the indicated times.

S4 Fig

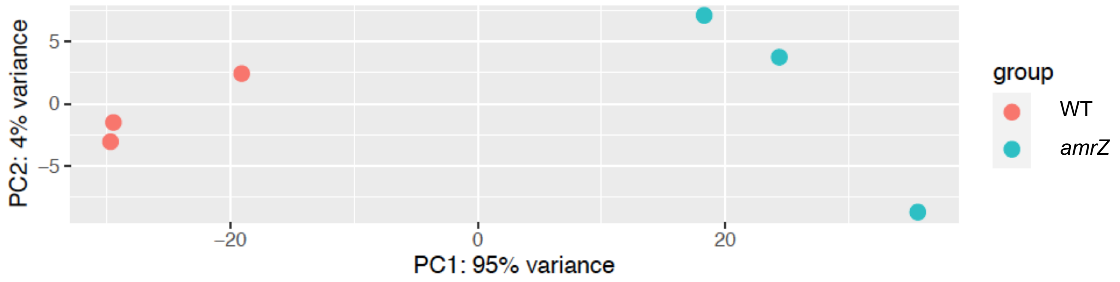


**S4 Fig.** Principal component analysis conducted with the RNAseq expression data from the wild type strain AEIV (WT) and its derivative mutant *amrZ.*

S5 Fig

```
                                                   EMSA RS11610Fw
  1 GCGGCACCTG CTCGACATCC GCAATTGCGG CCTGGCCGGC GCCCTGCAGC TCGCCCCACG
 61 CGACGGCGAC GCCCTGGTGC GCCCTTTCGA AGCGGCCATG CGGCTCTGGC GCCGGGGCTT
121 CTACGTGCGT TGCGGCGGCG ACACCCTGCA GTTCGGTCCG CCCTTCAGCA GTTCCGCCGC
181 CGAACTGGAT CGCCTGTTCG ACGCCGTGGG CGAAGTGCTC GGCGGACTCG ACTAAAGAAG
                                             AmrZ BS
241 CCGGGCTGCC CGGCGCAAAG CGCCCCGGAT GCTAGCGCTT TGATCTATCC TGCTCGCAGG
301 CTCCGTCGAC CACCTCCGGT ACAGACGGGT GTATTCCCGC CGGCCGGAGT GCCGGCCCAG
                      AVAEIV_RS11610 →        EMSA RS11610Rv
361 CAGTAGACCG GACATGCATT CGCCAGGAAG CAAGCCAGAA GAAACCAGCC GCGAGCCAGG
421 GACGGCCTGC GACTGTAGCC GTACCGCCGG CTTCGAAATC ATCCTCGACA GCATCGACGC
481 CATCGTCTAT GTGGCGGACA TGCGTACCCA CGAACTGTTG TACCTCAACA GCTACGGGCG
541 CCAGCACTAC GGCGACCCCG AGGCGCTGAA GTGCTGGCAG GTGCTGCAGG CCGGACAGGC
601 CGGCCCCTGC GATTTCTGCA GCAACCCCCA
```

**S5 Fig. Regulatory region of *AVAEIV_11610*.** The structural region of *AVAEIV_RS11610* is shown in red, along with the detected AmrZ binding site (green). The DNA sequence to design oligonucleotides EMSARS11610Fw and EMSARS11610Rv, used for PCR amplification of the *amrZ* regulatory region is shown. Part of the structural region of ORF AVAEIV_RS11615 located upstream is also shown in red.

**S1 Table. Strains and plasmids used in this work.**

| Name | Characteristics | Source |
|---|---|---|
| **Strains** | | |
| AEIV | Wild type strain | (Larsen y Haug, 1971) |
| JG521 | Mutant Δ*amrZ*::Km | This work |
| MG06 | JG521 derivative; carries a wild type copy of *amrZ*. Δ*amrZ*::Km, *amrZ*+. Kmr, Tcr | This work |
| AEalgU | AEIV derivative, carries an *algU*::Km mutation | (Chowdhury-Paul et al., 2023) |
| AED-gusA | Carries an *algD-gusA* transcriptional fusion. Gmr | (Quiroz-Rocha et al., 2017) |
| JG522 | AED-*gusA* derivative, carries and Δ*amrZ*::Km mutation. Gmr, Kmr | This work |
| CLAM31 | Mutant Δ*amrZ*::Gm | This work |
| CLAM17 | Mutant *avGReg*::Gm | (Ahumada-Manuel et al., 2020) |
| CLAM04 | *mucG*::Sp mutant, Spr | (Ahumada-Manuel et al., 2020) |
| CLAM36 | AEIV derivative. Carries the c-di-GMP biosensor and the Turbo-RFP encoding gene, under the control of a σ70 promoter, contained in pLA68; integrated into the chromosome. Kmr | This work |
| CLAM37 | CLAM36 derivative, carries the *mucG*::Sp mutation. Spr, Kmr | This work |
| MG07 | CLAM36 derivative, carries the *avGReg*::Gm mutation. Gmr, Kmr | This work |
| MG08 | CLAM36 derivative, carries the Δ*amrZ*::Gm mutation. Gmr, Kmr | This work |
| MG01 | AEIV derivative; carries a P*avGReg-gusA* transcriptional fusion. Tcr | This work |
| MG03 | JG521 derivative; carries the P*avGReg-gusA* construction. Δ*amrZ*::Km. Kmr, Tcr | This work |
| MG02 | AEIV derivative; carries a P*amrZ-gusA* transcriptional fusion. Tcr | This work |
| MG04 | JG521 derivative; carries the P*amrZ-gusA* construction. Δ*amrZ*::Km. Kmr, Tcr | This work |
| MG05 | AEalgU derivative, carries the P*amrZ-gusA* transcriptional fusion. *algU*::Km. Kmr, Tcr | This work |

| | | |
|---|---|---|
| **Plasmids** | | |
| pJET1.2/Blunt | PCR clonning vector. Apr | Thermo Scientific |
| pUMATc (5'-3') | Vector enabling the integration into the *A. vinelandii* chromosome of DNA fragments within the *melA locus* by double homologous recombination. Tcr | Cocotl-Yañez et al., 2014 |
| pUMAgusATcT | Used for the construction of transcriptional fusions with the *gusA* reporter gene. Tcr | Muriel-Millán et al., 2015 |
| pUMA-Km(5'-3') | Plasmid used for the integration of DNA into the *A. vinelandii* genome within the *melA locus*. Kmr | This work |
| pBSL141 | Source of the Gm cassette | Alexeyev et al., 1995 |

| | | |
|---|---|---|
| pBSL98 | Source of the Km cassette | Alexeyev et al., 1995 |
| pFY4357 | Plasmid carrying a c-di-GMP biosensor | Zamorano-Sanchez et al., 2019 |
| pET-28a+ | Expression vector | Novagen |
| pET-amrZ | pET-28 a+ derivative, carries the *amrZ* gene as a NdeI-BamHI fragment of 384 bp. Expression of AmrZ | This work |
| pJETamrZ | pJET1.2/Blunt derivative carrying the *amrZ* gene, Ampr | This work |
| pJETamrZ-Gm* | pJETamrZ derivative. An internal *amrZ* ScaI fragment of 96 pb was replaced by a Gmr cassette (Δ*amrZ*::Gm). Used for the construction of mutants CLAM31 and MG08. | (Ahumada-Manuel et al., 2020) |
| pJETamrZ-Km* | pJETamrZ derivative. An internal amrZ ScaI fragment of 96 pb was replaced by a Kmr cassette (Δ*amrZ*::Km). Used for the construction of mutants JG521 and JG522 and | This study |
| pUMAamrZ* | pUMATc 5'-3' derivative; carries a copy of the wild type *amrZ* gene. Used for construction of strain MG06. Tcr | This work |
| pJETPamrZ | pJET1.2/Blunt derivative, carries the regulatory region of *amrZ* (P*amrZ*). Ampr | This work |
| pJETP420 | pJET1.2/Blunt derivative, carries the regulatory region of *avGReg* (P*avGReg*) Ampr | This work |
| pUMApamrZ-gusA* | pUMAgusATcT derivative, carries the P*amrZ-gusA* construction. PamrZ was cloned as a XbaI-EcoRI fragment. Used for the construction of strains MG02, MG04 and MG05. Tcr | This work |
| pUMAp420-gusA* | pUMAgusATcT derivative, carries the P*avGReg-gusA* construction. P*avGReg* was cloned as a XbaI-EcoRI fragment. Used for the construction of strains MG01 and MG03. Tcr | This work |
| pLA05* | Carries a Δ*mucG*::Sp mutation. Used for construction of mutant CLAM37 | (Ahumada-Manuel et al., 2020) |
| pLA33* | Carries a Δ*avGReg*::Gm mutation. Used for construction of mutant MG07 | (Ahumada-Manuel et al., 2020) |
| pJB2 | pJET1.2/Blunt derivative. Carries a 2.2 kb fragment with the dual tandem riboswitch Bc4-5 and the gene encoding the Turbo RFP protein | This work |
| pLA65 | pJET1.2/Blunt, carries a 154 pb fragment with a σ70 promoter. Apr. | This work |
| pLA66 | pLA65 derivative. Carries the c-di-GMP biosensor (Bc4-5-TurboRFP) under the control of a σ70 promoter. Apr | This work |
| pLA68* | pUMAKm(5'-3') derivative, carries the Pσ70-Bc4-5-TurboRFP construction from pLA66. Apr, Kmr. Used for construction of strain CLAM36 | This work |

* These plasmids were previously linearized with ScaI before transformation to allow double reciprocal recombination and allelic exchange

**S2 Table. Oligonucleotides used in the present study**

| Name | Sequence (5'-3') | Use of the amplicon | DNA template |
|---|---|---|---|
| AmrZ-F | GATACTCTCGCCAAGCTTC | *locus amrZ,* 1099 bp. Construction of pJETamrZ | AEIV chromosomal DNA |
| AmrZ-R | TGATGACGGGCGCAATGAC | | |
| pAmrZXbaF | ACTCTAGATTTCCACGGCGCTCCGCACAC | *amrZ* regulatory region. 799 bp, construction of pUMApamrZ-gusA | AEIV chromosomal DNA |
| pAmrZEcoRRv | ACGAATTCCACGCATGCCATCGGGTAG | | |
| pGm-F | TTTGCCCATGGACGCACAC | $\sigma^{70}$ promoter from the Gm cassette. 154 bp. Construction of pLA65 | pBSL141 |
| PRIMER2 | AGTAGGATCCGTTACCACCGCTGCGTTC | | |
| p00420XbaIFw | ACTCTAGAAGTGCGCAGTGCCTGGAAG | *avGReg* regulatory region. 268bp, construction of pUMAp420-gusA | AEIV chromosomal DNA |
| p00420EcoRIRv | ACGAATTCTGCCGCGCTGGGTGTTACG | | |
| algD-EMSA-F1.21 | TACGGCAATCCCATTGCTG | Amplification of *algD* DNA fragments for EMSA assay | AEIV chromosomal DNA |
| palgD-EMSA-F2 | TATTCCACACTGGCCAAGG | | |
| palgD-EMSA-F3 | CGAACATGGATCGCTTGAG | | |
| palgD-EMSA-R1 | CTTGGCCAGTGTGGAATAG | | |
| palgD-EMSA-R2 | TGGTCCTTCCATGGGTTAC | | |
| palgD-R | CCGAAAATGCTGATACGC | | |
| AmrZ-NdeI | CGCATATGCAACCTATGAAACAGGCAATC | AmrZ expression. 384 bp, construction of pET-amrZ | AEIV chromosomal DNA |
| AmrZ-BamhIRv | ACGGATCCTGGCTGAAGGCAGTGCATGTC | | |
| algD-RT3-F | TTCGGACTGGGCTATGTAGG | *algD* qPCR | cDNA |
| algD-RT3.2-R | ATCTTGGCGGCGGAAATATC | | |
| qgyrAL-F | CAA GCT GGC CCA TGA ACT GCT C | *gyrA* qPCR | cDNA |
| qgyrAL-R | TGG GAA TCC TGG TCG GCA TCA C | | |
| qamrZ_F | ACTCTAGCCGTACAGCTGAC | *amrZ* qPCR | cDNA |
| qAmrZ_R | AGTTCATGCTGCGGTGATGG | | |
| qPCR Fw avin38420 | GTT TCC ACC GAC GAA CTG AC | AVAEIV_RS18795 qPCR | cDNA |
| qPCR Rv avin38420 | GCG CTT GAA ATG GTC CAG AT | | |
| qPCR Fw avin22780 | AAC TCA CCG GCC TGA ACA A | AVAEIV_RS11610 qPCR | cDNA |
| qPCR Rv avin22780 | TTG AAG TGG TCC AGG TCG AA | | |
| qRecA F | TCA CCG GCA ACA TCA AGA AC | *recA* qPCR | cDNA |
| qRecA R | ACC GAG GCG TAG AAC TTC AG | | |
| pMMB-Rv | CTG GCA GTT CCC TAC TCT C | Bc4-5-TurboRFP. 2228 bp. Construction of pJB2 | pFY4357 |
| pMMB-dual-SecF | CTG GCT TTG CTT CCA GAT GT | | |
| 33420-F | AAG TCT CGC TCG TCA CTT C | Negative control in EMSA assay. 605 bp | AEIV chromosomal DNA |
| 33420-R | CTC CAG GAG CCG GTA TAT G | | |
| qgyrAL-F | CAA GCT GGC CCA TGA ACT GCT C | Negative control in EMSA assay. 110 bp | AEIV chromosomal DNA |
| qgyrAL-R | TGG GAA TCC TGG TCG GCA TCA C | | |
| pApAmrZXbaIFw | ACT CTA GAT ACT CTC GCC AAG CTT CCG | Amplification of *amrZ* DNA fragments for EMSA assay. 404 bp | AEIV chromosomal DNA |
| pApAmrZEcoRIRv | ACG AAT TCC ACG CAT GCC ATC GGG TAG | | |
| 33420B-F | CCC TTT CAG CCT GTT CAC | Negative control in EMSA assay. 251 bp | AEIV chromosomal DNA |
| 33420B-R | GAC AAA CCC TGC CTT TCG | | |
| EMSA RS11610 Fw | ACA CCC TGC AGT TCG GTC | Amplification of AVAEIV_RS11610 DNA fragment for EMSA assay. 270 | AEIV chromosomal DNA |
| EMSA RS11610 Rv | GGC TGG TTT CTT CTG GCT TG | | |

**S3 Table. Predicted AmrZ binding sites by MEME/FIMO analysis 400 nt upstream and 50 nt downstream each gene of the *A. vinelandii* AEIV genome.**

| locus_tag | Start coordinate | End coordinate | Distance to ATG codon | Strand | Score | *p-val* | q-val | Sequence | Genes | Product |
|---|---|---|---|---|---|---|---|---|---|---|
| AVAEIV_RS12210 | 2586370 | 2586382 | 196 | + | 18.6286 | 1.27E-08 | 0.0174 | CAGATCGCCATCA | - | flagellar basal body P-ring protein FlgI |
| AVAEIV_RS01190 | 233047 | 233059 | 372 | + | 18.0571 | 2.31E-08 | 0.0191 | CAGATCGCCAGCA | - | ATP-binding cassette domain-containing protein |
| AVAEIV_RS07210 | 1486556 | 1486568 | 135 | + | 18.0571 | 2.31E-08 | 0.0191 | CAGATCGCCAGCA | - | iron-containing alcohol dehydrogenase |
| AVAEIV_RS24210 | 5147484 | 5147496 | 211 | - | 17.5 | 8.42E-08 | 0.0464 | CAGAACGCCATCA | *ftsE* | cell division ATP-binding protein FtsE |
| AVAEIV_RS07035 | 1449144 | 1449156 | 354 | - | 17.3143 | 1.10E-07 | 0.043 | CCGATCGCCATCA | - | co-chaperone GroES |
| AVAEIV_RS11080 | 2335171 | 2335183 | 111 | + | 17.3 | 1.20E-07 | 0.043 | CAGATCGCCACCA | - | amino acid ABC transporter ATP-binding protein |
| AVAEIV_RS03095 | 637084 | 637096 | 129 | - | 17.2571 | 1.67E-07 | 0.043 | CAGTTCGCCAGCA | *fusA* | elongation factor G |
| AVAEIV_RS00785 | 153731 | 153743 | 102 | + | 17.1714 | 1.78E-07 | 0.043 | CAGATGGCCAGCA | *nifS* | cysteine desulfurase NifS |
| AVAEIV_RS25630 | 5479602 | 5479614 | 88 | + | 17.1143 | 1.88E-07 | 0.043 | CACATCGCCAGCA | - | SMP-30/gluconolactonase/LRE family protein |
| AVAEIV_RS10385 | 2170456 | 2170468 | 377 | + | 16.9286 | 2.30E-07 | 0.0451 | CAGATCGCCAGCG | - | FAD-dependent oxidoreductase |
| AVAEIV_RS06660 | 1377181 | 1377193 | 190 | - | 16.8429 | 2.55E-07 | 0.0649 | CAGATTGCCAGCA | *rapZ* | RNase adapter RapZ |
| AVAEIV_RS17335 | 3754351 | 3754363 | 353 | + | 16.7429 | 2.77E-07 | 0.0649 | CCGATCGCCAGCA | - | hypothetical protein |
| AVAEIV_RS06320 | 1290611 | 1290623 | 344 | - | 16.4571 | 4.62E-07 | 0.0649 | CAGAGCGCCAGCA | - | DUF3094 domain-containing protein |
| AVAEIV_RS06330 | 1290611 | 1290623 | 369 | - | 16.4571 | 4.62E-07 | 0.0563 | CAGAGCGCCAGCA | - | hypothetical protein |
| AVAEIV_RS00720 | 145878 | 145890 | 202 | - | 16.4286 | 4.72E-07 | 0.0563 | CCGATGGCCATCA | - | NifX-associated nitrogen fixation protein |
| AVAEIV_RS11020 | 2320732 | 2320744 | 79 | - | 16.4143 | 4.83E-07 | 0.0563 | CAGATGGCCACCA | - | FAD-dependent oxidoreductase |
| AVAEIV_RS21080 | 4483546 | 4483558 | 252 | + | 16.3714 | 5.60E-07 | 0.0649 | CAGTTGGCCAGCA | - | 2-keto-4-pentenoate hydratase |
| AVAEIV_RS24270 | 5160750 | 5160762 | 287 | + | 16.3714 | 5.60E-07 | 0.0649 | CCCATCGCCATCA | - | sensor domain-containing diguanylate cyclase |
| AVAEIV_RS03005 | 619905 | 619917 | 98 | - | 16.3429 | 5.81E-07 | 0.0563 | CACCTCGCCATCA | *birA* | bifunctional biotin--[acetyl-CoA-carboxylase] ligase/biotin operon repressor BirA |
| AVAEIV_RS18710 | 4030327 | 4030339 | 308 | - | 16.3143 | 6.02E-07 | 0.0649 | CGCATCGCCATCA | - | IclR family transcriptional regulator |
| AVAEIV_RS05450 | 1128031 | 1128043 | 346 | - | 16.2857 | 6.30E-07 | 0.0563 | CAGATTGCCATCG | *ispH* | 4-hydroxy-3-methylbut-2-enyl diphosphate reductase |
| AVAEIV_RS03520 | 709711 | 709723 | 369 | + | 16.1857 | 6.70E-07 | 0.0563 | CCGATCGCCATCG | - | glycoside hydrolase family 5 protein |
| AVAEIV_RS04250 | 867134 | 867146 | 212 | + | 16.1857 | 6.70E-07 | 0.0649 | CCGATCGCCATCG | - | hypothetical protein |
| AVAEIV_RS07625 | 1589588 | 1589600 | 360 | - | 16.1857 | 6.70E-07 | 0.0563 | CCGATCGCCATCG | - | ComEC/Rec2 |
| AVAEIV_RS15595 | 3417847 | 3417859 | 244 | - | 16.1857 | 6.70E-07 | 0.0649 | CCGAACGCCATCA | *smc* | chromosome segregation protein SMC |
| AVAEIV_RS19880 | 4234233 | 4234245 | 387 | - | 16.1857 | 6.70E-07 | 0.0649 | CCGATCGCCATCG | - | YqfO family protein |
| AVAEIV_RS16005 | 3488230 | 3488242 | 323 | + | 16.1571 | 7.36E-07 | 0.0563 | CAGCTCGCCATCG | - | hypothetical protein |
| AVAEIV_RS16560 | 3592014 | 3592026 | 222 | + | 16.1571 | 7.36E-07 | 0.0649 | CAGCTCGCCATCG | - | MarR family transcriptional regulator |
| AVAEIV_RS25255 | 5377611 | 5377623 | 89 | + | 16.1571 | 7.36E-07 | 0.0563 | CAAAACGCCATCA | - | kinase/response regulator |
| AVAEIV_RS02780 | 569627 | 569639 | 284 | + | 16.1286 | 7.84E-07 | 0.0563 | CGGATCGCCATCG | *epd* | erythrose-4-phosphate dehydrogenase |
| AVAEIV_RS07590 | 1582939 | 1582951 | 361 | + | 16.1286 | 7.84E-07 | 0.0563 | CGGATCGCCATCG | *nqrM* | (Na+)-NQR maturation NqrM |
| AVAEIV_RS15295 | 3340630 | 3340642 | 324 | + | 16.1286 | 7.84E-07 | 0.0563 | CGGATCGCCATCG | *fabI* | enoyl-ACP reductase FabI |
| AVAEIV_RS08330 | 1735552 | 1735564 | 170 | - | 16.0714 | 8.52E-07 | 0.0563 | CAAATTGCCATCA | - | type I secretion system permease/ATPase |
| AVAEIV_RS11780 | 2495467 | 2495479 | 326 | - | 16.0571 | 8.64E-07 | 0.0649 | CTGATCGCCATCA | - | alpha-L-glutamate ligase-like protein |
| AVAEIV_RS05080 | 1046640 | 1046652 | 381 | + | 16.0429 | 9.09E-07 | 0.0649 | CAGTTTGCCAGCA | - | nucleotide sugar dehydrogenase |
| AVAEIV_RS05190 | 1067482 | 1067494 | 249 | - | 15.9857 | 9.59E-07 | 0.0649 | CCGATCGCCACCA | *yaaA* | peroxide stress protein YaaA |
| AVAEIV_RS20470 | 4353056 | 4353068 | 359 | + | 15.8857 | 1.11E-06 | 0.0649 | CGGTTCGCCAGCA | - | FecR domain-containing protein |
| AVAEIV_RS02100 | 419209 | 419221 | 330 | - | 15.8571 | 1.11E-06 | 0.0649 | CCGATGGCCAGCA | *istA* | IS21 family transposase |
| AVAEIV_RS02110 | 419209 | 419221 | 219 | - | 15.8571 | 1.11E-06 | 0.0563 | CCGATGGCCAGCA | - | hypothetical protein |
| AVAEIV_RS02280 | 455353 | 455365 | 178 | - | 15.8143 | 1.18E-06 | 0.0649 | CAGTTGGCCATCG | - | oxidative damage protection protein |
| AVAEIV_RS05895 | 1208221 | 1208233 | 48 | - | 15.8143 | 1.18E-06 | 0.0649 | CAGTTGGCCATCG | - | TIGR03751 family conjugal transfer lipoprotein |
| AVAEIV_RS05910 | 1210071 | 1210083 | 216 | - | 15.8143 | 1.18E-06 | 0.0649 | CAGTTGGCCATCG | - | hypothetical protein |
| AVAEIV_RS05915 | 1210071 | 1210083 | 48 | - | 15.8143 | 1.18E-06 | 0.0649 | CAGTTGGCCATCG | - | hypothetical protein |
| AVAEIV_RS15050 | 3277592 | 3277604 | 11 | + | 15.8143 | 1.18E-06 | 0.0649 | CAGGTCGCCAGCA | - | GspMb/PilO family protein |
| AVAEIV_RS03500 | 706694 | 706706 | 186 | + | 15.8 | 1.22E-06 | 0.0649 | CCCATCGCCAGCA | - | hypothetical protein |
| AVAEIV_RS05430 | 1123275 | 1123287 | 307 | - | 15.8 | 1.22E-06 | 0.0563 | CCCATCGCCAGCA | *ribF* | bifunctional riboflavin kinase/FAD synthetase |
| AVAEIV_RS07375 | 1529976 | 1529988 | 228 | - | 15.8 | 1.22E-06 | 0.0563 | CCCATCGCCAGCA | - | OprD family porin |
| AVAEIV_RS11780 | 2495205 | 2495217 | 64 | + | 15.8 | 1.22E-06 | 0.0649 | CCCATCGCCAGCA | - | alpha-L-glutamate ligase-like protein |

| | | | | | | | | | | |
|---|---|---|---|---|---|---|---|---|---|---|
| AVAEIV_RS21055 | 4477374 | 4477386 | 163 | + | 15.8 | 1.22E-06 | 0.0649 | CCCATCGCCAGCA | - | CDP-6-deoxy-delta-3,4-glucoseen reductase |
| AVAEIV_RS15935 | 3475028 | 3475040 | 173 | - | 15.7571 | 1.25E-06 | 0.0563 | CACTTCGCCATCG | - | NADPH-dependent F420 reductase |
| AVAEIV_RS21815 | 4645984 | 4645996 | 109 | - | 15.7571 | 1.25E-06 | 0.0563 | CACTTCGCCATCG | - | BON domain-containing protein |
| AVAEIV_RS05415 | 1120250 | 1120262 | 283 | - | 15.7429 | 1.26E-06 | 0.0563 | CGCATCGCCAGCA | - | DUF2790 domain-containing protein |
| AVAEIV_RS25195 | 5365157 | 5365169 | 385 | - | 15.7429 | 1.26E-06 | 0.0649 | CGCATCGCCAGCA | *hybE* | [NiFe]-hydrogenase assembly chaperone HybE |
| AVAEIV_RS25200 | 5365157 | 5365169 | 160 | - | 15.7429 | 1.26E-06 | 0.0649 | CGCATCGCCAGCA | - | rubredoxin |
| AVAEIV_RS18285 | 3940707 | 3940719 | 219 | - | 15.6714 | 1.37E-06 | 0.0649 | CACATGGCCATCG | - | hypothetical protein |
| AVAEIV_RS18290 | 3940707 | 3940719 | 31 | - | 15.6714 | 1.37E-06 | 0.0649 | CACATGGCCATCG | - | hypothetical protein |
| AVAEIV_RS01965 | 394540 | 394552 | 297 | + | 15.6143 | 1.42E-06 | 0.0649 | CCGATCGCCAGCG | - | hypothetical protein |
| AVAEIV_RS01975 | 394540 | 394552 | 260 | + | 15.6143 | 1.42E-06 | 0.0563 | CCGATCGCCAGCG | - | helix-turn-helix transcriptional regulator |
| AVAEIV_RS18495 | 3982240 | 3982252 | 41 | + | 15.6143 | 1.42E-06 | 0.0649 | CCGAACGCCAGCA | - | ACP phosphodiesterase |
| AVAEIV_RS18500 | 3982240 | 3982252 | 66 | + | 15.6143 | 1.42E-06 | 0.0563 | CCGAACGCCAGCA | - | ArsR/SmtB family transcription factor |
| AVAEIV_RS05765 | 1181966 | 1181978 | 307 | - | 15.5857 | 1.48E-06 | 0.0649 | CAGCTCGCCAGCG | - | NAD(P)H-dependent flavin oxidoreductase |
| AVAEIV_RS10835 | 2275049 | 2275061 | 262 | - | 15.5857 | 1.48E-06 | 0.0649 | CAGCTCGCCAGCG | - | RAMP superfamily CRISPR-associated protein |
| AVAEIV_RS00865 | 169088 | 169100 | -26 | + | 15.5714 | 1.52E-06 | 0.0649 | CAGAGGGCCAGCA | - | AEC family transporter |
| AVAEIV_RS03700 | 745557 | 745569 | 374 | - | 15.5571 | 1.54E-06 | 0.0563 | CGGATCGCCAGCG | *miaA* | MiaA |
| AVAEIV_RS07780 | 1619586 | 1619598 | 296 | + | 15.5571 | 1.54E-06 | 0.0563 | CACTTCGCCACCA | - | flavin reductase family protein |
| AVAEIV_RS16935 | 3668561 | 3668573 | 64 | + | 15.5571 | 1.54E-06 | 0.0649 | CGGATCGCCAGCG | - | cytochrome c |
| AVAEIV_RS24680 | 5244170 | 5244182 | 253 | + | 15.5571 | 1.54E-06 | 0.0563 | CGGAACGCCAGCA | - | OsmC family protein |
| AVAEIV_RS04530 | 927810 | 927822 | 269 | + | 15.5143 | 1.57E-06 | 0.0649 | CACAGCGCCAGCA | - | dicarboxylate/amino acid:cation symporter |
| AVAEIV_RS10470 | 2188848 | 2188860 | -22 | - | 15.5143 | 1.57E-06 | 0.0649 | CGCTTCGCCATCA | - | GNAT family N-acetyltransferase |
| AVAEIV_RS10475 | 2188848 | 2188860 | 314 | - | 15.5143 | 1.57E-06 | 0.0563 | CGCTTCGCCATCA | - | IS5 family transposase |
| AVAEIV_RS15660 | 3428335 | 3428347 | 67 | + | 15.5143 | 1.57E-06 | 0.0649 | CACAGCGCCAGCA | - | MinD/ParA family protein |
| AVAEIV_RS15665 | 3428335 | 3428347 | 79 | + | 15.5143 | 1.57E-06 | 0.0563 | CACAGCGCCAGCA | *sodC* | superoxide dismutase family protein |
| AVAEIV_RS18310 | 3942846 | 3942858 | 185 | - | 15.5143 | 1.57E-06 | 0.0649 | CACAGCGCCAGCA | - | hypothetical protein |
| AVAEIV_RS07270 | 1508040 | 1508052 | 329 | + | 15.5 | 1.61E-06 | 0.0649 | CAAATTGCCAGCA | *argF* | ornithine carbamoyltransferase |
| AVAEIV_RS07275 | 1508040 | 1508052 | -40 | + | 15.5 | 1.61E-06 | 0.0563 | CAAATTGCCAGCA | *grxD* | Grx4 family monothiol glutaredoxin |
| AVAEIV_RS07410 | 1539355 | 1539367 | 119 | - | 15.5 | 1.61E-06 | 0.0563 | CAGCTTGCCAGCA | - | protein |
| AVAEIV_RS24585 | 5222559 | 5222571 | 254 | + | 15.4857 | 1.67E-06 | 0.0563 | CTGATCGCCAGCA | *aguA* | agmatine deiminase |
| AVAEIV_RS06615 | 1364731 | 1364743 | 239 | + | 15.3857 | 1.79E-06 | 0.0563 | CCGTTCGCCATCG | *mreD* | rod shape-determining protein MreD |
| AVAEIV_RS24750 | 5258485 | 5258497 | 182 | + | 15.3857 | 1.79E-06 | 0.0563 | CCGTTCGCCATCG | *araD1* | AraD1 family protein |
| AVAEIV_RS21495 | 4574486 | 4574498 | 357 | + | 15.3714 | 1.82E-06 | 0.0649 | CAGTTCGCCACCG | *dapB* | 4-hydroxy-tetrahydrodipicolinate reductase |
| AVAEIV_RS04345 | 884433 | 884445 | 96 | - | 15.3286 | 1.94E-06 | 0.0649 | CAGAGCGCCAGCG | *leuS* | leucine--tRNA ligase |
| AVAEIV_RS04350 | 884433 | 884445 | 88 | - | 15.3286 | 1.94E-06 | 0.0563 | CAGAGCGCCAGCG | - | hypothetical protein |
| AVAEIV_RS16050 | 3497035 | 3497047 | 46 | - | 15.3286 | 1.94E-06 | 0.0563 | CAGAGCGCCAGCG | - | gallate dioxygenase |
| AVAEIV_RS10015 | 2097080 | 2097092 | 181 | + | 15.3 | 1.97E-06 | 0.0563 | CCGATGGCCATCG | - | K+/H+ antiporter subunit F |
| AVAEIV_RS07725 | 1609614 | 1609626 | -11 | - | 15.2714 | 2.07E-06 | 0.0563 | CAGCAGGCCATCA | *pabC* | aminodeoxychorismate lyase |
| AVAEIV_RS20910 | 4445615 | 4445627 | -34 | + | 15.2714 | 2.07E-06 | 0.0563 | CAGCAGGCCATCA | - | alpha-ketoacid dehydrogenase subunit beta |
| AVAEIV_RS02375 | 472892 | 472904 | 350 | - | 15.2571 | 2.11E-06 | 0.0649 | CTGTTCGCCATCA | - | LysR family transcriptional regulator |
| AVAEIV_RS07510 | 1566888 | 1566900 | 215 | + | 15.2571 | 2.11E-06 | 0.0649 | CAGGTCGCCATCG | *sulA* | SOS-induced cell division inhibitor SulA |
| AVAEIV_RS10000 | 2094973 | 2094985 | -22 | + | 15.2571 | 2.11E-06 | 0.0563 | CTGTTCGCCATCA | - | Na+/H+ antiporter subunit C |
| AVAEIV_RS10005 | 2094973 | 2094985 | 304 | + | 15.2571 | 2.11E-06 | 0.0563 | CTGTTCGCCATCA | - | monovalent cation/H+ antiporter subunit D |
| AVAEIV_RS16450 | 3571810 | 3571822 | 100 | - | 15.2571 | 2.11E-06 | 0.0563 | CAGGTCGCCATCG | - | flavodoxin |
| AVAEIV_RS15955 | 3478087 | 3478099 | 90 | + | 15.2429 | 2.19E-06 | 0.0563 | CAGTTGGCCAGCG | - | helix-turn-helix domain-containing protein |
| AVAEIV_RS16575 | 3594523 | 3594535 | 343 | + | 15.2429 | 2.19E-06 | 0.0563 | CAGTTGGCCAGCG | - | YbaN family protein |
| AVAEIV_RS02995 | 618258 | 618270 | 12 | + | 15.2286 | 2.22E-06 | 0.0563 | CATATCGCCAGCA | - | lysophospholipid acyltransferase family protein |
| AVAEIV_RS03115 | 641689 | 641701 | 47 | - | 15.2286 | 2.22E-06 | 0.0563 | CACATCGCCACCG | *rplD* | 50S ribosomal protein L4 |
| AVAEIV_RS24085 | 5125339 | 5125351 | 141 | + | 15.2286 | 2.22E-06 | 0.0649 | CACAACGCCACCA | - | LysR family transcriptional regulator |
| AVAEIV_RS24090 | 5125339 | 5125351 | -40 | + | 15.2286 | 2.22E-06 | 0.0563 | CACAACGCCACCA | - | DoxX family protein |
| AVAEIV_RS23175 | 4948980 | 4948992 | 194 | - | 15.2143 | 2.24E-06 | 0.0563 | CACCTCGCCATCG | - | hypothetical protein |
| AVAEIV_RS09875 | 2068094 | 2068106 | 97 | - | 15.1857 | 2.30E-06 | 0.0649 | CGCATCGCCATCG | *rsxB* | electron transport complex subunit RsxB |
| AVAEIV_RS11800 | 2498092 | 2498104 | 101 | - | 15.1857 | 2.30E-06 | 0.0563 | CAGTTCGCCGTCA | *prpB* | methylisocitrate lyase |

| | | | | | | | | | | |
|---|---|---|---|---|---|---|---|---|---|---|
| AVAEIV_RS14280 | 3109001 | 3109013 | 42 | - | 15.1857 | 2.30E-06 | 0.0563 | CGCATCGCCATCG | - | HigA family addiction module antitoxin |
| AVAEIV_RS16990 | 3683219 | 3683231 | 84 | + | 15.1857 | 2.30E-06 | 0.0649 | CGCATCGCCATCG | - | LysR family transcriptional regulator |
| AVAEIV_RS16995 | 3683219 | 3683231 | 32 | + | 15.1857 | 2.30E-06 | 0.0563 | CGCATCGCCATCG | - | YbfB/YjiJ family MFS transporter |
| AVAEIV_RS23975 | 5107245 | 5107257 | 370 | - | 15.1857 | 2.30E-06 | 0.0649 | CGCATCGCCATCG | - | uroporphyrinogen-III synthase |
| AVAEIV_RS00655 | 133159 | 133171 | 295 | - | 15.1429 | 2.41E-06 | 0.0563 | CAGATCGCCAGCC | - | helix-turn-helix domain-containing protein |
| AVAEIV_RS04475 | 914001 | 914013 | 309 | + | 15.1429 | 2.41E-06 | 0.0649 | CCGAGCGCCAGCA | - | IucA/IucC family protein |
| AVAEIV_RS07725 | 1609298 | 1609310 | 305 | - | 15.1429 | 2.41E-06 | 0.0563 | CAGATCGCCAGCC | *pabC* | aminodeoxychorismate lyase |
| AVAEIV_RS14880 | 3236922 | 3236934 | -41 | + | 15.1429 | 2.41E-06 | 0.0649 | CCGAGCGCCAGCA | - | fatty acid cis/trans isomerase |
| AVAEIV_RS19085 | 4113627 | 4113639 | 118 | + | 15.1429 | 2.41E-06 | 0.0649 | CCGAGCGCCAGCA | - | Na+/H+ antiporter |
| AVAEIV_RS22285 | 4758023 | 4758035 | 75 | + | 15.1429 | 2.41E-06 | 0.0649 | CCGAGCGCCAGCA | - | GNAT family N-acetyltransferase |
| AVAEIV_RS24045 | 5117245 | 5117257 | 50 | + | 15.1429 | 2.41E-06 | 0.0563 | CAGATCGCCAGCC | - | CPBP family intramembrane glutamic endopeptidase |
| AVAEIV_RS13235 | 2871150 | 2871162 | 45 | - | 15.1286 | 2.43E-06 | 0.0563 | CGGTTCGCCACCA | - | DUF1016 N-terminal domain-containing protein |
| AVAEIV_RS04585 | 938424 | 938436 | 178 | + | 15.1143 | 2.48E-06 | 0.0649 | CAGCGCGCCAGCA | - | urease subunit gamma |
| AVAEIV_RS11405 | 2413507 | 2413519 | 161 | - | 15.1143 | 2.48E-06 | 0.0563 | CAGCGCGCCAGCA | - | GNAT family N-acetyltransferase |
| AVAEIV_RS12005 | 2551126 | 2551138 | 59 | - | 15.1143 | 2.48E-06 | 0.0649 | CAGTCCGCCATCA | *phaC* | class I poly(R)-hydroxyalkanoic acid synthase |
| AVAEIV_RS25100 | 5345348 | 5345360 | 346 | + | 15.1143 | 2.48E-06 | 0.0649 | CAGCGCGCCAGCA | - | ABC transporter permease |
| AVAEIV_RS00860 | 168286 | 168298 | 243 | + | 15.1 | 2.56E-06 | 0.0649 | CAGATGGCCGTCA | - | Na/Pi cotransporter family protein |
| AVAEIV_RS04450 | 907597 | 907609 | 200 | + | 15.1 | 2.56E-06 | 0.0649 | CCGATGGCCACCA | *hemL* | glutamate-1-semialdehyde 2,1-aminomutase |
| AVAEIV_RS07120 | 1467341 | 1467353 | 373 | + | 15.1 | 2.56E-06 | 0.0563 | CGCATTGCCATCA | - | sigma-E factor negative regulatory protein |
| AVAEIV_RS18685 | 4024227 | 4024239 | -33 | + | 15.1 | 2.56E-06 | 0.0649 | CAGTGCGCCATCG | *pcaD* | 3-oxoadipate enol-lactonase |
| AVAEIV_RS19755 | 4209449 | 4209461 | -46 | - | 15.1 | 2.56E-06 | 0.0649 | CACATGGCCAGCG | - | TetR/AcrR family transcriptional regulator |
| AVAEIV_RS04535 | 928570 | 928582 | 44 | + | 15.0857 | 2.58E-06 | 0.0649 | CCGCTGGCCATCA | - | alpha/beta fold hydrolase |
| AVAEIV_RS04540 | 928570 | 928582 | 203 | + | 15.0857 | 2.58E-06 | 0.0563 | CCGCTGGCCATCA | - | LysR substrate-binding domain-containing protein |
| AVAEIV_RS24165 | 5137251 | 5137263 | 353 | - | 15.0857 | 2.58E-06 | 0.0563 | CCGCTGGCCATCA | - | Lrp/AsnC ligand binding domain-containing protein |
| AVAEIV_RS25095 | 5344091 | 5344103 | 162 | + | 15.0857 | 2.58E-06 | 0.0649 | CCGCTGGCCATCA | - | Gfo/Idh/MocA family protein |
| AVAEIV_RS18585 | 4003186 | 4003198 | 307 | - | 15.0571 | 2.65E-06 | 0.0568 | CCGTTGGCCAGCA | - | O-methyltransferase |
| AVAEIV_RS20725 | 4406300 | 4406312 | 162 | + | 15.0571 | 2.65E-06 | 0.0649 | CCGTTGGCCAGCA | *prfA* | peptide chain release factor 1 |
| AVAEIV_RS20580 | 4375137 | 4375149 | 257 | + | 15.0429 | 2.70E-06 | 0.0568 | CCCATCGCCACCA | - | thermonuclease family protein |
| AVAEIV_RS13860 | 3005827 | 3005839 | 376 | - | 15.0286 | 2.77E-06 | 0.0649 | CAGCACGCCATCG | - | DotU family type VI secretion system protein |
| AVAEIV_RS16305 | 3549713 | 3549725 | 377 | - | 15.0286 | 2.77E-06 | 0.0649 | CAGCACGCCATCG | - | antitoxin |
| AVAEIV_RS19385 | 4158096 | 4158108 | 82 | + | 15.0286 | 2.77E-06 | 0.0649 | CAGCACGCCATCG | - | tRNA-Met |
| AVAEIV_RS19725 | 4204941 | 4204953 | 101 | + | 15.0286 | 2.77E-06 | 0.0649 | CAGCACGCCATCG | - | O-acetyl-ADP-ribose deacetylase |
| AVAEIV_RS06020 | 1226736 | 1226748 | 180 | - | 15.0143 | 2.81E-06 | 0.0649 | CAGAGGGCCATCG | - | methyl-accepting chemotaxis protein |
| AVAEIV_RS06025 | 1226736 | 1226748 | 277 | - | 15.0143 | 2.81E-06 | 0.0568 | CAGAGGGCCATCG | - | containing protein |
| AVAEIV_RS05465 | 1130372 | 1130384 | 243 | - | 15 | 2.86E-06 | 0.0568 | CGGTTGGCCAGCA | *pilV* | type IV pilus modification protein PilV |
| AVAEIV_RS15245 | 3329817 | 3329829 | 60 | + | 15 | 2.86E-06 | 0.0649 | CGGTTGGCCAGCA | *dnaQ* | DNA polymerase III subunit epsilon |
| AVAEIV_RS26100 | 5612316 | 5612328 | -40 | - | 15 | 2.86E-06 | 0.0649 | CCCTTCGCCAGCA | - | tyrosine-type recombinase/integrase |
| AVAEIV_RS01865 | 369442 | 369454 | 349 | + | 14.9857 | 2.87E-06 | 0.0568 | CGCATCGCCACCA | - | FecR family protein |
| AVAEIV_RS08015 | 1666206 | 1666218 | 356 | - | 14.9857 | 2.87E-06 | 0.0649 | CGCATCGCCACCA | - | TrkH family potassium uptake protein |
| AVAEIV_RS08025 | 1666206 | 1666218 | 276 | - | 14.9857 | 2.87E-06 | 0.0568 | CGCATCGCCACCA | - | NAD(P)H nitroreductase |
| AVAEIV_RS16875 | 3653816 | 3653828 | 240 | + | 14.9571 | 2.94E-06 | 0.0568 | CAGAATGCCACCA | - | hypothetical protein |
| AVAEIV_RS05635 | 1158398 | 1158410 | 31 | - | 14.9429 | 3.01E-06 | 0.0568 | CAAAATGCCATCA | - | hypothetical protein |
| AVAEIV_RS21160 | 4500238 | 4500250 | 373 | - | 14.9429 | 3.01E-06 | 0.0649 | CGCTTCGCCAGCA | - | amidohydrolase |
| AVAEIV_RS11580 | 2448221 | 2448233 | 235 | + | 14.9286 | 3.04E-06 | 0.0568 | CTGATCGCCATCG | - | Dps family protein |
| AVAEIV_RS19165 | 4132363 | 4132375 | 261 | - | 14.9286 | 3.04E-06 | 0.0649 | CTGATCGCCATCG | - | phage minor tail protein L |
| AVAEIV_RS20825 | 4426160 | 4426172 | 26 | - | 14.9286 | 3.04E-06 | 0.0568 | CTGATCGCCATCG | - | HNH endonuclease |
| AVAEIV_RS22045 | 4705148 | 4705160 | 369 | - | 14.9286 | 3.04E-06 | 0.0649 | CTGATCGCCATCG | - | DUF2955 domain-containing protein |
| AVAEIV_RS01005 | 198653 | 198665 | 363 | + | 14.9143 | 3.14E-06 | 0.0649 | CAGTTCGCCATCC | *zwf* | glucose-6-phosphate dehydrogenase |
| AVAEIV_RS01015 | 199155 | 199167 | 295 | - | 14.9143 | 3.14E-06 | 0.0568 | CCCATGGCCAGCA | - | PA3496 family putative envelope integrity protein |
| AVAEIV_RS11415 | 2416200 | 2416212 | 229 | + | 14.9143 | 3.14E-06 | 0.0649 | CAGAAGGCCAGCG | - | containing protein |
| AVAEIV_RS12300 | 2603494 | 2603506 | 206 | + | 14.9143 | 3.14E-06 | 0.0568 | CAGAAGGCCAGCG | *fliD* | flagellar filament capping protein FliD |
| AVAEIV_RS19680 | 4196287 | 4196299 | 313 | + | 14.9143 | 3.14E-06 | 0.0649 | CGGAATGCCATCA | - | ABC transporter ATP-binding protein |

| | | | | | | | | | | |
|---|---|---|---|---|---|---|---|---|---|---|
| AVAEIV_RS19685 | 4196287 | 4196299 | -1 | + | 14.9143 | 3.14E-06 | 0.0568 | CGGAATGCCATCA | - | DUF4197 domain-containing protein |
| AVAEIV_RS08445 | 1764325 | 1764337 | 188 | - | 14.8571 | 3.27E-06 | 0.0569 | CAGATCGCCGTCG | - | LysR family transcriptional regulator |
| AVAEIV_RS10030 | 2098421 | 2098433 | 299 | + | 14.8571 | 3.27E-06 | 0.0569 | CAGATCGCCGTCG | - | DUF5064 family protein |
| AVAEIV_RS10770 | 2257153 | 2257165 | 191 | - | 14.8571 | 3.27E-06 | 0.0649 | CGCATGGCCAGCA | - | hypothetical protein |
| AVAEIV_RS15665 | 3428130 | 3428142 | 284 | + | 14.8571 | 3.27E-06 | 0.0569 | CAGATCGCCGTCG | *sodC* | superoxide dismutase family protein |
| AVAEIV_RS20685 | 4398117 | 4398129 | 303 | + | 14.8571 | 3.27E-06 | 0.0649 | CGCATGGCCAGCA | - | TIGR01777 family oxidoreductase |
| AVAEIV_RS22720 | 4864555 | 4864567 | 214 | + | 14.8571 | 3.27E-06 | 0.0649 | CCGAACGCCACCA | - | class 1 fructose-bisphosphatase |
| AVAEIV_RS23215 | 4955231 | 4955243 | 251 | + | 14.8429 | 3.33E-06 | 0.0571 | CCAATCGCCATCG | - | hypothetical protein |
| AVAEIV_RS16765 | 3629273 | 3629285 | -34 | - | 14.8286 | 3.39E-06 | 0.0649 | CAGCTCGCCACCG | - | 3-deoxy-7-phosphoheptulonate synthase |
| AVAEIV_RS24305 | 5168277 | 5168289 | 176 | - | 14.8143 | 3.43E-06 | 0.0649 | CCGTTCGCCAGCG | - | SdiA-regulated domain-containing protein |
| AVAEIV_RS01990 | 399007 | 399019 | 154 | - | 14.8 | 3.47E-06 | 0.0649 | CGGATCGCCACCG | - | sugar MFS transporter |
| AVAEIV_RS02370 | 471076 | 471088 | 75 | - | 14.7857 | 3.56E-06 | 0.0649 | CGGCTCGCCATCG | *trmL* | carboxymethylaminomethyluridine(34)-2'-O)-methyltransferase TrmL |
| AVAEIV_RS11635 | 2462705 | 2462717 | 348 | - | 14.7857 | 3.56E-06 | 0.0649 | CAGGGCGCCATCA | - | SdiA-regulated domain-containing protein |
| AVAEIV_RS23535 | 5016417 | 5016429 | 60 | + | 14.7857 | 3.56E-06 | 0.0649 | CAGACCGCCATCG | - | MFS transporter |
| AVAEIV_RS12235 | 2593015 | 2593027 | 128 | + | 14.7714 | 3.64E-06 | 0.0649 | CCCAGCGCCATCA | *fliQ* | flagellar biosynthesis protein FliQ |
| AVAEIV_RS14260 | 3106670 | 3106682 | 287 | + | 14.7714 | 3.64E-06 | 0.0649 | CACATCGCCATCC | - | flagellar motor protein MotB |
| AVAEIV_RS03485 | 702062 | 702074 | 68 | + | 14.7571 | 3.69E-06 | 0.0601 | CACAGCGCCACCA | - | translation initiation factor Sui1 |
| AVAEIV_RS09470 | 1973719 | 1973731 | -48 | - | 14.7429 | 3.74E-06 | 0.0601 | CAGATCGCCATGA | - | HU family DNA-binding protein |
| AVAEIV_RS05160 | 1060377 | 1060389 | 271 | - | 14.7286 | 3.79E-06 | 0.0649 | CCGAAGGCCAGCA | - | DUF2231 domain-containing protein |
| AVAEIV_RS08975 | 1866730 | 1866742 | 264 | - | 14.7286 | 3.79E-06 | 0.0601 | CCGAAGGCCAGCA | - | ParA family protein |
| AVAEIV_RS20040 | 4274617 | 4274629 | 307 | - | 14.7286 | 3.79E-06 | 0.0601 | CTGATCGCCACCA | - | glycerol dehydrogenase |
| AVAEIV_RS15310 | 3344897 | 3344909 | -43 | - | 14.7143 | 3.86E-06 | 0.0649 | CTGCTCGCCATCA | - | DUF599 domain-containing protein |
| AVAEIV_RS24405 | 5186423 | 5186435 | 271 | - | 14.7 | 3.95E-06 | 0.0601 | CGGCTTGCCATCA | *phoR* | phosphate regulon sensor histidine kinase PhoR |
| AVAEIV_RS05885 | 1204848 | 1204860 | 113 | + | 14.6857 | 4.01E-06 | 0.0649 | CCCTTGGCCATCA | - | hypothetical protein |
| AVAEIV_RS22170 | 4730871 | 4730883 | 48 | - | 14.6857 | 4.01E-06 | 0.0601 | CAGGTCGCCAGCG | - | hypothetical protein |
| AVAEIV_RS02580 | 524012 | 524024 | 365 | + | 14.6714 | 4.11E-06 | 0.0649 | CCGATCGCCGTCA | - | hypothetical protein |
| AVAEIV_RS02585 | 524012 | 524024 | 9 | + | 14.6714 | 4.11E-06 | 0.0601 | CCGATCGCCGTCA | *ompR* | two-component system response regulator OmpR |
| AVAEIV_RS19945 | 4251754 | 4251766 | 226 | - | 14.6714 | 4.11E-06 | 0.0601 | CGGATGGCCAGCG | - | peptidoglycan DD-metalloendopeptidase family protein |
| AVAEIV_RS02285 | 456357 | 456369 | 97 | - | 14.6571 | 4.12E-06 | 0.0649 | CAGATCGCCGCCA | *mutY* | A/G-specific adenine glycosylase |
| AVAEIV_RS08650 | 1804587 | 1804599 | 258 | - | 14.6571 | 4.12E-06 | 0.0649 | CAGATCGCCGCCA | - | ABC-2 transporter permease |
| AVAEIV_RS09560 | 1992634 | 1992646 | 52 | + | 14.6571 | 4.12E-06 | 0.0601 | CAGATCGCCGCCA | - | SlyX family protein |
| AVAEIV_RS23950 | 5102434 | 5102446 | 143 | - | 14.6571 | 4.12E-06 | 0.0649 | CAGATCGCCGCCA | - | AlgP family protein |
| AVAEIV_RS03150 | 645208 | 645220 | 25 | + | 14.6429 | 4.19E-06 | 0.0601 | CCGCTCGCCACCA | *rpmC* | 50S ribosomal protein L29 |
| AVAEIV_RS03155 | 645208 | 645220 | 219 | + | 14.6429 | 4.19E-06 | 0.0601 | CCGCTCGCCACCA | *rpsQ* | 30S ribosomal protein S17 |
| AVAEIV_RS23430 | 4995918 | 4995930 | 319 | - | 14.6429 | 4.19E-06 | 0.0649 | CACCACGCCAGCA | - | tetratricopeptide repeat protein |
| AVAEIV_RS24520 | 5211318 | 5211330 | 190 | - | 14.6429 | 4.19E-06 | 0.0649 | CCAATCGCCACCA | - | polyamine ABC transporter substrate-binding protein |
| AVAEIV_RS04430 | 901134 | 901146 | 309 | - | 14.6286 | 4.23E-06 | 0.0601 | CACAGGGCCAGCA | *rhmD* | L-rhamnonate dehydratase |
| AVAEIV_RS11250 | 2375306 | 2375318 | 315 | + | 14.6286 | 4.23E-06 | 0.0601 | CACAGGGCCAGCA | - | Crp/Fnr family transcriptional regulator |
| AVAEIV_RS14535 | 3165644 | 3165656 | 381 | + | 14.6286 | 4.23E-06 | 0.0649 | CACAGGGCCAGCA | *cysG* | siroheme synthase CysG |
| AVAEIV_RS10190 | 2126961 | 2126973 | 223 | - | 14.6143 | 4.27E-06 | 0.0649 | CGCATCGCCAGCG | *hemN* | oxygen-independent coproporphyrinogen III oxidase |
| AVAEIV_RS13500 | 2919337 | 2919349 | 238 | + | 14.6143 | 4.27E-06 | 0.0601 | CGCATCGCCAGCG | - | 3-oxoadipyl-CoA thiolase |
| AVAEIV_RS14405 | 3136927 | 3136939 | 129 | - | 14.6143 | 4.27E-06 | 0.0649 | CGCATCGCCAGCG | *flhB* | flagellar biosynthesis protein FlhB |
| AVAEIV_RS09200 | 1920003 | 1920015 | 324 | + | 14.6 | 4.33E-06 | 0.0601 | CTGATGGCCAGCA | - | DUF2334 domain-containing protein |
| AVAEIV_RS03435 | 693591 | 693603 | -31 | - | 14.5714 | 4.45E-06 | 0.0649 | CGACTCGCCATCA | *prmA* | 50S ribosomal protein L11 methyltransferase |
| AVAEIV_RS00195 | 40275 | 40287 | 330 | - | 14.5571 | 4.49E-06 | 0.0649 | CAGCGCGCCATCG | - | SMI1/KNR4 family protein |
| AVAEIV_RS25790 | 5526288 | 5526300 | -40 | + | 14.5571 | 4.49E-06 | 0.0601 | CAGCGCGCCATCG | - | glycoside hydrolase family 32 protein |
| AVAEIV_RS21795 | 4642003 | 4642015 | 363 | + | 14.5429 | 4.56E-06 | 0.0649 | CAGTCCGCCAGCA | - | sugar ABC transporter substrate-binding protein |
| AVAEIV_RS21800 | 4642003 | 4642015 | 155 | + | 14.5429 | 4.56E-06 | 0.0601 | CAGTCCGCCAGCA | - | acyl-CoA dehydrogenase family protein |
| AVAEIV_RS06660 | 1377260 | 1377272 | 269 | + | 14.5286 | 4.61E-06 | 0.0649 | CGGAGCGCCATCG | *rapZ* | RNase adapter RapZ |
| AVAEIV_RS18080 | 3907544 | 3907556 | 357 | + | 14.5286 | 4.61E-06 | 0.0649 | CGCATTGCCAGCA | *queE* | 7-carboxy-7-deazaguanine synthase QueE |
| AVAEIV_RS03955 | 797669 | 797681 | 193 | + | 14.5143 | 4.63E-06 | 0.0649 | CCGCTGGCCAGCA | - | MotA/TolQ/ExbB proton channel family protein |

| | | | | | | | | | | |
|---|---|---|---|---|---|---|---|---|---|---|
| AVAEIV_RS04585 | 938454 | 938466 | 208 | + | 14.5143 | 4.63E-06 | 0.0649 | CCGCTGGCCAGCA | - | urease subunit gamma |
| AVAEIV_RS11165 | 2358046 | 2358058 | 193 | + | 14.5143 | 4.63E-06 | 0.0649 | CCGCTGGCCAGCA | - | MotA/TolQ/ExbB proton channel family protein |
| AVAEIV_RS13285 | 2877290 | 2877302 | 362 | - | 14.5143 | 4.63E-06 | 0.0601 | CCGCTGGCCAGCA | *ltrA* | group II intron reverse transcriptase/maturase |
| AVAEIV_RS13775 | 2981511 | 2981523 | 366 | + | 14.5143 | 4.63E-06 | 0.0649 | CCGCTGGCCAGCA | - | DUF4150 domain-containing protein |
| AVAEIV_RS26100 | 5612370 | 5612382 | 14 | - | 14.5143 | 4.63E-06 | 0.0649 | CCGCTGGCCAGCA | - | tyrosine-type recombinase/integrase |
| AVAEIV_RS00575 | 118455 | 118467 | 84 | - | 14.5 | 4.68E-06 | 0.0601 | CCGGTCGCCAGCA | - | hypothetical protein |
| AVAEIV_RS21225 | 4512902 | 4512914 | -47 | - | 14.5 | 4.68E-06 | 0.0649 | CCGAGTGCCATCA | - | paraquat-inducible protein A |
| AVAEIV_RS01365 | 268123 | 268135 | 227 | - | 14.4857 | 4.74E-06 | 0.0649 | CAGTTGGCCACCG | *spoT* | bifunctional GTP diphosphokinase/guanosine-3',5'-bis pyrophosphate 3'-pyrophosphohydrolase |
| AVAEIV_RS10150 | 2119260 | 2119272 | 113 | + | 14.4857 | 4.74E-06 | 0.0601 | CCGAACGCCAGCG | - | YbaB/EbfC family nucleoid-associated protein |
| AVAEIV_RS05550 | 1149808 | 1149820 | 86 | + | 14.4714 | 4.82E-06 | 0.0649 | CAAAGTGCCATCA | *mobH* | MobH family relaxase |
| AVAEIV_RS05555 | 1149808 | 1149820 | 107 | + | 14.4714 | 4.82E-06 | 0.0601 | CAAAGTGCCATCA | - | hypothetical protein |
| AVAEIV_RS05560 | 1149808 | 1149820 | 372 | + | 14.4714 | 4.82E-06 | 0.0601 | CAAAGTGCCATCA | - | hypothetical protein |
| AVAEIV_RS18050 | 3902046 | 3902058 | 372 | - | 14.4714 | 4.82E-06 | 0.0649 | CAAAGTGCCATCA | - | hypothetical protein |
| AVAEIV_RS18055 | 3902046 | 3902058 | 107 | - | 14.4714 | 4.82E-06 | 0.0649 | CAAAGTGCCATCA | - | hypothetical protein |
| AVAEIV_RS18060 | 3902046 | 3902058 | 86 | - | 14.4714 | 4.82E-06 | 0.0601 | CAAAGTGCCATCA | *mobH* | MobH family relaxase |
| AVAEIV_RS02060 | 409366 | 409378 | 117 | + | 14.4571 | 4.90E-06 | 0.0601 | CTGAGCGCCATCA | - | TIGR04255 family protein |
| AVAEIV_RS02280 | 455538 | 455550 | 363 | + | 14.4571 | 4.90E-06 | 0.0649 | CGGCTGGCCAGCA | - | oxidative damage protection protein |
| AVAEIV_RS06145 | 1253192 | 1253204 | 209 | + | 14.4571 | 4.90E-06 | 0.0649 | CGGCTGGCCAGCA | - | type 4 pilus major pilin |
| AVAEIV_RS07785 | 1620370 | 1620382 | 83 | - | 14.4571 | 4.90E-06 | 0.0601 | CGGCTGGCCAGCA | - | alpha/beta fold hydrolase |
| AVAEIV_RS12040 | 2558031 | 2558043 | 312 | + | 14.4571 | 4.90E-06 | 0.0649 | CAAAACGCCAGCG | - | Smr/MutS family protein |
| AVAEIV_RS15180 | 3304191 | 3304203 | 159 | - | 14.4571 | 4.90E-06 | 0.0601 | CGGCTGGCCAGCA | - | LLM class flavin-dependent oxidoreductase |
| AVAEIV_RS19690 | 4198088 | 4198100 | 321 | - | 14.4571 | 4.90E-06 | 0.0649 | CAGCACGCCAGCG | - | class I SAM-dependent methyltransferase |
| AVAEIV_RS01170 | 227889 | 227901 | 337 | + | 14.4429 | 4.97E-06 | 0.0649 | CCCTTCGCCATCG | - | alpha/beta fold hydrolase |
| AVAEIV_RS01320 | 257826 | 257838 | 49 | + | 14.4429 | 4.97E-06 | 0.0601 | CCCTTCGCCATCG | - | HesA/MoeB/ThiF family protein |
| AVAEIV_RS04795 | 974980 | 974992 | 318 | - | 14.4429 | 4.97E-06 | 0.0601 | CGGGTCGCCAGCA | *mdcE* | gamma |
| AVAEIV_RS00085 | 19827 | 19839 | 171 | + | 14.4 | 5.07E-06 | 0.0649 | CACACCGCCAGCA | *fmt* | methionyl-tRNA formyltransferase |
| AVAEIV_RS05035 | 1033623 | 1033635 | 278 | - | 14.3857 | 5.12E-06 | 0.0649 | CAGATCGCCACCC | - | alginate O-acetyltransferase |
| AVAEIV_RS06550 | 1352937 | 1352949 | 294 | - | 14.3857 | 5.12E-06 | 0.0649 | CGCTTCGCCATCG | - | ABC transporter substrate-binding protein |
| AVAEIV_RS08865 | 1841841 | 1841853 | 93 | - | 14.3857 | 5.12E-06 | 0.0601 | CACAGCGCCAGCG | *casB* | type I-E CRISPR-associated protein Cse2/CasB |
| AVAEIV_RS11310 | 2389126 | 2389138 | 87 | + | 14.3857 | 5.12E-06 | 0.0601 | CGCTTCGCCATCG | *tal* | transaldolase |
| AVAEIV_RS14120 | 3072272 | 3072284 | 87 | - | 14.3857 | 5.12E-06 | 0.0649 | CGCTTCGCCATCG | *tal* | transaldolase |
| AVAEIV_RS05680 | 1164469 | 1164481 | 323 | - | 14.3714 | 5.24E-06 | 0.0601 | CAGGTGGCCATCG | - | hypothetical protein |
| AVAEIV_RS19655 | 4190687 | 4190699 | -6 | - | 14.3714 | 5.24E-06 | 0.0601 | CAGCATGCCAGCA | - | putative RNA methyltransferase |
| AVAEIV_RS08880 | 1843888 | 1843900 | 378 | + | 14.3571 | 5.32E-06 | 0.0601 | CTGATCGCCAGCG | *cas6e* | type I-E CRISPR-associated protein Cas6/Cse3/CasE |
| AVAEIV_RS09870 | 2067453 | 2067465 | 16 | - | 14.3571 | 5.32E-06 | 0.0649 | CTGATCGCCAGCG | *rsxC* | electron transport complex subunit RsxC |
| AVAEIV_RS10360 | 2166181 | 2166193 | 184 | - | 14.3571 | 5.32E-06 | 0.0649 | CTGATCGCCAGCG | - | helix-turn-helix domain-containing protein |
| AVAEIV_RS13880 | 3007930 | 3007942 | 191 | - | 14.3571 | 5.32E-06 | 0.0601 | CTGAACGCCAGCA | *tagH* | protein TagH |
| AVAEIV_RS20280 | 4325623 | 4325635 | 59 | - | 14.3571 | 5.32E-06 | 0.0649 | CCCAAGGCCATCA | - | transposase |
| AVAEIV_RS21915 | 4672686 | 4672698 | -28 | - | 14.3571 | 5.32E-06 | 0.0649 | CTGATCGCCAGCG | - | HU family DNA-binding protein |
| AVAEIV_RS23095 | 4937804 | 4937816 | 243 | - | 14.3571 | 5.32E-06 | 0.0601 | CTGATCGCCAGCG | - | DUF1353 domain-containing protein |
| AVAEIV_RS25315 | 5395178 | 5395190 | 384 | + | 14.3571 | 5.32E-06 | 0.0649 | CTGATCGCCAGCG | - | domain-containing protein |
| AVAEIV_RS17340 | 3755225 | 3755237 | 35 | + | 14.3429 | 5.42E-06 | 0.0649 | CACAAGGCCACCA | *astA* | arginine N-succinyltransferase |
| AVAEIV_RS16740 | 3623929 | 3623941 | 349 | - | 14.3286 | 5.45E-06 | 0.0601 | CACCTGGCCATCG | - | glutamine amidotransferase |
| AVAEIV_RS20290 | 4326565 | 4326577 | 329 | + | 14.3286 | 5.45E-06 | 0.0649 | CACCTGGCCATCG | - | IS110 family transposase |
| AVAEIV_RS05935 | 1213736 | 1213748 | 317 | - | 14.3143 | 5.50E-06 | 0.0649 | CACGTCGCCATCG | - | TIGR03750 family conjugal transfer protein |
| AVAEIV_RS05940 | 1213736 | 1213748 | -40 | - | 14.3143 | 5.50E-06 | 0.0649 | CACGTCGCCATCG | - | membrane protein |
| AVAEIV_RS10070 | 2105711 | 2105723 | 328 | + | 14.3143 | 5.50E-06 | 0.0601 | CACGTCGCCATCG | - | F0F1 ATP synthase subunit epsilon |
| AVAEIV_RS09495 | 1978116 | 1978128 | 124 | - | 14.3 | 5.58E-06 | 0.0601 | CCGTTGGCCACCA | - | DUF2946 family protein |
| AVAEIV_RS19540 | 4174235 | 4174247 | 127 | + | 14.3 | 5.58E-06 | 0.0601 | CGCAAGGCCATCA | - | LytTR family DNA-binding domain-containing protein |
| AVAEIV_RS21780 | 4638403 | 4638415 | 281 | + | 14.3 | 5.58E-06 | 0.0649 | CACTTGGCCAGCG | - | ABC transporter permease |
| AVAEIV_RS03880 | 782426 | 782438 | 342 | + | 14.2857 | 5.64E-06 | 0.0649 | CAGAACGCCGGCA | *ispA* | (2E,6E)-farnesyl diphosphate synthase |

| | | | | | | | | | | |
|---|---|---|---|---|---|---|---|---|---|---|
| AVAEIV_RS03885 | 782426 | 782438 | 103 | + | 14.2857 | 5.64E-06 | 0.0649 | CAGAACGCCGGCA | - | exodeoxyribonuclease VII small subunit |
| AVAEIV_RS07735 | 1611113 | 1611125 | 385 | + | 14.2857 | 5.64E-06 | 0.0601 | CAGATCGCCGGCG | *tmk* | dTMP kinase |
| AVAEIV_RS12790 | 2718374 | 2718386 | 103 | + | 14.2857 | 5.64E-06 | 0.0601 | CAGAACGCCGGCA | - | formylglycine-generating enzyme family protein |
| AVAEIV_RS13805 | 2990839 | 2990851 | 31 | + | 14.2857 | 5.64E-06 | 0.0649 | CAGATCGCCGGCG | *tssH* | type VI secretion system ATPase TssH |
| AVAEIV_RS15210 | 3313068 | 3313080 | 379 | - | 14.2857 | 5.64E-06 | 0.0649 | CAGATCGCCGGCG | - | ABC transporter permease |
| AVAEIV_RS18860 | 4062020 | 4062032 | 220 | - | 14.2857 | 5.64E-06 | 0.0649 | CGGTGCGCCAGCA | *recA* | recombinase RecA |
| AVAEIV_RS19060 | 4106117 | 4106129 | 190 | - | 14.2857 | 5.64E-06 | 0.0649 | CAGAACGCCGGCA | *tsf* | translation elongation factor Ts |
| AVAEIV_RS23110 | 4938885 | 4938897 | 91 | + | 14.2857 | 5.64E-06 | 0.0601 | CAGATCGCCGGCG | - | DUF6890 family protein |
| AVAEIV_RS01350 | 265092 | 265104 | 373 | - | 14.2714 | 5.69E-06 | 0.0649 | CCGCACGCCAGCA | *vnfA1* | regulator VnfA1 |
| AVAEIV_RS01355 | 265092 | 265104 | 33 | - | 14.2714 | 5.69E-06 | 0.0649 | CCGCACGCCAGCA | - | NifU family protein |
| AVAEIV_RS18905 | 4072879 | 4072891 | 341 | + | 14.2714 | 5.69E-06 | 0.0649 | CCAAACGCCAGCA | - | protein |
| AVAEIV_RS21835 | 4652731 | 4652743 | -50 | + | 14.2571 | 5.72E-06 | 0.0649 | CCGAGGGCCAGCA | - | ABC transporter substrate-binding protein |
| AVAEIV_RS18320 | 3944065 | 3944077 | 167 | + | 14.2286 | 5.85E-06 | 0.0649 | CAGTCGGCCATCA | - | DUF6387 family protein |
| AVAEIV_RS22695 | 4860298 | 4860310 | 269 | - | 14.2286 | 5.85E-06 | 0.0601 | CTCATGGCCATCA | - | ketopantoate reductase family protein |
| AVAEIV_RS03685 | 741912 | 741924 | 102 | - | 14.2143 | 5.94E-06 | 0.0601 | CAGACCGCCAGCG | *tsaE* | threonylcarbamoyltransferase complex ATPase subunit type 1 TsaE |
| AVAEIV_RS05940 | 1213988 | 1214000 | 212 | - | 14.2143 | 5.94E-06 | 0.0649 | CAGACCGCCAGCG | - | membrane protein |
| AVAEIV_RS05945 | 1213988 | 1214000 | -46 | - | 14.2143 | 5.94E-06 | 0.0649 | CAGACCGCCAGCG | - | protein |
| AVAEIV_RS11420 | 2417221 | 2417233 | 230 | + | 14.2143 | 5.94E-06 | 0.0649 | CAGGGCGCCAGCA | - | TauD/TfdA dioxygenase family protein |
| AVAEIV_RS12300 | 2603416 | 2603428 | 284 | + | 14.2143 | 5.94E-06 | 0.0601 | CAGACCGCCAGCG | *fliD* | flagellar filament capping protein FliD |
| AVAEIV_RS16115 | 3511950 | 3511962 | 226 | + | 14.2143 | 5.94E-06 | 0.0601 | CAGTGGGCCATCG | - | ABC transporter substrate-binding protein |
| AVAEIV_RS24775 | 5267689 | 5267701 | 377 | + | 14.2143 | 5.94E-06 | 0.0649 | CAGGGCGCCAGCA | - | cation:dicarboxylate symporter family transporter |
| AVAEIV_RS00285 | 62049 | 62061 | 123 | + | 14.2 | 6.00E-06 | 0.0649 | CCCAGCGCCAGCA | - | ABC transporter permease |
| AVAEIV_RS08480 | 1772167 | 1772179 | 152 | + | 14.2 | 6.00E-06 | 0.0601 | CAGATTGCCGGCA | - | DUF2933 domain-containing protein |
| AVAEIV_RS18250 | 3936605 | 3936617 | 143 | - | 14.2 | 6.00E-06 | 0.0649 | CCCAGCGCCAGCA | - | hypothetical protein |
| AVAEIV_RS19300 | 4147123 | 4147135 | 306 | + | 14.2 | 6.00E-06 | 0.0649 | CACATCGCCAGCC | - | hypothetical protein |
| AVAEIV_RS24430 | 5194273 | 5194285 | 287 | + | 14.1857 | 6.04E-06 | 0.0649 | CCGGTGGCCATCA | *pstA* | phosphate ABC transporter permease PstA |
| AVAEIV_RS07850 | 1634274 | 1634286 | 320 | - | 14.1714 | 6.10E-06 | 0.0649 | CCGAAGGCCATCG | - | DUF2897 family protein |
| AVAEIV_RS03545 | 715306 | 715318 | 131 | - | 14.1571 | 6.19E-06 | 0.0601 | CACTGCGCCATCG | - | YidH family protein |
| AVAEIV_RS15870 | 3462023 | 3462035 | 0 | - | 14.1571 | 6.19E-06 | 0.0601 | CAGAAGGCCACCG | - | ABC transporter substrate-binding protein |
| AVAEIV_RS22755 | 4871651 | 4871663 | 2 | + | 14.1571 | 6.19E-06 | 0.0601 | CAGAAGGCCACCG | *glpK* | glycerol kinase GlpK |
| AVAEIV_RS02520 | 505362 | 505374 | 308 | + | 14.1429 | 6.29E-06 | 0.0601 | CTGCTCGCCAGCA | - | ExbD/TolR family protein |
| AVAEIV_RS04470 | 911901 | 911913 | -40 | - | 14.1429 | 6.29E-06 | 0.0649 | CAGCAGGCCATCG | - | type III PLP-dependent enzyme |
| AVAEIV_RS07895 | 1642067 | 1642079 | 275 | - | 14.1429 | 6.29E-06 | 0.0649 | CGCAGCGCCAGCA | - | LysR family transcriptional regulator |
| AVAEIV_RS10005 | 2095226 | 2095238 | 51 | - | 14.1429 | 6.29E-06 | 0.0601 | CGCAGCGCCAGCA | - | monovalent cation/H+ antiporter subunit D |
| AVAEIV_RS10100 | 2108455 | 2108467 | 265 | - | 14.1429 | 6.29E-06 | 0.0601 | CAGCAGGCCATCG | - | F0F1 ATP synthase subunit alpha |
| AVAEIV_RS14755 | 3208371 | 3208383 | 52 | + | 14.1429 | 6.29E-06 | 0.0601 | CGCAGCGCCAGCA | *nuoG* | NADH-quinone oxidoreductase subunit NuoG |
| AVAEIV_RS17625 | 3817778 | 3817790 | 328 | + | 14.1429 | 6.29E-06 | 0.0601 | CTGCTCGCCAGCA | *minE* | cell division topological specificity factor MinE |
| AVAEIV_RS09170 | 1908995 | 1909007 | 296 | - | 14.1286 | 6.38E-06 | 0.0601 | CAGGACGCCATCG | - | PEP/pyruvate-binding domain-containing protein |
| AVAEIV_RS20440 | 4345582 | 4345594 | 208 | + | 14.1286 | 6.38E-06 | 0.0649 | CTGTTCGCCATCG | - | TraR/DksA family transcriptional regulator |
| AVAEIV_RS20445 | 4345582 | 4345594 | -37 | + | 14.1286 | 6.38E-06 | 0.0601 | CTGTTCGCCATCG | - | DUF3015 domain-containing protein |
| AVAEIV_RS00450 | 91999 | 92011 | 72 | - | 14.1143 | 6.48E-06 | 0.0601 | CAGTAGGCCAGCG | - | cytochrome c |
| AVAEIV_RS03735 | 752202 | 752214 | 56 | - | 14.1143 | 6.48E-06 | 0.0601 | CAGAGTGCCAGCG | - | adenylosuccinate synthase |
| AVAEIV_RS07170 | 1478746 | 1478758 | 65 | - | 14.1143 | 6.48E-06 | 0.0601 | CCCTTGGCCAGCA | *recO* | DNA repair protein RecO |
| AVAEIV_RS00885 | 172889 | 172901 | 304 | - | 14.1 | 6.54E-06 | 0.0649 | CCGATCGCCGGCA | - | DUF2388 domain-containing protein |
| AVAEIV_RS09785 | 2047178 | 2047190 | 298 | - | 14.1 | 6.54E-06 | 0.0649 | CCGATCGCCGGCA | *egtD* | L-histidine N(alpha)-methyltransferase |
| AVAEIV_RS13185 | 2860297 | 2860309 | 359 | - | 14.1 | 6.54E-06 | 0.0601 | CCGATCGCCGGCA | - | hypothetical protein |
| AVAEIV_RS13310 | 2882426 | 2882438 | 239 | - | 14.1 | 6.54E-06 | 0.0601 | CCGATCGCCGGCA | - | LysR family transcriptional regulator |
| AVAEIV_RS21195 | 4506614 | 4506626 | 260 | - | 14.1 | 6.54E-06 | 0.0649 | CCGATCGCCGGCA | *otsB* | trehalose-phosphatase |
| AVAEIV_RS23010 | 4926979 | 4926991 | -28 | + | 14.1 | 6.54E-06 | 0.0601 | CATATCGCCAGCG | - | protein |
| AVAEIV_RS26130 | 5617235 | 5617247 | 76 | - | 14.1 | 6.54E-06 | 0.0649 | CGCATGGCCACCA | *rnpA* | ribonuclease P protein component |
| AVAEIV_RS26140 | 5617235 | 5617247 | 58 | - | 14.1 | 6.54E-06 | 0.0601 | CGCATGGCCACCA | - | hypothetical protein |

| | | | | | | | | | | |
|---|---|---|---|---|---|---|---|---|---|---|
| AVAEIV_RS05835 | 1195371 | 1195383 | 279 | + | 14.0714 | 6.63E-06 | 0.0601 | CAGCTCGCCGGCA | - | PaaI family thioesterase |
| AVAEIV_RS12690 | 2694788 | 2694800 | -49 | - | 14.0714 | 6.63E-06 | 0.0649 | CGCGTCGCCATCA | - | 3-hydroxybutyryl-CoA dehydrogenase |
| AVAEIV_RS12990 | 2815275 | 2815287 | 262 | - | 14.0714 | 6.63E-06 | 0.0649 | CAGCTCGCCGGCA | - | (2Fe-2S)-binding protein |
| AVAEIV_RS18300 | 3941824 | 3941836 | 298 | - | 14.0714 | 6.63E-06 | 0.0649 | CAGATCGGCAGCA | - | hypothetical protein |
| AVAEIV_RS00080 | 18731 | 18743 | 22 | - | 14.0571 | 6.69E-06 | 0.0649 | CAGTTCGCCGTCG | *rsmB* | RsmB |
| AVAEIV_RS02035 | 407567 | 407579 | 379 | + | 14.0571 | 6.69E-06 | 0.0649 | CGCTTGGCCAGCA | - | hypothetical protein |
| AVAEIV_RS05305 | 1095711 | 1095723 | 16 | - | 14.0571 | 6.69E-06 | 0.0649 | CCGTTCGCCACCG | - | TAXI family TRAP transporter solute-binding subunit |
| AVAEIV_RS07605 | 1585703 | 1585715 | 369 | + | 14.0571 | 6.69E-06 | 0.0601 | CAGTTCGCCGTCG | - | subunit |
| AVAEIV_RS08610 | 1795827 | 1795839 | 90 | + | 14.0571 | 6.69E-06 | 0.0601 | CGCAACGCCATCG | - | cysteine desulfurase |
| AVAEIV_RS09300 | 1942292 | 1942304 | 384 | - | 14.0571 | 6.69E-06 | 0.0649 | CAGTTCGCCGTCG | - | DUF3144 domain-containing protein |
| AVAEIV_RS10870 | 2281872 | 2281884 | 90 | + | 14.0571 | 6.69E-06 | 0.0601 | CGCAACGCCATCG | - | cysteine desulfurase |
| AVAEIV_RS13460 | 2912051 | 2912063 | -44 | + | 14.0571 | 6.69E-06 | 0.0649 | CGCTTGGCCAGCA | - | hypothetical protein |
| AVAEIV_RS24510 | 5208959 | 5208971 | 359 | + | 14.0571 | 6.69E-06 | 0.0649 | CAGTTCGCCGTCG | - | ABC transporter permease subunit |
| AVAEIV_RS03535 | 714098 | 714110 | 13 | - | 14.0429 | 6.79E-06 | 0.0601 | CAGGTTGCCATCG | - | cytochrome P450 |
| AVAEIV_RS10960 | 2305639 | 2305651 | 167 | + | 14.0429 | 6.79E-06 | 0.0649 | CCATTCGCCATCG | - | (2,3-dihydroxybenzoyl)adenylate synthase |
| AVAEIV_RS15385 | 3359919 | 3359931 | 158 | - | 14.0429 | 6.79E-06 | 0.0601 | CGGATCGCCGGCA | - | PA1571 family protein |
| AVAEIV_RS16790 | 3633385 | 3633397 | 347 | - | 14.0429 | 6.79E-06 | 0.0601 | CTGATGGCCATCG | - | dienelactone hydrolase family protein |
| AVAEIV_RS22345 | 4772257 | 4772269 | 27 | - | 14.0429 | 6.79E-06 | 0.0649 | CGGATCGCCGGCA | *rfaP* | lipopolysaccharide core heptose(I) kinase RfaP |
| AVAEIV_RS04815 | 978071 | 978083 | 126 | - | 14.0286 | 6.87E-06 | 0.0601 | CCGACCGCCAGCA | *madM* | malonate transporter subunit MadM |
| AVAEIV_RS06075 | 1239714 | 1239726 | -31 | - | 14.0286 | 6.87E-06 | 0.0649 | CAGTTGGCCATCC | - | DUF3275 family protein |
| AVAEIV_RS20360 | 4331351 | 4331363 | 234 | - | 14.0286 | 6.87E-06 | 0.0601 | CCGACCGCCAGCA | - | hypothetical protein |
| AVAEIV_RS21675 | 4615126 | 4615138 | 356 | - | 14.0286 | 6.87E-06 | 0.0601 | GAGATCGCCAGCA | *gcl* | glyoxylate carboligase |
| AVAEIV_RS22610 | 4842628 | 4842640 | 36 | + | 14.0286 | 6.87E-06 | 0.0601 | CCGACCGCCAGCA | *ubiE* | methyltransferase/2-methoxy-6-polyprenyl-1,4-benzoquinol methylase UbiE |
| AVAEIV_RS22740 | 4869527 | 4869539 | -41 | + | 14.0286 | 6.87E-06 | 0.0649 | CCGACCGCCAGCA | - | choice-of-anchor I family protein |
| AVAEIV_RS22745 | 4869527 | 4869539 | 377 | + | 14.0286 | 6.87E-06 | 0.0601 | CCGACCGCCAGCA | - | hypothetical protein |
| AVAEIV_RS22965 | 4919443 | 4919455 | 272 | + | 14.0286 | 6.87E-06 | 0.0649 | CCCATTGCCATCG | - | OsmC family protein |
| AVAEIV_RS24035 | 5115567 | 5115579 | 124 | - | 14.0286 | 6.87E-06 | 0.0601 | CCGACCGCCAGCA | *xerC* | tyrosine recombinase XerC |
| AVAEIV_RS00055 | 12540 | 12552 | 132 | - | 14.0143 | 6.93E-06 | 0.0651 | CCGAGCGCCAGCG | *glyS* | glycine--tRNA ligase subunit beta |
| AVAEIV_RS20235 | 4316507 | 4316519 | 365 | + | 14.0143 | 6.93E-06 | 0.0602 | CCGAGCGCCAGCG | *suhB* | inositol-phosphate phosphatase |
| AVAEIV_RS17325 | 3750936 | 3750948 | 286 | + | 14 | 7.01E-06 | 0.0654 | CACCTTGCCATCG | - | aspartate kinase |
| AVAEIV_RS00275 | 58666 | 58678 | 211 | - | 13.9857 | 7.10E-06 | 0.0602 | CTCATCGCCATCG | - | hypothetical protein |
| AVAEIV_RS01655 | 319503 | 319515 | 192 | + | 13.9857 | 7.10E-06 | 0.0602 | CAGCGCGCCAGCG | - | hypothetical protein |
| AVAEIV_RS06615 | 1364821 | 1364833 | 149 | + | 13.9857 | 7.10E-06 | 0.0602 | CAGCGCGCCAGCG | *mreD* | rod shape-determining protein MreD |
| AVAEIV_RS20635 | 4388648 | 4388660 | 364 | + | 13.9857 | 7.10E-06 | 0.0602 | CAGCGCGCCAGCG | - | 1-acyl-sn-glycerol-3-phosphate acyltransferase |
| AVAEIV_RS10045 | 2101889 | 2101901 | 66 | + | 13.9714 | 7.24E-06 | 0.0668 | CCGAAGGCCACCA | - | transporter permease LivM |
| AVAEIV_RS10515 | 2199154 | 2199166 | 148 | - | 13.9714 | 7.24E-06 | 0.0602 | CCGATGGCCACCG | - | helix-turn-helix transcriptional regulator |
| AVAEIV_RS00885 | 172825 | 172837 | 240 | + | 13.9571 | 7.33E-06 | 0.0668 | CCGCTGGCCATCG | - | DUF2388 domain-containing protein |
| AVAEIV_RS00890 | 174411 | 174423 | 255 | + | 13.9571 | 7.33E-06 | 0.0668 | TAGATCGCCAGCA | - | beta |
| AVAEIV_RS08610 | 1795563 | 1795575 | 354 | + | 13.9571 | 7.33E-06 | 0.0602 | CCGCTGGCCATCG | - | cysteine desulfurase |
| AVAEIV_RS10870 | 2281608 | 2281620 | 354 | + | 13.9571 | 7.33E-06 | 0.0602 | CCGCTGGCCATCG | - | cysteine desulfurase |
| AVAEIV_RS15240 | 3329061 | 3329073 | 95 | + | 13.9571 | 7.33E-06 | 0.0668 | CGGAGCGCCAGCG | - | containing protein |
| AVAEIV_RS03770 | 758269 | 758281 | 94 | + | 13.9429 | 7.41E-06 | 0.0602 | CAGCTGGCCACCG | - | hypothetical protein |
| AVAEIV_RS10175 | 2123727 | 2123739 | -49 | - | 13.9429 | 7.41E-06 | 0.0668 | CAGTTCGCCATGA | - | cytochrome ubiquinol oxidase subunit I |
| AVAEIV_RS18800 | 4047771 | 4047783 | 297 | + | 13.9429 | 7.41E-06 | 0.0602 | CCGGTCGCCATCG | - | fumarate hydratase |
| AVAEIV_RS01350 | 264921 | 264933 | 202 | + | 13.9286 | 7.49E-06 | 0.0668 | CAGGTCGCCACCG | *vnfA1* | regulator VnfA1 |
| AVAEIV_RS04715 | 959953 | 959965 | 324 | - | 13.9286 | 7.49E-06 | 0.0602 | CAACTGGCCATCG | - | AI-2E family transporter |
| AVAEIV_RS08075 | 1673626 | 1673638 | 363 | + | 13.9286 | 7.49E-06 | 0.0602 | CTGTTCGCCACCA | *edd* | phosphogluconate dehydratase |
| AVAEIV_RS13830 | 2996662 | 2996674 | 274 | + | 13.9286 | 7.49E-06 | 0.0668 | CAACTGGCCATCG | *tssC* | subunit |
| AVAEIV_RS18530 | 3989240 | 3989252 | -44 | - | 13.9286 | 7.49E-06 | 0.0602 | CCGAGTGCCAGCA | - | di-heme oxidoredictase family protein |
| AVAEIV_RS21570 | 4588396 | 4588408 | 45 | - | 13.9286 | 7.49E-06 | 0.0602 | CCGTAGGCCAGCA | - | protein |
| AVAEIV_RS24200 | 5145775 | 5145787 | 282 | + | 13.9286 | 7.49E-06 | 0.0668 | CCGTAGGCCAGCA | *rpoH* | RNA polymerase sigma factor RpoH |

| | | | | | | | | | | |
|---|---|---|---|---|---|---|---|---|---|---|
| AVAEIV_RS04380 | 891552 | 891564 | 349 | + | 13.9143 | 7.62E-06 | 0.0668 | CGGATGGCCACCG | *miaB* | methylthiotransferase MiaB |
| AVAEIV_RS11555 | 2442393 | 2442405 | 312 | + | 13.9143 | 7.62E-06 | 0.0604 | CCTATCGCCAGCA | - | heavy metal translocating P-type ATPase |
| AVAEIV_RS25650 | 5485688 | 5485700 | 336 | + | 13.9143 | 7.62E-06 | 0.0668 | CAGTTCGCCATTA | - | hypothetical protein |
| AVAEIV_RS05840 | 1196037 | 1196049 | 96 | - | 13.9 | 7.80E-06 | 0.0604 | CCCCTCGCCATCG | - | protein |
| AVAEIV_RS10335 | 2158196 | 2158208 | 359 | + | 13.9 | 7.80E-06 | 0.0604 | CGGCTGGCCATCG | - | FAD-dependent oxidoreductase |
| AVAEIV_RS17330 | 3753540 | 3753552 | 171 | + | 13.9 | 7.80E-06 | 0.0668 | CGGCTGGCCATCG | *alaS* | alanine--tRNA ligase |
| AVAEIV_RS20160 | 4306113 | 4306125 | 65 | + | 13.9 | 7.80E-06 | 0.0668 | CGGCTGGCCATCG | *ispG* | diphosphate synthase |
| AVAEIV_RS22420 | 4794656 | 4794668 | 329 | + | 13.9 | 7.80E-06 | 0.0668 | CGGCTGGCCATCG | - | antitoxin |
| AVAEIV_RS24760 | 5261297 | 5261309 | 322 | - | 13.9 | 7.80E-06 | 0.0604 | CGGCTGGCCATCG | - | OprD family porin |
| AVAEIV_RS00215 | 44021 | 44033 | 58 | - | 13.8857 | 7.88E-06 | 0.0668 | CACCTCGCCACCG | - | hypothetical protein |
| AVAEIV_RS15210 | 3312948 | 3312960 | 259 | - | 13.8857 | 7.88E-06 | 0.0668 | CGGGTCGCCATCG | - | ABC transporter permease |
| AVAEIV_RS17330 | 3753743 | 3753755 | 374 | + | 13.8857 | 7.88E-06 | 0.0668 | CACATGGCCATCC | *alaS* | alanine--tRNA ligase |
| AVAEIV_RS25080 | 5338651 | 5338663 | -10 | + | 13.8857 | 7.88E-06 | 0.0604 | CACATGGCCATCC | - | TIM barrel protein |
| AVAEIV_RS08260 | 1719066 | 1719078 | 36 | - | 13.8714 | 7.98E-06 | 0.0604 | CCCTACGCCAGCA | - | glycosyltransferase family 2 protein |
| AVAEIV_RS01945 | 389628 | 389640 | 367 | + | 13.8571 | 8.04E-06 | 0.0604 | CAGTTCGCCGCCA | - | metallophosphoesterase |
| AVAEIV_RS06470 | 1332542 | 1332554 | 297 | - | 13.8571 | 8.04E-06 | 0.0604 | CAGTTCGCCGCCA | - | alpha-hydroxy acid oxidase |
| AVAEIV_RS09885 | 2070924 | 2070936 | 153 | - | 13.8571 | 8.04E-06 | 0.0668 | CAGATGGCCATGA | *metG* | methionine--tRNA ligase |
| AVAEIV_RS09890 | 2070924 | 2070936 | 27 | - | 13.8571 | 8.04E-06 | 0.0604 | CAGATGGCCATGA | *apbC* | iron-sulfur cluster carrier protein ApbC |
| AVAEIV_RS24590 | 5224501 | 5224513 | 234 | - | 13.8571 | 8.04E-06 | 0.0668 | CGCAACGCCACCA | - | ArsR/SmtB family transcription factor |
| AVAEIV_RS02520 | 505315 | 505327 | 355 | - | 13.8429 | 8.16E-06 | 0.0604 | CTGATGGCCACCA | - | ExbD/TolR family protein |
| AVAEIV_RS09435 | 1967164 | 1967176 | 379 | + | 13.8429 | 8.16E-06 | 0.0668 | CAATTCGCCGTCA | *istA* | IS21 family transposase |
| AVAEIV_RS16300 | 3549135 | 3549147 | 41 | + | 13.8429 | 8.16E-06 | 0.0668 | CAGTTCGGCATCA | - | type II toxin-antitoxin system VapC family toxin |
| AVAEIV_RS20190 | 4310304 | 4310316 | 204 | + | 13.8429 | 8.16E-06 | 0.0668 | CGCCTCGCCATCG | *fdx* | ISC system 2Fe-2S type ferredoxin |
| AVAEIV_RS08305 | 1728717 | 1728729 | 235 | + | 13.8286 | 8.27E-06 | 0.0604 | CCCATTGCCACCA | *cysC* | adenylyl-sulfate kinase |
| AVAEIV_RS09200 | 1920004 | 1920016 | 323 | - | 13.8286 | 8.27E-06 | 0.0604 | CTGCTGGCCATCA | - | DUF2334 domain-containing protein |
| AVAEIV_RS14730 | 3203143 | 3203155 | 329 | + | 13.8286 | 8.27E-06 | 0.0604 | CCGATCGCCAGCC | - | NADH-quinone oxidoreductase subunit A |
| AVAEIV_RS26100 | 5612502 | 5612514 | 146 | + | 13.8286 | 8.27E-06 | 0.0668 | CACTGGGCCAGCA | - | tyrosine-type recombinase/integrase |
| AVAEIV_RS26105 | 5612502 | 5612514 | -41 | + | 13.8286 | 8.27E-06 | 0.0668 | CACTGGGCCAGCA | - | tyrosine-type recombinase/integrase |
| AVAEIV_RS00945 | 187718 | 187730 | 241 | + | 13.8143 | 8.35E-06 | 0.0604 | CAGCATGCCATCG | - | I78 family peptidase inhibitor |
| AVAEIV_RS13180 | 2860067 | 2860079 | 372 | - | 13.8143 | 8.35E-06 | 0.0668 | CGCTTCGCCAGCG | - | hypothetical protein |
| AVAEIV_RS14440 | 3149402 | 3149414 | 224 | - | 13.8143 | 8.35E-06 | 0.0668 | CTGGTCGCCATCA | *treS* | maltose alpha-D-glucosyltransferase |
| AVAEIV_RS19920 | 4247947 | 4247959 | 319 | + | 13.8143 | 8.35E-06 | 0.0668 | CGCTTCGCCAGCG | - | 3'-5' exonuclease |
| AVAEIV_RS22175 | 4733633 | 4733645 | 387 | + | 13.8143 | 8.35E-06 | 0.0668 | CAGAGCGCCGGCA | - | diguanylate cyclase domain-containing protein |
| AVAEIV_RS22950 | 4916365 | 4916377 | 230 | + | 13.8143 | 8.35E-06 | 0.0604 | CGCTTCGCCAGCG | - | histidine triad nucleotide-binding protein |
| AVAEIV_RS03855 | 775695 | 775707 | 242 | - | 13.8 | 8.47E-06 | 0.0604 | CCGCGCGCCAGCA | - | phosphonate metabolism protein PhnP |
| AVAEIV_RS04370 | 888701 | 888713 | -31 | - | 13.8 | 8.47E-06 | 0.0668 | CAGGTGGCCAGCG | *ybeY* | rRNA maturation RNase YbeY |
| AVAEIV_RS05215 | 1070924 | 1070936 | 128 | - | 13.8 | 8.47E-06 | 0.0604 | CCGCGCGCCAGCA | *moaE* | molybdopterin synthase catalytic subunit MoaE |
| AVAEIV_RS07660 | 1595616 | 1595628 | 140 | - | 13.8 | 8.47E-06 | 0.0604 | CAGCTCGCCAGCC | *murB* | UDP-N-acetylmuramate dehydrogenase |
| AVAEIV_RS09000 | 1873401 | 1873413 | 229 | - | 13.8 | 8.47E-06 | 0.0668 | GAGTTCGCCATCA | - | DUF2243 domain-containing protein |
| AVAEIV_RS10755 | 2255244 | 2255256 | 343 | - | 13.8 | 8.47E-06 | 0.0668 | CAGCTCGCCAGCC | *cas2* | CRISPR-associated endonuclease Cas2 |
| AVAEIV_RS17250 | 3738596 | 3738608 | 63 | - | 13.8 | 8.47E-06 | 0.0668 | GAGTTCGCCATCA | - | bifunctional 4-hydroxy-2-oxoglutarate aldolase/2-dehydro-3-deoxy-phosphogluconate aldolase |
| AVAEIV_RS21400 | 4552650 | 4552662 | 174 | - | 13.8 | 8.47E-06 | 0.0604 | CAGGAGGCCAGCA | - | DUF2845 domain-containing protein |
| AVAEIV_RS02075 | 412998 | 413010 | 169 | + | 13.7857 | 8.63E-06 | 0.0668 | CATTTTGCCATCA | - | hypothetical protein |
| AVAEIV_RS08220 | 1707698 | 1707710 | 314 | - | 13.7857 | 8.63E-06 | 0.0604 | CCCAAGGCCAGCA | - | ABC transporter ATP-binding protein |
| AVAEIV_RS13110 | 2842316 | 2842328 | 279 | - | 13.7857 | 8.63E-06 | 0.0604 | CCCATGGCCAGCG | - | (2Fe-2S)-binding protein |
| AVAEIV_RS15325 | 3348932 | 3348944 | 228 | - | 13.7857 | 8.63E-06 | 0.0668 | CCGATGGCCGTCA | *lpdA* | dihydrolipoyl dehydrogenase |
| AVAEIV_RS19705 | 4200398 | 4200410 | 258 | - | 13.7857 | 8.63E-06 | 0.0668 | CCGTGCGCCATCG | *tsaB* | threonylcarbamoyltransferase complex dimerization subunit type 1 TsaB |
| AVAEIV_RS22565 | 4831312 | 4831324 | 178 | + | 13.7857 | 8.63E-06 | 0.0604 | CCCATGGCCAGCG | - | IS4-like element ISAzvi6 family transposase |
| AVAEIV_RS13200 | 2861890 | 2861902 | 260 | - | 13.7714 | 8.73E-06 | 0.0604 | CGGATCGCCAGCC | - | MbcA/ParS/Xre antitoxin family protein |
| AVAEIV_RS19340 | 4153210 | 4153222 | 25 | - | 13.7714 | 8.73E-06 | 0.0668 | CGGATCGCCAGCC | - | hypothetical protein |

| | | | | | | | | | | |
|---|---|---|---|---|---|---|---|---|---|---|
| AVAEIV_RS14910 | 3249038 | 3249050 | 330 | + | 13.7571 | 8.83E-06 | 0.0668 | CCGCTGGCCACCA | - | alanine/glycine:cation symporter family protein |
| AVAEIV_RS17205 | 3731360 | 3731372 | 238 | - | 13.7571 | 8.83E-06 | 0.0668 | CCGCTGGCCACCA | *asd* | aspartate-semialdehyde dehydrogenase |
| AVAEIV_RS20310 | 4327542 | 4327554 | 41 | - | 13.7571 | 8.83E-06 | 0.0604 | CACCAGGCCAGCA | - | hypothetical protein |
| AVAEIV_RS21995 | 4693582 | 4693594 | 109 | + | 13.7571 | 8.83E-06 | 0.0604 | CACCTGGCCAGCG | - | paraquat-inducible protein A |
| AVAEIV_RS05360 | 1108398 | 1108410 | -36 | - | 13.7429 | 8.92E-06 | 0.0668 | CACGTCGCCAGCG | *rlmF* | RlmF |
| AVAEIV_RS10085 | 2107011 | 2107023 | 30 | - | 13.7429 | 8.92E-06 | 0.0604 | CGGCGCGCCAGCA | - | F0F1 ATP synthase subunit A |
| AVAEIV_RS12315 | 2605724 | 2605736 | 123 | - | 13.7429 | 8.92E-06 | 0.0604 | CGGCGCGCCAGCA | - | hypothetical protein |
| AVAEIV_RS15635 | 3424988 | 3425000 | 48 | + | 13.7429 | 8.92E-06 | 0.0668 | CGGCGCGCCAGCA | *ccmE* | cytochrome c maturation protein CcmE |
| AVAEIV_RS16715 | 3620109 | 3620121 | 272 | + | 13.7429 | 8.92E-06 | 0.0668 | CCGGTCGCCACCA | - | cobyrinate a,c-diamide synthase |
| AVAEIV_RS18570 | 4001576 | 4001588 | -26 | + | 13.7429 | 8.92E-06 | 0.0668 | CGGCGCGCCAGCA | - | LysR substrate-binding domain-containing protein |
| AVAEIV_RS20600 | 4383637 | 4383649 | 262 | + | 13.7429 | 8.92E-06 | 0.0668 | CACGACGCCAGCA | - | DUF1302 domain-containing protein |
| AVAEIV_RS01530 | 294625 | 294637 | 146 | + | 13.7286 | 9.05E-06 | 0.0668 | CAGTTGGCCGGCA | *ruvX* | Holliday junction resolvase RuvX |
| AVAEIV_RS01860 | 368991 | 369003 | 255 | - | 13.7286 | 9.05E-06 | 0.0604 | CGGATGGCCGTCA | - | RNA polymerase sigma factor |
| AVAEIV_RS02655 | 541356 | 541368 | 376 | + | 13.7286 | 9.05E-06 | 0.0668 | CGCATGGCCAGCG | - | GNAT family N-acetyltransferase |
| AVAEIV_RS03095 | 637017 | 637029 | 196 | - | 13.7286 | 9.05E-06 | 0.0604 | CGCATGGCCAGCG | *fusA* | elongation factor G |
| AVAEIV_RS05055 | 1039397 | 1039409 | 162 | + | 13.7286 | 9.05E-06 | 0.0668 | CAGTTGGCCGGCA | *algG* | mannuronan 5-epimerase AlgG |
| AVAEIV_RS08085 | 1677859 | 1677871 | 191 | + | 13.7286 | 9.05E-06 | 0.0668 | CGCATGGCCAGCG | - | MurR/RpiR family transcriptional regulator |
| AVAEIV_RS08090 | 1677859 | 1677871 | -30 | + | 13.7286 | 9.05E-06 | 0.0604 | CGCATGGCCAGCG | *pgl* | 6-phosphogluconolactonase |
| AVAEIV_RS12250 | 2594827 | 2594839 | 285 | - | 13.7286 | 9.05E-06 | 0.0668 | CGCATGGCCAGCG | *fliN* | flagellar motor switch protein FliN |
| AVAEIV_RS06880 | 1415107 | 1415119 | 265 | - | 13.7143 | 9.17E-06 | 0.0604 | CTGATTGCCATCG | *rnpB* | RNase P RNA component class A |
| AVAEIV_RS07145 | 1472995 | 1473007 | 121 | - | 13.7143 | 9.17E-06 | 0.0604 | CACATCGCCGCCA | *lepB* | signal peptidase I |
| AVAEIV_RS09170 | 1908937 | 1908949 | 354 | + | 13.7143 | 9.17E-06 | 0.0604 | CCGACGGCCATCA | - | PEP/pyruvate-binding domain-containing protein |
| AVAEIV_RS10285 | 2146002 | 2146014 | 367 | + | 13.7143 | 9.17E-06 | 0.0604 | CAGATCGCCTTCA | *gltX* | glutamate--tRNA ligase |
| AVAEIV_RS16295 | 3548955 | 3548967 | 252 | - | 13.7143 | 9.17E-06 | 0.0671 | GAGATGGCCATCA | *gcvT* | glycine cleavage system aminomethyltransferase GcvT |
| AVAEIV_RS18805 | 4049384 | 4049396 | 370 | - | 13.7143 | 9.17E-06 | 0.0604 | CCGACGGCCATCA | - | hypothetical protein |
| AVAEIV_RS19130 | 4123903 | 4123915 | 217 | + | 13.7143 | 9.17E-06 | 0.0671 | CCATTGGCCAGCA | - | glycoside hydrolase family 104 protein |
| AVAEIV_RS22390 | 4787568 | 4787580 | 189 | + | 13.7143 | 9.17E-06 | 0.0604 | CAGATCGCCTTCA | - | universal stress protein |
| AVAEIV_RS22120 | 4720743 | 4720755 | 41 | + | 13.7 | 9.28E-06 | 0.0604 | CCGAGGGCCATCG | - | diacylglycerol/lipid kinase family protein |
| AVAEIV_RS15460 | 3378788 | 3378800 | 231 | - | 13.6857 | 9.37E-06 | 0.0604 | CGGGTCGCCACCA | - | glycosyltransferase family 4 protein |
| AVAEIV_RS00225 | 45083 | 45095 | 371 | - | 13.6714 | 9.47E-06 | 0.0604 | CGGAACGCCACCG | - | ABC transporter substrate-binding protein |
| AVAEIV_RS15125 | 3292442 | 3292454 | 178 | + | 13.6714 | 9.47E-06 | 0.0604 | CAGCGGGCCATCG | - | ATP-binding protein |
| AVAEIV_RS19405 | 4159913 | 4159925 | 281 | + | 13.6714 | 9.47E-06 | 0.0682 | CAGATCGCCAACA | - | KilA-N domain-containing protein |
| AVAEIV_RS01390 | 271162 | 271174 | 20 | - | 13.6571 | 9.60E-06 | 0.0604 | CAGTCGGCCAGCA | - | DUF4870 domain-containing protein |
| AVAEIV_RS03355 | 675857 | 675869 | 361 | + | 13.6571 | 9.60E-06 | 0.0604 | CAGGGCGCCATCG | *nusB* | transcription antitermination factor NusB |
| AVAEIV_RS10040 | 2100886 | 2100898 | 316 | - | 13.6571 | 9.60E-06 | 0.0682 | CCCACCGCCATCA | *livG* | high-affinity branched-chain amino acid ABC transporter ATP-binding protein LivG |
| AVAEIV_RS12785 | 2716691 | 2716703 | 353 | + | 13.6571 | 9.60E-06 | 0.0604 | CGATTGGCCAGCA | - | dipeptidase |
| AVAEIV_RS13935 | 3020999 | 3021011 | 260 | - | 13.6571 | 9.60E-06 | 0.0604 | CGGCACGCCATCG | - | MFS transporter |
| AVAEIV_RS16100 | 3509108 | 3509120 | 324 | + | 13.6571 | 9.60E-06 | 0.0604 | CAGGGCGCCATCG | - | ABC transporter substrate-binding protein |
| AVAEIV_RS19350 | 4156056 | 4156068 | 188 | + | 13.6571 | 9.60E-06 | 0.0682 | CGGCACGCCATCG | - | terminase large subunit domain-containing protein |
| AVAEIV_RS22605 | 4842239 | 4842251 | -22 | + | 13.6571 | 9.60E-06 | 0.0604 | CCCACCGCCATCA | - | containing protein |
| AVAEIV_RS23915 | 5096072 | 5096084 | 194 | - | 13.6571 | 9.60E-06 | 0.0682 | CAGGGCGCCATCG | - | YaiI/YqxD family protein |
| AVAEIV_RS23920 | 5096072 | 5096084 | -31 | - | 13.6571 | 9.60E-06 | 0.0682 | CAGGGCGCCATCG | - | FeoB-associated Cys-rich membrane protein |
| AVAEIV_RS25700 | 5499057 | 5499069 | 344 | - | 13.6571 | 9.60E-06 | 0.0682 | CAGTCGGCCAGCA | - | CoA transferase subunit A |
| AVAEIV_RS00780 | 152794 | 152806 | 99 | + | 13.6429 | 9.74E-06 | 0.0604 | CCGAATGCCACCA | *nifU* | Fe-S cluster assembly protein NifU |
| AVAEIV_RS07620 | 1590034 | 1590046 | 208 | - | 13.6429 | 9.74E-06 | 0.0682 | CCCAGCGCCATCG | - | DUF2062 domain-containing protein |
| AVAEIV_RS10950 | 2302924 | 2302936 | 29 | + | 13.6429 | 9.74E-06 | 0.0682 | CAGTGGGCCAGCG | - | amino acid adenylation domain-containing protein |
| AVAEIV_RS15345 | 3355711 | 3355723 | 103 | + | 13.6429 | 9.74E-06 | 0.0682 | CCCAGCGCCATCG | *sdhA* | succinate dehydrogenase flavoprotein subunit |
| AVAEIV_RS02925 | 603244 | 603256 | 237 | - | 13.6286 | 9.83E-06 | 0.0682 | CGGTACGCCAGCG | - | PolC-type DNA polymerase III |
| AVAEIV_RS11670 | 2475116 | 2475128 | 246 | - | 13.6286 | 9.83E-06 | 0.0682 | CGGTACGCCAGCG | - | cation-transporting P-type ATPase |
| AVAEIV_RS11675 | 2475116 | 2475128 | 9 | - | 13.6286 | 9.83E-06 | 0.0604 | CGGTACGCCAGCG | - | transglutaminase domain-containing protein |
| AVAEIV_RS12590 | 2674720 | 2674732 | 242 | + | 13.6286 | 9.83E-06 | 0.0604 | CGTTTCGCCATCA | *pgsA* | phosphatidyltransferase |

| | | | | | | | | | | |
|---|---|---|---|---|---|---|---|---|---|---|
| AVAEIV_RS18935 | 4078026 | 4078038 | -3 | + | 13.6286 | 9.83E-06 | 0.0682 | CATAGCGCCAGCA | *truD* | tRNA pseudouridine(13) synthase TruD |
| AVAEIV_RS03810 | 768234 | 768246 | 149 | - | 13.6143 | 9.91E-06 | 0.0604 | CAGATCGCCATGG | - | HAD family hydrolase |
| AVAEIV_RS06040 | 1233237 | 1233249 | 274 | - | 13.6143 | 9.91E-06 | 0.0682 | CAGATCGCCATGG | - | IS1595-like element ISAzvi1 family transposase |
| AVAEIV_RS06045 | 1233237 | 1233249 | -48 | - | 13.6143 | 9.91E-06 | 0.0604 | CAGATCGCCATGG | - | HU family DNA-binding protein |
| AVAEIV_RS06050 | 1233237 | 1233249 | 308 | - | 13.6143 | 9.91E-06 | 0.0604 | CAGATCGCCATGG | - | hypothetical protein |
| AVAEIV_RS15390 | 3361670 | 3361682 | 240 | + | 13.6143 | 9.91E-06 | 0.0682 | CAGATCGCCATGG | *pdxB* | 4-phosphoerythronate dehydrogenase PdxB |
| AVAEIV_RS02520 | 505500 | 505512 | 170 | + | 13.6 | 1.00E-05 | 0.0604 | CTGATCGCCACCG | - | ExbD/TolR family protein |
| AVAEIV_RS03250 | 657326 | 657338 | 271 | - | 13.6 | 1.00E-05 | 0.0604 | CCGAAGGCCAGCG | - | MFS transporter |
| AVAEIV_RS04200 | 857607 | 857619 | 25 | + | 13.6 | 1.00E-05 | 0.0604 | CCGTTCGCCATCC | *dmpE* | 2-oxopent-4-enoate hydratase |
| AVAEIV_RS07595 | 1583456 | 1583468 | 209 | - | 13.6 | 1.00E-05 | 0.0604 | CTGATCGCCACCG | *sthA* | Si-specific NAD(P)(+) transhydrogenase |
| AVAEIV_RS11640 | 2462753 | 2462765 | -46 | - | 13.6 | 1.00E-05 | 0.0682 | CTGATCGCCACCG | - | YgiW/YdeI family stress tolerance OB fold protein |
| AVAEIV_RS18105 | 3911979 | 3911991 | 157 | - | 13.6 | 1.00E-05 | 0.0682 | CTGATCGCCACCG | *tolR* | protein TolR |
| AVAEIV_RS22320 | 4766386 | 4766398 | 231 | - | 13.6 | 1.00E-05 | 0.0682 | CTGATCGCCACCG | - | hypothetical protein |
| AVAEIV_RS25025 | 5327501 | 5327513 | -37 | + | 13.6 | 1.00E-05 | 0.0604 | CTGATCGCCACCG | - | sugar ABC transporter substrate-binding protein |
| AVAEIV_RS05945 | 1214320 | 1214332 | 286 | - | 13.5857 | 1.02E-05 | 0.0684 | CTGCTCGCCATCG | - | protein |
| AVAEIV_RS05950 | 1214320 | 1214332 | -28 | - | 13.5857 | 1.02E-05 | 0.0684 | CTGCTCGCCATCG | - | protein |
| AVAEIV_RS07185 | 1481570 | 1481582 | 118 | - | 13.5857 | 1.02E-05 | 0.0684 | CGCAGCGCCATCG | *cmoB* | acid(34) synthase CmoB |
| AVAEIV_RS18525 | 3987856 | 3987868 | -43 | + | 13.5857 | 1.02E-05 | 0.0607 | CTGCTCGCCATCG | - | imelysin family protein |
| AVAEIV_RS20595 | 4379584 | 4379596 | 289 | - | 13.5857 | 1.02E-05 | 0.0607 | CAGTGCGCCGTCA | - | L,D-transpeptidase family protein |
| AVAEIV_RS25725 | 5502603 | 5502615 | 246 | + | 13.5857 | 1.02E-05 | 0.0607 | CTGCTCGCCATCG | - | bifunctional diguanylate cyclase/phosphodiesterase |
| AVAEIV_RS03715 | 748275 | 748287 | 383 | - | 13.5714 | 1.04E-05 | 0.061 | CGGCTTGCCATCG | *hflK* | FtsH protease activity modulator HflK |
| AVAEIV_RS06720 | 1385120 | 1385132 | 169 | - | 13.5714 | 1.04E-05 | 0.061 | CAGCAGGCCAGCG | - | phospholipid-binding protein MlaC |
| AVAEIV_RS07745 | 1612811 | 1612823 | 336 | + | 13.5714 | 1.04E-05 | 0.061 | CAGCAGGCCAGCG | - | PilZ domain-containing protein |
| AVAEIV_RS00970 | 192043 | 192055 | 173 | + | 13.5571 | 1.05E-05 | 0.0684 | CTGTTCGCCAGCG | - | YeeE/YedE family protein |
| AVAEIV_RS00975 | 192043 | 192055 | -40 | + | 13.5571 | 1.05E-05 | 0.0611 | CTGTTCGCCAGCG | - | DUF3365 domain-containing protein |
| AVAEIV_RS01535 | 295364 | 295376 | 322 | - | 13.5571 | 1.05E-05 | 0.0684 | CCCGTCGCCAGCA | - | YqgE/AlgH family protein |
| AVAEIV_RS05785 | 1186538 | 1186550 | 68 | - | 13.5571 | 1.05E-05 | 0.0684 | CTGTTCGCCAGCG | - | 3-hydroxyacyl-CoA dehydrogenase |
| AVAEIV_RS14645 | 3187198 | 3187210 | 159 | + | 13.5571 | 1.05E-05 | 0.0611 | CTGTTCGCCAGCG | - | GNAT family N-acetyltransferase |
| AVAEIV_RS23435 | 4996912 | 4996924 | 378 | - | 13.5571 | 1.05E-05 | 0.0684 | CAGGACGCCAGCG | - | sterol desaturase family protein |
| AVAEIV_RS23440 | 4996912 | 4996924 | 59 | - | 13.5571 | 1.05E-05 | 0.0684 | CAGGACGCCAGCG | - | EF-hand domain-containing protein |
| AVAEIV_RS00255 | 54505 | 54517 | 323 | - | 13.5429 | 1.07E-05 | 0.0684 | CAGAGCGCCAGCC | *uvrD* | DNA helicase II |
| AVAEIV_RS08150 | 1692597 | 1692609 | 211 | + | 13.5429 | 1.07E-05 | 0.0612 | CCGATCGCCGTCG | *pheA* | prephenate dehydratase |
| AVAEIV_RS09515 | 1981467 | 1981479 | 225 | + | 13.5429 | 1.07E-05 | 0.0612 | CAGAGCGCCAGCC | - | efflux RND transporter permease subunit |
| AVAEIV_RS14140 | 3076677 | 3076689 | 247 | - | 13.5429 | 1.07E-05 | 0.0684 | CCGATCGCCGTCG | - | carbohydrate ABC transporter permease |
| AVAEIV_RS06010 | 1225113 | 1225125 | 29 | + | 13.5286 | 1.07E-05 | 0.0684 | CAGATCGCCGCCG | - | lytic transglycosylase |
| AVAEIV_RS17885 | 3872067 | 3872079 | 29 | - | 13.5286 | 1.07E-05 | 0.0612 | CAGATCGCCGCCG | - | lytic transglycosylase domain-containing protein |
| AVAEIV_RS25725 | 5502465 | 5502477 | 384 | + | 13.5286 | 1.07E-05 | 0.0612 | CAGATCGCCGCCG | - | bifunctional diguanylate cyclase/phosphodiesterase |
| AVAEIV_RS04620 | 944421 | 944433 | -46 | - | 13.5143 | 1.09E-05 | 0.0684 | CGCCTGGCCAGCA | - | cation diffusion facilitator family transporter |
| AVAEIV_RS05055 | 1039512 | 1039524 | 277 | - | 13.5143 | 1.09E-05 | 0.0684 | CAGAACGGCATCA | *algG* | mannuronan 5-epimerase AlgG |
| AVAEIV_RS20540 | 4366594 | 4366606 | -16 | - | 13.5143 | 1.09E-05 | 0.0684 | CTGATTGCCACCA | - | FKBP-type peptidyl-prolyl cis-trans isomerase |
| AVAEIV_RS20545 | 4366594 | 4366606 | 250 | - | 13.5143 | 1.09E-05 | 0.0614 | CTGATTGCCACCA | - | acetate kinase |
| AVAEIV_RS21810 | 4644425 | 4644437 | 368 | + | 13.5143 | 1.09E-05 | 0.0614 | CAGCTCGCCGTCG | - | 2-hydroxyacid dehydrogenase |
| AVAEIV_RS22125 | 4721690 | 4721702 | 72 | - | 13.5143 | 1.09E-05 | 0.0614 | CAGCTCGCCGTCG | - | hypothetical protein |
| AVAEIV_RS22360 | 4775856 | 4775868 | 225 | - | 13.5143 | 1.09E-05 | 0.0684 | CAGATCGGCATCG | *waaF* | lipopolysaccharide heptosyltransferase II |
| AVAEIV_RS22365 | 4776631 | 4776643 | -48 | + | 13.5143 | 1.09E-05 | 0.0684 | CAGCTCGCCGTCG | - | branched-chain amino acid transaminase |
| AVAEIV_RS25035 | 5330857 | 5330869 | 299 | + | 13.5143 | 1.09E-05 | 0.0684 | CAGAACGGCATCA | - | LysR family transcriptional regulator |
| AVAEIV_RS25925 | 5569297 | 5569309 | 94 | + | 13.5143 | 1.09E-05 | 0.0684 | CCGCTCGCCACCG | - | SrfA family protein |
| AVAEIV_RS03570 | 721345 | 721357 | 327 | - | 13.4857 | 1.11E-05 | 0.0684 | CAGTTCGCCGGCG | - | DUF4136 domain-containing protein |
| AVAEIV_RS06920 | 1422994 | 1423006 | 376 | - | 13.4857 | 1.11E-05 | 0.0616 | CAGTTCGCCGGCG | *murD* | UDP-N-acetylmuramoyl-L-alanine--D-glutamate ligase |
| AVAEIV_RS08080 | 1675618 | 1675630 | 194 | - | 13.4857 | 1.11E-05 | 0.0616 | CGGATCGCCGTCG | - | glucokinase |
| AVAEIV_RS12405 | 2627980 | 2627992 | 198 | - | 13.4857 | 1.11E-05 | 0.0616 | CGGATCGCCGTCG | - | DEAD/DEAH box helicase family protein |
| AVAEIV_RS14540 | 3166813 | 3166825 | 195 | + | 13.4857 | 1.11E-05 | 0.0684 | CGCAACGCCAGCG | *serS* | serine--tRNA ligase |

| | | | | | | | | | | |
|---|---|---|---|---|---|---|---|---|---|---|
| AVAEIV_RS16675 | 3612377 | 3612389 | 298 | - | 13.4857 | 1.11E-05 | 0.0684 | CGCAACGCCAGCG | - | adenosylcobinamide-GDP ribazoletransferase |
| AVAEIV_RS16735 | 3624404 | 3624416 | 213 | - | 13.4857 | 1.11E-05 | 0.0684 | CAGTTCGCCGGCG | - | SIR2 family NAD-dependent protein deacylase |
| AVAEIV_RS20230 | 4316926 | 4316938 | 183 | - | 13.4857 | 1.11E-05 | 0.0684 | CAGTTCGCCGGCG | *trmJ* | methyltransferase TrmJ |
| AVAEIV_RS21355 | 4539580 | 4539592 | 108 | + | 13.4857 | 1.11E-05 | 0.0616 | CGGATCGCCGTCG | - | sigma-54-dependent transcriptional regulator |
| AVAEIV_RS23060 | 4934090 | 4934102 | 196 | - | 13.4857 | 1.11E-05 | 0.0616 | CGGATCGCCGTCG | - | head completion/stabilization protein |
| AVAEIV_RS25405 | 5412172 | 5412184 | 102 | - | 13.4857 | 1.11E-05 | 0.0684 | CAGTTCGCCGGCG | - | electron transport complex subunit E |
| AVAEIV_RS25950 | 5576875 | 5576887 | 124 | + | 13.4857 | 1.11E-05 | 0.0684 | CAGTTCGCCGGCG | *atpD* | F0F1 ATP synthase subunit beta |
| AVAEIV_RS00285 | 62250 | 62262 | 324 | + | 13.4714 | 1.13E-05 | 0.0684 | CCGACCGCCATCG | - | ABC transporter permease |
| AVAEIV_RS02045 | 408061 | 408073 | 118 | - | 13.4714 | 1.13E-05 | 0.0684 | CTGATGGCCAGCG | - | helix-turn-helix domain-containing protein |
| AVAEIV_RS16785 | 3633773 | 3633785 | 327 | - | 13.4714 | 1.13E-05 | 0.0684 | CTGAAGGCCAGCA | - | dicarboxylate/amino acid:cation symporter |
| AVAEIV_RS16790 | 3633773 | 3633785 | -41 | - | 13.4714 | 1.13E-05 | 0.0616 | CTGAAGGCCAGCA | - | dienelactone hydrolase family protein |
| AVAEIV_RS22300 | 4761724 | 4761736 | 44 | - | 13.4714 | 1.13E-05 | 0.0684 | CCGACCGCCATCG | - | O-antigen ligase family protein |
| AVAEIV_RS24165 | 5137250 | 5137262 | 354 | + | 13.4714 | 1.13E-05 | 0.0616 | CTGATGGCCAGCG | - | Lrp/AsnC ligand binding domain-containing protein |
| AVAEIV_RS24785 | 5268575 | 5268587 | 246 | + | 13.4714 | 1.13E-05 | 0.0616 | CCGGGCGCCATCA | - | containing protein |
| AVAEIV_RS10075 | 2106116 | 2106128 | 360 | - | 13.4571 | 1.14E-05 | 0.0616 | CGAAACGCCACCA | - | AtpZ/AtpI family protein |
| AVAEIV_RS11700 | 2478643 | 2478655 | 240 | - | 13.4571 | 1.14E-05 | 0.0616 | CGAAACGCCACCA | - | DUF1826 domain-containing protein |
| AVAEIV_RS17045 | 3698085 | 3698097 | 117 | + | 13.4571 | 1.14E-05 | 0.0688 | CCGTGGGCCAGCA | - | cupin |
| AVAEIV_RS21795 | 4641943 | 4641955 | 303 | + | 13.4571 | 1.14E-05 | 0.0688 | CGGATGGCCATCC | - | sugar ABC transporter substrate-binding protein |
| AVAEIV_RS21795 | 4641944 | 4641956 | 304 | - | 13.4571 | 1.14E-05 | 0.0688 | CGGATGGCCATCC | - | sugar ABC transporter substrate-binding protein |
| AVAEIV_RS21800 | 4641944 | 4641956 | 214 | - | 13.4571 | 1.14E-05 | 0.0616 | CGGATGGCCATCC | - | acyl-CoA dehydrogenase family protein |
| AVAEIV_RS21800 | 4641943 | 4641955 | 215 | + | 13.4571 | 1.14E-05 | 0.0616 | CGGATGGCCATCC | - | acyl-CoA dehydrogenase family protein |
| AVAEIV_RS07585 | 1582340 | 1582352 | 1 | - | 13.4429 | 1.16E-05 | 0.0617 | CAGCCCGCCATCG | - | FAD:protein FMN transferase |
| AVAEIV_RS14300 | 3117696 | 3117708 | 383 | + | 13.4429 | 1.16E-05 | 0.0689 | CGACTCGCCATCG | - | phytanoyl-CoA dioxygenase family protein |
| AVAEIV_RS22395 | 4788527 | 4788539 | 301 | - | 13.4429 | 1.16E-05 | 0.0617 | CCCAGCGCCACCA | *msrA* | peptide-methionine (S)-S-oxide reductase MsrA |
| AVAEIV_RS24735 | 5255244 | 5255256 | 73 | - | 13.4429 | 1.16E-05 | 0.0689 | CAGCCCGCCATCG | - | FadR/GntR family transcriptional regulator |
| AVAEIV_RS04075 | 827705 | 827717 | 8 | - | 13.4286 | 1.17E-05 | 0.0617 | CACGTGGCCATCG | *nadD* | nicotinate-nucleotide adenylyltransferase |
| AVAEIV_RS13665 | 2955882 | 2955894 | 368 | + | 13.4286 | 1.17E-05 | 0.0693 | CGGCGGGCCATCA | - | chromosome partitioning protein ParB |
| AVAEIV_RS21555 | 4585394 | 4585406 | 110 | + | 13.4286 | 1.17E-05 | 0.0617 | CACCTTGCCAGCG | - | (Fe-S)-binding protein |
| AVAEIV_RS02375 | 472703 | 472715 | 161 | - | 13.4143 | 1.19E-05 | 0.0693 | CAGTTCGACATCA | - | LysR family transcriptional regulator |
| AVAEIV_RS02380 | 472703 | 472715 | 46 | - | 13.4143 | 1.19E-05 | 0.0617 | CAGTTCGACATCA | - | aspartate aminotransferase family protein |
| AVAEIV_RS13650 | 2953945 | 2953957 | 242 | + | 13.4143 | 1.19E-05 | 0.0693 | CAGTCCGCCAGCG | - | type VI secretion protein |
| AVAEIV_RS15815 | 3450256 | 3450268 | 69 | - | 13.4143 | 1.19E-05 | 0.0693 | CGATTCGCCAGCG | - | phenol hydroxylase subunit P4 |
| AVAEIV_RS18115 | 3913250 | 3913262 | 272 | + | 13.4143 | 1.19E-05 | 0.0693 | CGGACCGCCATCG | *ybgC* | tol-pal system-associated acyl-CoA thioesterase |
| AVAEIV_RS21605 | 4599749 | 4599761 | 157 | + | 13.4143 | 1.19E-05 | 0.0693 | CAGTCCGCCAGCG | - | sugar diacid recognition domain-containing protein |
| AVAEIV_RS21610 | 4599749 | 4599761 | 159 | + | 13.4143 | 1.19E-05 | 0.0617 | CAGTCCGCCAGCG | *pyk* | pyruvate kinase |
| AVAEIV_RS23175 | 4949174 | 4949186 | 0 | - | 13.4143 | 1.19E-05 | 0.0617 | CGGATCGCCATCT | - | hypothetical protein |
| AVAEIV_RS07145 | 1473035 | 1473047 | 81 | + | 13.4 | 1.20E-05 | 0.0617 | CAGAAGGCCGGCA | *lepB* | signal peptidase I |
| AVAEIV_RS10510 | 2197381 | 2197393 | 385 | - | 13.4 | 1.20E-05 | 0.0617 | CCCTGCGCCAGCA | - | hypothetical protein |
| AVAEIV_RS13045 | 2826565 | 2826577 | 159 | - | 13.4 | 1.20E-05 | 0.0699 | CGCATCGCCATCC | - | YnfA family protein |
| AVAEIV_RS15910 | 3471523 | 3471535 | -19 | - | 13.4 | 1.20E-05 | 0.0699 | CGCATCGCCATCC | - | rhodanese-related sulfurtransferase |
| AVAEIV_RS22995 | 4924020 | 4924032 | 50 | - | 13.4 | 1.20E-05 | 0.0617 | CAGTTTGCCGGCA | - | DUF4411 family protein |
| AVAEIV_RS25380 | 5408618 | 5408630 | 171 | + | 13.4 | 1.20E-05 | 0.0617 | CAGAAGGCCGGCA | - | toxin-antitoxin system YwqK family antitoxin |
| AVAEIV_RS01600 | 309256 | 309268 | 314 | - | 13.3857 | 1.22E-05 | 0.07 | CTGCTCGCCACCA | - | metallophosphoesterase family protein |
| AVAEIV_RS03695 | 743848 | 743860 | 81 | - | 13.3857 | 1.22E-05 | 0.0617 | CGCAGCGCCACCA | *mutL* | DNA mismatch repair endonuclease MutL |
| AVAEIV_RS15660 | 3428289 | 3428301 | 21 | + | 13.3857 | 1.22E-05 | 0.07 | CTGCTCGCCACCA | - | MinD/ParA family protein |
| AVAEIV_RS15665 | 3428289 | 3428301 | 125 | + | 13.3857 | 1.22E-05 | 0.0617 | CTGCTCGCCACCA | *sodC* | superoxide dismutase family protein |
| AVAEIV_RS23150 | 4945258 | 4945270 | 330 | - | 13.3857 | 1.22E-05 | 0.0617 | CCGCTGGCCAGCG | - | hypothetical protein |
| AVAEIV_RS24035 | 5115435 | 5115447 | 256 | - | 13.3857 | 1.22E-05 | 0.0617 | CCGCAGGCCAGCA | *xerC* | tyrosine recombinase XerC |
| AVAEIV_RS10390 | 2172307 | 2172319 | 271 | + | 13.3714 | 1.23E-05 | 0.0617 | CCGGTCGCCAGCG | - | NIPSNAP family protein |
| AVAEIV_RS14110 | 3069529 | 3069541 | 342 | + | 13.3714 | 1.23E-05 | 0.0701 | CCGGACGCCAGCA | *xylB* | xylulokinase |
| AVAEIV_RS15270 | 3333378 | 3333390 | 166 | - | 13.3714 | 1.23E-05 | 0.0617 | CAGTTCGCCAGGA | - | extracellular solute-binding protein |
| AVAEIV_RS18065 | 3904012 | 3904024 | 12 | - | 13.3714 | 1.23E-05 | 0.0617 | CCGGTCGCCAGCG | *xerC* | tyrosine recombinase XerC |

| | | | | | | | | | | |
|---|---|---|---|---|---|---|---|---|---|---|
| AVAEIV_RS22320 | 4766160 | 4766172 | 5 | + | 13.3714 | 1.23E-05 | 0.0701 | CAGTTCGCCAGGA | - | hypothetical protein |
| AVAEIV_RS23120 | 4940304 | 4940316 | 368 | - | 13.3714 | 1.23E-05 | 0.0701 | CCGTAGGCCATCG | - | type II toxin-antitoxin system RelE/ParE family toxin |
| AVAEIV_RS25590 | 5469271 | 5469283 | 136 | - | 13.3571 | 1.24E-05 | 0.0617 | CCCTTGGCCACCA | *eno* | phosphopyruvate hydratase |
| AVAEIV_RS06390 | 1303717 | 1303729 | 35 | - | 13.3429 | 1.26E-05 | 0.0705 | CGCTGCGCCAGCA | - | UDP-glucose dehydrogenase family protein |
| AVAEIV_RS06395 | 1303717 | 1303729 | 131 | - | 13.3429 | 1.26E-05 | 0.0617 | CGCTGCGCCAGCA | - | alkaline phosphatase D family protein |
| AVAEIV_RS07745 | 1613022 | 1613034 | 125 | - | 13.3429 | 1.26E-05 | 0.0617 | CGCTGCGCCAGCA | - | PilZ domain-containing protein |
| AVAEIV_RS07845 | 1632901 | 1632913 | 15 | + | 13.3429 | 1.26E-05 | 0.0617 | CGCTGCGCCAGCA | - | OmpA family protein |
| AVAEIV_RS10720 | 2243729 | 2243741 | 59 | + | 13.3429 | 1.26E-05 | 0.0705 | CCGATCGCCGCCA | - | hypothetical protein |
| AVAEIV_RS11685 | 2477353 | 2477365 | 330 | - | 13.3429 | 1.26E-05 | 0.0705 | CGCTGCGCCAGCA | - | hypothetical protein |
| AVAEIV_RS11695 | 2477353 | 2477365 | 323 | - | 13.3429 | 1.26E-05 | 0.0617 | CGCTGCGCCAGCA | *zigA* | zinc metallochaperone GTPase ZigA |
| AVAEIV_RS14570 | 3173214 | 3173226 | 138 | - | 13.3429 | 1.26E-05 | 0.0617 | CACATCGCCGGCG | *aat* | leucyl/phenylalanyl-tRNA--protein transferase |
| AVAEIV_RS18760 | 4038485 | 4038497 | 261 | + | 13.3429 | 1.26E-05 | 0.0617 | CTGACCGCCATCA | - | DUF4129 domain-containing protein |
| AVAEIV_RS22700 | 4861451 | 4861463 | 138 | - | 13.3429 | 1.26E-05 | 0.0617 | CCGATCGCCGCCA | - | thioesterase family protein |
| AVAEIV_RS23270 | 4964442 | 4964454 | 321 | + | 13.3429 | 1.26E-05 | 0.0705 | CACATCGCCGGCG | *pdhA* | component subunit alpha |
| AVAEIV_RS23275 | 4964442 | 4964454 | 59 | + | 13.3429 | 1.26E-05 | 0.0705 | CACATCGCCGGCG | - | hypothetical protein |
| AVAEIV_RS03440 | 695327 | 695339 | 200 | - | 13.3286 | 1.28E-05 | 0.0705 | CCGCTCGCCGTCA | *accC* | acetyl-CoA carboxylase biotin carboxylase subunit |
| AVAEIV_RS04245 | 864859 | 864871 | 72 | - | 13.3286 | 1.28E-05 | 0.0623 | CCCCTCGCCAGCG | - | LysR family transcriptional regulator |
| AVAEIV_RS04945 | 1012351 | 1012363 | 270 | + | 13.3286 | 1.28E-05 | 0.0705 | CAGACGGCCAGCG | - | acyl-CoA dehydrogenase family protein |
| AVAEIV_RS07450 | 1552023 | 1552035 | 255 | - | 13.3286 | 1.28E-05 | 0.0623 | CGGCAGGCCAGCA | - | efflux transporter outer membrane subunit |
| AVAEIV_RS08485 | 1774560 | 1774572 | 252 | + | 13.3286 | 1.28E-05 | 0.0705 | CAGGGGGCCAGCA | - | ABC1 kinase family protein |
| AVAEIV_RS13615 | 2946023 | 2946035 | 377 | - | 13.3286 | 1.28E-05 | 0.0705 | CGAATGGCCAGCG | - | LysR family transcriptional regulator |
| AVAEIV_RS20075 | 4286058 | 4286070 | 130 | + | 13.3286 | 1.28E-05 | 0.0705 | CGAAAGGCCAGCA | - | sulfonate ABC transporter substrate-binding protein |
| AVAEIV_RS24965 | 5312786 | 5312798 | 113 | - | 13.3286 | 1.28E-05 | 0.0705 | CGGCTGGCCAGCG | *pyk* | pyruvate kinase |
| AVAEIV_RS12775 | 2715385 | 2715397 | 205 | + | 13.3143 | 1.30E-05 | 0.0705 | CAGCTCGCCGCCA | - | cupin-like domain-containing protein |
| AVAEIV_RS18985 | 4088639 | 4088651 | 351 | - | 13.3143 | 1.30E-05 | 0.0705 | CGGGTCGCCAGCG | - | alpha |
| AVAEIV_RS19055 | 4105068 | 4105080 | 306 | - | 13.3143 | 1.30E-05 | 0.0705 | CAGCTCGCCGCCA | *pyrH* | UMP kinase |
| AVAEIV_RS24650 | 5237096 | 5237108 | 85 | - | 13.3143 | 1.30E-05 | 0.0625 | CCCAGGGCCAGCA | - | pyridoxamine 5'-phosphate oxidase family protein |
| AVAEIV_RS00785 | 153624 | 153636 | 209 | - | 13.2857 | 1.32E-05 | 0.0625 | CGGATCGCCGCCA | *nifS* | cysteine desulfurase NifS |
| AVAEIV_RS02370 | 471221 | 471233 | 220 | - | 13.2857 | 1.32E-05 | 0.0705 | GCGATCGCCATCA | *trmL* | carboxymethylaminomethyluridine(34)-2'-O)-methyltransferase TrmL |
| AVAEIV_RS03910 | 788004 | 788016 | 74 | + | 13.2857 | 1.32E-05 | 0.0705 | CGGATCGCCGCCA | *adhP* | alcohol dehydrogenase AdhP |
| AVAEIV_RS07405 | 1537383 | 1537395 | 316 | - | 13.2857 | 1.32E-05 | 0.0625 | CGGATCGCCGCCA | - | acyl-CoA dehydrogenase family protein |
| AVAEIV_RS08410 | 1757210 | 1757222 | 140 | - | 13.2857 | 1.32E-05 | 0.0625 | CGGATCGCCGCCA | - | protein |
| AVAEIV_RS20360 | 4331458 | 4331470 | 127 | + | 13.2857 | 1.32E-05 | 0.0625 | CGGATCGCCGCCA | - | hypothetical protein |
| AVAEIV_RS23900 | 5093521 | 5093533 | 142 | + | 13.2857 | 1.32E-05 | 0.0705 | CGGATCGCCGCCA | *ppk1* | polyphosphate kinase 1 |
| AVAEIV_RS02805 | 575724 | 575736 | -37 | - | 13.2714 | 1.33E-05 | 0.0705 | CGGCTCGCCGTCA | - | DeoR/GlpR family DNA-binding transcription regulator |
| AVAEIV_RS02810 | 575724 | 575736 | 284 | - | 13.2714 | 1.33E-05 | 0.0625 | CGGCTCGCCGTCA | - | M48 family metallopeptidase |
| AVAEIV_RS03215 | 651034 | 651046 | 218 | - | 13.2714 | 1.33E-05 | 0.0625 | CAGACCGCCGTCA | *rpmJ* | 50S ribosomal protein L36 |
| AVAEIV_RS06535 | 1349364 | 1349376 | 275 | + | 13.2714 | 1.33E-05 | 0.0705 | CAGTTCGGCAGCA | - | ABC transporter ATP-binding protein |
| AVAEIV_RS10330 | 2157515 | 2157527 | 218 | - | 13.2714 | 1.33E-05 | 0.0625 | CCGAACGCCATCC | - | shikimate dehydrogenase family protein |
| AVAEIV_RS14210 | 3096048 | 3096060 | 368 | + | 13.2714 | 1.33E-05 | 0.0705 | CACACCGCCAGCG | *sohB* | protease SohB |
| AVAEIV_RS20685 | 4398009 | 4398021 | 195 | + | 13.2714 | 1.33E-05 | 0.0705 | CCTATTGCCATCA | - | TIGR01777 family oxidoreductase |
| AVAEIV_RS20690 | 4398009 | 4398021 | -22 | + | 13.2714 | 1.33E-05 | 0.0625 | CCTATTGCCATCA | - | NAD(P)/FAD-dependent oxidoreductase |
| AVAEIV_RS24090 | 5125023 | 5125035 | 276 | - | 13.2714 | 1.33E-05 | 0.0625 | CGGCTCGCCGTCA | - | DoxX family protein |
| AVAEIV_RS00310 | 68371 | 68383 | 202 | + | 13.2571 | 1.35E-05 | 0.0626 | CGCAGGGCCAGCA | - | DUF2782 domain-containing protein |
| AVAEIV_RS00775 | 152138 | 152150 | 367 | - | 13.2571 | 1.35E-05 | 0.0626 | CGCAGGGCCAGCA | - | HesB/IscA family protein |
| AVAEIV_RS03605 | 726721 | 726733 | 260 | + | 13.2571 | 1.35E-05 | 0.0705 | CGCAGGGCCAGCA | - | binding protein |
| AVAEIV_RS08400 | 1753051 | 1753063 | 185 | + | 13.2571 | 1.35E-05 | 0.0626 | CGCTACGCCATCG | *fadB* | fatty acid oxidation complex subunit alpha FadB |
| AVAEIV_RS13540 | 2931301 | 2931313 | 283 | - | 13.2571 | 1.35E-05 | 0.0705 | CCGAGCGCCACCG | - | NAD(P)-dependent alcohol dehydrogenase |
| AVAEIV_RS00040 | 9137 | 9149 | 338 | - | 13.2429 | 1.37E-05 | 0.0705 | CCGCGCGCCATCG | - | hypothetical protein |
| AVAEIV_RS00320 | 71135 | 71147 | 365 | - | 13.2429 | 1.37E-05 | 0.0705 | CCGCGCGCCATCG | - | alpha/beta fold hydrolase |
| AVAEIV_RS09015 | 1876387 | 1876399 | 141 | - | 13.2429 | 1.37E-05 | 0.0705 | CTGGTCGCCAGCA | - | phosphonate ABC transporter ATP-binding protein |

| | | | | | | | | | | |
|---|---|---|---|---|---|---|---|---|---|---|
| AVAEIV_RS11190 | 2362315 | 2362327 | 33 | + | 13.2429 | 1.37E-05 | 0.0705 | CGGTTCGCCGGCA | - | peroxiredoxin-like family protein |
| AVAEIV_RS13930 | 3020159 | 3020171 | 93 | + | 13.2429 | 1.37E-05 | 0.063 | CCGCGCGCCATCG | - | sugar kinase |
| AVAEIV_RS16390 | 3560510 | 3560522 | 371 | - | 13.2429 | 1.37E-05 | 0.0705 | CTGTTGGCCATCG | - | holin family protein |
| AVAEIV_RS16395 | 3560510 | 3560522 | 51 | - | 13.2429 | 1.37E-05 | 0.0705 | CTGTTGGCCATCG | - | hypothetical protein |
| AVAEIV_RS25050 | 5332058 | 5332070 | 368 | - | 13.2429 | 1.37E-05 | 0.063 | CTGGTCGCCAGCA | *fahA* | fumarylacetoacetase |
| AVAEIV_RS25390 | 5410440 | 5410452 | 121 | - | 13.2429 | 1.37E-05 | 0.0705 | CAGGAGGCCATCG | - | protein |
| AVAEIV_RS00450 | 91779 | 91791 | 292 | + | 13.2286 | 1.38E-05 | 0.0631 | CCCAAGGCCATCG | - | cytochrome c |
| AVAEIV_RS10165 | 2121318 | 2121330 | 381 | + | 13.2286 | 1.38E-05 | 0.0708 | CAGCGCGCCACCG | *cydX* | cytochrome bd-I oxidase subunit CydX |
| AVAEIV_RS15960 | 3479962 | 3479974 | 197 | + | 13.2286 | 1.38E-05 | 0.0708 | CCCTTTGCCATCG | - | malate dehydrogenase |
| AVAEIV_RS21355 | 4539466 | 4539478 | 222 | + | 13.2286 | 1.38E-05 | 0.0631 | CCCAAGGCCATCG | - | sigma-54-dependent transcriptional regulator |
| AVAEIV_RS03760 | 757419 | 757431 | 239 | + | 13.2143 | 1.40E-05 | 0.0631 | CCGATGGCCGGCA | *rpsF* | 30S ribosomal protein S6 |
| AVAEIV_RS06380 | 1300290 | 1300302 | 160 | + | 13.2143 | 1.40E-05 | 0.0631 | CAGTACGCCAGCC | *ampE* | regulatory signaling modulator protein AmpE |
| AVAEIV_RS17220 | 3734427 | 3734439 | -10 | - | 13.2143 | 1.40E-05 | 0.071 | CCGATGGCCGGCA | *leuC* | 3-isopropylmalate dehydratase large subunit |
| AVAEIV_RS17225 | 3734427 | 3734439 | 169 | - | 13.2143 | 1.40E-05 | 0.0631 | CCGATGGCCGGCA | - | LysR family transcriptional regulator |
| AVAEIV_RS21740 | 4632239 | 4632251 | 347 | - | 13.2143 | 1.40E-05 | 0.071 | CCGATGGCCGGCA | - | hypothetical protein |
| AVAEIV_RS25480 | 5427109 | 5427121 | 184 | + | 13.2143 | 1.40E-05 | 0.071 | CAACGCGCCATCG | *norR* | nitric oxide reductase transcriptional regulator NorR |
| AVAEIV_RS25485 | 5427109 | 5427121 | -28 | + | 13.2143 | 1.40E-05 | 0.0631 | CAACGCGCCATCG | *hmpA* | NO-inducible flavohemoprotein |
| AVAEIV_RS18085 | 3908357 | 3908369 | 240 | + | 13.2 | 1.41E-05 | 0.071 | CAGATCGCGAGCA | *ybgF* | tol-pal system protein YbgF |
| AVAEIV_RS04455 | 908054 | 908066 | -28 | - | 13.1857 | 1.42E-05 | 0.071 | CTCTACGCCATCA | *thiE* | thiamine phosphate synthase |
| AVAEIV_RS13295 | 2880658 | 2880670 | 361 | - | 13.1857 | 1.42E-05 | 0.071 | CTCTTCGCCATCG | *maiA* | maleylacetoacetate isomerase |
| AVAEIV_RS18675 | 4023000 | 4023012 | 146 | - | 13.1857 | 1.42E-05 | 0.071 | CAGAGCGCCATCT | - | DMT family transporter |
| AVAEIV_RS19865 | 4232544 | 4232556 | 274 | + | 13.1857 | 1.42E-05 | 0.071 | CGGCGCGCCATCG | - | CoA pyrophosphatase |
| AVAEIV_RS24735 | 5255305 | 5255317 | 134 | - | 13.1857 | 1.42E-05 | 0.071 | CGAAGCGCCATCG | - | FadR/GntR family transcriptional regulator |
| AVAEIV_RS01640 | 318163 | 318175 | 286 | - | 13.1714 | 1.44E-05 | 0.0711 | CAGTTGGCCGTCG | - | methyl-accepting chemotaxis protein |
| AVAEIV_RS05435 | 1124380 | 1124392 | 155 | - | 13.1714 | 1.44E-05 | 0.0635 | CAGGTCGCCGGCA | *ileS* | isoleucine--tRNA ligase |
| AVAEIV_RS16890 | 3657794 | 3657806 | 41 | + | 13.1714 | 1.44E-05 | 0.0711 | CGGTCCGCCAGCA | - | alpha/beta hydrolase family esterase |
| AVAEIV_RS18095 | 3910298 | 3910310 | 283 | - | 13.1714 | 1.44E-05 | 0.0711 | CAGGTCGCCGGCA | *tolB* | Tol-Pal system beta propeller repeat protein TolB |
| AVAEIV_RS11005 | 2316323 | 2316335 | 238 | + | 13.1571 | 1.46E-05 | 0.0635 | CAAATTGCCATCC | - | OprD family porin |
| AVAEIV_RS24015 | 5112427 | 5112439 | 274 | + | 13.1571 | 1.46E-05 | 0.0635 | CAGAACGCCGGCG | *lptM* | LPS translocon maturation chaperone LptM |
| AVAEIV_RS13285 | 2877469 | 2877481 | 183 | - | 13.1429 | 1.47E-05 | 0.0635 | CCGACGGCCAGCA | *ltrA* | group II intron reverse transcriptase/maturase |
| AVAEIV_RS16655 | 3609233 | 3609245 | 359 | - | 13.1429 | 1.47E-05 | 0.0635 | CAATAGGCCACCA | - | hypothetical protein |
| AVAEIV_RS25890 | 5554952 | 5554964 | 105 | - | 13.1429 | 1.47E-05 | 0.0713 | CTGATCGCCATCC | - | formylglycine-generating enzyme family protein |
| AVAEIV_RS04995 | 1022063 | 1022075 | 63 | - | 13.1286 | 1.48E-05 | 0.0635 | CCGAGGGCCAGCG | - | alpha-ketoacid dehydrogenase subunit beta |
| AVAEIV_RS05780 | 1185088 | 1185100 | 172 | + | 13.1286 | 1.48E-05 | 0.0713 | CCGAGGGCCAGCG | - | 3-oxoadipyl-CoA thiolase |
| AVAEIV_RS07180 | 1480471 | 1480483 | 12 | - | 13.1286 | 1.48E-05 | 0.0713 | CACCTCGCCGGCA | - | DUF2905 domain-containing protein |
| AVAEIV_RS08955 | 1862071 | 1862083 | 159 | - | 13.1286 | 1.48E-05 | 0.0635 | CTGAGCGCCACCA | - | acyl-CoA dehydrogenase |
| AVAEIV_RS12895 | 2782725 | 2782737 | 72 | - | 13.1286 | 1.48E-05 | 0.0713 | CAGAAGGCCAGCC | - | aspartate aminotransferase family protein |
| AVAEIV_RS19215 | 4139076 | 4139088 | 301 | - | 13.1286 | 1.48E-05 | 0.0713 | CACATCGGCAGCA | - | hypothetical protein |
| AVAEIV_RS21890 | 4667075 | 4667087 | 318 | - | 13.1286 | 1.48E-05 | 0.0713 | CCGAGGGCCAGCG | - | amidohydrolase family protein |
| AVAEIV_RS26105 | 5612840 | 5612852 | 297 | - | 13.1286 | 1.48E-05 | 0.0713 | CACATCGGCAGCA | - | tyrosine-type recombinase/integrase |
| AVAEIV_RS00575 | 118568 | 118580 | -29 | - | 13.1143 | 1.49E-05 | 0.0635 | CAGCCGGCCAGCA | - | hypothetical protein |
| AVAEIV_RS00980 | 193335 | 193347 | -47 | + | 13.1143 | 1.49E-05 | 0.0713 | CAGCCGGCCAGCA | - | N-acetylmuramoyl-L-alanine amidase |
| AVAEIV_RS03830 | 772449 | 772461 | 294 | - | 13.1143 | 1.49E-05 | 0.0713 | CCCTTCGCCACCG | *trmA* | tRNA (uridine(54)-C5)-methyltransferase TrmA |
| AVAEIV_RS12405 | 2627926 | 2627938 | 252 | - | 13.1143 | 1.49E-05 | 0.0635 | CACTTCGCCGTCG | - | DEAD/DEAH box helicase family protein |
| AVAEIV_RS16965 | 3675821 | 3675833 | 202 | + | 13.1143 | 1.49E-05 | 0.0713 | CAGCCGGCCAGCA | - | sugar phosphate isomerase/epimerase family protein |
| AVAEIV_RS18770 | 4041036 | 4041048 | 345 | - | 13.1143 | 1.49E-05 | 0.0635 | CAGCCGGCCAGCA | - | AAA family ATPase |
| AVAEIV_RS20520 | 4364969 | 4364981 | 365 | + | 13.1143 | 1.49E-05 | 0.0713 | CAGCCGGCCAGCA | *radA* | DNA repair protein RadA |
| AVAEIV_RS20525 | 4364969 | 4364981 | 46 | + | 13.1143 | 1.49E-05 | 0.0713 | CAGCCGGCCAGCA | - | YdcH family protein |
| AVAEIV_RS20530 | 4364969 | 4364981 | 167 | + | 13.1143 | 1.49E-05 | 0.0635 | CAGCCGGCCAGCA | *mscL* | MscL |
| AVAEIV_RS20910 | 4445459 | 4445471 | 122 | - | 13.1143 | 1.49E-05 | 0.0635 | CACTTCGCCGTCG | - | alpha-ketoacid dehydrogenase subunit beta |
| AVAEIV_RS00090 | 20395 | 20407 | 176 | - | 13.1 | 1.51E-05 | 0.0713 | CAGGCCGCCAGCA | *def* | peptide deformylase |
| AVAEIV_RS00095 | 20395 | 20407 | -38 | - | 13.1 | 1.51E-05 | 0.0635 | CAGGCCGCCAGCA | - | LysM peptidoglycan-binding domain-containing protein |

| | | | | | | | | | | |
|---|---|---|---|---|---|---|---|---|---|---|
| AVAEIV_RS03640 | 733184 | 733196 | 252 | - | 13.1 | 1.51E-05 | 0.0713 | CAGCGGGCCAGCG | - | AhpA/YtjB family protein |
| AVAEIV_RS12920 | 2801495 | 2801507 | 237 | + | 13.1 | 1.51E-05 | 0.0713 | CAGTCGGCCATCG | - | DNA-binding protein |
| AVAEIV_RS18765 | 4040209 | 4040221 | 30 | - | 13.1 | 1.51E-05 | 0.0635 | CAGGCCGCCAGCA | - | DUF4350 domain-containing protein |
| AVAEIV_RS22125 | 4721401 | 4721413 | 361 | - | 13.1 | 1.51E-05 | 0.0635 | CGCATCGCCGGCA | - | hypothetical protein |
| AVAEIV_RS00765 | 151205 | 151217 | 42 | + | 13.0857 | 1.52E-05 | 0.0713 | CAGGGCGCCAGCG | *modC* | molybdenum ABC transporter ATP-binding protein |
| AVAEIV_RS00770 | 151205 | 151217 | 316 | + | 13.0857 | 1.52E-05 | 0.0635 | CAGGGCGCCAGCG | - | hypothetical protein |
| AVAEIV_RS22040 | 4704048 | 4704060 | 325 | + | 13.0857 | 1.52E-05 | 0.0713 | CCCACCGCCAGCA | - | ion channel |
| AVAEIV_RS22735 | 4868039 | 4868051 | 234 | + | 13.0857 | 1.52E-05 | 0.0713 | CCCACCGCCAGCA | - | hypothetical protein |
| AVAEIV_RS01055 | 206590 | 206602 | 325 | + | 13.0714 | 1.54E-05 | 0.0635 | CAGAATGCCGGCA | *trpB* | tryptophan synthase subunit beta |
| AVAEIV_RS04240 | 864106 | 864118 | 289 | - | 13.0714 | 1.54E-05 | 0.0635 | CCCAGCGCCAGCG | - | GlcG/HbpS family heme-binding protein |
| AVAEIV_RS06920 | 1423098 | 1423110 | 272 | - | 13.0714 | 1.54E-05 | 0.0635 | CCCAGCGCCAGCG | *murD* | UDP-N-acetylmuramoyl-L-alanine--D-glutamate ligase |
| AVAEIV_RS07440 | 1546101 | 1546113 | -43 | + | 13.0714 | 1.54E-05 | 0.0635 | CCGATCGCCACCC | - | efflux RND transporter permease subunit |
| AVAEIV_RS07810 | 1625114 | 1625126 | 246 | - | 13.0714 | 1.54E-05 | 0.0635 | CCCAGCGCCAGCG | *mhpT* | 3-(3-hydroxy-phenyl)propionate transporter MhpT |
| AVAEIV_RS08880 | 1844036 | 1844048 | 230 | - | 13.0714 | 1.54E-05 | 0.0635 | CCCAGCGCCAGCG | *cas6e* | type I-E CRISPR-associated protein Cas6/Cse3/CasE |
| AVAEIV_RS11025 | 2322284 | 2322296 | 91 | - | 13.0714 | 1.54E-05 | 0.0635 | CAGATTGCCGGCG | - | TauD/TfdA dioxygenase family protein |
| AVAEIV_RS12385 | 2623534 | 2623546 | 111 | + | 13.0714 | 1.54E-05 | 0.0713 | CCCAGCGCCAGCG | - | ABC transporter permease |
| AVAEIV_RS18840 | 4057398 | 4057410 | 163 | - | 13.0714 | 1.54E-05 | 0.0713 | CAGATTGCCGGCG | - | phosphoketolase |
| AVAEIV_RS18845 | 4057398 | 4057410 | 29 | - | 13.0714 | 1.54E-05 | 0.0635 | CAGATTGCCGGCG | - | containing protein |
| AVAEIV_RS21060 | 4478739 | 4478751 | 238 | + | 13.0714 | 1.54E-05 | 0.0713 | CCCAGCGCCAGCG | *mhpT* | 3-(3-hydroxy-phenyl)propionate transporter MhpT |
| AVAEIV_RS21320 | 4532437 | 4532449 | 193 | + | 13.0714 | 1.54E-05 | 0.0635 | CCCAGCGCCAGCG | - | Rieske (2Fe-2S) protein |
| AVAEIV_RS21730 | 4630890 | 4630902 | 352 | + | 13.0714 | 1.54E-05 | 0.0713 | CCCAGCGCCAGCG | - | branched-chain amino acid ABC transporter permease |
| AVAEIV_RS02425 | 482081 | 482093 | 23 | + | 13.0571 | 1.55E-05 | 0.0713 | CCGCATGCCAGCA | - | GspE/PulE family protein |
| AVAEIV_RS09580 | 1996619 | 1996631 | 385 | + | 13.0571 | 1.55E-05 | 0.0635 | CCGGTGGCCATCG | - | dipeptide ABC transporter ATP-binding protein |
| AVAEIV_RS14385 | 3130952 | 3130964 | 301 | + | 13.0571 | 1.55E-05 | 0.0713 | CCAATTGCCAGCG | - | flagellar protein FlhE |
| AVAEIV_RS23050 | 4932222 | 4932234 | 256 | + | 13.0571 | 1.55E-05 | 0.0635 | CAGAGCGCCGCCA | - | phage major capsid protein, P2 family |
| AVAEIV_RS23845 | 5082376 | 5082388 | 30 | - | 13.0571 | 1.55E-05 | 0.0713 | CAGAGCGCCGCCA | - | YecA family protein |
| AVAEIV_RS23850 | 5082376 | 5082388 | 144 | - | 13.0571 | 1.55E-05 | 0.0635 | CAGAGCGCCGCCA | - | TIGR02449 family protein |
| AVAEIV_RS23855 | 5082376 | 5082388 | 350 | - | 13.0571 | 1.55E-05 | 0.0635 | CAGAGCGCCGCCA | - | cell division protein ZapA |
| AVAEIV_RS03130 | 643292 | 643304 | 192 | - | 13.0429 | 1.56E-05 | 0.0635 | CACCGCGCCAGCG | *rpsS* | 30S ribosomal protein S19 |
| AVAEIV_RS05055 | 1039384 | 1039396 | 149 | + | 13.0429 | 1.56E-05 | 0.0714 | CAGCTCGCCACCC | *algG* | mannuronan 5-epimerase AlgG |
| AVAEIV_RS06550 | 1352853 | 1352865 | 210 | - | 13.0429 | 1.56E-05 | 0.0714 | CAGCTCGCCACCC | - | ABC transporter substrate-binding protein |
| AVAEIV_RS10010 | 2096518 | 2096530 | 255 | - | 13.0429 | 1.56E-05 | 0.0635 | CAGATCGCCAGGG | - | Na+/H+ antiporter subunit E |
| AVAEIV_RS12650 | 2681738 | 2681750 | 205 | - | 13.0429 | 1.56E-05 | 0.0635 | CAGCTCGCCACCC | - | hypothetical protein |
| AVAEIV_RS18395 | 3962843 | 3962855 | 313 | + | 13.0429 | 1.56E-05 | 0.0714 | CAGCGCGCCGTCA | *relA* | GTP diphosphokinase |
| AVAEIV_RS22520 | 4822198 | 4822210 | 153 | - | 13.0286 | 1.58E-05 | 0.0721 | CACATGGCCGTCG | - | AAA family ATPase |
| AVAEIV_RS02965 | 613666 | 613678 | 204 | + | 13.0143 | 1.60E-05 | 0.0721 | TACATCGCCAGCA | - | ComF family protein |
| AVAEIV_RS05165 | 1059955 | 1059967 | 159 | - | 13.0143 | 1.60E-05 | 0.0635 | CTGCTCGCCAGCG | *coxB* | cytochrome c oxidase subunit II |
| AVAEIV_RS14170 | 3084163 | 3084175 | 348 | + | 13.0143 | 1.60E-05 | 0.0721 | CAGATCGCCAGTG | - | DUF2149 domain-containing protein |
| AVAEIV_RS14985 | 3264728 | 3264740 | -40 | + | 13.0143 | 1.60E-05 | 0.0635 | CTGCTCGCCAGCG | - | peptidylprolyl isomerase |
| AVAEIV_RS18645 | 4013152 | 4013164 | 168 | + | 13.0143 | 1.60E-05 | 0.0635 | CTGCTCGCCAGCG | - | shikimate dehydrogenase |
| AVAEIV_RS19510 | 4171439 | 4171451 | 160 | + | 13.0143 | 1.60E-05 | 0.0635 | CCCCTGGCCATCG | - | hypothetical protein |
| AVAEIV_RS21990 | 4692852 | 4692864 | 228 | + | 13.0143 | 1.60E-05 | 0.0635 | CCCCTGGCCATCG | - | paraquat-inducible protein A |
| AVAEIV_RS22095 | 4715496 | 4715508 | 140 | - | 13.0143 | 1.60E-05 | 0.0635 | CGCAGCGCCAGCG | - | nucleotidyltransferase family protein |
| AVAEIV_RS23575 | 5024915 | 5024927 | 328 | - | 13.0143 | 1.60E-05 | 0.0635 | TACATCGCCAGCA | *rsmA* | 16S rRNA (adenine(1518)-N(6)/adenine(1519)-N(6))-dimethyltransferase RsmA |
| AVAEIV_RS25195 | 5364833 | 5364845 | 61 | + | 13.0143 | 1.60E-05 | 0.0721 | CAGTGCGCCGGCA | *hybE* | [NiFe]-hydrogenase assembly chaperone HybE |
| AVAEIV_RS03825 | 770312 | 770324 | 233 | - | 13 | 1.62E-05 | 0.0635 | CCCGTCGCCATCG | - | DUF4345 family protein |
| AVAEIV_RS09130 | 1902588 | 1902600 | 349 | - | 13 | 1.62E-05 | 0.0723 | CCCGTCGCCATCG | - | lipopolysaccharide kinase InaA family protein |
| AVAEIV_RS09785 | 2046911 | 2046923 | 31 | + | 13 | 1.62E-05 | 0.0723 | CGAATTGCCAGCG | *egtD* | L-histidine N(alpha)-methyltransferase |
| AVAEIV_RS10970 | 2306549 | 2306561 | 329 | + | 13 | 1.62E-05 | 0.0635 | CCATGCGCCAGCA | - | MFS transporter |
| AVAEIV_RS15325 | 3348971 | 3348983 | 267 | - | 13 | 1.62E-05 | 0.0723 | CAGACCGCCATCC | *lpdA* | dihydrolipoyl dehydrogenase |
| AVAEIV_RS15385 | 3359697 | 3359709 | 380 | - | 13 | 1.62E-05 | 0.0635 | CCCGACGCCATCA | - | PA1571 family protein |

| | | | | | | | | | | |
|---|---|---|---|---|---|---|---|---|---|---|
| AVAEIV_RS18570 | 4001591 | 4001603 | -11 | + | 13 | 1.62E-05 | 0.0723 | CGGGTGGCCATCG | - | LysR substrate-binding domain-containing protein |
| AVAEIV_RS00040 | 8785 | 8797 | -14 | + | 12.9857 | 1.64E-05 | 0.0723 | CCGTTGGCCGTCA | - | hypothetical protein |
| AVAEIV_RS02865 | 588250 | 588262 | 20 | + | 12.9857 | 1.64E-05 | 0.0635 | CGGCGCGCCACCA | - | containing protein |
| AVAEIV_RS07360 | 1525197 | 1525209 | 302 | + | 12.9857 | 1.64E-05 | 0.0635 | CCCTTGGCCAGCG | - | ATP-binding cassette domain-containing protein |
| AVAEIV_RS14835 | 3227761 | 3227773 | -38 | + | 12.9857 | 1.64E-05 | 0.0723 | CGGCGCGCCACCA | - | antibiotic biosynthesis monooxygenase |
| AVAEIV_RS14840 | 3227761 | 3227773 | 277 | + | 12.9857 | 1.64E-05 | 0.0635 | CGGCGCGCCACCA | - | SOS response-associated peptidase family protein |
| AVAEIV_RS00590 | 120711 | 120723 | -33 | - | 12.9714 | 1.65E-05 | 0.0635 | CAGTTGGCCGCCA | - | 5-(carboxyamino)imidazole ribonucleotide synthase |
| AVAEIV_RS01490 | 287398 | 287410 | 184 | - | 12.9714 | 1.65E-05 | 0.0723 | CGCATGGCCACCG | - | YggT family protein |
| AVAEIV_RS14570 | 3173242 | 3173254 | 110 | + | 12.9571 | 1.66E-05 | 0.0635 | CGCCAGGCCATCA | *aat* | leucyl/phenylalanyl-tRNA--protein transferase |
| AVAEIV_RS05365 | 1110450 | 1110462 | 179 | - | 12.9429 | 1.68E-05 | 0.0723 | CAAATAGCCATCA | - | cyclase/phosphodiesterase |
| AVAEIV_RS07635 | 1592678 | 1592690 | 181 | + | 12.9429 | 1.68E-05 | 0.0635 | CAGCTCGCCGGCG | - | ExbD/TolR family protein |
| AVAEIV_RS10945 | 2299154 | 2299166 | 261 | + | 12.9429 | 1.68E-05 | 0.0723 | CAGCACGCCGGCA | - | 2,3-dihydro-2,3-dihydroxybenzoate dehydrogenase |
| AVAEIV_RS11025 | 2322242 | 2322254 | 133 | - | 12.9429 | 1.68E-05 | 0.0635 | CAGCTCGCCGGCG | - | TauD/TfdA dioxygenase family protein |
| AVAEIV_RS11725 | 2485843 | 2485855 | 218 | - | 12.9429 | 1.68E-05 | 0.0723 | CGCGTCGCCATCG | - | ABC transporter permease |
| AVAEIV_RS12290 | 2603370 | 2603382 | 375 | - | 12.9429 | 1.68E-05 | 0.0723 | CAGCTCGCCGGCG | *fliF* | flagellar basal-body MS-ring/collar protein FliF |
| AVAEIV_RS12300 | 2603370 | 2603382 | 330 | - | 12.9429 | 1.68E-05 | 0.0635 | CAGCTCGCCGGCG | *fliD* | flagellar filament capping protein FliD |
| AVAEIV_RS17330 | 3753541 | 3753553 | 172 | - | 12.9429 | 1.68E-05 | 0.0723 | CCGATGGCCAGCC | *alaS* | alanine--tRNA ligase |
| AVAEIV_RS19785 | 4216943 | 4216955 | 125 | + | 12.9429 | 1.68E-05 | 0.0723 | CTCAGCGCCAGCA | - | YaeQ family protein |
| AVAEIV_RS21320 | 4532275 | 4532287 | 355 | + | 12.9429 | 1.68E-05 | 0.0635 | CTCAGCGCCAGCA | - | Rieske (2Fe-2S) protein |
| AVAEIV_RS21805 | 4643380 | 4643392 | 63 | + | 12.9429 | 1.68E-05 | 0.0635 | CAGATCGGCAGCG | - | SfnB family sulfur acquisition oxidoreductase |
| AVAEIV_RS23570 | 5023936 | 5023948 | 191 | - | 12.9429 | 1.68E-05 | 0.0635 | CAGCTCGCCGGCG | *pdxA* | 4-hydroxythreonine-4-phosphate dehydrogenase PdxA |
| AVAEIV_RS25975 | 5580783 | 5580795 | 177 | + | 12.9429 | 1.68E-05 | 0.0723 | CAGAACGGCAGCA | *atpE* | F0F1 ATP synthase subunit C |
| AVAEIV_RS02520 | 505314 | 505326 | 356 | + | 12.9286 | 1.69E-05 | 0.0635 | CTGGTGGCCATCA | - | ExbD/TolR family protein |
| AVAEIV_RS17285 | 3745629 | 3745641 | 312 | + | 12.9286 | 1.69E-05 | 0.0723 | CTAATTGCCAGCA | - | tRNA-Arg |
| AVAEIV_RS17290 | 3745629 | 3745641 | 27 | + | 12.9286 | 1.69E-05 | 0.0635 | CTAATTGCCAGCA | - | Arc family DNA-binding protein |
| AVAEIV_RS17740 | 3842194 | 3842206 | 378 | - | 12.9286 | 1.69E-05 | 0.0723 | CGCTTGGCCAGCG | *dapA* | 4-hydroxy-tetrahydrodipicolinate synthase |
| AVAEIV_RS17750 | 3842194 | 3842206 | 340 | - | 12.9286 | 1.69E-05 | 0.0635 | CGCTTGGCCAGCG | - | peroxiredoxin |
| AVAEIV_RS03185 | 647849 | 647861 | 30 | - | 12.9143 | 1.71E-05 | 0.0635 | CACTTCGCCGCCA | *rplF* | 50S ribosomal protein L6 |
| AVAEIV_RS05115 | 1053014 | 1053026 | -49 | - | 12.9143 | 1.71E-05 | 0.0723 | CAGTTCGCCTTCA | - | cytochrome ubiquinol oxidase subunit I |
| AVAEIV_RS08165 | 1697113 | 1697125 | 246 | - | 12.9143 | 1.71E-05 | 0.0635 | CCGTCGGCCATCA | *cmk* | (d)CMP kinase |
| AVAEIV_RS09255 | 1932339 | 1932351 | 168 | - | 12.9143 | 1.71E-05 | 0.0635 | CAGGATGCCATCG | - | MFS family transporter |
| AVAEIV_RS11025 | 2322318 | 2322330 | 57 | + | 12.9143 | 1.71E-05 | 0.0635 | CGGATCGCCGGCG | - | TauD/TfdA dioxygenase family protein |
| AVAEIV_RS14760 | 3210751 | 3210763 | 380 | - | 12.9143 | 1.71E-05 | 0.0635 | CGGATCGCCGGCG | *nuoH* | NADH-quinone oxidoreductase subunit NuoH |
| AVAEIV_RS17420 | 3772344 | 3772356 | -48 | - | 12.9143 | 1.71E-05 | 0.0635 | CGGATCGCCGGCG | *ggpS* | glucosylglycerol-phosphate synthase |
| AVAEIV_RS18420 | 3969776 | 3969788 | 280 | - | 12.9143 | 1.71E-05 | 0.0723 | CACATGGCCATGA | - | FxLYD domain-containing protein |
| AVAEIV_RS18430 | 3969776 | 3969788 | 199 | - | 12.9143 | 1.71E-05 | 0.0635 | CACATGGCCATGA | - | tRNA-Met |
| AVAEIV_RS18830 | 4054362 | 4054374 | 172 | + | 12.9143 | 1.71E-05 | 0.0723 | CGGATCGCCGGCG | - | tRNA-uridine aminocarboxypropyltransferase |
| AVAEIV_RS18835 | 4054362 | 4054374 | 19 | + | 12.9143 | 1.71E-05 | 0.0635 | CGGATCGCCGGCG | - | protein |
| AVAEIV_RS19870 | 4232102 | 4232114 | 267 | + | 12.9143 | 1.71E-05 | 0.0635 | CACTTCGCCGCCA | - | NUDIX hydrolase |
| AVAEIV_RS25205 | 5366165 | 5366177 | 320 | + | 12.9143 | 1.71E-05 | 0.0723 | CGGATCGCCGGCG | - | hydrogenase expression/formation protein |
| AVAEIV_RS25415 | 5414041 | 5414053 | 180 | - | 12.9143 | 1.71E-05 | 0.0723 | CCAAGGGCCAGCA | - | RnfABCDGE type electron transport complex subunit D |
| AVAEIV_RS01465 | 284249 | 284261 | 274 | - | 12.9 | 1.73E-05 | 0.0723 | GAGATCGCCAGCG | *rdgB* | pyrophosphatase |
| AVAEIV_RS03705 | 746735 | 746747 | 258 | - | 12.9 | 1.73E-05 | 0.0635 | CAGTAGGCCATCC | *hfq* | RNA chaperone Hfq |
| AVAEIV_RS03725 | 750465 | 750477 | 330 | + | 12.9 | 1.73E-05 | 0.0635 | CCGACCGCCAGCG | - | DUF2065 domain-containing protein |
| AVAEIV_RS06505 | 1339457 | 1339469 | 339 | - | 12.9 | 1.73E-05 | 0.0635 | GAGATCGCCAGCG | - | TonB-dependent receptor |
| AVAEIV_RS09700 | 2033306 | 2033318 | 252 | + | 12.9 | 1.73E-05 | 0.0723 | CCGACCGCCAGCG | - | deoxyguanosinetriphosphate triphosphohydrolase |
| AVAEIV_RS13300 | 2881827 | 2881839 | 133 | + | 12.9 | 1.73E-05 | 0.0723 | CAGGTCGCCAGCC | - | MFS transporter |
| AVAEIV_RS15935 | 3475092 | 3475104 | 109 | + | 12.9 | 1.73E-05 | 0.0635 | GAGATCGCCAGCG | - | NADPH-dependent F420 reductase |
| AVAEIV_RS16620 | 3604405 | 3604417 | 139 | + | 12.9 | 1.73E-05 | 0.0723 | CCGGGCGCCAGCA | - | YihY/virulence factor BrkB family protein |
| AVAEIV_RS17325 | 3750879 | 3750891 | 229 | + | 12.9 | 1.73E-05 | 0.0723 | CAGGTCGCCAGCC | - | aspartate kinase |
| AVAEIV_RS21495 | 4574363 | 4574375 | 234 | + | 12.9 | 1.73E-05 | 0.0723 | CAGGTCGCCAGCC | *dapB* | 4-hydroxy-tetrahydrodipicolinate reductase |
| AVAEIV_RS22490 | 4810822 | 4810834 | 192 | - | 12.9 | 1.73E-05 | 0.0723 | CCGGGCGCCAGCA | - | ParA family protein |

| | | | | | | | | | | |
|---|---|---|---|---|---|---|---|---|---|---|
| AVAEIV_RS23350 | 4977723 | 4977735 | -50 | + | 12.9 | 1.73E-05 | 0.0723 | CCGACCGCCAGCG | - | S1C family serine protease |
| AVAEIV_RS24410 | 5187934 | 5187946 | 140 | + | 12.9 | 1.73E-05 | 0.0635 | GAGATCGCCAGCG | - | hemolysin family protein |
| AVAEIV_RS03745 | 754181 | 754193 | 206 | - | 12.8857 | 1.75E-05 | 0.0724 | CAATCGGCCATCA | - | tRNA-Leu |
| AVAEIV_RS03750 | 754181 | 754193 | -11 | - | 12.8857 | 1.75E-05 | 0.0638 | CAATCGGCCATCA | *rnr* | ribonuclease R |
| AVAEIV_RS18325 | 3945246 | 3945258 | -3 | + | 12.8857 | 1.75E-05 | 0.0724 | CATATCGCCATCC | - | tyrosine-type recombinase/integrase |
| AVAEIV_RS18330 | 3945246 | 3945258 | 338 | + | 12.8857 | 1.75E-05 | 0.0638 | CATATCGCCATCC | - | antitoxin |
| AVAEIV_RS06935 | 1426712 | 1426724 | 319 | - | 12.8714 | 1.76E-05 | 0.0638 | CAAGGCGCCAGCA | *murC* | UDP-N-acetylmuramate--L-alanine ligase |
| AVAEIV_RS07525 | 1568124 | 1568136 | 80 | + | 12.8714 | 1.76E-05 | 0.0638 | CAGCCCGCCAGCG | - | L,D-transpeptidase |
| AVAEIV_RS15830 | 3453222 | 3453234 | 10 | - | 12.8714 | 1.76E-05 | 0.0724 | CAAGGCGCCAGCA | - | beta |
| AVAEIV_RS20435 | 4345127 | 4345139 | 212 | + | 12.8714 | 1.76E-05 | 0.0724 | CACAGCGCCGGCA | - | Lon protease family protein |
| AVAEIV_RS25805 | 5530736 | 5530748 | -41 | - | 12.8714 | 1.76E-05 | 0.0724 | CGACTCGCCAGCG | - | TetR/AcrR family transcriptional regulator |
| AVAEIV_RS25810 | 5530736 | 5530748 | 146 | - | 12.8714 | 1.76E-05 | 0.0638 | CGACTCGCCAGCG | - | glutathione S-transferase family protein |
| AVAEIV_RS01780 | 352143 | 352155 | 146 | - | 12.8571 | 1.78E-05 | 0.0724 | CAGCTTGCCGGCA | - | sensor histidine kinase |
| AVAEIV_RS01785 | 352143 | 352155 | 31 | - | 12.8571 | 1.78E-05 | 0.0638 | CAGCTTGCCGGCA | - | LysR family transcriptional regulator |
| AVAEIV_RS01840 | 364829 | 364841 | 327 | + | 12.8571 | 1.78E-05 | 0.0638 | CTCAACGCCATCG | *fdxH* | formate dehydrogenase subunit beta |
| AVAEIV_RS05945 | 1214414 | 1214426 | 380 | + | 12.8571 | 1.78E-05 | 0.0724 | CAGCTTGCCGGCA | - | protein |
| AVAEIV_RS05950 | 1214414 | 1214426 | 66 | + | 12.8571 | 1.78E-05 | 0.0724 | CAGCTTGCCGGCA | - | protein |
| AVAEIV_RS05960 | 1215679 | 1215691 | 72 | + | 12.8571 | 1.78E-05 | 0.0724 | CAGCTTGCCGGCA | - | hypothetical protein |
| AVAEIV_RS06640 | 1372263 | 1372275 | 102 | - | 12.8571 | 1.78E-05 | 0.0638 | CGGCGGGCCAGCA | *tldD* | metalloprotease TldD |
| AVAEIV_RS10755 | 2255153 | 2255165 | 252 | + | 12.8571 | 1.78E-05 | 0.0724 | CGGCGGGCCAGCA | *cas2* | CRISPR-associated endonuclease Cas2 |
| AVAEIV_RS17930 | 3878856 | 3878868 | 72 | - | 12.8571 | 1.78E-05 | 0.0638 | CAGCTTGCCGGCA | - | hypothetical protein |
| AVAEIV_RS20030 | 4270874 | 4270886 | 299 | + | 12.8571 | 1.78E-05 | 0.0638 | CTCAACGCCATCG | - | putative quinol monooxygenase |
| AVAEIV_RS21425 | 4559873 | 4559885 | 60 | + | 12.8571 | 1.78E-05 | 0.0724 | CAGCTTGCCGGCA | *infB* | translation initiation factor IF-2 |
| AVAEIV_RS02490 | 496056 | 496068 | 218 | - | 12.8429 | 1.81E-05 | 0.064 | CCGAAGGCCACCG | *argA* | amino-acid N-acetyltransferase |
| AVAEIV_RS05060 | 1041053 | 1041065 | 298 | - | 12.8429 | 1.81E-05 | 0.0724 | CAGAAGGCCGTCG | - | alginate export family protein |
| AVAEIV_RS11090 | 2336716 | 2336728 | 50 | - | 12.8429 | 1.81E-05 | 0.064 | CAGTTTGCCGTCG | - | containing protein |
| AVAEIV_RS15175 | 3303204 | 3303216 | 373 | + | 12.8429 | 1.81E-05 | 0.064 | CTGATCGCCGGCA | - | ABC transporter ATP-binding protein |
| AVAEIV_RS20715 | 4404421 | 4404433 | 359 | - | 12.8429 | 1.81E-05 | 0.0724 | CTGATCGCCGGCA | - | molybdopterin-synthase adenylyltransferase MoeB |
| AVAEIV_RS08430 | 1762481 | 1762493 | 293 | - | 12.8286 | 1.82E-05 | 0.064 | CAGATGGCCTTCA | - | transposase |
| AVAEIV_RS08905 | 1851134 | 1851146 | 267 | + | 12.8286 | 1.82E-05 | 0.0724 | CCGCAGGCCATCG | - | amylo-alpha-1,6-glucosidase |
| AVAEIV_RS09200 | 1920073 | 1920085 | 254 | - | 12.8286 | 1.82E-05 | 0.064 | CCGCAGGCCATCG | - | DUF2334 domain-containing protein |
| AVAEIV_RS14355 | 3126229 | 3126241 | 284 | - | 12.8286 | 1.82E-05 | 0.0724 | CACATGGCCGCCA | - | type II toxin-antitoxin system RelE/ParE family toxin |
| AVAEIV_RS20245 | 4320653 | 4320665 | 65 | + | 12.8286 | 1.82E-05 | 0.0724 | CAGATGGCCTTCA | *secD* | protein translocase subunit SecD |
| AVAEIV_RS21390 | 4547852 | 4547864 | -47 | + | 12.8286 | 1.82E-05 | 0.064 | CATTGCGCCAGCA | *acs* | acetate--CoA ligase |
| AVAEIV_RS01515 | 292395 | 292407 | 184 | - | 12.8143 | 1.84E-05 | 0.0724 | CCGGACGCCATCG | - | dihydroorotase |
| AVAEIV_RS09030 | 1879298 | 1879310 | 259 | - | 12.8143 | 1.84E-05 | 0.0724 | CAGCAGGCCACCG | - | MBL fold metallo-hydrolase |
| AVAEIV_RS15645 | 3426130 | 3426142 | 268 | + | 12.8143 | 1.84E-05 | 0.0724 | CAGCAGGCCACCG | - | heme ABC transporter permease |
| AVAEIV_RS21505 | 4577402 | 4577414 | 22 | + | 12.8143 | 1.84E-05 | 0.0724 | CAGCAGGCCACCG | *dnaK* | molecular chaperone DnaK |
| AVAEIV_RS03445 | 695951 | 695963 | 345 | - | 12.8 | 1.85E-05 | 0.0724 | CGCCGCGCCAGCA | *accB* | acetyl-CoA carboxylase biotin carboxyl carrier protein |
| AVAEIV_RS05430 | 1123338 | 1123350 | 244 | - | 12.8 | 1.85E-05 | 0.064 | CGCTCCGCCATCA | *ribF* | bifunctional riboflavin kinase/FAD synthetase |
| AVAEIV_RS09100 | 1897118 | 1897130 | 88 | - | 12.8 | 1.85E-05 | 0.0724 | CAACAGGCCATCG | - | FecR family protein |
| AVAEIV_RS10855 | 2279233 | 2279245 | -9 | + | 12.8 | 1.85E-05 | 0.064 | CAGAATGCCAGCC | - | rhodanese-like domain-containing protein |
| AVAEIV_RS13160 | 2856192 | 2856204 | 253 | - | 12.8 | 1.85E-05 | 0.0724 | CGCCGCGCCAGCA | - | LacI family DNA-binding transcriptional regulator |
| AVAEIV_RS18600 | 4005399 | 4005411 | 171 | + | 12.8 | 1.85E-05 | 0.064 | CTGTTCGCCACCG | - | MFS transporter |
| AVAEIV_RS18970 | 4084469 | 4084481 | 368 | - | 12.8 | 1.85E-05 | 0.0724 | CAGGACGCCACCG | *kdsA* | 3-deoxy-8-phosphooctulonate synthase |
| AVAEIV_RS21140 | 4492825 | 4492837 | 357 | + | 12.8 | 1.85E-05 | 0.0724 | CAACAGGCCATCG | - | IS630 family transposase |
| AVAEIV_RS21145 | 4492825 | 4492837 | -31 | + | 12.8 | 1.85E-05 | 0.064 | CAACAGGCCATCG | - | hypothetical protein |
| AVAEIV_RS23550 | 5019013 | 5019025 | 63 | + | 12.8 | 1.85E-05 | 0.0724 | CGCCGCGCCAGCA | *murU* | uridylyltransferase MurU |
| AVAEIV_RS25165 | 5359804 | 5359816 | 341 | + | 12.8 | 1.85E-05 | 0.0724 | CGCCGCGCCAGCA | *hypD* | hydrogenase formation protein HypD |
| AVAEIV_RS25170 | 5359804 | 5359816 | 90 | + | 12.8 | 1.85E-05 | 0.0724 | CGCCGCGCCAGCA | - | chaperone |
| AVAEIV_RS15945 | 3476386 | 3476398 | 323 | + | 12.7857 | 1.88E-05 | 0.064 | CACTTGGCCGGCA | - | LysR family transcriptional regulator |
| AVAEIV_RS17510 | 3790467 | 3790479 | 245 | + | 12.7857 | 1.88E-05 | 0.064 | CTATTCGCCATCG | *ppk2* | polyphosphate kinase 2 |

| | | | | | | | | | | |
|---|---|---|---|---|---|---|---|---|---|---|
| AVAEIV_RS18605 | 4007462 | 4007474 | 347 | - | 12.7857 | 1.88E-05 | 0.0724 | CGCTGCGCCATCG | - | carboxymuconolactone decarboxylase family protein |
| AVAEIV_RS24615 | 5230017 | 5230029 | 223 | - | 12.7857 | 1.88E-05 | 0.0724 | CAGATGGCCAACA | *anfO* | Fe-only nitrogenase accessory protein AnfO |
| AVAEIV_RS04105 | 833496 | 833508 | 155 | - | 12.7714 | 1.90E-05 | 0.064 | CGGCAGGCCATCG | - | septal ring lytic transglycosylase RlpA family protein |
| AVAEIV_RS06830 | 1406816 | 1406828 | 343 | - | 12.7714 | 1.90E-05 | 0.064 | CACATAGCCAGCA | *petA* | ubiquinol-cytochrome c reductase iron-sulfur subunit |
| AVAEIV_RS07725 | 1609435 | 1609447 | 168 | - | 12.7714 | 1.90E-05 | 0.064 | CGGATCGCCAGTA | *pabC* | aminodeoxychorismate lyase |
| AVAEIV_RS08980 | 1868331 | 1868343 | -40 | + | 12.7714 | 1.90E-05 | 0.064 | CGGCAGGCCATCG | - | hypothetical protein |
| AVAEIV_RS11860 | 2515671 | 2515683 | 35 | - | 12.7714 | 1.90E-05 | 0.0724 | CCCCACGCCATCG | - | nitrate reductase |
| AVAEIV_RS23350 | 4978136 | 4978148 | 363 | + | 12.7714 | 1.90E-05 | 0.0724 | CTCATTGCCATCG | - | S1C family serine protease |
| AVAEIV_RS23355 | 4978136 | 4978148 | 36 | + | 12.7714 | 1.90E-05 | 0.064 | CTCATTGCCATCG | - | hypothetical protein |
| AVAEIV_RS02870 | 591212 | 591224 | 306 | - | 12.7571 | 1.92E-05 | 0.0724 | CCGCTCGCCGGCA | - | diguanylate cyclase |
| AVAEIV_RS10715 | 2241756 | 2241768 | 19 | + | 12.7571 | 1.92E-05 | 0.0724 | CCCAGGGCCATCG | - | tetratricopeptide repeat protein |
| AVAEIV_RS10800 | 2265861 | 2265873 | 258 | - | 12.7571 | 1.92E-05 | 0.0724 | CGGGACGCCATCG | - | hypothetical protein |
| AVAEIV_RS15775 | 3444924 | 3444936 | 166 | - | 12.7571 | 1.92E-05 | 0.064 | CATCTCGCCAGCG | - | lysozyme inhibitor LprI family protein |
| AVAEIV_RS21155 | 4497885 | 4497897 | 97 | - | 12.7571 | 1.92E-05 | 0.0724 | CTGAGCGCCAGCG | - | MFS transporter |
| AVAEIV_RS22585 | 4837313 | 4837325 | 122 | + | 12.7571 | 1.92E-05 | 0.064 | CTGAGCGCCAGCG | *argS* | arginine--tRNA ligase |
| AVAEIV_RS24040 | 5116235 | 5116247 | 352 | + | 12.7571 | 1.92E-05 | 0.064 | CCGATCGGCAGCA | - | HAD family hydrolase |
| AVAEIV_RS03730 | 750883 | 750895 | 132 | - | 12.7429 | 1.94E-05 | 0.064 | CCCTACGCCAGCG | - | ATP phosphoribosyltransferase regulatory subunit |
| AVAEIV_RS06375 | 1299519 | 1299531 | 371 | + | 12.7429 | 1.94E-05 | 0.064 | CCGTTCGCCGTCG | *ampD* | AmpD |
| AVAEIV_RS15160 | 3300400 | 3300412 | 239 | - | 12.7429 | 1.94E-05 | 0.064 | CCGTTCGCCGTCG | - | VOC family protein |
| AVAEIV_RS21215 | 4510125 | 4510137 | 324 | + | 12.7429 | 1.94E-05 | 0.0724 | CCGTTCGCCGTCG | - | 2-isopropylmalate synthase |
| AVAEIV_RS00815 | 158328 | 158340 | 135 | + | 12.7286 | 1.95E-05 | 0.064 | CAGCTCGGCAGCA | *nifM* | nitrogen fixation protein NifM |
| AVAEIV_RS03660 | 737976 | 737988 | 287 | + | 12.7286 | 1.95E-05 | 0.0724 | CAGTTCGCCGCCG | - | rhodanese-like domain-containing protein |
| AVAEIV_RS06350 | 1294378 | 1294390 | 292 | - | 12.7286 | 1.95E-05 | 0.0724 | CGCAACGCCACCG | - | prepilin peptidase |
| AVAEIV_RS12310 | 2605309 | 2605321 | 158 | - | 12.7286 | 1.95E-05 | 0.064 | CAGTTCGCCGCCG | *fliT* | flagellar protein FliT |
| AVAEIV_RS12430 | 2634005 | 2634017 | 272 | - | 12.7286 | 1.95E-05 | 0.064 | CAGTTCGCCGCCG | - | response regulator |
| AVAEIV_RS15380 | 3359444 | 3359456 | 158 | + | 12.7286 | 1.95E-05 | 0.064 | CAACTCGCCGGCA | - | pyrimidine/purine nucleoside phosphorylase |
| AVAEIV_RS17405 | 3769280 | 3769292 | 260 | - | 12.7286 | 1.95E-05 | 0.064 | CACCGGGCCATCG | *phhA* | phenylalanine 4-monooxygenase |
| AVAEIV_RS19825 | 4225086 | 4225098 | 357 | + | 12.7286 | 1.95E-05 | 0.0724 | CAGTTCGCCGCCG | *rplS* | 50S ribosomal protein L19 |
| AVAEIV_RS22080 | 4712729 | 4712741 | 385 | + | 12.7286 | 1.95E-05 | 0.0724 | CAGTTCGCCGCCG | - | NAD-dependent epimerase/dehydratase family protein |
| AVAEIV_RS22605 | 4841843 | 4841855 | 374 | + | 12.7286 | 1.95E-05 | 0.064 | CAGTTCGCCGCCG | - | containing protein |
| AVAEIV_RS23620 | 5035304 | 5035316 | 65 | - | 12.7286 | 1.95E-05 | 0.064 | CAACTCGCCGGCA | - | SulP family inorganic anion transporter |
| AVAEIV_RS24705 | 5248966 | 5248978 | 269 | - | 12.7286 | 1.95E-05 | 0.064 | CAGTTCGCCGCCG | - | hypothetical protein |
| AVAEIV_RS01610 | 312470 | 312482 | -10 | - | 12.7143 | 1.97E-05 | 0.0724 | CTGATGGCCACCG | - | type II toxin-antitoxin system HipA family toxin |
| AVAEIV_RS07125 | 1468059 | 1468071 | 251 | - | 12.7143 | 1.97E-05 | 0.064 | CAGTTCGGCATCG | - | MucB/RseB C-terminal domain-containing protein |
| AVAEIV_RS07210 | 1486650 | 1486662 | 229 | - | 12.7143 | 1.97E-05 | 0.0724 | CAGTTCGGCATCG | - | iron-containing alcohol dehydrogenase |
| AVAEIV_RS07960 | 1654793 | 1654805 | 298 | + | 12.7143 | 1.97E-05 | 0.064 | CAGTTCGGCATCG | - | ATP-binding protein |
| AVAEIV_RS08645 | 1802681 | 1802693 | 311 | - | 12.7143 | 1.97E-05 | 0.064 | GCGATCGCCAGCA | - | hypothetical protein |
| AVAEIV_RS10175 | 2124008 | 2124020 | 232 | - | 12.7143 | 1.97E-05 | 0.0724 | CCGTTGGCCATCC | - | cytochrome ubiquinol oxidase subunit I |
| AVAEIV_RS10180 | 2124008 | 2124020 | 20 | - | 12.7143 | 1.97E-05 | 0.0724 | CCGTTGGCCATCC | *cydP* | cytochrome oxidase putative small subunit CydP |
| AVAEIV_RS12955 | 2808908 | 2808920 | 366 | + | 12.7143 | 1.97E-05 | 0.0724 | CAGTTCGGCATCG | *groL* | chaperonin GroEL |
| AVAEIV_RS17755 | 3843254 | 3843266 | -45 | + | 12.7143 | 1.97E-05 | 0.0724 | CAGTTCGGCATCG | - | sulfurtransferase TusA family protein |
| AVAEIV_RS17760 | 3843254 | 3843266 | 170 | + | 12.7143 | 1.97E-05 | 0.064 | CAGTTCGGCATCG | - | M48 family metalloprotease |
| AVAEIV_RS18790 | 4044894 | 4044906 | 360 | - | 12.7143 | 1.97E-05 | 0.064 | CAGTTCGGCATCG | *pyk* | pyruvate kinase |
| AVAEIV_RS20480 | 4355386 | 4355398 | 370 | - | 12.7143 | 1.97E-05 | 0.0724 | CAGTTCGGCATCG | *ettA* | energy-dependent translational throttle protein EttA |
| AVAEIV_RS20485 | 4355386 | 4355398 | -45 | - | 12.7143 | 1.97E-05 | 0.064 | CAGTTCGGCATCG | *glyA* | serine hydroxymethyltransferase |
| AVAEIV_RS21640 | 4606293 | 4606305 | 177 | - | 12.7143 | 1.97E-05 | 0.064 | CAGTTCGGCATCG | *glcF* | glycolate oxidase subunit GlcF |
| AVAEIV_RS22810 | 4883191 | 4883203 | -25 | + | 12.7143 | 1.97E-05 | 0.064 | CAGTTCGGCATCG | - | carbohydrate porin |
| AVAEIV_RS24295 | 5166638 | 5166650 | 284 | - | 12.7143 | 1.97E-05 | 0.0724 | CACGGCGCCATCG | *ilvA* | threonine ammonia-lyase, biosynthetic |
| AVAEIV_RS24430 | 5194274 | 5194286 | 288 | - | 12.7143 | 1.97E-05 | 0.0724 | CTGATGGCCACCG | *pstA* | phosphate ABC transporter permease PstA |
| AVAEIV_RS25190 | 5364577 | 5364589 | 263 | + | 12.7143 | 1.97E-05 | 0.0724 | CAGTTCGGCATCG | - | nickel-dependent hydrogenase large subunit |
| AVAEIV_RS25230 | 5369953 | 5369965 | 112 | - | 12.7143 | 1.97E-05 | 0.0724 | CAGTTCGGCATCG | - | nickel-dependent hydrogenase large subunit |
| AVAEIV_RS01530 | 294693 | 294705 | 214 | - | 12.7 | 2.00E-05 | 0.0724 | CTGCTGGCCATCG | *ruvX* | Holliday junction resolvase RuvX |

| | | | | | | | | | | |
|---|---|---|---|---|---|---|---|---|---|---|
| AVAEIV_RS03610 | 727485 | 727497 | 213 | - | 12.7 | 2.00E-05 | 0.0724 | CCGAACGCCAGCC | *msrQ* | subunit MsrQ |
| AVAEIV_RS15775 | 3445130 | 3445142 | -40 | + | 12.7 | 2.00E-05 | 0.064 | CTGCTGGCCATCG | - | lysozyme inhibitor LprI family protein |
| AVAEIV_RS16680 | 3613010 | 3613022 | 359 | + | 12.7 | 2.00E-05 | 0.0724 | CGGCTCGCCGGCA | *cobC* | alpha-ribazole phosphatase family protein |
| AVAEIV_RS18405 | 3965135 | 3965147 | 196 | + | 12.7 | 2.00E-05 | 0.0724 | CTGCAGGCCATCA | *cysM* | cysteine synthase CysM |
| AVAEIV_RS18410 | 3965135 | 3965147 | 145 | + | 12.7 | 2.00E-05 | 0.064 | CTGCAGGCCATCA | - | response regulator |
| AVAEIV_RS22270 | 4754555 | 4754567 | 348 | - | 12.7 | 2.00E-05 | 0.0724 | CTGCTGGCCATCG | - | acyltransferase family protein |
| AVAEIV_RS22390 | 4787538 | 4787550 | 219 | + | 12.7 | 2.00E-05 | 0.064 | CTGCTGGCCATCG | - | universal stress protein |
| AVAEIV_RS22950 | 4916437 | 4916449 | 158 | + | 12.7 | 2.00E-05 | 0.064 | CTGCTGGCCATCG | - | histidine triad nucleotide-binding protein |
| AVAEIV_RS24270 | 5160640 | 5160652 | 177 | - | 12.7 | 2.00E-05 | 0.0724 | CTAATGGCCATCG | - | sensor domain-containing diguanylate cyclase |
| AVAEIV_RS24625 | 5231881 | 5231893 | 123 | + | 12.7 | 2.00E-05 | 0.0724 | CAAGTTGCCATCG | *anfG* | Fe-only nitrogenase subunit delta |
| AVAEIV_RS25230 | 5369816 | 5369828 | -25 | - | 12.7 | 2.00E-05 | 0.0724 | CCGAACGCCAGCC | - | nickel-dependent hydrogenase large subunit |
| AVAEIV_RS25570 | 5466211 | 5466223 | 371 | - | 12.7 | 2.00E-05 | 0.0724 | CGCAGGGCCATCG | - | 2-dehydro-3-deoxy-6-phosphogalactonate aldolase |
| AVAEIV_RS06925 | 1424444 | 1424456 | 272 | - | 12.6857 | 2.02E-05 | 0.064 | CCCCTTGCCATCG | *ftsW* | putative lipid II flippase FtsW |
| AVAEIV_RS08240 | 1715677 | 1715689 | 261 | - | 12.6857 | 2.02E-05 | 0.064 | CCAACCGCCAGCA | - | NAD(P)/FAD-dependent oxidoreductase |
| AVAEIV_RS09715 | 2034333 | 2034345 | 44 | - | 12.6857 | 2.02E-05 | 0.0724 | CTGGTCGCCATCG | - | DUF883 family protein |
| AVAEIV_RS09720 | 2034333 | 2034345 | 91 | - | 12.6857 | 2.02E-05 | 0.064 | CTGGTCGCCATCG | - | glutaredoxin family protein |
| AVAEIV_RS12010 | 2552636 | 2552648 | 145 | - | 12.6857 | 2.02E-05 | 0.0724 | GAAATCGCCAGCA | - | acetyl-CoA C-acetyltransferase |
| AVAEIV_RS12700 | 2697075 | 2697087 | 86 | + | 12.6857 | 2.02E-05 | 0.0724 | CTGGACGCCATCA | - | DUF485 domain-containing protein |
| AVAEIV_RS16245 | 3541937 | 3541949 | 236 | + | 12.6857 | 2.02E-05 | 0.0724 | CGGTTCGCCGTCG | - | DUF6566 family protein |
| AVAEIV_RS08060 | 1671547 | 1671559 | 135 | + | 12.6714 | 2.04E-05 | 0.064 | CTGTTGGCCAGCG | - | L-threonylcarbamoyladenylate synthase |
| AVAEIV_RS17195 | 3728776 | 3728788 | 77 | + | 12.6714 | 2.04E-05 | 0.0724 | CAGCACGCCAGCC | - | FimV/HubP family polar landmark protein |
| AVAEIV_RS17950 | 3884116 | 3884128 | 135 | + | 12.6714 | 2.04E-05 | 0.0724 | CAGGAGGCCAGCG | - | type II toxin-antitoxin system RelE/ParE family toxin |
| AVAEIV_RS20070 | 4284717 | 4284729 | 346 | - | 12.6714 | 2.04E-05 | 0.0724 | CATCTTGCCAGCA | - | OprD family porin |
| AVAEIV_RS20080 | 4288718 | 4288730 | 332 | + | 12.6714 | 2.04E-05 | 0.0724 | GAGTTCGCCATCG | - | TonB-dependent copper receptor |
| AVAEIV_RS21365 | 4543020 | 4543032 | 124 | + | 12.6714 | 2.04E-05 | 0.064 | CAGGAGGCCAGCG | *folK* | diphosphokinase |
| AVAEIV_RS22490 | 4810945 | 4810957 | 315 | - | 12.6714 | 2.04E-05 | 0.0724 | CAGGAGGCCAGCG | - | ParA family protein |
| AVAEIV_RS01265 | 247645 | 247657 | -34 | - | 12.6571 | 2.06E-05 | 0.0726 | AAGATCGCCATCA | - | protein |
| AVAEIV_RS01525 | 294135 | 294147 | 128 | - | 12.6571 | 2.06E-05 | 0.0726 | CCCAAGGCCAGCG | *pyrR* | bifunctional pyr operon transcriptional regulator/uracil phosphoribosyltransferase PyrR |
| AVAEIV_RS02840 | 583151 | 583163 | 155 | - | 12.6571 | 2.06E-05 | 0.064 | CAGATGGCTATCG | - | NfeD family protein |
| AVAEIV_RS04675 | 954810 | 954822 | 308 | - | 12.6571 | 2.06E-05 | 0.0726 | CCGAAGGCCGTCA | - | TetR family transcriptional regulator |
| AVAEIV_RS04680 | 954810 | 954822 | -47 | - | 12.6571 | 2.06E-05 | 0.064 | CCGAAGGCCGTCA | - | multidrug effflux MFS transporter |
| AVAEIV_RS07895 | 1641966 | 1641978 | 174 | - | 12.6571 | 2.06E-05 | 0.0726 | CAGTCCGCCACCG | - | LysR family transcriptional regulator |
| AVAEIV_RS19655 | 4190596 | 4190608 | 85 | + | 12.6571 | 2.06E-05 | 0.064 | CTCAACGCCACCA | - | putative RNA methyltransferase |
| AVAEIV_RS20695 | 4399238 | 4399250 | -31 | + | 12.6571 | 2.06E-05 | 0.064 | AAGATCGCCATCA | - | YbgA family protein |
| AVAEIV_RS23555 | 5020252 | 5020264 | 283 | + | 12.6571 | 2.06E-05 | 0.0726 | CCCAAGGCCAGCG | - | aminoglycoside phosphotransferase family protein |
| AVAEIV_RS25655 | 5485232 | 5485244 | 266 | + | 12.6571 | 2.06E-05 | 0.064 | CCCTTCGCCATCC | - | carbohydrate porin |
| AVAEIV_RS00430 | 87336 | 87348 | 230 | + | 12.6429 | 2.09E-05 | 0.064 | CAGATGGCCGCCG | - | hypothetical protein |
| AVAEIV_RS03775 | 759088 | 759100 | 187 | - | 12.6429 | 2.09E-05 | 0.064 | CCCTGCGCCACCA | *rplI* | 50S ribosomal protein L9 |
| AVAEIV_RS08185 | 1701179 | 1701191 | 81 | - | 12.6429 | 2.09E-05 | 0.064 | CGGAACGCCAGCC | - | containing protein |
| AVAEIV_RS11645 | 2463596 | 2463608 | -33 | - | 12.6429 | 2.09E-05 | 0.0726 | CAATCCGCCATCG | - | MliC family protein |
| AVAEIV_RS13880 | 3007992 | 3008004 | 129 | + | 12.6429 | 2.09E-05 | 0.064 | CTCCTCGCCATCG | *tagH* | protein TagH |
| AVAEIV_RS20870 | 4438665 | 4438677 | 285 | - | 12.6429 | 2.09E-05 | 0.0726 | CACTTCGCCACCC | *pqqD* | chaperone PqqD |
| AVAEIV_RS21020 | 4469405 | 4469417 | 299 | - | 12.6429 | 2.09E-05 | 0.064 | CAGATGGCCGCCG | - | type II toxin-antitoxin system RelE/ParE family toxin |
| AVAEIV_RS23610 | 5032740 | 5032752 | 297 | + | 12.6429 | 2.09E-05 | 0.064 | CCCTGCGCCACCA | - | multifunctional CCA addition/repair protein |
| AVAEIV_RS04915 | 1003970 | 1003982 | 295 | - | 12.6286 | 2.11E-05 | 0.0726 | CCGCTGGCCACCG | - | energy transducer TonB |
| AVAEIV_RS07295 | 1511834 | 1511846 | 149 | - | 12.6286 | 2.11E-05 | 0.0726 | CGCCATGCCATCA | *pyrC* | dihydroorotase |
| AVAEIV_RS07300 | 1511834 | 1511846 | -6 | - | 12.6286 | 2.11E-05 | 0.064 | CGCCATGCCATCA | - | argininosuccinate synthase |
| AVAEIV_RS07950 | 1651938 | 1651950 | 167 | - | 12.6286 | 2.11E-05 | 0.064 | CAGATGGGCATCG | - | glutathione peroxidase |
| AVAEIV_RS08320 | 1731961 | 1731973 | 260 | - | 12.6286 | 2.11E-05 | 0.064 | CACCAGGCCAGCG | - | DUF1972 domain-containing protein |
| AVAEIV_RS08350 | 1744614 | 1744626 | 13 | + | 12.6286 | 2.11E-05 | 0.064 | CGGCCCGCCAGCA | - | family antitoxin |
| AVAEIV_RS08355 | 1744614 | 1744626 | 225 | + | 12.6286 | 2.11E-05 | 0.064 | CGGCCCGCCAGCA | - | Txe/YoeB family addiction module toxin |

| | | | | | | | | | | |
|---|---|---|---|---|---|---|---|---|---|---|
| AVAEIV_RS13280 | 2876475 | 2876487 | 342 | + | 12.6286 | 2.11E-05 | 0.064 | CAGCAGGCCGTCA | - | recombinase family protein |
| AVAEIV_RS18875 | 4064814 | 4064826 | 363 | + | 12.6286 | 2.11E-05 | 0.0726 | CCGTTCGCCATGA | - | alpha/beta fold hydrolase |
| AVAEIV_RS22590 | 4839087 | 4839099 | 109 | + | 12.6286 | 2.11E-05 | 0.064 | CGGCCCGCCAGCA | - | SPOR domain-containing protein |
| AVAEIV_RS24890 | 5296486 | 5296498 | 384 | + | 12.6286 | 2.11E-05 | 0.0726 | CCAATGGCCACCG | - | OprD family porin |
| AVAEIV_RS25480 | 5427232 | 5427244 | 307 | + | 12.6286 | 2.11E-05 | 0.0726 | CACCAGGCCAGCG | *norR* | nitric oxide reductase transcriptional regulator NorR |
| AVAEIV_RS25680 | 5493621 | 5493633 | 169 | + | 12.6286 | 2.11E-05 | 0.064 | CCGCTGGCCACCG | - | efflux transporter outer membrane subunit |
| AVAEIV_RS04010 | 811339 | 811351 | 146 | - | 12.6143 | 2.13E-05 | 0.064 | CCACTGGCCATCG | - | aryl-sulfate sulfotransferase |
| AVAEIV_RS04275 | 871158 | 871170 | 77 | + | 12.6143 | 2.13E-05 | 0.0726 | CAGGTCGCCGTCG | - | phenol hydroxylase subunit P4 |
| AVAEIV_RS04490 | 918162 | 918174 | 375 | + | 12.6143 | 2.13E-05 | 0.0726 | CGAAGCGCCAGCG | - | siderophore biosynthesis protein PvsA |
| AVAEIV_RS04675 | 954824 | 954836 | 322 | + | 12.6143 | 2.13E-05 | 0.0726 | CCACTGGCCATCG | - | TetR family transcriptional regulator |
| AVAEIV_RS09185 | 1914865 | 1914877 | 311 | - | 12.6143 | 2.13E-05 | 0.064 | CAGGTCGCCGTCG | - | proton-conducting transporter membrane subunit |
| AVAEIV_RS14380 | 3130452 | 3130464 | 362 | + | 12.6143 | 2.13E-05 | 0.0726 | CGAAGCGCCAGCG | - | RNA polymerase sigma factor FliA |
| AVAEIV_RS15600 | 3419151 | 3419163 | 366 | + | 12.6143 | 2.13E-05 | 0.0726 | CTCTTCGCCAGCG | *moaA* | GTP 3',8-cyclase MoaA |
| AVAEIV_RS18235 | 3935461 | 3935473 | 376 | - | 12.6143 | 2.13E-05 | 0.0726 | CACGACGCCAGCG | - | hypothetical protein |
| AVAEIV_RS20365 | 4332314 | 4332326 | 373 | - | 12.6143 | 2.13E-05 | 0.064 | CGAAGCGCCAGCG | - | hypothetical protein |
| AVAEIV_RS22040 | 4703964 | 4703976 | 241 | + | 12.6143 | 2.13E-05 | 0.0726 | CGGCGCGCCAGCG | - | ion channel |
| AVAEIV_RS24320 | 5170297 | 5170309 | 325 | - | 12.6143 | 2.13E-05 | 0.064 | CCACTGGCCATCG | *serA* | phosphoglycerate dehydrogenase |
| AVAEIV_RS02830 | 580352 | 580364 | 164 | + | 12.6 | 2.15E-05 | 0.0726 | CAGTTGGCCGGCG | - | cytochrome B6 |
| AVAEIV_RS05655 | 1163243 | 1163255 | 159 | - | 12.6 | 2.15E-05 | 0.0726 | CGCAAGGCCAGCG | - | protein |
| AVAEIV_RS05800 | 1188010 | 1188022 | 362 | - | 12.6 | 2.15E-05 | 0.064 | CCCATCGCCGTCG | - | acyl-CoA dehydrogenase family protein |
| AVAEIV_RS06560 | 1353171 | 1353183 | 359 | - | 12.6 | 2.15E-05 | 0.064 | CAGTTGGCCGGCG | - | ATP-binding protein |
| AVAEIV_RS06905 | 1419188 | 1419200 | 265 | + | 12.6 | 2.15E-05 | 0.064 | CGCAAGGCCAGCG | - | diaminopimelate ligase |
| AVAEIV_RS14350 | 3125595 | 3125607 | -28 | - | 12.6 | 2.15E-05 | 0.0726 | CGCTTTGCCAGCG | - | helix-turn-helix domain-containing protein |
| AVAEIV_RS18970 | 4084158 | 4084170 | 57 | + | 12.6 | 2.15E-05 | 0.0726 | CGGATGGCCGTCG | *kdsA* | 3-deoxy-8-phosphooctulonate synthase |
| AVAEIV_RS21725 | 4629596 | 4629608 | 190 | + | 12.6 | 2.15E-05 | 0.0726 | CAGTTGGCCGGCG | - | ABC transporter substrate-binding protein |
| AVAEIV_RS25855 | 5544121 | 5544133 | 289 | + | 12.6 | 2.15E-05 | 0.064 | CACAGCGCCAGCC | - | ABC transporter substrate-binding protein |
| AVAEIV_RS25865 | 5546678 | 5546690 | 196 | + | 12.6 | 2.15E-05 | 0.064 | CAGTTGGCCGGCG | - | ABC transporter permease |
| AVAEIV_RS02865 | 588215 | 588227 | 55 | - | 12.5857 | 2.18E-05 | 0.064 | CAGAAAGCCAGCA | - | containing protein |
| AVAEIV_RS03940 | 794591 | 794603 | 345 | + | 12.5857 | 2.18E-05 | 0.064 | CCGACGGCCATCG | *ubiX* | flavin prenyltransferase UbiX |
| AVAEIV_RS05165 | 1059862 | 1059874 | 252 | - | 12.5857 | 2.18E-05 | 0.064 | TGGATCGCCAGCA | *coxB* | cytochrome c oxidase subunit II |
| AVAEIV_RS12035 | 2557442 | 2557454 | 375 | - | 12.5857 | 2.18E-05 | 0.0726 | CAGATCGCCTTCG | *folE* | GTP cyclohydrolase I FolE |
| AVAEIV_RS19045 | 4103776 | 4103788 | 333 | - | 12.5857 | 2.18E-05 | 0.0726 | GAGAAGGCCATCA | *uppS* | polyprenyl diphosphate synthase |
| AVAEIV_RS00395 | 81314 | 81326 | 120 | + | 12.5714 | 2.21E-05 | 0.0726 | CACCTCGCCGTCG | - | hypothetical protein |
| AVAEIV_RS04105 | 833372 | 833384 | 279 | - | 12.5714 | 2.21E-05 | 0.064 | CGGCTGGCCACCG | - | septal ring lytic transglycosylase RlpA family protein |
| AVAEIV_RS04185 | 853303 | 853315 | 160 | - | 12.5714 | 2.21E-05 | 0.064 | CACCTCGCCGTCG | - | dehydrogenase |
| AVAEIV_RS05930 | 1213177 | 1213189 | 147 | + | 12.5714 | 2.21E-05 | 0.0726 | CGGCTGGCCACCG | - | protein |
| AVAEIV_RS09670 | 2025076 | 2025088 | 379 | - | 12.5714 | 2.21E-05 | 0.0726 | CAGACGGCCACCG | - | hypothetical protein |
| AVAEIV_RS15350 | 3356133 | 3356145 | 153 | + | 12.5714 | 2.21E-05 | 0.0726 | CCCCACGCCACCA | *sdhD* | anchor protein |
| AVAEIV_RS18325 | 3945266 | 3945278 | 17 | + | 12.5714 | 2.21E-05 | 0.0726 | CTGATCGCCAGCC | - | tyrosine-type recombinase/integrase |
| AVAEIV_RS18330 | 3945266 | 3945278 | 318 | + | 12.5714 | 2.21E-05 | 0.064 | CTGATCGCCAGCC | - | antitoxin |
| AVAEIV_RS19915 | 4245875 | 4245887 | 158 | - | 12.5714 | 2.21E-05 | 0.0726 | CACATCGGCATCG | *guaB* | IMP dehydrogenase |
| AVAEIV_RS21175 | 4502222 | 4502234 | 22 | + | 12.5714 | 2.21E-05 | 0.0726 | CGAATGGCCACCG | - | helix-turn-helix domain-containing protein |
| AVAEIV_RS22010 | 4696301 | 4696313 | 303 | + | 12.5714 | 2.21E-05 | 0.064 | CACAACGGCATCA | - | integrase core domain-containing protein |
| AVAEIV_RS24205 | 5146690 | 5146702 | 85 | - | 12.5714 | 2.21E-05 | 0.0726 | CTGATCGCCAGCC | *ftsX* | permease-like cell division protein FtsX |
| AVAEIV_RS24875 | 5292881 | 5292893 | -10 | - | 12.5714 | 2.21E-05 | 0.0726 | CCCATGGCCATCC | - | carboxyl transferase domain-containing protein |
| AVAEIV_RS00145 | 30189 | 30201 | 316 | + | 12.5571 | 2.23E-05 | 0.064 | CCCAGGGCCACCA | - | hypothetical protein |
| AVAEIV_RS03810 | 768333 | 768345 | 50 | - | 12.5571 | 2.23E-05 | 0.064 | CAGCCGGCCATCG | - | HAD family hydrolase |
| AVAEIV_RS11365 | 2403947 | 2403959 | 197 | - | 12.5571 | 2.23E-05 | 0.0726 | CAGCCGGCCATCG | *xdhC* | xanthine dehydrogenase accessory protein XdhC |
| AVAEIV_RS14060 | 3057507 | 3057519 | 221 | + | 12.5571 | 2.23E-05 | 0.0726 | CGGGTCGCCACCG | - | dehydrogenase |
| AVAEIV_RS22785 | 4877706 | 4877718 | 153 | - | 12.5571 | 2.23E-05 | 0.064 | CGGGTCGCCACCG | *tpiA* | triose-phosphate isomerase |
| AVAEIV_RS23500 | 5007865 | 5007877 | -19 | - | 12.5571 | 2.23E-05 | 0.0726 | CCTTTCGCCATCG | *rpe* | ribulose-phosphate 3-epimerase |
| AVAEIV_RS24950 | 5309919 | 5309931 | 153 | + | 12.5571 | 2.23E-05 | 0.0726 | CGGGTCGCCACCG | *tpiA* | triose-phosphate isomerase |

| | | | | | | | | | | |
|---|---|---|---|---|---|---|---|---|---|---|
| AVAEIV_RS01140 | 224328 | 224340 | 380 | - | 12.5429 | 2.25E-05 | 0.0726 | CTGCGCGCCAGCA | - | endonuclease I family protein |
| AVAEIV_RS01150 | 224328 | 224340 | 184 | - | 12.5429 | 2.25E-05 | 0.064 | CTGCGCGCCAGCA | - | CsbD family protein |
| AVAEIV_RS01265 | 247851 | 247863 | 172 | - | 12.5429 | 2.25E-05 | 0.0726 | CGCATCGCCGTCG | - | protein |
| AVAEIV_RS01415 | 276463 | 276475 | 161 | - | 12.5429 | 2.25E-05 | 0.0726 | CGCATCGCCGTCG | - | phosphomannomutase/phosphoglucomutase |
| AVAEIV_RS11775 | 2494294 | 2494306 | 225 | - | 12.5429 | 2.25E-05 | 0.0726 | CAGATCGCCAACG | *pabB* | aminodeoxychorismate synthase component I |
| AVAEIV_RS15485 | 3383965 | 3383977 | 334 | + | 12.5429 | 2.25E-05 | 0.064 | CTGCGCGCCAGCA | - | acetyltransferase |
| AVAEIV_RS15985 | 3484415 | 3484427 | 30 | + | 12.5429 | 2.25E-05 | 0.064 | CTGCGCGCCAGCA | - | ABC transporter permease |
| AVAEIV_RS16450 | 3571826 | 3571838 | 84 | + | 12.5429 | 2.25E-05 | 0.064 | CAGATCGCCAACG | - | flavodoxin |
| AVAEIV_RS16995 | 3682946 | 3682958 | 305 | + | 12.5429 | 2.25E-05 | 0.064 | CACTTCGCCGGCG | - | YbfB/YjiJ family MFS transporter |
| AVAEIV_RS18095 | 3910242 | 3910254 | 227 | - | 12.5429 | 2.25E-05 | 0.0726 | CAGGCCGCCATCG | *tolB* | Tol-Pal system beta propeller repeat protein TolB |
| AVAEIV_RS19175 | 4132835 | 4132847 | 71 | + | 12.5429 | 2.25E-05 | 0.0726 | CACCTGGCCATCC | - | hypothetical protein |
| AVAEIV_RS21075 | 4482708 | 4482720 | 307 | - | 12.5429 | 2.25E-05 | 0.0726 | CAGGCCGCCATCG | - | feruloyl-CoA synthase |
| AVAEIV_RS24175 | 5140839 | 5140851 | 281 | - | 12.5429 | 2.25E-05 | 0.0726 | CAGATCGCCAACG | - | MFS transporter |
| AVAEIV_RS24770 | 5264443 | 5264455 | -37 | + | 12.5429 | 2.25E-05 | 0.064 | CACCTGGCCATCC | - | argininosuccinate lyase |
| AVAEIV_RS03550 | 715824 | 715836 | 27 | - | 12.5286 | 2.28E-05 | 0.064 | CCCGGCGCCATCA | - | eCIS core domain-containing protein |
| AVAEIV_RS09535 | 1987881 | 1987893 | 39 | + | 12.5286 | 2.28E-05 | 0.064 | CGGACGGCCATCG | - | TenA family transcriptional regulator |
| AVAEIV_RS15330 | 3350312 | 3350324 | 323 | - | 12.5286 | 2.28E-05 | 0.0726 | GACATCGCCATCG | *sucB* | dihydrolipoyllysine-residue succinyltransferase |
| AVAEIV_RS18545 | 3992647 | 3992659 | 332 | - | 12.5286 | 2.28E-05 | 0.064 | CAGTCGGCCAGCG | - | protein |
| AVAEIV_RS20035 | 4274938 | 4274950 | 242 | - | 12.5286 | 2.28E-05 | 0.0726 | CCGTTCGGCATCA | - | LuxR C-terminal-related transcriptional regulator |
| AVAEIV_RS20040 | 4274938 | 4274950 | -14 | - | 12.5286 | 2.28E-05 | 0.064 | CCGTTCGGCATCA | - | glycerol dehydrogenase |
| AVAEIV_RS21970 | 4686320 | 4686332 | 276 | + | 12.5286 | 2.28E-05 | 0.064 | GACAACGCCATCA | - | acetate/propionate family kinase |
| AVAEIV_RS24585 | 5222460 | 5222472 | 353 | + | 12.5286 | 2.28E-05 | 0.064 | CCCACCGCCATCG | *aguA* | agmatine deiminase |
| AVAEIV_RS00330 | 71702 | 71714 | -28 | + | 12.5143 | 2.31E-05 | 0.064 | CGCCTCGCCACCG | - | phosphatase PAP2 family protein |
| AVAEIV_RS03060 | 625365 | 625377 | 340 | + | 12.5143 | 2.31E-05 | 0.064 | CCCTGGGCCAGCA | *rplA* | 50S ribosomal protein L1 |
| AVAEIV_RS03290 | 666848 | 666860 | 175 | + | 12.5143 | 2.31E-05 | 0.0726 | CCCTGGGCCAGCA | - | DUF3299 domain-containing protein |
| AVAEIV_RS03685 | 741876 | 741888 | 138 | - | 12.5143 | 2.31E-05 | 0.064 | CCCTGGGCCAGCA | *tsaE* | threonylcarbamoyltransferase complex ATPase subunit type 1 TsaE |
| AVAEIV_RS06755 | 1392546 | 1392558 | 108 | + | 12.5143 | 2.31E-05 | 0.0726 | TAGATGGCCATCG | *algW* | Do family serine endopeptidase AlgW |
| AVAEIV_RS06760 | 1392546 | 1392558 | -10 | + | 12.5143 | 2.31E-05 | 0.064 | TAGATGGCCATCG | - | Nif3-like dinuclear metal center hexameric protein |
| AVAEIV_RS07025 | 1447521 | 1447533 | 269 | + | 12.5143 | 2.31E-05 | 0.064 | CGCCTCGCCACCG | - | HugZ family protein |
| AVAEIV_RS07445 | 1549012 | 1549024 | 156 | - | 12.5143 | 2.31E-05 | 0.064 | CCCTGGGCCAGCA | - | efflux RND transporter permease subunit |
| AVAEIV_RS09870 | 2067772 | 2067784 | 335 | + | 12.5143 | 2.31E-05 | 0.0726 | CGCCTCGCCACCG | *rsxC* | electron transport complex subunit RsxC |
| AVAEIV_RS13230 | 2869626 | 2869638 | 373 | + | 12.5143 | 2.31E-05 | 0.064 | CACGGCGCCACCA | - | tyrosine-type recombinase/integrase |
| AVAEIV_RS19995 | 4262431 | 4262443 | 178 | - | 12.5143 | 2.31E-05 | 0.0726 | CACGGCGCCACCA | - | ABC transporter substrate-binding protein |
| AVAEIV_RS23975 | 5106920 | 5106932 | 45 | + | 12.5143 | 2.31E-05 | 0.0726 | CCCTGGGCCAGCA | - | uroporphyrinogen-III synthase |
| AVAEIV_RS24190 | 5143015 | 5143027 | 364 | - | 12.5143 | 2.31E-05 | 0.0726 | CCCTGGGCCAGCA | *rpmB* | 50S ribosomal protein L28 |
| AVAEIV_RS24195 | 5143015 | 5143027 | -50 | - | 12.5143 | 2.31E-05 | 0.064 | CCCTGGGCCAGCA | - | ABC transporter substrate-binding protein |
| AVAEIV_RS25480 | 5427150 | 5427162 | 225 | - | 12.5143 | 2.31E-05 | 0.0726 | CGCCTCGCCACCG | *norR* | nitric oxide reductase transcriptional regulator NorR |
| AVAEIV_RS00750 | 148440 | 148452 | 3 | - | 12.5 | 2.33E-05 | 0.064 | CCGAGCGCCGGCA | - | DUF364 domain-containing protein |
| AVAEIV_RS00810 | 157810 | 157822 | 184 | - | 12.5 | 2.33E-05 | 0.064 | CAGATCGCCGGCC | - | nitrogen fixation protein NifZ |
| AVAEIV_RS01395 | 272515 | 272527 | 107 | + | 12.5 | 2.33E-05 | 0.0726 | CGTTTCGCCATCG | - | exodeoxyribonuclease III |
| AVAEIV_RS01400 | 272515 | 272527 | -40 | + | 12.5 | 2.33E-05 | 0.064 | CGTTTCGCCATCG | *pyrE* | orotate phosphoribosyltransferase |
| AVAEIV_RS02635 | 535378 | 535390 | 293 | + | 12.5 | 2.33E-05 | 0.064 | CAGATCGCCGGCC | *garD* | galactarate dehydratase |
| AVAEIV_RS04155 | 844912 | 844924 | -31 | + | 12.5 | 2.33E-05 | 0.064 | CGTTTCGCCATCG | - | efflux RND transporter permease subunit |
| AVAEIV_RS05255 | 1082439 | 1082451 | 141 | - | 12.5 | 2.33E-05 | 0.0726 | CAGATCGCCGGCC | - | hydantoinase B/oxoprolinase family protein |
| AVAEIV_RS06475 | 1336022 | 1336034 | 261 | - | 12.5 | 2.33E-05 | 0.0726 | CCGAGCGCCGGCA | - | sulfite reductase subunit alpha |
| AVAEIV_RS06495 | 1339182 | 1339194 | 241 | + | 12.5 | 2.33E-05 | 0.0726 | CAGATCGCCGGCC | - | tetratricopeptide repeat protein |
| AVAEIV_RS06510 | 1342044 | 1342056 | 222 | + | 12.5 | 2.33E-05 | 0.064 | CTGCTGGCCACCA | - | PepSY domain-containing protein |
| AVAEIV_RS06935 | 1426776 | 1426788 | 255 | + | 12.5 | 2.33E-05 | 0.064 | CCGCATGCCATCG | *murC* | UDP-N-acetylmuramate--L-alanine ligase |
| AVAEIV_RS11720 | 2483208 | 2483220 | 136 | + | 12.5 | 2.33E-05 | 0.0726 | CCGAGCGCCGGCA | *greB* | transcription elongation factor GreB |
| AVAEIV_RS14930 | 3254814 | 3254826 | 323 | + | 12.5 | 2.33E-05 | 0.064 | CCGAGCGCCGGCA | *xthA* | exodeoxyribonuclease III |
| AVAEIV_RS15390 | 3361790 | 3361802 | 360 | + | 12.5 | 2.33E-05 | 0.0726 | CTGCTGGCCACCA | *pdxB* | 4-phosphoerythronate dehydrogenase PdxB |

| | | | | | | | | | | |
|---|---|---|---|---|---|---|---|---|---|---|
| AVAEIV_RS15470 | 3381335 | 3381347 | 30 | + | 12.5 | 2.33E-05 | 0.064 | CCGAGCGCCGGCA | - | hypothetical protein |
| AVAEIV_RS16220 | 3536628 | 3536640 | 110 | - | 12.5 | 2.33E-05 | 0.064 | CACCCCGCCATCG | *cysT* | sulfate ABC transporter permease subunit CysT |
| AVAEIV_RS18365 | 3951538 | 3951550 | 165 | + | 12.5 | 2.33E-05 | 0.0726 | CGCAGGGCCACCA | - | alpha/beta hydrolase |
| AVAEIV_RS20330 | 4329741 | 4329753 | 90 | - | 12.5 | 2.33E-05 | 0.0726 | CATAGCGCCAGCG | - | type II toxin-antitoxin system RelE/ParE family toxin |
| AVAEIV_RS20335 | 4329741 | 4329753 | 251 | - | 12.5 | 2.33E-05 | 0.064 | CATAGCGCCAGCG | - | hypothetical protein |
| AVAEIV_RS20860 | 4436923 | 4436935 | 48 | - | 12.5 | 2.33E-05 | 0.0726 | CGTTTCGCCATCG | - | alpha/beta hydrolase family protein |
| AVAEIV_RS22490 | 4810624 | 4810636 | -6 | + | 12.5 | 2.33E-05 | 0.0726 | CCGCATGCCATCG | - | ParA family protein |
| AVAEIV_RS03540 | 714690 | 714702 | 251 | + | 12.4857 | 2.35E-05 | 0.0643 | CGGTTCGCCGCCA | - | hypothetical protein |
| AVAEIV_RS04955 | 1014278 | 1014290 | 204 | - | 12.4857 | 2.35E-05 | 0.0727 | CCCATCGCCATGA | *mmsB* | 3-hydroxyisobutyrate dehydrogenase |
| AVAEIV_RS11170 | 2358797 | 2358809 | 80 | - | 12.4857 | 2.35E-05 | 0.0727 | CGCCGGGCCATCA | - | energy transducer TonB |
| AVAEIV_RS13100 | 2839627 | 2839639 | 232 | + | 12.4857 | 2.35E-05 | 0.0643 | CCGCTCGCCAGCC | - | L-serine ammonia-lyase |
| AVAEIV_RS21080 | 4483574 | 4483586 | 280 | - | 12.4857 | 2.35E-05 | 0.0727 | CTGGTCGCCACCA | - | 2-keto-4-pentenoate hydratase |
| AVAEIV_RS25685 | 5496232 | 5496244 | 236 | - | 12.4857 | 2.35E-05 | 0.0727 | CAGTTGGCCAGGA | - | 3-hydroxyacyl-CoA dehydrogenase family protein |
| AVAEIV_RS02760 | 565038 | 565050 | 55 | - | 12.4714 | 2.38E-05 | 0.0729 | CACTCCGCCAGCG | - | beta-ketoacyl synthase |
| AVAEIV_RS04200 | 857405 | 857417 | 227 | + | 12.4714 | 2.38E-05 | 0.0649 | CGCACCGCCATCG | *dmpE* | 2-oxopent-4-enoate hydratase |
| AVAEIV_RS06135 | 1251674 | 1251686 | 168 | - | 12.4714 | 2.38E-05 | 0.0729 | CTCATCGCCGTCA | *pilV* | adhesin PilV |
| AVAEIV_RS12975 | 2811546 | 2811558 | 298 | - | 12.4714 | 2.38E-05 | 0.0729 | CGCACCGCCATCG | - | nucleotidyltransferase family protein |
| AVAEIV_RS19060 | 4106285 | 4106297 | 358 | - | 12.4714 | 2.38E-05 | 0.0729 | CGCATCGCCATCT | *tsf* | translation elongation factor Ts |
| AVAEIV_RS25690 | 5497178 | 5497190 | -49 | - | 12.4714 | 2.38E-05 | 0.0729 | CGCACCGCCATCG | - | acetyl-CoA C-acetyltransferase |
| AVAEIV_RS07470 | 1556382 | 1556394 | 230 | - | 12.4571 | 2.42E-05 | 0.0653 | CACAAGGCCGGCA | - | helix-turn-helix domain-containing protein |
| AVAEIV_RS08145 | 1691705 | 1691717 | 18 | - | 12.4571 | 2.42E-05 | 0.0653 | CCGATGGCCGCCA | *serC* | transaminase |
| AVAEIV_RS12115 | 2571468 | 2571480 | 348 | + | 12.4571 | 2.42E-05 | 0.0731 | CGCTGGGCCAGCA | *zapE* | cell division protein ZapE |
| AVAEIV_RS14645 | 3187131 | 3187143 | 226 | - | 12.4571 | 2.42E-05 | 0.0653 | CAGTACGCCACCC | - | GNAT family N-acetyltransferase |
| AVAEIV_RS16550 | 3591091 | 3591103 | 216 | + | 12.4571 | 2.42E-05 | 0.0731 | CTGCACGCCATCG | - | UDP-2,3-diacylglucosamine diphosphatase |
| AVAEIV_RS17090 | 3707180 | 3707192 | 218 | + | 12.4571 | 2.42E-05 | 0.0731 | CACTTTGCCGGCA | - | hypothetical protein |
| AVAEIV_RS17800 | 3851148 | 3851160 | 329 | - | 12.4571 | 2.42E-05 | 0.0653 | CACATGGCCGGCG | - | NAD(P)-dependent alcohol dehydrogenase |
| AVAEIV_RS19700 | 4199835 | 4199847 | 380 | + | 12.4571 | 2.42E-05 | 0.0731 | CGCTGGGCCAGCA | - | DUF72 domain-containing protein |
| AVAEIV_RS21625 | 4603945 | 4603957 | 294 | - | 12.4571 | 2.42E-05 | 0.0731 | CAGTGCGCCGTCG | *glcC* | transcriptional regulator GlcC |
| AVAEIV_RS21630 | 4603945 | 4603957 | -39 | - | 12.4571 | 2.42E-05 | 0.0653 | CAGTGCGCCGTCG | *glcD* | glycolate oxidase subunit GlcD |
| AVAEIV_RS22190 | 4736734 | 4736746 | 191 | + | 12.4571 | 2.42E-05 | 0.0731 | CACATGGCCGGCG | - | acyl-CoA dehydrogenase family protein |
| AVAEIV_RS26070 | 5606081 | 5606093 | 86 | + | 12.4571 | 2.42E-05 | 0.0731 | CTGCACGCCATCG | - | protein |
| AVAEIV_RS04755 | 970143 | 970155 | 361 | + | 12.4429 | 2.45E-05 | 0.0733 | CCGCTGGCCGTCA | - | HopJ type III effector protein |
| AVAEIV_RS04760 | 970143 | 970155 | 71 | + | 12.4429 | 2.45E-05 | 0.0733 | CCGCTGGCCGTCA | - | DUF1244 domain-containing protein |
| AVAEIV_RS13140 | 2850921 | 2850933 | 226 | + | 12.4429 | 2.45E-05 | 0.0733 | CCCCAGGCCAGCA | - | binding protein |
| AVAEIV_RS16825 | 3646135 | 3646147 | -1 | - | 12.4429 | 2.45E-05 | 0.0733 | CATCAGGCCATCA | - | class I SAM-dependent DNA methyltransferase |
| AVAEIV_RS17600 | 3812885 | 3812897 | 329 | + | 12.4429 | 2.45E-05 | 0.0657 | CCCCAGGCCAGCA | - | serine protein kinase PrkA |
| AVAEIV_RS23085 | 4937152 | 4937164 | 294 | - | 12.4429 | 2.45E-05 | 0.0657 | CCGCTGGCCGTCA | - | TraR/DksA family transcriptional regulator |
| AVAEIV_RS02040 | 407884 | 407896 | 246 | - | 12.4286 | 2.48E-05 | 0.0733 | CGCATCGCCATGA | - | Rha family transcriptional regulator |
| AVAEIV_RS02765 | 566580 | 566592 | 97 | - | 12.4286 | 2.48E-05 | 0.0733 | CACTTCGCCAGGA | *metK* | methionine adenosyltransferase |
| AVAEIV_RS06440 | 1326590 | 1326602 | 81 | + | 12.4286 | 2.48E-05 | 0.0733 | CTTTTCGCCATCA | - | MBG domain-containing protein |
| AVAEIV_RS07215 | 1488096 | 1488108 | 260 | - | 12.4286 | 2.48E-05 | 0.0733 | CACTTCGCCAGGA | - | GntP family permease |
| AVAEIV_RS13225 | 2868006 | 2868018 | 188 | - | 12.4286 | 2.48E-05 | 0.0662 | CTTTTCGCCATCA | *ahpF* | alkyl hydroperoxide reductase subunit F |
| AVAEIV_RS17575 | 3810480 | 3810492 | 243 | + | 12.4286 | 2.48E-05 | 0.0733 | CGCCTTGCCACCA | - | cation acetate symporter |
| AVAEIV_RS17585 | 3810480 | 3810492 | 359 | + | 12.4286 | 2.48E-05 | 0.0662 | CGCCTTGCCACCA | - | DUF2061 domain-containing protein |
| AVAEIV_RS20125 | 4298609 | 4298621 | 329 | - | 12.4286 | 2.48E-05 | 0.0733 | CGGCTCGCCAGCC | - | TonB-dependent siderophore receptor |
| AVAEIV_RS23665 | 5044625 | 5044637 | -22 | + | 12.4286 | 2.48E-05 | 0.0662 | CCCGACGCCAGCA | - | Crp/Fnr family transcriptional regulator |
| AVAEIV_RS25875 | 5552110 | 5552122 | 41 | - | 12.4286 | 2.48E-05 | 0.0733 | CGGCTCGCCAGCC | - | VWA domain-containing protein |
| AVAEIV_RS02465 | 492810 | 492822 | 278 | - | 12.4143 | 2.50E-05 | 0.0737 | CCGAACGCCGTCG | - | TolC family protein |
| AVAEIV_RS19040 | 4103028 | 4103040 | 340 | - | 12.4143 | 2.50E-05 | 0.0737 | CAGGTCGCCGCCA | - | phosphatidate cytidylyltransferase |
| AVAEIV_RS20665 | 4394747 | 4394759 | 358 | + | 12.4143 | 2.50E-05 | 0.0737 | CAGGTCGCCGCCA | - | PA4642 family protein |
| AVAEIV_RS25490 | 5428445 | 5428457 | 35 | + | 12.4143 | 2.50E-05 | 0.0667 | CGGTCCGCCACCA | - | CBS domain-containing protein |
| AVAEIV_RS04960 | 1015139 | 1015151 | 242 | - | 12.4 | 2.52E-05 | 0.0739 | CAGGTCGGCATCA | - | enoyl-CoA hydratase |

| | | | | | | | | | | |
|---|---|---|---|---|---|---|---|---|---|---|
| AVAEIV_RS07175 | 1479354 | 1479366 | 142 | - | 12.4 | 2.52E-05 | 0.067 | TAGTTCGCCACCA | *pdxJ* | pyridoxine 5'-phosphate synthase |
| AVAEIV_RS14430 | 3143411 | 3143423 | 25 | + | 12.4 | 2.52E-05 | 0.0739 | CCCATCGCCGCCA | - | glutaminase |
| AVAEIV_RS16790 | 3633384 | 3633396 | 348 | + | 12.4 | 2.52E-05 | 0.067 | GCGATGGCCATCA | - | dienelactone hydrolase family protein |
| AVAEIV_RS17490 | 3787958 | 3787970 | 228 | + | 12.4 | 2.52E-05 | 0.0739 | CGGTGCGCCACCG | - | DsbA family protein |
| AVAEIV_RS00515 | 107348 | 107360 | 275 | - | 12.3857 | 2.56E-05 | 0.074 | CGCCTGGCCAGCG | - | SulP family inorganic anion transporter |
| AVAEIV_RS01880 | 378307 | 378319 | 383 | + | 12.3857 | 2.56E-05 | 0.074 | CACACGGCCAGCG | - | DUF3141 domain-containing protein |
| AVAEIV_RS04080 | 828278 | 828290 | 100 | + | 12.3857 | 2.56E-05 | 0.067 | CGCCTGGCCAGCG | *rsfS* | ribosome silencing factor |
| AVAEIV_RS05570 | 1150912 | 1150924 | 167 | + | 12.3857 | 2.56E-05 | 0.067 | CGCCAGGCCAGCA | - | IS5 family transposase |
| AVAEIV_RS10200 | 2128032 | 2128044 | 244 | + | 12.3857 | 2.56E-05 | 0.074 | CAGAACGGCATCG | *ccoS* | cbb3-type cytochrome oxidase assembly protein CcoS |
| AVAEIV_RS11110 | 2345012 | 2345024 | 351 | - | 12.3857 | 2.56E-05 | 0.074 | CAGATCGCCTCCA | - | sigma-70 family RNA polymerase sigma factor |
| AVAEIV_RS11115 | 2345012 | 2345024 | 77 | - | 12.3857 | 2.56E-05 | 0.067 | CAGATCGCCTCCA | - | acyl-CoA dehydrogenase family protein |
| AVAEIV_RS12980 | 2812187 | 2812199 | -47 | + | 12.3857 | 2.56E-05 | 0.074 | CGCCAGGCCAGCA | - | XdhC family protein |
| AVAEIV_RS14035 | 3049801 | 3049813 | 290 | - | 12.3857 | 2.56E-05 | 0.074 | CAATTGGCCGGCA | - | phytanoyl-CoA dioxygenase family protein |
| AVAEIV_RS14040 | 3049801 | 3049813 | 115 | - | 12.3857 | 2.56E-05 | 0.067 | CAATTGGCCGGCA | - | FagA protein |
| AVAEIV_RS17475 | 3782656 | 3782668 | 21 | - | 12.3857 | 2.56E-05 | 0.067 | CAGAACGGCATCG | - | PHB depolymerase family esterase |
| AVAEIV_RS22170 | 4730817 | 4730829 | 102 | - | 12.3857 | 2.56E-05 | 0.067 | CGCCTGGCCAGCG | - | hypothetical protein |
| AVAEIV_RS24025 | 5114134 | 5114146 | 14 | - | 12.3857 | 2.56E-05 | 0.067 | CAGTTGGGCAGCA | *dapF* | diaminopimelate epimerase |
| AVAEIV_RS04580 | 938062 | 938074 | 201 | - | 12.3714 | 2.59E-05 | 0.074 | CACATCGGCACCA | - | urease subunit beta |
| AVAEIV_RS06775 | 1397044 | 1397056 | 122 | + | 12.3714 | 2.59E-05 | 0.074 | CAGTGTGCCGTCA | - | YhcB family protein |
| AVAEIV_RS06780 | 1397044 | 1397056 | 134 | + | 12.3714 | 2.59E-05 | 0.0672 | CAGTGTGCCGTCA | - | alpha/beta hydrolase |
| AVAEIV_RS09325 | 1944932 | 1944944 | 146 | + | 12.3714 | 2.59E-05 | 0.0672 | CACTGCGCCATCC | - | TSUP family transporter |
| AVAEIV_RS15330 | 3350156 | 3350168 | 167 | - | 12.3714 | 2.59E-05 | 0.074 | CGCGTCGCCAGCG | *sucB* | dihydrolipoyllysine-residue succinyltransferase |
| AVAEIV_RS17150 | 3717953 | 3717965 | 373 | - | 12.3714 | 2.59E-05 | 0.074 | CACTGTGCCAGCG | - | SDR family oxidoreductase |
| AVAEIV_RS19330 | 4151951 | 4151963 | 108 | - | 12.3714 | 2.59E-05 | 0.074 | CGCAGTGCCATCG | - | hypothetical protein |
| AVAEIV_RS20060 | 4280468 | 4280480 | 211 | + | 12.3714 | 2.59E-05 | 0.074 | CCGCCGGCCATCA | - | hypothetical protein |
| AVAEIV_RS22100 | 4717619 | 4717631 | 360 | - | 12.3714 | 2.59E-05 | 0.074 | CCGAGGGCCACCG | - | NTP transferase domain-containing protein |
| AVAEIV_RS22270 | 4754193 | 4754205 | -14 | + | 12.3714 | 2.59E-05 | 0.074 | CACCTCGCCGCCA | - | acyltransferase family protein |
| AVAEIV_RS23295 | 4966271 | 4966283 | 273 | - | 12.3714 | 2.59E-05 | 0.0672 | CGCGTCGCCAGCG | - | ribose-phosphate diphosphokinase |
| AVAEIV_RS23725 | 5055108 | 5055120 | 317 | + | 12.3714 | 2.59E-05 | 0.0672 | CCGCCGGCCATCA | - | protein |
| AVAEIV_RS25935 | 5571727 | 5571739 | 222 | - | 12.3714 | 2.59E-05 | 0.0672 | GAGCTGGCCATCA | - | ABC transporter substrate-binding protein |
| AVAEIV_RS01775 | 350223 | 350235 | 34 | - | 12.3571 | 2.62E-05 | 0.074 | CAGCCGGCCACCA | - | sigma-54-dependent transcriptional regulator |
| AVAEIV_RS07740 | 1612104 | 1612116 | 22 | - | 12.3571 | 2.62E-05 | 0.0675 | CAGCAGGCCATCC | - | DNA polymerase III subunit delta' |
| AVAEIV_RS14165 | 3082223 | 3082235 | 23 | + | 12.3571 | 2.62E-05 | 0.0675 | CCCTTCGCCGGCA | - | lactonase family protein |
| AVAEIV_RS17420 | 3772021 | 3772033 | 275 | - | 12.3571 | 2.62E-05 | 0.0675 | TAGAGCGCCAGCA | *ggpS* | glucosylglycerol-phosphate synthase |
| AVAEIV_RS20225 | 4316181 | 4316193 | 214 | - | 12.3571 | 2.62E-05 | 0.074 | CAGCCGGCCACCA | *cysE* | serine O-acetyltransferase |
| AVAEIV_RS21600 | 4598527 | 4598539 | 172 | + | 12.3571 | 2.62E-05 | 0.074 | CAGCAGGCCATCC | - | glycerate kinase |
| AVAEIV_RS01585 | 304731 | 304743 | 145 | + | 12.3429 | 2.64E-05 | 0.074 | CCGGGCGCCATCG | - | hypothetical protein |
| AVAEIV_RS01590 | 304731 | 304743 | 166 | + | 12.3429 | 2.64E-05 | 0.0675 | CCGGGCGCCATCG | - | ATP-binding protein |
| AVAEIV_RS04840 | 985054 | 985066 | 263 | + | 12.3429 | 2.64E-05 | 0.0675 | CCGGGCGCCATCG | *nhaA* | Na+/H+ antiporter NhaA |
| AVAEIV_RS05190 | 1067449 | 1067461 | 216 | - | 12.3429 | 2.64E-05 | 0.074 | CCGTCGGCCAGCA | *yaaA* | peroxide stress protein YaaA |
| AVAEIV_RS07430 | 1544829 | 1544841 | 98 | - | 12.3429 | 2.64E-05 | 0.074 | CCGGGCGCCATCG | - | ATP-binding protein |
| AVAEIV_RS07435 | 1544829 | 1544841 | 21 | - | 12.3429 | 2.64E-05 | 0.0675 | CCGGGCGCCATCG | - | MdtA/MuxA family multidrug efflux RND transporter periplasmic adaptor subunit |
| AVAEIV_RS10110 | 2112241 | 2112253 | 346 | - | 12.3429 | 2.64E-05 | 0.074 | GCCATCGCCATCA | *phnE* | phosphonate ABC transporter, permease protein PhnE |
| AVAEIV_RS13155 | 2854761 | 2854773 | 83 | - | 12.3429 | 2.64E-05 | 0.074 | CCGGGCGCCATCG | *hutU* | urocanate hydratase |
| AVAEIV_RS15725 | 3435892 | 3435904 | 246 | + | 12.3429 | 2.64E-05 | 0.0675 | CGCATCGCCGCCA | - | 2-hydroxyacid dehydrogenase |
| AVAEIV_RS16630 | 3606469 | 3606481 | 285 | - | 12.3429 | 2.64E-05 | 0.074 | CGCATCGCCGCCA | - | YcgN family cysteine cluster protein |
| AVAEIV_RS18655 | 4017266 | 4017278 | 304 | + | 12.3429 | 2.64E-05 | 0.074 | CGCATCGCCGCCA | *quiC* | 3-dehydroshikimate dehydratase QuiC |
| AVAEIV_RS18660 | 4017266 | 4017278 | 144 | + | 12.3429 | 2.64E-05 | 0.0675 | CGCATCGCCGCCA | - | OprD family porin |
| AVAEIV_RS19180 | 4133393 | 4133405 | 282 | + | 12.3429 | 2.64E-05 | 0.074 | CCGAACGCCATCT | - | hypothetical protein |
| AVAEIV_RS19750 | 4208903 | 4208915 | 178 | - | 12.3429 | 2.64E-05 | 0.074 | CAGGATGCCAGCG | - | hypothetical protein |
| AVAEIV_RS24250 | 5157199 | 5157211 | 350 | - | 12.3429 | 2.64E-05 | 0.074 | CTGTTCGCCATCC | - | thymidylate synthase |

| | | | | | | | | | | |
|---|---|---|---|---|---|---|---|---|---|---|
| AVAEIV_RS25675 | 5490378 | 5490390 | 275 | + | 12.3429 | 2.64E-05 | 0.0675 | CAGGACGCCATCC | - | efflux RND transporter permease subunit |
| AVAEIV_RS01655 | 319628 | 319640 | 67 | + | 12.3286 | 2.67E-05 | 0.068 | TCGATGGCCATCA | - | hypothetical protein |
| AVAEIV_RS12010 | 2552543 | 2552555 | 52 | + | 12.3286 | 2.67E-05 | 0.0743 | CCGAATGCCGTCA | - | acetyl-CoA C-acetyltransferase |
| AVAEIV_RS15790 | 3446936 | 3446948 | 250 | + | 12.3286 | 2.67E-05 | 0.0743 | CAGTAGGCCAGCC | *arsC* | arsenate reductase (glutaredoxin) |
| AVAEIV_RS18485 | 3980050 | 3980062 | -50 | + | 12.3286 | 2.67E-05 | 0.068 | CAGAACGCCACCT | *olsB* | L-ornithine N(alpha)-acyltransferase |
| AVAEIV_RS19445 | 4164184 | 4164196 | -28 | - | 12.3286 | 2.67E-05 | 0.0743 | CAAATCGCCAACA | - | phage antirepressor KilAC domain-containing protein |
| AVAEIV_RS19675 | 4195381 | 4195393 | 324 | + | 12.3286 | 2.67E-05 | 0.0743 | CACACCGCCGTCA | - | ABC transporter permease subunit |
| AVAEIV_RS00660 | 134935 | 134947 | 44 | - | 12.3143 | 2.70E-05 | 0.0743 | CCCAGCGCCACCG | - | hypothetical protein |
| AVAEIV_RS12615 | 2678233 | 2678245 | 351 | + | 12.3143 | 2.70E-05 | 0.0743 | CGAACGGCCATCA | - | MFS transporter |
| AVAEIV_RS15390 | 3361698 | 3361710 | 268 | - | 12.3143 | 2.70E-05 | 0.0743 | CGGCCGGCCATCA | *pdxB* | 4-phosphoerythronate dehydrogenase PdxB |
| AVAEIV_RS15460 | 3378943 | 3378955 | 76 | + | 12.3143 | 2.70E-05 | 0.0682 | CTCCTGGCCAGCA | - | glycosyltransferase family 4 protein |
| AVAEIV_RS20140 | 4301835 | 4301847 | 298 | + | 12.3143 | 2.70E-05 | 0.0743 | CAGATTGCCGCCG | *der* | ribosome biogenesis GTPase Der |
| AVAEIV_RS20450 | 4345981 | 4345993 | 123 | + | 12.3143 | 2.70E-05 | 0.0682 | CACAACGCCACCC | - | DUF4105 domain-containing protein |
| AVAEIV_RS21275 | 4523635 | 4523647 | 20 | + | 12.3143 | 2.70E-05 | 0.0743 | CAGATTGCCGCCG | - | acetolactate synthase 3 large subunit |
| AVAEIV_RS21280 | 4523635 | 4523647 | 365 | + | 12.3143 | 2.70E-05 | 0.0682 | CAGATTGCCGCCG | - | DUF4124 domain-containing protein |
| AVAEIV_RS02665 | 544643 | 544655 | 283 | - | 12.3 | 2.73E-05 | 0.0744 | CACGAGGCCATCG | - | lipopolysaccharide biosynthesis protein |
| AVAEIV_RS03040 | 623990 | 624002 | 130 | + | 12.3 | 2.73E-05 | 0.0685 | CCGATCGCCATGG | - | tRNA-Trp |
| AVAEIV_RS03045 | 623990 | 624002 | 248 | + | 12.3 | 2.73E-05 | 0.0685 | CCGATCGCCATGG | *secE* | preprotein translocase subunit SecE |
| AVAEIV_RS03105 | 640463 | 640475 | 227 | + | 12.3 | 2.73E-05 | 0.0685 | CCGATCGCCATGG | *rpsJ* | 30S ribosomal protein S10 |
| AVAEIV_RS03385 | 681312 | 681324 | 234 | + | 12.3 | 2.73E-05 | 0.0744 | CGCTTCGCCGGCA | - | termination factor Rho |
| AVAEIV_RS06590 | 1360820 | 1360832 | 18 | - | 12.3 | 2.73E-05 | 0.0744 | CCGCATGCCACCA | *gatB* | subunit GatB |
| AVAEIV_RS06710 | 1383734 | 1383746 | 268 | + | 12.3 | 2.73E-05 | 0.0685 | CCGATCGCCATGG | *mlaE* | permease subunit MlaE |
| AVAEIV_RS14540 | 3166946 | 3166958 | 328 | - | 12.3 | 2.73E-05 | 0.0744 | CGTTTCGCCACCA | *serS* | serine--tRNA ligase |
| AVAEIV_RS20190 | 4310273 | 4310285 | 173 | - | 12.3 | 2.73E-05 | 0.0744 | CGGGCCGCCATCA | *fdx* | ISC system 2Fe-2S type ferredoxin |
| AVAEIV_RS02990 | 617373 | 617385 | 128 | - | 12.2857 | 2.76E-05 | 0.0685 | CAGGTGGCCGGCA | - | CDP-alcohol phosphatidyltransferase family protein |
| AVAEIV_RS04770 | 971399 | 971411 | 71 | + | 12.2857 | 2.76E-05 | 0.0744 | CACCGCGCCACCG | *folM* | dihydromonapterin reductase |
| AVAEIV_RS04775 | 971399 | 971411 | 217 | + | 12.2857 | 2.76E-05 | 0.0685 | CACCGCGCCACCG | *mdcA* | malonate decarboxylase subunit alpha |
| AVAEIV_RS05345 | 1104031 | 1104043 | 292 | + | 12.2857 | 2.76E-05 | 0.0685 | CTCAACGCCAGCG | - | hypothetical protein |
| AVAEIV_RS05530 | 1145613 | 1145625 | 319 | + | 12.2857 | 2.76E-05 | 0.0685 | CTGATCGCCGTCG | - | tRNA-Asn |
| AVAEIV_RS12885 | 2781299 | 2781311 | 371 | - | 12.2857 | 2.76E-05 | 0.0744 | CAGGTGGCCGGCA | - | thioesterase II family protein |
| AVAEIV_RS12890 | 2781299 | 2781311 | 117 | - | 12.2857 | 2.76E-05 | 0.0744 | CAGGTGGCCGGCA | - | MbtH family protein |
| AVAEIV_RS17740 | 3842142 | 3842154 | 326 | + | 12.2857 | 2.76E-05 | 0.0744 | CAGGTGGCCGGCA | *dapA* | 4-hydroxy-tetrahydrodipicolinate synthase |
| AVAEIV_RS18265 | 3939801 | 3939813 | 333 | - | 12.2857 | 2.76E-05 | 0.0744 | CACCGCGCCACCG | - | hypothetical protein |
| AVAEIV_RS18270 | 3939801 | 3939813 | 115 | - | 12.2857 | 2.76E-05 | 0.0744 | CACCGCGCCACCG | - | hypothetical protein |
| AVAEIV_RS20320 | 4328429 | 4328441 | 202 | + | 12.2857 | 2.76E-05 | 0.0685 | CTGAACGCCGTCA | - | hypothetical protein |
| AVAEIV_RS20650 | 4391680 | 4391692 | -43 | - | 12.2857 | 2.76E-05 | 0.0744 | CTGATCGCCGTCG | - | YajG family lipoprotein |
| AVAEIV_RS22245 | 4747408 | 4747420 | 301 | + | 12.2857 | 2.76E-05 | 0.0685 | CAGATCGCCACGG | - | aldo/keto reductase |
| AVAEIV_RS03480 | 701360 | 701372 | 256 | - | 12.2714 | 2.79E-05 | 0.0687 | CAGTTTGCCGGCG | - | nucleotidyltransferase family protein |
| AVAEIV_RS07575 | 1580237 | 1580249 | 191 | - | 12.2714 | 2.79E-05 | 0.0687 | CAGTTTGCCGGCG | *nqrE* | subunit E |
| AVAEIV_RS07890 | 1640944 | 1640956 | 190 | - | 12.2714 | 2.79E-05 | 0.0746 | CCCTGCGCCAGCG | - | regulator |
| AVAEIV_RS12410 | 2629331 | 2629343 | 4 | - | 12.2714 | 2.79E-05 | 0.0687 | CGGATTGCCGTCG | - | hypothetical protein |
| AVAEIV_RS12800 | 2720885 | 2720897 | 232 | + | 12.2714 | 2.79E-05 | 0.0746 | TAGTTGGCCAGCA | - | integrase core domain-containing protein |
| AVAEIV_RS12920 | 2801581 | 2801593 | 323 | - | 12.2714 | 2.79E-05 | 0.0746 | CAGTTTGCCGGCG | - | DNA-binding protein |
| AVAEIV_RS22195 | 4737508 | 4737520 | 12 | + | 12.2714 | 2.79E-05 | 0.0746 | CCCTGCGCCAGCG | - | protein |
| AVAEIV_RS23675 | 5045736 | 5045748 | 1 | - | 12.2714 | 2.79E-05 | 0.0687 | CAGAAGGCCGGCG | - | tRNA-Ile |
| AVAEIV_RS23680 | 5045736 | 5045748 | 257 | - | 12.2714 | 2.79E-05 | 0.0687 | CAGAAGGCCGGCG | - | LemA family protein |
| AVAEIV_RS25395 | 5411368 | 5411380 | 321 | + | 12.2714 | 2.79E-05 | 0.0746 | CAGCTCGCCATGG | *nifX* | nitrogen fixation protein NifX |
| AVAEIV_RS25400 | 5411368 | 5411380 | 36 | + | 12.2714 | 2.79E-05 | 0.0746 | CAGCTCGCCATGG | - | RnfH family protein |
| AVAEIV_RS05325 | 1098484 | 1098496 | 106 | + | 12.2571 | 2.82E-05 | 0.0688 | CCGCAGGCCAGCG | - | DUF2218 domain-containing protein |
| AVAEIV_RS05845 | 1196892 | 1196904 | -10 | + | 12.2571 | 2.82E-05 | 0.0688 | CTGATGGCCATCC | - | family protein |
| AVAEIV_RS08885 | 1845084 | 1845096 | -33 | - | 12.2571 | 2.82E-05 | 0.0688 | CAGGTTGCCATCC | *cas2e* | type I-E CRISPR-associated endoribonuclease Cas2e |
| AVAEIV_RS15090 | 3283658 | 3283670 | 199 | + | 12.2571 | 2.82E-05 | 0.0748 | CTGCTCGCCACCG | - | carbonate dehydratase |

| | | | | | | | | | | |
|---|---|---|---|---|---|---|---|---|---|---|
| AVAEIV_RS16930 | 3666897 | 3666909 | 353 | + | 12.2571 | 2.82E-05 | 0.0748 | CCGACTGCCATCG | - | maltoporin |
| AVAEIV_RS18415 | 3969146 | 3969158 | 47 | + | 12.2571 | 2.82E-05 | 0.0748 | CAGTGCGCCGCCA | *dinB* | DNA polymerase IV |
| AVAEIV_RS21495 | 4574248 | 4574260 | 119 | - | 12.2571 | 2.82E-05 | 0.0748 | CCAAAGGCCAGCG | *dapB* | 4-hydroxy-tetrahydrodipicolinate reductase |
| AVAEIV_RS21870 | 4661268 | 4661280 | -42 | + | 12.2571 | 2.82E-05 | 0.0748 | CGCAGCGCCACCG | - | acyl-CoA dehydrogenase family protein |
| AVAEIV_RS23060 | 4934004 | 4934016 | 282 | - | 12.2571 | 2.82E-05 | 0.0688 | CCATTCGCCATCC | - | head completion/stabilization protein |
| AVAEIV_RS23740 | 5059631 | 5059643 | -29 | - | 12.2571 | 2.82E-05 | 0.0688 | CCGCAGGCCAGCG | - | ABC transporter ATP-binding protein |
| AVAEIV_RS01150 | 224522 | 224534 | -10 | + | 12.2429 | 2.85E-05 | 0.0689 | CACATGGGCAGCA | - | CsbD family protein |
| AVAEIV_RS05995 | 1221763 | 1221775 | 27 | - | 12.2429 | 2.85E-05 | 0.0753 | CACCTGGCCGGCA | - | membrane protein |
| AVAEIV_RS10105 | 2110301 | 2110313 | -43 | + | 12.2429 | 2.85E-05 | 0.0689 | CATCTGGCCACCA | - | F0F1 ATP synthase subunit gamma |
| AVAEIV_RS11665 | 2471969 | 2471981 | 186 | + | 12.2429 | 2.85E-05 | 0.0753 | CAAAACGCCATTA | - | DEAD/DEAH box helicase |
| AVAEIV_RS17900 | 3875236 | 3875248 | 27 | + | 12.2429 | 2.85E-05 | 0.0689 | CACCTGGCCGGCA | - | membrane protein |
| AVAEIV_RS23150 | 4945481 | 4945493 | 107 | - | 12.2429 | 2.85E-05 | 0.0689 | CGCCGCGCCATCG | - | hypothetical protein |
| AVAEIV_RS24670 | 5242800 | 5242812 | 333 | - | 12.2429 | 2.85E-05 | 0.0689 | CGCCGCGCCATCG | - | c-type cytochrome |
| AVAEIV_RS02675 | 546621 | 546633 | 48 | - | 12.2286 | 2.88E-05 | 0.0753 | CACGTCGCCGGCA | - | glycosyltransferase |
| AVAEIV_RS17730 | 3839926 | 3839938 | 233 | - | 12.2286 | 2.88E-05 | 0.0753 | CCGAACGCTATCA | - | synthase |
| AVAEIV_RS22115 | 4719373 | 4719385 | 312 | - | 12.2286 | 2.88E-05 | 0.0693 | CCGAGCGCCAGCC | - | CDP-alcohol phosphatidyltransferase family protein |
| AVAEIV_RS23385 | 4985459 | 4985471 | -44 | - | 12.2286 | 2.88E-05 | 0.0693 | CCGAGCGCCAGCC | - | efflux transporter outer membrane subunit |
| AVAEIV_RS04570 | 935754 | 935766 | 306 | + | 12.2143 | 2.91E-05 | 0.0753 | CGCTGCGCCAGCG | - | domain-containing protein |
| AVAEIV_RS06410 | 1309402 | 1309414 | 57 | - | 12.2143 | 2.91E-05 | 0.0694 | CGCTGCGCCAGCG | *pfkB* | 1-phosphofructokinase |
| AVAEIV_RS15780 | 3445311 | 3445323 | 226 | + | 12.2143 | 2.91E-05 | 0.0694 | CCGAACGCCGCCA | - | type II toxin-antitoxin system HicA family toxin |
| AVAEIV_RS16755 | 3627675 | 3627687 | 299 | - | 12.2143 | 2.91E-05 | 0.0753 | CGCATGGCCGGCA | - | NAD-dependent deacylase |
| AVAEIV_RS17255 | 3739167 | 3739179 | 312 | + | 12.2143 | 2.91E-05 | 0.0753 | CACATAGCCATCG | - | Hpt domain-containing protein |
| AVAEIV_RS21760 | 4635473 | 4635485 | 158 | - | 12.2143 | 2.91E-05 | 0.0753 | CGCTGCGCCAGCG | - | hypothetical protein |
| AVAEIV_RS21770 | 4635473 | 4635485 | 164 | - | 12.2143 | 2.91E-05 | 0.0694 | CGCTGCGCCAGCG | - | HipA N-terminal domain-containing protein |
| AVAEIV_RS22135 | 4723656 | 4723668 | 225 | + | 12.2143 | 2.91E-05 | 0.0694 | CTGACCGCCATCG | - | KGG domain-containing protein |
| AVAEIV_RS22710 | 4863171 | 4863183 | 201 | + | 12.2143 | 2.91E-05 | 0.0753 | CAGGCGGCCAGCA | - | lipocalin family protein |
| AVAEIV_RS22715 | 4863171 | 4863183 | -50 | + | 12.2143 | 2.91E-05 | 0.0753 | CAGGCGGCCAGCA | - | hypothetical protein |
| AVAEIV_RS23160 | 4946948 | 4946960 | 84 | - | 12.2143 | 2.91E-05 | 0.0694 | CCGAACGCCGCCA | - | hypothetical protein |
| AVAEIV_RS02695 | 551822 | 551834 | 212 | + | 12.2 | 2.95E-05 | 0.0753 | CCGCTCGCCGTCG | - | GumC family protein |
| AVAEIV_RS03960 | 798545 | 798557 | 205 | + | 12.2 | 2.95E-05 | 0.0753 | CGGGGTGCCATCA | - | energy transducer TonB |
| AVAEIV_RS04135 | 841965 | 841977 | 223 | - | 12.2 | 2.95E-05 | 0.0753 | GACATGGCCAGCA | - | lytic murein transglycosylase |
| AVAEIV_RS04145 | 841965 | 841977 | 375 | - | 12.2 | 2.95E-05 | 0.0696 | GACATGGCCAGCA | - | TolC family protein |
| AVAEIV_RS04440 | 903349 | 903361 | 211 | + | 12.2 | 2.95E-05 | 0.0696 | CATCACGCCATCG | *aldA* | aldehyde dehydrogenase |
| AVAEIV_RS08370 | 1747808 | 1747820 | 297 | - | 12.2 | 2.95E-05 | 0.0753 | CGAAAGGCCAGCG | - | hypothetical protein |
| AVAEIV_RS08765 | 1825874 | 1825886 | 112 | - | 12.2 | 2.95E-05 | 0.0753 | CTCATCGCCATCC | - | tyrosine-type recombinase/integrase |
| AVAEIV_RS09895 | 2072332 | 2072344 | 55 | - | 12.2 | 2.95E-05 | 0.0696 | CAAAGCGCCAGCC | *dcd* | dCTP deaminase |
| AVAEIV_RS10140 | 2115777 | 2115789 | 277 | - | 12.2 | 2.95E-05 | 0.0696 | CAGCGCGCCAGCC | *ffs* | signal recognition particle sRNA small type |
| AVAEIV_RS10925 | 2295234 | 2295246 | 195 | - | 12.2 | 2.95E-05 | 0.0753 | CCGCTCGCCGTCG | *puuE* | allantoinase PuuE |
| AVAEIV_RS13335 | 2887394 | 2887406 | 260 | + | 12.2 | 2.95E-05 | 0.0753 | CACAGCGCCATGA | - | transposase |
| AVAEIV_RS13725 | 2967638 | 2967650 | 376 | - | 12.2 | 2.95E-05 | 0.0696 | GACATGGCCAGCA | - | ABC transporter ATP-binding protein |
| AVAEIV_RS15520 | 3396779 | 3396791 | 344 | + | 12.2 | 2.95E-05 | 0.0753 | CAGAAGGCCAGCT | - | long-chain fatty acid--CoA ligase |
| AVAEIV_RS18245 | 3936263 | 3936275 | 167 | - | 12.2 | 2.95E-05 | 0.0753 | CCAATCGCCGTCG | - | hypothetical protein |
| AVAEIV_RS20820 | 4424295 | 4424307 | 53 | - | 12.2 | 2.95E-05 | 0.0696 | CAGCGCGCCAGCC | - | restriction system-associated AAA family ATPase |
| AVAEIV_RS21005 | 4467363 | 4467375 | 106 | + | 12.2 | 2.95E-05 | 0.0696 | CTCATCGCCATCC | - | GNAT family N-acetyltransferase |
| AVAEIV_RS22475 | 4806836 | 4806848 | 380 | - | 12.2 | 2.95E-05 | 0.0696 | CAGCGCGCCAGCC | - | sulfite exporter TauE/SafE family protein |
| AVAEIV_RS24310 | 5169064 | 5169076 | 180 | - | 12.2 | 2.95E-05 | 0.0753 | CCGAACGGCATCA | - | fumarylacetoacetate hydrolase family protein |
| AVAEIV_RS24540 | 5214824 | 5214836 | 277 | - | 12.2 | 2.95E-05 | 0.0753 | CAAATCGCTATCG | - | IS5/IS1182 family transposase |
| AVAEIV_RS00425 | 86305 | 86317 | 385 | - | 12.1857 | 2.98E-05 | 0.0696 | CAGCTCGCCGCCG | *znuB* | zinc ABC transporter permease subunit ZnuB |
| AVAEIV_RS06495 | 1339236 | 1339248 | 295 | + | 12.1857 | 2.98E-05 | 0.0753 | CAGCTCGCCGCCG | - | tetratricopeptide repeat protein |
| AVAEIV_RS13285 | 2877542 | 2877554 | 110 | + | 12.1857 | 2.98E-05 | 0.0696 | CGATGCGCCACCA | *ltrA* | group II intron reverse transcriptase/maturase |
| AVAEIV_RS18280 | 3940525 | 3940537 | 288 | - | 12.1857 | 2.98E-05 | 0.0753 | CAGCTCGCCGCCG | - | hypothetical protein |
| AVAEIV_RS18285 | 3940525 | 3940537 | 37 | - | 12.1857 | 2.98E-05 | 0.0753 | CAGCTCGCCGCCG | - | hypothetical protein |

| | | | | | | | | | | |
|---|---|---|---|---|---|---|---|---|---|---|
| AVAEIV_RS19890 | 4237771 | 4237783 | 191 | + | 12.1857 | 2.98E-05 | 0.0696 | CAGCTTGCCATGA | *mltF* | membrane-bound lytic murein transglycosylase MltF |
| AVAEIV_RS20615 | 4385412 | 4385424 | 358 | - | 12.1857 | 2.98E-05 | 0.0696 | CAGCTCGCCGCCG | - | TMEM165/GDT1 family protein |
| AVAEIV_RS22300 | 4762033 | 4762045 | 353 | + | 12.1857 | 2.98E-05 | 0.0753 | CAGATGGCCGTCC | - | O-antigen ligase family protein |
| AVAEIV_RS00740 | 147605 | 147617 | -33 | - | 12.1714 | 3.01E-05 | 0.0696 | CCGTTCGCCGGCG | - | hypothetical protein |
| AVAEIV_RS01490 | 287471 | 287483 | 257 | + | 12.1714 | 3.01E-05 | 0.0753 | CCGTTCGCCGGCG | - | YggT family protein |
| AVAEIV_RS02840 | 583069 | 583081 | 237 | + | 12.1714 | 3.01E-05 | 0.0696 | CCGTTCGCCGGCG | - | NfeD family protein |
| AVAEIV_RS04415 | 897443 | 897455 | -43 | + | 12.1714 | 3.01E-05 | 0.0696 | CTGATCGCCATGA | - | amidohydrolase family protein |
| AVAEIV_RS14300 | 3117323 | 3117335 | 10 | + | 12.1714 | 3.01E-05 | 0.0753 | CAAATCGGCATCG | - | phytanoyl-CoA dioxygenase family protein |
| AVAEIV_RS18255 | 3937038 | 3937050 | 205 | - | 12.1714 | 3.01E-05 | 0.0753 | CAGCTCGGCATCG | - | hypothetical protein |
| AVAEIV_RS22460 | 4801472 | 4801484 | 307 | - | 12.1714 | 3.01E-05 | 0.0696 | CAGCTCGGCATCG | *mscK* | mechanosensitive channel MscK |
| AVAEIV_RS23945 | 5100820 | 5100832 | 310 | + | 12.1714 | 3.01E-05 | 0.0696 | CAGCTCGGCATCG | - | TIGR02444 family protein |
| AVAEIV_RS02260 | 453850 | 453862 | 89 | - | 12.1571 | 3.04E-05 | 0.0753 | CGGAACGCCGCCA | *cooS* | subunit |
| AVAEIV_RS02775 | 567472 | 567484 | 321 | + | 12.1571 | 3.04E-05 | 0.0696 | CGGATCGCCGCCG | *tkt* | transketolase |
| AVAEIV_RS05430 | 1123502 | 1123514 | 80 | - | 12.1571 | 3.04E-05 | 0.0696 | CGGATCGCCGCCG | *ribF* | bifunctional riboflavin kinase/FAD synthetase |
| AVAEIV_RS07160 | 1475091 | 1475103 | 14 | - | 12.1571 | 3.04E-05 | 0.0696 | CCAAGGGCCACCA | *era* | GTPase Era |
| AVAEIV_RS07620 | 1589871 | 1589883 | 45 | - | 12.1571 | 3.04E-05 | 0.0753 | CAGCGGGCCGTCA | - | DUF2062 domain-containing protein |
| AVAEIV_RS07625 | 1589871 | 1589883 | 77 | - | 12.1571 | 3.04E-05 | 0.0696 | CAGCGGGCCGTCA | - | ComEC/Rec2 |
| AVAEIV_RS09915 | 2078589 | 2078601 | 217 | - | 12.1571 | 3.04E-05 | 0.0753 | CGGAACGCCGCCA | - | penicillin acylase family protein |
| AVAEIV_RS10085 | 2106896 | 2106908 | 145 | + | 12.1571 | 3.04E-05 | 0.0696 | CGGATCGCCGCCG | - | F0F1 ATP synthase subunit A |
| AVAEIV_RS10100 | 2108760 | 2108772 | -40 | + | 12.1571 | 3.04E-05 | 0.0696 | CGGATCGCCGCCG | - | F0F1 ATP synthase subunit alpha |
| AVAEIV_RS10195 | 2127729 | 2127741 | 155 | + | 12.1571 | 3.04E-05 | 0.0753 | CAGATGGCCAGGG | - | sulfite exporter TauE/SafE family protein |
| AVAEIV_RS16275 | 3546655 | 3546667 | 164 | - | 12.1571 | 3.04E-05 | 0.0753 | CGGAACGCCGCCA | - | hypothetical protein |
| AVAEIV_RS16280 | 3546655 | 3546667 | -47 | - | 12.1571 | 3.04E-05 | 0.0753 | CGGAACGCCGCCA | - | DNA-binding protein |
| AVAEIV_RS16290 | 3546655 | 3546667 | 333 | - | 12.1571 | 3.04E-05 | 0.0696 | CGGAACGCCGCCA | - | IS5 family transposase |
| AVAEIV_RS24985 | 5317014 | 5317026 | 336 | - | 12.1571 | 3.04E-05 | 0.0696 | CGGATCGCCGCCG | *iolB* | 5-deoxy-glucuronate isomerase |
| AVAEIV_RS25170 | 5359939 | 5359951 | 225 | + | 12.1571 | 3.04E-05 | 0.0753 | CAGATGGCCAGGG | - | chaperone |
| AVAEIV_RS26015 | 5590137 | 5590149 | 120 | - | 12.1571 | 3.04E-05 | 0.0753 | CAGTTTGCCAGGA | - | 2OG-Fe(II) oxygenase |
| AVAEIV_RS01195 | 233343 | 233355 | -33 | - | 12.1429 | 3.07E-05 | 0.0753 | CGGCTCGCCGTCG | - | GTP-binding protein |
| AVAEIV_RS01795 | 354060 | 354072 | 252 | - | 12.1429 | 3.07E-05 | 0.0696 | CGAATCGCCGTCG | - | ketopantoate reductase family protein |
| AVAEIV_RS03190 | 648056 | 648068 | 367 | - | 12.1429 | 3.07E-05 | 0.0696 | CTGATCGCCATTA | *rplR* | 50S ribosomal protein L18 |
| AVAEIV_RS03715 | 748478 | 748490 | 180 | - | 12.1429 | 3.07E-05 | 0.0696 | CGCACGGCCAGCA | *hflK* | FtsH protease activity modulator HflK |
| AVAEIV_RS09640 | 2013826 | 2013838 | 77 | - | 12.1429 | 3.07E-05 | 0.0753 | CAGTACGGCAGCA | - | AAA family ATPase |
| AVAEIV_RS15200 | 3310560 | 3310572 | 327 | - | 12.1429 | 3.07E-05 | 0.0753 | CAGTACGGCAGCA | - | hypothetical protein |
| AVAEIV_RS17565 | 3807418 | 3807430 | 286 | + | 12.1429 | 3.07E-05 | 0.0753 | CGGCTCGCCGTCG | - | DUF2784 domain-containing protein |
| AVAEIV_RS19265 | 4143700 | 4143712 | 286 | + | 12.1429 | 3.07E-05 | 0.0753 | CGCCACGCCAGCG | - | DUF1799 domain-containing protein |
| AVAEIV_RS21055 | 4477428 | 4477440 | 217 | + | 12.1429 | 3.07E-05 | 0.0753 | CCGACCGCCACCG | - | CDP-6-deoxy-delta-3,4-glucoseen reductase |
| AVAEIV_RS22465 | 4804999 | 4805011 | 189 | + | 12.1429 | 3.07E-05 | 0.0696 | CACATCGACATCG | *selO* | protein adenylyltransferase SelO |
| AVAEIV_RS24025 | 5113981 | 5113993 | 167 | - | 12.1429 | 3.07E-05 | 0.0696 | CGCACGGCCAGCA | *dapF* | diaminopimelate epimerase |
| AVAEIV_RS24450 | 5198030 | 5198042 | 274 | - | 12.1429 | 3.07E-05 | 0.0696 | CCGACCGCCACCG | - | D-hexose-6-phosphate mutarotase |
| AVAEIV_RS25165 | 5359779 | 5359791 | 316 | + | 12.1429 | 3.07E-05 | 0.0753 | CAGACCGCCGTCG | *hypD* | hydrogenase formation protein HypD |
| AVAEIV_RS25170 | 5359779 | 5359791 | 65 | + | 12.1429 | 3.07E-05 | 0.0753 | CAGACCGCCGTCG | - | chaperone |
| AVAEIV_RS00750 | 148149 | 148161 | 294 | - | 12.1286 | 3.11E-05 | 0.0698 | CTGCTGGCCAGCG | - | DUF364 domain-containing protein |
| AVAEIV_RS01465 | 283929 | 283941 | -46 | - | 12.1286 | 3.11E-05 | 0.0753 | CACAATGCCGGCA | *rdgB* | pyrophosphatase |
| AVAEIV_RS02780 | 569768 | 569780 | 143 | + | 12.1286 | 3.11E-05 | 0.0698 | CTGCTGGCCAGCG | *epd* | erythrose-4-phosphate dehydrogenase |
| AVAEIV_RS04585 | 938455 | 938467 | 209 | - | 12.1286 | 3.11E-05 | 0.0753 | CTGCTGGCCAGCG | - | urease subunit gamma |
| AVAEIV_RS04900 | 999690 | 999702 | 310 | - | 12.1286 | 3.11E-05 | 0.0753 | GAAAACGCCATCA | - | response regulator |
| AVAEIV_RS05080 | 1046314 | 1046326 | 55 | + | 12.1286 | 3.11E-05 | 0.0753 | CACATTGCCGGCG | - | nucleotide sugar dehydrogenase |
| AVAEIV_RS06275 | 1279643 | 1279655 | 367 | - | 12.1286 | 3.11E-05 | 0.0753 | CCCAGCGCCGTCA | - | hypothetical protein |
| AVAEIV_RS07445 | 1548792 | 1548804 | 376 | + | 12.1286 | 3.11E-05 | 0.0698 | CTGCTGGCCAGCG | - | efflux RND transporter permease subunit |
| AVAEIV_RS09205 | 1920923 | 1920935 | 168 | + | 12.1286 | 3.11E-05 | 0.0698 | CCGCCCGCCATCG | - | domain-containing protein |
| AVAEIV_RS10265 | 2142171 | 2142183 | -49 | + | 12.1286 | 3.11E-05 | 0.0698 | CTGCTGGCCAGCG | - | ComEA family DNA-binding protein |
| AVAEIV_RS13285 | 2877289 | 2877301 | 363 | + | 12.1286 | 3.11E-05 | 0.0698 | CTGCTGGCCAGCG | *ltrA* | group II intron reverse transcriptase/maturase |

| | | | | | | | | | | |
|---|---|---|---|---|---|---|---|---|---|---|
| AVAEIV_RS13775 | 2981512 | 2981524 | 367 | - | 12.1286 | 3.11E-05 | 0.0753 | CTGCTGGCCAGCG | - | DUF4150 domain-containing protein |
| AVAEIV_RS19415 | 4160627 | 4160639 | -10 | - | 12.1286 | 3.11E-05 | 0.0753 | CAGATGGCCAGTG | - | recombination protein NinB |
| AVAEIV_RS19895 | 4240236 | 4240248 | 305 | - | 12.1286 | 3.11E-05 | 0.0753 | CCCATCGCCACCC | *tadA* | tRNA adenosine(34) deaminase TadA |
| AVAEIV_RS21975 | 4687467 | 4687479 | 325 | + | 12.1286 | 3.11E-05 | 0.0698 | CTGCTGGCCAGCG | *fabI* | enoyl-ACP reductase FabI |
| AVAEIV_RS24730 | 5254632 | 5254644 | 356 | + | 12.1286 | 3.11E-05 | 0.0753 | CGCAGGGCCAGCG | - | substrate-binding domain-containing protein |
| AVAEIV_RS26100 | 5612369 | 5612381 | 13 | + | 12.1286 | 3.11E-05 | 0.0753 | CTGCTGGCCAGCG | - | tyrosine-type recombinase/integrase |
| AVAEIV_RS01315 | 256344 | 256356 | 318 | + | 12.1143 | 3.15E-05 | 0.0701 | CGAATGGCCATCC | *moeB* | molybdopterin-synthase adenylyltransferase MoeB |
| AVAEIV_RS01665 | 323072 | 323084 | 11 | - | 12.1143 | 3.15E-05 | 0.0753 | CAGCCCGCCACCG | - | DUF1615 domain-containing protein |
| AVAEIV_RS02360 | 469589 | 469601 | 113 | + | 12.1143 | 3.15E-05 | 0.0701 | CCATGGGCCAGCA | *grxC* | glutaredoxin 3 |
| AVAEIV_RS09130 | 1902504 | 1902516 | 265 | - | 12.1143 | 3.15E-05 | 0.0753 | CGGCTGGCCATCC | - | lipopolysaccharide kinase InaA family protein |
| AVAEIV_RS10495 | 2196058 | 2196070 | 80 | + | 12.1143 | 3.15E-05 | 0.0753 | CATCTTGCCATCG | - | AAA domain-containing protein |
| AVAEIV_RS10500 | 2196058 | 2196070 | 217 | + | 12.1143 | 3.15E-05 | 0.0701 | CATCTTGCCATCG | - | hypothetical protein |
| AVAEIV_RS10780 | 2259375 | 2259387 | 314 | - | 12.1143 | 3.15E-05 | 0.0753 | CTGGACGCCAGCA | - | Card1-like endonuclease domain-containing protein |
| AVAEIV_RS12725 | 2702561 | 2702573 | 370 | - | 12.1143 | 3.15E-05 | 0.0753 | CCCGTGGCCATCG | - | DUF3325 domain-containing protein |
| AVAEIV_RS17165 | 3721275 | 3721287 | 327 | - | 12.1143 | 3.15E-05 | 0.0753 | CAATTGGCCAGCC | - | CvpA family protein |
| AVAEIV_RS20470 | 4352892 | 4352904 | 195 | - | 12.1143 | 3.15E-05 | 0.0753 | CGGCTGGCCATCC | - | FecR domain-containing protein |
| AVAEIV_RS21105 | 4487668 | 4487680 | 183 | + | 12.1143 | 3.15E-05 | 0.0753 | CACAGCGCCGCCA | - | non-heme iron oxygenase ferredoxin subunit |
| AVAEIV_RS23490 | 5006370 | 5006382 | 45 | - | 12.1143 | 3.15E-05 | 0.0753 | CGGTTCGCCGGCG | *trpE* | anthranilate synthase component I |
| AVAEIV_RS24075 | 5121369 | 5121381 | 180 | + | 12.1143 | 3.15E-05 | 0.0701 | CAGAGCGCCTTCA | - | YifB family Mg chelatase-like AAA ATPase |
| AVAEIV_RS24255 | 5157727 | 5157739 | -37 | - | 12.1143 | 3.15E-05 | 0.0753 | CCCGTGGCCATCG | *lgt* | prolipoprotein diacylglyceryl transferase |
| AVAEIV_RS04385 | 892797 | 892809 | 196 | + | 12.1 | 3.18E-05 | 0.0753 | CGAAGGGCCACCA | - | IS481-like element ISAzvi11 family transposase |
| AVAEIV_RS04415 | 897237 | 897249 | 163 | - | 12.1 | 3.18E-05 | 0.0705 | CAACCCGCCATCG | - | amidohydrolase family protein |
| AVAEIV_RS05800 | 1188000 | 1188012 | 372 | + | 12.1 | 3.18E-05 | 0.0705 | CCGATCGCCACGA | - | acyl-CoA dehydrogenase family protein |
| AVAEIV_RS09400 | 1958742 | 1958754 | 220 | - | 12.1 | 3.18E-05 | 0.0705 | GAGTACGCCAGCA | - | IS110-like element ISAzvi3 family transposase |
| AVAEIV_RS14885 | 3238698 | 3238710 | 133 | - | 12.1 | 3.18E-05 | 0.0753 | GGGATCGCCATCG | - | efflux transporter outer membrane subunit |
| AVAEIV_RS00475 | 98554 | 98566 | 105 | - | 12.0857 | 3.22E-05 | 0.0753 | CCGCTCGCCATGA | - | hypothetical protein |
| AVAEIV_RS00945 | 187767 | 187779 | 192 | - | 12.0857 | 3.22E-05 | 0.0707 | AAGATCGCCAGCA | - | I78 family peptidase inhibitor |
| AVAEIV_RS01725 | 338383 | 338395 | 234 | + | 12.0857 | 3.22E-05 | 0.0753 | CCCTTCGCCAGCC | *ahcY* | adenosylhomocysteinase |
| AVAEIV_RS01730 | 338383 | 338395 | 15 | + | 12.0857 | 3.22E-05 | 0.0707 | CCCTTCGCCAGCC | - | c-type cytochrome |
| AVAEIV_RS06665 | 1377617 | 1377629 | 66 | + | 12.0857 | 3.22E-05 | 0.0753 | CAGTTGGCGATCA | *ptsN* | PTS IIA-like nitrogen regulatory protein PtsN |
| AVAEIV_RS09640 | 2013837 | 2013849 | 88 | - | 12.0857 | 3.22E-05 | 0.0753 | CGACGGGCCATCA | - | AAA family ATPase |
| AVAEIV_RS12340 | 2609041 | 2609053 | 345 | - | 12.0857 | 3.22E-05 | 0.0707 | CCGAAGGCCGGCA | *motB* | flagellar motor protein MotB |
| AVAEIV_RS14020 | 3043559 | 3043571 | 81 | + | 12.0857 | 3.22E-05 | 0.0707 | CCGAAGGCCGGCA | - | hypothetical protein |
| AVAEIV_RS15975 | 3482045 | 3482057 | 48 | - | 12.0857 | 3.22E-05 | 0.0707 | CGGTTGGCCAGCC | - | acyl-CoA dehydrogenase family protein |
| AVAEIV_RS18690 | 4025929 | 4025941 | 24 | - | 12.0857 | 3.22E-05 | 0.0753 | CGTATTGCCATCG | - | 3-carboxy-cis,cis-muconate cycloisomerase |
| AVAEIV_RS18895 | 4067489 | 4067501 | 143 | + | 12.0857 | 3.22E-05 | 0.0707 | CATTTTGCCAGCG | *mutS* | DNA mismatch repair protein MutS |
| AVAEIV_RS22205 | 4739157 | 4739169 | 13 | - | 12.0857 | 3.22E-05 | 0.0753 | CCCTTCGCCAGCC | - | ABC transporter ATP-binding protein |
| AVAEIV_RS01840 | 364967 | 364979 | 189 | + | 12.0714 | 3.25E-05 | 0.0709 | CGCATCGCCACCC | *fdxH* | formate dehydrogenase subunit beta |
| AVAEIV_RS13780 | 2982157 | 2982169 | 365 | - | 12.0714 | 3.25E-05 | 0.0753 | CGGATTGCCGCCA | - | DUF3540 domain-containing protein |
| AVAEIV_RS15530 | 3397079 | 3397091 | 236 | + | 12.0714 | 3.25E-05 | 0.0709 | CGCATCGCCACCC | *codA* | cytosine deaminase |
| AVAEIV_RS16605 | 3600720 | 3600732 | 109 | + | 12.0714 | 3.25E-05 | 0.0753 | CTCCTCGCCAGCG | - | class II glutamine amidotransferase |
| AVAEIV_RS19825 | 4224753 | 4224765 | 24 | + | 12.0714 | 3.25E-05 | 0.0753 | CGGCCCGCCATCG | *rplS* | 50S ribosomal protein L19 |
| AVAEIV_RS20475 | 4353488 | 4353500 | 286 | - | 12.0714 | 3.25E-05 | 0.0753 | CGCATCGCCACCC | - | sigma-70 family RNA polymerase sigma factor |
| AVAEIV_RS23155 | 4946146 | 4946158 | 380 | - | 12.0714 | 3.25E-05 | 0.0709 | CGGCCCGCCATCG | - | hypothetical protein |
| AVAEIV_RS24470 | 5201026 | 5201038 | 263 | + | 12.0714 | 3.25E-05 | 0.0709 | CGCATCGCCACCC | - | class I SAM-dependent methyltransferase |
| AVAEIV_RS25895 | 5556204 | 5556216 | 96 | - | 12.0714 | 3.25E-05 | 0.0753 | CGCATCGCCACCC | - | ABC transporter permease |
| AVAEIV_RS03245 | 657858 | 657870 | 380 | - | 12.0571 | 3.29E-05 | 0.0753 | CAGCAGGCCGGCA | *uvrA* | excinuclease ABC subunit UvrA |
| AVAEIV_RS09635 | 2013048 | 2013060 | 259 | + | 12.0571 | 3.29E-05 | 0.0753 | CGCGAGGCCATCA | - | DUF58 domain-containing protein |
| AVAEIV_RS11915 | 2530251 | 2530263 | 162 | + | 12.0571 | 3.29E-05 | 0.0753 | CGGCTTGCCGTCA | - | protein |
| AVAEIV_RS19020 | 4096589 | 4096601 | -45 | + | 12.0571 | 3.29E-05 | 0.0753 | CAGAAGGGCAGCA | - | OmpH family outer membrane protein |
| AVAEIV_RS19750 | 4208687 | 4208699 | -38 | + | 12.0571 | 3.29E-05 | 0.0753 | CAGAAGGGCAGCA | - | hypothetical protein |
| AVAEIV_RS20540 | 4366855 | 4366867 | 245 | - | 12.0571 | 3.29E-05 | 0.0753 | CGCGTGGCCATCG | - | FKBP-type peptidyl-prolyl cis-trans isomerase |

| | | | | | | | | | | |
|---|---|---|---|---|---|---|---|---|---|---|
| AVAEIV_RS20545 | 4366855 | 4366867 | -11 | - | 12.0571 | 3.29E-05 | 0.0712 | CGCGTGGCCATCG | - | acetate kinase |
| AVAEIV_RS00475 | 98692 | 98704 | 243 | - | 12.0429 | 3.32E-05 | 0.0753 | CAGGTCGCCGGCG | - | hypothetical protein |
| AVAEIV_RS04555 | 932676 | 932688 | -43 | - | 12.0429 | 3.32E-05 | 0.0753 | CTGTTCGCCGGCA | - | TorF family putative porin |
| AVAEIV_RS04560 | 932676 | 932688 | 269 | - | 12.0429 | 3.32E-05 | 0.0712 | CTGTTCGCCGGCA | - | DMT family transporter |
| AVAEIV_RS08200 | 1704598 | 1704610 | 154 | + | 12.0429 | 3.32E-05 | 0.0712 | CGCTATGCCATCG | *rfbD* | dTDP-4-dehydrorhamnose reductase |
| AVAEIV_RS10005 | 2095326 | 2095338 | -49 | + | 12.0429 | 3.32E-05 | 0.0712 | CTGTTCGCCGGCA | - | monovalent cation/H+ antiporter subunit D |
| AVAEIV_RS10160 | 2120893 | 2120905 | 47 | + | 12.0429 | 3.32E-05 | 0.0753 | CCACAGGCCAGCA | - | cyd operon YbgE family protein |
| AVAEIV_RS10165 | 2120893 | 2120905 | -44 | + | 12.0429 | 3.32E-05 | 0.0753 | CCACAGGCCAGCA | *cydX* | cytochrome bd-I oxidase subunit CydX |
| AVAEIV_RS11135 | 2348681 | 2348693 | 345 | + | 12.0429 | 3.32E-05 | 0.0712 | CTGTTCGCCGGCA | - | ABC transporter ATP-binding protein |
| AVAEIV_RS13220 | 2866906 | 2866918 | 262 | + | 12.0429 | 3.32E-05 | 0.0712 | CTGTTCGCCGGCA | - | alpha/beta fold hydrolase |
| AVAEIV_RS16005 | 3488193 | 3488205 | 360 | - | 12.0429 | 3.32E-05 | 0.0712 | CGCTATGCCATCG | - | hypothetical protein |
| AVAEIV_RS22605 | 4841971 | 4841983 | 246 | - | 12.0429 | 3.32E-05 | 0.0712 | CAGGTCGCCGGCG | - | containing protein |
| AVAEIV_RS23055 | 4933153 | 4933165 | 344 | - | 12.0429 | 3.32E-05 | 0.0712 | CAGGTCGCCGGCG | *gpM* | phage terminase small subunit |
| AVAEIV_RS23310 | 4970616 | 4970628 | 138 | - | 12.0429 | 3.32E-05 | 0.0753 | CAGGTCGCCGGCG | - | nicotinate phosphoribosyltransferase |
| AVAEIV_RS23315 | 4970616 | 4970628 | -36 | - | 12.0429 | 3.32E-05 | 0.0712 | CAGGTCGCCGGCG | - | CapA family protein |
| AVAEIV_RS23805 | 5073552 | 5073564 | 151 | + | 12.0429 | 3.32E-05 | 0.0753 | CAGGTCGCCGGCG | *gcvP* | aminomethyl-transferring glycine dehydrogenase |
| AVAEIV_RS25960 | 5579316 | 5579328 | 75 | + | 12.0429 | 3.32E-05 | 0.0753 | CAGGTCGCCGGCG | *atpA* | F0F1 ATP synthase subunit alpha |
| AVAEIV_RS00740 | 147303 | 147315 | 269 | - | 12.0286 | 3.36E-05 | 0.0712 | CGGATGGCCGGCG | - | hypothetical protein |
| AVAEIV_RS03870 | 779125 | 779137 | 52 | + | 12.0286 | 3.36E-05 | 0.0753 | CGGAAGGCCGGCA | - | TonB-dependent receptor domain-containing protein |
| AVAEIV_RS09155 | 1906700 | 1906712 | 298 | - | 12.0286 | 3.36E-05 | 0.0753 | CGCTTCGCCAGCC | - | ATP-dependent zinc protease |
| AVAEIV_RS15895 | 3468268 | 3468280 | 239 | - | 12.0286 | 3.36E-05 | 0.0753 | CCTATCGCCACCG | - | ABC transporter substrate-binding protein |
| AVAEIV_RS16505 | 3582021 | 3582033 | 345 | - | 12.0286 | 3.36E-05 | 0.0753 | CAATTCGCCAGGA | - | site-specific integrase |
| AVAEIV_RS16570 | 3593901 | 3593913 | 366 | + | 12.0286 | 3.36E-05 | 0.0712 | CGACTTGCCACCA | - | YecA family protein |
| AVAEIV_RS23420 | 4992692 | 4992704 | 159 | - | 12.0286 | 3.36E-05 | 0.0753 | CCGCGTGCCATCG | - | LysR family transcriptional regulator |
| AVAEIV_RS23425 | 4992692 | 4992704 | 268 | - | 12.0286 | 3.36E-05 | 0.0712 | CCGCGTGCCATCG | - | alginate lyase family protein |
| AVAEIV_RS23760 | 5064834 | 5064846 | 289 | - | 12.0286 | 3.36E-05 | 0.0753 | CCCATCGCCGGCG | - | LysR family transcriptional regulator |
| AVAEIV_RS02800 | 575193 | 575205 | 204 | - | 12.0143 | 3.39E-05 | 0.0753 | CCGACGGCCAGCG | - | oxidoreductase |
| AVAEIV_RS03810 | 768202 | 768214 | 181 | + | 12.0143 | 3.39E-05 | 0.0712 | CAGGTGGCCAGCC | - | HAD family hydrolase |
| AVAEIV_RS08095 | 1678362 | 1678374 | 213 | - | 12.0143 | 3.39E-05 | 0.0712 | CCGGGGGCCAGCA | - | bifunctional 4-hydroxy-2-oxoglutarate aldolase/2-dehydro-3-deoxy-phosphogluconate aldolase |
| AVAEIV_RS08910 | 1852126 | 1852138 | 180 | - | 12.0143 | 3.39E-05 | 0.0753 | CACATAGCCACCA | - | glycosyltransferase family 4 protein |
| AVAEIV_RS20470 | 4352915 | 4352927 | 218 | + | 12.0143 | 3.39E-05 | 0.0753 | GAGATGGCCAGCG | - | FecR domain-containing protein |
| AVAEIV_RS24495 | 5205477 | 5205489 | 178 | + | 12.0143 | 3.39E-05 | 0.0712 | CAGAGCGCCATGG | - | hypothetical protein |
| AVAEIV_RS24680 | 5244260 | 5244272 | 163 | + | 12.0143 | 3.39E-05 | 0.0712 | CCGACGGCCAGCG | - | OsmC family protein |
| AVAEIV_RS25275 | 5384742 | 5384754 | 313 | + | 12.0143 | 3.39E-05 | 0.0753 | CCGATGGCCAGCT | *modA* | molybdate ABC transporter substrate-binding protein |
| AVAEIV_RS01580 | 303223 | 303235 | 134 | - | 12 | 3.43E-05 | 0.0712 | CACCTCGCCGGCG | - | YceI family protein |
| AVAEIV_RS02645 | 538102 | 538114 | 307 | + | 12 | 3.43E-05 | 0.0712 | CACATCGGCAGCG | - | DUF6491 family protein |
| AVAEIV_RS04650 | 952214 | 952226 | 29 | - | 12 | 3.43E-05 | 0.0753 | CTGCCCGCCATCA | - | histone deacetylase |
| AVAEIV_RS05525 | 1143176 | 1143188 | 24 | + | 12 | 3.43E-05 | 0.0712 | CAGTATGCCAGCC | *clpB* | ATP-dependent chaperone ClpB |
| AVAEIV_RS07875 | 1637688 | 1637700 | 215 | - | 12 | 3.43E-05 | 0.0712 | CACCACGCCGGCA | *earP* | rhamnosyltransferase EarP |
| AVAEIV_RS11360 | 2403000 | 2403012 | 191 | + | 12 | 3.43E-05 | 0.0753 | CACCTCGCCGGCG | *guaD* | guanine deaminase |
| AVAEIV_RS11560 | 2444752 | 2444764 | 136 | + | 12 | 3.43E-05 | 0.0712 | CACAACGGCAGCA | - | FeoA family protein |
| AVAEIV_RS11565 | 2444752 | 2444764 | 360 | + | 12 | 3.43E-05 | 0.0712 | CACAACGGCAGCA | *feoB* | Fe(2+) transporter permease subunit FeoB |
| AVAEIV_RS12690 | 2694941 | 2694953 | 104 | - | 12 | 3.43E-05 | 0.0753 | CCGCTCGCCGCCA | - | 3-hydroxybutyryl-CoA dehydrogenase |
| AVAEIV_RS12690 | 2694916 | 2694928 | 79 | + | 12 | 3.43E-05 | 0.0753 | CCGCTCGCCGCCA | - | 3-hydroxybutyryl-CoA dehydrogenase |
| AVAEIV_RS17450 | 3780607 | 3780619 | 352 | - | 12 | 3.43E-05 | 0.0753 | CACCTCGCCGGCG | *hppD* | 4-hydroxyphenylpyruvate dioxygenase |
| AVAEIV_RS17880 | 3871412 | 3871424 | -35 | - | 12 | 3.43E-05 | 0.0712 | CAGCTCGCTACCA | - | protein |
| AVAEIV_RS18600 | 4005207 | 4005219 | 363 | + | 12 | 3.43E-05 | 0.0712 | CCGCTCGCCGCCA | - | MFS transporter |
| AVAEIV_RS20600 | 4383651 | 4383663 | 276 | - | 12 | 3.43E-05 | 0.0753 | CACCTCGCCGGCG | - | DUF1302 domain-containing protein |
| AVAEIV_RS21555 | 4585139 | 4585151 | 365 | + | 12 | 3.43E-05 | 0.0712 | CAGATCGGCGTCA | - | (Fe-S)-binding protein |
| AVAEIV_RS23805 | 5073654 | 5073666 | 253 | + | 12 | 3.43E-05 | 0.0753 | CACCTCGCCGGCG | *gcvP* | aminomethyl-transferring glycine dehydrogenase |
| AVAEIV_RS24425 | 5192636 | 5192648 | 347 | + | 12 | 3.43E-05 | 0.0753 | CACCTCGCCGGCG | *pstB* | phosphate ABC transporter ATP-binding protein PstB |

| | | | | | | | | | | |
|---|---|---|---|---|---|---|---|---|---|---|
| AVAEIV_RS00065 | 13198 | 13210 | 225 | - | 11.9857 | 3.46E-05 | 0.0712 | CTGCGCGCCATCG | - | DNA-3-methyladenine glycosylase I |
| AVAEIV_RS01685 | 328315 | 328327 | 350 | - | 11.9857 | 3.46E-05 | 0.0753 | CAGTTGGCCAACA | - | YfhL family 4Fe-4S dicluster ferredoxin |
| AVAEIV_RS02245 | 449967 | 449979 | 165 | + | 11.9857 | 3.46E-05 | 0.0753 | CAGCCGGCCAGCG | - | hypothetical protein |
| AVAEIV_RS02745 | 561544 | 561556 | 48 | + | 11.9857 | 3.46E-05 | 0.0753 | CTGCGCGCCATCG | *yrfG* | GMP/IMP nucleotidase |
| AVAEIV_RS02750 | 561544 | 561556 | 97 | + | 11.9857 | 3.46E-05 | 0.0712 | CTGCGCGCCATCG | *nudE* | ADP compounds hydrolase NudE |
| AVAEIV_RS03200 | 649081 | 649093 | 199 | + | 11.9857 | 3.46E-05 | 0.0712 | CAGCCGGCCAGCG | *rpmD* | 50S ribosomal protein L30 |
| AVAEIV_RS03205 | 649081 | 649093 | 379 | + | 11.9857 | 3.46E-05 | 0.0712 | CAGCCGGCCAGCG | *rplO* | 50S ribosomal protein L15 |
| AVAEIV_RS03465 | 699029 | 699041 | 220 | + | 11.9857 | 3.46E-05 | 0.0712 | CTGCGCGCCATCG | - | NUDIX hydrolase |
| AVAEIV_RS03635 | 731338 | 731350 | 334 | - | 11.9857 | 3.46E-05 | 0.0753 | CCCTACGCCACCG | - | hypothetical protein |
| AVAEIV_RS09620 | 2010532 | 2010544 | 163 | - | 11.9857 | 3.46E-05 | 0.0753 | CTGCGCGCCATCG | - | VWA domain-containing protein |
| AVAEIV_RS09620 | 2010592 | 2010604 | 223 | - | 11.9857 | 3.46E-05 | 0.0753 | CTGCGCGCCATCG | - | VWA domain-containing protein |
| AVAEIV_RS11315 | 2389926 | 2389938 | 252 | - | 11.9857 | 3.46E-05 | 0.0712 | CAGCCGGCCAGCG | *tkt* | transketolase |
| AVAEIV_RS12705 | 2699073 | 2699085 | 324 | - | 11.9857 | 3.46E-05 | 0.0753 | CAGCCGGCCAGCG | - | acyl-CoA synthetase |
| AVAEIV_RS14115 | 3071472 | 3071484 | 252 | + | 11.9857 | 3.46E-05 | 0.0753 | CAGCCGGCCAGCG | *tkt* | transketolase |
| AVAEIV_RS14400 | 3135921 | 3135933 | 328 | - | 11.9857 | 3.46E-05 | 0.0753 | CACTACGCCGTCG | *flhA* | flagellar biosynthesis protein FlhA |
| AVAEIV_RS17475 | 3782614 | 3782626 | 63 | + | 11.9857 | 3.46E-05 | 0.0712 | CGACTGGCCAGCG | - | PHB depolymerase family esterase |
| AVAEIV_RS19050 | 4104290 | 4104302 | 267 | - | 11.9857 | 3.46E-05 | 0.0753 | CTGCGCGCCATCG | *frr* | ribosome recycling factor |
| AVAEIV_RS00305 | 68739 | 68751 | 228 | + | 11.9714 | 3.50E-05 | 0.0753 | CTCTATGCCATCA | *polA* | DNA polymerase I |
| AVAEIV_RS00315 | 68739 | 68751 | 138 | + | 11.9714 | 3.50E-05 | 0.0712 | CTCTATGCCATCA | - | homoserine kinase |
| AVAEIV_RS04635 | 947373 | 947385 | 260 | + | 11.9714 | 3.50E-05 | 0.0712 | CGCATCGCCGGCG | - | SDR family oxidoreductase |
| AVAEIV_RS05805 | 1189238 | 1189250 | 288 | - | 11.9714 | 3.50E-05 | 0.0712 | CTGTCCGCCAGCA | - | class I adenylate-forming enzyme family protein |
| AVAEIV_RS07005 | 1444064 | 1444076 | 313 | + | 11.9714 | 3.50E-05 | 0.0712 | CAGGCCGCCAGCG | - | YajQ family cyclic di-GMP-binding protein |
| AVAEIV_RS07255 | 1505432 | 1505444 | 303 | - | 11.9714 | 3.50E-05 | 0.0753 | CGCAACGCCGGCA | *glpD* | glycerol-3-phosphate dehydrogenase |
| AVAEIV_RS08350 | 1744277 | 1744289 | 350 | + | 11.9714 | 3.50E-05 | 0.0712 | CTCAAGGCCATCG | - | family antitoxin |
| AVAEIV_RS09020 | 1877269 | 1877281 | 182 | + | 11.9714 | 3.50E-05 | 0.0753 | CACGATGCCATCG | - | PhnD/SsuA/transferrin family substrate-binding protein |
| AVAEIV_RS09025 | 1877269 | 1877281 | -22 | + | 11.9714 | 3.50E-05 | 0.0712 | CACGATGCCATCG | - | FCD domain-containing protein |
| AVAEIV_RS11020 | 2320509 | 2320521 | 302 | + | 11.9714 | 3.50E-05 | 0.0712 | CGCATCGCCGGCG | - | FAD-dependent oxidoreductase |
| AVAEIV_RS11420 | 2417327 | 2417339 | 336 | - | 11.9714 | 3.50E-05 | 0.0753 | CGCATCGCCGGCG | - | TauD/TfdA dioxygenase family protein |
| AVAEIV_RS12360 | 2614717 | 2614729 | 227 | + | 11.9714 | 3.50E-05 | 0.0712 | CGCAACGCCGGCA | - | methyl-accepting chemotaxis protein |
| AVAEIV_RS15625 | 3422575 | 3422587 | 75 | + | 11.9714 | 3.50E-05 | 0.0753 | CAGGCCGCCAGCG | - | DsbE family thiol:disulfide interchange protein |
| AVAEIV_RS19175 | 4132925 | 4132937 | 161 | + | 11.9714 | 3.50E-05 | 0.0753 | CAACTCGCCGCCA | - | hypothetical protein |
| AVAEIV_RS19390 | 4158456 | 4158468 | 54 | + | 11.9714 | 3.50E-05 | 0.0753 | CGGATCGCTACCA | - | antiterminator Q family protein |
| AVAEIV_RS21830 | 4651658 | 4651670 | 309 | - | 11.9714 | 3.50E-05 | 0.0753 | CTCAAGGCCATCG | - | ABC transporter substrate-binding protein |
| AVAEIV_RS24135 | 5132528 | 5132540 | 203 | - | 11.9714 | 3.50E-05 | 0.0712 | CCCTCGGCCATCA | - | xanthine phosphoribosyltransferase |
| AVAEIV_RS25680 | 5493543 | 5493555 | 247 | + | 11.9714 | 3.50E-05 | 0.0712 | CAGGCCGCCAGCG | - | efflux transporter outer membrane subunit |
| AVAEIV_RS25715 | 5502220 | 5502232 | 226 | - | 11.9714 | 3.50E-05 | 0.0753 | CAGGCCGCCAGCG | *lpxO* | lipid A hydroxylase LpxO |
| AVAEIV_RS00070 | 14114 | 14126 | 113 | - | 11.9571 | 3.54E-05 | 0.0712 | GCGATCGCCACCA | - | bifunctional metallophosphatase/5'-nucleotidase |
| AVAEIV_RS00490 | 100776 | 100788 | 295 | - | 11.9571 | 3.54E-05 | 0.0753 | CCCGGCGCCAGCA | - | cytochrome c oxidase subunit 3 |
| AVAEIV_RS00645 | 131566 | 131578 | 346 | - | 11.9571 | 3.54E-05 | 0.0753 | GACATCGCCAGCG | - | MFS transporter |
| AVAEIV_RS03935 | 793438 | 793450 | 141 | + | 11.9571 | 3.54E-05 | 0.0712 | CAACTCGGCATCA | *mpl* | UDP-N-acetylmuramate:L-alanyl-gamma-D-glutamyl-meso-diaminopimelate ligase |
| AVAEIV_RS06240 | 1276227 | 1276239 | 184 | - | 11.9571 | 3.54E-05 | 0.0753 | CGAATCGCTATCA | - | ParB family protein |
| AVAEIV_RS06525 | 1346049 | 1346061 | 60 | - | 11.9571 | 3.54E-05 | 0.0712 | CCCACCGCCAGCG | - | biotin-dependent carboxyltransferase family protein |
| AVAEIV_RS07130 | 1469174 | 1469186 | 86 | - | 11.9571 | 3.54E-05 | 0.0712 | CATATCGCCATCT | - | SoxR reducing system RseC family protein |
| AVAEIV_RS08840 | 1833885 | 1833897 | 294 | - | 11.9571 | 3.54E-05 | 0.0712 | CCCGGCGCCAGCA | - | hypothetical protein |
| AVAEIV_RS11030 | 2323340 | 2323352 | 63 | + | 11.9571 | 3.54E-05 | 0.0712 | CGGACGGCCAGCG | - | LLM class flavin-dependent oxidoreductase |
| AVAEIV_RS12625 | 2679597 | 2679609 | 340 | - | 11.9571 | 3.54E-05 | 0.0753 | GACATCGCCAGCG | - | NAD-dependent epimerase/dehydratase family protein |
| AVAEIV_RS12630 | 2679597 | 2679609 | -49 | - | 11.9571 | 3.54E-05 | 0.0753 | GACATCGCCAGCG | - | hypothetical protein |
| AVAEIV_RS12940 | 2804250 | 2804262 | 322 | + | 11.9571 | 3.54E-05 | 0.0712 | CCCATCGCCAGCT | - | BCAM0308 family protein |
| AVAEIV_RS13050 | 2827445 | 2827457 | 182 | + | 11.9571 | 3.54E-05 | 0.0753 | CGGTCTGCCAGCA | - | arsenate reductase ArsC |
| AVAEIV_RS17535 | 3798416 | 3798428 | 289 | + | 11.9571 | 3.54E-05 | 0.0753 | CCCTGGGCCATCG | - | regulator |
| AVAEIV_RS20360 | 4331340 | 4331352 | 245 | - | 11.9571 | 3.54E-05 | 0.0712 | CAGGTTGCCGGCA | - | hypothetical protein |

| | | | | | | | | | | |
|---|---|---|---|---|---|---|---|---|---|---|
| AVAEIV_RS21080 | 4483460 | 4483472 | 166 | - | 11.9571 | 3.54E-05 | 0.0753 | GACATCGCCAGCG | - | 2-keto-4-pentenoate hydratase |
| AVAEIV_RS23295 | 4966394 | 4966406 | 150 | + | 11.9571 | 3.54E-05 | 0.0712 | CCCGGCGCCAGCA | - | ribose-phosphate diphosphokinase |
| AVAEIV_RS24065 | 5121055 | 5121067 | 164 | + | 11.9571 | 3.54E-05 | 0.0753 | CATACCGCCATCG | *glnK* | P-II family nitrogen regulator |
| AVAEIV_RS24070 | 5121055 | 5121067 | 187 | + | 11.9571 | 3.54E-05 | 0.0712 | CATACCGCCATCG | - | accessory factor UbiK family protein |
| AVAEIV_RS01640 | 318135 | 318147 | 258 | - | 11.9429 | 3.57E-05 | 0.0753 | CAGAATGCCGGCG | - | methyl-accepting chemotaxis protein |
| AVAEIV_RS04375 | 889809 | 889821 | 74 | - | 11.9429 | 3.57E-05 | 0.0753 | CCGAGCGCCGTCG | - | PhoH family protein |
| AVAEIV_RS10360 | 2166327 | 2166339 | 330 | + | 11.9429 | 3.57E-05 | 0.0753 | CAGACCGCCGCCA | - | helix-turn-helix domain-containing protein |
| AVAEIV_RS18345 | 3948347 | 3948359 | 259 | + | 11.9429 | 3.57E-05 | 0.0753 | TAGAAGGCCAGCA | *mazG* | nucleoside triphosphate pyrophosphohydrolase |
| AVAEIV_RS01805 | 357245 | 357257 | 361 | - | 11.9286 | 3.60E-05 | 0.0753 | CCGGAGGCCATCG | - | substrate binding protein |
| AVAEIV_RS09180 | 1912647 | 1912659 | -17 | - | 11.9286 | 3.60E-05 | 0.0717 | CCGGAGGCCATCG | - | YbcC family protein |
| AVAEIV_RS11540 | 2441405 | 2441417 | 56 | - | 11.9286 | 3.60E-05 | 0.0717 | CCGCCCGCCACCA | - | hypothetical protein |
| AVAEIV_RS12465 | 2643793 | 2643805 | -31 | + | 11.9286 | 3.60E-05 | 0.0717 | GAAATCGCCACCA | - | GNAT family N-acetyltransferase |
| AVAEIV_RS13105 | 2840926 | 2840938 | 386 | - | 11.9286 | 3.60E-05 | 0.0717 | CCGGAGGCCATCG | *gcvT* | glycine cleavage system aminomethyltransferase GcvT |
| AVAEIV_RS16660 | 3610798 | 3610810 | 298 | + | 11.9286 | 3.60E-05 | 0.0753 | CGCTACGCCACCG | - | hypothetical protein |
| AVAEIV_RS16670 | 3610798 | 3610810 | 204 | + | 11.9286 | 3.60E-05 | 0.0717 | CGCTACGCCACCG | - | four-helix bundle copper-binding protein |
| AVAEIV_RS20380 | 4334469 | 4334481 | 184 | + | 11.9286 | 3.60E-05 | 0.0717 | CCAACCGCCACCA | *rpsT* | 30S ribosomal protein S20 |
| AVAEIV_RS25735 | 5506320 | 5506332 | 304 | + | 11.9286 | 3.60E-05 | 0.0717 | CAGTTGGCCATGG | - | hypothetical protein |
| AVAEIV_RS12075 | 2563296 | 2563308 | 174 | + | 11.9143 | 3.63E-05 | 0.0717 | GCGTTCGCCAGCA | - | methylthioribulose 1-phosphate dehydratase |
| AVAEIV_RS13045 | 2826763 | 2826775 | 357 | - | 11.9143 | 3.63E-05 | 0.0753 | CACCTTGCCGGCA | - | YnfA family protein |
| AVAEIV_RS14885 | 3238576 | 3238588 | 11 | + | 11.9143 | 3.63E-05 | 0.0717 | CAGAGCGGCATCG | - | efflux transporter outer membrane subunit |
| AVAEIV_RS15395 | 3361271 | 3361283 | 296 | + | 11.9143 | 3.63E-05 | 0.0717 | CAGAGCGGCATCG | - | MATE family efflux transporter |
| AVAEIV_RS15410 | 3366304 | 3366316 | 158 | + | 11.9143 | 3.63E-05 | 0.0717 | CAGAGCGGCATCG | *tviB* | Vi polysaccharide biosynthesis UDP-N-acetylglucosamine C-6 dehydrogenase TviB |
| AVAEIV_RS16940 | 3670445 | 3670457 | 175 | - | 11.9143 | 3.63E-05 | 0.0753 | CAGAACGCCAGGG | - | GMC family oxidoreductase |
| AVAEIV_RS17530 | 3795494 | 3795506 | 78 | + | 11.9143 | 3.63E-05 | 0.0717 | CCCATCGCCAGGA | *mnmC* | thiouridine)(34)-methyltransferase MnmD/FAD-dependent 5-carboxymethylaminomethyl-2-thiouridine(34) oxidoreductase MnmC |
| AVAEIV_RS23850 | 5082293 | 5082305 | 227 | + | 11.9143 | 3.63E-05 | 0.0717 | CTACTGGCCAGCA | - | TIGR02449 family protein |
| AVAEIV_RS06530 | 1348217 | 1348229 | 102 | + | 11.9 | 3.67E-05 | 0.0753 | CGCGGCGCCAGCA | - | peptide ABC transporter ATP-binding protein |
| AVAEIV_RS11075 | 2334274 | 2334286 | 227 | - | 11.9 | 3.67E-05 | 0.0718 | CGCACCGCCAGCG | - | amino acid ABC transporter permease |
| AVAEIV_RS12665 | 2686975 | 2686987 | 119 | - | 11.9 | 3.67E-05 | 0.0718 | CCTTTTGCCAGCA | *dosP* | oxygen-sensing cyclic-di-GMP phosphodiesterase DosP |
| AVAEIV_RS14940 | 3257331 | 3257343 | 323 | - | 11.9 | 3.67E-05 | 0.0718 | CGCACCGCCAGCG | - | Arm DNA-binding domain-containing protein |
| AVAEIV_RS16405 | 3563225 | 3563237 | 349 | - | 11.9 | 3.67E-05 | 0.0753 | CCCAAGGCCACCG | - | IS630 family transposase |
| AVAEIV_RS20965 | 4457300 | 4457312 | -18 | + | 11.9 | 3.67E-05 | 0.0753 | CGCATCGACATCA | - | cation-transporting P-type ATPase |
| AVAEIV_RS21190 | 4505831 | 4505843 | 235 | - | 11.9 | 3.67E-05 | 0.0753 | CGCGGCGCCAGCA | *otsA* | forming) |
| AVAEIV_RS21425 | 4559964 | 4559976 | 151 | + | 11.9 | 3.67E-05 | 0.0753 | CGCATCGCCAGCT | *infB* | translation initiation factor IF-2 |
| AVAEIV_RS23945 | 5100902 | 5100914 | 228 | + | 11.9 | 3.67E-05 | 0.0718 | CAGTCCGCCGGCA | - | TIGR02444 family protein |
| AVAEIV_RS24685 | 5246074 | 5246086 | 222 | - | 11.9 | 3.67E-05 | 0.0753 | CGATTCGCCGGCA | - | aldo/keto reductase |
| AVAEIV_RS24695 | 5246074 | 5246086 | 360 | - | 11.9 | 3.67E-05 | 0.0718 | CGATTCGCCGGCA | - | cyclase/phosphodiesterase |
| AVAEIV_RS25270 | 5383705 | 5383717 | 56 | - | 11.9 | 3.67E-05 | 0.0753 | CCCAAGGCCACCG | *modB* | molybdate ABC transporter permease subunit |
| AVAEIV_RS01190 | 232628 | 232640 | -47 | - | 11.8857 | 3.70E-05 | 0.0753 | CGGAACGCCACCC | - | ATP-binding cassette domain-containing protein |
| AVAEIV_RS01495 | 288279 | 288291 | 232 | + | 11.8857 | 3.70E-05 | 0.0753 | GGAATCGCCATCA | *proC* | pyrroline-5-carboxylate reductase |
| AVAEIV_RS02340 | 466230 | 466242 | 75 | - | 11.8857 | 3.70E-05 | 0.0753 | CCATCCGCCAGCA | - | S41 family peptidase |
| AVAEIV_RS03075 | 627459 | 627471 | 304 | - | 11.8857 | 3.70E-05 | 0.0718 | CGGAGCGCCGTCG | *rpoB* | DNA-directed RNA polymerase subunit beta |
| AVAEIV_RS03650 | 735751 | 735763 | 200 | + | 11.8857 | 3.70E-05 | 0.0753 | CAGTGCGCCGGCG | - | NAD(P)-dependent alcohol dehydrogenase |
| AVAEIV_RS07180 | 1480448 | 1480460 | -11 | + | 11.8857 | 3.70E-05 | 0.0753 | CAGCGGGCCATCC | - | DUF2905 domain-containing protein |
| AVAEIV_RS07180 | 1480605 | 1480617 | 146 | - | 11.8857 | 3.70E-05 | 0.0753 | CAACGCGCCACCG | - | DUF2905 domain-containing protein |
| AVAEIV_RS09185 | 1914969 | 1914981 | 207 | + | 11.8857 | 3.70E-05 | 0.0718 | CCATCCGCCAGCA | - | proton-conducting transporter membrane subunit |
| AVAEIV_RS14430 | 3143590 | 3143602 | 204 | - | 11.8857 | 3.70E-05 | 0.0753 | CCGATCGCGAGCA | - | glutaminase |
| AVAEIV_RS17045 | 3698301 | 3698313 | 333 | - | 11.8857 | 3.70E-05 | 0.0753 | TCGATCGCCACCA | - | cupin |
| AVAEIV_RS18720 | 4031988 | 4032000 | 213 | - | 11.8857 | 3.70E-05 | 0.0753 | TCGATCGCCACCA | *pcaG* | protocatechuate 3,4-dioxygenase subunit alpha |
| AVAEIV_RS20240 | 4318940 | 4318952 | 231 | - | 11.8857 | 3.70E-05 | 0.0753 | CAGCGGGCCATCC | *secF* | protein translocase subunit SecF |

| | | | | | | | | | | |
|---|---|---|---|---|---|---|---|---|---|---|
| AVAEIV_RS20935 | 4452177 | 4452189 | 342 | + | 11.8857 | 3.70E-05 | 0.0753 | CATATCGCCATTA | - | STAS domain-containing protein |
| AVAEIV_RS21155 | 4498004 | 4498016 | 216 | - | 11.8857 | 3.70E-05 | 0.0753 | CAGTGCGCCGGCG | - | MFS transporter |
| AVAEIV_RS26025 | 5591619 | 5591631 | 335 | + | 11.8857 | 3.70E-05 | 0.0718 | TCGATCGCCACCA | - | IS5 family transposase |
| AVAEIV_RS03410 | 688491 | 688503 | 311 | + | 11.8714 | 3.74E-05 | 0.0753 | TCGCTCGCCATCA | *purD* | phosphoribosylamine--glycine ligase |
| AVAEIV_RS06865 | 1412521 | 1412533 | 8 | + | 11.8714 | 3.74E-05 | 0.0753 | CCGCTGGCCGGCA | - | YraN family protein |
| AVAEIV_RS07440 | 1546044 | 1546056 | 14 | - | 11.8714 | 3.74E-05 | 0.0718 | CGGCATGCCAGCG | - | efflux RND transporter permease subunit |
| AVAEIV_RS07515 | 1567556 | 1567568 | 266 | - | 11.8714 | 3.74E-05 | 0.0753 | CAGATTGACATCG | *lexA* | transcriptional repressor LexA |
| AVAEIV_RS09580 | 1996899 | 1996911 | 105 | + | 11.8714 | 3.74E-05 | 0.0718 | CCAATGGCCGGCA | - | dipeptide ABC transporter ATP-binding protein |
| AVAEIV_RS13815 | 2993962 | 2993974 | 262 | - | 11.8714 | 3.74E-05 | 0.0753 | CCGCTGGCCGGCA | *tssF* | type VI secretion system baseplate subunit TssF |
| AVAEIV_RS19750 | 4208951 | 4208963 | 226 | - | 11.8714 | 3.74E-05 | 0.0753 | CAGGGCGCCATCC | - | hypothetical protein |
| AVAEIV_RS22615 | 4843256 | 4843268 | 169 | + | 11.8714 | 3.74E-05 | 0.0718 | CCGCTGGCCGGCA | - | SCP2 domain-containing protein |
| AVAEIV_RS24825 | 5280852 | 5280864 | 114 | - | 11.8714 | 3.74E-05 | 0.0753 | CCCGACGCCATCG | - | sensor histidine kinase |
| AVAEIV_RS24865 | 5290449 | 5290461 | 85 | - | 11.8714 | 3.74E-05 | 0.0753 | GAGAGCGCCATCG | - | subunit alpha |
| AVAEIV_RS25190 | 5364566 | 5364578 | 252 | + | 11.8714 | 3.74E-05 | 0.0753 | CCGCTGGCCGGCA | - | nickel-dependent hydrogenase large subunit |
| AVAEIV_RS25545 | 5459011 | 5459023 | -48 | + | 11.8714 | 3.74E-05 | 0.0753 | CGTCTTGCCATCA | - | glycosyl hydrolase family 28-related protein |
| AVAEIV_RS01410 | 274999 | 275011 | 347 | - | 11.8571 | 3.77E-05 | 0.0753 | AAGTTCGCCATCA | *argB* | acetylglutamate kinase |
| AVAEIV_RS03935 | 793256 | 793268 | 323 | - | 11.8571 | 3.77E-05 | 0.0718 | CAGAGCGCCACCT | *mpl* | UDP-N-acetylmuramate:L-alanyl-gamma-D-glutamyl-meso-diaminopimelate ligase |
| AVAEIV_RS07750 | 1613352 | 1613364 | 157 | + | 11.8571 | 3.77E-05 | 0.0718 | CCGGTCGCCGGCA | - | TatD family hydrolase |
| AVAEIV_RS11260 | 2379684 | 2379696 | 327 | - | 11.8571 | 3.77E-05 | 0.0753 | CCCAGTGCCAGCG | - | family protein |
| AVAEIV_RS12920 | 2801538 | 2801550 | 280 | - | 11.8571 | 3.77E-05 | 0.0753 | CCGGTCGCCGGCA | - | DNA-binding protein |
| AVAEIV_RS15645 | 3426038 | 3426050 | 176 | + | 11.8571 | 3.77E-05 | 0.0753 | CCGGTCGCCGGCA | - | heme ABC transporter permease |
| AVAEIV_RS16025 | 3492588 | 3492600 | 295 | - | 11.8571 | 3.77E-05 | 0.0753 | CCGGTCGCCGGCA | *ttdA* | L(+)-tartrate dehydratase subunit alpha |
| AVAEIV_RS16935 | 3668461 | 3668473 | -36 | + | 11.8571 | 3.77E-05 | 0.0753 | CACGACGCCACCG | - | cytochrome c |
| AVAEIV_RS20905 | 4444502 | 4444514 | 75 | + | 11.8571 | 3.77E-05 | 0.0718 | CGGCGCGCCACCG | - | thiamine pyrophosphate-dependent dehydrogenase E1 component subunit alpha |
| AVAEIV_RS23570 | 5024000 | 5024012 | 127 | + | 11.8571 | 3.77E-05 | 0.0718 | CGGCGCGCCACCG | *pdxA* | 4-hydroxythreonine-4-phosphate dehydrogenase PdxA |
| AVAEIV_RS23840 | 5081986 | 5081998 | 230 | + | 11.8571 | 3.77E-05 | 0.0753 | CCGGTCGCCGGCA | *pepP* | Xaa-Pro aminopeptidase |
| AVAEIV_RS24940 | 5307479 | 5307491 | 313 | - | 11.8571 | 3.77E-05 | 0.0753 | CCCAGTGCCAGCG | - | DMT family transporter |
| AVAEIV_RS05050 | 1037864 | 1037876 | 230 | + | 11.8429 | 3.81E-05 | 0.0753 | CAGTTGGCCGCCG | - | alginate O-acetyltransferase AlgX-related protein |
| AVAEIV_RS06145 | 1253031 | 1253043 | 48 | - | 11.8429 | 3.81E-05 | 0.0753 | CAACTGGCCGGCA | - | type 4 pilus major pilin |
| AVAEIV_RS06210 | 1268041 | 1268053 | 103 | - | 11.8429 | 3.81E-05 | 0.0753 | CACATGGCCAACA | - | single-stranded DNA-binding protein |
| AVAEIV_RS07930 | 1649223 | 1649235 | 275 | - | 11.8429 | 3.81E-05 | 0.0753 | CAACTGGCCGGCA | - | DUF2959 domain-containing protein |
| AVAEIV_RS10055 | 2104395 | 2104407 | 359 | - | 11.8429 | 3.81E-05 | 0.0753 | CAGTTGGCCGCCG | - | binding protein |
| AVAEIV_RS10065 | 2104395 | 2104407 | 226 | - | 11.8429 | 3.81E-05 | 0.0718 | CAGTTGGCCGCCG | *atpD* | F0F1 ATP synthase subunit beta |
| AVAEIV_RS11210 | 2366433 | 2366445 | 302 | + | 11.8429 | 3.81E-05 | 0.0753 | CAGTTGGCCGCCG | - | ferredoxin family protein |
| AVAEIV_RS15625 | 3422593 | 3422605 | 93 | + | 11.8429 | 3.81E-05 | 0.0753 | CCGAAAGCCATCA | - | DsbE family thiol:disulfide interchange protein |
| AVAEIV_RS16350 | 3552736 | 3552748 | 355 | - | 11.8429 | 3.81E-05 | 0.0753 | CGCAAGGCCACCG | - | BrnT family toxin |
| AVAEIV_RS22455 | 4799687 | 4799699 | 264 | + | 11.8429 | 3.81E-05 | 0.0718 | CCGAACGCCGGCG | - | potassium/proton antiporter |
| AVAEIV_RS01530 | 294692 | 294704 | 213 | + | 11.8286 | 3.85E-05 | 0.0753 | GCGATGGCCAGCA | *ruvX* | Holliday junction resolvase RuvX |
| AVAEIV_RS03445 | 695791 | 695803 | 185 | + | 11.8286 | 3.85E-05 | 0.0753 | CTCACCGCCAGCA | *accB* | acetyl-CoA carboxylase biotin carboxyl carrier protein |
| AVAEIV_RS03510 | 707847 | 707859 | 235 | - | 11.8286 | 3.85E-05 | 0.0718 | CACGGGGCCATCG | - | SdiA-regulated domain-containing protein |
| AVAEIV_RS04930 | 1006893 | 1006905 | -44 | + | 11.8286 | 3.85E-05 | 0.0753 | GCGATGGCCAGCA | - | OprD family outer membrane porin |
| AVAEIV_RS10290 | 2147783 | 2147795 | 358 | + | 11.8286 | 3.85E-05 | 0.0718 | CGCCAGGCCATCG | - | mechanosensitive ion channel family protein |
| AVAEIV_RS10745 | 2251300 | 2251312 | 280 | + | 11.8286 | 3.85E-05 | 0.0718 | CGCCAGGCCATCG | *csx16* | CRISPR-associated protein Csx16 |
| AVAEIV_RS12045 | 2558437 | 2558449 | 339 | + | 11.8286 | 3.85E-05 | 0.0753 | CCGATCGCCTGCA | - | hypothetical protein |
| AVAEIV_RS12640 | 2681121 | 2681133 | 16 | + | 11.8286 | 3.85E-05 | 0.0753 | CAAAGTGCCGTCA | - | protein |
| AVAEIV_RS13240 | 2873256 | 2873268 | 292 | - | 11.8286 | 3.85E-05 | 0.0753 | CGCCAGGCCATCG | - | IS701-like element ISAzvi8 family transposase |
| AVAEIV_RS13535 | 2930174 | 2930186 | 323 | - | 11.8286 | 3.85E-05 | 0.0753 | CGCCAGGCCATCG | - | aldehyde dehydrogenase |
| AVAEIV_RS14690 | 3195728 | 3195740 | 212 | + | 11.8286 | 3.85E-05 | 0.0718 | CGCCAGGCCATCG | - | ATP-binding protein |
| AVAEIV_RS15340 | 3353888 | 3353900 | 64 | - | 11.8286 | 3.85E-05 | 0.0753 | CAGCGTGCCGTCA | - | succinate dehydrogenase iron-sulfur subunit |
| AVAEIV_RS15545 | 3401597 | 3401609 | 367 | - | 11.8286 | 3.85E-05 | 0.0753 | CGCCAGGCCATCG | - | Gfo/Idh/MocA family protein |

| | | | | | | | | | | |
|---|---|---|---|---|---|---|---|---|---|---|
| AVAEIV_RS15775 | 3445131 | 3445143 | -41 | - | 11.8286 | 3.85E-05 | 0.0718 | GCGATGGCCAGCA | - | lysozyme inhibitor LprI family protein |
| AVAEIV_RS16600 | 3598294 | 3598306 | 50 | + | 11.8286 | 3.85E-05 | 0.0718 | CACCGCGCCATCC | - | vWA domain-containing protein |
| AVAEIV_RS22335 | 4770835 | 4770847 | 234 | + | 11.8286 | 3.85E-05 | 0.0753 | CGATCCGCCAGCA | - | lipopolysaccharide kinase InaA family protein |
| AVAEIV_RS22390 | 4787539 | 4787551 | 218 | - | 11.8286 | 3.85E-05 | 0.0718 | GCGATGGCCAGCA | - | universal stress protein |
| AVAEIV_RS22805 | 4882109 | 4882121 | 283 | + | 11.8286 | 3.85E-05 | 0.0718 | CGCCAGGCCATCG | - | IS5 family transposase |
| AVAEIV_RS23530 | 5013407 | 5013419 | -49 | + | 11.8286 | 3.85E-05 | 0.0718 | CAGGTCGGCAGCA | - | GlcG/HbpS family heme-binding protein |
| AVAEIV_RS23905 | 5094624 | 5094636 | 213 | + | 11.8286 | 3.85E-05 | 0.0753 | CATATCGCCGCCA | *hemB* | porphobilinogen synthase |
| AVAEIV_RS23910 | 5094624 | 5094636 | 131 | + | 11.8286 | 3.85E-05 | 0.0718 | CATATCGCCGCCA | *elbB* | isoprenoid biosynthesis glyoxalase ElbB |
| AVAEIV_RS24385 | 5184312 | 5184324 | 38 | + | 11.8286 | 3.85E-05 | 0.0753 | CAAGTCGCCGGCA | - | rubredoxin |
| AVAEIV_RS24390 | 5184312 | 5184324 | 88 | + | 11.8286 | 3.85E-05 | 0.0718 | CAAGTCGCCGGCA | - | chorismate--pyruvate lyase family protein |
| AVAEIV_RS25970 | 5580516 | 5580528 | 240 | + | 11.8286 | 3.85E-05 | 0.0753 | CAGCTTGCCACCC | - | F0F1 ATP synthase subunit B |
| AVAEIV_RS00290 | 63995 | 64007 | 291 | + | 11.8143 | 3.88E-05 | 0.0753 | CTCAGCGCCAGCG | - | ATP-binding cassette domain-containing protein |
| AVAEIV_RS02250 | 450733 | 450745 | 104 | + | 11.8143 | 3.88E-05 | 0.0753 | CAGCACGCCGGCG | - | AAA family ATPase |
| AVAEIV_RS04705 | 958162 | 958174 | 213 | - | 11.8143 | 3.88E-05 | 0.0718 | CGCGACGCCATCG | - | alpha/beta fold hydrolase |
| AVAEIV_RS05640 | 1160665 | 1160677 | 260 | + | 11.8143 | 3.88E-05 | 0.0753 | CAGCACGCCGGCG | - | containing protein |
| AVAEIV_RS05995 | 1222109 | 1222121 | 373 | - | 11.8143 | 3.88E-05 | 0.0753 | CAGCACGCCGGCG | - | membrane protein |
| AVAEIV_RS07925 | 1647347 | 1647359 | 253 | + | 11.8143 | 3.88E-05 | 0.0718 | CGCGACGCCATCG | - | thiopurine S-methyltransferase |
| AVAEIV_RS07995 | 1660319 | 1660331 | 304 | + | 11.8143 | 3.88E-05 | 0.0718 | CATATCGGCATCA | - | ABC transporter permease |
| AVAEIV_RS08050 | 1671139 | 1671151 | 387 | + | 11.8143 | 3.88E-05 | 0.0753 | CGGCTGGCCGGCA | - | septation protein A |
| AVAEIV_RS11750 | 2488990 | 2489002 | 285 | - | 11.8143 | 3.88E-05 | 0.0753 | CGGATGGGCAGCA | - | Lpp/OprI family alanine-zipper lipoprotein |
| AVAEIV_RS11755 | 2488990 | 2489002 | 160 | - | 11.8143 | 3.88E-05 | 0.0718 | CGGATGGGCAGCA | - | GNAT family N-acetyltransferase |
| AVAEIV_RS12320 | 2606984 | 2606996 | -44 | - | 11.8143 | 3.88E-05 | 0.0718 | CTCAGCGCCAGCG | - | EscU/YscU/HrcU family type III secretion system export apparatus switch protein |
| AVAEIV_RS14775 | 3212859 | 3212871 | 343 | - | 11.8143 | 3.88E-05 | 0.0718 | CAGCACGCCGGCG | *nuoK* | NADH-quinone oxidoreductase subunit NuoK |
| AVAEIV_RS14885 | 3238648 | 3238660 | 83 | - | 11.8143 | 3.88E-05 | 0.0753 | CAGCACGCCGGCG | - | efflux transporter outer membrane subunit |
| AVAEIV_RS17900 | 3874890 | 3874902 | 373 | + | 11.8143 | 3.88E-05 | 0.0718 | CAGCACGCCGGCG | - | membrane protein |
| AVAEIV_RS17980 | 3887388 | 3887400 | 260 | - | 11.8143 | 3.88E-05 | 0.0718 | CAGCACGCCGGCG | - | containing protein |
| AVAEIV_RS18310 | 3942935 | 3942947 | 274 | - | 11.8143 | 3.88E-05 | 0.0753 | CAAATAGCCATCG | - | hypothetical protein |
| AVAEIV_RS18315 | 3942935 | 3942947 | 36 | - | 11.8143 | 3.88E-05 | 0.0753 | CAAATAGCCATCG | - | AlpA family phage regulatory protein |
| AVAEIV_RS18530 | 3989044 | 3989056 | 152 | + | 11.8143 | 3.88E-05 | 0.0718 | CGCGACGCCATCG | - | di-heme oxidoredictase family protein |
| AVAEIV_RS19150 | 4130212 | 4130224 | 270 | + | 11.8143 | 3.88E-05 | 0.0753 | CGATGCGCCAGCG | - | phage tail protein |
| AVAEIV_RS20495 | 4358479 | 4358491 | 358 | + | 11.8143 | 3.88E-05 | 0.0753 | CAGACGGCCGGCA | - | YbdD/YjiX family protein |
| AVAEIV_RS22940 | 4916224 | 4916236 | 307 | + | 11.8143 | 3.88E-05 | 0.0753 | CTGATCGCCACCC | *argC* | N-acetyl-gamma-glutamyl-phosphate reductase |
| AVAEIV_RS22950 | 4916224 | 4916236 | 371 | + | 11.8143 | 3.88E-05 | 0.0718 | CTGATCGCCACCC | - | histidine triad nucleotide-binding protein |
| AVAEIV_RS23345 | 4975780 | 4975792 | 366 | - | 11.8143 | 3.88E-05 | 0.0718 | CGCGACGCCATCG | - | DUF2267 domain-containing protein |
| AVAEIV_RS24870 | 5291529 | 5291541 | 368 | + | 11.8143 | 3.88E-05 | 0.0753 | CAGCACGCCGGCG | - | gamma-carboxygeranoyl-CoA hydratase |
| AVAEIV_RS01230 | 243244 | 243256 | 36 | - | 11.8 | 3.92E-05 | 0.0753 | CTGCTTGCCAGCG | - | nucleotidyltransferase substrate binding protein |
| AVAEIV_RS03495 | 706302 | 706314 | 84 | + | 11.8 | 3.92E-05 | 0.0753 | CCGCCGGCCAGCA | - | MATE family efflux transporter |
| AVAEIV_RS04450 | 907598 | 907610 | 201 | - | 11.8 | 3.92E-05 | 0.0753 | CTGGTGGCCATCG | *hemL* | glutamate-1-semialdehyde 2,1-aminomutase |
| AVAEIV_RS05110 | 1051943 | 1051955 | 315 | - | 11.8 | 3.92E-05 | 0.0753 | CTGCTCGCCATCC | *cydB* | cytochrome d ubiquinol oxidase subunit II |
| AVAEIV_RS08660 | 1808090 | 1808102 | -49 | - | 11.8 | 3.92E-05 | 0.0753 | CTGGTGGCCATCG | - | HlyD family secretion protein |
| AVAEIV_RS09555 | 1992153 | 1992165 | 111 | - | 11.8 | 3.92E-05 | 0.0718 | CGGGTCGCCGGCA | - | HIT family protein |
| AVAEIV_RS10505 | 2197845 | 2197857 | 234 | + | 11.8 | 3.92E-05 | 0.0753 | CTGGAGGCCATCA | - | hypothetical protein |
| AVAEIV_RS11130 | 2347840 | 2347852 | 327 | - | 11.8 | 3.92E-05 | 0.0718 | CCGCCGGCCAGCA | - | ABC transporter permease |
| AVAEIV_RS11540 | 2441489 | 2441501 | -28 | + | 11.8 | 3.92E-05 | 0.0718 | CCCTTCGCCGTCG | - | hypothetical protein |
| AVAEIV_RS11545 | 2441489 | 2441501 | 358 | + | 11.8 | 3.92E-05 | 0.0718 | CCCTTCGCCGTCG | - | hypothetical protein |
| AVAEIV_RS16980 | 3681874 | 3681886 | 144 | + | 11.8 | 3.92E-05 | 0.0753 | CGGTTGGCCGTCG | - | YMGG-like glycine zipper-containing protein |
| AVAEIV_RS16985 | 3681874 | 3681886 | 65 | + | 11.8 | 3.92E-05 | 0.0718 | CGGTTGGCCGTCG | - | transposase |
| AVAEIV_RS17115 | 3712666 | 3712678 | 201 | + | 11.8 | 3.92E-05 | 0.0753 | CGCAGCGCCATCC | - | NAD(+) kinase |
| AVAEIV_RS17170 | 3721827 | 3721839 | 92 | - | 11.8 | 3.92E-05 | 0.0753 | CGCAGTGCCAGCG | - | SPOR domain-containing protein |
| AVAEIV_RS18540 | 3991729 | 3991741 | -29 | - | 11.8 | 3.92E-05 | 0.0718 | CCGCCGGCCAGCA | - | DUF1513 domain-containing protein |
| AVAEIV_RS18685 | 4024348 | 4024360 | 88 | + | 11.8 | 3.92E-05 | 0.0753 | CGGGTCGCCGGCA | *pcaD* | 3-oxoadipate enol-lactonase |

| | | | | | | | | | | |
|---|---|---|---|---|---|---|---|---|---|---|
| AVAEIV_RS22010 | 4696427 | 4696439 | 177 | + | 11.8 | 3.92E-05 | 0.0718 | CGGGTCGCCGGCA | - | integrase core domain-containing protein |
| AVAEIV_RS22145 | 4725519 | 4725531 | 308 | - | 11.8 | 3.92E-05 | 0.0753 | CGGGTCGCCGGCA | - | containing protein |
| AVAEIV_RS22550 | 4827859 | 4827871 | 178 | + | 11.8 | 3.92E-05 | 0.0753 | CCGCCGGCCAGCA | - | PilN domain-containing protein |
| AVAEIV_RS22625 | 4845547 | 4845559 | 152 | - | 11.8 | 3.92E-05 | 0.0718 | CCGCCGGCCAGCA | *hisI* | phosphoribosyl-AMP cyclohydrolase |
| AVAEIV_RS23055 | 4933163 | 4933175 | 334 | + | 11.8 | 3.92E-05 | 0.0718 | CTGGTGGCCATCG | *gpM* | phage terminase small subunit |
| AVAEIV_RS23755 | 5061642 | 5061654 | 384 | + | 11.8 | 3.92E-05 | 0.0718 | CCGCCGGCCAGCA | *fecA* | TonB-dependent Fe(3+) dicitrate receptor FecA |
| AVAEIV_RS24335 | 5174671 | 5174683 | 94 | + | 11.8 | 3.92E-05 | 0.0753 | CGGGTCGCCGGCA | - | DUF523 domain-containing protein |
| AVAEIV_RS24350 | 5177458 | 5177470 | 145 | + | 11.8 | 3.92E-05 | 0.0753 | CTGCTCGCCATCC | - | alpha/beta hydrolase |
| AVAEIV_RS24355 | 5177458 | 5177470 | -49 | + | 11.8 | 3.92E-05 | 0.0718 | CTGCTCGCCATCC | - | DUF6436 domain-containing protein |
| AVAEIV_RS24515 | 5210005 | 5210017 | 257 | - | 11.8 | 3.92E-05 | 0.0753 | CAGCGCGCCATGA | - | ABC transporter ATP-binding protein |
| AVAEIV_RS24735 | 5255226 | 5255238 | 55 | + | 11.8 | 3.92E-05 | 0.0753 | CAGAGGGCCGGCG | - | FadR/GntR family transcriptional regulator |
| AVAEIV_RS25550 | 5461137 | 5461149 | 77 | + | 11.8 | 3.92E-05 | 0.0753 | CCGCCGGCCAGCA | - | 2-hydroxyacid dehydrogenase |
| AVAEIV_RS02220 | 443689 | 443701 | 17 | - | 11.7857 | 3.96E-05 | 0.0718 | CGGAACGCCGGCG | - | Ni/Fe hydrogenase subunit alpha |
| AVAEIV_RS02675 | 546812 | 546824 | 239 | + | 11.7857 | 3.96E-05 | 0.0753 | CCGGCCGCCAGCA | - | glycosyltransferase |
| AVAEIV_RS05295 | 1093793 | 1093805 | 358 | + | 11.7857 | 3.96E-05 | 0.0753 | CCGGCCGCCAGCA | - | hypothetical protein |
| AVAEIV_RS05605 | 1156009 | 1156021 | 164 | - | 11.7857 | 3.96E-05 | 0.0718 | CGGCTTGCCATCC | - | family |
| AVAEIV_RS10030 | 2098364 | 2098376 | 356 | + | 11.7857 | 3.96E-05 | 0.0718 | GAGGTCGCCAGCA | - | DUF5064 family protein |
| AVAEIV_RS11015 | 2318855 | 2318867 | 210 | + | 11.7857 | 3.96E-05 | 0.0718 | CCGCGGGCCAGCG | - | ABC transporter permease |
| AVAEIV_RS12085 | 2564517 | 2564529 | 182 | - | 11.7857 | 3.96E-05 | 0.0718 | CACTTCGCCGCCG | *mtnC* | acireductone synthase |
| AVAEIV_RS12975 | 2811210 | 2811222 | -38 | + | 11.7857 | 3.96E-05 | 0.0753 | CCGGCCGCCAGCA | - | nucleotidyltransferase family protein |
| AVAEIV_RS14700 | 3197035 | 3197047 | 305 | - | 11.7857 | 3.96E-05 | 0.0718 | GAGTTGGCCATCG | - | IS630 family transposase |
| AVAEIV_RS15050 | 3277878 | 3277890 | 297 | - | 11.7857 | 3.96E-05 | 0.0753 | CAGCAGGCCAGCC | - | GspMb/PilO family protein |
| AVAEIV_RS15715 | 3435098 | 3435110 | 244 | - | 11.7857 | 3.96E-05 | 0.0753 | CCGGCCGCCAGCA | - | peroxiredoxin |
| AVAEIV_RS15720 | 3435098 | 3435110 | -38 | - | 11.7857 | 3.96E-05 | 0.0718 | CCGGCCGCCAGCA | - | sulfate ABC transporter substrate-binding protein |
| AVAEIV_RS16070 | 3501097 | 3501109 | 155 | - | 11.7857 | 3.96E-05 | 0.0718 | CAGTTCGCCTTCG | - | 4-oxalomesaconate tautomerase |
| AVAEIV_RS19420 | 4161146 | 4161158 | 334 | + | 11.7857 | 3.96E-05 | 0.0753 | GAGGTCGCCAGCA | - | hypothetical protein |
| AVAEIV_RS20310 | 4327620 | 4327632 | -37 | + | 11.7857 | 3.96E-05 | 0.0718 | CAGCAGGCCAGCC | - | hypothetical protein |
| AVAEIV_RS20315 | 4327620 | 4327632 | 343 | + | 11.7857 | 3.96E-05 | 0.0718 | CAGCAGGCCAGCC | - | Rha family transcriptional regulator |
| AVAEIV_RS21805 | 4643203 | 4643215 | 240 | + | 11.7857 | 3.96E-05 | 0.0718 | CTGCGCGCCACCA | - | SfnB family sulfur acquisition oxidoreductase |
| AVAEIV_RS22365 | 4776936 | 4776948 | 257 | + | 11.7857 | 3.96E-05 | 0.0753 | CAGCAGGCCAGCC | - | branched-chain amino acid transaminase |
| AVAEIV_RS22980 | 4922308 | 4922320 | 379 | - | 11.7857 | 3.96E-05 | 0.0753 | CATTTCGCCGGCA | *trpD* | anthranilate phosphoribosyltransferase |
| AVAEIV_RS24505 | 5207653 | 5207665 | -48 | + | 11.7857 | 3.96E-05 | 0.0753 | CAGCAGGCCAGCC | - | ABC transporter permease subunit |
| AVAEIV_RS24835 | 5281822 | 5281834 | -38 | - | 11.7857 | 3.96E-05 | 0.0718 | CCGGCCGCCAGCA | - | substrate binding protein |
| AVAEIV_RS25940 | 5573073 | 5573085 | 188 | - | 11.7857 | 3.96E-05 | 0.0718 | CGTATCGCCGTCA | - | di-heme oxidoredictase family protein |
| AVAEIV_RS03570 | 721161 | 721173 | 143 | - | 11.7714 | 4.00E-05 | 0.0753 | CCTACCGCCATCA | - | DUF4136 domain-containing protein |
| AVAEIV_RS06160 | 1256252 | 1256264 | -50 | + | 11.7714 | 4.00E-05 | 0.0753 | CCGAACGACATCA | *pilP* | type IV pilus biogenesis protein PilP |
| AVAEIV_RS11420 | 2417071 | 2417083 | 80 | - | 11.7714 | 4.00E-05 | 0.0753 | CCGGGCGCCAGCG | - | TauD/TfdA dioxygenase family protein |
| AVAEIV_RS12510 | 2657501 | 2657513 | 75 | - | 11.7714 | 4.00E-05 | 0.0753 | CTGTTCGCCAGCC | *malQ* | 4-alpha-glucanotransferase |
| AVAEIV_RS18900 | 4070099 | 4070111 | 235 | + | 11.7714 | 4.00E-05 | 0.072 | CTGTTCGCCAGCC | *fdxA* | ferredoxin FdxA |
| AVAEIV_RS19050 | 4104257 | 4104269 | 234 | - | 11.7714 | 4.00E-05 | 0.0753 | CTCAAGGCCACCA | *frr* | ribosome recycling factor |
| AVAEIV_RS20145 | 4302931 | 4302943 | -5 | + | 11.7714 | 4.00E-05 | 0.0753 | GCCATCGCCAGCA | *bamB* | outer membrane protein assembly factor BamB |
| AVAEIV_RS24900 | 5298168 | 5298180 | 333 | + | 11.7714 | 4.00E-05 | 0.072 | CACTACGGCATCA | - | TerC family protein |
| AVAEIV_RS25290 | 5385855 | 5385867 | 132 | + | 11.7714 | 4.00E-05 | 0.072 | CCGGGCGCCAGCG | - | ATP-binding cassette domain-containing protein |
| AVAEIV_RS03235 | 652665 | 652677 | 273 | + | 11.7571 | 4.03E-05 | 0.072 | CACAAGGCCATTA | - | DNA-directed RNA polymerase subunit alpha |
| AVAEIV_RS03695 | 743827 | 743839 | 102 | + | 11.7571 | 4.03E-05 | 0.072 | CCGAGCGCTATCA | *mutL* | DNA mismatch repair endonuclease MutL |
| AVAEIV_RS04410 | 896421 | 896433 | 63 | + | 11.7571 | 4.03E-05 | 0.072 | CTCCTGGCCATCG | *lra1* | NAD(P)+-dependent L-rhamnose 1-dehydrogenase Lra1 |
| AVAEIV_RS06245 | 1276468 | 1276480 | 156 | + | 11.7571 | 4.03E-05 | 0.0753 | CGGTTTGCCAGCC | - | Arc family DNA-binding protein |
| AVAEIV_RS07185 | 1481412 | 1481424 | -40 | - | 11.7571 | 4.03E-05 | 0.0753 | CGCCTCGCCGGCA | *cmoB* | acid(34) synthase CmoB |
| AVAEIV_RS12975 | 2811569 | 2811581 | 321 | + | 11.7571 | 4.03E-05 | 0.0753 | TCGATGGCCAGCA | - | nucleotidyltransferase family protein |
| AVAEIV_RS13890 | 3010510 | 3010522 | 164 | - | 11.7571 | 4.03E-05 | 0.072 | CAGATGGCGATCG | - | DUF6152 family protein |
| AVAEIV_RS18825 | 4053587 | 4053599 | 298 | + | 11.7571 | 4.03E-05 | 0.0753 | CGCCTCGCCGGCA | *erdR* | response regulator transcription factor ErdR |
| AVAEIV_RS19030 | 4100440 | 4100452 | -44 | + | 11.7571 | 4.03E-05 | 0.0753 | CCGAGCGCTATCA | *rseP* | RIP metalloprotease RseP |

| | | | | | | | | | | |
|---|---|---|---|---|---|---|---|---|---|---|
| AVAEIV_RS22270 | 4754554 | 4754566 | 347 | + | 11.7571 | 4.03E-05 | 0.0753 | TCGATGGCCAGCA | - | acyltransferase family protein |
| AVAEIV_RS03270 | 661492 | 661504 | 328 | - | 11.7429 | 4.06E-05 | 0.072 | CGGCCGGCCAGCA | - | sugar nucleotide-binding protein |
| AVAEIV_RS04710 | 959100 | 959112 | 259 | - | 11.7429 | 4.06E-05 | 0.072 | CGGCCGGCCAGCA | - | DUF4892 domain-containing protein |
| AVAEIV_RS07770 | 1617559 | 1617571 | 71 | - | 11.7429 | 4.06E-05 | 0.072 | CACAGCGCCGGCG | - | 3-carboxyethylcatechol 2,3-dioxygenase |
| AVAEIV_RS07875 | 1637726 | 1637738 | 177 | + | 11.7429 | 4.06E-05 | 0.072 | CACAGCGCCGGCG | *earP* | rhamnosyltransferase EarP |
| AVAEIV_RS09600 | 2002528 | 2002540 | 318 | + | 11.7429 | 4.06E-05 | 0.072 | CAGATCGCCGCCC | - | acetyl-CoA hydrolase/transferase family protein |
| AVAEIV_RS09710 | 2033900 | 2033912 | 29 | + | 11.7429 | 4.06E-05 | 0.0753 | CGGCCGGCCAGCA | - | phage holin family protein |
| AVAEIV_RS10105 | 2110066 | 2110078 | 192 | - | 11.7429 | 4.06E-05 | 0.072 | CCGAGCGCCGCCA | - | F0F1 ATP synthase subunit gamma |
| AVAEIV_RS11130 | 2347911 | 2347923 | 256 | + | 11.7429 | 4.06E-05 | 0.072 | CAGATCGCCGCCC | - | ABC transporter permease |
| AVAEIV_RS13160 | 2855893 | 2855905 | -46 | - | 11.7429 | 4.06E-05 | 0.0753 | CAGATCGCCGCCC | - | LacI family DNA-binding transcriptional regulator |
| AVAEIV_RS13165 | 2855893 | 2855905 | 195 | - | 11.7429 | 4.06E-05 | 0.072 | CAGATCGCCGCCC | - | SDR family oxidoreductase |
| AVAEIV_RS13215 | 2865707 | 2865719 | -44 | - | 11.7429 | 4.06E-05 | 0.072 | CCCCCCGCCAGCA | - | formimidoylglutamate deiminase |
| AVAEIV_RS15650 | 3426832 | 3426844 | 216 | - | 11.7429 | 4.06E-05 | 0.0753 | CGGCCGGCCAGCA | *ccmB* | heme exporter protein CcmB |
| AVAEIV_RS17050 | 3699267 | 3699279 | 364 | - | 11.7429 | 4.06E-05 | 0.0753 | CAGTTCGCCAACG | - | NAD(P)-dependent oxidoreductase |
| AVAEIV_RS24155 | 5136445 | 5136457 | 235 | + | 11.7429 | 4.06E-05 | 0.0753 | CAGATCGCCGCCC | - | RidA family protein |
| AVAEIV_RS25835 | 5537812 | 5537824 | 285 | + | 11.7429 | 4.06E-05 | 0.0753 | CGGCCGGCCAGCA | - | DeoR/GlpR family DNA-binding transcription regulator |
| AVAEIV_RS25870 | 5547500 | 5547512 | 159 | - | 11.7429 | 4.06E-05 | 0.072 | CCGAGCGCCGCCA | - | ABC transporter ATP-binding protein |
| AVAEIV_RS26010 | 5587762 | 5587774 | 207 | - | 11.7429 | 4.06E-05 | 0.0753 | CCGAGCGCCGCCA | - | hypothetical protein |
| AVAEIV_RS26055 | 5603886 | 5603898 | 262 | - | 11.7429 | 4.06E-05 | 0.0753 | CAGCACGACATCA | - | protein |
| AVAEIV_RS01640 | 318094 | 318106 | 217 | + | 11.7286 | 4.10E-05 | 0.0753 | CAGGTGGCCGTCG | - | methyl-accepting chemotaxis protein |
| AVAEIV_RS02470 | 493952 | 493964 | 231 | - | 11.7286 | 4.10E-05 | 0.0753 | CGCGTTGCCATCG | - | ABC transporter permease |
| AVAEIV_RS02685 | 549117 | 549129 | 186 | - | 11.7286 | 4.10E-05 | 0.0753 | CACGAGGCCAGCG | - | O-antigen ligase family protein |
| AVAEIV_RS03670 | 738576 | 738588 | 269 | + | 11.7286 | 4.10E-05 | 0.0723 | CAGCTCGCCGTCC | *orn* | oligoribonuclease |
| AVAEIV_RS04925 | 1005511 | 1005523 | 331 | - | 11.7286 | 4.10E-05 | 0.0753 | CCGATCGCCAGGG | *exbB* | tonB-system energizer ExbB |
| AVAEIV_RS05125 | 1054554 | 1054566 | 358 | - | 11.7286 | 4.10E-05 | 0.0753 | CAGCTTGCCGGCG | - | hypothetical protein |
| AVAEIV_RS06925 | 1424414 | 1424426 | 302 | - | 11.7286 | 4.10E-05 | 0.0723 | CAGCTTGCCGGCG | *ftsW* | putative lipid II flippase FtsW |
| AVAEIV_RS11320 | 2392441 | 2392453 | 74 | - | 11.7286 | 4.10E-05 | 0.0723 | CGGGCCGCCAGCA | *rpiA* | ribose-5-phosphate isomerase RpiA |
| AVAEIV_RS13155 | 2854735 | 2854747 | 57 | + | 11.7286 | 4.10E-05 | 0.0753 | CGGCGGGCCAGCG | *hutU* | urocanate hydratase |
| AVAEIV_RS14365 | 3126906 | 3126918 | 201 | + | 11.7286 | 4.10E-05 | 0.0753 | CCCCGCGCCAGCG | - | XRE family transcriptional regulator |
| AVAEIV_RS15830 | 3453446 | 3453458 | 234 | - | 11.7286 | 4.10E-05 | 0.0753 | GACTTCGCCATCG | - | beta |
| AVAEIV_RS20675 | 4394828 | 4394840 | 357 | + | 11.7286 | 4.10E-05 | 0.0723 | CAGCTTGCCGGCG | *upp* | uracil phosphoribosyltransferase |
| AVAEIV_RS22345 | 4772252 | 4772264 | 22 | + | 11.7286 | 4.10E-05 | 0.0753 | CAGCTTGCCGGCG | *rfaP* | lipopolysaccharide core heptose(I) kinase RfaP |
| AVAEIV_RS22365 | 4776662 | 4776674 | -17 | + | 11.7286 | 4.10E-05 | 0.0753 | CGGTCGGCCATCG | - | branched-chain amino acid transaminase |
| AVAEIV_RS22980 | 4922303 | 4922315 | 374 | + | 11.7286 | 4.10E-05 | 0.0753 | CAGCTTGCCGGCG | *trpD* | anthranilate phosphoribosyltransferase |
| AVAEIV_RS23310 | 4970846 | 4970858 | 368 | - | 11.7286 | 4.10E-05 | 0.0753 | CCGCGCGCCGTCA | - | nicotinate phosphoribosyltransferase |
| AVAEIV_RS23455 | 4998205 | 4998217 | 133 | + | 11.7286 | 4.10E-05 | 0.0723 | CACGAGGCCAGCG | - | ATP-binding protein |
| AVAEIV_RS25530 | 5452018 | 5452030 | 288 | - | 11.7286 | 4.10E-05 | 0.0753 | CCGGTGGCCACCG | - | glycosyl hydrolase family 28-related protein |
| AVAEIV_RS25695 | 5498244 | 5498256 | 259 | - | 11.7286 | 4.10E-05 | 0.0753 | GACTTCGCCATCG | - | CoA transferase subunit B |
| AVAEIV_RS05510 | 1141529 | 1141541 | 334 | - | 11.7143 | 4.15E-05 | 0.0753 | CAGCGCGCCGCCA | - | outer membrane protein assembly factor BamD |
| AVAEIV_RS07955 | 1655210 | 1655222 | 207 | - | 11.7143 | 4.15E-05 | 0.0753 | CAGCGCGCCGCCA | - | response regulator |
| AVAEIV_RS08720 | 1818629 | 1818641 | 285 | - | 11.7143 | 4.15E-05 | 0.0726 | CTGATCGCCGGCG | - | S1 RNA-binding domain-containing protein |
| AVAEIV_RS09645 | 2015193 | 2015205 | 25 | - | 11.7143 | 4.15E-05 | 0.0753 | CTGATCGCCGGCG | - | virulence factor family protein |
| AVAEIV_RS10910 | 2292594 | 2292606 | 62 | + | 11.7143 | 4.15E-05 | 0.0753 | CACTTCGCCACCT | - | ureidoglycolate lyase |
| AVAEIV_RS12345 | 2610071 | 2610083 | 264 | + | 11.7143 | 4.15E-05 | 0.0726 | CTGATCGCCGGCG | *cheA* | chemotaxis protein CheA |
| AVAEIV_RS19900 | 4241552 | 4241564 | 208 | - | 11.7143 | 4.15E-05 | 0.0753 | CCCATGGCCGTCG | - | multicopper oxidase family protein |
| AVAEIV_RS20560 | 4373120 | 4373132 | 354 | - | 11.7143 | 4.15E-05 | 0.0753 | CTGATCGCCGGCG | - | OmpA family protein |
| AVAEIV_RS22825 | 4888092 | 4888104 | -31 | - | 11.7143 | 4.15E-05 | 0.0753 | AACATCGCCATCA | *typA* | translational GTPase TypA |
| AVAEIV_RS23690 | 5047214 | 5047226 | 116 | + | 11.7143 | 4.15E-05 | 0.0726 | CTGATCGCCGGCG | - | TPM domain-containing protein |
| AVAEIV_RS04135 | 842071 | 842083 | 329 | - | 11.7 | 4.19E-05 | 0.0753 | CGGATCGCCGTCC | - | lytic murein transglycosylase |
| AVAEIV_RS04145 | 842071 | 842083 | 269 | - | 11.7 | 4.19E-05 | 0.0726 | CGGATCGCCGTCC | - | TolC family protein |
| AVAEIV_RS07160 | 1474852 | 1474864 | 253 | - | 11.7 | 4.19E-05 | 0.0726 | CAGTTCGCCGGCC | *era* | GTPase Era |
| AVAEIV_RS11410 | 2415415 | 2415427 | 153 | - | 11.7 | 4.19E-05 | 0.0753 | CAGATCGCCCGCA | - | NADP(H)-dependent aldo-keto reductase |

| | | | | | | | | | | |
|---|---|---|---|---|---|---|---|---|---|---|
| AVAEIV_RS12010 | 2552673 | 2552685 | 182 | + | 11.7 | 4.19E-05 | 0.0753 | CGGAATGCCGGCA | - | acetyl-CoA C-acetyltransferase |
| AVAEIV_RS13120 | 2846654 | 2846666 | 317 | + | 11.7 | 4.19E-05 | 0.0753 | CAGTTCGCCGGCC | - | ABC transporter ATP-binding protein |
| AVAEIV_RS13675 | 2957091 | 2957103 | 266 | - | 11.7 | 4.19E-05 | 0.0753 | CGCAACGCCAGCC | - | replication initiator protein A |
| AVAEIV_RS13825 | 2995077 | 2995089 | 238 | + | 11.7 | 4.19E-05 | 0.0753 | CGGATCGCCGTCC | - | Hcp family type VI secretion system effector |
| AVAEIV_RS17490 | 3788080 | 3788092 | 350 | - | 11.7 | 4.19E-05 | 0.0753 | CAGTTCGCCGGCC | - | DsbA family protein |
| AVAEIV_RS18495 | 3982362 | 3982374 | 163 | + | 11.7 | 4.19E-05 | 0.0753 | CCGTGCGCCGGCA | - | ACP phosphodiesterase |
| AVAEIV_RS18505 | 3982362 | 3982374 | 330 | + | 11.7 | 4.19E-05 | 0.0726 | CCGTGCGCCGGCA | - | alkene reductase |
| AVAEIV_RS19730 | 4205264 | 4205276 | 104 | + | 11.7 | 4.19E-05 | 0.0753 | CACATGGCCGCCG | - | hypothetical protein |
| AVAEIV_RS20480 | 4355094 | 4355106 | 78 | - | 11.7 | 4.19E-05 | 0.0753 | CGGATCGCCGTCC | *ettA* | energy-dependent translational throttle protein EttA |
| AVAEIV_RS20485 | 4355094 | 4355106 | 247 | - | 11.7 | 4.19E-05 | 0.0726 | CGGATCGCCGTCC | *glyA* | serine hydroxymethyltransferase |
| AVAEIV_RS22165 | 4729549 | 4729561 | 300 | + | 11.7 | 4.19E-05 | 0.0726 | CGGATCGCCGTCC | - | glycosyltransferase family 2 protein |
| AVAEIV_RS22770 | 4875601 | 4875613 | 299 | + | 11.7 | 4.19E-05 | 0.0726 | CGCTGGGCCACCA | - | pyruvate kinase |
| AVAEIV_RS25260 | 5380457 | 5380469 | 278 | - | 11.7 | 4.19E-05 | 0.0726 | CGGATTGCCGGCG | - | HD-GYP domain-containing protein |
| AVAEIV_RS26110 | 5613529 | 5613541 | 295 | + | 11.7 | 4.19E-05 | 0.0753 | CAGTTCGCCGGCC | - | tyrosine-type recombinase/integrase |
| AVAEIV_RS04650 | 952184 | 952196 | -1 | - | 11.6857 | 4.21E-05 | 0.0753 | CGGAGCGCCGCCA | - | histone deacetylase |
| AVAEIV_RS06180 | 1262110 | 1262122 | 333 | + | 11.6857 | 4.21E-05 | 0.0753 | CACCTGGCCGTCG | - | hypothetical protein |
| AVAEIV_RS09790 | 2048118 | 2048130 | -42 | - | 11.6857 | 4.21E-05 | 0.0753 | CGCCCCGCCAGCA | *egtB* | ergothioneine biosynthesis protein EgtB |
| AVAEIV_RS09795 | 2048118 | 2048130 | 293 | - | 11.6857 | 4.21E-05 | 0.0726 | CGCCCCGCCAGCA | - | protein/permease |
| AVAEIV_RS12170 | 2581577 | 2581589 | 217 | + | 11.6857 | 4.21E-05 | 0.0753 | GCGAGCGCCATCA | *flgA* | flagellar basal body P-ring formation chaperone FlgA |
| AVAEIV_RS12180 | 2581577 | 2581589 | 351 | + | 11.6857 | 4.21E-05 | 0.0726 | GCGAGCGCCATCA | *flgC* | flagellar basal body rod protein FlgC |
| AVAEIV_RS20700 | 4399802 | 4399814 | 362 | + | 11.6857 | 4.21E-05 | 0.0726 | CTGATGGCCAGCC | - | MerR family transcriptional regulator |
| AVAEIV_RS22915 | 4909900 | 4909912 | 387 | + | 11.6857 | 4.21E-05 | 0.0753 | CCCTTCGCCATGA | *tyrS* | tyrosine--tRNA ligase |
| AVAEIV_RS22920 | 4909900 | 4909912 | -4 | + | 11.6857 | 4.21E-05 | 0.0726 | CCCTTCGCCATGA | - | peptidoglycan DD-metalloendopeptidase family protein |
| AVAEIV_RS24010 | 5112611 | 5112623 | 67 | - | 11.6857 | 4.21E-05 | 0.0753 | CGGAGCGCCGCCA | *cyaY* | iron donor protein CyaY |
| AVAEIV_RS24015 | 5112611 | 5112623 | 90 | - | 11.6857 | 4.21E-05 | 0.0726 | CGGAGCGCCGCCA | *lptM* | LPS translocon maturation chaperone LptM |
| AVAEIV_RS24020 | 5112611 | 5112623 | 276 | - | 11.6857 | 4.21E-05 | 0.0726 | CGGAGCGCCGCCA | *lysA* | diaminopimelate decarboxylase |
| AVAEIV_RS24840 | 5282682 | 5282694 | 128 | + | 11.6857 | 4.21E-05 | 0.0726 | CCCTTCGCCATGA | - | tripartite tricarboxylate transporter TctB family protein |
| AVAEIV_RS01500 | 288931 | 288943 | 161 | - | 11.6714 | 4.25E-05 | 0.0753 | CACGTCGCCGTCG | - | YggS family pyridoxal phosphate-dependent enzyme |
| AVAEIV_RS04345 | 884618 | 884630 | 281 | - | 11.6714 | 4.25E-05 | 0.0753 | CGCCGCGCCAGCG | *leuS* | leucine--tRNA ligase |
| AVAEIV_RS04355 | 884618 | 884630 | 358 | - | 11.6714 | 4.25E-05 | 0.0728 | CGCCGCGCCAGCG | - | YdcF family protein |
| AVAEIV_RS10090 | 2107427 | 2107439 | 307 | - | 11.6714 | 4.25E-05 | 0.0728 | CGGCTCGCCACCC | - | F0F1 ATP synthase subunit C |
| AVAEIV_RS10975 | 2309255 | 2309267 | 120 | + | 11.6714 | 4.25E-05 | 0.0753 | CGGCGCGCCGTCA | - | substrate binding protein |
| AVAEIV_RS12215 | 2587366 | 2587378 | 345 | - | 11.6714 | 4.25E-05 | 0.0728 | CGCTCCGCCATCG | *flgJ* | flagellar assembly peptidoglycan hydrolase FlgJ |
| AVAEIV_RS14885 | 3238596 | 3238608 | 31 | + | 11.6714 | 4.25E-05 | 0.0753 | CGCCGCGCCAGCG | - | efflux transporter outer membrane subunit |
| AVAEIV_RS15890 | 3467302 | 3467314 | 309 | - | 11.6714 | 4.25E-05 | 0.0753 | CTGTACGCCACCG | - | ABC transporter ATP-binding protein |
| AVAEIV_RS16755 | 3627348 | 3627360 | -28 | - | 11.6714 | 4.25E-05 | 0.0753 | CGGATCGCCAGGG | - | NAD-dependent deacylase |
| AVAEIV_RS18655 | 4017303 | 4017315 | 341 | + | 11.6714 | 4.25E-05 | 0.0753 | CGGCGCGCCGTCA | *quiC* | 3-dehydroshikimate dehydratase QuiC |
| AVAEIV_RS18660 | 4017303 | 4017315 | 107 | + | 11.6714 | 4.25E-05 | 0.0728 | CGGCGCGCCGTCA | - | OprD family porin |
| AVAEIV_RS18935 | 4078382 | 4078394 | 353 | + | 11.6714 | 4.25E-05 | 0.0753 | CACGTCGCCGTCG | *truD* | tRNA pseudouridine(13) synthase TruD |
| AVAEIV_RS21030 | 4471697 | 4471709 | 15 | - | 11.6714 | 4.25E-05 | 0.0753 | CGGCTCGCCACCC | - | Fic family protein |
| AVAEIV_RS22120 | 4720691 | 4720703 | 93 | - | 11.6714 | 4.25E-05 | 0.0728 | CGCCGCGCCAGCG | - | diacylglycerol/lipid kinase family protein |
| AVAEIV_RS22315 | 4765151 | 4765163 | 295 | + | 11.6714 | 4.25E-05 | 0.0753 | CGCCGCGCCAGCG | - | glycosyltransferase family 25 protein |
| AVAEIV_RS25235 | 5370971 | 5370983 | 57 | - | 11.6714 | 4.25E-05 | 0.0753 | CGGCGCGCCGTCA | - | hydrogenase small subunit |
| AVAEIV_RS00535 | 111693 | 111705 | 379 | - | 11.6571 | 4.30E-05 | 0.0756 | CGCATGGCCGTCG | *prlC* | oligopeptidase A |
| AVAEIV_RS00545 | 111693 | 111705 | 373 | - | 11.6571 | 4.30E-05 | 0.073 | CGCATGGCCGTCG | - | FecR domain-containing protein |
| AVAEIV_RS02320 | 462190 | 462202 | 52 | + | 11.6571 | 4.30E-05 | 0.073 | CGCATGGCCGTCG | *hisH* | imidazole glycerol phosphate synthase subunit HisH |
| AVAEIV_RS02340 | 466538 | 466550 | 383 | + | 11.6571 | 4.30E-05 | 0.0756 | CAGGCGGCCATCG | - | S41 family peptidase |
| AVAEIV_RS04715 | 960012 | 960024 | 265 | - | 11.6571 | 4.30E-05 | 0.073 | CTGCGGGCCAGCA | - | AI-2E family transporter |
| AVAEIV_RS08085 | 1677947 | 1677959 | 279 | + | 11.6571 | 4.30E-05 | 0.0756 | CAGGCGGCCATCG | - | MurR/RpiR family transcriptional regulator |
| AVAEIV_RS09710 | 2033991 | 2034003 | 120 | - | 11.6571 | 4.30E-05 | 0.0756 | CAGGCGGCCATCG | - | phage holin family protein |
| AVAEIV_RS11130 | 2347959 | 2347971 | 208 | + | 11.6571 | 4.30E-05 | 0.073 | CAGGCGGCCATCG | - | ABC transporter permease |
| AVAEIV_RS19200 | 4137599 | 4137611 | 64 | + | 11.6571 | 4.30E-05 | 0.0756 | CAAACCGCCATCC | - | superinfection immunity protein |

| | | | | | | | | | | |
|---|---|---|---|---|---|---|---|---|---|---|
| AVAEIV_RS19205 | 4137599 | 4137611 | 97 | + | 11.6571 | 4.30E-05 | 0.073 | CAAACCGCCATCC | - | type II toxin-antitoxin system RelE/ParE family toxin |
| AVAEIV_RS20590 | 4380137 | 4380149 | 387 | - | 11.6571 | 4.30E-05 | 0.0756 | CAGGCGGCCATCG | - | protein |
| AVAEIV_RS25330 | 5397501 | 5397513 | 60 | + | 11.6571 | 4.30E-05 | 0.0756 | CTCGTTGCCATCA | - | hypothetical protein |
| AVAEIV_RS25335 | 5397501 | 5397513 | 114 | + | 11.6571 | 4.30E-05 | 0.073 | CTCGTTGCCATCA | *tssB* | subunit |
| AVAEIV_RS04040 | 820987 | 820999 | 66 | - | 11.6429 | 4.34E-05 | 0.0756 | CGGAACGCCAGTA | - | hypothetical protein |
| AVAEIV_RS04045 | 820987 | 820999 | 187 | - | 11.6429 | 4.34E-05 | 0.073 | CGGAACGCCAGTA | - | CidA/LrgA family protein |
| AVAEIV_RS04435 | 902767 | 902779 | -25 | + | 11.6429 | 4.34E-05 | 0.073 | CACAACGCCTTCA | - | aldolase/citrate lyase family protein |
| AVAEIV_RS04480 | 914803 | 914815 | -25 | - | 11.6429 | 4.34E-05 | 0.0756 | CTGACCGCCAGCG | - | MFS transporter |
| AVAEIV_RS05820 | 1193261 | 1193273 | 33 | - | 11.6429 | 4.34E-05 | 0.073 | CCGTTGGGCATCA | - | branched-chain amino acid ABC transporter permease |
| AVAEIV_RS06945 | 1429189 | 1429201 | 214 | - | 11.6429 | 4.34E-05 | 0.073 | CACAACGCCTTCA | - | cell division protein FtsQ/DivIB |
| AVAEIV_RS09325 | 1944850 | 1944862 | 228 | + | 11.6429 | 4.34E-05 | 0.073 | CGGTGCGCCGGCA | - | TSUP family transporter |
| AVAEIV_RS10000 | 2094997 | 2095009 | -46 | + | 11.6429 | 4.34E-05 | 0.073 | CTGACCGCCAGCG | - | Na+/H+ antiporter subunit C |
| AVAEIV_RS10005 | 2094997 | 2095009 | 280 | + | 11.6429 | 4.34E-05 | 0.073 | CTGACCGCCAGCG | - | monovalent cation/H+ antiporter subunit D |
| AVAEIV_RS12210 | 2586332 | 2586344 | 234 | - | 11.6429 | 4.34E-05 | 0.073 | CCGTTGGGCATCA | - | flagellar basal body P-ring protein FlgI |
| AVAEIV_RS14880 | 3237139 | 3237151 | 176 | - | 11.6429 | 4.34E-05 | 0.0756 | CTGACCGCCAGCG | - | fatty acid cis/trans isomerase |
| AVAEIV_RS16330 | 3551490 | 3551502 | 289 | - | 11.6429 | 4.34E-05 | 0.0756 | CTGACCGCCAGCG | - | type II toxin-antitoxin system HigB family toxin |
| AVAEIV_RS17065 | 3702042 | 3702054 | 127 | + | 11.6429 | 4.34E-05 | 0.0756 | CCCGGGGCCATCA | - | LacI family DNA-binding transcriptional regulator |
| AVAEIV_RS18715 | 4031127 | 4031139 | -38 | - | 11.6429 | 4.34E-05 | 0.0756 | CGGTGCGCCGGCA | *zapE* | cell division protein ZapE |
| AVAEIV_RS21365 | 4542816 | 4542828 | 328 | - | 11.6429 | 4.34E-05 | 0.073 | CGGTGCGCCGGCA | *folK* | diphosphokinase |
| AVAEIV_RS22275 | 4755680 | 4755692 | 131 | - | 11.6429 | 4.34E-05 | 0.0756 | CGGTGCGCCGGCA | *hldE* | phosphate kinase/D-glycero-beta-D-manno-heptose 1-phosphate adenylyltransferase HldE |
| AVAEIV_RS00560 | 116981 | 116993 | 180 | + | 11.6286 | 4.39E-05 | 0.0756 | CCGAACGGCAGCA | - | DUF1161 domain-containing protein |
| AVAEIV_RS05160 | 1060263 | 1060275 | 157 | - | 11.6286 | 4.39E-05 | 0.0756 | CGCCAGGCCACCA | - | DUF2231 domain-containing protein |
| AVAEIV_RS05940 | 1214146 | 1214158 | 370 | - | 11.6286 | 4.39E-05 | 0.0756 | CGCCTGGCCACCG | - | membrane protein |
| AVAEIV_RS05945 | 1214146 | 1214158 | 112 | - | 11.6286 | 4.39E-05 | 0.0756 | CGCCTGGCCACCG | - | protein |
| AVAEIV_RS06415 | 1310285 | 1310297 | 133 | + | 11.6286 | 4.39E-05 | 0.0733 | CGCCTGGCCACCG | - | PTS fructose-like transporter subunit IIB |
| AVAEIV_RS06695 | 1382180 | 1382192 | 174 | - | 11.6286 | 4.39E-05 | 0.0756 | CGCCAGGCCACCA | - | KdsC family phosphatase |
| AVAEIV_RS13445 | 2909107 | 2909119 | 323 | - | 11.6286 | 4.39E-05 | 0.0756 | GGGAGCGCCATCA | - | 2,3-butanediol dehydrogenase |
| AVAEIV_RS13595 | 2941025 | 2941037 | 277 | - | 11.6286 | 4.39E-05 | 0.0733 | CCGCACGCCGGCA | - | IS630 family transposase |
| AVAEIV_RS14895 | 3244984 | 3244996 | 131 | + | 11.6286 | 4.39E-05 | 0.0756 | CCGCTCGCCGGCG | - | permease subunit |
| AVAEIV_RS16975 | 3679882 | 3679894 | 339 | - | 11.6286 | 4.39E-05 | 0.0733 | CCGCACGCCGGCA | - | DUF3313 domain-containing protein |
| AVAEIV_RS18590 | 4003828 | 4003840 | 307 | - | 11.6286 | 4.39E-05 | 0.0733 | CAGTCCGCCAGCC | *yghX* | YghX family hydrolase |
| AVAEIV_RS18755 | 4037464 | 4037476 | 318 | - | 11.6286 | 4.39E-05 | 0.0733 | CCGAACGGCAGCA | - | stage II sporulation protein M |
| AVAEIV_RS20005 | 4265884 | 4265896 | 242 | - | 11.6286 | 4.39E-05 | 0.0756 | CCAATCGCCGGCG | - | LysR family transcriptional regulator |
| AVAEIV_RS20885 | 4440688 | 4440700 | 368 | - | 11.6286 | 4.39E-05 | 0.0756 | CGCCTGGCCACCG | *pqqA* | pyrroloquinoline quinone precursor peptide PqqA |
| AVAEIV_RS21480 | 4568822 | 4568834 | 74 | + | 11.6286 | 4.39E-05 | 0.0756 | CGCCTGGCCACCG | *greA* | transcription elongation factor GreA |
| AVAEIV_RS22710 | 4863189 | 4863201 | 219 | + | 11.6286 | 4.39E-05 | 0.0756 | CAGAAGGACAGCA | - | lipocalin family protein |
| AVAEIV_RS22715 | 4863189 | 4863201 | -32 | + | 11.6286 | 4.39E-05 | 0.0756 | CAGAAGGACAGCA | - | hypothetical protein |
| AVAEIV_RS02565 | 518327 | 518339 | 371 | - | 11.6 | 4.46E-05 | 0.076 | CAACTCGCCGGCG | - | cellulose biosynthesis protein BcsC |
| AVAEIV_RS03200 | 648973 | 648985 | 307 | + | 11.6 | 4.46E-05 | 0.0734 | CAGAAGGCCATGG | *rpmD* | 50S ribosomal protein L30 |
| AVAEIV_RS05650 | 1162343 | 1162355 | 214 | + | 11.6 | 4.46E-05 | 0.076 | CAACTCGCCGGCG | - | integrating conjugative element protein |
| AVAEIV_RS10285 | 2146137 | 2146149 | 232 | + | 11.6 | 4.46E-05 | 0.0734 | CAGTACGCCGCCG | *gltX* | glutamate--tRNA ligase |
| AVAEIV_RS11040 | 2326361 | 2326373 | -45 | - | 11.6 | 4.46E-05 | 0.0734 | CCCTTCGCCGCCA | - | RNA-binding domain-containing protein |
| AVAEIV_RS11815 | 2502890 | 2502902 | 146 | - | 11.6 | 4.46E-05 | 0.0734 | CAGTACGCCGCCG | *prpF* | 2-methylaconitate cis-trans isomerase PrpF |
| AVAEIV_RS13780 | 2982110 | 2982122 | 318 | - | 11.6 | 4.46E-05 | 0.076 | CAACTCGCCGGCG | - | DUF3540 domain-containing protein |
| AVAEIV_RS17970 | 3885710 | 3885722 | 214 | - | 11.6 | 4.46E-05 | 0.0734 | CAACTCGCCGGCG | - | integrating conjugative element protein |
| AVAEIV_RS23130 | 4942289 | 4942301 | 166 | + | 11.6 | 4.46E-05 | 0.0734 | CCGCTGGCCAGCC | - | DUF2590 family protein |
| AVAEIV_RS24965 | 5312785 | 5312797 | 112 | + | 11.6 | 4.46E-05 | 0.076 | CCGCTGGCCAGCC | *pyk* | pyruvate kinase |
| AVAEIV_RS05065 | 1042467 | 1042479 | 330 | - | 11.5857 | 4.50E-05 | 0.076 | CAGTTCGCCTCCA | - | sel1 repeat family protein |
| AVAEIV_RS06565 | 1354612 | 1354624 | 173 | - | 11.5857 | 4.50E-05 | 0.0734 | GCGATCGCCAGCG | - | response regulator transcription factor |
| AVAEIV_RS08310 | 1729359 | 1729371 | 183 | - | 11.5857 | 4.50E-05 | 0.0734 | CACTCGGCCAGCG | - | FAD/NAD(P)-binding protein |
| AVAEIV_RS08390 | 1751968 | 1751980 | 243 | + | 11.5857 | 4.50E-05 | 0.076 | CGCACGGCCATCG | - | AraC family transcriptional regulator |

| | | | | | | | | | | |
|---|---|---|---|---|---|---|---|---|---|---|
| AVAEIV_RS09315 | 1943713 | 1943725 | 84 | - | 11.5857 | 4.50E-05 | 0.076 | CAAATTGCCAGTA | - | DUF3597 domain-containing protein |
| AVAEIV_RS09320 | 1943713 | 1943725 | 143 | - | 11.5857 | 4.50E-05 | 0.0734 | CAAATTGCCAGTA | - | DUF1176 domain-containing protein |
| AVAEIV_RS11935 | 2535576 | 2535588 | 214 | - | 11.5857 | 4.50E-05 | 0.076 | CACTCGGCCAGCG | *lpxH* | UDP-2,3-diacylglucosamine diphosphatase |
| AVAEIV_RS17150 | 3717934 | 3717946 | 354 | + | 11.5857 | 4.50E-05 | 0.076 | CACTCGGCCAGCG | - | SDR family oxidoreductase |
| AVAEIV_RS18565 | 4000027 | 4000039 | 76 | + | 11.5857 | 4.50E-05 | 0.0734 | CCGGTCGCCAGCC | - | substrate-binding domain-containing protein |
| AVAEIV_RS19120 | 4123001 | 4123013 | 173 | + | 11.5857 | 4.50E-05 | 0.076 | GCGAACGCCAGCA | - | HNH endonuclease |
| AVAEIV_RS19675 | 4195336 | 4195348 | 279 | + | 11.5857 | 4.50E-05 | 0.076 | CCGTCGGCCACCA | - | ABC transporter permease subunit |
| AVAEIV_RS19845 | 4227879 | 4227891 | 40 | + | 11.5857 | 4.50E-05 | 0.076 | CAGCGGGCCGGCA | *ffh* | signal recognition particle protein |
| AVAEIV_RS19850 | 4227879 | 4227891 | 102 | + | 11.5857 | 4.50E-05 | 0.0734 | CAGCGGGCCGGCA | - | inner membrane protein YpjD |
| AVAEIV_RS22390 | 4787496 | 4787508 | 261 | + | 11.5857 | 4.50E-05 | 0.0734 | CCGGTCGCCAGCC | - | universal stress protein |
| AVAEIV_RS23700 | 5050142 | 5050154 | 227 | + | 11.5857 | 4.50E-05 | 0.076 | CGCATGGCCATCT | - | CinA family protein |
| AVAEIV_RS24880 | 5294434 | 5294446 | 229 | + | 11.5857 | 4.50E-05 | 0.076 | CACTCGGCCAGCG | - | isovaleryl-CoA dehydrogenase |
| AVAEIV_RS01100 | 217673 | 217685 | 168 | + | 11.5714 | 4.55E-05 | 0.076 | CGGATCGGCAGCG | - | TonB-dependent siderophore receptor |
| AVAEIV_RS04420 | 898421 | 898433 | 103 | - | 11.5714 | 4.55E-05 | 0.0734 | CAGGGCGCCGGCA | - | maltoporin |
| AVAEIV_RS05275 | 1089798 | 1089810 | 342 | - | 11.5714 | 4.55E-05 | 0.076 | CGGCACGCCGGCA | - | regulator |
| AVAEIV_RS05280 | 1089798 | 1089810 | 94 | - | 11.5714 | 4.55E-05 | 0.0734 | CGGCACGCCGGCA | - | CobW family GTP-binding protein |
| AVAEIV_RS06140 | 1252805 | 1252817 | 362 | - | 11.5714 | 4.55E-05 | 0.076 | CTGATCGCCAGTA | - | ATPase, T2SS/T4P/T4SS family |
| AVAEIV_RS06415 | 1310215 | 1310227 | 203 | - | 11.5714 | 4.55E-05 | 0.0734 | CGAATCGCCGGCG | - | PTS fructose-like transporter subunit IIB |
| AVAEIV_RS06745 | 1388798 | 1388810 | -45 | - | 11.5714 | 4.55E-05 | 0.0734 | CGGATCGGCAGCG | *hisD* | histidinol dehydrogenase |
| AVAEIV_RS06750 | 1390012 | 1390024 | 166 | - | 11.5714 | 4.55E-05 | 0.0734 | CCACGGGCCAGCA | *hisC* | histidinol-phosphate transaminase |
| AVAEIV_RS07285 | 1509904 | 1509916 | 253 | - | 11.5714 | 4.55E-05 | 0.076 | CCGATCGACACCA | - | bacterioferritin-associated ferredoxin |
| AVAEIV_RS10150 | 2119080 | 2119092 | 293 | + | 11.5714 | 4.55E-05 | 0.0734 | CCGATGGCGATCA | - | YbaB/EbfC family nucleoid-associated protein |
| AVAEIV_RS10925 | 2294991 | 2295003 | -48 | + | 11.5714 | 4.55E-05 | 0.076 | CGGCTCGCCGGCG | *puuE* | allantoinase PuuE |
| AVAEIV_RS10930 | 2294991 | 2295003 | 150 | + | 11.5714 | 4.55E-05 | 0.0734 | CGGCTCGCCGGCG | *uraH* | hydroxyisourate hydrolase |
| AVAEIV_RS11415 | 2416058 | 2416070 | 87 | + | 11.5714 | 4.55E-05 | 0.076 | CGGCTCGCCGGCG | - | containing protein |
| AVAEIV_RS11540 | 2441207 | 2441219 | 254 | - | 11.5714 | 4.55E-05 | 0.0734 | CGAAACGCCGGCA | - | hypothetical protein |
| AVAEIV_RS11645 | 2463844 | 2463856 | 215 | - | 11.5714 | 4.55E-05 | 0.076 | CGAATCGCCGGCG | - | MliC family protein |
| AVAEIV_RS11985 | 2542913 | 2542925 | -39 | - | 11.5714 | 4.55E-05 | 0.0734 | CAGTTTGCCATTG | *clpX* | ATP-dependent Clp protease ATP-binding subunit ClpX |
| AVAEIV_RS12100 | 2566703 | 2566715 | 207 | + | 11.5714 | 4.55E-05 | 0.0734 | CTGCAGGCCATCG | - | ribonuclease E inhibitor RraB |
| AVAEIV_RS12170 | 2581478 | 2581490 | 118 | + | 11.5714 | 4.55E-05 | 0.076 | TACTTTGCCATCA | *flgA* | flagellar basal body P-ring formation chaperone FlgA |
| AVAEIV_RS12175 | 2581478 | 2581490 | 28 | + | 11.5714 | 4.55E-05 | 0.0734 | TACTTTGCCATCA | *flgB* | flagellar basal body rod protein FlgB |
| AVAEIV_RS12925 | 2801193 | 2801205 | 229 | + | 11.5714 | 4.55E-05 | 0.0734 | CCACGGGCCAGCA | - | site-specific integrase |
| AVAEIV_RS16675 | 3612130 | 3612142 | 51 | + | 11.5714 | 4.55E-05 | 0.076 | CCGTGGGCCACCG | - | adenosylcobinamide-GDP ribazoletransferase |
| AVAEIV_RS16740 | 3624031 | 3624043 | 247 | - | 11.5714 | 4.55E-05 | 0.0734 | CAACTGGCCAGCC | - | glutamine amidotransferase |
| AVAEIV_RS20730 | 4407652 | 4407664 | 249 | + | 11.5714 | 4.55E-05 | 0.076 | CAGACCGCCGGCG | *hemA* | glutamyl-tRNA reductase |
| AVAEIV_RS21050 | 4476290 | 4476302 | 319 | - | 11.5714 | 4.55E-05 | 0.076 | CGGCTCGCCGGCG | - | OprD family porin |
| AVAEIV_RS21470 | 4567843 | 4567855 | 123 | + | 11.5714 | 4.55E-05 | 0.076 | CCATCGGCCATCA | *rlmE* | 23S rRNA (uridine(2552)-2'-O)-methyltransferase RlmE |
| AVAEIV_RS21475 | 4567843 | 4567855 | -50 | + | 11.5714 | 4.55E-05 | 0.0734 | CCATCGGCCATCA | *yhbY* | ribosome assembly RNA-binding protein YhbY |
| AVAEIV_RS22420 | 4794504 | 4794516 | 177 | + | 11.5714 | 4.55E-05 | 0.076 | CGGATCGGCAGCG | - | antitoxin |
| AVAEIV_RS22450 | 4799434 | 4799446 | 62 | - | 11.5714 | 4.55E-05 | 0.0734 | CAGGGCGCCGGCA | - | 2-hydroxymuconate tautomerase family protein |
| AVAEIV_RS23580 | 5026134 | 5026146 | -31 | + | 11.5714 | 4.55E-05 | 0.0734 | CAGATCGACGTCA | *apaG* | Co2+/Mg2+ efflux protein ApaG |
| AVAEIV_RS23585 | 5026134 | 5026146 | 349 | + | 11.5714 | 4.55E-05 | 0.0734 | CAGATCGACGTCA | - | symmetrical bis(5'-nucleosyl)-tetraphosphatase |
| AVAEIV_RS25455 | 5422120 | 5422132 | 86 | + | 11.5714 | 4.55E-05 | 0.0734 | CGGCACGCCGGCA | - | ArsC/Spx/MgsR family protein |
| AVAEIV_RS04620 | 944808 | 944820 | 341 | - | 11.5571 | 4.59E-05 | 0.076 | CTGGACGCCATCG | - | cation diffusion facilitator family transporter |
| AVAEIV_RS04630 | 944808 | 944820 | 242 | - | 11.5571 | 4.59E-05 | 0.0735 | CTGGACGCCATCG | *hrpB* | ATP-dependent helicase HrpB |
| AVAEIV_RS07675 | 1601847 | 1601859 | -45 | - | 11.5571 | 4.59E-05 | 0.0735 | CAGCGTGCCATCC | - | HAD-IA family hydrolase |
| AVAEIV_RS10140 | 2115785 | 2115797 | 269 | - | 11.5571 | 4.59E-05 | 0.0735 | CCGCCCGCCAGCG | *ffs* | signal recognition particle sRNA small type |
| AVAEIV_RS11060 | 2330946 | 2330958 | 205 | + | 11.5571 | 4.59E-05 | 0.0735 | CTGGACGCCATCG | - | FAD-binding oxidoreductase |
| AVAEIV_RS12250 | 2594684 | 2594696 | 142 | + | 11.5571 | 4.59E-05 | 0.076 | CGGTACGCCGTCG | *fliN* | flagellar motor switch protein FliN |
| AVAEIV_RS16755 | 3627549 | 3627561 | 173 | + | 11.5571 | 4.59E-05 | 0.076 | CCGCCCGCCAGCG | - | NAD-dependent deacylase |
| AVAEIV_RS17110 | 3711849 | 3711861 | 274 | - | 11.5571 | 4.59E-05 | 0.076 | CTGGACGCCATCG | - | metallophosphoesterase |
| AVAEIV_RS17150 | 3717850 | 3717862 | 270 | - | 11.5571 | 4.59E-05 | 0.076 | CCAGGCGCCAGCA | - | SDR family oxidoreductase |

| | | | | | | | | | | |
|---|---|---|---|---|---|---|---|---|---|---|
| AVAEIV_RS20690 | 4397753 | 4397765 | 234 | + | 11.5571 | 4.59E-05 | 0.0735 | CCCAGCGCCGGCA | - | NAD(P)/FAD-dependent oxidoreductase |
| AVAEIV_RS21220 | 4512554 | 4512566 | 221 | - | 11.5571 | 4.59E-05 | 0.076 | CCGCCCGCCAGCG | - | PqiB family protein |
| AVAEIV_RS21640 | 4606094 | 4606106 | 376 | + | 11.5571 | 4.59E-05 | 0.0735 | CCGCCCGCCAGCG | *glcF* | glycolate oxidase subunit GlcF |
| AVAEIV_RS23555 | 5020216 | 5020228 | 247 | - | 11.5571 | 4.59E-05 | 0.076 | CGGAGGGCCGGCA | - | aminoglycoside phosphotransferase family protein |
| AVAEIV_RS25095 | 5344188 | 5344200 | 259 | - | 11.5571 | 4.59E-05 | 0.076 | CTGGACGCCATCG | - | Gfo/Idh/MocA family protein |
| AVAEIV_RS02215 | 442766 | 442778 | 180 | + | 11.5429 | 4.64E-05 | 0.0736 | CGCTTCGCCGCCA | - | oxidoreductase |
| AVAEIV_RS04220 | 860147 | 860159 | 314 | + | 11.5429 | 4.64E-05 | 0.0736 | CGGGATGCCATCG | - | 2Fe-2S iron-sulfur cluster-binding protein |
| AVAEIV_RS06655 | 1376270 | 1376282 | 158 | - | 11.5429 | 4.64E-05 | 0.076 | CGCTTCGCCGCCA | - | HPr family phosphocarrier protein |
| AVAEIV_RS06765 | 1393206 | 1393218 | 214 | + | 11.5429 | 4.64E-05 | 0.0736 | CACCATGCCACCG | *cysD* | sulfate adenylyltransferase subunit CysD |
| AVAEIV_RS09520 | 1985883 | 1985895 | 231 | + | 11.5429 | 4.64E-05 | 0.076 | CGCTTCGCCGCCA | - | ParA family protein |
| AVAEIV_RS12275 | 2599781 | 2599793 | 283 | - | 11.5429 | 4.64E-05 | 0.076 | CCGCTTGCCGGCA | *fliI* | flagellar protein export ATPase FliI |
| AVAEIV_RS13475 | 2915029 | 2915041 | 251 | - | 11.5429 | 4.64E-05 | 0.076 | GAGTACGCCATCG | - | MFS transporter |
| AVAEIV_RS13580 | 2939087 | 2939099 | 15 | + | 11.5429 | 4.64E-05 | 0.076 | CATGAGGCCATCA | - | IS3 family transposase |
| AVAEIV_RS13965 | 3032208 | 3032220 | 351 | + | 11.5429 | 4.64E-05 | 0.0736 | CCGCTTGCCGGCA | - | sugar dehydrogenase complex small subunit |
| AVAEIV_RS14980 | 3264366 | 3264378 | 149 | + | 11.5429 | 4.64E-05 | 0.076 | CCGCTTGCCGGCA | - | DUF3363 domain-containing protein |
| AVAEIV_RS14985 | 3264366 | 3264378 | 322 | + | 11.5429 | 4.64E-05 | 0.0736 | CCGCTTGCCGGCA | - | peptidylprolyl isomerase |
| AVAEIV_RS19005 | 4094558 | 4094570 | -28 | - | 11.5429 | 4.64E-05 | 0.076 | CCTCGCGCCATCA | *lpxA* | acyl-ACP--UDP-N-acetylglucosamine O-acyltransferase |
| AVAEIV_RS21435 | 4561904 | 4561916 | 76 | + | 11.5429 | 4.64E-05 | 0.076 | CCGCTTGCCGGCA | *rimP* | ribosome maturation factor RimP |
| AVAEIV_RS22345 | 4772275 | 4772287 | 45 | + | 11.5429 | 4.64E-05 | 0.076 | CACCATGCCACCG | *rfaP* | lipopolysaccharide core heptose(I) kinase RfaP |
| AVAEIV_RS02035 | 407172 | 407184 | -16 | - | 11.5286 | 4.68E-05 | 0.076 | GGGATCGCCAGCG | - | hypothetical protein |
| AVAEIV_RS04870 | 993427 | 993439 | 1 | + | 11.5286 | 4.68E-05 | 0.0736 | CGGAGTGCCATCC | - | protein |
| AVAEIV_RS05815 | 1192287 | 1192299 | 132 | + | 11.5286 | 4.68E-05 | 0.0736 | AAGATCGCCATCG | - | branched-chain amino acid ABC transporter permease |
| AVAEIV_RS06785 | 1397448 | 1397460 | 385 | - | 11.5286 | 4.68E-05 | 0.0736 | CGCTTCGGCATCA | - | tryptophan--tRNA ligase |
| AVAEIV_RS07445 | 1548864 | 1548876 | 304 | + | 11.5286 | 4.68E-05 | 0.0736 | AAGAACGCCATCA | - | efflux RND transporter permease subunit |
| AVAEIV_RS07450 | 1551947 | 1551959 | 331 | + | 11.5286 | 4.68E-05 | 0.0736 | AAGAACGCCATCA | - | efflux transporter outer membrane subunit |
| AVAEIV_RS07485 | 1560510 | 1560522 | 257 | - | 11.5286 | 4.68E-05 | 0.0736 | CACAGCGGCAGCA | *fadA* | acetyl-CoA C-acyltransferase FadA |
| AVAEIV_RS12975 | 2811285 | 2811297 | 37 | - | 11.5286 | 4.68E-05 | 0.076 | GGGATCGCCAGCG | - | nucleotidyltransferase family protein |
| AVAEIV_RS15215 | 3313770 | 3313782 | 332 | + | 11.5286 | 4.68E-05 | 0.076 | CCCTCCGCCACCA | - | ABC transporter ATP-binding protein |
| AVAEIV_RS19120 | 4123012 | 4123024 | 184 | + | 11.5286 | 4.68E-05 | 0.076 | CAGGTGGCCGCCA | - | HNH endonuclease |
| AVAEIV_RS03910 | 787985 | 787997 | 55 | + | 11.5143 | 4.72E-05 | 0.076 | CCGCTCGCCAGGA | *adhP* | alcohol dehydrogenase AdhP |
| AVAEIV_RS05250 | 1080313 | 1080325 | 288 | + | 11.5143 | 4.72E-05 | 0.076 | TCGATCGCCAGCG | - | hypothetical protein |
| AVAEIV_RS06665 | 1377582 | 1377594 | 31 | - | 11.5143 | 4.72E-05 | 0.076 | TCGAACGCCAGCA | *ptsN* | PTS IIA-like nitrogen regulatory protein PtsN |
| AVAEIV_RS08165 | 1696999 | 1697011 | 360 | - | 11.5143 | 4.72E-05 | 0.0736 | TCGATCGCCAGCG | *cmk* | (d)CMP kinase |
| AVAEIV_RS08845 | 1834336 | 1834348 | 253 | + | 11.5143 | 4.72E-05 | 0.0736 | CATTTCGCCAGCC | - | CitMHS family transporter |
| AVAEIV_RS10365 | 2167124 | 2167136 | 27 | - | 11.5143 | 4.72E-05 | 0.076 | TCGAACGCCAGCA | - | NAD(P)-dependent alcohol dehydrogenase |
| AVAEIV_RS13100 | 2839484 | 2839496 | 375 | - | 11.5143 | 4.72E-05 | 0.0736 | CGGATGGCGATCA | - | L-serine ammonia-lyase |
| AVAEIV_RS14170 | 3083796 | 3083808 | -19 | + | 11.5143 | 4.72E-05 | 0.076 | TCGATCGCCAGCG | - | DUF2149 domain-containing protein |
| AVAEIV_RS18805 | 4049441 | 4049453 | 313 | - | 11.5143 | 4.72E-05 | 0.0736 | TCGAACGCCAGCA | - | hypothetical protein |
| AVAEIV_RS22975 | 4921052 | 4921064 | 166 | - | 11.5143 | 4.72E-05 | 0.076 | CAGAAGGCCGCCG | *trpC* | indole-3-glycerol phosphate synthase TrpC |
| AVAEIV_RS23935 | 5099441 | 5099453 | 301 | - | 11.5143 | 4.72E-05 | 0.076 | CCTTGCGCCAGCA | - | TerC family protein |
| AVAEIV_RS01005 | 198260 | 198272 | -30 | + | 11.5 | 4.77E-05 | 0.076 | CAGATCGCCGGTA | *zwf* | glucose-6-phosphate dehydrogenase |
| AVAEIV_RS01010 | 198260 | 198272 | 137 | + | 11.5 | 4.77E-05 | 0.0736 | CAGATCGCCGGTA | *hexR* | transcriptional regulator HexR |
| AVAEIV_RS01140 | 223902 | 223914 | -46 | - | 11.5 | 4.77E-05 | 0.076 | CTGCTCGCCGGCA | - | endonuclease I family protein |
| AVAEIV_RS01145 | 223902 | 223914 | 208 | - | 11.5 | 4.77E-05 | 0.0736 | CTGCTCGCCGGCA | - | hypothetical protein |
| AVAEIV_RS01360 | 265960 | 265972 | 217 | - | 11.5 | 4.77E-05 | 0.076 | CTGCTCGCCGGCA | - | RidA family protein |
| AVAEIV_RS03395 | 682744 | 682756 | 232 | + | 11.5 | 4.77E-05 | 0.076 | CGCAGCGCCGGCA | - | hypothetical protein |
| AVAEIV_RS03420 | 690649 | 690661 | 320 | + | 11.5 | 4.77E-05 | 0.076 | CAGCAGGCCGTCG | *fis* | DNA-binding transcriptional regulator Fis |
| AVAEIV_RS04785 | 974208 | 974220 | -50 | + | 11.5 | 4.77E-05 | 0.0736 | CGGCCCGCCAGCG | - | malonate decarboxylase subunit delta |
| AVAEIV_RS04790 | 974208 | 974220 | 242 | + | 11.5 | 4.77E-05 | 0.0736 | CGGCCCGCCAGCG | - | beta |
| AVAEIV_RS05045 | 1036559 | 1036571 | 385 | - | 11.5 | 4.77E-05 | 0.076 | CTGCTCGCCGGCA | - | mannuronate-specific alginate lyase |
| AVAEIV_RS06995 | 1443243 | 1443255 | -5 | + | 11.5 | 4.77E-05 | 0.076 | CCCATGGGCATCA | - | ATP-binding protein |
| AVAEIV_RS11405 | 2413499 | 2413511 | 169 | - | 11.5 | 4.77E-05 | 0.0736 | CAGCAGGCCGTCG | - | GNAT family N-acetyltransferase |

| | | | | | | | | | | |
|---|---|---|---|---|---|---|---|---|---|---|
| AVAEIV_RS12265 | 2597859 | 2597871 | 202 | - | 11.5 | 4.77E-05 | 0.076 | CGCAGCGCCGGCA | - | flagellar hook-length control protein FliK |
| AVAEIV_RS12635 | 2680281 | 2680293 | 104 | + | 11.5 | 4.77E-05 | 0.076 | CGCAGCGCCGGCA | - | protein |
| AVAEIV_RS14135 | 3075849 | 3075861 | 265 | + | 11.5 | 4.77E-05 | 0.076 | CTGATCGGCAGCA | - | ABC transporter ATP-binding protein |
| AVAEIV_RS14635 | 3184544 | 3184556 | 205 | + | 11.5 | 4.77E-05 | 0.0736 | CTGCTCGCCGGCA | *purB* | adenylosuccinate lyase |
| AVAEIV_RS15275 | 3335161 | 3335173 | 209 | - | 11.5 | 4.77E-05 | 0.0736 | CAGCAGGCCGTCG | - | extracellular solute-binding protein |
| AVAEIV_RS15340 | 3354149 | 3354161 | 325 | - | 11.5 | 4.77E-05 | 0.076 | CAGAAGGGCATCG | - | succinate dehydrogenase iron-sulfur subunit |
| AVAEIV_RS17035 | 3692545 | 3692557 | 317 | + | 11.5 | 4.77E-05 | 0.0736 | CTGCTCGCCGGCA | - | efflux RND transporter periplasmic adaptor subunit |
| AVAEIV_RS17525 | 3794539 | 3794551 | 218 | + | 11.5 | 4.77E-05 | 0.0736 | CGCAGCGCCGGCA | - | EAL domain-containing protein |
| AVAEIV_RS17805 | 3852551 | 3852563 | 50 | + | 11.5 | 4.77E-05 | 0.0736 | CCCATGGGCATCA | - | SDR family NAD(P)-dependent oxidoreductase |
| AVAEIV_RS19990 | 4261546 | 4261558 | 368 | + | 11.5 | 4.77E-05 | 0.076 | CGGCCCGCCAGCG | - | nitrogenase component 1 |
| AVAEIV_RS19995 | 4262306 | 4262318 | 53 | + | 11.5 | 4.77E-05 | 0.076 | CAGCAGGCCGTCG | - | ABC transporter substrate-binding protein |
| AVAEIV_RS21205 | 4507649 | 4507661 | 167 | + | 11.5 | 4.77E-05 | 0.076 | CGCAGCGCCGGCA | *rimI* | ribosomal protein S18-alanine N-acetyltransferase |
| AVAEIV_RS21780 | 4638452 | 4638464 | 330 | - | 11.5 | 4.77E-05 | 0.076 | CCGCAGGCCACCG | - | ABC transporter permease |
| AVAEIV_RS21825 | 4650209 | 4650221 | 298 | + | 11.5 | 4.77E-05 | 0.076 | CAGCAGGCCGTCG | - | ABC transporter ATP-binding protein |
| AVAEIV_RS21855 | 4657519 | 4657531 | -46 | - | 11.5 | 4.77E-05 | 0.076 | CTGCTCGCCGGCA | - | branched-chain amino acid ABC transporter permease |
| AVAEIV_RS22240 | 4746167 | 4746179 | 370 | + | 11.5 | 4.77E-05 | 0.0736 | CGCAGCGCCGGCA | - | NAD(P)/FAD-dependent oxidoreductase |
| AVAEIV_RS24405 | 5186381 | 5186393 | 313 | - | 11.5 | 4.77E-05 | 0.0736 | CCGCAGGCCACCG | *phoR* | phosphate regulon sensor histidine kinase PhoR |
| AVAEIV_RS25145 | 5354546 | 5354558 | 163 | + | 11.5 | 4.77E-05 | 0.0736 | CTGCTCGCCGGCA | - | HupE/UreJ family protein |
| AVAEIV_RS25810 | 5530531 | 5530543 | 351 | + | 11.5 | 4.77E-05 | 0.0736 | CCGCAGGCCACCG | - | glutathione S-transferase family protein |
| AVAEIV_RS00865 | 169436 | 169448 | 322 | - | 11.4857 | 4.81E-05 | 0.076 | GGCTTCGCCATCA | - | AEC family transporter |
| AVAEIV_RS01455 | 281144 | 281156 | 121 | + | 11.4857 | 4.81E-05 | 0.0736 | CTCTACGCCAGCG | *trmB* | tRNA (guanosine(46)-N7)-methyltransferase TrmB |
| AVAEIV_RS02045 | 408294 | 408306 | 351 | - | 11.4857 | 4.81E-05 | 0.076 | CACATTGACAGCA | - | helix-turn-helix domain-containing protein |
| AVAEIV_RS03185 | 647927 | 647939 | -48 | - | 11.4857 | 4.81E-05 | 0.0736 | CTCTACGCCAGCG | *rplF* | 50S ribosomal protein L6 |
| AVAEIV_RS05425 | 1121859 | 1121871 | 43 | - | 11.4857 | 4.81E-05 | 0.0736 | CCGAATGCCAGCC | *murJ* | murein biosynthesis integral membrane protein MurJ |
| AVAEIV_RS05475 | 1132099 | 1132111 | 230 | - | 11.4857 | 4.81E-05 | 0.0736 | CGGTACGCCATCT | - | hypothetical protein |
| AVAEIV_RS12015 | 2553443 | 2553455 | 146 | - | 11.4857 | 4.81E-05 | 0.076 | CCAATCGCCAGTA | *phbB* | acetoacetyl-CoA reductase |
| AVAEIV_RS12020 | 2553443 | 2553455 | 290 | - | 11.4857 | 4.81E-05 | 0.0736 | CCAATCGCCAGTA | - | AraC family transcriptional regulator |
| AVAEIV_RS15170 | 3302384 | 3302396 | 339 | + | 11.4857 | 4.81E-05 | 0.0736 | CTCTACGCCAGCG | - | ABC transporter permease |
| AVAEIV_RS17090 | 3707185 | 3707197 | 223 | - | 11.4857 | 4.81E-05 | 0.076 | CAGACTGCCGGCA | - | hypothetical protein |
| AVAEIV_RS17200 | 3730182 | 3730194 | 330 | - | 11.4857 | 4.81E-05 | 0.076 | CGCGAGGCCAGCA | - | aspartate-semialdehyde dehydrogenase |
| AVAEIV_RS22275 | 4755862 | 4755874 | 313 | + | 11.4857 | 4.81E-05 | 0.076 | CCACAGGCCATCG | *hldE* | phosphate kinase/D-glycero-beta-D-manno-heptose 1-phosphate adenylyltransferase HldE |
| AVAEIV_RS22350 | 4773430 | 4773442 | -16 | - | 11.4857 | 4.81E-05 | 0.076 | CAGCTGGCCTTCA | - | glycosyltransferase family 4 protein |
| AVAEIV_RS22590 | 4838852 | 4838864 | 344 | + | 11.4857 | 4.81E-05 | 0.0736 | CGCGTGGCCAGCG | - | SPOR domain-containing protein |
| AVAEIV_RS23065 | 4934617 | 4934629 | 127 | + | 11.4857 | 4.81E-05 | 0.0736 | CAGTTCGCCACGG | - | phage tail protein |
| AVAEIV_RS24465 | 5200412 | 5200424 | -41 | - | 11.4857 | 4.81E-05 | 0.0736 | GACAGCGCCAGCA | - | OmpA family protein |
| AVAEIV_RS25360 | 5402424 | 5402436 | 52 | + | 11.4857 | 4.81E-05 | 0.0736 | CTCTACGCCAGCG | *tssG* | type VI secretion system baseplate subunit TssG |
| AVAEIV_RS25880 | 5552851 | 5552863 | 269 | - | 11.4857 | 4.81E-05 | 0.076 | CAGAGCGACATCG | - | hypothetical protein |
| AVAEIV_RS00600 | 123115 | 123127 | 66 | - | 11.4714 | 4.85E-05 | 0.076 | CCGCCTGCCAGCA | *pbpG* | D-alanyl-D-alanine endopeptidase |
| AVAEIV_RS00605 | 123115 | 123127 | 112 | - | 11.4714 | 4.85E-05 | 0.0738 | CCGCCTGCCAGCA | - | DUF3299 domain-containing protein |
| AVAEIV_RS02365 | 469816 | 469828 | 166 | - | 11.4714 | 4.85E-05 | 0.0738 | CGGTTTGCCGTCG | *secB* | protein-export chaperone SecB |
| AVAEIV_RS05170 | 1060921 | 1060933 | 137 | - | 11.4714 | 4.85E-05 | 0.0738 | CAGAGTGCCGGCG | *ctaD* | cytochrome c oxidase subunit I |
| AVAEIV_RS12060 | 2561213 | 2561225 | 178 | + | 11.4714 | 4.85E-05 | 0.076 | CTGGTTGCCATCG | - | alpha/beta hydrolase |
| AVAEIV_RS15240 | 3329008 | 3329020 | 42 | - | 11.4714 | 4.85E-05 | 0.076 | CGGAAGGCCGTCG | - | containing protein |
| AVAEIV_RS15655 | 3427546 | 3427558 | 247 | + | 11.4714 | 4.85E-05 | 0.076 | CGGAAGGCCGTCG | *ccmA* | CcmA |
| AVAEIV_RS16390 | 3560509 | 3560521 | 370 | + | 11.4714 | 4.85E-05 | 0.076 | CCGATGGCCAACA | - | holin family protein |
| AVAEIV_RS16395 | 3560509 | 3560521 | 50 | + | 11.4714 | 4.85E-05 | 0.076 | CCGATGGCCAACA | - | hypothetical protein |
| AVAEIV_RS16410 | 3564384 | 3564396 | 270 | + | 11.4714 | 4.85E-05 | 0.076 | CACGTCGCCGCCA | - | alpha/beta hydrolase |
| AVAEIV_RS16700 | 3617150 | 3617162 | 179 | + | 11.4714 | 4.85E-05 | 0.076 | CCGGCGGCCATCA | *cobD* | threonine-phosphate decarboxylase CobD |
| AVAEIV_RS16720 | 3620921 | 3620933 | 312 | + | 11.4714 | 4.85E-05 | 0.076 | CAGTAGGCCGGCG | *cobO* | cob(I)yrinic acid a,c-diamide adenosyltransferase |
| AVAEIV_RS16945 | 3671294 | 3671306 | 278 | + | 11.4714 | 4.85E-05 | 0.076 | CAATTCGCCATGG | - | gluconate 2-dehydrogenase subunit 3 family protein |
| AVAEIV_RS19005 | 4094540 | 4094552 | -46 | - | 11.4714 | 4.85E-05 | 0.076 | CCGAGCGCCACCC | *lpxA* | acyl-ACP--UDP-N-acetylglucosamine O-acyltransferase |

| | | | | | | | | | | |
|---|---|---|---|---|---|---|---|---|---|---|
| AVAEIV_RS22560 | 4828617 | 4828629 | 297 | + | 11.4714 | 4.85E-05 | 0.0738 | CGGTTCGACAGCA | - | penicillin-binding protein 1A |
| AVAEIV_RS22935 | 4915115 | 4915127 | 366 | - | 11.4714 | 4.85E-05 | 0.076 | CGCTATGCCAGCG | *erpA* | iron-sulfur cluster insertion protein ErpA |
| AVAEIV_RS23040 | 4931564 | 4931576 | 143 | + | 11.4714 | 4.85E-05 | 0.076 | CCGAGCGCCACCC | - | terminase large subunit domain-containing protein |
| AVAEIV_RS23045 | 4931564 | 4931576 | 3 | + | 11.4714 | 4.85E-05 | 0.0738 | CCGAGCGCCACCC | - | GPO family capsid scaffolding protein |
| AVAEIV_RS24455 | 5200054 | 5200066 | 350 | + | 11.4714 | 4.85E-05 | 0.076 | CAGTTCGCCCTCA | - | catalase |
| AVAEIV_RS24465 | 5200054 | 5200066 | 317 | + | 11.4714 | 4.85E-05 | 0.0738 | CAGTTCGCCCTCA | - | OmpA family protein |
| AVAEIV_RS26120 | 5616706 | 5616718 | 180 | + | 11.4714 | 4.85E-05 | 0.076 | TGGTTGGCCATCA | *yidC* | membrane protein insertase YidC |
| AVAEIV_RS00620 | 127174 | 127186 | 357 | - | 11.4571 | 4.90E-05 | 0.076 | CAGAACGCCTTCG | *modC* | molybdenum ABC transporter ATP-binding protein |
| AVAEIV_RS05900 | 1208772 | 1208784 | 379 | - | 11.4571 | 4.90E-05 | 0.076 | CGCATCGCGATCA | - | protein |
| AVAEIV_RS07335 | 1520815 | 1520827 | 254 | - | 11.4571 | 4.90E-05 | 0.076 | CAGGCGGCCACCA | *mobA* | molybdenum cofactor guanylyltransferase MobA |
| AVAEIV_RS07345 | 1520815 | 1520827 | 350 | - | 11.4571 | 4.90E-05 | 0.0741 | CAGGCGGCCACCA | *glp* | gephyrin-like molybdotransferase Glp |
| AVAEIV_RS10295 | 2150527 | 2150539 | 321 | - | 11.4571 | 4.90E-05 | 0.076 | CAGAAAGCCAGCG | - | isocitrate lyase/PEP mutase family protein |
| AVAEIV_RS13980 | 3033848 | 3033860 | 297 | + | 11.4571 | 4.90E-05 | 0.0741 | TGGAACGCCAGCA | - | hypothetical protein |
| AVAEIV_RS15050 | 3277700 | 3277712 | 119 | + | 11.4571 | 4.90E-05 | 0.076 | TGGAACGCCAGCA | - | GspMb/PilO family protein |
| AVAEIV_RS15145 | 3298611 | 3298623 | 381 | + | 11.4571 | 4.90E-05 | 0.076 | GCCATGGCCATCA | - | ArsC/Spx/MgsR family protein |
| AVAEIV_RS20980 | 4463207 | 4463219 | -23 | + | 11.4571 | 4.90E-05 | 0.076 | CAGACCGCCAGGA | - | TolC family protein |
| AVAEIV_RS21485 | 4572267 | 4572279 | 301 | - | 11.4571 | 4.90E-05 | 0.076 | GCCATGGCCATCA | *carB* | carbamoyl-phosphate synthase large subunit |
| AVAEIV_RS22190 | 4736636 | 4736648 | 93 | + | 11.4571 | 4.90E-05 | 0.076 | CGTTGCGCCAGCA | - | acyl-CoA dehydrogenase family protein |
| AVAEIV_RS22805 | 4882182 | 4882194 | 210 | - | 11.4571 | 4.90E-05 | 0.0741 | GAGAAGGCCATCG | - | IS5 family transposase |
| AVAEIV_RS24575 | 5220043 | 5220055 | 338 | - | 11.4571 | 4.90E-05 | 0.0741 | CATATCGCCGGCG | - | glutamine synthetase family protein |
| AVAEIV_RS25265 | 5383303 | 5383315 | 351 | - | 11.4571 | 4.90E-05 | 0.076 | CAGAACGCCTTCG | *modC* | molybdenum ABC transporter ATP-binding protein |
| AVAEIV_RS25385 | 5409749 | 5409761 | 9 | - | 11.4571 | 4.90E-05 | 0.076 | CGAATCGCCAGGA | - | DUF4280 domain-containing protein |
| AVAEIV_RS25470 | 5423360 | 5423372 | 245 | + | 11.4571 | 4.90E-05 | 0.0741 | CCGCGCGCCATCC | *grxD* | Grx4 family monothiol glutaredoxin |
| AVAEIV_RS00275 | 58927 | 58939 | -50 | - | 11.4429 | 4.94E-05 | 0.0743 | CCGATGGACAGCA | - | hypothetical protein |
| AVAEIV_RS01280 | 251065 | 251077 | 348 | + | 11.4429 | 4.94E-05 | 0.0761 | GACATGGCCACCA | *vnfG* | V-containing nitrogenase subunit delta |
| AVAEIV_RS01810 | 358248 | 358260 | 321 | + | 11.4429 | 4.94E-05 | 0.0761 | CAGCGCGCCACCC | - | ABC transporter substrate-binding protein |
| AVAEIV_RS06175 | 1260476 | 1260488 | 32 | - | 11.4429 | 4.94E-05 | 0.0761 | CAGTCTGCCACCG | - | TcpQ domain-containing protein |
| AVAEIV_RS09940 | 2082237 | 2082249 | 213 | - | 11.4429 | 4.94E-05 | 0.0761 | CTGAACGCCAGCC | - | YchJ family protein |
| AVAEIV_RS10135 | 2114823 | 2114835 | 187 | - | 11.4429 | 4.94E-05 | 0.0743 | CTCAATGCCACCA | - | hydroxypyruvate isomerase family protein |
| AVAEIV_RS10920 | 2294266 | 2294278 | 153 | + | 11.4429 | 4.94E-05 | 0.0761 | CCGATGGACAGCA | *uraD* | decarboxylase |
| AVAEIV_RS11645 | 2463833 | 2463845 | 204 | + | 11.4429 | 4.94E-05 | 0.0761 | CACCACGCCGTCG | - | MliC family protein |
| AVAEIV_RS11720 | 2483149 | 2483161 | 77 | - | 11.4429 | 4.94E-05 | 0.0761 | CTGAACGCCAGCC | *greB* | transcription elongation factor GreB |
| AVAEIV_RS13255 | 2874104 | 2874116 | 299 | + | 11.4429 | 4.94E-05 | 0.0743 | CGAAAGGCCACCG | - | aminotransferase class V-fold PLP-dependent enzyme |
| AVAEIV_RS17335 | 3754225 | 3754237 | 227 | + | 11.4429 | 4.94E-05 | 0.0761 | CGCCTGGCCGTCA | - | hypothetical protein |
| AVAEIV_RS20905 | 4444517 | 4444529 | 60 | - | 11.4429 | 4.94E-05 | 0.0743 | CCGATGGACAGCA | - | thiamine pyrophosphate-dependent dehydrogenase E1 component subunit alpha |
| AVAEIV_RS20980 | 4463285 | 4463297 | 55 | + | 11.4429 | 4.94E-05 | 0.0761 | CAAAGCGCCACCC | - | TolC family protein |
| AVAEIV_RS21475 | 4567461 | 4567473 | 332 | - | 11.4429 | 4.94E-05 | 0.0743 | CCGATGGACAGCA | *yhbY* | ribosome assembly RNA-binding protein YhbY |
| AVAEIV_RS24485 | 5204139 | 5204151 | 362 | + | 11.4429 | 4.94E-05 | 0.0743 | CAGCGCGCCACCC | - | ankyrin repeat domain-containing protein |
| AVAEIV_RS00020 | 3608 | 3620 | 70 | - | 11.4286 | 4.99E-05 | 0.0748 | CCGTGCGCCAGCC | *gyrB* | DNA topoisomerase (ATP-hydrolyzing) subunit B |
| AVAEIV_RS01765 | 348106 | 348118 | 152 | - | 11.4286 | 4.99E-05 | 0.0765 | CGACAGGCCATCG | *dctP* | C4-dicarboxylate TRAP substrate-binding protein DctP |
| AVAEIV_RS08515 | 1779529 | 1779541 | 46 | + | 11.4286 | 4.99E-05 | 0.0765 | CAGTGGGCCATTA | - | LemA family protein |
| AVAEIV_RS08520 | 1779529 | 1779541 | 93 | + | 11.4286 | 4.99E-05 | 0.0748 | CAGTGGGCCATTA | - | hypothetical protein |
| AVAEIV_RS15325 | 3348858 | 3348870 | 154 | + | 11.4286 | 4.99E-05 | 0.0765 | CTCCTTGCCATCG | *lpdA* | dihydrolipoyl dehydrogenase |
| AVAEIV_RS19860 | 4231478 | 4231490 | 96 | + | 11.4286 | 4.99E-05 | 0.0765 | CACAATGCCATTA | *purT* | formyltransferase |
| AVAEIV_RS01170 | 227576 | 227588 | 24 | - | 11.4143 | 5.03E-05 | 0.0765 | CCCGCCGCCATCA | - | alpha/beta fold hydrolase |
| AVAEIV_RS01175 | 227576 | 227588 | 156 | - | 11.4143 | 5.03E-05 | 0.0752 | CCCGCCGCCATCA | - | AsmA family protein |
| AVAEIV_RS04565 | 934511 | 934523 | 209 | - | 11.4143 | 5.03E-05 | 0.0765 | CGCAACGCCGTCG | - | tellurite resistance TerB family protein |
| AVAEIV_RS04765 | 970856 | 970868 | 298 | - | 11.4143 | 5.03E-05 | 0.0765 | TACAGCGCCAGCA | *folX* | dihydroneopterin triphosphate 2'-epimerase |
| AVAEIV_RS16335 | 3551979 | 3551991 | 263 | - | 11.4143 | 5.03E-05 | 0.0765 | CCGTTCGCCGCCG | - | helix-turn-helix domain-containing protein |
| AVAEIV_RS16340 | 3551979 | 3551991 | 138 | - | 11.4143 | 5.03E-05 | 0.0765 | CCGTTCGCCGCCG | - | type II toxin-antitoxin system HigB family toxin |
| AVAEIV_RS18340 | 3947346 | 3947358 | 358 | - | 11.4143 | 5.03E-05 | 0.0765 | CTGCGCGCCAGCG | - | DUF2058 domain-containing protein |

| | | | | | | | | | | |
|---|---|---|---|---|---|---|---|---|---|---|
| AVAEIV_RS19645 | 4186463 | 4186475 | 348 | + | 11.4143 | 5.03E-05 | 0.0752 | CTGCGCGCCAGCG | - | glycosyltransferase |
| AVAEIV_RS19775 | 4214440 | 4214452 | 140 | - | 11.4143 | 5.03E-05 | 0.0765 | CCGTTCGCCGCCG | - | NADH:flavin oxidoreductase/NADH oxidase |
| AVAEIV_RS23550 | 5019146 | 5019158 | 196 | - | 11.4143 | 5.03E-05 | 0.0765 | CTGCGCGCCAGCG | *murU* | uridylyltransferase MurU |
| AVAEIV_RS00320 | 71114 | 71126 | 344 | - | 11.4 | 5.09E-05 | 0.0765 | CTCAAGGCCAGCG | - | alpha/beta fold hydrolase |
| AVAEIV_RS00560 | 117049 | 117061 | 248 | + | 11.4 | 5.09E-05 | 0.0765 | CCATTCGCCGTCG | - | DUF1161 domain-containing protein |
| AVAEIV_RS01325 | 258271 | 258283 | 385 | - | 11.4 | 5.09E-05 | 0.0752 | CGGCGCGCCATCC | - | MoaD/ThiS family protein |
| AVAEIV_RS01495 | 288022 | 288034 | -25 | - | 11.4 | 5.09E-05 | 0.0765 | CGCATCGCCTTCA | *proC* | pyrroline-5-carboxylate reductase |
| AVAEIV_RS04245 | 864560 | 864572 | 371 | - | 11.4 | 5.09E-05 | 0.0752 | CGGCGCGCCATCC | - | LysR family transcriptional regulator |
| AVAEIV_RS06400 | 1306669 | 1306681 | 244 | - | 11.4 | 5.09E-05 | 0.0765 | CCCTCGGCCAGCA | *cra* | catabolite repressor/activator |
| AVAEIV_RS06500 | 1339873 | 1339885 | 241 | - | 11.4 | 5.09E-05 | 0.0765 | CTGAAGGCCGTCA | - | Fe2+-dependent dioxygenase |
| AVAEIV_RS07235 | 1499101 | 1499113 | 66 | - | 11.4 | 5.09E-05 | 0.0765 | CCCGGCGCCATCG | - | transglycosylase domain-containing protein |
| AVAEIV_RS07380 | 1531302 | 1531314 | 250 | - | 11.4 | 5.09E-05 | 0.0752 | CCGGTTGCCACCG | - | acyl-CoA thioesterase |
| AVAEIV_RS09035 | 1880186 | 1880198 | 278 | + | 11.4 | 5.09E-05 | 0.0765 | CCATTCGCCGTCG | - | CoA transferase subunit B |
| AVAEIV_RS11065 | 2332306 | 2332318 | 260 | - | 11.4 | 5.09E-05 | 0.0752 | CAGGTTGCCGTCG | - | dependent enzyme |
| AVAEIV_RS12840 | 2770574 | 2770586 | 227 | + | 11.4 | 5.09E-05 | 0.0765 | CGGCGCGCCATCC | - | non-ribosomal peptide synthase/polyketide synthase |
| AVAEIV_RS13145 | 2851756 | 2851768 | 53 | + | 11.4 | 5.09E-05 | 0.0765 | CAGTAGGCCAGCT | *hutG* | N-formylglutamate deformylase |
| AVAEIV_RS15090 | 3283479 | 3283491 | 20 | - | 11.4 | 5.09E-05 | 0.0765 | CGAAGCGCCATCC | - | carbonate dehydratase |
| AVAEIV_RS18010 | 3896280 | 3896292 | 206 | + | 11.4 | 5.09E-05 | 0.0765 | CGGCGCGCCATCC | - | hypothetical protein |
| AVAEIV_RS18805 | 4049383 | 4049395 | 371 | + | 11.4 | 5.09E-05 | 0.0752 | CTGATGGCCGTCG | - | hypothetical protein |
| AVAEIV_RS19865 | 4232358 | 4232370 | 88 | - | 11.4 | 5.09E-05 | 0.0765 | CAGGTTGCCGTCG | - | CoA pyrophosphatase |
| AVAEIV_RS19870 | 4232358 | 4232370 | 11 | - | 11.4 | 5.09E-05 | 0.0752 | CAGGTTGCCGTCG | - | NUDIX hydrolase |
| AVAEIV_RS20095 | 4290860 | 4290872 | 212 | + | 11.4 | 5.09E-05 | 0.0752 | CCCTCGGCCAGCA | - | endonuclease I family protein |
| AVAEIV_RS20240 | 4318893 | 4318905 | 184 | + | 11.4 | 5.09E-05 | 0.0765 | CAGGTTGCCGTCG | *secF* | protein translocase subunit SecF |
| AVAEIV_RS20765 | 4411998 | 4412010 | 172 | + | 11.4 | 5.09E-05 | 0.0752 | CAGTTGGACATCG | - | 50S ribosomal protein L25/general stress protein Ctc |
| AVAEIV_RS21570 | 4588272 | 4588284 | 169 | + | 11.4 | 5.09E-05 | 0.0752 | CTCAAGGCCAGCG | - | protein |
| AVAEIV_RS25040 | 5330349 | 5330361 | 345 | + | 11.4 | 5.09E-05 | 0.0752 | CCCTCGGCCAGCA | *hppD* | 4-hydroxyphenylpyruvate dioxygenase |
| AVAEIV_RS25440 | 5418195 | 5418207 | 315 | + | 11.4 | 5.09E-05 | 0.0752 | GACAACGCCATCG | *nifA* | nif-specific transcriptional activator NifA |
| AVAEIV_RS06235 | 1274464 | 1274476 | 208 | - | 11.3857 | 5.15E-05 | 0.0767 | CAATTCGCCGCCG | - | DUF2857 domain-containing protein |
| AVAEIV_RS06535 | 1349157 | 1349169 | 68 | + | 11.3857 | 5.15E-05 | 0.0767 | TTGATCGCCAGCA | - | ABC transporter ATP-binding protein |
| AVAEIV_RS08100 | 1679007 | 1679019 | 293 | - | 11.3857 | 5.15E-05 | 0.0755 | CGCCACGCCACCG | *rluB* | 23S rRNA pseudouridine(2605) synthase RluB |
| AVAEIV_RS08190 | 1702661 | 1702673 | 289 | - | 11.3857 | 5.15E-05 | 0.0755 | CAGTTCGGCACCG | *rfaH* | protein RfaH |
| AVAEIV_RS12275 | 2599520 | 2599532 | 22 | - | 11.3857 | 5.15E-05 | 0.0767 | CCCATCGACAGCA | *fliI* | flagellar protein export ATPase FliI |
| AVAEIV_RS18695 | 4027713 | 4027725 | 370 | - | 11.3857 | 5.15E-05 | 0.0767 | GAGATCGCCGGCA | *pcaF* | 3-oxoadipyl-CoA thiolase |
| AVAEIV_RS19455 | 4164955 | 4164967 | 219 | + | 11.3857 | 5.15E-05 | 0.0767 | CGCCACGCCACCG | - | hypothetical protein |
| AVAEIV_RS19460 | 4164955 | 4164967 | -45 | + | 11.3857 | 5.15E-05 | 0.0767 | CGCCACGCCACCG | - | hypothetical protein |
| AVAEIV_RS19465 | 4164955 | 4164967 | 73 | + | 11.3857 | 5.15E-05 | 0.0755 | CGCCACGCCACCG | - | S24 family peptidase |
| AVAEIV_RS20440 | 4345449 | 4345461 | 75 | + | 11.3857 | 5.15E-05 | 0.0767 | CGCACGGCCACCA | - | TraR/DksA family transcriptional regulator |
| AVAEIV_RS20445 | 4345449 | 4345461 | 96 | + | 11.3857 | 5.15E-05 | 0.0755 | CGCACGGCCACCA | - | DUF3015 domain-containing protein |
| AVAEIV_RS21970 | 4686270 | 4686282 | 326 | + | 11.3857 | 5.15E-05 | 0.0755 | CCGACCGCCGGCA | - | acetate/propionate family kinase |
| AVAEIV_RS23555 | 5020102 | 5020114 | 133 | + | 11.3857 | 5.15E-05 | 0.0767 | CAGATGGCCCTCA | - | aminoglycoside phosphotransferase family protein |
| AVAEIV_RS23560 | 5020102 | 5020114 | -10 | + | 11.3857 | 5.15E-05 | 0.0755 | CAGATGGCCCTCA | - | LPS-assembly protein LptD |
| AVAEIV_RS23820 | 5076886 | 5076898 | 135 | - | 11.3857 | 5.15E-05 | 0.0767 | GAGATCGCCGGCA | - | ABC transporter permease |
| AVAEIV_RS24220 | 5149787 | 5149799 | 375 | - | 11.3857 | 5.15E-05 | 0.0767 | CAGTTGGCCGTCC | - | dihydrofolate reductase |
| AVAEIV_RS24690 | 5245703 | 5245715 | 232 | + | 11.3857 | 5.15E-05 | 0.0755 | CCGACCGCCGGCA | - | hypothetical protein |
| AVAEIV_RS25355 | 5400525 | 5400537 | 194 | + | 11.3857 | 5.15E-05 | 0.0755 | CCGACCGCCGGCA | *tssF* | type VI secretion system baseplate subunit TssF |
| AVAEIV_RS00010 | 1255 | 1267 | 198 | + | 11.3714 | 5.19E-05 | 0.0758 | CAGGTCGCCATGG | *dnaN* | DNA polymerase III subunit beta |
| AVAEIV_RS04920 | 1004291 | 1004303 | 194 | - | 11.3714 | 5.19E-05 | 0.0769 | CTGCTGGCCACCG | *exbD* | TonB system transport protein ExbD |
| AVAEIV_RS05220 | 1073352 | 1073364 | 300 | + | 11.3714 | 5.19E-05 | 0.0769 | CCGAGCGCCGGCG | *rhlB* | ATP-dependent RNA helicase RhlB |
| AVAEIV_RS12245 | 2594382 | 2594394 | 304 | - | 11.3714 | 5.19E-05 | 0.0769 | CAGAACGCCGGCC | *fliO* | flagellar biosynthetic protein FliO |
| AVAEIV_RS12620 | 2678812 | 2678824 | 232 | - | 11.3714 | 5.19E-05 | 0.0769 | CGCCCGGCCATCA | - | MFS transporter |
| AVAEIV_RS16915 | 3663121 | 3663133 | 102 | - | 11.3714 | 5.19E-05 | 0.0769 | CAGATTGCCTTCG | - | integrase core domain-containing protein |
| AVAEIV_RS16920 | 3663121 | 3663133 | 3 | - | 11.3714 | 5.19E-05 | 0.0758 | CAGATTGCCTTCG | - | containing protein |

| | | | | | | | | | | |
|---|---|---|---|---|---|---|---|---|---|---|
| AVAEIV_RS21555 | 4585118 | 4585130 | 386 | + | 11.3714 | 5.19E-05 | 0.0758 | CTGCAGGCCACCA | - | (Fe-S)-binding protein |
| AVAEIV_RS21665 | 4612279 | 4612291 | 303 | + | 11.3714 | 5.19E-05 | 0.0758 | CTGCAGGCCACCA | - | malic enzyme-like NAD(P)-binding protein |
| AVAEIV_RS26055 | 5603808 | 5603820 | 184 | - | 11.3714 | 5.19E-05 | 0.0769 | CTGCTGGCCACCG | - | protein |
| AVAEIV_RS02415 | 480055 | 480067 | 216 | - | 11.3571 | 5.24E-05 | 0.0769 | CTGGTCGCCACCG | - | DUF2388 domain-containing protein |
| AVAEIV_RS03005 | 619713 | 619725 | 290 | + | 11.3571 | 5.24E-05 | 0.0758 | GCGTTCGCCATCG | *birA* | bifunctional biotin--[acetyl-CoA-carboxylase] ligase/biotin operon repressor BirA |
| AVAEIV_RS03405 | 687091 | 687103 | 323 | + | 11.3571 | 5.24E-05 | 0.0769 | CACCTTGCCGTCG | - | hybrid sensor histidine kinase/response regulator |
| AVAEIV_RS03720 | 749649 | 749661 | 172 | + | 11.3571 | 5.24E-05 | 0.0758 | CTGGTCGCCACCG | *hflC* | protease modulator HflC |
| AVAEIV_RS03950 | 796962 | 796974 | 198 | - | 11.3571 | 5.24E-05 | 0.0769 | CTGGTCGCCACCG | - | ExbD/TolR family protein |
| AVAEIV_RS04990 | 1020747 | 1020759 | 298 | - | 11.3571 | 5.24E-05 | 0.0758 | CAGCCCGCCGGCA | *pdhA* | component subunit alpha |
| AVAEIV_RS06505 | 1339539 | 1339551 | 257 | + | 11.3571 | 5.24E-05 | 0.0758 | CACCTTGCCGTCG | - | TonB-dependent receptor |
| AVAEIV_RS06940 | 1428180 | 1428192 | 272 | - | 11.3571 | 5.24E-05 | 0.0758 | CACATTGGCATCG | - | D-alanine--D-alanine ligase |
| AVAEIV_RS08550 | 1785194 | 1785206 | 83 | + | 11.3571 | 5.24E-05 | 0.0758 | CCATGGGCCACCA | - | copper resistance protein B |
| AVAEIV_RS11160 | 2357339 | 2357351 | 198 | - | 11.3571 | 5.24E-05 | 0.0769 | CTGGTCGCCACCG | - | ExbD/TolR family protein |
| AVAEIV_RS12095 | 2565721 | 2565733 | 141 | - | 11.3571 | 5.24E-05 | 0.0758 | CAGCCCGCCGGCA | *speE* | polyamine aminopropyltransferase |
| AVAEIV_RS15405 | 3364004 | 3364016 | 199 | + | 11.3571 | 5.24E-05 | 0.0758 | CACCTCGACAGCA | - | polysaccharide biosynthesis tyrosine autokinase |
| AVAEIV_RS15615 | 3421786 | 3421798 | 283 | - | 11.3571 | 5.24E-05 | 0.0769 | CCCATCGCCATGG | *ccmI* | c-type cytochrome biogenesis protein CcmI |
| AVAEIV_RS17650 | 3824300 | 3824312 | 118 | - | 11.3571 | 5.24E-05 | 0.0769 | CAGTTGGCCAGGG | - | hypothetical protein |
| AVAEIV_RS19275 | 4144624 | 4144636 | 116 | + | 11.3571 | 5.24E-05 | 0.0769 | CAGACCGGCAGCA | - | phage tail protein |
| AVAEIV_RS20785 | 4415894 | 4415906 | 377 | + | 11.3571 | 5.24E-05 | 0.0769 | CGCCGGGCCATCG | - | ATP-binding protein |
| AVAEIV_RS21715 | 4627056 | 4627068 | 68 | + | 11.3571 | 5.24E-05 | 0.0769 | CGGTTCGCCGCCG | - | YqiA/YcfP family alpha/beta fold hydrolase |
| AVAEIV_RS23555 | 5020324 | 5020336 | 355 | + | 11.3571 | 5.24E-05 | 0.0769 | CAGCCCGCCGGCA | - | aminoglycoside phosphotransferase family protein |
| AVAEIV_RS25245 | 5374049 | 5374061 | 257 | - | 11.3571 | 5.24E-05 | 0.0758 | CGCATCGCCAACA | *ltrA* | group II intron reverse transcriptase/maturase |
| AVAEIV_RS26065 | 5605487 | 5605499 | 121 | + | 11.3571 | 5.24E-05 | 0.0769 | CCGCACGCCAGCC | - | DUF4433 domain-containing protein |
| AVAEIV_RS02550 | 512725 | 512737 | 378 | + | 11.3429 | 5.29E-05 | 0.0769 | CCGAGGGCCAGCC | - | hypothetical protein |
| AVAEIV_RS02900 | 595859 | 595871 | 241 | + | 11.3429 | 5.29E-05 | 0.0761 | CGGTTCGGCATCG | - | substrate binding protein |
| AVAEIV_RS04510 | 923399 | 923411 | 175 | - | 11.3429 | 5.29E-05 | 0.0769 | CGCTCGGCCAGCA | *ureG* | urease accessory protein UreG |
| AVAEIV_RS04580 | 938181 | 938193 | 320 | + | 11.3429 | 5.29E-05 | 0.0769 | CGCTCGGCCAGCA | - | urease subunit beta |
| AVAEIV_RS05030 | 1032442 | 1032454 | 299 | + | 11.3429 | 5.29E-05 | 0.0769 | CAGCGCGCCGGCG | - | alginate O-acetyltransferase AlgF |
| AVAEIV_RS05510 | 1141493 | 1141505 | 298 | - | 11.3429 | 5.29E-05 | 0.0769 | CGCGGCGCCATCG | - | outer membrane protein assembly factor BamD |
| AVAEIV_RS06255 | 1277294 | 1277306 | 171 | - | 11.3429 | 5.29E-05 | 0.0769 | GAACTCGCCAGCA | - | putative protein CP11 |
| AVAEIV_RS07450 | 1551978 | 1551990 | 300 | - | 11.3429 | 5.29E-05 | 0.0761 | CGCTCGGCCAGCA | - | efflux transporter outer membrane subunit |
| AVAEIV_RS14190 | 3089899 | 3089911 | 338 | - | 11.3429 | 5.29E-05 | 0.0761 | CAGGTCGCCATTG | - | DUF2970 domain-containing protein |
| AVAEIV_RS15995 | 3487927 | 3487939 | 275 | + | 11.3429 | 5.29E-05 | 0.0769 | CAGCGCGCCGGCG | - | ATP-binding protein |
| AVAEIV_RS19755 | 4209721 | 4209733 | 226 | - | 11.3429 | 5.29E-05 | 0.0769 | CGATTCGCCGTCG | - | TetR/AcrR family transcriptional regulator |
| AVAEIV_RS20885 | 4440668 | 4440680 | 348 | + | 11.3429 | 5.29E-05 | 0.0769 | CAGCGCGCCGGCG | *pqqA* | pyrroloquinoline quinone precursor peptide PqqA |
| AVAEIV_RS22205 | 4739473 | 4739485 | 329 | + | 11.3429 | 5.29E-05 | 0.0769 | CAGTCCGCCGTCG | - | ABC transporter ATP-binding protein |
| AVAEIV_RS23135 | 4942610 | 4942622 | 159 | - | 11.3429 | 5.29E-05 | 0.0761 | CGCTCGGCCAGCA | - | baseplate J/gp47 family protein |
| AVAEIV_RS24410 | 5187856 | 5187868 | 218 | + | 11.3429 | 5.29E-05 | 0.0761 | CGCTCGGCCAGCA | - | hemolysin family protein |
| AVAEIV_RS01555 | 298995 | 299007 | 209 | - | 11.3286 | 5.35E-05 | 0.0775 | AAGATCGCCACCA | - | 16S rRNA (uracil(1498)-N(3))-methyltransferase |
| AVAEIV_RS05790 | 1187620 | 1187632 | 373 | - | 11.3286 | 5.35E-05 | 0.0775 | CCACGCGCCAGCG | - | enoyl-CoA hydratase |
| AVAEIV_RS06375 | 1299643 | 1299655 | 247 | - | 11.3286 | 5.35E-05 | 0.0762 | CCGAAGGCCGCCA | *ampD* | AmpD |
| AVAEIV_RS08845 | 1834484 | 1834496 | 105 | - | 11.3286 | 5.35E-05 | 0.0762 | CGCATCGACAGCA | - | CitMHS family transporter |
| AVAEIV_RS08860 | 1840220 | 1840232 | 167 | + | 11.3286 | 5.35E-05 | 0.0762 | CACTTTGCCGGCG | *casA* | type I-E CRISPR-associated protein Cse1/CasA |
| AVAEIV_RS09975 | 2089652 | 2089664 | -49 | + | 11.3286 | 5.35E-05 | 0.0762 | TGCATGGCCATCA | - | OmpA family protein |
| AVAEIV_RS13470 | 2913630 | 2913642 | 376 | + | 11.3286 | 5.35E-05 | 0.0775 | CCGATGGCCGCCG | - | hypothetical protein |
| AVAEIV_RS17285 | 3745394 | 3745406 | 77 | - | 11.3286 | 5.35E-05 | 0.0775 | CCCATCGCCATTG | - | tRNA-Arg |
| AVAEIV_RS17290 | 3745394 | 3745406 | 262 | - | 11.3286 | 5.35E-05 | 0.0762 | CCCATCGCCATTG | - | Arc family DNA-binding protein |
| AVAEIV_RS18955 | 4081712 | 4081724 | 188 | + | 11.3286 | 5.35E-05 | 0.0775 | CAGGCTGCCATCG | *ispD* | cytidylyltransferase |
| AVAEIV_RS23580 | 5025975 | 5025987 | 128 | - | 11.3286 | 5.35E-05 | 0.0762 | CAGGCTGCCATCG | *apaG* | Co2+/Mg2+ efflux protein ApaG |
| AVAEIV_RS02250 | 450607 | 450619 | -22 | - | 11.3143 | 5.41E-05 | 0.0779 | AAAATCGCCATCA | - | AAA family ATPase |
| AVAEIV_RS04485 | 916746 | 916758 | 167 | + | 11.3143 | 5.41E-05 | 0.0779 | CCAATGGCCGTCG | - | IucA/IucC family protein |

| | | | | | | | | | | |
|---|---|---|---|---|---|---|---|---|---|---|
| AVAEIV_RS08510 | 1779117 | 1779129 | 246 | + | 11.3143 | 5.41E-05 | 0.0779 | GAGACCGCCAGCA | - | TPM domain-containing protein |
| AVAEIV_RS08955 | 1861856 | 1861868 | 374 | - | 11.3143 | 5.41E-05 | 0.0762 | CCGCTGGCCGTCG | - | acyl-CoA dehydrogenase |
| AVAEIV_RS09380 | 1957066 | 1957078 | 344 | - | 11.3143 | 5.41E-05 | 0.0779 | CCCACTGCCATCG | - | IS4-like element ISAzvi7 family transposase |
| AVAEIV_RS10520 | 2199686 | 2199698 | 182 | - | 11.3143 | 5.41E-05 | 0.0762 | CCCCAGGCCAGCG | - | hypothetical protein |
| AVAEIV_RS11575 | 2447538 | 2447550 | 160 | - | 11.3143 | 5.41E-05 | 0.0762 | CCGCTGGCCGTCG | - | hypothetical protein |
| AVAEIV_RS13785 | 2983122 | 2983134 | 222 | + | 11.3143 | 5.41E-05 | 0.0779 | CCGCTGGCCGTCG | - | pentapeptide repeat-containing protein |
| AVAEIV_RS17655 | 3824857 | 3824869 | 334 | + | 11.3143 | 5.41E-05 | 0.0779 | CTCCACGCCACCA | - | helix-turn-helix domain-containing protein |
| AVAEIV_RS18215 | 3929540 | 3929552 | 0 | - | 11.3143 | 5.41E-05 | 0.0762 | CAGTGCGCCTTCA | *purN* | phosphoribosylglycinamide formyltransferase |
| AVAEIV_RS25710 | 5499721 | 5499733 | 276 | - | 11.3143 | 5.41E-05 | 0.0762 | CCGATGGGCATCG | - | substrate-binding domain-containing protein |
| AVAEIV_RS00800 | 157035 | 157047 | 56 | + | 11.3 | 5.46E-05 | 0.0762 | CCGGTCGCCGTCG | - | hypothetical protein |
| AVAEIV_RS08465 | 1769182 | 1769194 | 292 | - | 11.3 | 5.46E-05 | 0.0762 | CAGCTGGCCGCCG | - | heavy metal translocating P-type ATPase |
| AVAEIV_RS08705 | 1815734 | 1815746 | 339 | + | 11.3 | 5.46E-05 | 0.0762 | CCGGTCGCCGTCG | - | SDR family NAD(P)-dependent oxidoreductase |
| AVAEIV_RS09260 | 1934705 | 1934717 | 234 | - | 11.3 | 5.46E-05 | 0.078 | CCCGACGCCAGCG | - | group II truncated hemoglobin |
| AVAEIV_RS11125 | 2346981 | 2346993 | 296 | - | 11.3 | 5.46E-05 | 0.0762 | CAGCAGGCCGCCA | - | class II aldolase/adducin family protein |
| AVAEIV_RS12375 | 2621583 | 2621595 | 52 | + | 11.3 | 5.46E-05 | 0.078 | CTCAGGGCCACCA | - | dipeptide ABC transporter ATP-binding protein |
| AVAEIV_RS15215 | 3313613 | 3313625 | 175 | + | 11.3 | 5.46E-05 | 0.078 | CCGGTCGCCGTCG | - | ABC transporter ATP-binding protein |
| AVAEIV_RS18355 | 3950476 | 3950488 | 373 | + | 11.3 | 5.46E-05 | 0.078 | CAGCAGGCCGCCA | - | PepSY-associated TM helix domain-containing protein |
| AVAEIV_RS18360 | 3950476 | 3950488 | 158 | + | 11.3 | 5.46E-05 | 0.078 | CAGCAGGCCGCCA | - | hypothetical protein |
| AVAEIV_RS19660 | 4191194 | 4191206 | 323 | - | 11.3 | 5.46E-05 | 0.0762 | CAGCAGGCCGCCA | - | hypothetical protein |
| AVAEIV_RS23815 | 5075141 | 5075153 | 126 | - | 11.3 | 5.46E-05 | 0.078 | CGGCACGCCAGCC | *gcvT* | glycine cleavage system aminomethyltransferase GcvT |
| AVAEIV_RS24145 | 5134917 | 5134929 | 295 | - | 11.3 | 5.46E-05 | 0.078 | CCGGTCGCCGTCG | - | cupin domain-containing protein |
| AVAEIV_RS24190 | 5143023 | 5143035 | 372 | + | 11.3 | 5.46E-05 | 0.078 | CAGGGCGCCAGCC | *rpmB* | 50S ribosomal protein L28 |
| AVAEIV_RS25430 | 5416673 | 5416685 | 158 | + | 11.3 | 5.46E-05 | 0.078 | CGGCACGCCAGCC | *rsxA* | electron transport complex subunit RsxA |
| AVAEIV_RS25435 | 5416673 | 5416685 | 258 | + | 11.3 | 5.46E-05 | 0.0762 | CGGCACGCCAGCC | *nifL* | nitrogen fixation negative regulator NifL |
| AVAEIV_RS04190 | 854231 | 854243 | 91 | - | 11.2857 | 5.51E-05 | 0.0762 | CAGGTCGCCGCCG | - | MFS transporter |
| AVAEIV_RS04570 | 935795 | 935807 | 347 | + | 11.2857 | 5.51E-05 | 0.078 | CAACTGGCCGTCG | - | domain-containing protein |
| AVAEIV_RS04985 | 1019627 | 1019639 | 265 | - | 11.2857 | 5.51E-05 | 0.0762 | CAGGTCGCCGCCG | - | Leu/Phe/Val dehydrogenase |
| AVAEIV_RS05110 | 1051618 | 1051630 | -10 | - | 11.2857 | 5.51E-05 | 0.078 | CAAATGGGCATCG | *cydB* | cytochrome d ubiquinol oxidase subunit II |
| AVAEIV_RS07230 | 1492346 | 1492358 | 381 | - | 11.2857 | 5.51E-05 | 0.0762 | CAGGTCGCCGCCG | *putA* | bifunctional proline dehydrogenase/L-glutamate gamma-semialdehyde dehydrogenase PutA |
| AVAEIV_RS08010 | 1663162 | 1663174 | 371 | + | 11.2857 | 5.51E-05 | 0.0762 | CGGTCCGCCACCG | *queF* | QueF |
| AVAEIV_RS08075 | 1673761 | 1673773 | 228 | + | 11.2857 | 5.51E-05 | 0.0762 | CCGTTGGCCGGCG | *edd* | phosphogluconate dehydratase |
| AVAEIV_RS08640 | 1803175 | 1803187 | 385 | + | 11.2857 | 5.51E-05 | 0.078 | CGGTTCGCCACCT | - | SRPBCC family protein |
| AVAEIV_RS09280 | 1938689 | 1938701 | 120 | - | 11.2857 | 5.51E-05 | 0.078 | CAGGTCGCCGCCG | - | hypothetical protein |
| AVAEIV_RS09285 | 1938689 | 1938701 | 149 | - | 11.2857 | 5.51E-05 | 0.0762 | CAGGTCGCCGCCG | - | hypothetical protein |
| AVAEIV_RS09765 | 2042584 | 2042596 | -49 | - | 11.2857 | 5.51E-05 | 0.078 | CTGTTCGCCGCCA | - | peptidylprolyl isomerase |
| AVAEIV_RS09875 | 2068115 | 2068127 | 118 | - | 11.2857 | 5.51E-05 | 0.078 | CTGTTCGCCGCCA | *rsxB* | electron transport complex subunit RsxB |
| AVAEIV_RS12985 | 2814568 | 2814580 | 22 | - | 11.2857 | 5.51E-05 | 0.078 | CAGGTCGCCGCCG | - | binding subunit |
| AVAEIV_RS13925 | 3019139 | 3019151 | 338 | + | 11.2857 | 5.51E-05 | 0.0762 | CTGTTCGCCGCCA | - | sugar phosphate isomerase/epimerase family protein |
| AVAEIV_RS15275 | 3335207 | 3335219 | 163 | + | 11.2857 | 5.51E-05 | 0.0762 | CAGGTCGCCGCCG | - | extracellular solute-binding protein |
| AVAEIV_RS16720 | 3620900 | 3620912 | 291 | + | 11.2857 | 5.51E-05 | 0.078 | CCGTAGGCCGGCA | *cobO* | cob(I)yrinic acid a,c-diamide adenosyltransferase |
| AVAEIV_RS17000 | 3686139 | 3686151 | -11 | + | 11.2857 | 5.51E-05 | 0.078 | CAGGTTGCCATGA | - | short-chain fatty acid transporter |
| AVAEIV_RS17615 | 3816158 | 3816170 | 288 | + | 11.2857 | 5.51E-05 | 0.0762 | CCCAGCGCCAGCC | *minC* | septum site-determining protein MinC |
| AVAEIV_RS20480 | 4355396 | 4355408 | 380 | + | 11.2857 | 5.51E-05 | 0.078 | CTGTTCGCCGCCA | *ettA* | energy-dependent translational throttle protein EttA |
| AVAEIV_RS22015 | 4699731 | 4699743 | 324 | + | 11.2857 | 5.51E-05 | 0.078 | CAACTGGCCGTCG | - | TonB-dependent siderophore receptor |
| AVAEIV_RS24275 | 5162949 | 5162961 | 124 | + | 11.2857 | 5.51E-05 | 0.078 | CCCAGCGCCAGCC | *ptsP* | phosphoenolpyruvate--protein phosphotransferase |
| AVAEIV_RS02205 | 441625 | 441637 | 29 | - | 11.2714 | 5.57E-05 | 0.0762 | CGGATGGCCGCCG | - | cyclic nucleotide-binding domain-containing protein |
| AVAEIV_RS03680 | 740393 | 740405 | 134 | - | 11.2714 | 5.57E-05 | 0.0762 | CAGGTCGGCATCG | - | bifunctional ADP-dependent NAD(P)H-hydrate dehydratase/NAD(P)H-hydrate epimerase |
| AVAEIV_RS04380 | 891553 | 891565 | 350 | - | 11.2714 | 5.57E-05 | 0.078 | CCGGTGGCCATCC | *miaB* | methylthiotransferase MiaB |
| AVAEIV_RS05260 | 1084521 | 1084533 | 44 | - | 11.2714 | 5.57E-05 | 0.078 | CCCAACGCCGCCA | - | hydantoinase/oxoprolinase family protein |
| AVAEIV_RS06510 | 1342312 | 1342324 | -46 | + | 11.2714 | 5.57E-05 | 0.0762 | CTGTTCGGCATCA | - | PepSY domain-containing protein |

| | | | | | | | | | | |
|---|---|---|---|---|---|---|---|---|---|---|
| AVAEIV_RS10005 | 2095180 | 2095192 | 97 | + | 11.2714 | 5.57E-05 | 0.0762 | CTCACCGCCATCG | - | monovalent cation/H+ antiporter subunit D |
| AVAEIV_RS10715 | 2241869 | 2241881 | 132 | - | 11.2714 | 5.57E-05 | 0.078 | CAGTACGCGAGCA | - | tetratricopeptide repeat protein |
| AVAEIV_RS10825 | 2272924 | 2272936 | 13 | - | 11.2714 | 5.57E-05 | 0.078 | CTCACCGCCATCG | - | TIGR03986 family CRISPR-associated RAMP protein |
| AVAEIV_RS11355 | 2399938 | 2399950 | 33 | - | 11.2714 | 5.57E-05 | 0.0762 | TAGTTCGCCACCG | - | class I adenylate-forming enzyme family protein |
| AVAEIV_RS11680 | 2475797 | 2475809 | 78 | - | 11.2714 | 5.57E-05 | 0.0762 | CGGAACGCGATCA | - | CHAD domain-containing protein |
| AVAEIV_RS11855 | 2512934 | 2512946 | 102 | + | 11.2714 | 5.57E-05 | 0.078 | CTATTCGCCGTCA | *cobA* | uroporphyrinogen-III C-methyltransferase |
| AVAEIV_RS11875 | 2520768 | 2520780 | 226 | - | 11.2714 | 5.57E-05 | 0.078 | CTCGGCGCCATCA | - | kinase/phosphatase |
| AVAEIV_RS18765 | 4040171 | 4040183 | 68 | - | 11.2714 | 5.57E-05 | 0.0762 | CGGATGGCCGCCG | - | DUF4350 domain-containing protein |
| AVAEIV_RS19050 | 4104273 | 4104285 | 250 | + | 11.2714 | 5.57E-05 | 0.078 | CAGGTCGGCATCG | *frr* | ribosome recycling factor |
| AVAEIV_RS20470 | 4352871 | 4352883 | 174 | - | 11.2714 | 5.57E-05 | 0.078 | CTCACCGCCATCG | - | FecR domain-containing protein |
| AVAEIV_RS20975 | 4462279 | 4462291 | 290 | + | 11.2714 | 5.57E-05 | 0.078 | CAGGTCGGCATCG | - | efflux RND transporter periplasmic adaptor subunit |
| AVAEIV_RS22110 | 4718284 | 4718296 | 178 | + | 11.2714 | 5.57E-05 | 0.0762 | CGCTTCGCCACCC | *lptG* | LPS export ABC transporter permease LptG |
| AVAEIV_RS22380 | 4782642 | 4782654 | -42 | - | 11.2714 | 5.57E-05 | 0.0762 | CCCATCGCCGCCG | *aceF* | dihydrolipoyllysine-residue acetyltransferase |
| AVAEIV_RS22460 | 4801568 | 4801580 | 211 | - | 11.2714 | 5.57E-05 | 0.0762 | CGGATGGCCGCCG | *mscK* | mechanosensitive channel MscK |
| AVAEIV_RS22555 | 4828957 | 4828969 | 211 | + | 11.2714 | 5.57E-05 | 0.078 | CTCATCGCCATCT | - | pilus assembly protein PilM |
| AVAEIV_RS22560 | 4828957 | 4828969 | -43 | + | 11.2714 | 5.57E-05 | 0.0762 | CTCATCGCCATCT | - | penicillin-binding protein 1A |
| AVAEIV_RS23710 | 5053013 | 5053025 | 19 | - | 11.2714 | 5.57E-05 | 0.078 | CGACGCGCCAGCG | *vnfA3* | regulator VnfA3 |
| AVAEIV_RS25125 | 5349291 | 5349303 | 241 | + | 11.2714 | 5.57E-05 | 0.0762 | CAGGTCGGCATCG | *fhuB* | Fe(3+)-hydroxamate ABC transporter permease FhuB |
| AVAEIV_RS25425 | 5415995 | 5416007 | 102 | - | 11.2714 | 5.57E-05 | 0.078 | CCGATCGCCTTCG | - | RnfABCDGE type electron transport complex subunit B |
| AVAEIV_RS00075 | 17685 | 17697 | 315 | - | 11.2571 | 5.63E-05 | 0.0782 | CGCCAGGCCAGCG | *trkA* | Trk system potassium transporter TrkA |
| AVAEIV_RS00465 | 97015 | 97027 | 351 | + | 11.2571 | 5.63E-05 | 0.0782 | CGCCAGGCCAGCG | *cyoE* | heme o synthase |
| AVAEIV_RS01715 | 335799 | 335811 | 262 | + | 11.2571 | 5.63E-05 | 0.0782 | CAGATCGCCAGGC | - | DEAD/DEAH box helicase |
| AVAEIV_RS02255 | 452043 | 452055 | 191 | + | 11.2571 | 5.63E-05 | 0.0782 | CCGAGCGCCAGGA | - | NAD(P)/FAD-dependent oxidoreductase |
| AVAEIV_RS04705 | 958177 | 958189 | 198 | - | 11.2571 | 5.63E-05 | 0.0762 | CAGATCGCCAGGC | - | alpha/beta fold hydrolase |
| AVAEIV_RS07190 | 1482291 | 1482303 | 99 | - | 11.2571 | 5.63E-05 | 0.0782 | GACTGCGCCATCA | *cmoA* | carboxy-S-adenosyl-L-methionine synthase CmoA |
| AVAEIV_RS10320 | 2156806 | 2156818 | 77 | - | 11.2571 | 5.63E-05 | 0.0782 | CGGCAGGCCGTCA | - | SDR family oxidoreductase |
| AVAEIV_RS11480 | 2429598 | 2429610 | 166 | - | 11.2571 | 5.63E-05 | 0.0762 | CGCCAGGCCAGCG | - | ABC transporter ATP-binding protein |
| AVAEIV_RS12430 | 2634133 | 2634145 | 144 | - | 11.2571 | 5.63E-05 | 0.0762 | CAGATCGCCAGGC | - | response regulator |
| AVAEIV_RS18325 | 3945594 | 3945606 | 345 | + | 11.2571 | 5.63E-05 | 0.0782 | GAGATGGCCACCG | - | tyrosine-type recombinase/integrase |
| AVAEIV_RS18330 | 3945594 | 3945606 | -10 | + | 11.2571 | 5.63E-05 | 0.0782 | GAGATGGCCACCG | - | antitoxin |
| AVAEIV_RS18335 | 3945594 | 3945606 | 237 | + | 11.2571 | 5.63E-05 | 0.0762 | GAGATGGCCACCG | - | type II toxin-antitoxin system YafQ family toxin |
| AVAEIV_RS18840 | 4057387 | 4057399 | 152 | + | 11.2571 | 5.63E-05 | 0.0782 | CCGACGGCCACCG | - | phosphoketolase |
| AVAEIV_RS18845 | 4057387 | 4057399 | 40 | + | 11.2571 | 5.63E-05 | 0.0762 | CCGACGGCCACCG | - | containing protein |
| AVAEIV_RS19585 | 4177674 | 4177686 | 339 | + | 11.2571 | 5.63E-05 | 0.0762 | CAGCGTGCCGGCA | - | DUF1643 domain-containing protein |
| AVAEIV_RS20535 | 4366175 | 4366187 | 132 | + | 11.2571 | 5.63E-05 | 0.0782 | CGGATGGGCATCG | - | glutathione peroxidase |
| AVAEIV_RS20695 | 4399019 | 4399031 | 188 | + | 11.2571 | 5.63E-05 | 0.0762 | CCGACGGCCACCG | - | YbgA family protein |
| AVAEIV_RS23685 | 5046278 | 5046290 | 326 | - | 11.2571 | 5.63E-05 | 0.0762 | CAGCGTGCCGGCA | - | TPM domain-containing protein |
| AVAEIV_RS24675 | 5244507 | 5244519 | 176 | - | 11.2571 | 5.63E-05 | 0.0782 | CGGCTGGCCGTCG | - | cytochrome b/b6 domain-containing protein |
| AVAEIV_RS25335 | 5397376 | 5397388 | 239 | + | 11.2571 | 5.63E-05 | 0.0762 | CCGAGCGCCAGGA | *tssB* | subunit |
| AVAEIV_RS25860 | 5545863 | 5545875 | 76 | + | 11.2571 | 5.63E-05 | 0.0762 | CAGACGGCCGTCG | - | ABC transporter permease |
| AVAEIV_RS01330 | 258925 | 258937 | 137 | - | 11.2429 | 5.69E-05 | 0.0762 | CCGCCGGCCATCG | - | aspartate carbamoyltransferase |
| AVAEIV_RS03405 | 686833 | 686845 | 65 | + | 11.2429 | 5.69E-05 | 0.0782 | GAAAAGGCCATCA | - | hybrid sensor histidine kinase/response regulator |
| AVAEIV_RS04635 | 947262 | 947274 | 371 | + | 11.2429 | 5.69E-05 | 0.0762 | CCGCCGGCCATCG | - | SDR family oxidoreductase |
| AVAEIV_RS08380 | 1748255 | 1748267 | 114 | - | 11.2429 | 5.69E-05 | 0.0762 | GAGCTGGCCATCG | - | hypothetical protein |
| AVAEIV_RS10110 | 2111972 | 2111984 | 77 | + | 11.2429 | 5.69E-05 | 0.0782 | CACCTCGCCGCCG | *phnE* | phosphonate ABC transporter, permease protein PhnE |
| AVAEIV_RS13415 | 2900743 | 2900755 | 317 | - | 11.2429 | 5.69E-05 | 0.0782 | CAGCTCGCCTTCG | - | TIGR03571 family LLM class oxidoreductase |
| AVAEIV_RS14405 | 3136813 | 3136825 | 15 | + | 11.2429 | 5.69E-05 | 0.0782 | CCAGGGGCCATCA | *flhB* | flagellar biosynthesis protein FlhB |
| AVAEIV_RS16515 | 3582560 | 3582572 | 356 | - | 11.2429 | 5.69E-05 | 0.0762 | CTGAGCGCCGGCA | *tal* | transaldolase |
| AVAEIV_RS18320 | 3944040 | 3944052 | 142 | - | 11.2429 | 5.69E-05 | 0.0782 | CACCACGCCGCCA | - | DUF6387 family protein |
| AVAEIV_RS20470 | 4352916 | 4352928 | 219 | - | 11.2429 | 5.69E-05 | 0.0782 | CCGCTGGCCATCT | - | FecR domain-containing protein |
| AVAEIV_RS21265 | 4521473 | 4521485 | 115 | - | 11.2429 | 5.69E-05 | 0.0782 | CTGCATGCCATCG | *ilvC* | ketol-acid reductoisomerase |
| AVAEIV_RS22650 | 4847756 | 4847768 | 114 | - | 11.2429 | 5.69E-05 | 0.0762 | CACATCGGCACCG | - | 16S rRNA (uracil(1498)-N(3))-methyltransferase |

| | | | | | | | | | | |
|---|---|---|---|---|---|---|---|---|---|---|
| AVAEIV_RS23420 | 4992611 | 4992623 | 78 | - | 11.2429 | 5.69E-05 | 0.0782 | CCGCCGGCCATCG | - | LysR family transcriptional regulator |
| AVAEIV_RS23425 | 4992611 | 4992623 | 349 | - | 11.2429 | 5.69E-05 | 0.0762 | CCGCCGGCCATCG | - | alginate lyase family protein |
| AVAEIV_RS25275 | 5384743 | 5384755 | 314 | - | 11.2429 | 5.69E-05 | 0.0782 | GAGCTGGCCATCG | *modA* | molybdate ABC transporter substrate-binding protein |
| AVAEIV_RS00700 | 141562 | 141574 | 198 | + | 11.2286 | 5.75E-05 | 0.0762 | CTGCTCGCCAGCC | - | 4Fe4S-binding leucine-rich repeat protein |
| AVAEIV_RS01035 | 205294 | 205306 | 224 | - | 11.2286 | 5.75E-05 | 0.0782 | GAGGACGCCATCA | - | LLM class flavin-dependent oxidoreductase |
| AVAEIV_RS01045 | 205294 | 205306 | 112 | - | 11.2286 | 5.75E-05 | 0.0762 | GAGGACGCCATCA | - | hypothetical protein |
| AVAEIV_RS01290 | 252032 | 252044 | 201 | - | 11.2286 | 5.75E-05 | 0.0762 | CCGATCGCCAACG | - | hypothetical protein |
| AVAEIV_RS02340 | 466454 | 466466 | 299 | + | 11.2286 | 5.75E-05 | 0.0782 | CGGAGTGCCGGCA | - | S41 family peptidase |
| AVAEIV_RS05585 | 1154545 | 1154557 | -11 | - | 11.2286 | 5.75E-05 | 0.0762 | CGACAGGCCACCA | - | DUF6012 family protein |
| AVAEIV_RS05590 | 1154545 | 1154557 | 383 | - | 11.2286 | 5.75E-05 | 0.0762 | CGACAGGCCACCA | - | CopG family ribbon-helix-helix protein |
| AVAEIV_RS07455 | 1553734 | 1553746 | 293 | + | 11.2286 | 5.75E-05 | 0.0762 | CTGGTGGCCAGCG | - | integrating conjugative element protein |
| AVAEIV_RS08065 | 1672304 | 1672316 | 59 | + | 11.2286 | 5.75E-05 | 0.0762 | CCCTTCGCCGGCG | - | segregation and condensation protein A |
| AVAEIV_RS08340 | 1738773 | 1738785 | 55 | + | 11.2286 | 5.75E-05 | 0.0762 | CAGCCGGCCACCG | - | TolC family outer membrane protein |
| AVAEIV_RS08710 | 1817857 | 1817869 | 258 | + | 11.2286 | 5.75E-05 | 0.0782 | CGGTTGGCCGGCG | - | rRNA pseudouridine synthase |
| AVAEIV_RS09020 | 1877088 | 1877100 | 1 | + | 11.2286 | 5.75E-05 | 0.0782 | CAAAGCGCCAGGA | - | PhnD/SsuA/transferrin family substrate-binding protein |
| AVAEIV_RS09025 | 1877088 | 1877100 | 159 | + | 11.2286 | 5.75E-05 | 0.0762 | CAAAGCGCCAGGA | - | FCD domain-containing protein |
| AVAEIV_RS10115 | 2112906 | 2112918 | 184 | - | 11.2286 | 5.75E-05 | 0.0782 | CTGCTCGCCAGCC | - | PhnE/PtxC family ABC transporter permease |
| AVAEIV_RS10795 | 2264366 | 2264378 | 255 | - | 11.2286 | 5.75E-05 | 0.0782 | CTGCTCGCCAGCC | *csm6* | CRISPR-associated ring nuclease Csm6 |
| AVAEIV_RS10855 | 2279253 | 2279265 | -29 | + | 11.2286 | 5.75E-05 | 0.0762 | CAGTCCGCCATGA | - | rhodanese-like domain-containing protein |
| AVAEIV_RS11475 | 2428668 | 2428680 | 172 | - | 11.2286 | 5.75E-05 | 0.0762 | CTGGAGGCCAGCA | - | ABC transporter permease |
| AVAEIV_RS12105 | 2568074 | 2568086 | -43 | - | 11.2286 | 5.75E-05 | 0.0782 | CTGCTCGCCAGCC | - | c-type cytochrome |
| AVAEIV_RS12110 | 2568074 | 2568086 | 165 | - | 11.2286 | 5.75E-05 | 0.0762 | CTGCTCGCCAGCC | - | VRR-NUC domain-containing protein |
| AVAEIV_RS12340 | 2609259 | 2609271 | 127 | + | 11.2286 | 5.75E-05 | 0.0762 | CTGCTCGCCAGCC | *motB* | flagellar motor protein MotB |
| AVAEIV_RS14775 | 3212854 | 3212866 | 348 | + | 11.2286 | 5.75E-05 | 0.0762 | CCCTTCGCCGGCG | *nuoK* | NADH-quinone oxidoreductase subunit NuoK |
| AVAEIV_RS18025 | 3898629 | 3898641 | 383 | + | 11.2286 | 5.75E-05 | 0.0782 | CGACAGGCCACCA | - | CopG family ribbon-helix-helix protein |
| AVAEIV_RS18030 | 3898629 | 3898641 | -11 | + | 11.2286 | 5.75E-05 | 0.0782 | CGACAGGCCACCA | - | DUF6012 family protein |
| AVAEIV_RS18035 | 3898629 | 3898641 | 272 | + | 11.2286 | 5.75E-05 | 0.0762 | CGACAGGCCACCA | - | hypothetical protein |
| AVAEIV_RS19325 | 4151617 | 4151629 | 159 | - | 11.2286 | 5.75E-05 | 0.0782 | CCGATCGCCAACG | - | DUF6651 domain-containing protein |
| AVAEIV_RS19595 | 4178978 | 4178990 | 1 | + | 11.2286 | 5.75E-05 | 0.0762 | CTGGAGGCCAGCA | - | hypothetical protein |
| AVAEIV_RS21000 | 4466958 | 4466970 | 14 | + | 11.2286 | 5.75E-05 | 0.0782 | CGGTTGGCCGGCG | - | SulP family inorganic anion transporter |
| AVAEIV_RS21705 | 4624440 | 4624452 | -26 | + | 11.2286 | 5.75E-05 | 0.0782 | CACAGGGCCGCCA | - | esterase-like activity of phytase family protein |
| AVAEIV_RS22585 | 4837165 | 4837177 | 270 | + | 11.2286 | 5.75E-05 | 0.0762 | CCGGCCGCCATCG | *argS* | arginine--tRNA ligase |
| AVAEIV_RS24030 | 5114895 | 5114907 | 97 | - | 11.2286 | 5.75E-05 | 0.0762 | CAGCCGGCCACCG | - | DUF484 family protein |
| AVAEIV_RS24840 | 5282811 | 5282823 | -1 | + | 11.2286 | 5.75E-05 | 0.0762 | GAGGACGCCATCA | - | tripartite tricarboxylate transporter TctB family protein |
| AVAEIV_RS25075 | 5337754 | 5337766 | 96 | + | 11.2286 | 5.75E-05 | 0.0762 | CGATTCGCCATGA | - | DUF86 domain-containing protein |
| AVAEIV_RS25475 | 5424696 | 5424708 | 219 | + | 11.2286 | 5.75E-05 | 0.0762 | CTAATCGCCAGCC | - | site-specific integrase |
| AVAEIV_RS26065 | 5605570 | 5605582 | 204 | + | 11.2286 | 5.75E-05 | 0.0762 | CGCAGCGCCAGCC | - | DUF4433 domain-containing protein |
| AVAEIV_RS00110 | 24805 | 24817 | 382 | + | 11.2143 | 5.81E-05 | 0.0782 | CCGTCGGCCAGCG | - | NADPH:quinone reductase |
| AVAEIV_RS00180 | 37504 | 37516 | 191 | - | 11.2143 | 5.81E-05 | 0.0782 | CTTTTTGCCATCA | - | sensor domain-containing diguanylate cyclase |
| AVAEIV_RS01030 | 202088 | 202100 | 83 | + | 11.2143 | 5.81E-05 | 0.0762 | GCCATCGCCATCG | *oadA* | alpha |
| AVAEIV_RS03290 | 666701 | 666713 | 28 | + | 11.2143 | 5.81E-05 | 0.0782 | CCGTCGGCCAGCG | - | DUF3299 domain-containing protein |
| AVAEIV_RS04515 | 924250 | 924262 | 336 | + | 11.2143 | 5.81E-05 | 0.0782 | CCGTCGGCCAGCG | - | urease accessory protein UreF |
| AVAEIV_RS04855 | 991178 | 991190 | 158 | + | 11.2143 | 5.81E-05 | 0.0782 | CCGTTGGCCAGCT | - | RNA polymerase sigma factor |
| AVAEIV_RS04960 | 1015241 | 1015253 | 344 | - | 11.2143 | 5.81E-05 | 0.0782 | GCCATCGCCATCG | - | enoyl-CoA hydratase |
| AVAEIV_RS05455 | 1129940 | 1129952 | 55 | - | 11.2143 | 5.81E-05 | 0.0782 | CCCGTCGCCATCC | - | GspH/FimT family pseudopilin |
| AVAEIV_RS05460 | 1129940 | 1129952 | 244 | - | 11.2143 | 5.81E-05 | 0.0762 | CCCGTCGCCATCC | - | type IV pilin protein |
| AVAEIV_RS09065 | 1885404 | 1885416 | 355 | - | 11.2143 | 5.81E-05 | 0.0762 | CAGGCCGCCACCG | *zwf* | glucose-6-phosphate dehydrogenase |
| AVAEIV_RS09570 | 1995079 | 1995091 | 73 | + | 11.2143 | 5.81E-05 | 0.0762 | CAGGCCGCCACCG | - | ABC transporter permease |
| AVAEIV_RS10850 | 2278887 | 2278899 | 271 | + | 11.2143 | 5.81E-05 | 0.0782 | CGGAAAGCCAGCA | - | Cas10/Cmr2 second palm domain-containing protein |
| AVAEIV_RS10855 | 2278887 | 2278899 | 337 | + | 11.2143 | 5.81E-05 | 0.0762 | CGGAAAGCCAGCA | - | rhodanese-like domain-containing protein |
| AVAEIV_RS11070 | 2333749 | 2333761 | 31 | + | 11.2143 | 5.81E-05 | 0.0762 | CGCATCGCCGCCG | - | amino acid ABC transporter permease |
| AVAEIV_RS12190 | 2582900 | 2582912 | 195 | + | 11.2143 | 5.81E-05 | 0.0762 | CTGTCCGCCACCA | *flgE* | flagellar hook protein FlgE |

| | | | | | | | | | | |
|---|---|---|---|---|---|---|---|---|---|---|
| AVAEIV_RS12930 | 2803890 | 2803902 | 132 | + | 11.2143 | 5.81E-05 | 0.0782 | CCGATGGCCACGA | - | dienelactone hydrolase family protein |
| AVAEIV_RS15990 | 3485184 | 3485196 | 250 | + | 11.2143 | 5.81E-05 | 0.0762 | CGGGTGGCCATCC | - | ABC transporter ATP-binding protein |
| AVAEIV_RS16075 | 3502351 | 3502363 | 225 | + | 11.2143 | 5.81E-05 | 0.0762 | CGCATCGCCGCCG | - | OprD family porin |
| AVAEIV_RS16790 | 3633379 | 3633391 | 353 | - | 11.2143 | 5.81E-05 | 0.0762 | GCCATCGCCATCG | - | dienelactone hydrolase family protein |
| AVAEIV_RS18525 | 3987738 | 3987750 | 75 | + | 11.2143 | 5.81E-05 | 0.0762 | CAACCGGCCATCG | - | imelysin family protein |
| AVAEIV_RS19445 | 4164435 | 4164447 | 223 | + | 11.2143 | 5.81E-05 | 0.0782 | GCCAACGCCATCA | - | phage antirepressor KilAC domain-containing protein |
| AVAEIV_RS19660 | 4191191 | 4191203 | 326 | - | 11.2143 | 5.81E-05 | 0.0762 | CAGGCCGCCACCG | - | hypothetical protein |
| AVAEIV_RS20160 | 4306194 | 4306206 | 146 | + | 11.2143 | 5.81E-05 | 0.0782 | CGTATCGCCGGCA | *ispG* | diphosphate synthase |
| AVAEIV_RS20620 | 4386287 | 4386299 | 290 | + | 11.2143 | 5.81E-05 | 0.0762 | CGCATCGCCGCCG | - | TMEM165/GDT1 family protein |
| AVAEIV_RS22210 | 4740222 | 4740234 | 29 | - | 11.2143 | 5.81E-05 | 0.0782 | CGGATCGCCTTCG | - | ABC transporter substrate-binding protein |
| AVAEIV_RS22290 | 4759895 | 4759907 | 157 | + | 11.2143 | 5.81E-05 | 0.0782 | CCCGTCGCCATCC | *msbA* | lipid A export permease/ATP-binding protein MsbA |
| AVAEIV_RS23635 | 5038668 | 5038680 | 198 | + | 11.2143 | 5.81E-05 | 0.0782 | CGTATCGCCGGCA | *folB* | dihydroneopterin aldolase |
| AVAEIV_RS25465 | 5423261 | 5423273 | -28 | + | 11.2143 | 5.81E-05 | 0.0762 | CGCATCGCCGCCG | - | rhodanese-like domain-containing protein |
| AVAEIV_RS25470 | 5423261 | 5423273 | 344 | + | 11.2143 | 5.81E-05 | 0.0762 | CGCATCGCCGCCG | *grxD* | Grx4 family monothiol glutaredoxin |
| AVAEIV_RS25835 | 5537495 | 5537507 | -32 | - | 11.2143 | 5.81E-05 | 0.0782 | CGCAACGCCGCCA | - | DeoR/GlpR family DNA-binding transcription regulator |
| AVAEIV_RS00445 | 89962 | 89974 | 322 | + | 11.2 | 5.87E-05 | 0.0762 | CGCATCGGCATCG | - | GMC family oxidoreductase |
| AVAEIV_RS04205 | 858383 | 858395 | 48 | + | 11.2 | 5.87E-05 | 0.0762 | CGCATCGGCATCG | *dmpH* | 2-oxo-3-hexenedioate decarboxylase |
| AVAEIV_RS04490 | 917747 | 917759 | -40 | - | 11.2 | 5.87E-05 | 0.0784 | CACGTCGCCACCC | - | siderophore biosynthesis protein PvsA |
| AVAEIV_RS05040 | 1035109 | 1035121 | 252 | + | 11.2 | 5.87E-05 | 0.0784 | CACGTCGCCACCC | - | MBOAT family O-acyltransferase |
| AVAEIV_RS07110 | 1466885 | 1466897 | 126 | - | 11.2 | 5.87E-05 | 0.0784 | CGGTATGCCATCC | *nadB* | L-aspartate oxidase |
| AVAEIV_RS07115 | 1466885 | 1466897 | 215 | - | 11.2 | 5.87E-05 | 0.0762 | CGGTATGCCATCC | *rpoE* | RNA polymerase sigma factor RpoE |
| AVAEIV_RS07655 | 1594937 | 1594949 | 358 | + | 11.2 | 5.87E-05 | 0.0762 | CCGATCGACAGCG | - | low molecular weight protein-tyrosine-phosphatase |
| AVAEIV_RS10015 | 2097081 | 2097093 | 180 | - | 11.2 | 5.87E-05 | 0.0762 | TCGATGGCCATCG | - | K+/H+ antiporter subunit F |
| AVAEIV_RS10645 | 2226506 | 2226518 | 273 | - | 11.2 | 5.87E-05 | 0.0784 | CAAGCCGCCATCG | - | MMPL family transporter |
| AVAEIV_RS10985 | 2312241 | 2312253 | 253 | + | 11.2 | 5.87E-05 | 0.0784 | CCTACCGCCAGCA | - | MmgE/PrpD family protein |
| AVAEIV_RS11120 | 2346330 | 2346342 | -42 | - | 11.2 | 5.87E-05 | 0.0762 | CGCCTCGCCGTCG | - | protein |
| AVAEIV_RS13215 | 2865583 | 2865595 | 80 | + | 11.2 | 5.87E-05 | 0.0762 | GACATCGCCACCG | - | formimidoylglutamate deiminase |
| AVAEIV_RS14200 | 3092251 | 3092263 | 21 | + | 11.2 | 5.87E-05 | 0.0762 | CGCATCGGCATCG | - | DUF934 domain-containing protein |
| AVAEIV_RS14330 | 3122360 | 3122372 | 90 | + | 11.2 | 5.87E-05 | 0.0784 | TCGAAGGCCATCA | *rfbG* | CDP-glucose 4,6-dehydratase |
| AVAEIV_RS15425 | 3371113 | 3371125 | 298 | + | 11.2 | 5.87E-05 | 0.0762 | CGCATCGGCATCG | - | phosphoribosyltransferase family protein |
| AVAEIV_RS15530 | 3397123 | 3397135 | 192 | - | 11.2 | 5.87E-05 | 0.0762 | CGCCTCGCCGTCG | *codA* | cytosine deaminase |
| AVAEIV_RS16135 | 3517226 | 3517238 | 196 | - | 11.2 | 5.87E-05 | 0.0762 | CCGAATGCCGTCG | - | TIM barrel protein |
| AVAEIV_RS20725 | 4406357 | 4406369 | 219 | - | 11.2 | 5.87E-05 | 0.0784 | CCTACCGCCAGCA | *prfA* | peptide chain release factor 1 |
| AVAEIV_RS21530 | 4581684 | 4581696 | 307 | + | 11.2 | 5.87E-05 | 0.0784 | CACTTCGGCAGCG | - | RnfH family protein |
| AVAEIV_RS21540 | 4581684 | 4581696 | 338 | + | 11.2 | 5.87E-05 | 0.0762 | CACTTCGGCAGCG | *smpB* | SsrA-binding protein SmpB |
| AVAEIV_RS22200 | 4738281 | 4738293 | 13 | + | 11.2 | 5.87E-05 | 0.0784 | CCCTTGGCCAGCC | - | ABC transporter permease |
| AVAEIV_RS25000 | 5321074 | 5321086 | 269 | - | 11.2 | 5.87E-05 | 0.0762 | CACACCGCCGTCG | - | TIM barrel protein |
| AVAEIV_RS01020 | 200799 | 200811 | 288 | - | 11.1857 | 5.93E-05 | 0.0788 | TAGATGGCCACCG | - | LysR family transcriptional regulator |
| AVAEIV_RS05640 | 1160576 | 1160588 | 171 | + | 11.1857 | 5.93E-05 | 0.0788 | CTCCAGGCCAGCA | - | containing protein |
| AVAEIV_RS06135 | 1251792 | 1251804 | 286 | + | 11.1857 | 5.93E-05 | 0.0788 | CAGATGGCCGTTA | *pilV* | adhesin PilV |
| AVAEIV_RS12640 | 2681059 | 2681071 | -46 | - | 11.1857 | 5.93E-05 | 0.0788 | CTGCTGGCCGTCA | - | protein |
| AVAEIV_RS13930 | 3020195 | 3020207 | 57 | + | 11.1857 | 5.93E-05 | 0.0763 | CTGCTGGCCGTCA | - | sugar kinase |
| AVAEIV_RS14515 | 3161654 | 3161666 | 206 | - | 11.1857 | 5.93E-05 | 0.0763 | CACTGGGCCGGCA | *tusC* | sulfurtransferase complex subunit TusC |
| AVAEIV_RS15410 | 3366187 | 3366199 | 275 | + | 11.1857 | 5.93E-05 | 0.0763 | CTGCTGGCCGTCA | *tviB* | Vi polysaccharide biosynthesis UDP-N-acetylglucosamine C-6 dehydrogenase TviB |
| AVAEIV_RS16110 | 3511112 | 3511124 | 93 | + | 11.1857 | 5.93E-05 | 0.0763 | GACCTCGCCATCG | - | ABC transporter substrate-binding protein |
| AVAEIV_RS16530 | 3586860 | 3586872 | 316 | + | 11.1857 | 5.93E-05 | 0.0788 | CAATTGGCCAGCT | - | VacJ family lipoprotein |
| AVAEIV_RS16535 | 3586860 | 3586872 | 107 | + | 11.1857 | 5.93E-05 | 0.0763 | CAATTGGCCAGCT | - | DUF4404 family protein |
| AVAEIV_RS17980 | 3887477 | 3887489 | 171 | - | 11.1857 | 5.93E-05 | 0.0763 | CTCCAGGCCAGCA | - | containing protein |
| AVAEIV_RS18860 | 4062041 | 4062053 | 241 | - | 11.1857 | 5.93E-05 | 0.0788 | CAGATGGCGAGCG | *recA* | recombinase RecA |
| AVAEIV_RS19405 | 4160012 | 4160024 | 380 | + | 11.1857 | 5.93E-05 | 0.0788 | CCGATGGCCACTA | - | KilA-N domain-containing protein |
| AVAEIV_RS21320 | 4532455 | 4532467 | 175 | + | 11.1857 | 5.93E-05 | 0.0763 | CAGAAGGCGAGCA | - | Rieske (2Fe-2S) protein |

| | | | | | | | | | | |
|---|---|---|---|---|---|---|---|---|---|---|
| AVAEIV_RS21605 | 4599779 | 4599791 | 187 | - | 11.1857 | 5.93E-05 | 0.0788 | CGGCCGGCCATCG | - | sugar diacid recognition domain-containing protein |
| AVAEIV_RS21610 | 4599779 | 4599791 | 129 | - | 11.1857 | 5.93E-05 | 0.0763 | CGGCCGGCCATCG | *pyk* | pyruvate kinase |
| AVAEIV_RS25050 | 5332151 | 5332163 | 275 | + | 11.1857 | 5.93E-05 | 0.0763 | CGAACGGCCATCG | *fahA* | fumarylacetoacetase |
| AVAEIV_RS25120 | 5348548 | 5348560 | 109 | + | 11.1857 | 5.93E-05 | 0.0763 | CTCCTGGCCAGCG | - | ABC transporter substrate-binding protein |
| AVAEIV_RS00710 | 144049 | 144061 | 143 | - | 11.1714 | 5.99E-05 | 0.0763 | CAGCATGCCGTCG | *nifN* | protein NifN |
| AVAEIV_RS03375 | 679094 | 679106 | 99 | - | 11.1714 | 5.99E-05 | 0.0763 | CTCGACGCCAGCA | *ribA* | GTP cyclohydrolase II |
| AVAEIV_RS05330 | 1098848 | 1098860 | 71 | - | 11.1714 | 5.99E-05 | 0.0763 | CGTTTCGCCACCG | - | cytochrome b |
| AVAEIV_RS09360 | 1950784 | 1950796 | 136 | + | 11.1714 | 5.99E-05 | 0.0763 | CTCGTCGCCAGCG | - | hypothetical protein |
| AVAEIV_RS11810 | 2499997 | 2500009 | 325 | - | 11.1714 | 5.99E-05 | 0.0763 | CGGATCGCCAACG | *acnD* | Fe/S-dependent 2-methylisocitrate dehydratase AcnD |
| AVAEIV_RS13815 | 2993905 | 2993917 | 205 | - | 11.1714 | 5.99E-05 | 0.079 | CGCCTGGCCATCC | *tssF* | type VI secretion system baseplate subunit TssF |
| AVAEIV_RS21080 | 4483484 | 4483496 | 190 | - | 11.1714 | 5.99E-05 | 0.079 | CGGGCCGCCATCG | - | 2-keto-4-pentenoate hydratase |
| AVAEIV_RS23670 | 5045050 | 5045062 | 288 | - | 11.1714 | 5.99E-05 | 0.0763 | CAGCATGCCGTCG | *cowN* | N(2)-fixation sustaining protein CowN |
| AVAEIV_RS23755 | 5061721 | 5061733 | 305 | - | 11.1714 | 5.99E-05 | 0.0763 | CAGCATGCCGTCG | *fecA* | TonB-dependent Fe(3+) dicitrate receptor FecA |
| AVAEIV_RS01220 | 238625 | 238637 | 120 | - | 11.1571 | 6.04E-05 | 0.0763 | CGGTCGGCCAGCG | *mgtA* | magnesium-translocating P-type ATPase |
| AVAEIV_RS01510 | 290113 | 290125 | 351 | + | 11.1571 | 6.04E-05 | 0.0763 | GACTTCGCCAGCG | - | NINE protein |
| AVAEIV_RS02700 | 552946 | 552958 | 329 | + | 11.1571 | 6.04E-05 | 0.079 | CCGAGCGGCAGCA | - | polysaccharide biosynthesis/export family protein |
| AVAEIV_RS03780 | 759542 | 759554 | 291 | - | 11.1571 | 6.04E-05 | 0.0763 | CAGCTTGCCTTCA | *dnaB* | replicative DNA helicase |
| AVAEIV_RS03880 | 782236 | 782248 | 152 | + | 11.1571 | 6.04E-05 | 0.079 | CAGCTCGCCGGCC | *ispA* | (2E,6E)-farnesyl diphosphate synthase |
| AVAEIV_RS08165 | 1697166 | 1697178 | 193 | + | 11.1571 | 6.04E-05 | 0.0763 | GGCATCGCCATCG | *cmk* | (d)CMP kinase |
| AVAEIV_RS08470 | 1771287 | 1771299 | 286 | + | 11.1571 | 6.04E-05 | 0.0763 | GGCATCGCCATCG | - | hypothetical protein |
| AVAEIV_RS08590 | 1792951 | 1792963 | 250 | + | 11.1571 | 6.04E-05 | 0.0763 | CGGTCGGCCAGCG | - | IS5/IS1182 family transposase |
| AVAEIV_RS09310 | 1943344 | 1943356 | 268 | + | 11.1571 | 6.04E-05 | 0.079 | CAGCTCGCCGGCC | - | regulator |
| AVAEIV_RS09380 | 1957015 | 1957027 | 293 | - | 11.1571 | 6.04E-05 | 0.079 | CAGCTCGCCGGCC | - | IS4-like element ISAzvi7 family transposase |
| AVAEIV_RS09510 | 1980296 | 1980308 | 282 | + | 11.1571 | 6.04E-05 | 0.0763 | GGCATCGCCATCG | - | efflux RND transporter periplasmic adaptor subunit |
| AVAEIV_RS09615 | 2008966 | 2008978 | 327 | - | 11.1571 | 6.04E-05 | 0.079 | CCGAGCGGCAGCA | - | BatD family protein |
| AVAEIV_RS10585 | 2212097 | 2212109 | 152 | + | 11.1571 | 6.04E-05 | 0.079 | CAGCTCGCCGGCC | *ispA* | (2E,6E)-farnesyl diphosphate synthase |
| AVAEIV_RS10925 | 2295208 | 2295220 | 169 | + | 11.1571 | 6.04E-05 | 0.079 | GGCATCGCCATCG | *puuE* | allantoinase PuuE |
| AVAEIV_RS11195 | 2363828 | 2363840 | 218 | + | 11.1571 | 6.04E-05 | 0.079 | CCCTCCGCCAGCG | - | LLM class flavin-dependent oxidoreductase |
| AVAEIV_RS11580 | 2448211 | 2448223 | 245 | - | 11.1571 | 6.04E-05 | 0.0763 | CAGCTCGCCGGCC | - | Dps family protein |
| AVAEIV_RS11610 | 2456492 | 2456504 | 91 | + | 11.1571 | 6.04E-05 | 0.079 | CAAAGCGCTAGCA | - | sensor domain-containing diguanylate cyclase |
| AVAEIV_RS12240 | 2593720 | 2593732 | 76 | - | 11.1571 | 6.04E-05 | 0.079 | GACTTCGCCAGCG | *fliP* | flagellar type III secretion system pore protein FliP |
| AVAEIV_RS14035 | 3049535 | 3049547 | 24 | + | 11.1571 | 6.04E-05 | 0.079 | CCAAGCGCCGGCA | - | phytanoyl-CoA dioxygenase family protein |
| AVAEIV_RS14040 | 3049535 | 3049547 | 381 | + | 11.1571 | 6.04E-05 | 0.0763 | CCAAGCGCCGGCA | - | FagA protein |
| AVAEIV_RS14060 | 3057625 | 3057637 | 339 | - | 11.1571 | 6.04E-05 | 0.079 | CAGATCGGCAGCC | - | dehydrogenase |
| AVAEIV_RS16930 | 3666919 | 3666931 | 375 | - | 11.1571 | 6.04E-05 | 0.079 | CCACATGCCATCG | - | maltoporin |
| AVAEIV_RS16990 | 3683310 | 3683322 | 175 | - | 11.1571 | 6.04E-05 | 0.079 | CAGATCGGCAGCC | - | LysR family transcriptional regulator |
| AVAEIV_RS22320 | 4766287 | 4766299 | 132 | - | 11.1571 | 6.04E-05 | 0.079 | CAGCTCGCCGGCC | - | hypothetical protein |
| AVAEIV_RS22785 | 4877588 | 4877600 | 271 | + | 11.1571 | 6.04E-05 | 0.0763 | CAGATCGGCAGCC | *tpiA* | triose-phosphate isomerase |
| AVAEIV_RS22955 | 4916954 | 4916966 | 62 | + | 11.1571 | 6.04E-05 | 0.0763 | GGCATCGCCATCG | *coq7* | monooxygenase |
| AVAEIV_RS23170 | 4948349 | 4948361 | 148 | + | 11.1571 | 6.04E-05 | 0.0763 | CGCCGGGCCACCA | - | hypothetical protein |
| AVAEIV_RS23945 | 5100826 | 5100838 | 304 | + | 11.1571 | 6.04E-05 | 0.0763 | GGCATCGCCATCG | - | TIGR02444 family protein |
| AVAEIV_RS24710 | 5250652 | 5250664 | 356 | + | 11.1571 | 6.04E-05 | 0.079 | CCGCGCGCCGGCA | - | hypothetical protein |
| AVAEIV_RS24950 | 5310037 | 5310049 | 271 | - | 11.1571 | 6.04E-05 | 0.079 | CAGATCGGCAGCC | *tpiA* | triose-phosphate isomerase |
| AVAEIV_RS25910 | 5561325 | 5561337 | 332 | - | 11.1571 | 6.04E-05 | 0.079 | CAGCTCGCCGGCC | - | hypothetical protein |
| AVAEIV_RS01420 | 276830 | 276842 | 30 | - | 11.1429 | 6.12E-05 | 0.0793 | CTGATCGCCTTCA | *dut* | dUTP diphosphatase |
| AVAEIV_RS05380 | 1114693 | 1114705 | 210 | - | 11.1429 | 6.12E-05 | 0.0793 | CCCAAGGCCGGCA | - | DNA polymerase III subunit chi |
| AVAEIV_RS06000 | 1224323 | 1224335 | 354 | - | 11.1429 | 6.12E-05 | 0.0793 | CGATTCGCCGCCA | *traD* | TraD |
| AVAEIV_RS12405 | 2627796 | 2627808 | 382 | - | 11.1429 | 6.12E-05 | 0.0767 | TAGTTGGCCAGCG | - | DEAD/DEAH box helicase family protein |
| AVAEIV_RS12860 | 2772524 | 2772536 | 247 | - | 11.1429 | 6.12E-05 | 0.0767 | CCCAAGGCCGGCA | - | containing protein |
| AVAEIV_RS14765 | 3211811 | 3211823 | 324 | + | 11.1429 | 6.12E-05 | 0.0767 | CCGATGGCTACCA | *nuoI* | NADH-quinone oxidoreductase subunit NuoI |
| AVAEIV_RS15130 | 3294818 | 3294830 | 279 | - | 11.1429 | 6.12E-05 | 0.0767 | AACATCGCCAGCA | - | diguanylate cyclase |
| AVAEIV_RS15965 | 3480559 | 3480571 | 258 | + | 11.1429 | 6.12E-05 | 0.0793 | TCCATCGCCATCG | - | helix-turn-helix domain-containing protein |

| | | | | | | | | | | |
|---|---|---|---|---|---|---|---|---|---|---|
| AVAEIV_RS16270 | 3546410 | 3546422 | 275 | + | 11.1429 | 6.12E-05 | 0.0793 | CGCACCGCCACCG | - | hypothetical protein |
| AVAEIV_RS16925 | 3663495 | 3663507 | 99 | - | 11.1429 | 6.12E-05 | 0.0767 | CCCATGGCCGGCG | - | RES family NAD+ phosphorylase |
| AVAEIV_RS17895 | 3872676 | 3872688 | 354 | + | 11.1429 | 6.12E-05 | 0.0767 | CGATTCGCCGCCA | *traD* | TraD |
| AVAEIV_RS19805 | 4221837 | 4221849 | 210 | + | 11.1429 | 6.12E-05 | 0.0793 | CCGTTGGCCAGTA | - | homoserine dehydrogenase |
| AVAEIV_RS22595 | 4840135 | 4840147 | -29 | + | 11.1429 | 6.12E-05 | 0.0767 | CAGTCCGCCGCCA | *hslV* | ATP-dependent protease subunit HslV |
| AVAEIV_RS23900 | 5093471 | 5093483 | 92 | - | 11.1429 | 6.12E-05 | 0.0793 | CTGATCGCCTTCA | *ppk1* | polyphosphate kinase 1 |
| AVAEIV_RS24025 | 5113970 | 5113982 | 178 | - | 11.1429 | 6.12E-05 | 0.0767 | CAGATCGCCCTCG | *dapF* | diaminopimelate epimerase |
| AVAEIV_RS01810 | 357965 | 357977 | 38 | - | 11.1286 | 6.18E-05 | 0.0793 | CGGAGCGCTAGCA | - | ABC transporter substrate-binding protein |
| AVAEIV_RS02595 | 527474 | 527486 | 340 | - | 11.1286 | 6.18E-05 | 0.0793 | CTGCACGCCACCG | *rimK* | 30S ribosomal protein S6--L-glutamate ligase |
| AVAEIV_RS02605 | 527474 | 527486 | 206 | - | 11.1286 | 6.18E-05 | 0.0769 | CTGCACGCCACCG | - | RNA-binding S4 domain-containing protein |
| AVAEIV_RS02845 | 584575 | 584587 | 151 | - | 11.1286 | 6.18E-05 | 0.0769 | CGGATCGCCGGCC | - | slipin family protein |
| AVAEIV_RS03945 | 795535 | 795547 | 342 | - | 11.1286 | 6.18E-05 | 0.0769 | CAGCGCGGCAGCA | - | YceK/YidQ family lipoprotein |
| AVAEIV_RS04665 | 953880 | 953892 | 310 | - | 11.1286 | 6.18E-05 | 0.0793 | CGGATCGCCGGCC | *cueR* | Cu(I)-responsive transcriptional regulator |
| AVAEIV_RS06175 | 1260487 | 1260499 | 43 | - | 11.1286 | 6.18E-05 | 0.0793 | GCCATTGCCATCA | - | TcpQ domain-containing protein |
| AVAEIV_RS07345 | 1521077 | 1521089 | 88 | - | 11.1286 | 6.18E-05 | 0.0769 | CAGGATGCCATCC | *glp* | gephyrin-like molybdotransferase Glp |
| AVAEIV_RS12495 | 2652254 | 2652266 | 135 | + | 11.1286 | 6.18E-05 | 0.0793 | CACATCGCGAGCG | *glgX* | glycogen debranching protein GlgX |
| AVAEIV_RS14405 | 3136757 | 3136769 | -41 | + | 11.1286 | 6.18E-05 | 0.0793 | CAGTGCGCCGCCG | *flhB* | flagellar biosynthesis protein FlhB |
| AVAEIV_RS17275 | 3743669 | 3743681 | 256 | - | 11.1286 | 6.18E-05 | 0.0793 | TATATCGCCAGCA | - | glycosyltransferase |
| AVAEIV_RS22160 | 4729916 | 4729928 | 193 | + | 11.1286 | 6.18E-05 | 0.0793 | TACATCGCCACCG | - | phosphotransferase |
| AVAEIV_RS23525 | 5011931 | 5011943 | -46 | + | 11.1286 | 6.18E-05 | 0.0769 | CTGCACGCCACCG | - | MFS transporter |
| AVAEIV_RS24100 | 5126523 | 5126535 | 76 | - | 11.1286 | 6.18E-05 | 0.0769 | CGGATCGCCGGCC | *rep* | DNA helicase Rep |
| AVAEIV_RS24535 | 5214447 | 5214459 | 68 | - | 11.1286 | 6.18E-05 | 0.0793 | CCAATGGCTATCA | - | SAM-dependent methyltransferase |
| AVAEIV_RS25750 | 5513556 | 5513568 | -41 | + | 11.1286 | 6.18E-05 | 0.0793 | CGTAGCGCCAGCG | *recB* | exodeoxyribonuclease V subunit beta |
| AVAEIV_RS00570 | 117786 | 117798 | 280 | - | 11.1143 | 6.24E-05 | 0.0769 | CACCTGGCCGGCG | - | hypothetical protein |
| AVAEIV_RS01155 | 225612 | 225624 | 44 | - | 11.1143 | 6.24E-05 | 0.0793 | CCAATCGCGATCA | - | carboxymuconolactone decarboxylase family protein |
| AVAEIV_RS03655 | 737238 | 737250 | 374 | - | 11.1143 | 6.24E-05 | 0.0793 | CCGACCGCCAGCC | *asd* | archaetidylserine decarboxylase |
| AVAEIV_RS03830 | 772470 | 772482 | 315 | - | 11.1143 | 6.24E-05 | 0.0793 | CCGCTGGCCGCCA | *trmA* | tRNA (uridine(54)-C5)-methyltransferase TrmA |
| AVAEIV_RS04730 | 963347 | 963359 | 215 | - | 11.1143 | 6.24E-05 | 0.0769 | CAGGGTGCCACCG | *mnmH* | tRNA 2-selenouridine(34) synthase MnmH |
| AVAEIV_RS06825 | 1406163 | 1406175 | 341 | + | 11.1143 | 6.24E-05 | 0.0769 | GAGATCGCCAGCC | *rpsI* | 30S ribosomal protein S9 |
| AVAEIV_RS07340 | 1520237 | 1520249 | 384 | + | 11.1143 | 6.24E-05 | 0.0769 | CAGTGCGGCATCG | *moaB* | molybdenum cofactor biosynthesis protein B |
| AVAEIV_RS07405 | 1537550 | 1537562 | 149 | + | 11.1143 | 6.24E-05 | 0.0769 | CCGCTGGCCGCCA | - | acyl-CoA dehydrogenase family protein |
| AVAEIV_RS07895 | 1642124 | 1642136 | 332 | - | 11.1143 | 6.24E-05 | 0.0793 | CTGCCGGCCATCA | - | LysR family transcriptional regulator |
| AVAEIV_RS10155 | 2119496 | 2119508 | 319 | - | 11.1143 | 6.24E-05 | 0.0769 | CACCAGGCCGGCA | *recR* | recombination mediator RecR |
| AVAEIV_RS11355 | 2399848 | 2399860 | 123 | - | 11.1143 | 6.24E-05 | 0.0769 | CGGAACGCCATGG | - | class I adenylate-forming enzyme family protein |
| AVAEIV_RS11520 | 2439566 | 2439578 | 79 | - | 11.1143 | 6.24E-05 | 0.0793 | GAGATCGCCAGCC | - | FecR family protein |
| AVAEIV_RS12045 | 2558255 | 2558267 | 157 | - | 11.1143 | 6.24E-05 | 0.0793 | GAGATCGCCAGCC | - | hypothetical protein |
| AVAEIV_RS12260 | 2596413 | 2596425 | 331 | - | 11.1143 | 6.24E-05 | 0.0793 | GCGAGCGCCAGCA | *fliL* | flagellar basal body-associated protein FliL |
| AVAEIV_RS12595 | 2675245 | 2675257 | 341 | + | 11.1143 | 6.24E-05 | 0.0769 | CTGCCGGCCATCA | - | tRNA-Gly |
| AVAEIV_RS15690 | 3430793 | 3430805 | 323 | + | 11.1143 | 6.24E-05 | 0.0793 | CACCAGGCCGGCA | *ssuB* | protein |
| AVAEIV_RS15895 | 3468209 | 3468221 | 180 | - | 11.1143 | 6.24E-05 | 0.0793 | CATCTGGCCACCG | - | ABC transporter substrate-binding protein |
| AVAEIV_RS17670 | 3826628 | 3826640 | 229 | + | 11.1143 | 6.24E-05 | 0.0793 | GAGATCGCCAGCC | - | tRNA-Asp |
| AVAEIV_RS17675 | 3826628 | 3826640 | 139 | + | 11.1143 | 6.24E-05 | 0.0793 | GAGATCGCCAGCC | - | tRNA-Val |
| AVAEIV_RS17680 | 3826628 | 3826640 | 2 | + | 11.1143 | 6.24E-05 | 0.0769 | GAGATCGCCAGCC | *rdgC* | recombination-associated protein RdgC |
| AVAEIV_RS18200 | 3927562 | 3927574 | 340 | + | 11.1143 | 6.24E-05 | 0.0793 | CCGCTGGCCGCCA | - | CDP-alcohol phosphatidyltransferase family protein |
| AVAEIV_RS19750 | 4209088 | 4209100 | 363 | + | 11.1143 | 6.24E-05 | 0.0793 | CCGACCGCCAGCC | - | hypothetical protein |
| AVAEIV_RS19930 | 4248869 | 4248881 | -11 | + | 11.1143 | 6.24E-05 | 0.0769 | CCATGCGCCACCG | - | DMT family transporter |
| AVAEIV_RS23080 | 4936835 | 4936847 | 162 | - | 11.1143 | 6.24E-05 | 0.0769 | CACCTGGCCGGCG | - | DUF2597 family protein |
| AVAEIV_RS24165 | 5137342 | 5137354 | 262 | + | 11.1143 | 6.24E-05 | 0.0769 | CACCTGGCCGGCG | - | Lrp/AsnC ligand binding domain-containing protein |
| AVAEIV_RS24825 | 5281043 | 5281055 | 305 | - | 11.1143 | 6.24E-05 | 0.0793 | GCGAGCGCCAGCA | - | sensor histidine kinase |
| AVAEIV_RS26105 | 5612751 | 5612763 | 208 | + | 11.1143 | 6.24E-05 | 0.0793 | CACCTGGCCGGCG | - | tyrosine-type recombinase/integrase |
| AVAEIV_RS00215 | 44011 | 44023 | 48 | - | 11.1 | 6.29E-05 | 0.0793 | CCGGTCGCCGCCA | - | hypothetical protein |
| AVAEIV_RS00225 | 45117 | 45129 | 337 | + | 11.1 | 6.29E-05 | 0.0769 | GGGTTCGCCACCA | - | ABC transporter substrate-binding protein |

| | | | | | | | | | | |
|---|---|---|---|---|---|---|---|---|---|---|
| AVAEIV_RS03450 | 696412 | 696424 | 340 | - | 11.1 | 6.29E-05 | 0.0793 | CGCTTCGACATCA | *aroQ* | type II 3-dehydroquinate dehydratase |
| AVAEIV_RS05585 | 1154169 | 1154181 | 365 | - | 11.1 | 6.29E-05 | 0.0769 | CACAGCGACAGCA | - | DUF6012 family protein |
| AVAEIV_RS06315 | 1290366 | 1290378 | 337 | - | 11.1 | 6.29E-05 | 0.0793 | CATATTGCCAGCC | - | NAD(P)/FAD-dependent oxidoreductase |
| AVAEIV_RS06320 | 1290366 | 1290378 | 99 | - | 11.1 | 6.29E-05 | 0.0793 | CATATTGCCAGCC | - | DUF3094 domain-containing protein |
| AVAEIV_RS06325 | 1290366 | 1290378 | 192 | - | 11.1 | 6.29E-05 | 0.0769 | CATATTGCCAGCC | - | GNAT family N-acetyltransferase |
| AVAEIV_RS07755 | 1615660 | 1615672 | 231 | - | 11.1 | 6.29E-05 | 0.0793 | CGGCGCGCCGGCA | - | IS5 family transposase |
| AVAEIV_RS12800 | 2720789 | 2720801 | 136 | + | 11.1 | 6.29E-05 | 0.0793 | CACAGCGACAGCA | - | integrase core domain-containing protein |
| AVAEIV_RS13720 | 2966626 | 2966638 | 377 | - | 11.1 | 6.29E-05 | 0.0769 | CGCATCGCCACGA | - | FecCD family ABC transporter permease |
| AVAEIV_RS13835 | 2997340 | 2997352 | 372 | - | 11.1 | 6.29E-05 | 0.0793 | CACGTCGCCGGCG | *tssB* | subunit |
| AVAEIV_RS14620 | 3182182 | 3182194 | 223 | - | 11.1 | 6.29E-05 | 0.0769 | CGGAGCGGCAGCA | - | NUDIX hydrolase |
| AVAEIV_RS15190 | 3307684 | 3307696 | 155 | + | 11.1 | 6.29E-05 | 0.0793 | CGCTCCGCCAGCG | - | ACP S-malonyltransferase |
| AVAEIV_RS15220 | 3315708 | 3315720 | 268 | - | 11.1 | 6.29E-05 | 0.0793 | CGGCGCGCCGGCA | - | protein |
| AVAEIV_RS18095 | 3910336 | 3910348 | 321 | - | 11.1 | 6.29E-05 | 0.0793 | CGGAGCGGCAGCA | *tolB* | Tol-Pal system beta propeller repeat protein TolB |
| AVAEIV_RS18525 | 3987497 | 3987509 | 316 | + | 11.1 | 6.29E-05 | 0.0769 | CACGACGCCGGCA | - | imelysin family protein |
| AVAEIV_RS18770 | 4041102 | 4041114 | 279 | + | 11.1 | 6.29E-05 | 0.0769 | CGGCGCGCCGGCA | - | AAA family ATPase |
| AVAEIV_RS20220 | 4315242 | 4315254 | 182 | + | 11.1 | 6.29E-05 | 0.0793 | CGAAGCGCCGGCA | *iscR* | Fe-S cluster assembly transcriptional regulator IscR |
| AVAEIV_RS22190 | 4736915 | 4736927 | 372 | + | 11.1 | 6.29E-05 | 0.0793 | CGCTCCGCCAGCG | - | acyl-CoA dehydrogenase family protein |
| AVAEIV_RS23080 | 4936813 | 4936825 | 184 | - | 11.1 | 6.29E-05 | 0.0769 | CTGCGGGCCATCG | - | DUF2597 family protein |
| AVAEIV_RS24175 | 5140720 | 5140732 | 162 | + | 11.1 | 6.29E-05 | 0.0793 | CATGTCGCCACCG | - | MFS transporter |
| AVAEIV_RS02205 | 441526 | 441538 | 128 | - | 11.0857 | 6.35E-05 | 0.0769 | CAGGTGGCCAGCT | - | cyclic nucleotide-binding domain-containing protein |
| AVAEIV_RS02465 | 492700 | 492712 | 168 | - | 11.0857 | 6.35E-05 | 0.0794 | CTGGGCGCCATCG | - | TolC family protein |
| AVAEIV_RS02850 | 585402 | 585414 | 245 | + | 11.0857 | 6.35E-05 | 0.0769 | CTGGGCGCCATCG | *ald* | alanine dehydrogenase |
| AVAEIV_RS03540 | 714910 | 714922 | 31 | + | 11.0857 | 6.35E-05 | 0.0769 | CGCAAGGCCGGCA | - | hypothetical protein |
| AVAEIV_RS05840 | 1195954 | 1195966 | 179 | + | 11.0857 | 6.35E-05 | 0.0769 | CAGGCGGCCAGCG | - | protein |
| AVAEIV_RS10590 | 2212433 | 2212445 | 249 | + | 11.0857 | 6.35E-05 | 0.0794 | CAGGCGGCCAGCG | - | exodeoxyribonuclease VII small subunit |
| AVAEIV_RS16185 | 3530346 | 3530358 | 150 | - | 11.0857 | 6.35E-05 | 0.0794 | CAGCCCGCCAGCC | - | IS4-like element ISAzvi7 family transposase |
| AVAEIV_RS16570 | 3593914 | 3593926 | 353 | + | 11.0857 | 6.35E-05 | 0.0769 | CCGGTCGGCATCA | - | YecA family protein |
| AVAEIV_RS16610 | 3601088 | 3601100 | -4 | - | 11.0857 | 6.35E-05 | 0.0794 | CACACCGCCATGA | - | hypothetical protein |
| AVAEIV_RS16615 | 3601088 | 3601100 | 95 | - | 11.0857 | 6.35E-05 | 0.0769 | CACACCGCCATGA | - | S9 family peptidase |
| AVAEIV_RS18040 | 3899289 | 3899301 | 285 | + | 11.0857 | 6.35E-05 | 0.0769 | GAGTCCGCCATCA | - | Fic family protein |
| AVAEIV_RS18995 | 4093008 | 4093020 | 342 | + | 11.0857 | 6.35E-05 | 0.0794 | CAGGCGGCCAGCG | *rnhB* | ribonuclease HII |
| AVAEIV_RS19185 | 4134296 | 4134308 | 379 | + | 11.0857 | 6.35E-05 | 0.0794 | CAGCCCGCCAGCC | - | pyocin knob domain-containing protein |
| AVAEIV_RS19190 | 4134296 | 4134308 | 29 | + | 11.0857 | 6.35E-05 | 0.0794 | CAGCCCGCCAGCC | - | phage tail protein |
| AVAEIV_RS19565 | 4176130 | 4176142 | 354 | - | 11.0857 | 6.35E-05 | 0.0769 | CAGGCGGCCAGCG | - | DUF551 domain-containing protein |
| AVAEIV_RS19575 | 4177402 | 4177414 | -47 | - | 11.0857 | 6.35E-05 | 0.0769 | CGCTTTGCCGGCA | - | hypothetical protein |
| AVAEIV_RS19580 | 4177402 | 4177414 | 156 | - | 11.0857 | 6.35E-05 | 0.0769 | CGCTTTGCCGGCA | - | hypothetical protein |
| AVAEIV_RS21025 | 4469934 | 4469946 | 53 | + | 11.0857 | 6.35E-05 | 0.0769 | CGCTTTGCCGGCA | - | HigA family addiction module antitoxin |
| AVAEIV_RS22070 | 4711317 | 4711329 | 172 | - | 11.0857 | 6.35E-05 | 0.0794 | CGGAACGCCATTG | *spt* | serine palmitoyltransferase |
| AVAEIV_RS23170 | 4948281 | 4948293 | 216 | - | 11.0857 | 6.35E-05 | 0.0769 | CAGGCGGCCAGCG | - | hypothetical protein |
| AVAEIV_RS24375 | 5182020 | 5182032 | -28 | + | 11.0857 | 6.35E-05 | 0.0769 | CCGAACGCCGCCG | - | containing protein |
| AVAEIV_RS25500 | 5431958 | 5431970 | 26 | - | 11.0857 | 6.35E-05 | 0.0794 | CGGATGGCTACCA | - | MFS transporter |
| AVAEIV_RS01500 | 288839 | 288851 | 69 | + | 11.0714 | 6.43E-05 | 0.0794 | CCCATGGACATCA | - | YggS family pyridoxal phosphate-dependent enzyme |
| AVAEIV_RS01505 | 288839 | 288851 | -10 | + | 11.0714 | 6.43E-05 | 0.0773 | CCCATGGACATCA | - | type IV pilus twitching motility protein PilT |
| AVAEIV_RS02240 | 449875 | 449887 | 326 | - | 11.0714 | 6.43E-05 | 0.0794 | CAAGTCGCCGCCA | - | HupE/UreJ family protein |
| AVAEIV_RS02245 | 449875 | 449887 | 73 | - | 11.0714 | 6.43E-05 | 0.0794 | CAAGTCGCCGCCA | - | hypothetical protein |
| AVAEIV_RS05255 | 1082545 | 1082557 | 247 | - | 11.0714 | 6.43E-05 | 0.0794 | CAAGTCGCCGCCA | - | hydantoinase B/oxoprolinase family protein |
| AVAEIV_RS07865 | 1635567 | 1635579 | 225 | - | 11.0714 | 6.43E-05 | 0.0773 | CTCAACGCCATCC | *pyrF* | orotidine-5'-phosphate decarboxylase |
| AVAEIV_RS08660 | 1808089 | 1808101 | -50 | + | 11.0714 | 6.43E-05 | 0.0794 | GCGATGGCCACCA | - | HlyD family secretion protein |
| AVAEIV_RS13755 | 2978112 | 2978124 | 164 | - | 11.0714 | 6.43E-05 | 0.0794 | CTGAATGCCGTCA | - | LysE/ArgO family amino acid transporter |
| AVAEIV_RS16280 | 3547088 | 3547100 | 386 | + | 11.0714 | 6.43E-05 | 0.0794 | CTCAACGCCATCC | - | DNA-binding protein |
| AVAEIV_RS16285 | 3547088 | 3547100 | 186 | + | 11.0714 | 6.43E-05 | 0.0794 | CTCAACGCCATCC | - | hypothetical protein |
| AVAEIV_RS18285 | 3940648 | 3940660 | 160 | - | 11.0714 | 6.43E-05 | 0.0794 | CTGATCGACAGCA | - | hypothetical protein |

| | | | | | | | | | | |
|---|---|---|---|---|---|---|---|---|---|---|
| AVAEIV_RS18290 | 3940648 | 3940660 | -28 | - | 11.0714 | 6.43E-05 | 0.0794 | CTGATCGACAGCA | - | hypothetical protein |
| AVAEIV_RS18815 | 4051959 | 4051971 | 386 | - | 11.0714 | 6.43E-05 | 0.0794 | CCCACGGCCAGCG | *fpr* | ferredoxin-NADP reductase |
| AVAEIV_RS21695 | 4621027 | 4621039 | 267 | + | 11.0714 | 6.43E-05 | 0.0794 | CATTCTGCCATCA | - | PqiC |
| AVAEIV_RS21775 | 4637334 | 4637346 | 194 | + | 11.0714 | 6.43E-05 | 0.0794 | GCGATGGCCACCA | - | ABC transporter permease |
| AVAEIV_RS23585 | 5026389 | 5026401 | 94 | + | 11.0714 | 6.43E-05 | 0.0773 | CAGGTCGGCACCA | - | symmetrical bis(5'-nucleosyl)-tetraphosphatase |
| AVAEIV_RS24145 | 5134663 | 5134675 | 41 | - | 11.0714 | 6.43E-05 | 0.0794 | GGCATTGCCATCA | - | cupin domain-containing protein |
| AVAEIV_RS02225 | 444670 | 444682 | 322 | + | 11.0571 | 6.49E-05 | 0.0773 | GCGCTGGCCATCA | - | hydrogenase maturation protease |
| AVAEIV_RS04315 | 877802 | 877814 | 120 | - | 11.0571 | 6.49E-05 | 0.0794 | CAAAACGCCGCCG | *arfA* | alternative ribosome rescue factor ArfA |
| AVAEIV_RS04320 | 877802 | 877814 | 365 | - | 11.0571 | 6.49E-05 | 0.0773 | CAAAACGCCGCCG | - | thermonuclease family protein |
| AVAEIV_RS05160 | 1060339 | 1060351 | 233 | - | 11.0571 | 6.49E-05 | 0.0794 | CAGCACGCCGCCG | - | DUF2231 domain-containing protein |
| AVAEIV_RS10075 | 2106252 | 2106264 | 224 | - | 11.0571 | 6.49E-05 | 0.0773 | CAGCACGCCGCCG | - | AtpZ/AtpI family protein |
| AVAEIV_RS15300 | 3341778 | 3341790 | 98 | - | 11.0571 | 6.49E-05 | 0.0773 | CGGACCGCCAGCC | - | uracil-xanthine permease family protein |
| AVAEIV_RS18705 | 4029207 | 4029219 | 219 | + | 11.0571 | 6.49E-05 | 0.0794 | CGGCTGGCCGCCA | - | CoA transferase subunit A |
| AVAEIV_RS22025 | 4701358 | 4701370 | 362 | - | 11.0571 | 6.49E-05 | 0.0794 | CGATGCGCCACCG | - | sigma-70 family RNA polymerase sigma factor |
| AVAEIV_RS22030 | 4701358 | 4701370 | 50 | - | 11.0571 | 6.49E-05 | 0.0773 | CGATGCGCCACCG | - | PqiC |
| AVAEIV_RS22625 | 4845554 | 4845566 | 145 | - | 11.0571 | 6.49E-05 | 0.0773 | CAGCACGCCGCCG | *hisI* | phosphoribosyl-AMP cyclohydrolase |
| AVAEIV_RS23430 | 4995880 | 4995892 | 281 | + | 11.0571 | 6.49E-05 | 0.0794 | CAGCACGCCGCCG | - | tetratricopeptide repeat protein |
| AVAEIV_RS25355 | 5400338 | 5400350 | 381 | + | 11.0571 | 6.49E-05 | 0.0773 | CGGCTGGCCGCCA | *tssF* | type VI secretion system baseplate subunit TssF |
| AVAEIV_RS25685 | 5496328 | 5496340 | 332 | + | 11.0571 | 6.49E-05 | 0.0794 | CGGCTGGCCGCCA | - | 3-hydroxyacyl-CoA dehydrogenase family protein |
| AVAEIV_RS00540 | 111135 | 111147 | 375 | + | 11.0429 | 6.55E-05 | 0.0773 | TCGAGCGCCAGCA | - | sigma-70 family RNA polymerase sigma factor |
| AVAEIV_RS00670 | 136253 | 136265 | -25 | + | 11.0429 | 6.55E-05 | 0.0773 | CAATGCGCCATCT | *nifH* | nitrogenase iron protein |
| AVAEIV_RS01025 | 200393 | 200405 | 311 | - | 11.0429 | 6.55E-05 | 0.0773 | CAGCCGGCCGTCA | - | acetyl-CoA carboxylase biotin carboxylase subunit |
| AVAEIV_RS05520 | 1142211 | 1142223 | 88 | + | 11.0429 | 6.55E-05 | 0.0773 | CACCCGGCCAGCG | *pgeF* | peptidoglycan editing factor PgeF |
| AVAEIV_RS06925 | 1424627 | 1424639 | 89 | - | 11.0429 | 6.55E-05 | 0.0773 | CAGCACGGCATCG | *ftsW* | putative lipid II flippase FtsW |
| AVAEIV_RS07990 | 1659415 | 1659427 | 279 | - | 11.0429 | 6.55E-05 | 0.0773 | CTGATCGCCATGG | - | ABC transporter ATP-binding protein |
| AVAEIV_RS08005 | 1663745 | 1663757 | 323 | - | 11.0429 | 6.55E-05 | 0.0794 | CCAACGGCCACCA | - | heavy metal response regulator transcription factor |
| AVAEIV_RS09045 | 1881705 | 1881717 | 193 | - | 11.0429 | 6.55E-05 | 0.0794 | CGGGTCGCCGCCA | - | 3-hydroxybutyrate dehydrogenase |
| AVAEIV_RS10385 | 2170620 | 2170632 | 213 | - | 11.0429 | 6.55E-05 | 0.0773 | CAGCACGGCATCG | - | FAD-dependent oxidoreductase |
| AVAEIV_RS12700 | 2697224 | 2697236 | 235 | - | 11.0429 | 6.55E-05 | 0.0794 | CACCTGGCCAGCT | - | DUF485 domain-containing protein |
| AVAEIV_RS12705 | 2698968 | 2698980 | 219 | - | 11.0429 | 6.55E-05 | 0.0794 | TCGAGCGCCAGCA | - | acyl-CoA synthetase |
| AVAEIV_RS12710 | 2698968 | 2698980 | -50 | - | 11.0429 | 6.55E-05 | 0.0773 | TCGAGCGCCAGCA | - | PAS domain S-box protein |
| AVAEIV_RS17485 | 3787158 | 3787170 | 53 | - | 11.0429 | 6.55E-05 | 0.0794 | CCAACGGCCACCA | - | ABC transporter ATP-binding protein |
| AVAEIV_RS18685 | 4024442 | 4024454 | 182 | + | 11.0429 | 6.55E-05 | 0.0794 | CAGCACGGCATCG | *pcaD* | 3-oxoadipate enol-lactonase |
| AVAEIV_RS19060 | 4106139 | 4106151 | 212 | + | 11.0429 | 6.55E-05 | 0.0794 | CAGCACGGCATCG | *tsf* | translation elongation factor Ts |
| AVAEIV_RS20015 | 4267548 | 4267560 | -35 | - | 11.0429 | 6.55E-05 | 0.0773 | CCGCCGGCCACCA | - | SDR family oxidoreductase |
| AVAEIV_RS20880 | 4440538 | 4440550 | 335 | - | 11.0429 | 6.55E-05 | 0.0794 | CAGCACGGCATCG | *pqqB* | pyrroloquinoline quinone biosynthesis protein PqqB |
| AVAEIV_RS20885 | 4440538 | 4440550 | 218 | - | 11.0429 | 6.55E-05 | 0.0794 | CAGCACGGCATCG | *pqqA* | pyrroloquinoline quinone precursor peptide PqqA |
| AVAEIV_RS21560 | 4586297 | 4586309 | 28 | + | 11.0429 | 6.55E-05 | 0.0773 | CACCTGGCCAGCT | - | LutB/LldF family L-lactate oxidation iron-sulfur protein |
| AVAEIV_RS22600 | 4840476 | 4840488 | 259 | + | 11.0429 | 6.55E-05 | 0.0773 | CTGATCGCCATGG | *hslU* | ATP-dependent protease ATPase subunit HslU |
| AVAEIV_RS23560 | 5019788 | 5019800 | 304 | - | 11.0429 | 6.55E-05 | 0.0773 | CTGATCGCCATGG | - | LPS-assembly protein LptD |
| AVAEIV_RS25395 | 5411329 | 5411341 | 282 | + | 11.0429 | 6.55E-05 | 0.0794 | CATTTCGCTATCA | *nifX* | nitrogen fixation protein NifX |
| AVAEIV_RS25400 | 5411329 | 5411341 | -3 | + | 11.0429 | 6.55E-05 | 0.0794 | CATTTCGCTATCA | - | RnfH family protein |
| AVAEIV_RS25715 | 5502181 | 5502193 | 187 | - | 11.0429 | 6.55E-05 | 0.0794 | CAGAGCGCGATCG | *lpxO* | lipid A hydroxylase LpxO |
| AVAEIV_RS01200 | 234701 | 234713 | 273 | + | 11.0286 | 6.62E-05 | 0.0794 | CGCCGTGCCATCG | - | ABC transporter substrate-binding protein |
| AVAEIV_RS01205 | 234701 | 234713 | 99 | + | 11.0286 | 6.62E-05 | 0.0773 | CGCCGTGCCATCG | - | hypothetical protein |
| AVAEIV_RS01380 | 270657 | 270669 | 343 | + | 11.0286 | 6.62E-05 | 0.0794 | CCGCGGGCCACCG | - | YicC/YloC family endoribonuclease |
| AVAEIV_RS01775 | 350513 | 350525 | 324 | + | 11.0286 | 6.62E-05 | 0.0794 | GCGTTGGCCAGCA | - | sigma-54-dependent transcriptional regulator |
| AVAEIV_RS02255 | 452216 | 452228 | 364 | + | 11.0286 | 6.62E-05 | 0.0794 | CAGCGGGCCGTCG | - | NAD(P)/FAD-dependent oxidoreductase |
| AVAEIV_RS06315 | 1289984 | 1289996 | -45 | + | 11.0286 | 6.62E-05 | 0.0794 | CACGCCGCCAGCG | - | NAD(P)/FAD-dependent oxidoreductase |
| AVAEIV_RS08180 | 1700244 | 1700256 | 349 | + | 11.0286 | 6.62E-05 | 0.0773 | GAACTGGCCATCA | - | containing protein |
| AVAEIV_RS08420 | 1759172 | 1759184 | 62 | - | 11.0286 | 6.62E-05 | 0.0773 | CGCCGTGCCATCG | - | oxidoreductase |
| AVAEIV_RS08965 | 1864703 | 1864715 | 62 | - | 11.0286 | 6.62E-05 | 0.0773 | CGCCGTGCCATCG | - | oxidoreductase |

| | | | | | | | | | | |
|---|---|---|---|---|---|---|---|---|---|---|
| AVAEIV_RS11155 | 2355443 | 2355455 | 338 | + | 11.0286 | 6.62E-05 | 0.0773 | CAGATCGCCGACA | - | LysR family transcriptional regulator |
| AVAEIV_RS15690 | 3430495 | 3430507 | 25 | + | 11.0286 | 6.62E-05 | 0.0794 | CCGGATGCCAGCG | *ssuB* | protein |
| AVAEIV_RS17170 | 3721819 | 3721831 | 84 | - | 11.0286 | 6.62E-05 | 0.0794 | CAGCGGGCCGTCG | - | SPOR domain-containing protein |
| AVAEIV_RS17390 | 3768770 | 3768782 | 326 | + | 11.0286 | 6.62E-05 | 0.0794 | GAACTGGCCATCA | - | hypothetical protein |
| AVAEIV_RS17845 | 3861796 | 3861808 | 253 | - | 11.0286 | 6.62E-05 | 0.0773 | CAGCAGGCCACCC | - | OprD family porin |
| AVAEIV_RS18320 | 3944037 | 3944049 | 139 | - | 11.0286 | 6.62E-05 | 0.0794 | CACGCCGCCAGCG | - | DUF6387 family protein |
| AVAEIV_RS18645 | 4013046 | 4013058 | 274 | - | 11.0286 | 6.62E-05 | 0.0773 | CAACTCGCCTTCA | - | shikimate dehydrogenase |
| AVAEIV_RS19035 | 4102142 | 4102154 | 263 | + | 11.0286 | 6.62E-05 | 0.0794 | CAGCAGGCCACCC | *ispC* | 1-deoxy-D-xylulose-5-phosphate reductoisomerase |
| AVAEIV_RS19065 | 4106903 | 4106915 | 98 | - | 11.0286 | 6.62E-05 | 0.0794 | CGGTTCGCGAGCA | *rpsB* | 30S ribosomal protein S2 |
| AVAEIV_RS19070 | 4106903 | 4106915 | 153 | - | 11.0286 | 6.62E-05 | 0.0773 | CGGTTCGCGAGCA | *map* | type I methionyl aminopeptidase |
| AVAEIV_RS20005 | 4265830 | 4265842 | 188 | - | 11.0286 | 6.62E-05 | 0.0794 | CCGGATGCCAGCG | - | LysR family transcriptional regulator |
| AVAEIV_RS20250 | 4321343 | 4321355 | 355 | + | 11.0286 | 6.62E-05 | 0.0794 | CGGGTCGGCATCA | *yajC* | preprotein translocase subunit YajC |
| AVAEIV_RS21685 | 4617946 | 4617958 | 176 | + | 11.0286 | 6.62E-05 | 0.0773 | CAGATCGCCGACA | - | 2-hydroxy-3-oxopropionate reductase |
| AVAEIV_RS21970 | 4686301 | 4686313 | 295 | - | 11.0286 | 6.62E-05 | 0.0773 | CAGCGGGCCGTCG | - | acetate/propionate family kinase |
| AVAEIV_RS22735 | 4867893 | 4867905 | 88 | - | 11.0286 | 6.62E-05 | 0.0794 | CAAAATGCCATTA | - | hypothetical protein |
| AVAEIV_RS24860 | 5288553 | 5288565 | 347 | + | 11.0286 | 6.62E-05 | 0.0794 | CCGGCCGCCACCA | - | hydroxymethylglutaryl-CoA lyase |
| AVAEIV_RS25185 | 5363384 | 5363396 | 109 | + | 11.0286 | 6.62E-05 | 0.0794 | CAGCGGGCCGTCG | *hypA* | hydrogenase maturation nickel metallochaperone HypA |
| AVAEIV_RS00235 | 47160 | 47172 | 11 | - | 11.0143 | 6.68E-05 | 0.0773 | CTGTTCGCCACCC | - | ATP-binding cassette domain-containing protein |
| AVAEIV_RS00470 | 97772 | 97784 | -49 | - | 11.0143 | 6.68E-05 | 0.0794 | CTGTTCGCCACCC | - | COX15/CtaA family protein |
| AVAEIV_RS01440 | 279245 | 279257 | 381 | - | 11.0143 | 6.68E-05 | 0.0773 | CGCGGGGCCAGCA | - | DUF423 domain-containing protein |
| AVAEIV_RS01500 | 288821 | 288833 | 51 | - | 11.0143 | 6.68E-05 | 0.0794 | CCAATCGCCAACA | - | YggS family pyridoxal phosphate-dependent enzyme |
| AVAEIV_RS01505 | 288821 | 288833 | 8 | - | 11.0143 | 6.68E-05 | 0.0773 | CCAATCGCCAACA | - | type IV pilus twitching motility protein PilT |
| AVAEIV_RS01765 | 348269 | 348281 | 315 | - | 11.0143 | 6.68E-05 | 0.0794 | GCCATCGCCACCA | *dctP* | C4-dicarboxylate TRAP substrate-binding protein DctP |
| AVAEIV_RS02555 | 513735 | 513747 | 339 | - | 11.0143 | 6.68E-05 | 0.0794 | CAGTCGGCCGGCA | - | glycosyl hydrolase family 8 |
| AVAEIV_RS03600 | 725705 | 725717 | 229 | + | 11.0143 | 6.68E-05 | 0.0794 | CAGGGCGCCGTCG | - | amino acid ABC transporter permease |
| AVAEIV_RS03660 | 737783 | 737795 | 94 | - | 11.0143 | 6.68E-05 | 0.0794 | CGCATGGCCAGCT | - | rhodanese-like domain-containing protein |
| AVAEIV_RS03940 | 794597 | 794609 | 339 | + | 11.0143 | 6.68E-05 | 0.0773 | GCCATCGCCACCA | *ubiX* | flavin prenyltransferase UbiX |
| AVAEIV_RS04490 | 918075 | 918087 | 288 | - | 11.0143 | 6.68E-05 | 0.0794 | GAGAACGCCACCG | - | siderophore biosynthesis protein PvsA |
| AVAEIV_RS04665 | 953826 | 953838 | 256 | - | 11.0143 | 6.68E-05 | 0.0794 | CGGATCGCCTCCA | *cueR* | Cu(I)-responsive transcriptional regulator |
| AVAEIV_RS04855 | 991334 | 991346 | 314 | + | 11.0143 | 6.68E-05 | 0.0794 | GGGATGGCCACCA | - | RNA polymerase sigma factor |
| AVAEIV_RS05390 | 1115891 | 1115903 | 386 | + | 11.0143 | 6.68E-05 | 0.0773 | CAGGGCGCCGTCG | *lptF* | LPS export ABC transporter permease LptF |
| AVAEIV_RS06435 | 1318615 | 1318627 | 263 | - | 11.0143 | 6.68E-05 | 0.0794 | CAGGGCGCCGTCG | - | protein |
| AVAEIV_RS06870 | 1414613 | 1414625 | 275 | + | 11.0143 | 6.68E-05 | 0.0794 | CATTTCGGCATCA | - | penicillin-binding protein activator |
| AVAEIV_RS07715 | 1607530 | 1607542 | 377 | - | 11.0143 | 6.68E-05 | 0.0773 | CCGGGCGCCACCG | *acpP* | acyl carrier protein |
| AVAEIV_RS09635 | 2013000 | 2013012 | 211 | + | 11.0143 | 6.68E-05 | 0.0794 | CCGTAGGCCAGCC | - | DUF58 domain-containing protein |
| AVAEIV_RS10320 | 2157059 | 2157071 | 330 | + | 11.0143 | 6.68E-05 | 0.0794 | CGGATCGCCTCCA | - | SDR family oxidoreductase |
| AVAEIV_RS10515 | 2199049 | 2199061 | 253 | - | 11.0143 | 6.68E-05 | 0.0773 | CTCATGGCCGGCA | - | helix-turn-helix transcriptional regulator |
| AVAEIV_RS11100 | 2343187 | 2343199 | 138 | + | 11.0143 | 6.68E-05 | 0.0794 | CCCTTCGGCAGCA | - | TonB-dependent siderophore receptor |
| AVAEIV_RS14060 | 3057600 | 3057612 | 314 | + | 11.0143 | 6.68E-05 | 0.0794 | CGGATCGCCTCCA | - | dehydrogenase |
| AVAEIV_RS15050 | 3277815 | 3277827 | 234 | - | 11.0143 | 6.68E-05 | 0.0794 | CTGTTCGCCACCC | - | GspMb/PilO family protein |
| AVAEIV_RS15185 | 3306442 | 3306454 | 286 | - | 11.0143 | 6.68E-05 | 0.0794 | CAGGGCGCCGTCG | - | DUF2147 domain-containing protein |
| AVAEIV_RS15480 | 3383619 | 3383631 | 87 | - | 11.0143 | 6.68E-05 | 0.0773 | CGCACGGCCAGCG | - | sugar transferase |
| AVAEIV_RS17505 | 3788670 | 3788682 | 219 | - | 11.0143 | 6.68E-05 | 0.0773 | CTCTGCGCCAGCG | - | class II fumarate hydratase |
| AVAEIV_RS17590 | 3811177 | 3811189 | -49 | + | 11.0143 | 6.68E-05 | 0.0773 | CTGTTCGCCACCC | - | beta (1-6) glucans synthase |
| AVAEIV_RS18970 | 4084248 | 4084260 | 147 | + | 11.0143 | 6.68E-05 | 0.0794 | CAGGGCGCCGTCG | *kdsA* | 3-deoxy-8-phosphooctulonate synthase |
| AVAEIV_RS19730 | 4205513 | 4205525 | 353 | + | 11.0143 | 6.68E-05 | 0.0794 | CGGCACGCCGTCG | - | hypothetical protein |
| AVAEIV_RS19890 | 4237606 | 4237618 | 356 | + | 11.0143 | 6.68E-05 | 0.0773 | CAGTCGGCCGGCA | *mltF* | membrane-bound lytic murein transglycosylase MltF |
| AVAEIV_RS20450 | 4345764 | 4345776 | 340 | - | 11.0143 | 6.68E-05 | 0.0773 | CAGGGCGCCGTCG | - | DUF4105 domain-containing protein |
| AVAEIV_RS20565 | 4373600 | 4373612 | 20 | + | 11.0143 | 6.68E-05 | 0.0794 | CGGTGCGCCATCT | - | MBL fold metallo-hydrolase |
| AVAEIV_RS22450 | 4799210 | 4799222 | 286 | + | 11.0143 | 6.68E-05 | 0.0773 | CCGGGCGCCACCG | - | 2-hydroxymuconate tautomerase family protein |
| AVAEIV_RS22785 | 4877613 | 4877625 | 246 | - | 11.0143 | 6.68E-05 | 0.0773 | CGGATCGCCTCCA | *tpiA* | triose-phosphate isomerase |
| AVAEIV_RS24040 | 5116303 | 5116315 | 284 | - | 11.0143 | 6.68E-05 | 0.0773 | CAGGGCGCCGTCG | - | HAD family hydrolase |

| | | | | | | | | | | |
|---|---|---|---|---|---|---|---|---|---|---|
| AVAEIV_RS24065 | 5121226 | 5121238 | 335 | - | 11.0143 | 6.68E-05 | 0.0794 | CAGTCGGCCGGCA | *glnK* | P-II family nitrogen regulator |
| AVAEIV_RS24070 | 5121226 | 5121238 | 16 | - | 11.0143 | 6.68E-05 | 0.0773 | CAGTCGGCCGGCA | - | accessory factor UbiK family protein |
| AVAEIV_RS24075 | 5121226 | 5121238 | 323 | - | 11.0143 | 6.68E-05 | 0.0773 | CAGTCGGCCGGCA | - | YifB family Mg chelatase-like AAA ATPase |
| AVAEIV_RS24240 | 5154544 | 5154556 | 157 | - | 11.0143 | 6.68E-05 | 0.0794 | CGGATCGCCTCCA | - | DMT family transporter |
| AVAEIV_RS24245 | 5154544 | 5154556 | -47 | - | 11.0143 | 6.68E-05 | 0.0773 | CGGATCGCCTCCA | - | PLP-dependent aminotransferase family protein |
| AVAEIV_RS24950 | 5310012 | 5310024 | 246 | + | 11.0143 | 6.68E-05 | 0.0794 | CGGATCGCCTCCA | *tpiA* | triose-phosphate isomerase |
| AVAEIV_RS25010 | 5324191 | 5324203 | 148 | - | 11.0143 | 6.68E-05 | 0.0773 | CCGGGCGCCACCG | - | Gfo/Idh/MocA family protein |
| AVAEIV_RS25855 | 5544269 | 5544281 | 141 | - | 11.0143 | 6.68E-05 | 0.0773 | TAGCGCGCCAGCA | - | ABC transporter substrate-binding protein |
| AVAEIV_RS02460 | 491033 | 491045 | -31 | - | 11 | 6.75E-05 | 0.0794 | CGCCTCGCCGCCA | - | efflux RND transporter periplasmic adaptor subunit |
| AVAEIV_RS02950 | 610514 | 610526 | 219 | - | 11 | 6.75E-05 | 0.0794 | CCAGGCGCCATCG | *dsbG* | thiol:disulfide interchange protein DsbG |
| AVAEIV_RS09700 | 2033378 | 2033390 | 324 | - | 11 | 6.75E-05 | 0.0794 | CGCTGCGCCATCC | - | deoxyguanosinetriphosphate triphosphohydrolase |
| AVAEIV_RS12775 | 2715493 | 2715505 | 313 | + | 11 | 6.75E-05 | 0.0794 | GAGCACGCCATCG | - | cupin-like domain-containing protein |
| AVAEIV_RS12790 | 2718238 | 2718250 | 239 | - | 11 | 6.75E-05 | 0.0775 | CACACCGCCGCCA | - | formylglycine-generating enzyme family protein |
| AVAEIV_RS13150 | 2852864 | 2852876 | -43 | - | 11 | 6.75E-05 | 0.0794 | CAAGCCGCCACCA | *hutI* | imidazolonepropionase |
| AVAEIV_RS13810 | 2992144 | 2992156 | 303 | + | 11 | 6.75E-05 | 0.0794 | TACAAGGCCAGCA | *tssG* | type VI secretion system baseplate subunit TssG |
| AVAEIV_RS15390 | 3361498 | 3361510 | 68 | + | 11 | 6.75E-05 | 0.0794 | CCGAATGCCGCCA | *pdxB* | 4-phosphoerythronate dehydrogenase PdxB |
| AVAEIV_RS15395 | 3361498 | 3361510 | 69 | + | 11 | 6.75E-05 | 0.0775 | CCGAATGCCGCCA | - | MATE family efflux transporter |
| AVAEIV_RS17635 | 3818952 | 3818964 | 223 | - | 11 | 6.75E-05 | 0.0775 | CCGATGGCGAGCA | - | M18 family aminopeptidase |
| AVAEIV_RS19285 | 4145736 | 4145748 | 154 | - | 11 | 6.75E-05 | 0.0794 | CAGATCGACGGCA | - | hypothetical protein |
| AVAEIV_RS20095 | 4290712 | 4290724 | 360 | + | 11 | 6.75E-05 | 0.0775 | GAGCACGCCATCG | - | endonuclease I family protein |
| AVAEIV_RS21280 | 4524021 | 4524033 | -21 | - | 11 | 6.75E-05 | 0.0775 | CAGAAGGCCAGTG | - | DUF4124 domain-containing protein |
| AVAEIV_RS21305 | 4530842 | 4530854 | 136 | + | 11 | 6.75E-05 | 0.0794 | CCATCGGCCAGCA | - | ChaN family lipoprotein |
| AVAEIV_RS21555 | 4585544 | 4585556 | -40 | + | 11 | 6.75E-05 | 0.0775 | CCCAACGCCACCC | - | (Fe-S)-binding protein |
| AVAEIV_RS23065 | 4934426 | 4934438 | 318 | - | 11 | 6.75E-05 | 0.0775 | CCATCGGCCAGCA | - | phage tail protein |
| AVAEIV_RS25725 | 5502604 | 5502616 | 245 | - | 11 | 6.75E-05 | 0.0775 | CCGATGGCGAGCA | - | bifunctional diguanylate cyclase/phosphodiesterase |
| AVAEIV_RS26115 | 5614898 | 5614910 | 112 | - | 11 | 6.75E-05 | 0.0794 | CCATCGGCCAGCA | *mnmE* | synthesis GTPase MnmE |
| AVAEIV_RS02045 | 408060 | 408072 | 117 | + | 10.9857 | 6.82E-05 | 0.0794 | TCGCTGGCCATCA | - | helix-turn-helix domain-containing protein |
| AVAEIV_RS02555 | 513774 | 513786 | 378 | + | 10.9857 | 6.82E-05 | 0.0794 | CGAACGGCCACCA | - | glycosyl hydrolase family 8 |
| AVAEIV_RS02810 | 575699 | 575711 | 309 | - | 10.9857 | 6.82E-05 | 0.0775 | TGGAGCGCCAGCA | - | M48 family metallopeptidase |
| AVAEIV_RS03910 | 788310 | 788322 | 380 | + | 10.9857 | 6.82E-05 | 0.0794 | CCCCCCGCCACCA | *adhP* | alcohol dehydrogenase AdhP |
| AVAEIV_RS03915 | 788310 | 788322 | 46 | + | 10.9857 | 6.82E-05 | 0.0794 | CCCCCCGCCACCA | - | hypothetical protein |
| AVAEIV_RS05525 | 1143164 | 1143176 | 36 | + | 10.9857 | 6.82E-05 | 0.0775 | CGGCAGGCCATCC | *clpB* | ATP-dependent chaperone ClpB |
| AVAEIV_RS06255 | 1277144 | 1277156 | 21 | - | 10.9857 | 6.82E-05 | 0.0794 | CGACACGCCACCG | - | putative protein CP11 |
| AVAEIV_RS08730 | 1821214 | 1821226 | 136 | - | 10.9857 | 6.82E-05 | 0.0794 | CCGCATGCCGTCA | - | hypothetical protein |
| AVAEIV_RS11405 | 2413584 | 2413596 | 84 | + | 10.9857 | 6.82E-05 | 0.0775 | CGCCTCGGCATCA | - | GNAT family N-acetyltransferase |
| AVAEIV_RS12120 | 2571883 | 2571895 | 332 | - | 10.9857 | 6.82E-05 | 0.0794 | CGGCCGGCCACCA | - | hypothetical protein |
| AVAEIV_RS12125 | 2571883 | 2571895 | 70 | - | 10.9857 | 6.82E-05 | 0.0775 | CGGCCGGCCACCA | - | AEC family transporter |
| AVAEIV_RS12725 | 2702329 | 2702341 | 138 | + | 10.9857 | 6.82E-05 | 0.0794 | CCATGGGCCAGCG | - | DUF3325 domain-containing protein |
| AVAEIV_RS16095 | 3508290 | 3508302 | -47 | - | 10.9857 | 6.82E-05 | 0.0775 | CCATGGGCCAGCG | - | acyl-CoA dehydrogenase family protein |
| AVAEIV_RS25935 | 5571846 | 5571858 | 103 | + | 10.9857 | 6.82E-05 | 0.0775 | CTGGACGCCAGCG | - | ABC transporter substrate-binding protein |
| AVAEIV_RS00430 | 87192 | 87204 | 374 | + | 10.9714 | 6.88E-05 | 0.0775 | CCGGTGGCCGGCA | - | hypothetical protein |
| AVAEIV_RS01860 | 368909 | 368921 | 337 | - | 10.9714 | 6.88E-05 | 0.0775 | CACCGCGCCGTCG | - | RNA polymerase sigma factor |
| AVAEIV_RS03465 | 699192 | 699204 | 57 | - | 10.9714 | 6.88E-05 | 0.0775 | AAGTTGGCCATCA | - | NUDIX hydrolase |
| AVAEIV_RS03890 | 784884 | 784896 | 12 | + | 10.9714 | 6.88E-05 | 0.0794 | CAGCTGGCGAGCA | - | EAL domain-containing protein |
| AVAEIV_RS07450 | 1552158 | 1552170 | 120 | - | 10.9714 | 6.88E-05 | 0.0775 | CTGAGCGCCAGCC | - | efflux transporter outer membrane subunit |
| AVAEIV_RS08930 | 1856396 | 1856408 | 337 | + | 10.9714 | 6.88E-05 | 0.0794 | CGGTTCGCCTGCA | - | response regulator transcription factor |
| AVAEIV_RS11845 | 2509830 | 2509842 | 381 | - | 10.9714 | 6.88E-05 | 0.0775 | CCGATCGCCACGG | *sigX* | RNA polymerase sigma factor SigX |
| AVAEIV_RS14015 | 3041998 | 3042010 | 33 | + | 10.9714 | 6.88E-05 | 0.0775 | CGACCGGCCATCA | *ppnN* | nucleotide 5'-monophosphate nucleosidase PpnN |
| AVAEIV_RS14110 | 3069315 | 3069327 | 128 | + | 10.9714 | 6.88E-05 | 0.0794 | CACCGCGCCGTCG | *xylB* | xylulokinase |
| AVAEIV_RS16435 | 3570060 | 3570072 | 297 | + | 10.9714 | 6.88E-05 | 0.0794 | CAGATCGCCGTGG | - | LysR family transcriptional regulator |
| AVAEIV_RS16895 | 3658465 | 3658477 | 264 | + | 10.9714 | 6.88E-05 | 0.0794 | CGGGATGCCAGCG | - | CopG family transcriptional regulator |
| AVAEIV_RS18075 | 3906585 | 3906597 | 105 | - | 10.9714 | 6.88E-05 | 0.0794 | CGGTTCGCCTGCA | *queC* | 7-cyano-7-deazaguanine synthase QueC |

| | | | | | | | | | | |
|---|---|---|---|---|---|---|---|---|---|---|
| AVAEIV_RS18220 | 3930169 | 3930181 | 32 | - | 10.9714 | 6.88E-05 | 0.0775 | CCGGTGGCCGGCA | - | DUF3108 domain-containing protein |
| AVAEIV_RS18925 | 4076194 | 4076206 | -40 | - | 10.9714 | 6.88E-05 | 0.0794 | CACAGCGGCATCG | - | methyltransferase |
| AVAEIV_RS19420 | 4161162 | 4161174 | 350 | - | 10.9714 | 6.88E-05 | 0.0794 | CAGAGGGCCACCT | - | hypothetical protein |
| AVAEIV_RS19510 | 4171582 | 4171594 | 17 | - | 10.9714 | 6.88E-05 | 0.0775 | CCGGTGGCCGGCA | - | hypothetical protein |
| AVAEIV_RS20265 | 4322967 | 4322979 | 331 | + | 10.9714 | 6.88E-05 | 0.0775 | CAGCTTGCCGCCG | - | tRNA-Leu |
| AVAEIV_RS22330 | 4769353 | 4769365 | 224 | + | 10.9714 | 6.88E-05 | 0.0794 | CAGATCGCCGTGG | - | carbamoyltransferase |
| AVAEIV_RS23315 | 4970367 | 4970379 | 213 | + | 10.9714 | 6.88E-05 | 0.0775 | CACCGCGCCGTCG | - | CapA family protein |
| AVAEIV_RS24755 | 5259390 | 5259402 | 344 | - | 10.9714 | 6.88E-05 | 0.0775 | CACCGCGCCGTCG | - | aldehyde dehydrogenase (NADP(+)) |
| AVAEIV_RS25330 | 5397703 | 5397715 | 262 | + | 10.9714 | 6.88E-05 | 0.0794 | GGGAACGCCATCG | - | hypothetical protein |
| AVAEIV_RS25460 | 5422444 | 5422456 | 205 | + | 10.9714 | 6.88E-05 | 0.0775 | CTGAGCGCCAGCC | - | nitrogen fixation protein NifQ |
| AVAEIV_RS25765 | 5519221 | 5519233 | 163 | - | 10.9714 | 6.88E-05 | 0.0794 | CTGAGCGCCAGCC | - | Arc family DNA-binding protein |
| AVAEIV_RS25770 | 5519221 | 5519233 | 109 | - | 10.9714 | 6.88E-05 | 0.0775 | CTGAGCGCCAGCC | *trhA* | PAQR family membrane homeostasis protein TrhA |
| AVAEIV_RS00415 | 85287 | 85299 | 155 | - | 10.9571 | 6.95E-05 | 0.0775 | CTGATCGCCGCCG | - | Fur family transcriptional regulator |
| AVAEIV_RS01295 | 252962 | 252974 | -20 | - | 10.9571 | 6.95E-05 | 0.0794 | CCGTTCGCCGTCC | - | AAA family ATPase |
| AVAEIV_RS01475 | 285169 | 285181 | 123 | - | 10.9571 | 6.95E-05 | 0.0794 | CTGATCGCCGCCG | *metW* | methionine biosynthesis protein MetW |
| AVAEIV_RS01770 | 348940 | 348952 | 294 | - | 10.9571 | 6.95E-05 | 0.0794 | CTGATCGCCGCCG | - | glycine zipper domain-containing protein |
| AVAEIV_RS01775 | 350270 | 350282 | 81 | + | 10.9571 | 6.95E-05 | 0.0794 | TCGTTGGCCAGCA | - | sigma-54-dependent transcriptional regulator |
| AVAEIV_RS02290 | 458880 | 458892 | 377 | - | 10.9571 | 6.95E-05 | 0.0794 | CTGATCGCCGCCG | - | AsmA family protein |
| AVAEIV_RS02710 | 554394 | 554406 | 366 | - | 10.9571 | 6.95E-05 | 0.0794 | CCGATCGCCCTCA | - | sugar transferase |
| AVAEIV_RS03480 | 701494 | 701506 | 122 | - | 10.9571 | 6.95E-05 | 0.0775 | CGGTAGGCCAGCC | - | nucleotidyltransferase family protein |
| AVAEIV_RS03560 | 718283 | 718295 | 272 | + | 10.9571 | 6.95E-05 | 0.0775 | CATATGGCTATCA | - | patatin-like phospholipase family protein |
| AVAEIV_RS03675 | 740612 | 740624 | 131 | + | 10.9571 | 6.95E-05 | 0.0794 | CTGATCGCCGCCG | *queG* | tRNA epoxyqueuosine(34) reductase QueG |
| AVAEIV_RS06635 | 1371238 | 1371250 | 276 | - | 10.9571 | 6.95E-05 | 0.0775 | CAGCTTGGCATCG | - | carbon-nitrogen hydrolase family protein |
| AVAEIV_RS08000 | 1662890 | 1662902 | 148 | - | 10.9571 | 6.95E-05 | 0.0794 | CTGATCGCCGCCG | - | heavy metal sensor histidine kinase |
| AVAEIV_RS08160 | 1694850 | 1694862 | 254 | + | 10.9571 | 6.95E-05 | 0.0775 | CCGAAGGCCGGCG | - | bifunctional prephenate dehydrogenase/3-phosphoshikimate 1-carboxyvinyltransferase |
| AVAEIV_RS08375 | 1747369 | 1747381 | 254 | - | 10.9571 | 6.95E-05 | 0.0775 | CTGATCGCCGCCG | - | SIMPL domain-containing protein |
| AVAEIV_RS08720 | 1818849 | 1818861 | 65 | - | 10.9571 | 6.95E-05 | 0.0775 | CAACTTGCCGTCG | - | S1 RNA-binding domain-containing protein |
| AVAEIV_RS10370 | 2168465 | 2168477 | 44 | + | 10.9571 | 6.95E-05 | 0.0794 | CCGAAGGCCGGCG | - | MFS transporter |
| AVAEIV_RS11180 | 2360815 | 2360827 | 257 | + | 10.9571 | 6.95E-05 | 0.0794 | CAGAAGGCTAGCG | - | hypothetical protein |
| AVAEIV_RS12225 | 2590063 | 2590075 | 363 | + | 10.9571 | 6.95E-05 | 0.0775 | CTGATCGCCGCCG | *flgL* | flagellar hook-associated protein FlgL |
| AVAEIV_RS14915 | 3252879 | 3252891 | 299 | + | 10.9571 | 6.95E-05 | 0.0794 | CTGATCGCCGCCG | - | containing protein |
| AVAEIV_RS15400 | 3363568 | 3363580 | 156 | + | 10.9571 | 6.95E-05 | 0.0775 | AAGAACGCCAGCA | - | low molecular weight protein-tyrosine-phosphatase |
| AVAEIV_RS15695 | 3431380 | 3431392 | 119 | + | 10.9571 | 6.95E-05 | 0.0794 | TCGTTGGCCAGCA | *ssuC* | aliphatic sulfonate ABC transporter permease SsuC |
| AVAEIV_RS18315 | 3943058 | 3943070 | 159 | - | 10.9571 | 6.95E-05 | 0.0794 | AAGAACGCCAGCA | - | AlpA family phage regulatory protein |
| AVAEIV_RS18485 | 3980019 | 3980031 | -19 | + | 10.9571 | 6.95E-05 | 0.0775 | CACAGTGCCATTA | *olsB* | L-ornithine N(alpha)-acyltransferase |
| AVAEIV_RS18595 | 4005038 | 4005050 | 63 | + | 10.9571 | 6.95E-05 | 0.0775 | CCGCACGCCATGA | - | hypothetical protein |
| AVAEIV_RS19845 | 4228204 | 4228216 | 365 | + | 10.9571 | 6.95E-05 | 0.0794 | CTGATCGCCGCCG | *ffh* | signal recognition particle protein |
| AVAEIV_RS21080 | 4483646 | 4483658 | 352 | - | 10.9571 | 6.95E-05 | 0.0794 | CTGATCGCCGCCG | - | 2-keto-4-pentenoate hydratase |
| AVAEIV_RS22555 | 4829085 | 4829097 | 339 | - | 10.9571 | 6.95E-05 | 0.0794 | CAGCTTGGCATCG | - | pilus assembly protein PilM |
| AVAEIV_RS23835 | 5080732 | 5080744 | 313 | - | 10.9571 | 6.95E-05 | 0.0794 | CTGATCGCCGCCG | *ubiH* | 2-octaprenyl-6-methoxyphenyl hydroxylase |
| AVAEIV_RS24220 | 5149372 | 5149384 | -40 | - | 10.9571 | 6.95E-05 | 0.0794 | CTGATCGCCGCCG | - | dihydrofolate reductase |
| AVAEIV_RS24225 | 5149372 | 5149384 | 157 | - | 10.9571 | 6.95E-05 | 0.0775 | CTGATCGCCGCCG | - | glycerophosphodiester phosphodiesterase |
| AVAEIV_RS26070 | 5606071 | 5606083 | 76 | - | 10.9571 | 6.95E-05 | 0.0794 | CAGTTCGACACCG | - | protein |
| AVAEIV_RS01505 | 288666 | 288678 | 163 | + | 10.9429 | 7.01E-05 | 0.0775 | CTCACGGCCAGCA | - | type IV pilus twitching motility protein PilT |
| AVAEIV_RS01775 | 350284 | 350296 | 95 | + | 10.9429 | 7.01E-05 | 0.0794 | CAGTTCGCCGCCC | - | sigma-54-dependent transcriptional regulator |
| AVAEIV_RS03295 | 668190 | 668202 | 242 | - | 10.9429 | 7.01E-05 | 0.0794 | CGATCGGCCAGCA | - | ABC transporter permease |
| AVAEIV_RS03745 | 754355 | 754367 | 380 | + | 10.9429 | 7.01E-05 | 0.0794 | CACTGCGCCGGCG | - | tRNA-Leu |
| AVAEIV_RS05630 | 1158174 | 1158186 | 40 | - | 10.9429 | 7.01E-05 | 0.0775 | CTCACGGCCAGCA | - | CopG family antitoxin |
| AVAEIV_RS05635 | 1158174 | 1158186 | 255 | - | 10.9429 | 7.01E-05 | 0.0775 | CTCACGGCCAGCA | - | hypothetical protein |
| AVAEIV_RS06020 | 1226667 | 1226679 | 111 | - | 10.9429 | 7.01E-05 | 0.0794 | CCGATCGCCACTG | - | methyl-accepting chemotaxis protein |
| AVAEIV_RS06025 | 1226667 | 1226679 | 346 | - | 10.9429 | 7.01E-05 | 0.0775 | CCGATCGCCACTG | - | containing protein |

| | | | | | | | | | | |
|---|---|---|---|---|---|---|---|---|---|---|
| AVAEIV_RS07445 | 1548846 | 1548858 | 322 | + | 10.9429 | 7.01E-05 | 0.0775 | CTGATCGGCATCG | - | efflux RND transporter permease subunit |
| AVAEIV_RS07450 | 1551929 | 1551941 | 349 | + | 10.9429 | 7.01E-05 | 0.0775 | CTGATCGGCATCG | - | efflux transporter outer membrane subunit |
| AVAEIV_RS10880 | 2288727 | 2288739 | 361 | + | 10.9429 | 7.01E-05 | 0.0794 | TCCATCGCCACCA | - | IS481-like element ISAzvi11 family transposase |
| AVAEIV_RS10885 | 2288727 | 2288739 | 75 | + | 10.9429 | 7.01E-05 | 0.0775 | TCCATCGCCACCA | - | hypothetical protein |
| AVAEIV_RS11450 | 2424631 | 2424643 | 192 | - | 10.9429 | 7.01E-05 | 0.0794 | CAGTTCGCCGCCC | - | GntP family permease |
| AVAEIV_RS12100 | 2566704 | 2566716 | 206 | - | 10.9429 | 7.01E-05 | 0.0775 | CCGATGGCCTGCA | - | ribonuclease E inhibitor RraB |
| AVAEIV_RS13195 | 2862056 | 2862068 | 54 | - | 10.9429 | 7.01E-05 | 0.0794 | CAGTGCGCTACCA | - | hypothetical protein |
| AVAEIV_RS13200 | 2862056 | 2862068 | 94 | - | 10.9429 | 7.01E-05 | 0.0775 | CAGTGCGCTACCA | - | MbcA/ParS/Xre antitoxin family protein |
| AVAEIV_RS17105 | 3710792 | 3710804 | 184 | + | 10.9429 | 7.01E-05 | 0.0794 | CTGATCGGCATCG | - | rhomboid family intramembrane serine protease |
| AVAEIV_RS21000 | 4467158 | 4467170 | 214 | - | 10.9429 | 7.01E-05 | 0.0794 | CGAGGCGCCATCG | - | SulP family inorganic anion transporter |
| AVAEIV_RS21005 | 4467158 | 4467170 | 311 | - | 10.9429 | 7.01E-05 | 0.0775 | CGAGGCGCCATCG | - | GNAT family N-acetyltransferase |
| AVAEIV_RS22360 | 4776018 | 4776030 | 387 | - | 10.9429 | 7.01E-05 | 0.0794 | CTGAACGGCATCA | *waaF* | lipopolysaccharide heptosyltransferase II |
| AVAEIV_RS23165 | 4947973 | 4947985 | -27 | - | 10.9429 | 7.01E-05 | 0.0775 | CTGCTCGCCGTCG | - | DNA-binding protein |
| AVAEIV_RS23700 | 5050143 | 5050155 | 228 | - | 10.9429 | 7.01E-05 | 0.0794 | CAGATGGCCATGC | - | CinA family protein |
| AVAEIV_RS24865 | 5290691 | 5290703 | 327 | + | 10.9429 | 7.01E-05 | 0.0794 | CCGATGGCCTGCA | - | subunit alpha |
| AVAEIV_RS00500 | 102865 | 102877 | 195 | + | 10.9286 | 7.09E-05 | 0.0794 | CGGCTTGCCGTCG | *ctaD* | cytochrome c oxidase subunit I |
| AVAEIV_RS04245 | 864673 | 864685 | 258 | + | 10.9286 | 7.09E-05 | 0.0775 | CCGTTCGCCAGGG | - | LysR family transcriptional regulator |
| AVAEIV_RS04580 | 938189 | 938201 | 328 | + | 10.9286 | 7.09E-05 | 0.0794 | CAGCAGGCCGGCG | - | urease subunit beta |
| AVAEIV_RS04815 | 978063 | 978075 | 134 | - | 10.9286 | 7.09E-05 | 0.0775 | CAGCAGGCCGGCG | *madM* | malonate transporter subunit MadM |
| AVAEIV_RS05070 | 1043354 | 1043366 | 37 | + | 10.9286 | 7.09E-05 | 0.0794 | CAGACCGACAGCA | - | alginate biosynthesis protein Alg44 |
| AVAEIV_RS06390 | 1303908 | 1303920 | 226 | - | 10.9286 | 7.09E-05 | 0.0794 | CAGCAGGCCGGCG | - | UDP-glucose dehydrogenase family protein |
| AVAEIV_RS06475 | 1336143 | 1336155 | 382 | + | 10.9286 | 7.09E-05 | 0.0794 | CGGCTTGCCGTCG | - | sulfite reductase subunit alpha |
| AVAEIV_RS06520 | 1345343 | 1345355 | 95 | - | 10.9286 | 7.09E-05 | 0.0775 | CGGCTTGCCGTCG | *pxpB* | 5-oxoprolinase subunit PxpB |
| AVAEIV_RS09100 | 1897211 | 1897223 | 181 | - | 10.9286 | 7.09E-05 | 0.0794 | CGCGAGGCCATCG | - | FecR family protein |
| AVAEIV_RS09630 | 2012060 | 2012072 | 206 | + | 10.9286 | 7.09E-05 | 0.0794 | CAGCAGGCCGGCG | - | DUF4381 domain-containing protein |
| AVAEIV_RS10355 | 2164557 | 2164569 | 269 | - | 10.9286 | 7.09E-05 | 0.0775 | CAGCAGGCCGGCG | - | DUF2934 domain-containing protein |
| AVAEIV_RS10685 | 2235040 | 2235052 | 226 | - | 10.9286 | 7.09E-05 | 0.0794 | CAGGTTGCCACCC | - | LysR family transcriptional regulator |
| AVAEIV_RS14095 | 3065839 | 3065851 | 313 | - | 10.9286 | 7.09E-05 | 0.0794 | CGCGAGGCCATCG | *edd* | phosphogluconate dehydratase |
| AVAEIV_RS15615 | 3421470 | 3421482 | -33 | + | 10.9286 | 7.09E-05 | 0.0794 | CAGCAGGCCGGCG | *ccmI* | c-type cytochrome biogenesis protein CcmI |
| AVAEIV_RS18605 | 4007128 | 4007140 | 13 | + | 10.9286 | 7.09E-05 | 0.0794 | CGATGGGCCAGCG | - | carboxymuconolactone decarboxylase family protein |
| AVAEIV_RS19105 | 4120009 | 4120021 | 58 | - | 10.9286 | 7.09E-05 | 0.0794 | CAAATCGCCCTCA | - | ATP-binding protein |
| AVAEIV_RS20155 | 4305043 | 4305055 | 117 | + | 10.9286 | 7.09E-05 | 0.0794 | CGGCTTGCCGTCG | *hisS* | histidine--tRNA ligase |
| AVAEIV_RS20700 | 4399803 | 4399815 | 361 | - | 10.9286 | 7.09E-05 | 0.0775 | TGGCTGGCCATCA | - | MerR family transcriptional regulator |
| AVAEIV_RS21725 | 4629362 | 4629374 | -44 | + | 10.9286 | 7.09E-05 | 0.0794 | CTCAGGGCCAGCG | - | ABC transporter substrate-binding protein |
| AVAEIV_RS21725 | 4629626 | 4629638 | 220 | + | 10.9286 | 7.09E-05 | 0.0794 | CAGCAGGCCGGCG | - | ABC transporter substrate-binding protein |
| AVAEIV_RS25145 | 5354536 | 5354548 | 173 | - | 10.9286 | 7.09E-05 | 0.0775 | CAGCAGGCCGGCG | - | HupE/UreJ family protein |
| AVAEIV_RS25190 | 5364483 | 5364495 | 169 | - | 10.9286 | 7.09E-05 | 0.0794 | CGCCATGCCAGCG | - | nickel-dependent hydrogenase large subunit |
| AVAEIV_RS25425 | 5415978 | 5415990 | 85 | + | 10.9286 | 7.09E-05 | 0.0794 | CAGCAGGCCGGCG | - | RnfABCDGE type electron transport complex subunit B |
| AVAEIV_RS00725 | 146264 | 146276 | 318 | + | 10.9143 | 7.16E-05 | 0.0775 | CGCAGGGCCATCC | - | CCE_0567 family metalloprotein |
| AVAEIV_RS00740 | 147600 | 147612 | -28 | + | 10.9143 | 7.16E-05 | 0.0775 | CTGTTCGCCGGCG | - | hypothetical protein |
| AVAEIV_RS02730 | 556648 | 556660 | 221 | - | 10.9143 | 7.16E-05 | 0.0775 | CGCCGCGCCACCG | - | phosphoenolpyruvate carboxykinase |
| AVAEIV_RS02730 | 556580 | 556592 | 289 | + | 10.9143 | 7.16E-05 | 0.0775 | CTGTTCGCCGGCG | - | phosphoenolpyruvate carboxykinase |
| AVAEIV_RS03380 | 679527 | 679539 | 294 | + | 10.9143 | 7.16E-05 | 0.0775 | CAGGACGCCGGCG | - | cobalamin-binding protein |
| AVAEIV_RS03995 | 807796 | 807808 | -36 | + | 10.9143 | 7.16E-05 | 0.0794 | CAGGACGCCGGCG | - | ABC transporter substrate-binding protein |
| AVAEIV_RS05300 | 1094390 | 1094402 | 133 | + | 10.9143 | 7.16E-05 | 0.0794 | CTGTTCGCCGGCG | - | HAD-IA family hydrolase |
| AVAEIV_RS05335 | 1099205 | 1099217 | 320 | + | 10.9143 | 7.16E-05 | 0.0775 | CTGTACGCCGGCA | *sbcD* | exonuclease subunit SbcD |
| AVAEIV_RS05550 | 1149757 | 1149769 | 35 | + | 10.9143 | 7.16E-05 | 0.0794 | CCAGGTGCCATCA | *mobH* | MobH family relaxase |
| AVAEIV_RS05555 | 1149757 | 1149769 | 158 | + | 10.9143 | 7.16E-05 | 0.0775 | CCAGGTGCCATCA | - | hypothetical protein |
| AVAEIV_RS05640 | 1160556 | 1160568 | 151 | - | 10.9143 | 7.16E-05 | 0.0794 | CTGTACGCCGGCA | - | containing protein |
| AVAEIV_RS07900 | 1641676 | 1641688 | 203 | + | 10.9143 | 7.16E-05 | 0.0775 | CGCCGCGCCACCG | - | sulfite exporter TauE/SafE family protein |
| AVAEIV_RS09865 | 2065352 | 2065364 | 60 | + | 10.9143 | 7.16E-05 | 0.0794 | TAGAGGGCCATCG | - | RnfABCDGE type electron transport complex subunit D |
| AVAEIV_RS10190 | 2127021 | 2127033 | 283 | - | 10.9143 | 7.16E-05 | 0.0794 | CTACTCGCCACCG | *hemN* | oxygen-independent coproporphyrinogen III oxidase |

| | | | | | | | | | | |
|---|---|---|---|---|---|---|---|---|---|---|
| AVAEIV_RS12390 | 2625027 | 2625039 | -39 | + | 10.9143 | 7.16E-05 | 0.0794 | CGCCGCGCCACCG | - | ABC transporter substrate-binding protein |
| AVAEIV_RS13410 | 2899560 | 2899572 | 257 | - | 10.9143 | 7.16E-05 | 0.0794 | GGCTTCGCCAGCA | - | putative quinol monooxygenase |
| AVAEIV_RS14620 | 3182023 | 3182035 | 382 | + | 10.9143 | 7.16E-05 | 0.0775 | CAAATCGCCACTG | - | NUDIX hydrolase |
| AVAEIV_RS17980 | 3887497 | 3887509 | 151 | + | 10.9143 | 7.16E-05 | 0.0775 | CTGTACGCCGGCA | - | containing protein |
| AVAEIV_RS18055 | 3902097 | 3902109 | 158 | - | 10.9143 | 7.16E-05 | 0.0794 | CCAGGTGCCATCA | - | hypothetical protein |
| AVAEIV_RS18060 | 3902097 | 3902109 | 35 | - | 10.9143 | 7.16E-05 | 0.0775 | CCAGGTGCCATCA | *mobH* | MobH family relaxase |
| AVAEIV_RS23445 | 4997490 | 4997502 | 76 | - | 10.9143 | 7.16E-05 | 0.0794 | CCCGTCGCCGGCA | - | hypothetical protein |
| AVAEIV_RS23450 | 4997490 | 4997502 | 48 | - | 10.9143 | 7.16E-05 | 0.0775 | CCCGTCGCCGGCA | *ompR* | two-component system response regulator OmpR |
| AVAEIV_RS23685 | 5046451 | 5046463 | 153 | - | 10.9143 | 7.16E-05 | 0.0775 | CGCCGCGCCACCG | - | TPM domain-containing protein |
| AVAEIV_RS24100 | 5126447 | 5126459 | 152 | - | 10.9143 | 7.16E-05 | 0.0775 | CTGTTCGCCGGCG | *rep* | DNA helicase Rep |
| AVAEIV_RS24605 | 5228051 | 5228063 | 210 | + | 10.9143 | 7.16E-05 | 0.0794 | CACTGGGCCAGCC | - | acyl-CoA desaturase |
| AVAEIV_RS25400 | 5411469 | 5411481 | 137 | - | 10.9143 | 7.16E-05 | 0.0794 | CTGCTGGCCATCC | - | RnfH family protein |
| AVAEIV_RS00685 | 140550 | 140562 | -42 | - | 10.9 | 7.23E-05 | 0.0775 | CAGTTGGCCTTCG | *nifT* | putative nitrogen fixation protein NifT |
| AVAEIV_RS00690 | 140550 | 140562 | 181 | - | 10.9 | 7.23E-05 | 0.0775 | CAGTTGGCCTTCG | - | protein |
| AVAEIV_RS02085 | 416719 | 416731 | 385 | + | 10.9 | 7.23E-05 | 0.0794 | CGGCACGCCATGA | - | IS110-like element ISAzvi2 family transposase |
| AVAEIV_RS02120 | 420927 | 420939 | 385 | - | 10.9 | 7.23E-05 | 0.0775 | CGGCACGCCATGA | - | IS110-like element ISAzvi2 family transposase |
| AVAEIV_RS03465 | 699171 | 699183 | 78 | - | 10.9 | 7.23E-05 | 0.0775 | TGGTTGGCCAGCA | - | NUDIX hydrolase |
| AVAEIV_RS03585 | 723646 | 723658 | 320 | - | 10.9 | 7.23E-05 | 0.0794 | CCGGCGGCCAGCA | - | asparaginase |
| AVAEIV_RS05960 | 1215659 | 1215671 | 52 | - | 10.9 | 7.23E-05 | 0.0794 | CGACTTGCCACCG | - | hypothetical protein |
| AVAEIV_RS06260 | 1277525 | 1277537 | 173 | + | 10.9 | 7.23E-05 | 0.0794 | CGCATGGCTAGCA | - | hypothetical protein |
| AVAEIV_RS06800 | 1403157 | 1403169 | 385 | + | 10.9 | 7.23E-05 | 0.0794 | CGGCACGCCATGA | - | IS110-like element ISAzvi2 family transposase |
| AVAEIV_RS07075 | 1460965 | 1460977 | 385 | + | 10.9 | 7.23E-05 | 0.0794 | CGGCACGCCATGA | - | IS110-like element ISAzvi2 family transposase |
| AVAEIV_RS07350 | 1522266 | 1522278 | 179 | - | 10.9 | 7.23E-05 | 0.0775 | CAATACGCCAGGA | *yegS* | lipid kinase YegS |
| AVAEIV_RS08465 | 1769168 | 1769180 | 306 | + | 10.9 | 7.23E-05 | 0.0775 | CCGTTCGCCAGTG | - | heavy metal translocating P-type ATPase |
| AVAEIV_RS08630 | 1800259 | 1800271 | 251 | - | 10.9 | 7.23E-05 | 0.0775 | CCGGCGGCCAGCA | - | LysR substrate-binding domain-containing protein |
| AVAEIV_RS09415 | 1963725 | 1963737 | 385 | + | 10.9 | 7.23E-05 | 0.0794 | CGGCACGCCATGA | - | IS110-like element ISAzvi2 family transposase |
| AVAEIV_RS09620 | 2010408 | 2010420 | 39 | + | 10.9 | 7.23E-05 | 0.0794 | CTGCGGGCCACCA | - | VWA domain-containing protein |
| AVAEIV_RS09880 | 2068795 | 2068807 | 217 | + | 10.9 | 7.23E-05 | 0.0794 | CCGGCGGCCAGCA | *rsxA* | electron transport complex subunit RsxA |
| AVAEIV_RS10110 | 2111965 | 2111977 | 70 | - | 10.9 | 7.23E-05 | 0.0794 | GAGGTGGCCAGCA | *phnE* | phosphonate ABC transporter, permease protein PhnE |
| AVAEIV_RS10150 | 2119081 | 2119093 | 292 | - | 10.9 | 7.23E-05 | 0.0775 | GTGATCGCCATCG | - | YbaB/EbfC family nucleoid-associated protein |
| AVAEIV_RS10485 | 2192629 | 2192641 | 385 | + | 10.9 | 7.23E-05 | 0.0794 | CGGCACGCCATGA | - | IS110-like element ISAzvi2 family transposase |
| AVAEIV_RS11595 | 2452861 | 2452873 | 385 | + | 10.9 | 7.23E-05 | 0.0794 | CGGCACGCCATGA | - | IS110-like element ISAzvi2 family transposase |
| AVAEIV_RS12140 | 2576572 | 2576584 | 385 | + | 10.9 | 7.23E-05 | 0.0794 | CGGCACGCCATGA | - | IS110-like element ISAzvi2 family transposase |
| AVAEIV_RS13385 | 2895384 | 2895396 | 257 | - | 10.9 | 7.23E-05 | 0.0775 | GAGGTGGCCAGCA | - | integrase core domain-containing protein |
| AVAEIV_RS13645 | 2951195 | 2951207 | 385 | - | 10.9 | 7.23E-05 | 0.0775 | CGGCACGCCATGA | - | IS110-like element ISAzvi2 family transposase |
| AVAEIV_RS14655 | 3188530 | 3188542 | 385 | - | 10.9 | 7.23E-05 | 0.0775 | CGGCACGCCATGA | - | IS110-like element ISAzvi2 family transposase |
| AVAEIV_RS14680 | 3195327 | 3195339 | 385 | + | 10.9 | 7.23E-05 | 0.0794 | CGGCACGCCATGA | - | IS110-like element ISAzvi2 family transposase |
| AVAEIV_RS15210 | 3312965 | 3312977 | 276 | - | 10.9 | 7.23E-05 | 0.0794 | CCGGCGGCCAGCA | - | ABC transporter permease |
| AVAEIV_RS15555 | 3401971 | 3401983 | 385 | - | 10.9 | 7.23E-05 | 0.0775 | CGGCACGCCATGA | - | IS110-like element ISAzvi2 family transposase |
| AVAEIV_RS16390 | 3560310 | 3560322 | 171 | + | 10.9 | 7.23E-05 | 0.0794 | CCGGCGGCCAGCA | - | holin family protein |
| AVAEIV_RS16430 | 3568876 | 3568888 | 329 | - | 10.9 | 7.23E-05 | 0.0794 | CGTTGCGCCATCG | - | NAD(P)-dependent alcohol dehydrogenase |
| AVAEIV_RS17930 | 3878876 | 3878888 | 52 | + | 10.9 | 7.23E-05 | 0.0775 | CGACTTGCCACCG | - | hypothetical protein |
| AVAEIV_RS17940 | 3882787 | 3882799 | 385 | + | 10.9 | 7.23E-05 | 0.0794 | CGGCACGCCATGA | - | IS110-like element ISAzvi2 family transposase |
| AVAEIV_RS19175 | 4132885 | 4132897 | 121 | - | 10.9 | 7.23E-05 | 0.0794 | CCCAACGCCGGCG | - | hypothetical protein |
| AVAEIV_RS19695 | 4198787 | 4198799 | 224 | + | 10.9 | 7.23E-05 | 0.0794 | CAGGGCGCCATGA | - | extensin family protein |
| AVAEIV_RS19760 | 4211563 | 4211575 | 384 | - | 10.9 | 7.23E-05 | 0.0794 | CCGGCGGCCAGCA | *lysS* | lysine--tRNA ligase |
| AVAEIV_RS22260 | 4752329 | 4752341 | 385 | + | 10.9 | 7.23E-05 | 0.0794 | CGGCACGCCATGA | - | IS110-like element ISAzvi2 family transposase |
| AVAEIV_RS22390 | 4787647 | 4787659 | 110 | - | 10.9 | 7.23E-05 | 0.0775 | CCGGCGGCCAGCA | - | universal stress protein |
| AVAEIV_RS22685 | 4859797 | 4859809 | 172 | - | 10.9 | 7.23E-05 | 0.0794 | CGGTATGCCGGCA | *pip* | prolyl aminopeptidase |
| AVAEIV_RS22690 | 4859797 | 4859809 | 79 | - | 10.9 | 7.23E-05 | 0.0775 | CGGTATGCCGGCA | - | DUF2238 domain-containing protein |
| AVAEIV_RS22795 | 4881708 | 4881720 | 385 | + | 10.9 | 7.23E-05 | 0.0794 | CGGCACGCCATGA | - | IS110-like element ISAzvi2 family transposase |
| AVAEIV_RS23165 | 4947889 | 4947901 | 57 | - | 10.9 | 7.23E-05 | 0.0775 | CCGGCGGCCAGCA | - | DNA-binding protein |

| | | | | | | | | | | |
|---|---|---|---|---|---|---|---|---|---|---|
| AVAEIV_RS24130 | 5131652 | 5131664 | -47 | - | 10.9 | 7.23E-05 | 0.0775 | CCGGCGGCCAGCA | - | catalase family peroxidase |
| AVAEIV_RS24420 | 5191753 | 5191765 | 349 | - | 10.9 | 7.23E-05 | 0.0794 | CCGGCGGCCAGCA | *phoU* | phosphate signaling complex protein PhoU |
| AVAEIV_RS24845 | 5282959 | 5282971 | 311 | - | 10.9 | 7.23E-05 | 0.0775 | CCGGCGGCCAGCA | - | tripartite tricarboxylate transporter permease |
| AVAEIV_RS01340 | 261758 | 261770 | 246 | - | 10.8857 | 7.30E-05 | 0.0795 | CTGATCGCCACCT | *nifN* | protein NifN |
| AVAEIV_RS02830 | 580572 | 580584 | 384 | - | 10.8857 | 7.30E-05 | 0.0795 | CGCATCGCGAGCA | - | cytochrome B6 |
| AVAEIV_RS03960 | 798635 | 798647 | 295 | + | 10.8857 | 7.30E-05 | 0.0795 | CGGATGGCCTGCA | - | energy transducer TonB |
| AVAEIV_RS07680 | 1602362 | 1602374 | 123 | - | 10.8857 | 7.30E-05 | 0.0775 | CTGACCGCCACCG | - | S49 family peptidase |
| AVAEIV_RS08070 | 1672884 | 1672896 | 282 | - | 10.8857 | 7.30E-05 | 0.0775 | GCCATGGCCAGCA | *scpB* | SMC-Scp complex subunit ScpB |
| AVAEIV_RS12395 | 2626386 | 2626398 | 386 | - | 10.8857 | 7.30E-05 | 0.0795 | GCCATGGCCAGCA | - | hypothetical protein |
| AVAEIV_RS13145 | 2851918 | 2851930 | 215 | + | 10.8857 | 7.30E-05 | 0.0795 | GCCATGGCCAGCA | *hutG* | N-formylglutamate deformylase |
| AVAEIV_RS15220 | 3315593 | 3315605 | 153 | + | 10.8857 | 7.30E-05 | 0.0795 | CTGATCGCCACCT | - | protein |
| AVAEIV_RS15225 | 3316819 | 3316831 | 153 | + | 10.8857 | 7.30E-05 | 0.0795 | CTGATCGCCACCT | - | 3-oxoacyl-[acyl-carrier-protein] synthase III C-terminal domain-containing protein |
| AVAEIV_RS15245 | 3329943 | 3329955 | 186 | - | 10.8857 | 7.30E-05 | 0.0795 | CCGCGCGCCAGCC | *dnaQ* | DNA polymerase III subunit epsilon |
| AVAEIV_RS16680 | 3612876 | 3612888 | 225 | + | 10.8857 | 7.30E-05 | 0.0795 | CGCATCGCGAGCA | *cobC* | alpha-ribazole phosphatase family protein |
| AVAEIV_RS18120 | 3914167 | 3914179 | 77 | - | 10.8857 | 7.30E-05 | 0.0795 | CAGGAGGCCAGCC | *ruvB* | Holliday junction branch migration DNA helicase RuvB |
| AVAEIV_RS21775 | 4637457 | 4637469 | 317 | - | 10.8857 | 7.30E-05 | 0.0795 | CGGTGCGCCGCCA | - | ABC transporter permease |
| AVAEIV_RS21955 | 4683040 | 4683052 | -32 | - | 10.8857 | 7.30E-05 | 0.0795 | CGCATCGCGAGCA | - | DUF5064 family protein |
| AVAEIV_RS21960 | 4683040 | 4683052 | 261 | - | 10.8857 | 7.30E-05 | 0.0775 | CGCATCGCGAGCA | - | DUF3141 domain-containing protein |
| AVAEIV_RS25190 | 5364660 | 5364672 | 346 | + | 10.8857 | 7.30E-05 | 0.0795 | CCGCGCGCCAGCC | - | nickel-dependent hydrogenase large subunit |
| AVAEIV_RS00730 | 146925 | 146937 | 150 | - | 10.8714 | 7.37E-05 | 0.0775 | CTGCCCGCCATCG | - | nitrogen fixation (4Fe-4S) ferredoxin-like protein |
| AVAEIV_RS00735 | 146925 | 146937 | 318 | - | 10.8714 | 7.37E-05 | 0.0775 | CTGCCCGCCATCG | - | 2Fe-2S ferredoxin |
| AVAEIV_RS00875 | 172027 | 172039 | 286 | - | 10.8714 | 7.37E-05 | 0.0797 | CTGCCCGCCATCG | - | DUF4105 domain-containing protein |
| AVAEIV_RS00880 | 172027 | 172039 | -25 | - | 10.8714 | 7.37E-05 | 0.0797 | CTGCCCGCCATCG | - | DUF2388 domain-containing protein |
| AVAEIV_RS01490 | 287341 | 287353 | 127 | - | 10.8714 | 7.37E-05 | 0.0797 | CCGAACGGCACCA | - | YggT family protein |
| AVAEIV_RS03230 | 652217 | 652229 | 78 | + | 10.8714 | 7.37E-05 | 0.0775 | AAGATTGCCAGCA | *rpsD* | 30S ribosomal protein S4 |
| AVAEIV_RS04185 | 853156 | 853168 | 307 | - | 10.8714 | 7.37E-05 | 0.0775 | CAGATCGGCGTCG | - | dehydrogenase |
| AVAEIV_RS06005 | 1224843 | 1224855 | 379 | - | 10.8714 | 7.37E-05 | 0.0797 | CGATTCGCCACCC | - | integrating conjugative element protein |
| AVAEIV_RS06995 | 1443596 | 1443608 | 348 | + | 10.8714 | 7.37E-05 | 0.0797 | CAGACCGCCATTG | - | ATP-binding protein |
| AVAEIV_RS07440 | 1545930 | 1545942 | 128 | - | 10.8714 | 7.37E-05 | 0.0775 | CCGCTCGCCGCCG | - | efflux RND transporter permease subunit |
| AVAEIV_RS10950 | 2303163 | 2303175 | 268 | + | 10.8714 | 7.37E-05 | 0.0797 | CACCACGCCGGCG | - | amino acid adenylation domain-containing protein |
| AVAEIV_RS12725 | 2702349 | 2702361 | 158 | + | 10.8714 | 7.37E-05 | 0.0797 | CAGATGGCTATCC | - | DUF3325 domain-containing protein |
| AVAEIV_RS14970 | 3263408 | 3263420 | 382 | - | 10.8714 | 7.37E-05 | 0.0797 | CAGCTCGCTACCG | - | SDR family oxidoreductase |
| AVAEIV_RS16065 | 3500431 | 3500443 | 33 | + | 10.8714 | 7.37E-05 | 0.0775 | CCGAGGGCTATCA | - | aldolase/oxaloacetate decarboxylase |
| AVAEIV_RS17260 | 3740734 | 3740746 | 148 | + | 10.8714 | 7.37E-05 | 0.0797 | CCGCTCGCCGCCG | - | fused response regulator/phosphatase |
| AVAEIV_RS17445 | 3777698 | 3777710 | 274 | - | 10.8714 | 7.37E-05 | 0.0775 | CAGATCGGCGTCG | - | metal-dependent hydrolase |
| AVAEIV_RS17550 | 3801817 | 3801829 | 133 | + | 10.8714 | 7.37E-05 | 0.0775 | CCGCACGCCGCCA | - | osmoprotectant NAGGN system M42 family peptidase |
| AVAEIV_RS17890 | 3872331 | 3872343 | 255 | + | 10.8714 | 7.37E-05 | 0.0775 | CGATTCGCCACCC | - | integrating conjugative element protein |
| AVAEIV_RS21665 | 4612300 | 4612312 | 282 | + | 10.8714 | 7.37E-05 | 0.0775 | CAGATCGGCGTCG | - | malic enzyme-like NAD(P)-binding protein |
| AVAEIV_RS23705 | 5051131 | 5051143 | 183 | + | 10.8714 | 7.37E-05 | 0.0797 | CTCAGCGCCGTCA | - | manganese catalase family protein |
| AVAEIV_RS23890 | 5089634 | 5089646 | 333 | + | 10.8714 | 7.37E-05 | 0.0797 | CTGATTGCCGCCA | *trxA* | thioredoxin TrxA |
| AVAEIV_RS23895 | 5089634 | 5089646 | -43 | + | 10.8714 | 7.37E-05 | 0.0775 | CTGATTGCCGCCA | *ppx* | exopolyphosphatase |
| AVAEIV_RS24375 | 5181802 | 5181814 | 190 | - | 10.8714 | 7.37E-05 | 0.0775 | CAGATCGGCGTCG | - | containing protein |
| AVAEIV_RS25295 | 5388152 | 5388164 | 363 | - | 10.8714 | 7.37E-05 | 0.0797 | CCGCTCGCCGCCG | - | tungsten ABC transporter permease |
| AVAEIV_RS02260 | 453826 | 453838 | 65 | + | 10.8571 | 7.46E-05 | 0.0803 | CAGATGGCCGCCC | *cooS* | subunit |
| AVAEIV_RS07175 | 1479249 | 1479261 | 247 | - | 10.8571 | 7.46E-05 | 0.0775 | CAGATCGCGATCC | *pdxJ* | pyridoxine 5'-phosphate synthase |
| AVAEIV_RS07280 | 1509368 | 1509380 | 122 | - | 10.8571 | 7.46E-05 | 0.0803 | CTCCTCGCCATCC | *bfr* | bacterioferritin |
| AVAEIV_RS07925 | 1647446 | 1647458 | 154 | + | 10.8571 | 7.46E-05 | 0.0775 | CGCGTCGCCGGCA | - | thiopurine S-methyltransferase |
| AVAEIV_RS09990 | 2091163 | 2091175 | 239 | + | 10.8571 | 7.46E-05 | 0.0775 | CAGATCGCGATCC | - | glycine zipper domain-containing protein |
| AVAEIV_RS12315 | 2605605 | 2605617 | 242 | - | 10.8571 | 7.46E-05 | 0.0775 | CCGCTCGGCATCG | - | hypothetical protein |
| AVAEIV_RS12930 | 2803891 | 2803903 | 133 | - | 10.8571 | 7.46E-05 | 0.0803 | CTCGTGGCCATCG | - | dienelactone hydrolase family protein |
| AVAEIV_RS17685 | 3827799 | 3827811 | 296 | + | 10.8571 | 7.46E-05 | 0.0775 | CGAACTGCCATCG | - | hybrid sensor histidine kinase/response regulator |

| | | | | | | | | | | |
|---|---|---|---|---|---|---|---|---|---|---|
| AVAEIV_RS20385 | 4335811 | 4335823 | 207 | + | 10.8571 | 7.46E-05 | 0.0803 | CTCCTCGCCATCC | *ogt* | methyltransferase |
| AVAEIV_RS23040 | 4931610 | 4931622 | 189 | + | 10.8571 | 7.46E-05 | 0.0803 | CGTGTCGCCACCA | - | terminase large subunit domain-containing protein |
| AVAEIV_RS23045 | 4931610 | 4931622 | -43 | + | 10.8571 | 7.46E-05 | 0.0775 | CGTGTCGCCACCA | - | GPO family capsid scaffolding protein |
| AVAEIV_RS25420 | 5415598 | 5415610 | 250 | - | 10.8571 | 7.46E-05 | 0.0803 | CAGATGGCCGCCC | *rsxC* | electron transport complex subunit RsxC |
| AVAEIV_RS00490 | 100539 | 100551 | 58 | - | 10.8429 | 7.53E-05 | 0.0804 | CTGTTCGACATCA | - | cytochrome c oxidase subunit 3 |
| AVAEIV_RS00590 | 120517 | 120529 | 161 | - | 10.8429 | 7.53E-05 | 0.0775 | CCGATGGCCAGGG | - | 5-(carboxyamino)imidazole ribonucleotide synthase |
| AVAEIV_RS00645 | 131269 | 131281 | 49 | - | 10.8429 | 7.53E-05 | 0.0804 | CAGCTCGGCACCG | - | MFS transporter |
| AVAEIV_RS01985 | 397349 | 397361 | -34 | - | 10.8429 | 7.53E-05 | 0.0804 | CATGCCGCCATCA | - | VOC family protein |
| AVAEIV_RS02735 | 558474 | 558486 | 54 | + | 10.8429 | 7.53E-05 | 0.0775 | CTGTTCGCCAGCT | *cls* | cardiolipin synthase |
| AVAEIV_RS03100 | 639040 | 639052 | 306 | - | 10.8429 | 7.53E-05 | 0.0775 | CAGCACGGCACCA | *tuf* | elongation factor Tu |
| AVAEIV_RS03425 | 691610 | 691622 | 286 | + | 10.8429 | 7.53E-05 | 0.0804 | CAACTCGCCGCCG | *dusB* | tRNA dihydrouridine synthase DusB |
| AVAEIV_RS04150 | 843509 | 843521 | 102 | + | 10.8429 | 7.53E-05 | 0.0775 | CCCGCCGCCAGCA | - | efflux RND transporter periplasmic adaptor subunit |
| AVAEIV_RS04460 | 909262 | 909274 | 284 | - | 10.8429 | 7.53E-05 | 0.0804 | CCCGCCGCCAGCA | - | kinase/phosphomethylpyrimidine kinase |
| AVAEIV_RS05175 | 1063366 | 1063378 | 196 | + | 10.8429 | 7.53E-05 | 0.0775 | CTGATCGCCACGA | - | hypothetical protein |
| AVAEIV_RS05920 | 1211680 | 1211692 | 192 | - | 10.8429 | 7.53E-05 | 0.0804 | CGCAACGCCGGCG | - | protein |
| AVAEIV_RS09340 | 1946385 | 1946397 | 329 | + | 10.8429 | 7.53E-05 | 0.0775 | CCCGCCGCCAGCA | - | acyltransferase family protein |
| AVAEIV_RS10075 | 2106273 | 2106285 | 203 | - | 10.8429 | 7.53E-05 | 0.0775 | CAACTCGCCGCCG | - | AtpZ/AtpI family protein |
| AVAEIV_RS15070 | 3280989 | 3281001 | -11 | + | 10.8429 | 7.53E-05 | 0.0804 | CTGATCGCCACGA | - | response regulator |
| AVAEIV_RS15375 | 3358946 | 3358958 | 38 | + | 10.8429 | 7.53E-05 | 0.0775 | CTGTCCGCCAGCG | - | exonuclease domain-containing protein |
| AVAEIV_RS15515 | 3393498 | 3393510 | 65 | - | 10.8429 | 7.53E-05 | 0.0775 | CCGATGGCCAGGG | - | polysaccharide biosynthesis/export family protein |
| AVAEIV_RS19080 | 4110253 | 4110265 | 367 | + | 10.8429 | 7.53E-05 | 0.0775 | CACTTCGCCTTCG | *dapC* | succinyldiaminopimelate transaminase |
| AVAEIV_RS19510 | 4171390 | 4171402 | 209 | - | 10.8429 | 7.53E-05 | 0.0775 | CAGCGGGCTATCA | - | hypothetical protein |
| AVAEIV_RS21990 | 4693066 | 4693078 | 14 | + | 10.8429 | 7.53E-05 | 0.0775 | CGCAACGCCGGCG | - | paraquat-inducible protein A |
| AVAEIV_RS22130 | 4722501 | 4722513 | 316 | + | 10.8429 | 7.53E-05 | 0.0775 | CTGTCCGCCAGCG | - | containing protein |
| AVAEIV_RS22815 | 4883494 | 4883506 | 262 | + | 10.8429 | 7.53E-05 | 0.0775 | CTGTCCGCCAGCG | - | cache domain-containing protein |
| AVAEIV_RS23050 | 4932192 | 4932204 | 286 | + | 10.8429 | 7.53E-05 | 0.0775 | CAGCTCGGCACCG | - | phage major capsid protein, P2 family |
| AVAEIV_RS23545 | 5018378 | 5018390 | 98 | + | 10.8429 | 7.53E-05 | 0.0804 | CCCTCGGCCATCG | - | DnaJ domain-containing protein |
| AVAEIV_RS25020 | 5326149 | 5326161 | 200 | - | 10.8429 | 7.53E-05 | 0.0775 | CACTTCGCCTTCG | - | sugar ABC transporter substrate-binding protein |
| AVAEIV_RS00050 | 10592 | 10604 | 245 | - | 10.8286 | 7.61E-05 | 0.0804 | GACAACGCCAGCG | *gmhB* | phosphatase |
| AVAEIV_RS00285 | 62065 | 62077 | 139 | - | 10.8286 | 7.61E-05 | 0.0804 | CAGGATGCCGGCA | - | ABC transporter permease |
| AVAEIV_RS00590 | 120507 | 120519 | 171 | + | 10.8286 | 7.61E-05 | 0.0775 | CCGGTCGCCACCC | - | 5-(carboxyamino)imidazole ribonucleotide synthase |
| AVAEIV_RS00800 | 157062 | 157074 | 29 | - | 10.8286 | 7.61E-05 | 0.0775 | CACTCGGCCACCG | - | hypothetical protein |
| AVAEIV_RS05290 | 1093154 | 1093166 | 116 | - | 10.8286 | 7.61E-05 | 0.0804 | CCCGGCGCCAGCG | *yejK* | nucleoid-associated protein YejK |
| AVAEIV_RS05520 | 1141921 | 1141933 | 378 | + | 10.8286 | 7.61E-05 | 0.0775 | GGCATGGCCAGCA | *pgeF* | peptidoglycan editing factor PgeF |
| AVAEIV_RS05930 | 1213236 | 1213248 | 206 | + | 10.8286 | 7.61E-05 | 0.0804 | CAGGTTGCCGGCG | - | protein |
| AVAEIV_RS06695 | 1382223 | 1382235 | 217 | + | 10.8286 | 7.61E-05 | 0.0804 | CAGGTCGCCGTCC | - | KdsC family phosphatase |
| AVAEIV_RS07445 | 1549211 | 1549223 | -43 | + | 10.8286 | 7.61E-05 | 0.0775 | CCGGTCGCCACCC | - | efflux RND transporter permease subunit |
| AVAEIV_RS07805 | 1624181 | 1624193 | 76 | - | 10.8286 | 7.61E-05 | 0.0775 | CAGGTTGCCGGCG | *dmpG* | 4-hydroxy-2-oxovalerate aldolase |
| AVAEIV_RS08290 | 1725614 | 1725626 | -36 | - | 10.8286 | 7.61E-05 | 0.0775 | CATGGCGCCATCG | - | transposase |
| AVAEIV_RS09060 | 1884546 | 1884558 | 210 | + | 10.8286 | 7.61E-05 | 0.0775 | CGCATCGCCTGCA | - | enoyl-CoA hydratase/isomerase family protein |
| AVAEIV_RS09615 | 2008827 | 2008839 | 188 | - | 10.8286 | 7.61E-05 | 0.0804 | CGGCTCGCTATCG | - | BatD family protein |
| AVAEIV_RS09705 | 2033708 | 2033720 | 220 | - | 10.8286 | 7.61E-05 | 0.0804 | CTCTTCGCCAGCC | - | hypothetical protein |
| AVAEIV_RS12835 | 2752937 | 2752949 | 138 | - | 10.8286 | 7.61E-05 | 0.0804 | CAACTCGGCATCG | - | non-ribosomal peptide synthetase |
| AVAEIV_RS13775 | 2981525 | 2981537 | 380 | + | 10.8286 | 7.61E-05 | 0.0804 | CAGGTCGCCGTCC | - | DUF4150 domain-containing protein |
| AVAEIV_RS13925 | 3019432 | 3019444 | 45 | + | 10.8286 | 7.61E-05 | 0.0775 | CTGTTTGCCGGCA | - | sugar phosphate isomerase/epimerase family protein |
| AVAEIV_RS14150 | 3078863 | 3078875 | 63 | + | 10.8286 | 7.61E-05 | 0.0804 | CGAATCGCTATCG | - | ABC transporter substrate-binding protein |
| AVAEIV_RS14155 | 3078863 | 3078875 | 262 | + | 10.8286 | 7.61E-05 | 0.0775 | CGAATCGCTATCG | - | helix-turn-helix domain-containing protein |
| AVAEIV_RS14690 | 3195607 | 3195619 | 333 | - | 10.8286 | 7.61E-05 | 0.0775 | CATGGCGCCATCG | - | ATP-binding protein |
| AVAEIV_RS14710 | 3197843 | 3197855 | 383 | - | 10.8286 | 7.61E-05 | 0.0775 | CATGGCGCCATCG | - | secretin N-terminal domain-containing protein |
| AVAEIV_RS17175 | 3723168 | 3723180 | 148 | - | 10.8286 | 7.61E-05 | 0.0804 | CATGGCGCCATCG | *folC* | synthase |
| AVAEIV_RS17565 | 3807216 | 3807228 | 84 | + | 10.8286 | 7.61E-05 | 0.0804 | CGAAGCGCCAGCC | - | DUF2784 domain-containing protein |
| AVAEIV_RS17645 | 3821347 | 3821359 | 275 | - | 10.8286 | 7.61E-05 | 0.0775 | CCCGGCGCCAGCG | - | carboxy terminal-processing peptidase |

| | | | | | | | | | | |
|---|---|---|---|---|---|---|---|---|---|---|
| AVAEIV_RS18850 | 4058833 | 4058845 | 145 | - | 10.8286 | 7.61E-05 | 0.0775 | CAGATCGCTATCT | - | LOG family protein |
| AVAEIV_RS18925 | 4076350 | 4076362 | 116 | + | 10.8286 | 7.61E-05 | 0.0804 | CGGTCCGCCATCC | - | methyltransferase |
| AVAEIV_RS19505 | 4170695 | 4170707 | 285 | - | 10.8286 | 7.61E-05 | 0.0775 | CTGAAGGCCGGCA | - | hypothetical protein |
| AVAEIV_RS19765 | 4212369 | 4212381 | 50 | + | 10.8286 | 7.61E-05 | 0.0804 | CAGCGGGCCGCCA | *prfB* | peptide chain release factor 2 |
| AVAEIV_RS20130 | 4299596 | 4299608 | 317 | + | 10.8286 | 7.61E-05 | 0.0804 | CATGGCGCCATCG | - | FecR family protein |
| AVAEIV_RS21065 | 4479672 | 4479684 | 76 | + | 10.8286 | 7.61E-05 | 0.0804 | CAGGTTGCCGGCG | *dmpG* | 4-hydroxy-2-oxovalerate aldolase |
| AVAEIV_RS21980 | 4688240 | 4688252 | 349 | + | 10.8286 | 7.61E-05 | 0.0775 | CTGCTCGCCATGA | - | efflux RND transporter periplasmic adaptor subunit |
| AVAEIV_RS22705 | 4862709 | 4862721 | 329 | + | 10.8286 | 7.61E-05 | 0.0804 | CCATTCGCCGGCG | - | cupin domain-containing protein |
| AVAEIV_RS23305 | 4968117 | 4968129 | 83 | - | 10.8286 | 7.61E-05 | 0.0775 | CGCATCGCCTGCA | - | IS5 family transposase |
| AVAEIV_RS24475 | 5202513 | 5202525 | 22 | - | 10.8286 | 7.61E-05 | 0.0804 | CCGGGCGCCGTCA | - | RcnB family protein |
| AVAEIV_RS24480 | 5202513 | 5202525 | 290 | - | 10.8286 | 7.61E-05 | 0.0775 | CCGGGCGCCGTCA | - | SagB family peptide dehydrogenase |
| AVAEIV_RS24855 | 5286169 | 5286181 | -47 | - | 10.8286 | 7.61E-05 | 0.0775 | CAGCGGGCCGCCA | - | AbrB family transcriptional regulator |
| AVAEIV_RS25855 | 5544156 | 5544168 | 254 | - | 10.8286 | 7.61E-05 | 0.0775 | CAGCGGGCCGCCA | - | ABC transporter substrate-binding protein |
| AVAEIV_RS00100 | 21257 | 21269 | 191 | - | 10.8143 | 7.69E-05 | 0.0775 | CGGCTCGCCGCCG | *dprA* | DNA-processing protein DprA |
| AVAEIV_RS00495 | 101176 | 101188 | 126 | + | 10.8143 | 7.69E-05 | 0.0804 | CGGCTCGCCGCCG | - | cytochrome c oxidase assembly protein |
| AVAEIV_RS01190 | 232765 | 232777 | 90 | + | 10.8143 | 7.69E-05 | 0.0804 | CAGACCGCCGCCG | - | ATP-binding cassette domain-containing protein |
| AVAEIV_RS01635 | 316257 | 316269 | 13 | - | 10.8143 | 7.69E-05 | 0.0804 | GCGCACGCCATCA | - | ABC transporter ATP-binding protein |
| AVAEIV_RS06490 | 1338108 | 1338120 | 103 | + | 10.8143 | 7.69E-05 | 0.0804 | CGGCTCGCCGCCG | - | PepSY-associated TM helix domain-containing protein |
| AVAEIV_RS06950 | 1429944 | 1429956 | 341 | + | 10.8143 | 7.69E-05 | 0.0775 | CAGTTGGCCGGCC | *ftsA* | cell division protein FtsA |
| AVAEIV_RS07120 | 1467372 | 1467384 | 342 | - | 10.8143 | 7.69E-05 | 0.0775 | CCACGGGCCACCA | - | sigma-E factor negative regulatory protein |
| AVAEIV_RS10395 | 2172712 | 2172724 | 368 | - | 10.8143 | 7.69E-05 | 0.0775 | TAGAAGGCCAGCG | - | MFS transporter |
| AVAEIV_RS10705 | 2238811 | 2238823 | 190 | - | 10.8143 | 7.69E-05 | 0.0775 | CGGCACGCCGCCA | - | YodC family protein |
| AVAEIV_RS12090 | 2565341 | 2565353 | 109 | - | 10.8143 | 7.69E-05 | 0.0775 | CGGATGGCCGTCC | - | PLDc N-terminal domain-containing protein |
| AVAEIV_RS12255 | 2595724 | 2595736 | 137 | - | 10.8143 | 7.69E-05 | 0.0804 | CAGACCGCCGCCG | *fliM* | flagellar motor switch protein FliM |
| AVAEIV_RS12320 | 2606858 | 2606870 | 82 | - | 10.8143 | 7.69E-05 | 0.0775 | GCGCTCGCCATCG | - | EscU/YscU/HrcU family type III secretion system export apparatus switch protein |
| AVAEIV_RS12630 | 2679674 | 2679686 | 28 | - | 10.8143 | 7.69E-05 | 0.0804 | CCCAGCGCTATCA | - | hypothetical protein |
| AVAEIV_RS13265 | 2875730 | 2875742 | -18 | + | 10.8143 | 7.69E-05 | 0.0804 | CGGCTCGCCGCCG | - | HlyD family secretion protein |
| AVAEIV_RS13270 | 2875730 | 2875742 | 331 | + | 10.8143 | 7.69E-05 | 0.0775 | CGGCTCGCCGCCG | - | nucleotidyltransferase family protein |
| AVAEIV_RS13540 | 2931233 | 2931245 | 215 | + | 10.8143 | 7.69E-05 | 0.0804 | CGAATCGCCGCCG | - | NAD(P)-dependent alcohol dehydrogenase |
| AVAEIV_RS13940 | 3022197 | 3022209 | 374 | - | 10.8143 | 7.69E-05 | 0.0775 | TAGAAGGCCAGCG | - | 2-hydroxyacid dehydrogenase |
| AVAEIV_RS14635 | 3184595 | 3184607 | 154 | - | 10.8143 | 7.69E-05 | 0.0775 | CCAGTTGCCAGCG | *purB* | adenylosuccinate lyase |
| AVAEIV_RS15585 | 3412880 | 3412892 | -25 | - | 10.8143 | 7.69E-05 | 0.0804 | CAGACCGCCGCCG | *ligA* | NAD-dependent DNA ligase LigA |
| AVAEIV_RS15605 | 3418566 | 3418578 | 380 | + | 10.8143 | 7.69E-05 | 0.0775 | CGGCTCGCCGCCG | - | ion transporter |
| AVAEIV_RS19040 | 4102957 | 4102969 | 269 | + | 10.8143 | 7.69E-05 | 0.0804 | CAGTTGGCCGGCC | - | phosphatidate cytidylyltransferase |
| AVAEIV_RS19130 | 4123746 | 4123758 | 60 | - | 10.8143 | 7.69E-05 | 0.0804 | CAGACCGCCGCCG | - | glycoside hydrolase family 104 protein |
| AVAEIV_RS20015 | 4267371 | 4267383 | 142 | + | 10.8143 | 7.69E-05 | 0.0775 | CGGCACGCCGCCA | - | SDR family oxidoreductase |
| AVAEIV_RS20420 | 4341245 | 4341257 | 87 | + | 10.8143 | 7.69E-05 | 0.0804 | CCACGGGCCACCA | - | FKBP-type peptidyl-prolyl cis-trans isomerase |
| AVAEIV_RS22430 | 4796245 | 4796257 | 336 | - | 10.8143 | 7.69E-05 | 0.0804 | CGGATGGCCGTCC | - | RNA polymerase sigma factor |
| AVAEIV_RS22430 | 4796260 | 4796272 | 351 | - | 10.8143 | 7.69E-05 | 0.0804 | CGGATGGCCGTCC | - | RNA polymerase sigma factor |
| AVAEIV_RS22925 | 4911329 | 4911341 | 37 | - | 10.8143 | 7.69E-05 | 0.0775 | TCCATGGCCAGCA | - | anhydro-N-acetylmuramic acid kinase |
| AVAEIV_RS22935 | 4915008 | 4915020 | 259 | + | 10.8143 | 7.69E-05 | 0.0804 | CAGATCGCCGCCT | *erpA* | iron-sulfur cluster insertion protein ErpA |
| AVAEIV_RS24220 | 5149452 | 5149464 | 40 | + | 10.8143 | 7.69E-05 | 0.0804 | GCAATCGCCATCG | - | dihydrofolate reductase |
| AVAEIV_RS24225 | 5149452 | 5149464 | 77 | + | 10.8143 | 7.69E-05 | 0.0775 | GCAATCGCCATCG | - | glycerophosphodiester phosphodiesterase |
| AVAEIV_RS24705 | 5248998 | 5249010 | 237 | - | 10.8143 | 7.69E-05 | 0.0775 | CCGTGGGCCGGCA | - | hypothetical protein |
| AVAEIV_RS25425 | 5416229 | 5416241 | 336 | - | 10.8143 | 7.69E-05 | 0.0804 | CGCAAGGCCAGCC | - | RnfABCDGE type electron transport complex subunit B |
| AVAEIV_RS00135 | 28535 | 28547 | 263 | - | 10.8 | 7.76E-05 | 0.0775 | CGGCTCGGCATCG | - | copper resistance protein B |
| AVAEIV_RS00140 | 30626 | 30638 | 249 | + | 10.8 | 7.76E-05 | 0.0804 | CACATCGCCGCCC | - | MgtC/SapB family protein |
| AVAEIV_RS00705 | 142490 | 142502 | 263 | + | 10.8 | 7.76E-05 | 0.0775 | CGGCTCGGCATCG | *nifE* | protein NifE |
| AVAEIV_RS02285 | 456235 | 456247 | -25 | - | 10.8 | 7.76E-05 | 0.0804 | CAATTCGGCAGCG | *mutY* | A/G-specific adenine glycosylase |
| AVAEIV_RS02720 | 555718 | 555730 | 334 | + | 10.8 | 7.76E-05 | 0.0804 | CCATTGGCCAGCC | - | hypothetical protein |
| AVAEIV_RS02725 | 555718 | 555730 | 3 | + | 10.8 | 7.76E-05 | 0.0775 | CCATTGGCCAGCC | *hslO* | Hsp33 family molecular chaperone HslO |

| | | | | | | | | | | |
|---|---|---|---|---|---|---|---|---|---|---|
| AVAEIV_RS03615 | 728318 | 728330 | 36 | - | 10.8 | 7.76E-05 | 0.0804 | CGGAGGGCCGCCA | *msrP* | subunit MsrP |
| AVAEIV_RS03620 | 728318 | 728330 | 56 | - | 10.8 | 7.76E-05 | 0.0775 | CGGAGGGCCGCCA | - | SDR family NAD(P)-dependent oxidoreductase |
| AVAEIV_RS04470 | 912150 | 912162 | 209 | + | 10.8 | 7.76E-05 | 0.0804 | CACATCGCCGCCC | - | type III PLP-dependent enzyme |
| AVAEIV_RS05135 | 1056159 | 1056171 | 138 | - | 10.8 | 7.76E-05 | 0.0804 | CAGATAGCCAGCC | - | ferritin-like domain-containing protein |
| AVAEIV_RS05140 | 1056159 | 1056171 | 106 | - | 10.8 | 7.76E-05 | 0.0775 | CAGATAGCCAGCC | - | DUF4142 domain-containing protein |
| AVAEIV_RS05735 | 1174523 | 1174535 | 260 | + | 10.8 | 7.76E-05 | 0.0804 | CAGCCCGCCGTCG | - | family protein |
| AVAEIV_RS06145 | 1253298 | 1253310 | 315 | - | 10.8 | 7.76E-05 | 0.0804 | CTGAATGCCATCC | - | type 4 pilus major pilin |
| AVAEIV_RS07540 | 1571305 | 1571317 | 229 | - | 10.8 | 7.76E-05 | 0.0804 | CGGCTCGGCATCG | - | CsiV family protein |
| AVAEIV_RS08415 | 1757839 | 1757851 | 260 | - | 10.8 | 7.76E-05 | 0.0775 | CAGCCCGCCGTCG | - | family protein |
| AVAEIV_RS08545 | 1784641 | 1784653 | 299 | - | 10.8 | 7.76E-05 | 0.0775 | CGGCTCGGCATCG | - | hypothetical protein |
| AVAEIV_RS11555 | 2442475 | 2442487 | 230 | - | 10.8 | 7.76E-05 | 0.0775 | CAGACCGGCATCG | - | heavy metal translocating P-type ATPase |
| AVAEIV_RS12070 | 2562273 | 2562285 | 49 | + | 10.8 | 7.76E-05 | 0.0775 | CCGACGGCCATCC | - | MFS transporter |
| AVAEIV_RS13735 | 2970932 | 2970944 | 340 | - | 10.8 | 7.76E-05 | 0.0804 | CCCAGCGCCGCCA | - | LysR family transcriptional regulator |
| AVAEIV_RS14075 | 3060519 | 3060531 | 258 | + | 10.8 | 7.76E-05 | 0.0804 | CCGAGCGCCTTCA | *pyk* | pyruvate kinase |
| AVAEIV_RS15990 | 3485068 | 3485080 | 366 | - | 10.8 | 7.76E-05 | 0.0775 | CAGGTCGCCAGGG | - | ABC transporter ATP-binding protein |
| AVAEIV_RS17485 | 3787222 | 3787234 | 117 | - | 10.8 | 7.76E-05 | 0.0804 | CGGCTCGGCATCG | - | ABC transporter ATP-binding protein |
| AVAEIV_RS17845 | 3862092 | 3862104 | -43 | + | 10.8 | 7.76E-05 | 0.0775 | CCGCTCGCCACCT | - | OprD family porin |
| AVAEIV_RS18865 | 4062393 | 4062405 | -28 | - | 10.8 | 7.76E-05 | 0.0804 | CGACTCGCCGTCG | - | CinA family protein |
| AVAEIV_RS19490 | 4168845 | 4168857 | 30 | + | 10.8 | 7.76E-05 | 0.0775 | CAGGGCGGCATCA | - | HNH endonuclease |
| AVAEIV_RS20900 | 4443416 | 4443428 | 82 | + | 10.8 | 7.76E-05 | 0.0775 | CAGCTCGCCGTCT | - | ATP-NAD kinase family protein |
| AVAEIV_RS21225 | 4512922 | 4512934 | -27 | + | 10.8 | 7.76E-05 | 0.0804 | CAGAATGCCTGCA | - | paraquat-inducible protein A |
| AVAEIV_RS22790 | 4878552 | 4878564 | 144 | - | 10.8 | 7.76E-05 | 0.0775 | CAGCCCGCCGTCG | - | aldehyde dehydrogenase family protein |
| AVAEIV_RS23905 | 5094542 | 5094554 | 131 | + | 10.8 | 7.76E-05 | 0.0804 | CCGAGAGCCAGCA | *hemB* | porphobilinogen synthase |
| AVAEIV_RS23910 | 5094542 | 5094554 | 213 | + | 10.8 | 7.76E-05 | 0.0775 | CCGAGAGCCAGCA | *elbB* | isoprenoid biosynthesis glyoxalase ElbB |
| AVAEIV_RS24490 | 5204992 | 5205004 | 304 | - | 10.8 | 7.76E-05 | 0.0775 | CCGAGAGCCAGCA | - | protein |
| AVAEIV_RS24945 | 5309073 | 5309085 | 161 | + | 10.8 | 7.76E-05 | 0.0804 | CAGCCCGCCGTCG | - | dehydrogenase |
| AVAEIV_RS25455 | 5422073 | 5422085 | 133 | + | 10.8 | 7.76E-05 | 0.0775 | CAGTGCGCGAGCA | - | ArsC/Spx/MgsR family protein |
| AVAEIV_RS00110 | 24412 | 24424 | -11 | + | 10.7857 | 7.83E-05 | 0.0806 | CGCTTGGCCATCT | - | NADPH:quinone reductase |
| AVAEIV_RS00115 | 24412 | 24424 | 164 | + | 10.7857 | 7.83E-05 | 0.0775 | CGCTTGGCCATCT | *hemF* | oxygen-dependent coproporphyrinogen oxidase |
| AVAEIV_RS02675 | 546662 | 546674 | 89 | + | 10.7857 | 7.83E-05 | 0.0806 | CGCGCCGCCAGCA | - | glycosyltransferase |
| AVAEIV_RS04080 | 828168 | 828180 | 210 | - | 10.7857 | 7.83E-05 | 0.0775 | CGCGCCGCCAGCA | *rsfS* | ribosome silencing factor |
| AVAEIV_RS05180 | 1063798 | 1063810 | 90 | - | 10.7857 | 7.83E-05 | 0.0775 | CGCGCCGCCAGCA | - | cytochrome c oxidase assembly protein |
| AVAEIV_RS05555 | 1149591 | 1149603 | 324 | - | 10.7857 | 7.83E-05 | 0.0775 | CGCGCCGCCAGCA | - | hypothetical protein |
| AVAEIV_RS05585 | 1154509 | 1154521 | 25 | - | 10.7857 | 7.83E-05 | 0.0775 | CGGCGGGCCGTCA | - | DUF6012 family protein |
| AVAEIV_RS09955 | 2083686 | 2083698 | 300 | - | 10.7857 | 7.83E-05 | 0.0806 | CTGTAGGCCACCG | - | CopD family protein |
| AVAEIV_RS12990 | 2815343 | 2815355 | 330 | + | 10.7857 | 7.83E-05 | 0.0806 | CGGATGGCCAGGG | - | (2Fe-2S)-binding protein |
| AVAEIV_RS14200 | 3092047 | 3092059 | 225 | + | 10.7857 | 7.83E-05 | 0.0775 | CACATCGGCATCC | - | DUF934 domain-containing protein |
| AVAEIV_RS15000 | 3269148 | 3269160 | 77 | + | 10.7857 | 7.83E-05 | 0.0806 | CCCATCGCCAGGG | - | sensor histidine kinase |
| AVAEIV_RS15530 | 3397149 | 3397161 | 166 | + | 10.7857 | 7.83E-05 | 0.0775 | CGCGCCGCCAGCA | *codA* | cytosine deaminase |
| AVAEIV_RS18030 | 3898665 | 3898677 | 25 | + | 10.7857 | 7.83E-05 | 0.0806 | CGGCGGGCCGTCA | - | DUF6012 family protein |
| AVAEIV_RS18035 | 3898665 | 3898677 | 236 | + | 10.7857 | 7.83E-05 | 0.0775 | CGGCGGGCCGTCA | - | hypothetical protein |
| AVAEIV_RS18055 | 3902263 | 3902275 | 324 | + | 10.7857 | 7.83E-05 | 0.0806 | CGCGCCGCCAGCA | - | hypothetical protein |
| AVAEIV_RS19315 | 4149682 | 4149694 | 268 | - | 10.7857 | 7.83E-05 | 0.0806 | CGCTCGGCCATCG | - | hypothetical protein |
| AVAEIV_RS20780 | 4414553 | 4414565 | 166 | - | 10.7857 | 7.83E-05 | 0.0775 | CTGAGTGCCACCG | - | Arm DNA-binding domain-containing protein |
| AVAEIV_RS21400 | 4552703 | 4552715 | 121 | + | 10.7857 | 7.83E-05 | 0.0775 | CGCGCCGCCAGCA | - | DUF2845 domain-containing protein |
| AVAEIV_RS21485 | 4572009 | 4572021 | 43 | + | 10.7857 | 7.83E-05 | 0.0806 | CGCTTGGCCATCT | *carB* | carbamoyl-phosphate synthase large subunit |
| AVAEIV_RS21685 | 4617818 | 4617830 | 304 | - | 10.7857 | 7.83E-05 | 0.0775 | CGGATGGCCAGGG | - | 2-hydroxy-3-oxopropionate reductase |
| AVAEIV_RS23160 | 4946759 | 4946771 | 273 | - | 10.7857 | 7.83E-05 | 0.0775 | CGGAAGGCCAGGA | - | hypothetical protein |
| AVAEIV_RS23360 | 4978443 | 4978455 | 99 | + | 10.7857 | 7.83E-05 | 0.0775 | CCCCTCGCCACCC | - | S8 family serine peptidase |
| AVAEIV_RS24800 | 5275226 | 5275238 | 134 | - | 10.7857 | 7.83E-05 | 0.0806 | GAGAGGGCCACCA | - | DUF2877 domain-containing protein |
| AVAEIV_RS24865 | 5290418 | 5290430 | 54 | - | 10.7857 | 7.83E-05 | 0.0806 | CGGAAGGCCAGGA | - | subunit alpha |
| AVAEIV_RS00230 | 46485 | 46497 | -33 | + | 10.7714 | 7.91E-05 | 0.0775 | CGGACCGCCGTCG | - | GTP-binding protein |

| | | | | | | | | | | |
|---|---|---|---|---|---|---|---|---|---|---|
| AVAEIV_RS00710 | 143954 | 143966 | 238 | + | 10.7714 | 7.91E-05 | 0.0775 | CTCATCGCCGGCG | *nifN* | protein NifN |
| AVAEIV_RS04715 | 959970 | 959982 | 307 | - | 10.7714 | 7.91E-05 | 0.0775 | CTCAACGCCGGCA | - | AI-2E family transporter |
| AVAEIV_RS05630 | 1158047 | 1158059 | 167 | + | 10.7714 | 7.91E-05 | 0.0775 | CGCATCGACATCG | - | CopG family antitoxin |
| AVAEIV_RS05635 | 1158047 | 1158059 | 382 | + | 10.7714 | 7.91E-05 | 0.0775 | CGCATCGACATCG | - | hypothetical protein |
| AVAEIV_RS06745 | 1388535 | 1388547 | 218 | + | 10.7714 | 7.91E-05 | 0.0775 | CTGATCGCTACCA | *hisD* | histidinol dehydrogenase |
| AVAEIV_RS06940 | 1428095 | 1428107 | 357 | + | 10.7714 | 7.91E-05 | 0.0775 | CGGATCGCCGTCT | - | D-alanine--D-alanine ligase |
| AVAEIV_RS07410 | 1539129 | 1539141 | 345 | - | 10.7714 | 7.91E-05 | 0.0775 | CAGATCGCCGCGA | - | protein |
| AVAEIV_RS08000 | 1662863 | 1662875 | 121 | - | 10.7714 | 7.91E-05 | 0.0808 | CACTTCGACAGCG | - | heavy metal sensor histidine kinase |
| AVAEIV_RS10075 | 2106208 | 2106220 | 268 | - | 10.7714 | 7.91E-05 | 0.0775 | CGCGGCGCCAGCG | - | AtpZ/AtpI family protein |
| AVAEIV_RS10920 | 2294156 | 2294168 | 43 | - | 10.7714 | 7.91E-05 | 0.0808 | CGCATCGACATCG | *uraD* | decarboxylase |
| AVAEIV_RS11645 | 2463764 | 2463776 | 135 | + | 10.7714 | 7.91E-05 | 0.0808 | CGGACCGCCGTCG | - | MliC family protein |
| AVAEIV_RS11680 | 2475725 | 2475737 | 150 | + | 10.7714 | 7.91E-05 | 0.0775 | CGGTTCGGCAGCG | - | CHAD domain-containing protein |
| AVAEIV_RS12835 | 2753053 | 2753065 | 254 | + | 10.7714 | 7.91E-05 | 0.0808 | CAGATCGCCGCGA | - | non-ribosomal peptide synthetase |
| AVAEIV_RS13465 | 2913157 | 2913169 | 112 | - | 10.7714 | 7.91E-05 | 0.0808 | GAGCGGGCCATCA | - | efflux RND transporter permease subunit |
| AVAEIV_RS14380 | 3130167 | 3130179 | 77 | + | 10.7714 | 7.91E-05 | 0.0808 | CCTTATGCCAGCA | - | RNA polymerase sigma factor FliA |
| AVAEIV_RS15640 | 3425365 | 3425377 | 252 | + | 10.7714 | 7.91E-05 | 0.0808 | ACGATCGCCAGCA | *ccmD* | heme exporter protein CcmD |
| AVAEIV_RS19770 | 4213411 | 4213423 | 261 | - | 10.7714 | 7.91E-05 | 0.0808 | GGGATCGCCACCG | - | tellurite resistance TerB family protein |
| AVAEIV_RS20435 | 4345287 | 4345299 | 372 | + | 10.7714 | 7.91E-05 | 0.0808 | CAGATCGCCGCGA | - | Lon protease family protein |
| AVAEIV_RS20445 | 4345287 | 4345299 | 258 | + | 10.7714 | 7.91E-05 | 0.0775 | CAGATCGCCGCGA | - | DUF3015 domain-containing protein |
| AVAEIV_RS20900 | 4443354 | 4443366 | 144 | + | 10.7714 | 7.91E-05 | 0.0775 | GGGAACGCCACCA | - | ATP-NAD kinase family protein |
| AVAEIV_RS21150 | 4495901 | 4495913 | 24 | - | 10.7714 | 7.91E-05 | 0.0808 | CGCATCGACATCG | - | containing protein |
| AVAEIV_RS22220 | 4742032 | 4742044 | 101 | + | 10.7714 | 7.91E-05 | 0.0808 | CAGTCCGCCGGCG | - | NUDIX domain-containing protein |
| AVAEIV_RS22225 | 4742032 | 4742044 | 68 | + | 10.7714 | 7.91E-05 | 0.0775 | CAGTCCGCCGGCG | - | DUF3298 and DUF4163 domain-containing protein |
| AVAEIV_RS22760 | 4872907 | 4872919 | 357 | - | 10.7714 | 7.91E-05 | 0.0775 | CACCGCGCCGCCA | - | DeoR/GlpR family DNA-binding transcription regulator |
| AVAEIV_RS24220 | 5149686 | 5149698 | 274 | + | 10.7714 | 7.91E-05 | 0.0808 | CGCGGCGCCAGCG | - | dihydrofolate reductase |
| AVAEIV_RS26065 | 5605733 | 5605745 | 367 | - | 10.7714 | 7.91E-05 | 0.0808 | CAGCGCGCCTTCA | - | DUF4433 domain-containing protein |
| AVAEIV_RS01180 | 231294 | 231306 | 203 | + | 10.7571 | 7.99E-05 | 0.0809 | TGCATGGCCAGCA | - | glutamate-cysteine ligase family protein |
| AVAEIV_RS01190 | 232736 | 232748 | 61 | - | 10.7571 | 7.99E-05 | 0.0809 | GAGACCGCCATCG | - | ATP-binding cassette domain-containing protein |
| AVAEIV_RS04445 | 905800 | 905812 | 40 | - | 10.7571 | 7.99E-05 | 0.0809 | CACTTCGCCGGCC | - | tetratricopeptide repeat protein |
| AVAEIV_RS05785 | 1186726 | 1186738 | 256 | + | 10.7571 | 7.99E-05 | 0.0809 | CCCATCGCCAGTG | - | 3-hydroxyacyl-CoA dehydrogenase |
| AVAEIV_RS06650 | 1374163 | 1374175 | 257 | - | 10.7571 | 7.99E-05 | 0.0775 | CACATCGCCCGCA | *pmbA* | metalloprotease PmbA |
| AVAEIV_RS06720 | 1385183 | 1385195 | 106 | - | 10.7571 | 7.99E-05 | 0.0775 | CGAGTCGCCATCC | - | phospholipid-binding protein MlaC |
| AVAEIV_RS07265 | 1506920 | 1506932 | 126 | + | 10.7571 | 7.99E-05 | 0.0809 | CGGTGGGCCGGCA | - | ABC transporter ATP-binding protein |
| AVAEIV_RS09665 | 2024558 | 2024570 | 73 | - | 10.7571 | 7.99E-05 | 0.0809 | CCGAACGCGAGCA | - | alpha/beta fold hydrolase |
| AVAEIV_RS11360 | 2403017 | 2403029 | 208 | - | 10.7571 | 7.99E-05 | 0.0809 | GAGATCGCCATCT | *guaD* | guanine deaminase |
| AVAEIV_RS12545 | 2666375 | 2666387 | 318 | - | 10.7571 | 7.99E-05 | 0.0775 | CGGAAGGCCAGTA | - | SulP family inorganic anion transporter |
| AVAEIV_RS13450 | 2910144 | 2910156 | 363 | + | 10.7571 | 7.99E-05 | 0.0809 | CGCAATGCCGGCA | - | SDR family NAD(P)-dependent oxidoreductase |
| AVAEIV_RS15200 | 3310509 | 3310521 | 276 | - | 10.7571 | 7.99E-05 | 0.0809 | GAGATCGCCATCT | - | hypothetical protein |
| AVAEIV_RS15495 | 3387464 | 3387476 | -44 | + | 10.7571 | 7.99E-05 | 0.0775 | CCATCCGCCAGCG | - | polysaccharide biosynthesis protein |
| AVAEIV_RS15730 | 3437578 | 3437590 | 133 | + | 10.7571 | 7.99E-05 | 0.0809 | CCGATCGCGAGCG | - | protein |
| AVAEIV_RS15995 | 3487696 | 3487708 | 44 | + | 10.7571 | 7.99E-05 | 0.0809 | GGGCTCGCCATCG | - | ATP-binding protein |
| AVAEIV_RS16705 | 3618178 | 3618190 | 306 | + | 10.7571 | 7.99E-05 | 0.0809 | CCATCCGCCAGCG | *cbiB* | adenosylcobinamide-phosphate synthase CbiB |
| AVAEIV_RS18085 | 3908076 | 3908088 | -41 | + | 10.7571 | 7.99E-05 | 0.0809 | CCGAGCGCCAACA | *ybgF* | tol-pal system protein YbgF |
| AVAEIV_RS19175 | 4132951 | 4132963 | 187 | - | 10.7571 | 7.99E-05 | 0.0809 | CACTTCGCCGGCC | - | hypothetical protein |
| AVAEIV_RS19555 | 4175165 | 4175177 | -41 | + | 10.7571 | 7.99E-05 | 0.0775 | CCCTGCGCCGGCA | - | hypothetical protein |
| AVAEIV_RS19560 | 4175165 | 4175177 | 241 | + | 10.7571 | 7.99E-05 | 0.0775 | CCCTGCGCCGGCA | - | DNA cytosine methyltransferase |
| AVAEIV_RS24325 | 5172951 | 5172963 | 114 | + | 10.7571 | 7.99E-05 | 0.0809 | TGCATGGCCAGCA | - | AEC family transporter |
| AVAEIV_RS25935 | 5571726 | 5571738 | 223 | + | 10.7571 | 7.99E-05 | 0.0775 | CTGATGGCCAGCT | - | ABC transporter substrate-binding protein |
| AVAEIV_RS00590 | 120555 | 120567 | 123 | + | 10.7429 | 8.07E-05 | 0.0775 | CTGCTCGCCGCCA | - | 5-(carboxyamino)imidazole ribonucleotide synthase |
| AVAEIV_RS02995 | 617919 | 617931 | 351 | + | 10.7429 | 8.07E-05 | 0.0775 | CCGCTGGCCGGCG | - | lysophospholipid acyltransferase family protein |
| AVAEIV_RS03490 | 702347 | 702359 | 193 | + | 10.7429 | 8.07E-05 | 0.0775 | CCGCTGGCCGGCG | *speA* | arginine decarboxylase |
| AVAEIV_RS04530 | 927836 | 927848 | 295 | + | 10.7429 | 8.07E-05 | 0.081 | CCGCTGGCCGGCG | - | dicarboxylate/amino acid:cation symporter |

| | | | | | | | | | | |
|---|---|---|---|---|---|---|---|---|---|---|
| AVAEIV_RS04785 | 974100 | 974112 | 58 | + | 10.7429 | 8.07E-05 | 0.0775 | CTGCTCGCCGCCA | - | malonate decarboxylase subunit delta |
| AVAEIV_RS04790 | 974100 | 974112 | 350 | + | 10.7429 | 8.07E-05 | 0.0775 | CTGCTCGCCGCCA | - | beta |
| AVAEIV_RS04815 | 977938 | 977950 | 259 | + | 10.7429 | 8.07E-05 | 0.0775 | CTGCTCGCCGCCA | *madM* | malonate transporter subunit MadM |
| AVAEIV_RS06775 | 1397007 | 1397019 | 85 | - | 10.7429 | 8.07E-05 | 0.081 | CCATTTGCCGGCA | - | YhcB family protein |
| AVAEIV_RS06780 | 1397007 | 1397019 | 171 | - | 10.7429 | 8.07E-05 | 0.0775 | CCATTTGCCGGCA | - | alpha/beta hydrolase |
| AVAEIV_RS07850 | 1634199 | 1634211 | 245 | - | 10.7429 | 8.07E-05 | 0.081 | CTGCTCGCCGCCA | - | DUF2897 family protein |
| AVAEIV_RS08970 | 1866397 | 1866409 | 49 | - | 10.7429 | 8.07E-05 | 0.0775 | CGGCCCGCCACCG | - | hypothetical protein |
| AVAEIV_RS09810 | 2053109 | 2053121 | 286 | - | 10.7429 | 8.07E-05 | 0.0775 | TCGCTCGCCATCG | *htpG* | molecular chaperone HtpG |
| AVAEIV_RS10825 | 2273155 | 2273167 | 244 | - | 10.7429 | 8.07E-05 | 0.081 | CCGCTGGCCGGCG | - | TIGR03986 family CRISPR-associated RAMP protein |
| AVAEIV_RS11170 | 2358707 | 2358719 | -10 | - | 10.7429 | 8.07E-05 | 0.081 | CCGATGGGCAGCG | - | energy transducer TonB |
| AVAEIV_RS12485 | 2649584 | 2649596 | 288 | - | 10.7429 | 8.07E-05 | 0.081 | GACATCGCCATCC | - | zinc-dependent alcohol dehydrogenase |
| AVAEIV_RS12585 | 2672952 | 2672964 | 153 | - | 10.7429 | 8.07E-05 | 0.0775 | CAATTGGCGATCA | *uvrC* | excinuclease ABC subunit UvrC |
| AVAEIV_RS14335 | 3123128 | 3123140 | 94 | + | 10.7429 | 8.07E-05 | 0.081 | CTCATGGCCAGCC | *rfbF* | glucose-1-phosphate cytidylyltransferase |
| AVAEIV_RS16130 | 3516020 | 3516032 | 384 | + | 10.7429 | 8.07E-05 | 0.0775 | CGCAGCGCCGCCA | - | Gfo/Idh/MocA family oxidoreductase |
| AVAEIV_RS16600 | 3598206 | 3598218 | 138 | + | 10.7429 | 8.07E-05 | 0.0775 | CCGCTGGCCGGCG | - | vWA domain-containing protein |
| AVAEIV_RS17150 | 3717961 | 3717973 | 381 | + | 10.7429 | 8.07E-05 | 0.081 | CAGTGCGCCTGCA | - | SDR family oxidoreductase |
| AVAEIV_RS18305 | 3942347 | 3942359 | 66 | - | 10.7429 | 8.07E-05 | 0.081 | CGGAGAGCCAGCA | - | ORF6N domain-containing protein |
| AVAEIV_RS18805 | 4049754 | 4049766 | 0 | + | 10.7429 | 8.07E-05 | 0.0775 | CCAATGGCCGGCG | - | hypothetical protein |
| AVAEIV_RS18810 | 4049754 | 4049766 | 278 | + | 10.7429 | 8.07E-05 | 0.0775 | CCAATGGCCGGCG | *tsaA* | methyltransferase TrmO |
| AVAEIV_RS19695 | 4198809 | 4198821 | 246 | - | 10.7429 | 8.07E-05 | 0.081 | CTGCTCGCCGCCA | - | extensin family protein |
| AVAEIV_RS20315 | 4327897 | 4327909 | 66 | + | 10.7429 | 8.07E-05 | 0.0775 | CGGAGAGCCAGCA | - | Rha family transcriptional regulator |
| AVAEIV_RS21870 | 4661358 | 4661370 | 48 | + | 10.7429 | 8.07E-05 | 0.081 | CGCAGCGCCGCCA | - | acyl-CoA dehydrogenase family protein |
| AVAEIV_RS21970 | 4686369 | 4686381 | 227 | - | 10.7429 | 8.07E-05 | 0.0775 | CGGCCCGCCACCG | - | acetate/propionate family kinase |
| AVAEIV_RS22095 | 4715379 | 4715391 | 257 | - | 10.7429 | 8.07E-05 | 0.0775 | CCGAAGGGCAGCA | - | nucleotidyltransferase family protein |
| AVAEIV_RS23580 | 5025940 | 5025952 | 163 | + | 10.7429 | 8.07E-05 | 0.0775 | CTGCTCGCCGCCA | *apaG* | Co2+/Mg2+ efflux protein ApaG |
| AVAEIV_RS25570 | 5465870 | 5465882 | 30 | - | 10.7429 | 8.07E-05 | 0.081 | CTGCTCGCCGCCA | - | 2-dehydro-3-deoxy-6-phosphogalactonate aldolase |
| AVAEIV_RS01900 | 380531 | 380543 | 360 | + | 10.7286 | 8.14E-05 | 0.081 | CGGATCGCCGTGA | - | DUF2790 domain-containing protein |
| AVAEIV_RS03070 | 627071 | 627083 | 109 | - | 10.7286 | 8.14E-05 | 0.0775 | CGAATTGCCGCCA | *rplL* | 50S ribosomal protein L7/L12 |
| AVAEIV_RS05895 | 1208391 | 1208403 | 218 | + | 10.7286 | 8.14E-05 | 0.081 | GACAGCGCCACCA | - | TIGR03751 family conjugal transfer lipoprotein |
| AVAEIV_RS05900 | 1208391 | 1208403 | -2 | + | 10.7286 | 8.14E-05 | 0.081 | GACAGCGCCACCA | - | protein |
| AVAEIV_RS08485 | 1774548 | 1774560 | 240 | - | 10.7286 | 8.14E-05 | 0.081 | GGGTACGCCAGCA | - | ABC1 kinase family protein |
| AVAEIV_RS09725 | 2035511 | 2035523 | 326 | - | 10.7286 | 8.14E-05 | 0.081 | CAGCCCGCCATCT | - | thermonuclease family protein |
| AVAEIV_RS14405 | 3137147 | 3137159 | 349 | - | 10.7286 | 8.14E-05 | 0.081 | CCGAGCGACAGCA | *flhB* | flagellar biosynthesis protein FlhB |
| AVAEIV_RS15250 | 3330556 | 3330568 | 323 | - | 10.7286 | 8.14E-05 | 0.081 | CTTTTCGCCAGCG | *rnhA* | ribonuclease HI |
| AVAEIV_RS20295 | 4326033 | 4326045 | 339 | + | 10.7286 | 8.14E-05 | 0.0775 | CAGCTCGCGAGCG | *istB* | IS21-like element helper ATPase IstB |
| AVAEIV_RS22175 | 4733424 | 4733436 | 178 | - | 10.7286 | 8.14E-05 | 0.081 | CCGGTCGCCGGCG | - | diguanylate cyclase domain-containing protein |
| AVAEIV_RS22720 | 4864719 | 4864731 | 378 | + | 10.7286 | 8.14E-05 | 0.081 | CGGATCGCCGTGA | - | class 1 fructose-bisphosphatase |
| AVAEIV_RS22765 | 4874129 | 4874141 | 193 | + | 10.7286 | 8.14E-05 | 0.0775 | CCGTAGGCCGTCG | *glpD* | glycerol-3-phosphate dehydrogenase |
| AVAEIV_RS23805 | 5073659 | 5073671 | 258 | - | 10.7286 | 8.14E-05 | 0.081 | CCGGTCGCCGGCG | *gcvP* | aminomethyl-transferring glycine dehydrogenase |
| AVAEIV_RS24790 | 5271569 | 5271581 | 133 | - | 10.7286 | 8.14E-05 | 0.0775 | CGACCCGCCATCG | - | containing protein |
| AVAEIV_RS25870 | 5547710 | 5547722 | -51 | - | 10.7286 | 8.14E-05 | 0.0775 | CCGGTCGCCGGCG | - | ABC transporter ATP-binding protein |
| AVAEIV_RS00420 | 85806 | 85818 | 118 | + | 10.7143 | 8.22E-05 | 0.0775 | CACGCGGCCATCG | *znuC* | zinc ABC transporter ATP-binding protein ZnuC |
| AVAEIV_RS01465 | 284356 | 284368 | 381 | + | 10.7143 | 8.22E-05 | 0.081 | CAGAGGGCGAGCA | *rdgB* | pyrophosphatase |
| AVAEIV_RS01470 | 284356 | 284368 | -35 | + | 10.7143 | 8.22E-05 | 0.081 | CAGAGGGCGAGCA | - | DUF4426 domain-containing protein |
| AVAEIV_RS03040 | 624014 | 624026 | 106 | + | 10.7143 | 8.22E-05 | 0.0775 | CGTTTCGCCATCC | - | tRNA-Trp |
| AVAEIV_RS03045 | 624014 | 624026 | 224 | + | 10.7143 | 8.22E-05 | 0.0775 | CGTTTCGCCATCC | *secE* | preprotein translocase subunit SecE |
| AVAEIV_RS03440 | 695098 | 695110 | -29 | + | 10.7143 | 8.22E-05 | 0.081 | CGGTTGGCGATCA | *accC* | acetyl-CoA carboxylase biotin carboxylase subunit |
| AVAEIV_RS07565 | 1578895 | 1578907 | 74 | - | 10.7143 | 8.22E-05 | 0.0775 | AGGATCGCCAGCA | - | Na(+)-translocating NADH-quinone reductase subunit C |
| AVAEIV_RS09140 | 1904551 | 1904563 | 240 | + | 10.7143 | 8.22E-05 | 0.081 | AGGATCGCCAGCA | - | sensor histidine kinase |
| AVAEIV_RS12275 | 2599859 | 2599871 | 361 | - | 10.7143 | 8.22E-05 | 0.081 | CAACTGGCCGGCG | *fliI* | flagellar protein export ATPase FliI |
| AVAEIV_RS12715 | 2701365 | 2701377 | 83 | - | 10.7143 | 8.22E-05 | 0.081 | CCCTTGGCCGCCA | - | RNA polymerase factor sigma-70 |
| AVAEIV_RS12720 | 2701365 | 2701377 | 226 | - | 10.7143 | 8.22E-05 | 0.0775 | CCCTTGGCCGCCA | - | diaminobutyrate--2-oxoglutarate transaminase |

| | | | | | | | | | | |
|---|---|---|---|---|---|---|---|---|---|---|
| AVAEIV_RS14870 | 3233610 | 3233622 | -47 | - | 10.7143 | 8.22E-05 | 0.0775 | CGGTTGGCGATCA | - | hypothetical protein |
| AVAEIV_RS15315 | 3346011 | 3346023 | 75 | + | 10.7143 | 8.22E-05 | 0.081 | CTCCGGGCCAGCA | *sucD* | succinate--CoA ligase subunit alpha |
| AVAEIV_RS15635 | 3424893 | 3424905 | -47 | + | 10.7143 | 8.22E-05 | 0.081 | AGGATCGCCAGCA | *ccmE* | cytochrome c maturation protein CcmE |
| AVAEIV_RS15770 | 3443445 | 3443457 | 237 | + | 10.7143 | 8.22E-05 | 0.0775 | CCATTCGCCAGGA | - | HipA domain-containing protein |
| AVAEIV_RS18555 | 3999089 | 3999101 | -36 | - | 10.7143 | 8.22E-05 | 0.0775 | CATATCGCTATCG | - | hypothetical protein |
| AVAEIV_RS18900 | 4069967 | 4069979 | 367 | + | 10.7143 | 8.22E-05 | 0.0775 | CAACTGGCCGGCG | *fdxA* | ferredoxin FdxA |
| AVAEIV_RS19260 | 4143163 | 4143175 | 51 | + | 10.7143 | 8.22E-05 | 0.081 | CTCTCGGCCATCA | - | hypothetical protein |
| AVAEIV_RS19300 | 4147182 | 4147194 | 365 | - | 10.7143 | 8.22E-05 | 0.081 | CCCTTGGCCGCCA | - | hypothetical protein |
| AVAEIV_RS19735 | 4206204 | 4206216 | 160 | - | 10.7143 | 8.22E-05 | 0.081 | CAACTGGCCGGCG | - | alpha/beta hydrolase |
| AVAEIV_RS20540 | 4366992 | 4367004 | 382 | - | 10.7143 | 8.22E-05 | 0.081 | CCGCCTGCCACCA | - | FKBP-type peptidyl-prolyl cis-trans isomerase |
| AVAEIV_RS21180 | 4503726 | 4503738 | 124 | + | 10.7143 | 8.22E-05 | 0.081 | CCGACCGCCATGA | - | RNA-guided endonuclease InsQ/TnpB family protein |
| AVAEIV_RS21185 | 4503726 | 4503738 | 189 | + | 10.7143 | 8.22E-05 | 0.0775 | CCGACCGCCATGA | - | hypothetical protein |
| AVAEIV_RS21845 | 4655746 | 4655758 | 187 | + | 10.7143 | 8.22E-05 | 0.081 | CAACTGGCCGGCG | - | ABC transporter substrate-binding protein |
| AVAEIV_RS22425 | 4795491 | 4795503 | 106 | - | 10.7143 | 8.22E-05 | 0.081 | CACGCGGCCATCG | - | FecR family protein |
| AVAEIV_RS22595 | 4839947 | 4839959 | 159 | + | 10.7143 | 8.22E-05 | 0.0775 | CAACTGGCCGGCG | *hslV* | ATP-dependent protease subunit HslV |
| AVAEIV_RS23635 | 5038546 | 5038558 | 76 | - | 10.7143 | 8.22E-05 | 0.081 | AGGATCGCCAGCA | *folB* | dihydroneopterin aldolase |
| AVAEIV_RS23640 | 5038546 | 5038558 | -23 | - | 10.7143 | 8.22E-05 | 0.0775 | AGGATCGCCAGCA | *plsY* | glycerol-3-phosphate 1-O-acyltransferase PlsY |
| AVAEIV_RS24505 | 5207758 | 5207770 | 57 | + | 10.7143 | 8.22E-05 | 0.081 | AGGATCGCCAGCA | - | ABC transporter permease subunit |
| AVAEIV_RS24865 | 5290335 | 5290347 | -29 | + | 10.7143 | 8.22E-05 | 0.081 | CGGTTGGCGATCA | - | subunit alpha |
| AVAEIV_RS00795 | 156127 | 156139 | 138 | - | 10.7 | 8.29E-05 | 0.0775 | CAGTCCGCCAGCT | *cysE* | serine O-acetyltransferase |
| AVAEIV_RS01190 | 232966 | 232978 | 291 | + | 10.7 | 8.29E-05 | 0.081 | GTGATCGCCACCA | - | ATP-binding cassette domain-containing protein |
| AVAEIV_RS03000 | 618827 | 618839 | 135 | - | 10.7 | 8.29E-05 | 0.0775 | CCCTTGGGCATCA | - | phosphatidate cytidylyltransferase |
| AVAEIV_RS03145 | 644546 | 644558 | 274 | + | 10.7 | 8.29E-05 | 0.0775 | CAGAACGCCATGC | *rplP* | 50S ribosomal protein L16 |
| AVAEIV_RS03185 | 647771 | 647783 | 108 | - | 10.7 | 8.29E-05 | 0.0775 | CAGACCGCCACGA | *rplF* | 50S ribosomal protein L6 |
| AVAEIV_RS04105 | 833371 | 833383 | 280 | + | 10.7 | 8.29E-05 | 0.0775 | CCGGTGGCCAGCC | - | septal ring lytic transglycosylase RlpA family protein |
| AVAEIV_RS04985 | 1019699 | 1019711 | 193 | - | 10.7 | 8.29E-05 | 0.0775 | TGGATCGCCACCG | - | Leu/Phe/Val dehydrogenase |
| AVAEIV_RS06525 | 1345932 | 1345944 | 177 | - | 10.7 | 8.29E-05 | 0.0775 | GCGATGGCCAGCG | - | biotin-dependent carboxyltransferase family protein |
| AVAEIV_RS06985 | 1439631 | 1439643 | 47 | + | 10.7 | 8.29E-05 | 0.0775 | CCGGTGGCCAGCC | - | Nudix family hydrolase |
| AVAEIV_RS07580 | 1580904 | 1580916 | 148 | - | 10.7 | 8.29E-05 | 0.0775 | GCGATGGCCAGCG | *nqrF* | subunit F |
| AVAEIV_RS07710 | 1606626 | 1606638 | 350 | + | 10.7 | 8.29E-05 | 0.0775 | CAGCGTGCCGTCG | *fabG* | 3-oxoacyl-ACP reductase FabG |
| AVAEIV_RS08015 | 1665948 | 1665960 | 98 | - | 10.7 | 8.29E-05 | 0.081 | CTATTCGCCGGCA | - | TrkH family potassium uptake protein |
| AVAEIV_RS08020 | 1665948 | 1665960 | 112 | - | 10.7 | 8.29E-05 | 0.0775 | CTATTCGCCGGCA | - | YkgJ family cysteine cluster protein |
| AVAEIV_RS08170 | 1697936 | 1697948 | 232 | + | 10.7 | 8.29E-05 | 0.0775 | CAGCGTGCCGTCG | *rpsA* | 30S ribosomal protein S1 |
| AVAEIV_RS08610 | 1795564 | 1795576 | 353 | - | 10.7 | 8.29E-05 | 0.0775 | GCGATGGCCAGCG | - | cysteine desulfurase |
| AVAEIV_RS09855 | 2063865 | 2063877 | 258 | - | 10.7 | 8.29E-05 | 0.081 | CCATAGGCCACCG | - | electron transport complex subunit E |
| AVAEIV_RS10735 | 2248370 | 2248382 | 276 | + | 10.7 | 8.29E-05 | 0.0775 | CCGAACGCCTGCA | - | acetone carboxylase subunit gamma |
| AVAEIV_RS10870 | 2281609 | 2281621 | 353 | - | 10.7 | 8.29E-05 | 0.0775 | GCGATGGCCAGCG | - | cysteine desulfurase |
| AVAEIV_RS11060 | 2331102 | 2331114 | 49 | - | 10.7 | 8.29E-05 | 0.0775 | CGCTGCGCCGGCA | - | FAD-binding oxidoreductase |
| AVAEIV_RS11730 | 2486667 | 2486679 | 361 | - | 10.7 | 8.29E-05 | 0.081 | CGGATCGCGAGCG | - | ABC transporter ATP-binding protein |
| AVAEIV_RS11740 | 2486667 | 2486679 | 334 | - | 10.7 | 8.29E-05 | 0.0775 | CGGATCGCGAGCG | - | hypothetical protein |
| AVAEIV_RS11770 | 2492707 | 2492719 | 92 | - | 10.7 | 8.29E-05 | 0.081 | CAGAAAGCCACCG | - | phosphoadenylyl-sulfate reductase |
| AVAEIV_RS12500 | 2652837 | 2652849 | 266 | + | 10.7 | 8.29E-05 | 0.081 | GCGAAGGCCAGCA | - | DUF2934 domain-containing protein |
| AVAEIV_RS13000 | 2815890 | 2815902 | 354 | - | 10.7 | 8.29E-05 | 0.0775 | GTGATCGCCACCA | - | LysR family transcriptional regulator |
| AVAEIV_RS13105 | 2841153 | 2841165 | 159 | + | 10.7 | 8.29E-05 | 0.0775 | GTGATCGCCACCA | *gcvT* | glycine cleavage system aminomethyltransferase GcvT |
| AVAEIV_RS13145 | 2851999 | 2852011 | 296 | - | 10.7 | 8.29E-05 | 0.081 | CGCTGCGCCGGCA | *hutG* | N-formylglutamate deformylase |
| AVAEIV_RS14895 | 3244955 | 3244967 | 102 | - | 10.7 | 8.29E-05 | 0.081 | CCGATCGCCTGCG | - | permease subunit |
| AVAEIV_RS15035 | 3274879 | 3274891 | 211 | - | 10.7 | 8.29E-05 | 0.081 | CTGTTCGGCAGCA | - | type II secretion system protein |
| AVAEIV_RS15475 | 3382498 | 3382510 | 84 | + | 10.7 | 8.29E-05 | 0.0775 | CGCTGCGCCGGCA | - | glycosyltransferase family 4 protein |
| AVAEIV_RS15480 | 3383359 | 3383371 | 347 | - | 10.7 | 8.29E-05 | 0.0775 | CAGCGTGCCGTCG | - | sugar transferase |
| AVAEIV_RS16175 | 3526969 | 3526981 | 38 | + | 10.7 | 8.29E-05 | 0.0775 | CGCTGCGCCGGCA | - | methionine ABC transporter permease |
| AVAEIV_RS18630 | 4011925 | 4011937 | -19 | + | 10.7 | 8.29E-05 | 0.0775 | CGGCTCGCCACGA | - | cupin domain-containing protein |
| AVAEIV_RS18645 | 4013086 | 4013098 | 234 | + | 10.7 | 8.29E-05 | 0.0775 | CGCTGCGCCGGCA | - | shikimate dehydrogenase |

| | | | | | | | | | | |
|---|---|---|---|---|---|---|---|---|---|---|
| AVAEIV_RS18725 | 4032889 | 4032901 | 381 | + | 10.7 | 8.29E-05 | 0.081 | CGCTGCGCCGGCA | *pcaH* | protocatechuate 3,4-dioxygenase subunit beta |
| AVAEIV_RS19285 | 4145731 | 4145743 | 149 | + | 10.7 | 8.29E-05 | 0.081 | CAGCGTGCCGTCG | - | hypothetical protein |
| AVAEIV_RS21455 | 4564252 | 4564264 | 102 | + | 10.7 | 8.29E-05 | 0.081 | GCGATGGCCAGCG | *glmM* | phosphoglucosamine mutase |
| AVAEIV_RS21920 | 4674122 | 4674134 | 43 | - | 10.7 | 8.29E-05 | 0.0775 | CGCTGCGCCGGCA | - | hypothetical protein |
| AVAEIV_RS22585 | 4837458 | 4837470 | -23 | + | 10.7 | 8.29E-05 | 0.0775 | CGATTCGCCAGCT | *argS* | arginine--tRNA ligase |
| AVAEIV_RS23365 | 4981347 | 4981359 | -50 | + | 10.7 | 8.29E-05 | 0.081 | GCGAAGGCCAGCA | - | EamA family transporter |
| AVAEIV_RS23370 | 4981347 | 4981359 | 353 | + | 10.7 | 8.29E-05 | 0.0775 | GCGAAGGCCAGCA | - | LysR family transcriptional regulator |
| AVAEIV_RS23485 | 5004609 | 5004621 | 218 | + | 10.7 | 8.29E-05 | 0.081 | CATATCGCCGCCG | - | IS630 family transposase |
| AVAEIV_RS24010 | 5112920 | 5112932 | 376 | - | 10.7 | 8.29E-05 | 0.081 | CAGCGTGCCGTCG | *cyaY* | iron donor protein CyaY |
| AVAEIV_RS24020 | 5112920 | 5112932 | -33 | - | 10.7 | 8.29E-05 | 0.0775 | CAGCGTGCCGTCG | *lysA* | diaminopimelate decarboxylase |
| AVAEIV_RS25010 | 5324129 | 5324141 | 210 | + | 10.7 | 8.29E-05 | 0.0775 | CTCATCGACATCA | - | Gfo/Idh/MocA family protein |
| AVAEIV_RS25340 | 5397892 | 5397904 | 319 | + | 10.7 | 8.29E-05 | 0.0775 | CAGGACGGCAGCA | *tssC* | subunit |
| AVAEIV_RS25890 | 5554975 | 5554987 | 128 | + | 10.7 | 8.29E-05 | 0.081 | GCGAAGGCCAGCA | - | formylglycine-generating enzyme family protein |
| AVAEIV_RS00140 | 30744 | 30756 | 367 | + | 10.6857 | 8.38E-05 | 0.081 | CGGCTGGCCGGCG | - | MgtC/SapB family protein |
| AVAEIV_RS01350 | 264769 | 264781 | 50 | + | 10.6857 | 8.38E-05 | 0.081 | CAGGGGGCCGGCA | *vnfA1* | regulator VnfA1 |
| AVAEIV_RS02495 | 499328 | 499340 | 238 | - | 10.6857 | 8.38E-05 | 0.081 | CGGCTGGCCGGCG | - | AraC family transcriptional regulator |
| AVAEIV_RS03175 | 646695 | 646707 | 290 | + | 10.6857 | 8.38E-05 | 0.0775 | CCGATCGGCGTCA | *rpsN* | 30S ribosomal protein S14 |
| AVAEIV_RS04455 | 908101 | 908113 | 19 | + | 10.6857 | 8.38E-05 | 0.081 | CGAATGGCCGGCG | *thiE* | thiamine phosphate synthase |
| AVAEIV_RS04485 | 916791 | 916803 | 212 | + | 10.6857 | 8.38E-05 | 0.081 | CAGACGGCCGGCG | - | IucA/IucC family protein |
| AVAEIV_RS04785 | 974197 | 974209 | -39 | - | 10.6857 | 8.38E-05 | 0.0775 | CGGCTGGCCGGCG | - | malonate decarboxylase subunit delta |
| AVAEIV_RS04790 | 974197 | 974209 | 253 | - | 10.6857 | 8.38E-05 | 0.0775 | CGGCTGGCCGGCG | - | beta |
| AVAEIV_RS07800 | 1623012 | 1623024 | 299 | + | 10.6857 | 8.38E-05 | 0.0775 | CGGCTGGCCGGCG | - | acetaldehyde dehydrogenase (acetylating) |
| AVAEIV_RS10230 | 2134421 | 2134433 | 357 | + | 10.6857 | 8.38E-05 | 0.081 | CCGATGGACACCA | *ccoO* | cytochrome-c oxidase, cbb3-type subunit II |
| AVAEIV_RS10800 | 2265895 | 2265907 | 292 | + | 10.6857 | 8.38E-05 | 0.081 | CGGCTGGCCGGCG | - | hypothetical protein |
| AVAEIV_RS10955 | 2303738 | 2303750 | -25 | - | 10.6857 | 8.38E-05 | 0.081 | CGCATTGCCAGCT | - | isochorismatase |
| AVAEIV_RS14380 | 3130336 | 3130348 | 246 | + | 10.6857 | 8.38E-05 | 0.081 | CGGCTGGCCGGCG | - | RNA polymerase sigma factor FliA |
| AVAEIV_RS18730 | 4033667 | 4033679 | 101 | - | 10.6857 | 8.38E-05 | 0.081 | CGGCTGGCCGGCG | *pcaQ* | pca operon transcription factor PcaQ |
| AVAEIV_RS19930 | 4248741 | 4248753 | 117 | - | 10.6857 | 8.38E-05 | 0.0775 | CGGCTGGCCGGCG | - | DMT family transporter |
| AVAEIV_RS20090 | 4288847 | 4288859 | 361 | + | 10.6857 | 8.38E-05 | 0.0775 | CAGGGGGCCGGCA | *leuA* | 2-isopropylmalate synthase |
| AVAEIV_RS21355 | 4539477 | 4539489 | 211 | - | 10.6857 | 8.38E-05 | 0.0775 | CAGACGGCCGGCG | - | sigma-54-dependent transcriptional regulator |
| AVAEIV_RS23465 | 5001893 | 5001905 | 57 | - | 10.6857 | 8.38E-05 | 0.081 | CGGCAGGCCGGCA | *hpaI* | 4-hydroxy-2-oxoheptanedioate aldolase |
| AVAEIV_RS23470 | 5001893 | 5001905 | 75 | - | 10.6857 | 8.38E-05 | 0.0775 | CGGCAGGCCGGCA | - | GNAT family N-acetyltransferase |
| AVAEIV_RS24450 | 5197972 | 5197984 | 332 | + | 10.6857 | 8.38E-05 | 0.0775 | CGGCTGGCCGGCG | - | D-hexose-6-phosphate mutarotase |
| AVAEIV_RS24925 | 5304139 | 5304151 | 153 | + | 10.6857 | 8.38E-05 | 0.081 | CGGCTGGCCGGCG | *tpiA* | triose-phosphate isomerase |
| AVAEIV_RS25390 | 5410523 | 5410535 | 204 | + | 10.6857 | 8.38E-05 | 0.081 | CCGATGGACACCA | - | protein |
| AVAEIV_RS25575 | 5467070 | 5467082 | 269 | + | 10.6857 | 8.38E-05 | 0.081 | CGGCTGGCCGGCG | - | 2-dehydro-3-deoxygalactonokinase |
| AVAEIV_RS26065 | 5605435 | 5605447 | 69 | + | 10.6857 | 8.38E-05 | 0.081 | CGCGGTGCCAGCA | - | DUF4433 domain-containing protein |
| AVAEIV_RS01195 | 233645 | 233657 | 269 | + | 10.6714 | 8.45E-05 | 0.081 | CAGTGCGCCGTCC | - | GTP-binding protein |
| AVAEIV_RS02430 | 483187 | 483199 | 365 | - | 10.6714 | 8.45E-05 | 0.081 | CCGCCGGCCAGCG | - | DUF502 domain-containing protein |
| AVAEIV_RS02855 | 586704 | 586716 | 231 | + | 10.6714 | 8.45E-05 | 0.0775 | CAGCTCGCCTGCG | - | TetR/AcrR family transcriptional regulator |
| AVAEIV_RS03575 | 722118 | 722130 | 262 | - | 10.6714 | 8.45E-05 | 0.081 | CTGCACGCCATCC | - | methyltransferase domain-containing protein |
| AVAEIV_RS03580 | 722118 | 722130 | -40 | - | 10.6714 | 8.45E-05 | 0.081 | CTGCACGCCATCC | - | MazG-like family protein |
| AVAEIV_RS04460 | 909244 | 909256 | 266 | - | 10.6714 | 8.45E-05 | 0.081 | CCGCCGGCCAGCG | - | kinase/phosphomethylpyrimidine kinase |
| AVAEIV_RS04530 | 927837 | 927849 | 296 | - | 10.6714 | 8.45E-05 | 0.081 | CCGCCGGCCAGCG | - | dicarboxylate/amino acid:cation symporter |
| AVAEIV_RS05400 | 1119341 | 1119353 | 334 | - | 10.6714 | 8.45E-05 | 0.081 | GAGACTGCCATCA | - | RDD family protein |
| AVAEIV_RS05405 | 1119341 | 1119353 | 24 | - | 10.6714 | 8.45E-05 | 0.081 | GAGACTGCCATCA | - | cold-shock protein |
| AVAEIV_RS05410 | 1119341 | 1119353 | 202 | - | 10.6714 | 8.45E-05 | 0.0775 | GAGACTGCCATCA | - | hypothetical protein |
| AVAEIV_RS05965 | 1216014 | 1216026 | -46 | - | 10.6714 | 8.45E-05 | 0.081 | TACATCGCCATCC | - | hypothetical protein |
| AVAEIV_RS05970 | 1216014 | 1216026 | 192 | - | 10.6714 | 8.45E-05 | 0.0775 | TACATCGCCATCC | - | antitoxin |
| AVAEIV_RS05975 | 1216014 | 1216026 | 373 | - | 10.6714 | 8.45E-05 | 0.0775 | TACATCGCCATCC | - | type II toxin-antitoxin system RelE/ParE family toxin |
| AVAEIV_RS09300 | 1942210 | 1942222 | 302 | + | 10.6714 | 8.45E-05 | 0.081 | CCAACGGCCAGCG | - | DUF3144 domain-containing protein |
| AVAEIV_RS10950 | 2303125 | 2303137 | 230 | + | 10.6714 | 8.45E-05 | 0.081 | CCGCCGGCCAGCG | - | amino acid adenylation domain-containing protein |

| | | | | | | | | | | |
|---|---|---|---|---|---|---|---|---|---|---|
| AVAEIV_RS11270 | 2382344 | 2382356 | 237 | + | 10.6714 | 8.45E-05 | 0.081 | CACTACGCTATCG | *dhaL* | dihydroxyacetone kinase subunit DhaL |
| AVAEIV_RS11650 | 2464644 | 2464656 | 142 | + | 10.6714 | 8.45E-05 | 0.081 | CGGGTCGCCGGCG | - | general secretion pathway protein GspB |
| AVAEIV_RS11705 | 2480471 | 2480483 | 112 | - | 10.6714 | 8.45E-05 | 0.081 | CAGTGCGCCGTCC | - | TatD family hydrolase |
| AVAEIV_RS11710 | 2480471 | 2480483 | 96 | - | 10.6714 | 8.45E-05 | 0.0775 | CAGTGCGCCGTCC | - | transglycosylase SLT domain-containing protein |
| AVAEIV_RS12550 | 2668820 | 2668832 | 280 | + | 10.6714 | 8.45E-05 | 0.081 | CTGCCCGCCACCA | - | ribbon-helix-helix protein, CopG family |
| AVAEIV_RS13945 | 3023467 | 3023479 | 195 | - | 10.6714 | 8.45E-05 | 0.0775 | CAGCGCGCCATGG | - | OprD family porin |
| AVAEIV_RS15005 | 3270228 | 3270240 | 118 | - | 10.6714 | 8.45E-05 | 0.081 | CTGCACGCCATCC | *sixA* | phosphohistidine phosphatase SixA |
| AVAEIV_RS15415 | 3367594 | 3367606 | 142 | + | 10.6714 | 8.45E-05 | 0.0775 | CCGCCGGCCAGCG | - | NAD-dependent epimerase/dehydratase family protein |
| AVAEIV_RS16510 | 3581384 | 3581396 | 379 | - | 10.6714 | 8.45E-05 | 0.0775 | CTGGAGGCCATCG | *dusA* | tRNA dihydrouridine(20/20a) synthase DusA |
| AVAEIV_RS17915 | 3878521 | 3878533 | 373 | + | 10.6714 | 8.45E-05 | 0.081 | TACATCGCCATCC | - | type II toxin-antitoxin system RelE/ParE family toxin |
| AVAEIV_RS17920 | 3878521 | 3878533 | 192 | + | 10.6714 | 8.45E-05 | 0.081 | TACATCGCCATCC | - | antitoxin |
| AVAEIV_RS17925 | 3878521 | 3878533 | -46 | + | 10.6714 | 8.45E-05 | 0.0775 | TACATCGCCATCC | - | hypothetical protein |
| AVAEIV_RS19065 | 4106995 | 4107007 | 190 | - | 10.6714 | 8.45E-05 | 0.081 | CAAATCGCCTGCG | *rpsB* | 30S ribosomal protein S2 |
| AVAEIV_RS19070 | 4106995 | 4107007 | 61 | - | 10.6714 | 8.45E-05 | 0.0775 | CAAATCGCCTGCG | *map* | type I methionyl aminopeptidase |
| AVAEIV_RS20940 | 4451532 | 4451544 | 201 | + | 10.6714 | 8.45E-05 | 0.0775 | CGGGTCGCCGGCG | - | D-alanyl-D-alanine carboxypeptidase family protein |
| AVAEIV_RS20995 | 4465669 | 4465681 | 282 | + | 10.6714 | 8.45E-05 | 0.081 | CGGGTCGCCGGCG | - | LysR substrate-binding domain-containing protein |
| AVAEIV_RS00040 | 8925 | 8937 | 126 | - | 10.6571 | 8.53E-05 | 0.081 | CGGCATGCCATCC | - | hypothetical protein |
| AVAEIV_RS03335 | 672216 | 672228 | 184 | + | 10.6571 | 8.53E-05 | 0.0775 | CTGCGCGCCACCG | *ribD* | diaminohydroxyphosphoribosylaminopyrimidine deaminase/5-amino-6-(5-phosphoribosylamino)uracil |
| AVAEIV_RS04120 | 836226 | 836238 | 22 | + | 10.6571 | 8.53E-05 | 0.0775 | CTGCGCGCCACCG | *lipB* | lipoyl(octanoyl) transferase LipB |
| AVAEIV_RS06860 | 1412375 | 1412387 | 331 | + | 10.6571 | 8.53E-05 | 0.081 | CGTATCGCCGTCG | - | phosphoheptose isomerase |
| AVAEIV_RS09085 | 1890993 | 1891005 | 122 | - | 10.6571 | 8.53E-05 | 0.0775 | TACAGCGCCACCA | - | carbohydrate kinase family protein |
| AVAEIV_RS13775 | 2981181 | 2981193 | 36 | + | 10.6571 | 8.53E-05 | 0.081 | CCCTTCGCCTTCA | - | DUF4150 domain-containing protein |
| AVAEIV_RS17160 | 3720455 | 3720467 | 96 | - | 10.6571 | 8.53E-05 | 0.081 | TGGTTCGCCAGCG | *purF* | amidophosphoribosyltransferase |
| AVAEIV_RS17520 | 3794848 | 3794860 | 238 | + | 10.6571 | 8.53E-05 | 0.081 | GAGGTCGCCAGCG | *pap* | polyphosphate:AMP phosphotransferase |
| AVAEIV_RS17825 | 3857536 | 3857548 | 207 | + | 10.6571 | 8.53E-05 | 0.0775 | CAGTACGCCTTCG | - | hydroxymethylglutaryl-CoA reductase, degradative |
| AVAEIV_RS18825 | 4053537 | 4053549 | 248 | + | 10.6571 | 8.53E-05 | 0.081 | CGCTTGGCCGCCA | *erdR* | response regulator transcription factor ErdR |
| AVAEIV_RS21050 | 4476072 | 4476084 | 101 | - | 10.6571 | 8.53E-05 | 0.081 | CCATGCGCCATCC | - | OprD family porin |
| AVAEIV_RS21335 | 4534666 | 4534678 | 366 | + | 10.6571 | 8.53E-05 | 0.0775 | CCGGCCGCCAGCG | *dksA* | RNA polymerase-binding protein DksA |
| AVAEIV_RS21585 | 4595461 | 4595473 | 165 | + | 10.6571 | 8.53E-05 | 0.081 | CCCTTCGCCTTCA | - | molybdenum storage protein subunit alpha |
| AVAEIV_RS23980 | 5107879 | 5107891 | 69 | - | 10.6571 | 8.53E-05 | 0.081 | CGGCATGCCATCC | *hemC* | hydroxymethylbilane synthase |
| AVAEIV_RS25185 | 5363644 | 5363656 | 369 | - | 10.6571 | 8.53E-05 | 0.081 | CCGGCCGCCAGCG | *hypA* | hydrogenase maturation nickel metallochaperone HypA |
| AVAEIV_RS25320 | 5395774 | 5395786 | 219 | - | 10.6571 | 8.53E-05 | 0.081 | TGGTTCGCCAGCG | - | DotU family type IV/VI secretion system protein |
| AVAEIV_RS01180 | 231066 | 231078 | -25 | - | 10.6429 | 8.61E-05 | 0.081 | CACTACGGCATCG | - | glutamate-cysteine ligase family protein |
| AVAEIV_RS01340 | 261757 | 261769 | 245 | + | 10.6429 | 8.61E-05 | 0.081 | CAGGTGGCGATCA | *nifN* | protein NifN |
| AVAEIV_RS01860 | 369200 | 369212 | 46 | + | 10.6429 | 8.61E-05 | 0.0775 | CGGATCGCCTGCG | - | RNA polymerase sigma factor |
| AVAEIV_RS02645 | 538459 | 538471 | -50 | - | 10.6429 | 8.61E-05 | 0.0775 | CAACCGGCCAGCG | - | DUF6491 family protein |
| AVAEIV_RS02650 | 539887 | 539899 | 31 | - | 10.6429 | 8.61E-05 | 0.081 | CGGAGCGCCATGG | - | ChbG/HpnK family deacetylase |
| AVAEIV_RS04880 | 995288 | 995300 | 92 | + | 10.6429 | 8.61E-05 | 0.0775 | CACTACGGCATCG | - | FAD-dependent oxidoreductase |
| AVAEIV_RS07255 | 1505361 | 1505373 | 232 | + | 10.6429 | 8.61E-05 | 0.081 | CCGGTTGCCGGCA | *glpD* | glycerol-3-phosphate dehydrogenase |
| AVAEIV_RS08945 | 1859245 | 1859257 | 155 | - | 10.6429 | 8.61E-05 | 0.0775 | CATCGGGCCACCA | - | acetyl-CoA C-acyltransferase |
| AVAEIV_RS10760 | 2255631 | 2255643 | -49 | - | 10.6429 | 8.61E-05 | 0.081 | CGGGTGGCCAGCC | - | CRISPR-associated endonuclease Cas1 |
| AVAEIV_RS10995 | 2314020 | 2314032 | 54 | + | 10.6429 | 8.61E-05 | 0.081 | CAGGTCGACACCA | - | tripartite tricarboxylate transporter TctB family protein |
| AVAEIV_RS11335 | 2396341 | 2396353 | 106 | - | 10.6429 | 8.61E-05 | 0.0775 | CCCTCGGCCACCA | - | hypothetical protein |
| AVAEIV_RS11340 | 2396341 | 2396353 | 230 | - | 10.6429 | 8.61E-05 | 0.0775 | CCCTCGGCCACCA | *eno* | phosphopyruvate hydratase |
| AVAEIV_RS12940 | 2804236 | 2804248 | 336 | - | 10.6429 | 8.61E-05 | 0.0775 | GCCATCGCCAGCG | - | BCAM0308 family protein |
| AVAEIV_RS14065 | 3058906 | 3058918 | 230 | + | 10.6429 | 8.61E-05 | 0.081 | CCCTCGGCCACCA | *eno* | phosphopyruvate hydratase |
| AVAEIV_RS14070 | 3058906 | 3058918 | 106 | + | 10.6429 | 8.61E-05 | 0.081 | CCCTCGGCCACCA | - | hypothetical protein |
| AVAEIV_RS15220 | 3315594 | 3315606 | 154 | - | 10.6429 | 8.61E-05 | 0.081 | CAGGTGGCGATCA | - | protein |
| AVAEIV_RS15225 | 3316820 | 3316832 | 154 | - | 10.6429 | 8.61E-05 | 0.081 | CAGGTGGCGATCA | - | 3-oxoacyl-[acyl-carrier-protein] synthase III C-terminal domain-containing protein |
| AVAEIV_RS16460 | 3573043 | 3573055 | 362 | + | 10.6429 | 8.61E-05 | 0.0775 | CCACTCGGCATCA | - | aldo/keto reductase |

| | | | | | | | | | | |
|---|---|---|---|---|---|---|---|---|---|---|
| AVAEIV_RS17275 | 3743435 | 3743447 | 22 | + | 10.6429 | 8.61E-05 | 0.081 | CGATAGGCCACCG | - | glycosyltransferase |
| AVAEIV_RS17755 | 3843281 | 3843293 | -18 | + | 10.6429 | 8.61E-05 | 0.081 | CCACTCGGCATCA | - | sulfurtransferase TusA family protein |
| AVAEIV_RS17760 | 3843281 | 3843293 | 143 | + | 10.6429 | 8.61E-05 | 0.0775 | CCACTCGGCATCA | - | M48 family metalloprotease |
| AVAEIV_RS20285 | 4325392 | 4325404 | 151 | - | 10.6429 | 8.61E-05 | 0.0775 | CAGTGCGCCAGGG | - | IS110-like element ISAzvi4 family transposase |
| AVAEIV_RS21315 | 4532832 | 4532844 | 306 | + | 10.6429 | 8.61E-05 | 0.081 | CAATGCGCCACCC | - | FecCD family ABC transporter permease |
| AVAEIV_RS21560 | 4586254 | 4586266 | 71 | - | 10.6429 | 8.61E-05 | 0.0775 | CAGCGCGCCATTG | - | LutB/LldF family L-lactate oxidation iron-sulfur protein |
| AVAEIV_RS21615 | 4601263 | 4601275 | 106 | - | 10.6429 | 8.61E-05 | 0.0775 | CCCTCGGCCACCA | - | hypothetical protein |
| AVAEIV_RS21620 | 4601263 | 4601275 | 230 | - | 10.6429 | 8.61E-05 | 0.0775 | CCCTCGGCCACCA | *eno* | phosphopyruvate hydratase |
| AVAEIV_RS22115 | 4719543 | 4719555 | 142 | - | 10.6429 | 8.61E-05 | 0.0775 | GCGATCGCCGTCA | - | CDP-alcohol phosphatidyltransferase family protein |
| AVAEIV_RS22190 | 4736909 | 4736921 | 366 | - | 10.6429 | 8.61E-05 | 0.081 | CGGAGCGCCATGG | - | acyl-CoA dehydrogenase family protein |
| AVAEIV_RS22775 | 4876307 | 4876319 | 106 | - | 10.6429 | 8.61E-05 | 0.0775 | CCCTCGGCCACCA | - | hypothetical protein |
| AVAEIV_RS22780 | 4876307 | 4876319 | 230 | - | 10.6429 | 8.61E-05 | 0.0775 | CCCTCGGCCACCA | *eno* | phosphopyruvate hydratase |
| AVAEIV_RS23140 | 4943908 | 4943920 | 54 | - | 10.6429 | 8.61E-05 | 0.0775 | GGGATGGCCAGCG | - | phage tail protein |
| AVAEIV_RS24470 | 5201243 | 5201255 | 46 | - | 10.6429 | 8.61E-05 | 0.0775 | CGGATCGCCTGCG | - | class I SAM-dependent methyltransferase |
| AVAEIV_RS24955 | 5311318 | 5311330 | 230 | + | 10.6429 | 8.61E-05 | 0.081 | CCCTCGGCCACCA | *eno* | phosphopyruvate hydratase |
| AVAEIV_RS24960 | 5311318 | 5311330 | 106 | + | 10.6429 | 8.61E-05 | 0.081 | CCCTCGGCCACCA | - | hypothetical protein |
| AVAEIV_RS00830 | 162522 | 162534 | 365 | - | 10.6286 | 8.68E-05 | 0.081 | CGCATCGGCAGCG | - | hypothetical protein |
| AVAEIV_RS00840 | 162522 | 162534 | 365 | - | 10.6286 | 8.68E-05 | 0.0775 | CGCATCGGCAGCG | - | CobW family GTP-binding protein |
| AVAEIV_RS00885 | 172826 | 172838 | 241 | - | 10.6286 | 8.68E-05 | 0.081 | TCGATGGCCAGCG | - | DUF2388 domain-containing protein |
| AVAEIV_RS01320 | 257505 | 257517 | 370 | - | 10.6286 | 8.68E-05 | 0.0775 | CGCCTCGCCGGCG | - | HesA/MoeB/ThiF family protein |
| AVAEIV_RS02445 | 485525 | 485537 | 290 | + | 10.6286 | 8.68E-05 | 0.0775 | GAGATCGCCGCCA | - | inorganic phosphate transporter |
| AVAEIV_RS02860 | 587241 | 587253 | 330 | - | 10.6286 | 8.68E-05 | 0.0775 | CGCCTCGCCGGCG | - | glutathione S-transferase family protein |
| AVAEIV_RS03060 | 625739 | 625751 | -34 | + | 10.6286 | 8.68E-05 | 0.0775 | CAGAAGGCGATCG | *rplA* | 50S ribosomal protein L1 |
| AVAEIV_RS03800 | 766428 | 766440 | 211 | + | 10.6286 | 8.68E-05 | 0.0775 | TCGAAGGCCAGCA | - | ferredoxin--NADP reductase |
| AVAEIV_RS04205 | 858215 | 858227 | 216 | + | 10.6286 | 8.68E-05 | 0.0775 | GAGATCGCCGCCA | *dmpH* | 2-oxo-3-hexenedioate decarboxylase |
| AVAEIV_RS04820 | 979329 | 979341 | 262 | + | 10.6286 | 8.68E-05 | 0.0775 | CCGCTGGCCAGGA | - | sodium:proton antiporter |
| AVAEIV_RS05000 | 1022958 | 1022970 | 174 | + | 10.6286 | 8.68E-05 | 0.0775 | GAGATCGCCGCCA | - | dihydrolipoamide acetyltransferase family protein |
| AVAEIV_RS05185 | 1066092 | 1066104 | -46 | - | 10.6286 | 8.68E-05 | 0.081 | CGCCTCGCCGGCG | - | metal-dependent hydrolase family protein |
| AVAEIV_RS05760 | 1180824 | 1180836 | 331 | - | 10.6286 | 8.68E-05 | 0.081 | CACATCGCCGGCT | - | acyl-CoA dehydrogenase |
| AVAEIV_RS06900 | 1417717 | 1417729 | -3 | - | 10.6286 | 8.68E-05 | 0.0775 | CATGGCGCCACCA | - | peptidoglycan D,D-transpeptidase FtsI family protein |
| AVAEIV_RS07385 | 1531903 | 1531915 | 158 | - | 10.6286 | 8.68E-05 | 0.0775 | CGCCTCGCCGGCG | - | p-hydroxycinnamoyl CoA hydratase/lyase |
| AVAEIV_RS07990 | 1659414 | 1659426 | 280 | + | 10.6286 | 8.68E-05 | 0.0775 | CCCATGGCGATCA | - | ABC transporter ATP-binding protein |
| AVAEIV_RS08290 | 1725341 | 1725353 | 237 | - | 10.6286 | 8.68E-05 | 0.0775 | TCGAAGGCCAGCA | - | transposase |
| AVAEIV_RS08840 | 1833826 | 1833838 | 353 | - | 10.6286 | 8.68E-05 | 0.0775 | CACGGCGCCGGCA | - | hypothetical protein |
| AVAEIV_RS10400 | 2174303 | 2174315 | 246 | - | 10.6286 | 8.68E-05 | 0.0775 | CCCATGGCGATCA | - | PA2778 family cysteine peptidase |
| AVAEIV_RS10980 | 2310711 | 2310723 | 116 | + | 10.6286 | 8.68E-05 | 0.081 | TCGATGGCCAGCG | *purB* | adenylosuccinate lyase |
| AVAEIV_RS11955 | 2540367 | 2540379 | 349 | + | 10.6286 | 8.68E-05 | 0.081 | CAGTTCGGCGGCA | *folD* | dehydrogenase/methenyltetrahydrofolate cyclohydrolase FolD |
| AVAEIV_RS11965 | 2540367 | 2540379 | 16 | + | 10.6286 | 8.68E-05 | 0.0775 | CAGTTCGGCGGCA | - | tRNA-His |
| AVAEIV_RS11970 | 2540367 | 2540379 | 154 | + | 10.6286 | 8.68E-05 | 0.0775 | CAGTTCGGCGGCA | - | tRNA-Leu |
| AVAEIV_RS11975 | 2540367 | 2540379 | 369 | + | 10.6286 | 8.68E-05 | 0.0775 | CAGTTCGGCGGCA | *tig* | trigger factor |
| AVAEIV_RS12690 | 2695200 | 2695212 | 363 | + | 10.6286 | 8.68E-05 | 0.081 | CACACCGCCGGCG | - | 3-hydroxybutyryl-CoA dehydrogenase |
| AVAEIV_RS12950 | 2805746 | 2805758 | 377 | + | 10.6286 | 8.68E-05 | 0.0775 | CGCCTCGCCGGCG | - | Hsp20/alpha crystallin family protein |
| AVAEIV_RS14260 | 3106731 | 3106743 | 348 | + | 10.6286 | 8.68E-05 | 0.081 | CCGAATGCCGGCG | - | flagellar motor protein MotB |
| AVAEIV_RS14410 | 3138081 | 3138093 | 211 | + | 10.6286 | 8.68E-05 | 0.081 | CTCCAGGCCATCG | *cheZ* | protein phosphatase CheZ |
| AVAEIV_RS14915 | 3252941 | 3252953 | 361 | - | 10.6286 | 8.68E-05 | 0.081 | CAGTTCGGCGGCA | - | containing protein |
| AVAEIV_RS14990 | 3264835 | 3264847 | 290 | - | 10.6286 | 8.68E-05 | 0.0775 | GAGATCGCCGCCA | - | MFS transporter |
| AVAEIV_RS15230 | 3318189 | 3318201 | 306 | + | 10.6286 | 8.68E-05 | 0.081 | TCGATGGCCAGCG | - | type III polyketide synthase |
| AVAEIV_RS15650 | 3426883 | 3426895 | 267 | + | 10.6286 | 8.68E-05 | 0.081 | CCGACCGCCGCCA | *ccmB* | heme exporter protein CcmB |
| AVAEIV_RS16425 | 3567755 | 3567767 | 278 | + | 10.6286 | 8.68E-05 | 0.081 | CAGCGTGCCATCT | - | Bcr/CflA family multidrug efflux MFS transporter |
| AVAEIV_RS16440 | 3570808 | 3570820 | 139 | - | 10.6286 | 8.68E-05 | 0.081 | CGCAACGGCAGCA | - | LysR family transcriptional regulator |
| AVAEIV_RS16445 | 3570808 | 3570820 | 96 | - | 10.6286 | 8.68E-05 | 0.0775 | CGCAACGGCAGCA | - | aldo/keto reductase |

| | | | | | | | | | | |
|---|---|---|---|---|---|---|---|---|---|---|
| AVAEIV_RS18365 | 3951418 | 3951430 | 45 | + | 10.6286 | 8.68E-05 | 0.081 | TCGAAGGCCAGCA | - | alpha/beta hydrolase |
| AVAEIV_RS18690 | 4026147 | 4026159 | 242 | + | 10.6286 | 8.68E-05 | 0.081 | TCGATGGCCAGCG | - | 3-carboxy-cis,cis-muconate cycloisomerase |
| AVAEIV_RS19175 | 4132880 | 4132892 | 116 | + | 10.6286 | 8.68E-05 | 0.081 | CGCCTCGCCGGCG | - | hypothetical protein |
| AVAEIV_RS19300 | 4146902 | 4146914 | 85 | + | 10.6286 | 8.68E-05 | 0.081 | TCGATGGCCAGCG | - | hypothetical protein |
| AVAEIV_RS20585 | 4378463 | 4378475 | -38 | + | 10.6286 | 8.68E-05 | 0.081 | CTCCAGGCCATCG | - | hybrid sensor histidine kinase/response regulator |
| AVAEIV_RS22600 | 4840477 | 4840489 | 258 | - | 10.6286 | 8.68E-05 | 0.0775 | CCCATGGCGATCA | *hslU* | ATP-dependent protease ATPase subunit HslU |
| AVAEIV_RS23630 | 5038269 | 5038281 | 143 | + | 10.6286 | 8.68E-05 | 0.081 | CGCCTCGCCGGCG | *folK* | diphosphokinase |
| AVAEIV_RS23640 | 5038269 | 5038281 | 254 | + | 10.6286 | 8.68E-05 | 0.0775 | CGCCTCGCCGGCG | *plsY* | glycerol-3-phosphate 1-O-acyltransferase PlsY |
| AVAEIV_RS23880 | 5087459 | 5087471 | 170 | - | 10.6286 | 8.68E-05 | 0.081 | GAGATCGCCGCCA | *ubiD* | 4-hydroxy-3-polyprenylbenzoate decarboxylase |
| AVAEIV_RS24005 | 5111801 | 5111813 | 151 | + | 10.6286 | 8.68E-05 | 0.0775 | GAGATCGCCGCCA | - | TIGR02647 family protein |
| AVAEIV_RS25120 | 5348549 | 5348561 | 108 | - | 10.6286 | 8.68E-05 | 0.0775 | CCGCTGGCCAGGA | - | ABC transporter substrate-binding protein |
| AVAEIV_RS00845 | 164458 | 164470 | -46 | - | 10.6143 | 8.76E-05 | 0.081 | CTGCTGGCCGGCA | - | DUF3301 domain-containing protein |
| AVAEIV_RS01040 | 205465 | 205477 | 98 | + | 10.6143 | 8.76E-05 | 0.081 | CGGCCGGCCAGCG | - | dodecin |
| AVAEIV_RS01755 | 346559 | 346571 | 306 | + | 10.6143 | 8.76E-05 | 0.081 | CGGCTGGCCAGCT | *dctM* | protein DctM |
| AVAEIV_RS03580 | 722448 | 722460 | 290 | + | 10.6143 | 8.76E-05 | 0.081 | CGCAGGGCCGGCA | - | MazG-like family protein |
| AVAEIV_RS05450 | 1128291 | 1128303 | 86 | - | 10.6143 | 8.76E-05 | 0.0775 | CGGCCGGCCAGCG | *ispH* | 4-hydroxy-3-methylbut-2-enyl diphosphate reductase |
| AVAEIV_RS06150 | 1254300 | 1254312 | 137 | - | 10.6143 | 8.76E-05 | 0.081 | CCGCCCGCCGTCA | - | type II secretion system F family protein |
| AVAEIV_RS09045 | 1881822 | 1881834 | 310 | - | 10.6143 | 8.76E-05 | 0.081 | CTGCTGGCCGGCA | - | 3-hydroxybutyrate dehydrogenase |
| AVAEIV_RS09175 | 1911147 | 1911159 | 74 | - | 10.6143 | 8.76E-05 | 0.0775 | CGGCCGGCCAGCG | - | CHAD domain-containing protein |
| AVAEIV_RS09200 | 1920344 | 1920356 | -17 | + | 10.6143 | 8.76E-05 | 0.0775 | CTGAGCGCCATCT | - | DUF2334 domain-containing protein |
| AVAEIV_RS09390 | 1957446 | 1957458 | 343 | - | 10.6143 | 8.76E-05 | 0.0775 | CCGTTGGCCATGG | - | hypothetical protein |
| AVAEIV_RS09585 | 1998574 | 1998586 | 271 | + | 10.6143 | 8.76E-05 | 0.0775 | CGGCCGGCCAGCG | - | YeaC family protein |
| AVAEIV_RS10430 | 2182435 | 2182447 | 382 | + | 10.6143 | 8.76E-05 | 0.0775 | CACCGTGCCATCC | *infC* | translation initiation factor IF-3 |
| AVAEIV_RS12215 | 2587731 | 2587743 | -20 | - | 10.6143 | 8.76E-05 | 0.0775 | CCGAGCGCCGCCG | *flgJ* | flagellar assembly peptidoglycan hydrolase FlgJ |
| AVAEIV_RS13025 | 2821192 | 2821204 | 174 | + | 10.6143 | 8.76E-05 | 0.0775 | CGCTACGCCGTCG | - | methyltransferase family protein |
| AVAEIV_RS13465 | 2913130 | 2913142 | 85 | - | 10.6143 | 8.76E-05 | 0.081 | GACCTCGCCAGCG | - | efflux RND transporter permease subunit |
| AVAEIV_RS13775 | 2981461 | 2981473 | 316 | - | 10.6143 | 8.76E-05 | 0.081 | GACCTCGCCAGCG | - | DUF4150 domain-containing protein |
| AVAEIV_RS14360 | 3126628 | 3126640 | 140 | + | 10.6143 | 8.76E-05 | 0.081 | GCAATCGCCACCA | - | type II toxin-antitoxin system HicB family antitoxin |
| AVAEIV_RS14370 | 3126628 | 3126640 | 207 | + | 10.6143 | 8.76E-05 | 0.0775 | GCAATCGCCACCA | - | flagellar protein FlaG |
| AVAEIV_RS14825 | 3222471 | 3222483 | 350 | + | 10.6143 | 8.76E-05 | 0.0775 | CTGCTGGCCGGCA | - | type II secretion system F family protein |
| AVAEIV_RS15120 | 3291455 | 3291467 | -1 | + | 10.6143 | 8.76E-05 | 0.0775 | CGGCTGGACATCA | - | ABC transporter substrate-binding protein |
| AVAEIV_RS15125 | 3292520 | 3292532 | 100 | + | 10.6143 | 8.76E-05 | 0.0775 | CCGCCCGCCGTCA | - | ATP-binding protein |
| AVAEIV_RS15320 | 3347428 | 3347440 | 326 | - | 10.6143 | 8.76E-05 | 0.081 | CCGAGCGCCGCCG | *sucC* | ADP-forming succinate--CoA ligase subunit beta |
| AVAEIV_RS15540 | 3400446 | 3400458 | 286 | - | 10.6143 | 8.76E-05 | 0.081 | CTGCTGGCCGGCA | - | sugar phosphate isomerase/epimerase family protein |
| AVAEIV_RS16540 | 3587114 | 3587126 | 285 | - | 10.6143 | 8.76E-05 | 0.0775 | CGGTGCGCCACCC | - | adenylate/guanylate cyclase domain-containing protein |
| AVAEIV_RS19735 | 4206405 | 4206417 | 361 | - | 10.6143 | 8.76E-05 | 0.081 | GAGATCGGCATCA | - | alpha/beta hydrolase |
| AVAEIV_RS19775 | 4214623 | 4214635 | 323 | - | 10.6143 | 8.76E-05 | 0.081 | CTCGACGCCATCG | - | NADH:flavin oxidoreductase/NADH oxidase |
| AVAEIV_RS20350 | 4330567 | 4330579 | 149 | + | 10.6143 | 8.76E-05 | 0.0775 | CTGCTGGCCGGCA | - | hypothetical protein |
| AVAEIV_RS23185 | 4953232 | 4953244 | 297 | + | 10.6143 | 8.76E-05 | 0.081 | CGGCTGGCCAGCT | - | hypothetical protein |
| AVAEIV_RS02640 | 537179 | 537191 | 76 | + | 10.6 | 8.83E-05 | 0.0775 | CCGAGCGGCATCG | - | 2-hydroxyacid dehydrogenase |
| AVAEIV_RS03315 | 670805 | 670817 | 173 | - | 10.6 | 8.83E-05 | 0.081 | CGCCTGGCCAGCC | *trxA* | thioredoxin |
| AVAEIV_RS03710 | 747190 | 747202 | 71 | - | 10.6 | 8.83E-05 | 0.0775 | CGGCTCGGCACCA | *hflX* | ribosome rescue GTPase HflX |
| AVAEIV_RS03925 | 791373 | 791385 | 83 | + | 10.6 | 8.83E-05 | 0.081 | CAGAGTGCCACGA | - | aldehyde dehydrogenase family protein |
| AVAEIV_RS03930 | 791373 | 791385 | 213 | + | 10.6 | 8.83E-05 | 0.0775 | CAGAGTGCCACGA | - | regulator |
| AVAEIV_RS04820 | 979635 | 979647 | -44 | - | 10.6 | 8.83E-05 | 0.0775 | GACAGGGCCAGCA | - | sodium:proton antiporter |
| AVAEIV_RS05360 | 1108526 | 1108538 | 92 | + | 10.6 | 8.83E-05 | 0.081 | CGCCTGGCCAGCC | *rlmF* | RlmF |
| AVAEIV_RS05805 | 1189292 | 1189304 | 234 | + | 10.6 | 8.83E-05 | 0.0775 | CCGCGCGCCGTCG | - | class I adenylate-forming enzyme family protein |
| AVAEIV_RS06070 | 1239247 | 1239259 | 237 | + | 10.6 | 8.83E-05 | 0.081 | CAGAACGGCATCC | - | hypothetical protein |
| AVAEIV_RS06490 | 1338180 | 1338192 | 175 | + | 10.6 | 8.83E-05 | 0.081 | CAGGAGGCCGTCG | - | PepSY-associated TM helix domain-containing protein |
| AVAEIV_RS08395 | 1751657 | 1751669 | 256 | - | 10.6 | 8.83E-05 | 0.0775 | CCGCGCGCCGTCG | *bktB* | beta-ketothiolase BktB |
| AVAEIV_RS09200 | 1920032 | 1920044 | 295 | - | 10.6 | 8.83E-05 | 0.0775 | CTGGTCGCCGGCA | - | DUF2334 domain-containing protein |
| AVAEIV_RS10360 | 2166100 | 2166112 | 103 | - | 10.6 | 8.83E-05 | 0.081 | CGGGCCGCCAGCG | - | helix-turn-helix domain-containing protein |

| | | | | | | | | | | |
|---|---|---|---|---|---|---|---|---|---|---|
| AVAEIV_RS11655 | 2466302 | 2466314 | 86 | - | 10.6 | 8.83E-05 | 0.081 | CGATGCGCCATCC | - | ExeA family protein |
| AVAEIV_RS11930 | 2533702 | 2533714 | 165 | + | 10.6 | 8.83E-05 | 0.0775 | CGCGATGCCATCG | - | tRNA-(ms[2]io[6]A)-hydroxylase |
| AVAEIV_RS12080 | 2563924 | 2563936 | 227 | + | 10.6 | 8.83E-05 | 0.0775 | CGCCTGGCCAGCC | - | 1,2-dihydroxy-3-keto-5-methylthiopentene dioxygenase |
| AVAEIV_RS12355 | 2613000 | 2613012 | 258 | - | 10.6 | 8.83E-05 | 0.0775 | CCCCTTGCCGGCA | - | methyl-accepting chemotaxis protein |
| AVAEIV_RS13465 | 2913376 | 2913388 | 331 | - | 10.6 | 8.83E-05 | 0.081 | CAGAATGGCAGCG | - | efflux RND transporter permease subunit |
| AVAEIV_RS13470 | 2913376 | 2913388 | 122 | - | 10.6 | 8.83E-05 | 0.081 | CAGAATGGCAGCG | - | hypothetical protein |
| AVAEIV_RS14355 | 3126213 | 3126225 | 268 | + | 10.6 | 8.83E-05 | 0.081 | CGACTCGCCGCCA | - | type II toxin-antitoxin system RelE/ParE family toxin |
| AVAEIV_RS15200 | 3310222 | 3310234 | -11 | + | 10.6 | 8.83E-05 | 0.081 | CGATGCGCCATCC | - | hypothetical protein |
| AVAEIV_RS16135 | 3517169 | 3517181 | 253 | - | 10.6 | 8.83E-05 | 0.0775 | CAGCATGCCGGCG | - | TIM barrel protein |
| AVAEIV_RS18545 | 3992990 | 3993002 | -11 | + | 10.6 | 8.83E-05 | 0.0775 | CGATGCGCCATCC | - | protein |
| AVAEIV_RS21130 | 4490217 | 4490229 | -17 | + | 10.6 | 8.83E-05 | 0.0775 | CCGCACGCCACCC | - | TetR/AcrR family transcriptional regulator |
| AVAEIV_RS21325 | 4533677 | 4533689 | 0 | + | 10.6 | 8.83E-05 | 0.081 | CGGGTCGCCAGCT | *sfsA* | DNA/RNA nuclease SfsA |
| AVAEIV_RS22440 | 4798394 | 4798406 | 181 | + | 10.6 | 8.83E-05 | 0.081 | CGCTTCGCCTTCA | - | LysR substrate-binding domain-containing protein |
| AVAEIV_RS22640 | 4846631 | 4846643 | 89 | + | 10.6 | 8.83E-05 | 0.0775 | CAGACCGGCACCA | *tatB* | Sec-independent protein translocase protein TatB |
| AVAEIV_RS23255 | 4959769 | 4959781 | 203 | + | 10.6 | 8.83E-05 | 0.0775 | CAGCCCGCCGCCA | - | DedA family protein |
| AVAEIV_RS23740 | 5059227 | 5059239 | 375 | - | 10.6 | 8.83E-05 | 0.0775 | GACAGGGCCAGCA | - | ABC transporter ATP-binding protein |
| AVAEIV_RS23900 | 5093756 | 5093768 | 377 | - | 10.6 | 8.83E-05 | 0.081 | CGCGATGCCATCG | *ppk1* | polyphosphate kinase 1 |
| AVAEIV_RS24675 | 5244593 | 5244605 | 262 | - | 10.6 | 8.83E-05 | 0.081 | CAGTTGGCCACGG | - | cytochrome b/b6 domain-containing protein |
| AVAEIV_RS24905 | 5299073 | 5299085 | 370 | - | 10.6 | 8.83E-05 | 0.0775 | GACAGGGCCAGCA | - | tRNA-Arg |
| AVAEIV_RS01180 | 231451 | 231463 | 360 | - | 10.5857 | 8.91E-05 | 0.0812 | CAGCGCGCCGCCG | - | glutamate-cysteine ligase family protein |
| AVAEIV_RS01775 | 350251 | 350263 | 62 | + | 10.5857 | 8.91E-05 | 0.0812 | CAGCGCGCCGCCG | - | sigma-54-dependent transcriptional regulator |
| AVAEIV_RS02520 | 505569 | 505581 | 101 | + | 10.5857 | 8.91E-05 | 0.0775 | CGCGTCGCCAGCC | - | ExbD/TolR family protein |
| AVAEIV_RS03200 | 648912 | 648924 | 368 | - | 10.5857 | 8.91E-05 | 0.0775 | CCCTTTGCCGTCG | *rpmD* | 50S ribosomal protein L30 |
| AVAEIV_RS04195 | 855519 | 855531 | 347 | - | 10.5857 | 8.91E-05 | 0.0775 | CAGCGCGCCGCCG | - | OprD family porin |
| AVAEIV_RS04955 | 1014268 | 1014280 | 194 | + | 10.5857 | 8.91E-05 | 0.0812 | TCCTTGGCCATCA | *mmsB* | 3-hydroxyisobutyrate dehydrogenase |
| AVAEIV_RS06425 | 1314922 | 1314934 | 386 | - | 10.5857 | 8.91E-05 | 0.0812 | AACATCGCCATCG | - | IS4-like element ISAzvi7 family transposase |
| AVAEIV_RS08655 | 1807235 | 1807247 | 169 | - | 10.5857 | 8.91E-05 | 0.0812 | CCCAAGGCCGTCG | *rbbA* | RbbA |
| AVAEIV_RS08655 | 1807336 | 1807348 | 270 | + | 10.5857 | 8.91E-05 | 0.0812 | CCGAGGGCCACCC | *rbbA* | RbbA |
| AVAEIV_RS09280 | 1938814 | 1938826 | 245 | - | 10.5857 | 8.91E-05 | 0.0812 | CCGTCCGCCGGCA | - | hypothetical protein |
| AVAEIV_RS09285 | 1938814 | 1938826 | 24 | - | 10.5857 | 8.91E-05 | 0.0775 | CCGTCCGCCGGCA | - | hypothetical protein |
| AVAEIV_RS09330 | 1945769 | 1945781 | 241 | + | 10.5857 | 8.91E-05 | 0.0775 | CGTACCGCCATCG | - | hypothetical protein |
| AVAEIV_RS10200 | 2128021 | 2128033 | 233 | + | 10.5857 | 8.91E-05 | 0.0812 | CGGTTGGACAGCA | *ccoS* | cbb3-type cytochrome oxidase assembly protein CcoS |
| AVAEIV_RS10530 | 2202327 | 2202339 | 362 | - | 10.5857 | 8.91E-05 | 0.0812 | TTGTTCGCCAGCA | *istA* | IS21 family transposase |
| AVAEIV_RS10540 | 2202327 | 2202339 | 78 | - | 10.5857 | 8.91E-05 | 0.0775 | TTGTTCGCCAGCA | - | AAA family ATPase |
| AVAEIV_RS11085 | 2336091 | 2336103 | 76 | - | 10.5857 | 8.91E-05 | 0.0775 | CAGCGCGCCGCCG | - | cysteine dioxygenase family protein |
| AVAEIV_RS11900 | 2528055 | 2528067 | 320 | - | 10.5857 | 8.91E-05 | 0.0812 | CGCTCGGCCACCA | *alaC* | alanine transaminase |
| AVAEIV_RS12065 | 2560770 | 2560782 | 364 | - | 10.5857 | 8.91E-05 | 0.0775 | GGCATCGCCAGCG | *aroC* | chorismate synthase |
| AVAEIV_RS12855 | 2772621 | 2772633 | -31 | - | 10.5857 | 8.91E-05 | 0.0812 | GAGTTCGCCGGCA | - | IS5 family transposase |
| AVAEIV_RS12860 | 2772621 | 2772633 | 150 | - | 10.5857 | 8.91E-05 | 0.0775 | GAGTTCGCCGGCA | - | containing protein |
| AVAEIV_RS13935 | 3021007 | 3021019 | 252 | - | 10.5857 | 8.91E-05 | 0.0775 | CGGGTTGCCGGCA | - | MFS transporter |
| AVAEIV_RS14765 | 3212048 | 3212060 | 87 | - | 10.5857 | 8.91E-05 | 0.0775 | CTGAAAGCCATCA | *nuoI* | NADH-quinone oxidoreductase subunit NuoI |
| AVAEIV_RS17610 | 3816318 | 3816330 | 92 | + | 10.5857 | 8.91E-05 | 0.0812 | CGCGTCGCCAGCC | - | lipid A biosynthesis lauroyl acyltransferase |
| AVAEIV_RS17615 | 3816318 | 3816330 | 128 | + | 10.5857 | 8.91E-05 | 0.0775 | CGCGTCGCCAGCC | *minC* | septum site-determining protein MinC |
| AVAEIV_RS18120 | 3914227 | 3914239 | 137 | - | 10.5857 | 8.91E-05 | 0.0812 | GGCATCGCCAGCG | *ruvB* | Holliday junction branch migration DNA helicase RuvB |
| AVAEIV_RS19560 | 4175382 | 4175394 | 24 | - | 10.5857 | 8.91E-05 | 0.0775 | CAAAGCGCCGCCG | - | DNA cytosine methyltransferase |
| AVAEIV_RS20025 | 4269777 | 4269789 | -40 | + | 10.5857 | 8.91E-05 | 0.0775 | CGTACCGCCATCG | - | MFS transporter |
| AVAEIV_RS20605 | 4383313 | 4383325 | 212 | + | 10.5857 | 8.91E-05 | 0.0775 | GGCATCGCCAGCG | - | 2-hydroxyacid dehydrogenase |
| AVAEIV_RS24795 | 5275037 | 5275049 | 279 | - | 10.5857 | 8.91E-05 | 0.0812 | CAGCGTGCCATGA | - | DUF2790 domain-containing protein |
| AVAEIV_RS24805 | 5275037 | 5275049 | 136 | - | 10.5857 | 8.91E-05 | 0.0775 | CAGCGTGCCATGA | - | hypothetical protein |
| AVAEIV_RS25935 | 5572000 | 5572012 | -51 | - | 10.5857 | 8.91E-05 | 0.0775 | CAGCGCGCCGCCG | - | ABC transporter substrate-binding protein |
| AVAEIV_RS00945 | 187898 | 187910 | 61 | + | 10.5714 | 8.99E-05 | 0.0775 | TCCATCGCCAGCG | - | I78 family peptidase inhibitor |
| AVAEIV_RS01660 | 319760 | 319772 | 329 | - | 10.5714 | 8.99E-05 | 0.0775 | CCGTGCGCCGGCG | - | Na/Pi cotransporter family protein |

| | | | | | | | | | | |
|---|---|---|---|---|---|---|---|---|---|---|
| AVAEIV_RS01805 | 357185 | 357197 | 301 | - | 10.5714 | 8.99E-05 | 0.0813 | CAGAAGGCCTTCG | - | substrate binding protein |
| AVAEIV_RS02010 | 402988 | 403000 | 356 | - | 10.5714 | 8.99E-05 | 0.0775 | CGAATGGCCAGGA | - | C80 family cysteine peptidase |
| AVAEIV_RS03495 | 706547 | 706559 | 329 | + | 10.5714 | 8.99E-05 | 0.0813 | CCGTGCGCCGGCG | - | MATE family efflux transporter |
| AVAEIV_RS03500 | 706547 | 706559 | 39 | + | 10.5714 | 8.99E-05 | 0.0813 | CCGTGCGCCGGCG | - | hypothetical protein |
| AVAEIV_RS04680 | 954387 | 954399 | 376 | - | 10.5714 | 8.99E-05 | 0.0775 | CAGATCGCCCGCG | - | multidrug effflux MFS transporter |
| AVAEIV_RS09260 | 1934665 | 1934677 | 194 | + | 10.5714 | 8.99E-05 | 0.0813 | TACTTGGCCACCA | - | group II truncated hemoglobin |
| AVAEIV_RS11290 | 2386270 | 2386282 | 252 | + | 10.5714 | 8.99E-05 | 0.0813 | CACTTGGCGAGCA | - | ABC transporter permease |
| AVAEIV_RS12080 | 2563846 | 2563858 | 305 | + | 10.5714 | 8.99E-05 | 0.0775 | CAGAAGGCCTTCG | - | 1,2-dihydroxy-3-keto-5-methylthiopentene dioxygenase |
| AVAEIV_RS13235 | 2871170 | 2871182 | 25 | + | 10.5714 | 8.99E-05 | 0.0775 | TCCATCGCCAGCG | - | DUF1016 N-terminal domain-containing protein |
| AVAEIV_RS13720 | 2966645 | 2966657 | 358 | - | 10.5714 | 8.99E-05 | 0.0775 | CAGCGCGGCATCG | - | FecCD family ABC transporter permease |
| AVAEIV_RS15190 | 3307732 | 3307744 | 203 | + | 10.5714 | 8.99E-05 | 0.0813 | GCATTCGCCAGCA | - | ACP S-malonyltransferase |
| AVAEIV_RS18365 | 3951466 | 3951478 | 93 | + | 10.5714 | 8.99E-05 | 0.0813 | GCATTCGCCAGCA | - | alpha/beta hydrolase |
| AVAEIV_RS18380 | 3957950 | 3957962 | 262 | - | 10.5714 | 8.99E-05 | 0.0813 | TGGTTTGCCAGCA | - | TonB-dependent siderophore receptor |
| AVAEIV_RS20600 | 4383444 | 4383456 | 69 | + | 10.5714 | 8.99E-05 | 0.0813 | CGGACCGCCGCCA | - | DUF1302 domain-containing protein |
| AVAEIV_RS20605 | 4383444 | 4383456 | 81 | + | 10.5714 | 8.99E-05 | 0.0775 | CGGACCGCCGCCA | - | 2-hydroxyacid dehydrogenase |
| AVAEIV_RS20915 | 4446455 | 4446467 | 142 | + | 10.5714 | 8.99E-05 | 0.0775 | CAGAAGGCCTTCG | - | acetyltransferase subunit |
| AVAEIV_RS22385 | 4784737 | 4784749 | 253 | + | 10.5714 | 8.99E-05 | 0.0775 | CCCCTCGCCAGGA | - | cation-transporting P-type ATPase |
| AVAEIV_RS22735 | 4867755 | 4867767 | -50 | + | 10.5714 | 8.99E-05 | 0.0813 | CGAATGGCCAGGA | - | hypothetical protein |
| AVAEIV_RS23760 | 5064527 | 5064539 | -18 | + | 10.5714 | 8.99E-05 | 0.0813 | CAGCGCGGCATCG | - | LysR family transcriptional regulator |
| AVAEIV_RS23765 | 5064527 | 5064539 | -7 | + | 10.5714 | 8.99E-05 | 0.0775 | CAGCGCGGCATCG | - | 3-oxoacyl-ACP reductase |
| AVAEIV_RS24110 | 5129899 | 5129911 | 12 | - | 10.5714 | 8.99E-05 | 0.0813 | CAGCACGCCAGGG | - | hypothetical protein |
| AVAEIV_RS00685 | 140474 | 140486 | 34 | - | 10.5571 | 9.05E-05 | 0.0775 | CAGATCGCGGGCA | *nifT* | putative nitrogen fixation protein NifT |
| AVAEIV_RS00690 | 140474 | 140486 | 257 | - | 10.5571 | 9.05E-05 | 0.0775 | CAGATCGCGGGCA | - | protein |
| AVAEIV_RS01090 | 214277 | 214289 | 181 | - | 10.5571 | 9.05E-05 | 0.0815 | CGCCCCGCCAGCG | - | sulfite exporter TauE/SafE family protein |
| AVAEIV_RS01095 | 214277 | 214289 | -17 | - | 10.5571 | 9.05E-05 | 0.0775 | CGCCCCGCCAGCG | - | AraC family transcriptional regulator |
| AVAEIV_RS02730 | 556507 | 556519 | 362 | - | 10.5571 | 9.05E-05 | 0.0775 | CACCAGGCCGTCG | - | phosphoenolpyruvate carboxykinase |
| AVAEIV_RS03605 | 726672 | 726684 | 211 | + | 10.5571 | 9.05E-05 | 0.0815 | CGGAGCGCCGCCG | - | binding protein |
| AVAEIV_RS03620 | 728011 | 728023 | 363 | - | 10.5571 | 9.05E-05 | 0.0775 | CAGGATGCCAGCC | - | SDR family NAD(P)-dependent oxidoreductase |
| AVAEIV_RS04835 | 982698 | 982710 | 237 | + | 10.5571 | 9.05E-05 | 0.0775 | CGGCCCGCCGTCA | - | LTA synthase family protein |
| AVAEIV_RS05835 | 1195408 | 1195420 | 242 | - | 10.5571 | 9.05E-05 | 0.0775 | GCCATTGCCAGCA | - | PaaI family thioesterase |
| AVAEIV_RS06335 | 1291151 | 1291163 | 293 | - | 10.5571 | 9.05E-05 | 0.0775 | CCGGGCGCCATCC | - | energy-coupling factor ABC transporter permease |
| AVAEIV_RS06435 | 1318716 | 1318728 | 364 | + | 10.5571 | 9.05E-05 | 0.0815 | CTGAAGGCCAGCC | - | protein |
| AVAEIV_RS08490 | 1776372 | 1776384 | 115 | + | 10.5571 | 9.05E-05 | 0.0815 | CACAAGGGCATCG | - | DUF3141 domain-containing protein |
| AVAEIV_RS08920 | 1853317 | 1853329 | 362 | + | 10.5571 | 9.05E-05 | 0.0775 | GAGAATGCCAGCG | - | regulator |
| AVAEIV_RS11010 | 2317651 | 2317663 | 272 | + | 10.5571 | 9.05E-05 | 0.0775 | CTGAAGGCCAGCC | - | ABC transporter ATP-binding protein |
| AVAEIV_RS12440 | 2638046 | 2638058 | -40 | - | 10.5571 | 9.05E-05 | 0.0815 | CTCCTCGCCGGCA | - | tetratricopeptide repeat protein |
| AVAEIV_RS12495 | 2652146 | 2652158 | 27 | - | 10.5571 | 9.05E-05 | 0.0815 | CGGAGCGCCGCCG | *glgX* | glycogen debranching protein GlgX |
| AVAEIV_RS13370 | 2893705 | 2893717 | -25 | - | 10.5571 | 9.05E-05 | 0.0815 | GAGAACGCCATCC | - | transposase domain-containing protein |
| AVAEIV_RS14330 | 3122297 | 3122309 | 27 | + | 10.5571 | 9.05E-05 | 0.0815 | CACATGGCCTCCA | *rfbG* | CDP-glucose 4,6-dehydratase |
| AVAEIV_RS21155 | 4498081 | 4498093 | 293 | + | 10.5571 | 9.05E-05 | 0.0815 | CAGTCCGGCAGCA | - | MFS transporter |
| AVAEIV_RS22625 | 4845747 | 4845759 | -48 | - | 10.5571 | 9.05E-05 | 0.0775 | CACCAGGCCGTCG | *hisI* | phosphoribosyl-AMP cyclohydrolase |
| AVAEIV_RS22630 | 4845747 | 4845759 | 346 | - | 10.5571 | 9.05E-05 | 0.0775 | CACCAGGCCGTCG | - | phosphoribosyl-ATP diphosphatase |
| AVAEIV_RS22750 | 4870539 | 4870551 | 248 | + | 10.5571 | 9.05E-05 | 0.0775 | CGGAGCGCCGCCG | - | MIP/aquaporin family protein |
| AVAEIV_RS22945 | 4915741 | 4915753 | 296 | + | 10.5571 | 9.05E-05 | 0.0775 | CTGAAGGCCAGCC | *hemJ* | protoporphyrinogen oxidase HemJ |
| AVAEIV_RS23845 | 5082479 | 5082491 | 133 | + | 10.5571 | 9.05E-05 | 0.0815 | CGGAGCGCCGCCG | - | YecA family protein |
| AVAEIV_RS23850 | 5082479 | 5082491 | 41 | + | 10.5571 | 9.05E-05 | 0.0775 | CGGAGCGCCGCCG | - | TIGR02449 family protein |
| AVAEIV_RS23855 | 5082479 | 5082491 | 247 | + | 10.5571 | 9.05E-05 | 0.0775 | CGGAGCGCCGCCG | - | cell division protein ZapA |
| AVAEIV_RS03220 | 651321 | 651333 | 181 | - | 10.5429 | 9.13E-05 | 0.0776 | CACGACGCCGTCG | *rpsM* | 30S ribosomal protein S13 |
| AVAEIV_RS03685 | 741972 | 741984 | 42 | - | 10.5429 | 9.13E-05 | 0.0776 | TCGCTCGCCACCA | *tsaE* | threonylcarbamoyltransferase complex ATPase subunit type 1 TsaE |
| AVAEIV_RS05325 | 1098269 | 1098281 | 321 | - | 10.5429 | 9.13E-05 | 0.0776 | CTGCCGGCCAGCA | - | DUF2218 domain-containing protein |
| AVAEIV_RS10950 | 2303111 | 2303123 | 216 | - | 10.5429 | 9.13E-05 | 0.0819 | CTGCCGGCCAGCA | - | amino acid adenylation domain-containing protein |

| | | | | | | | | | | |
|---|---|---|---|---|---|---|---|---|---|---|
| AVAEIV_RS12430 | 2634204 | 2634216 | 73 | + | 10.5429 | 9.13E-05 | 0.0776 | CAGGGCGCCACCC | - | response regulator |
| AVAEIV_RS13140 | 2850873 | 2850885 | 178 | - | 10.5429 | 9.13E-05 | 0.0819 | CGGCGCGCCGTCG | - | binding protein |
| AVAEIV_RS13140 | 2850876 | 2850888 | 181 | + | 10.5429 | 9.13E-05 | 0.0819 | CGGCGCGCCGTCG | - | binding protein |
| AVAEIV_RS15270 | 3333588 | 3333600 | -44 | - | 10.5429 | 9.13E-05 | 0.0776 | CCGCTCGCGAGCA | - | extracellular solute-binding protein |
| AVAEIV_RS17570 | 3808471 | 3808483 | 27 | - | 10.5429 | 9.13E-05 | 0.0819 | CAACGGGCCATCC | - | hemolysin family protein |
| AVAEIV_RS18575 | 4002455 | 4002467 | 276 | + | 10.5429 | 9.13E-05 | 0.0819 | CTGCCGGCCAGCA | - | LysR family transcriptional regulator |
| AVAEIV_RS18580 | 4002455 | 4002467 | 18 | + | 10.5429 | 9.13E-05 | 0.0776 | CTGCCGGCCAGCA | - | zinc-binding dehydrogenase |
| AVAEIV_RS19995 | 4262559 | 4262571 | 306 | - | 10.5429 | 9.13E-05 | 0.0819 | CGGAACGCCAGGG | - | ABC transporter substrate-binding protein |
| AVAEIV_RS23150 | 4945371 | 4945383 | 217 | + | 10.5429 | 9.13E-05 | 0.0776 | TACCTCGCCAGCG | - | hypothetical protein |
| AVAEIV_RS24265 | 5158385 | 5158397 | 281 | - | 10.5429 | 9.13E-05 | 0.0776 | TCGCTCGCCACCA | - | NRDE family protein |
| AVAEIV_RS25045 | 5331412 | 5331424 | 344 | + | 10.5429 | 9.13E-05 | 0.0776 | CACCTGGCCTTCA | - | VOC family protein |
| AVAEIV_RS00280 | 61112 | 61124 | 340 | - | 10.5286 | 9.21E-05 | 0.0819 | CTGGCCGCCAGCA | - | ABC transporter permease |
| AVAEIV_RS01945 | 389925 | 389937 | 70 | + | 10.5286 | 9.21E-05 | 0.0776 | CCGTTGGCCGCCG | - | metallophosphoesterase |
| AVAEIV_RS03830 | 772467 | 772479 | 312 | - | 10.5286 | 9.21E-05 | 0.0819 | CTGGCCGCCAGCA | *trmA* | tRNA (uridine(54)-C5)-methyltransferase TrmA |
| AVAEIV_RS04485 | 916780 | 916792 | 201 | + | 10.5286 | 9.21E-05 | 0.0819 | CTGGCCGCCAGCA | - | IucA/IucC family protein |
| AVAEIV_RS07330 | 1518996 | 1519008 | 247 | + | 10.5286 | 9.21E-05 | 0.0776 | AAGTTCGCCACCA | - | tRNA-Pro |
| AVAEIV_RS09940 | 2082159 | 2082171 | 135 | - | 10.5286 | 9.21E-05 | 0.0819 | CTGCGGGCCAGCG | - | YchJ family protein |
| AVAEIV_RS10965 | 2306813 | 2306825 | 133 | - | 10.5286 | 9.21E-05 | 0.0819 | CAAAATGACATCA | - | isochorismate synthase |
| AVAEIV_RS10970 | 2306813 | 2306825 | 65 | - | 10.5286 | 9.21E-05 | 0.0776 | CAAAATGACATCA | - | MFS transporter |
| AVAEIV_RS11955 | 2539988 | 2540000 | -30 | + | 10.5286 | 9.21E-05 | 0.0819 | CGCTTTGCCGTCG | *folD* | dehydrogenase/methenyltetrahydrofolate cyclohydrolase FolD |
| AVAEIV_RS11960 | 2539988 | 2540000 | 284 | + | 10.5286 | 9.21E-05 | 0.0776 | CGCTTTGCCGTCG | - | tRNA-Arg |
| AVAEIV_RS12910 | 2797028 | 2797040 | 141 | - | 10.5286 | 9.21E-05 | 0.0776 | CGGTCCGCCGGCA | - | FMN-binding glutamate synthase family protein |
| AVAEIV_RS16935 | 3668811 | 3668823 | 314 | - | 10.5286 | 9.21E-05 | 0.0819 | GAGCGCGCCATCG | - | cytochrome c |
| AVAEIV_RS17555 | 3803471 | 3803483 | 271 | + | 10.5286 | 9.21E-05 | 0.0776 | CGGTCCGCCGGCA | - | hybrid sensor histidine kinase/response regulator |
| AVAEIV_RS18200 | 3927565 | 3927577 | 343 | + | 10.5286 | 9.21E-05 | 0.0819 | CTGGCCGCCAGCA | - | CDP-alcohol phosphatidyltransferase family protein |
| AVAEIV_RS18705 | 4029210 | 4029222 | 222 | + | 10.5286 | 9.21E-05 | 0.0819 | CTGGCCGCCAGCA | - | CoA transferase subunit A |
| AVAEIV_RS19520 | 4172196 | 4172208 | 61 | - | 10.5286 | 9.21E-05 | 0.0776 | CCGAGTGCCGCCA | - | hypothetical protein |
| AVAEIV_RS19525 | 4172196 | 4172208 | 352 | - | 10.5286 | 9.21E-05 | 0.0776 | CCGAGTGCCGCCA | - | hypothetical protein |
| AVAEIV_RS21205 | 4507759 | 4507771 | 277 | + | 10.5286 | 9.21E-05 | 0.0819 | CAGATTGCCGCCC | *rimI* | ribosomal protein S18-alanine N-acetyltransferase |
| AVAEIV_RS21730 | 4630561 | 4630573 | 23 | - | 10.5286 | 9.21E-05 | 0.0819 | GAGCGCGCCATCG | - | branched-chain amino acid ABC transporter permease |
| AVAEIV_RS21850 | 4656694 | 4656706 | 15 | - | 10.5286 | 9.21E-05 | 0.0819 | GAGCGCGCCATCG | - | branched-chain amino acid ABC transporter permease |
| AVAEIV_RS24680 | 5244223 | 5244235 | 200 | + | 10.5286 | 9.21E-05 | 0.0776 | CAGATTGCCGCCC | - | OsmC family protein |
| AVAEIV_RS25010 | 5324180 | 5324192 | 159 | - | 10.5286 | 9.21E-05 | 0.0776 | CGACCCGCCACCA | - | Gfo/Idh/MocA family protein |
| AVAEIV_RS25865 | 5546546 | 5546558 | 328 | + | 10.5286 | 9.21E-05 | 0.0776 | CACAGTGCCGGCG | - | ABC transporter permease |
| AVAEIV_RS01035 | 205450 | 205462 | 380 | - | 10.5143 | 9.30E-05 | 0.0819 | CGGTGCGCCGGCG | - | LLM class flavin-dependent oxidoreductase |
| AVAEIV_RS01040 | 205450 | 205462 | 83 | - | 10.5143 | 9.30E-05 | 0.0819 | CGGTGCGCCGGCG | - | dodecin |
| AVAEIV_RS01045 | 205450 | 205462 | -44 | - | 10.5143 | 9.30E-05 | 0.0776 | CGGTGCGCCGGCG | - | hypothetical protein |
| AVAEIV_RS02535 | 509444 | 509456 | 209 | + | 10.5143 | 9.30E-05 | 0.0819 | CAGCTTGCCGTCC | *gshA* | glutamate--cysteine ligase |
| AVAEIV_RS02940 | 608963 | 608975 | 277 | + | 10.5143 | 9.30E-05 | 0.0819 | GGATTCGCCAGCA | - | malate synthase G |
| AVAEIV_RS02945 | 608963 | 608975 | 13 | + | 10.5143 | 9.30E-05 | 0.0776 | GGATTCGCCAGCA | - | glycine cleavage system protein R |
| AVAEIV_RS03445 | 695678 | 695690 | 72 | - | 10.5143 | 9.30E-05 | 0.0819 | CTGGGCGCCAGCG | *accB* | acetyl-CoA carboxylase biotin carboxyl carrier protein |
| AVAEIV_RS04010 | 811220 | 811232 | 265 | + | 10.5143 | 9.30E-05 | 0.0776 | CTGAACGCCAGCT | - | aryl-sulfate sulfotransferase |
| AVAEIV_RS04055 | 821969 | 821981 | 290 | + | 10.5143 | 9.30E-05 | 0.0776 | CCGATCGCCATGC | - | protein |
| AVAEIV_RS05230 | 1074480 | 1074492 | 181 | - | 10.5143 | 9.30E-05 | 0.0776 | CGAAGGGCCATCC | - | IS481-like element ISAzvi11 family transposase |
| AVAEIV_RS08070 | 1672798 | 1672810 | 368 | - | 10.5143 | 9.30E-05 | 0.0776 | CAGTTCGCGACCG | *scpB* | SMC-Scp complex subunit ScpB |
| AVAEIV_RS09365 | 1953622 | 1953634 | 303 | + | 10.5143 | 9.30E-05 | 0.0819 | CCTATCGCCGCCA | - | peptidase U32 family protein |
| AVAEIV_RS09370 | 1953622 | 1953634 | 144 | + | 10.5143 | 9.30E-05 | 0.0776 | CCTATCGCCGCCA | - | HPF/RaiA family ribosome-associated protein |
| AVAEIV_RS09470 | 1973357 | 1973369 | 314 | - | 10.5143 | 9.30E-05 | 0.0776 | CGGCGGGCCATCC | - | HU family DNA-binding protein |
| AVAEIV_RS09625 | 2011400 | 2011412 | 39 | - | 10.5143 | 9.30E-05 | 0.0819 | GACAAGGCCATCG | - | VWA domain-containing protein |
| AVAEIV_RS11240 | 2374887 | 2374899 | 280 | - | 10.5143 | 9.30E-05 | 0.0819 | CTGGGCGCCAGCG | - | ABC transporter ATP-binding protein |
| AVAEIV_RS11390 | 2409598 | 2409610 | 181 | - | 10.5143 | 9.30E-05 | 0.0776 | CGAAGGGCCATCC | - | IS481-like element ISAzvi11 family transposase |

| | | | | | | | | | | |
|---|---|---|---|---|---|---|---|---|---|---|
| AVAEIV_RS12355 | 2612957 | 2612969 | 301 | - | 10.5143 | 9.30E-05 | 0.0776 | CGACGCGCCACCG | - | methyl-accepting chemotaxis protein |
| AVAEIV_RS12715 | 2701634 | 2701646 | 352 | + | 10.5143 | 9.30E-05 | 0.0819 | TGCAACGCCAGCA | - | RNA polymerase factor sigma-70 |
| AVAEIV_RS12720 | 2701634 | 2701646 | -43 | + | 10.5143 | 9.30E-05 | 0.0776 | TGCAACGCCAGCA | - | diaminobutyrate--2-oxoglutarate transaminase |
| AVAEIV_RS13230 | 2869689 | 2869701 | 310 | + | 10.5143 | 9.30E-05 | 0.0776 | CAGATCGTCATCG | - | tyrosine-type recombinase/integrase |
| AVAEIV_RS13475 | 2914957 | 2914969 | 179 | - | 10.5143 | 9.30E-05 | 0.0819 | CAGATCGTCATCG | - | MFS transporter |
| AVAEIV_RS13580 | 2939329 | 2939341 | 257 | + | 10.5143 | 9.30E-05 | 0.0819 | CCGAATGCCAGGA | - | IS3 family transposase |
| AVAEIV_RS13810 | 2991871 | 2991883 | 30 | + | 10.5143 | 9.30E-05 | 0.0819 | CAGCGCGCCACCT | *tssG* | type VI secretion system baseplate subunit TssG |
| AVAEIV_RS14480 | 3156556 | 3156568 | 341 | - | 10.5143 | 9.30E-05 | 0.0776 | CTGGGCGCCAGCG | - | aminotransferase class V-fold PLP-dependent enzyme |
| AVAEIV_RS15745 | 3440015 | 3440027 | 218 | - | 10.5143 | 9.30E-05 | 0.0819 | CTGGGCGCCAGCG | - | tautomerase family protein |
| AVAEIV_RS15990 | 3485154 | 3485166 | 280 | + | 10.5143 | 9.30E-05 | 0.0776 | CTGGGCGCCAGCG | - | ABC transporter ATP-binding protein |
| AVAEIV_RS16055 | 3499977 | 3499989 | 378 | + | 10.5143 | 9.30E-05 | 0.0819 | CTGGGCGCCAGCG | - | LysR family transcriptional regulator |
| AVAEIV_RS16105 | 3510081 | 3510093 | 334 | + | 10.5143 | 9.30E-05 | 0.0776 | GACAAGGCCATCG | - | class II aldolase/adducin family protein |
| AVAEIV_RS18980 | 4087255 | 4087267 | 49 | + | 10.5143 | 9.30E-05 | 0.0819 | CGGCGGGCCATCC | *tilS* | tRNA lysidine(34) synthetase TilS |
| AVAEIV_RS19045 | 4103518 | 4103530 | 75 | - | 10.5143 | 9.30E-05 | 0.0819 | GACAAGGCCATCG | *uppS* | polyprenyl diphosphate synthase |
| AVAEIV_RS19200 | 4137847 | 4137859 | 312 | - | 10.5143 | 9.30E-05 | 0.0819 | CAATGGGCCAGCC | - | superinfection immunity protein |
| AVAEIV_RS19210 | 4137847 | 4137859 | 202 | - | 10.5143 | 9.30E-05 | 0.0776 | CAATGGGCCAGCC | - | XRE family transcriptional regulator |
| AVAEIV_RS22245 | 4747335 | 4747347 | 374 | - | 10.5143 | 9.30E-05 | 0.0776 | CCATTGGCCGTCG | - | aldo/keto reductase |
| AVAEIV_RS22970 | 4919084 | 4919096 | 291 | + | 10.5143 | 9.30E-05 | 0.0776 | CGGCGGGCCATCC | *crp* | cAMP-activated global transcriptional regulator CRP |
| AVAEIV_RS24450 | 5198011 | 5198023 | 293 | + | 10.5143 | 9.30E-05 | 0.0776 | CGCATGGCCTTCA | - | D-hexose-6-phosphate mutarotase |
| AVAEIV_RS24605 | 5228163 | 5228175 | 322 | - | 10.5143 | 9.30E-05 | 0.0819 | CAGCGCGCCACCT | - | acyl-CoA desaturase |
| AVAEIV_RS25240 | 5371662 | 5371674 | 181 | - | 10.5143 | 9.30E-05 | 0.0776 | CGAAGGGCCATCC | - | IS481-like element ISAzvi11 family transposase |
| AVAEIV_RS25900 | 5557089 | 5557101 | 287 | + | 10.5143 | 9.30E-05 | 0.0819 | CGAAGGGCCATCC | - | ABC transporter ATP-binding protein |
| AVAEIV_RS25915 | 5564019 | 5564031 | 290 | + | 10.5143 | 9.30E-05 | 0.0819 | CGAAGGGCCATCC | - | putative virulence factor |
| AVAEIV_RS00565 | 117833 | 117845 | -32 | + | 10.5 | 9.38E-05 | 0.0822 | CAGACGGCTAGCA | - | aminopeptidase |
| AVAEIV_RS00570 | 117833 | 117845 | 233 | + | 10.5 | 9.38E-05 | 0.0778 | CAGACGGCTAGCA | - | hypothetical protein |
| AVAEIV_RS01245 | 244706 | 244718 | 93 | - | 10.5 | 9.38E-05 | 0.0822 | CTGATCGCCGTCC | - | Nif11-like leader peptide family natural precursor |
| AVAEIV_RS03895 | 785543 | 785555 | 387 | - | 10.5 | 9.38E-05 | 0.0822 | CTGATCGCCGTCC | - | hypothetical protein |
| AVAEIV_RS08445 | 1764163 | 1764175 | 350 | - | 10.5 | 9.38E-05 | 0.0778 | CAGCGTGCCGCCA | - | LysR family transcriptional regulator |
| AVAEIV_RS08465 | 1769316 | 1769328 | 158 | - | 10.5 | 9.38E-05 | 0.0778 | CAGTTTGACAGCG | - | heavy metal translocating P-type ATPase |
| AVAEIV_RS09920 | 2078923 | 2078935 | -22 | - | 10.5 | 9.38E-05 | 0.0822 | CTGATCGCCGTCC | - | DUF2489 domain-containing protein |
| AVAEIV_RS10415 | 2176989 | 2177001 | 247 | - | 10.5 | 9.38E-05 | 0.0778 | CGTCAGGCCAGCA | - | protein |
| AVAEIV_RS12095 | 2565577 | 2565589 | 285 | + | 10.5 | 9.38E-05 | 0.0778 | CTGATCGCCGTCC | *speE* | polyamine aminopropyltransferase |
| AVAEIV_RS14310 | 3119270 | 3119282 | 341 | + | 10.5 | 9.38E-05 | 0.0822 | CAGATCGCCACGC | - | NAD-dependent epimerase/dehydratase family protein |
| AVAEIV_RS14760 | 3211112 | 3211124 | 19 | + | 10.5 | 9.38E-05 | 0.0778 | CGTCTGGCCAGCG | *nuoH* | NADH-quinone oxidoreductase subunit NuoH |
| AVAEIV_RS16990 | 3683299 | 3683311 | 164 | + | 10.5 | 9.38E-05 | 0.0822 | CGCTTCGCCATGG | - | LysR family transcriptional regulator |
| AVAEIV_RS16995 | 3683299 | 3683311 | -48 | + | 10.5 | 9.38E-05 | 0.0778 | CGCTTCGCCATGG | - | YbfB/YjiJ family MFS transporter |
| AVAEIV_RS18405 | 3965303 | 3965315 | 364 | - | 10.5 | 9.38E-05 | 0.0822 | CAACTGGGCAGCA | *cysM* | cysteine synthase CysM |
| AVAEIV_RS18410 | 3965303 | 3965315 | -23 | - | 10.5 | 9.38E-05 | 0.0778 | CAACTGGGCAGCA | - | response regulator |
| AVAEIV_RS18685 | 4024560 | 4024572 | 300 | - | 10.5 | 9.38E-05 | 0.0822 | CACCGCGCCACCC | *pcaD* | 3-oxoadipate enol-lactonase |
| AVAEIV_RS20360 | 4331335 | 4331347 | 250 | + | 10.5 | 9.38E-05 | 0.0778 | CTGAATGCCGGCA | - | hypothetical protein |
| AVAEIV_RS21780 | 4638442 | 4638454 | 320 | - | 10.5 | 9.38E-05 | 0.0822 | CCGTGCGCCAGCT | - | ABC transporter permease |
| AVAEIV_RS23745 | 5060381 | 5060393 | 138 | - | 10.5 | 9.38E-05 | 0.0778 | CCGACGGCCGGCA | - | sigma-70 family RNA polymerase sigma factor |
| AVAEIV_RS25790 | 5526120 | 5526132 | 128 | - | 10.5 | 9.38E-05 | 0.0778 | CGGGGCGCCATCC | - | glycoside hydrolase family 32 protein |
| AVAEIV_RS26045 | 5597876 | 5597888 | -49 | - | 10.5 | 9.38E-05 | 0.0822 | CAGCGTGCCGCCA | - | hypothetical protein |
| AVAEIV_RS00750 | 148148 | 148160 | 295 | + | 10.4857 | 9.46E-05 | 0.0779 | GCGCTGGCCAGCA | - | DUF364 domain-containing protein |
| AVAEIV_RS02390 | 475417 | 475429 | 225 | + | 10.4857 | 9.46E-05 | 0.0825 | GCGCTGGCCAGCA | - | ABC-type transport auxiliary lipoprotein family protein |
| AVAEIV_RS02465 | 492552 | 492564 | 20 | - | 10.4857 | 9.46E-05 | 0.0825 | CCGCTCGCCTGCA | - | TolC family protein |
| AVAEIV_RS02780 | 569769 | 569781 | 142 | - | 10.4857 | 9.46E-05 | 0.0779 | GCGCTGGCCAGCA | *epd* | erythrose-4-phosphate dehydrogenase |
| AVAEIV_RS04835 | 982657 | 982669 | 278 | + | 10.4857 | 9.46E-05 | 0.0779 | CTGAGCGCCGCCA | - | LTA synthase family protein |
| AVAEIV_RS07840 | 1632527 | 1632539 | 39 | - | 10.4857 | 9.46E-05 | 0.0779 | CCCTGCGCCAGCC | - | DUF4398 domain-containing protein |
| AVAEIV_RS08570 | 1788616 | 1788628 | 96 | + | 10.4857 | 9.46E-05 | 0.0779 | GCGCTGGCCAGCA | *istA* | IS21 family transposase |
| AVAEIV_RS09470 | 1973375 | 1973387 | 296 | - | 10.4857 | 9.46E-05 | 0.0779 | GCGCTGGCCAGCA | - | HU family DNA-binding protein |

| | | | | | | | | | | |
|---|---|---|---|---|---|---|---|---|---|---|
| AVAEIV_RS09500 | 1978514 | 1978526 | 208 | + | 10.4857 | 9.46E-05 | 0.0779 | CTGAGCGCCGCCA | - | TolC family protein |
| AVAEIV_RS10140 | 2115702 | 2115714 | 352 | - | 10.4857 | 9.46E-05 | 0.0779 | CCGCTCGCCTGCA | *ffs* | signal recognition particle sRNA small type |
| AVAEIV_RS12060 | 2561059 | 2561071 | 24 | - | 10.4857 | 9.46E-05 | 0.0825 | CCCTGCGCCAGCC | - | alpha/beta hydrolase |
| AVAEIV_RS12065 | 2561059 | 2561071 | 75 | - | 10.4857 | 9.46E-05 | 0.0779 | CCCTGCGCCAGCC | *aroC* | chorismate synthase |
| AVAEIV_RS16725 | 3621412 | 3621424 | 240 | - | 10.4857 | 9.46E-05 | 0.0825 | CCGAGGGCCGGCG | - | hypothetical protein |
| AVAEIV_RS17670 | 3826739 | 3826751 | 340 | + | 10.4857 | 9.46E-05 | 0.0825 | CCCTGCGCCAGCC | - | tRNA-Asp |
| AVAEIV_RS17675 | 3826739 | 3826751 | 250 | + | 10.4857 | 9.46E-05 | 0.0825 | CCCTGCGCCAGCC | - | tRNA-Val |
| AVAEIV_RS17735 | 3841173 | 3841185 | 255 | - | 10.4857 | 9.46E-05 | 0.0825 | CCGCGCGCCATGA | *bamC* | outer membrane protein assembly factor BamC |
| AVAEIV_RS18000 | 3893456 | 3893468 | -49 | - | 10.4857 | 9.46E-05 | 0.0825 | CAATTGGGCATCG | - | hypothetical protein |
| AVAEIV_RS18860 | 4062161 | 4062173 | 361 | + | 10.4857 | 9.46E-05 | 0.0825 | CCGCGCGCCATGA | *recA* | recombinase RecA |
| AVAEIV_RS23945 | 5100999 | 5101011 | 131 | - | 10.4857 | 9.46E-05 | 0.0779 | CAGAAGGCCGGCC | - | TIGR02444 family protein |
| AVAEIV_RS24665 | 5243020 | 5243032 | 57 | - | 10.4857 | 9.46E-05 | 0.0825 | CCGAGGGCCGGCG | *gabD* | dehydrogenase |
| AVAEIV_RS24670 | 5243020 | 5243032 | 113 | - | 10.4857 | 9.46E-05 | 0.0779 | CCGAGGGCCGGCG | - | c-type cytochrome |
| AVAEIV_RS25535 | 5455589 | 5455601 | 114 | - | 10.4857 | 9.46E-05 | 0.0825 | GCAATGGCCAGCA | - | containing protein |
| AVAEIV_RS02465 | 492597 | 492609 | 65 | + | 10.4714 | 9.54E-05 | 0.0825 | CTGATCGCCAGGG | - | TolC family protein |
| AVAEIV_RS02750 | 561304 | 561316 | 337 | + | 10.4714 | 9.54E-05 | 0.0779 | CAGCCGGCCGGCA | *nudE* | ADP compounds hydrolase NudE |
| AVAEIV_RS02900 | 596138 | 596150 | -38 | - | 10.4714 | 9.54E-05 | 0.0779 | GCGGTCGCCAGCA | - | substrate binding protein |
| AVAEIV_RS03355 | 676012 | 676024 | 206 | - | 10.4714 | 9.54E-05 | 0.0779 | CGGAGTGCCGCCA | *nusB* | transcription antitermination factor NusB |
| AVAEIV_RS03365 | 677316 | 677328 | 362 | - | 10.4714 | 9.54E-05 | 0.0779 | CGCAGCGCCACCC | - | phosphatidylglycerophosphatase A |
| AVAEIV_RS04540 | 928451 | 928463 | 322 | + | 10.4714 | 9.54E-05 | 0.0779 | CCCTTCGCCGCCG | - | LysR substrate-binding domain-containing protein |
| AVAEIV_RS05335 | 1099224 | 1099236 | 301 | - | 10.4714 | 9.54E-05 | 0.0779 | CTGAGCGGCATCA | *sbcD* | exonuclease subunit SbcD |
| AVAEIV_RS07615 | 1587682 | 1587694 | 334 | + | 10.4714 | 9.54E-05 | 0.0779 | CTGATCGCCAGGG | - | subunit |
| AVAEIV_RS07935 | 1650125 | 1650137 | 208 | - | 10.4714 | 9.54E-05 | 0.0825 | CAGCCGGCCGGCA | *htpX* | protease HtpX |
| AVAEIV_RS08315 | 1730906 | 1730918 | 56 | - | 10.4714 | 9.54E-05 | 0.0779 | CTGATCGCCAGGG | - | hypothetical protein |
| AVAEIV_RS08375 | 1747396 | 1747408 | 227 | - | 10.4714 | 9.54E-05 | 0.0779 | CTGCTCGCCACCC | - | SIMPL domain-containing protein |
| AVAEIV_RS10990 | 2313650 | 2313662 | 156 | - | 10.4714 | 9.54E-05 | 0.0825 | CTGCTCGCCACCC | - | tripartite tricarboxylate transporter permease |
| AVAEIV_RS11910 | 2528969 | 2528981 | 112 | - | 10.4714 | 9.54E-05 | 0.0825 | CGCAGCGCCACCC | - | ANTAR domain-containing response regulator |
| AVAEIV_RS12460 | 2643145 | 2643157 | 301 | - | 10.4714 | 9.54E-05 | 0.0825 | CGCAGCGCCACCC | - | MDR family oxidoreductase |
| AVAEIV_RS13180 | 2859852 | 2859864 | 157 | + | 10.4714 | 9.54E-05 | 0.0825 | CCATTTGCCAGCC | - | hypothetical protein |
| AVAEIV_RS15180 | 3304307 | 3304319 | 43 | + | 10.4714 | 9.54E-05 | 0.0779 | CTGCTCGCCACCC | - | LLM class flavin-dependent oxidoreductase |
| AVAEIV_RS15480 | 3383475 | 3383487 | 231 | - | 10.4714 | 9.54E-05 | 0.0779 | CCCATGGCCATGG | - | sugar transferase |
| AVAEIV_RS15635 | 3425199 | 3425211 | 259 | + | 10.4714 | 9.54E-05 | 0.0825 | CAGCACGGCAGCG | *ccmE* | cytochrome c maturation protein CcmE |
| AVAEIV_RS15640 | 3425199 | 3425211 | 86 | + | 10.4714 | 9.54E-05 | 0.0825 | CAGCACGGCAGCG | *ccmD* | heme exporter protein CcmD |
| AVAEIV_RS17020 | 3690638 | 3690650 | 199 | + | 10.4714 | 9.54E-05 | 0.0825 | CAGCCGGCCGGCA | - | hypothetical protein |
| AVAEIV_RS18900 | 4070024 | 4070036 | 310 | + | 10.4714 | 9.54E-05 | 0.0779 | CTGGAGGCCACCA | *fdxA* | ferredoxin FdxA |
| AVAEIV_RS18930 | 4077046 | 4077058 | 63 | + | 10.4714 | 9.54E-05 | 0.0825 | CAGCCGGCCGGCA | *surE* | 5'/3'-nucleotidase SurE |
| AVAEIV_RS21305 | 4530716 | 4530728 | 10 | + | 10.4714 | 9.54E-05 | 0.0825 | CAGCGCGCCACGA | - | ChaN family lipoprotein |
| AVAEIV_RS21490 | 4573451 | 4573463 | 312 | + | 10.4714 | 9.54E-05 | 0.0825 | CCGTTCGCCTTCG | *carA* | small subunit |
| AVAEIV_RS22875 | 4898885 | 4898897 | 378 | - | 10.4714 | 9.54E-05 | 0.0779 | CCGTTCGCCTTCG | - | nitroreductase family protein |
| AVAEIV_RS23585 | 5026377 | 5026389 | 106 | + | 10.4714 | 9.54E-05 | 0.0779 | CTGCTCGCCACCC | - | symmetrical bis(5'-nucleosyl)-tetraphosphatase |
| AVAEIV_RS23975 | 5106877 | 5106889 | 2 | + | 10.4714 | 9.54E-05 | 0.0825 | CCGTTCGCCTTCG | - | uroporphyrinogen-III synthase |
| AVAEIV_RS24150 | 5135941 | 5135953 | 144 | + | 10.4714 | 9.54E-05 | 0.0825 | CAGCCGGCCGGCA | *alr* | alanine racemase |
| AVAEIV_RS25130 | 5353280 | 5353292 | 185 | + | 10.4714 | 9.54E-05 | 0.0825 | CAGCCGGCCGGCA | - | condensation domain-containing protein |
| AVAEIV_RS00285 | 62181 | 62193 | 255 | + | 10.4571 | 9.63E-05 | 0.0826 | CGCCGTGCCAGCG | - | ABC transporter permease |
| AVAEIV_RS01030 | 202075 | 202087 | 96 | - | 10.4571 | 9.63E-05 | 0.0782 | CAGTTCGCCAGGC | *oadA* | alpha |
| AVAEIV_RS01040 | 205531 | 205543 | 164 | + | 10.4571 | 9.63E-05 | 0.0826 | CCGTGCGCCAGGA | - | dodecin |
| AVAEIV_RS01515 | 292337 | 292349 | 126 | + | 10.4571 | 9.63E-05 | 0.0826 | CCGTCGGCCACCG | - | dihydroorotase |
| AVAEIV_RS09300 | 1942204 | 1942216 | 296 | - | 10.4571 | 9.63E-05 | 0.0826 | CCGTTGGCCACCT | - | DUF3144 domain-containing protein |
| AVAEIV_RS09595 | 1999928 | 1999940 | 82 | + | 10.4571 | 9.63E-05 | 0.0782 | CAGTTGGCCGTCT | *pepN* | aminopeptidase N |
| AVAEIV_RS10305 | 2152510 | 2152522 | 229 | + | 10.4571 | 9.63E-05 | 0.0782 | CAGCGGGCCGGCG | - | pseudouridine synthase |
| AVAEIV_RS11985 | 2542498 | 2542510 | 376 | - | 10.4571 | 9.63E-05 | 0.0782 | CCACCGGCCAGCA | *clpX* | ATP-dependent Clp protease ATP-binding subunit ClpX |
| AVAEIV_RS12995 | 2816332 | 2816344 | 192 | + | 10.4571 | 9.63E-05 | 0.0826 | CAGTCGGCCGTCG | - | aldo/keto reductase |

| | | | | | | | | | | |
|---|---|---|---|---|---|---|---|---|---|---|
| AVAEIV_RS13280 | 2876692 | 2876704 | 125 | + | 10.4571 | 9.63E-05 | 0.0782 | CCGTGCGCCAGGA | - | recombinase family protein |
| AVAEIV_RS15400 | 3363451 | 3363463 | 273 | + | 10.4571 | 9.63E-05 | 0.0782 | CCGGACGCCAGCC | - | low molecular weight protein-tyrosine-phosphatase |
| AVAEIV_RS16640 | 3607486 | 3607498 | 306 | - | 10.4571 | 9.63E-05 | 0.0826 | CCGTGCGCCAGGA | - | YcgL domain-containing protein |
| AVAEIV_RS16695 | 3616290 | 3616302 | 377 | - | 10.4571 | 9.63E-05 | 0.0826 | CAGCGGGCCGGCG | - | cobyric acid synthase |
| AVAEIV_RS17645 | 3821250 | 3821262 | 372 | + | 10.4571 | 9.63E-05 | 0.0782 | CCACCGGCCAGCA | - | carboxy terminal-processing peptidase |
| AVAEIV_RS19005 | 4094812 | 4094824 | 226 | - | 10.4571 | 9.63E-05 | 0.0826 | CAGGCCGCCGGCA | *lpxA* | acyl-ACP--UDP-N-acetylglucosamine O-acyltransferase |
| AVAEIV_RS19175 | 4133127 | 4133139 | 363 | - | 10.4571 | 9.63E-05 | 0.0826 | CTCATGGCCGTCG | - | hypothetical protein |
| AVAEIV_RS19180 | 4133127 | 4133139 | 16 | - | 10.4571 | 9.63E-05 | 0.0826 | CTCATGGCCGTCG | - | hypothetical protein |
| AVAEIV_RS19405 | 4160013 | 4160025 | 381 | - | 10.4571 | 9.63E-05 | 0.0826 | CTAGTGGCCATCG | - | KilA-N domain-containing protein |
| AVAEIV_RS19640 | 4186693 | 4186705 | 90 | - | 10.4571 | 9.63E-05 | 0.0826 | CGGCTTGCCACCC | *tcdA* | synthase TcdA |
| AVAEIV_RS19645 | 4186693 | 4186705 | 118 | - | 10.4571 | 9.63E-05 | 0.0782 | CGGCTTGCCACCC | - | glycosyltransferase |
| AVAEIV_RS20325 | 4329421 | 4329433 | 70 | + | 10.4571 | 9.63E-05 | 0.0826 | CCACCGGCCAGCA | - | addiction module antidote protein |
| AVAEIV_RS20380 | 4334664 | 4334676 | -11 | - | 10.4571 | 9.63E-05 | 0.0782 | GAGTTGGCCACCG | *rpsT* | 30S ribosomal protein S20 |
| AVAEIV_RS24205 | 5146845 | 5146857 | 240 | - | 10.4571 | 9.63E-05 | 0.0826 | CCACCGGCCAGCA | *ftsX* | permease-like cell division protein FtsX |
| AVAEIV_RS24980 | 5316215 | 5316227 | 211 | + | 10.4571 | 9.63E-05 | 0.0782 | CAGGCCGCCGGCA | *iolE* | myo-inosose-2 dehydratase |
| AVAEIV_RS00545 | 111937 | 111949 | 129 | + | 10.4429 | 9.72E-05 | 0.0782 | TACAAGGCCATCG | - | FecR domain-containing protein |
| AVAEIV_RS01145 | 223876 | 223888 | 234 | + | 10.4429 | 9.72E-05 | 0.0782 | CAGGGCGCCGGCG | - | hypothetical protein |
| AVAEIV_RS02040 | 407906 | 407918 | 268 | - | 10.4429 | 9.72E-05 | 0.0826 | CAACAGGCCAGCC | - | Rha family transcriptional regulator |
| AVAEIV_RS02045 | 407906 | 407918 | -37 | - | 10.4429 | 9.72E-05 | 0.0826 | CAACAGGCCAGCC | - | helix-turn-helix domain-containing protein |
| AVAEIV_RS02080 | 413264 | 413276 | 34 | + | 10.4429 | 9.72E-05 | 0.0826 | CAACAGGCCAGCC | - | hypothetical protein |
| AVAEIV_RS02855 | 586808 | 586820 | 127 | - | 10.4429 | 9.72E-05 | 0.0782 | CGTCTCGCCGTCA | - | TetR/AcrR family transcriptional regulator |
| AVAEIV_RS03520 | 709712 | 709724 | 368 | - | 10.4429 | 9.72E-05 | 0.0782 | CCGATGGCGATCG | - | glycoside hydrolase family 5 protein |
| AVAEIV_RS03550 | 715529 | 715541 | 322 | - | 10.4429 | 9.72E-05 | 0.0782 | CCGAAGGCGATCA | - | eCIS core domain-containing protein |
| AVAEIV_RS05380 | 1114621 | 1114633 | 138 | - | 10.4429 | 9.72E-05 | 0.0826 | GACATCGCCGGCA | - | DNA polymerase III subunit chi |
| AVAEIV_RS05465 | 1130351 | 1130363 | 264 | + | 10.4429 | 9.72E-05 | 0.0782 | CCAGCCGCCAGCA | *pilV* | type IV pilus modification protein PilV |
| AVAEIV_RS07530 | 1568773 | 1568785 | -34 | + | 10.4429 | 9.72E-05 | 0.0782 | GACATCGCCGGCA | *nagZ* | beta-N-acetylhexosaminidase |
| AVAEIV_RS07625 | 1589587 | 1589599 | 361 | + | 10.4429 | 9.72E-05 | 0.0782 | CCGATGGCGATCG | - | ComEC/Rec2 |
| AVAEIV_RS10780 | 2259212 | 2259224 | 151 | - | 10.4429 | 9.72E-05 | 0.0826 | CCAGCCGCCAGCA | - | Card1-like endonuclease domain-containing protein |
| AVAEIV_RS11570 | 2447328 | 2447340 | 133 | + | 10.4429 | 9.72E-05 | 0.0782 | CCCTGGGCCGTCA | - | FeoC-like transcriptional regulator |
| AVAEIV_RS11575 | 2447328 | 2447340 | 370 | + | 10.4429 | 9.72E-05 | 0.0782 | CCCTGGGCCGTCA | - | hypothetical protein |
| AVAEIV_RS12470 | 2645955 | 2645967 | 66 | + | 10.4429 | 9.72E-05 | 0.0826 | CTGTGCGCCGGCA | - | protein |
| AVAEIV_RS12845 | 2771316 | 2771328 | 339 | - | 10.4429 | 9.72E-05 | 0.0826 | CTGTGCGCCGGCA | - | transposase |
| AVAEIV_RS13780 | 2981970 | 2981982 | 178 | + | 10.4429 | 9.72E-05 | 0.0826 | CAGGGCGCCGGCG | - | DUF3540 domain-containing protein |
| AVAEIV_RS13955 | 3027582 | 3027594 | -37 | + | 10.4429 | 9.72E-05 | 0.0782 | GACATCGCCGGCA | - | purine-cytosine permease family protein |
| AVAEIV_RS15045 | 3277252 | 3277264 | 219 | + | 10.4429 | 9.72E-05 | 0.0826 | CCACGGGCCAGCG | - | hypothetical protein |
| AVAEIV_RS15385 | 3359755 | 3359767 | 322 | - | 10.4429 | 9.72E-05 | 0.0782 | CAGGGCGCCGGCG | - | PA1571 family protein |
| AVAEIV_RS15705 | 3433753 | 3433765 | 237 | - | 10.4429 | 9.72E-05 | 0.0826 | CCGAAGGCGATCA | - | sulfonate ABC transporter substrate-binding protein |
| AVAEIV_RS16355 | 3553072 | 3553084 | 206 | + | 10.4429 | 9.72E-05 | 0.0826 | GACATCGCCGGCA | - | helix-turn-helix domain-containing protein |
| AVAEIV_RS16390 | 3560259 | 3560271 | 120 | + | 10.4429 | 9.72E-05 | 0.0826 | CCCACCGCCGGCA | - | holin family protein |
| AVAEIV_RS16700 | 3617137 | 3617149 | 166 | + | 10.4429 | 9.72E-05 | 0.0826 | CAGGGCGCCGGCG | *cobD* | threonine-phosphate decarboxylase CobD |
| AVAEIV_RS17205 | 3731387 | 3731399 | 265 | - | 10.4429 | 9.72E-05 | 0.0826 | GACATCGCCGGCA | *asd* | aspartate-semialdehyde dehydrogenase |
| AVAEIV_RS17215 | 3733101 | 3733113 | 100 | - | 10.4429 | 9.72E-05 | 0.0826 | CAGGGCGCCGGCG | *leuD* | 3-isopropylmalate dehydratase small subunit |
| AVAEIV_RS18445 | 3972542 | 3972554 | -36 | - | 10.4429 | 9.72E-05 | 0.0782 | CAGGGCGCCGGCG | - | hypothetical protein |
| AVAEIV_RS19075 | 4107726 | 4107738 | 126 | + | 10.4429 | 9.72E-05 | 0.0782 | TGGACCGCCATCA | - | [protein-PII] uridylyltransferase |
| AVAEIV_RS20915 | 4446263 | 4446275 | 334 | + | 10.4429 | 9.72E-05 | 0.0782 | CACCTGGCCATGG | - | acetyltransferase subunit |
| AVAEIV_RS21845 | 4655517 | 4655529 | -42 | + | 10.4429 | 9.72E-05 | 0.0826 | CAGGGCGCCGGCG | - | ABC transporter substrate-binding protein |
| AVAEIV_RS22320 | 4766399 | 4766411 | 244 | + | 10.4429 | 9.72E-05 | 0.0826 | CGGCACGCCGGCG | - | hypothetical protein |
| AVAEIV_RS22505 | 4814444 | 4814456 | 324 | + | 10.4429 | 9.72E-05 | 0.0826 | CCGAAGGCGATCA | *hemE* | uroporphyrinogen decarboxylase |
| AVAEIV_RS23690 | 5047219 | 5047231 | 111 | - | 10.4429 | 9.72E-05 | 0.0782 | CAGGGCGCCGGCG | - | TPM domain-containing protein |
| AVAEIV_RS24890 | 5296321 | 5296333 | 219 | + | 10.4429 | 9.72E-05 | 0.0826 | CGGAACGGCAGCG | - | OprD family porin |
| AVAEIV_RS02220 | 443328 | 443340 | 378 | + | 10.4286 | 9.80E-05 | 0.0782 | CCGATCGCCGCCC | - | Ni/Fe hydrogenase subunit alpha |
| AVAEIV_RS02265 | 454453 | 454465 | 133 | - | 10.4286 | 9.80E-05 | 0.0831 | CGCTTCGCTATCG | - | 4Fe-4S dicluster domain-containing protein |

| | | | | | | | | | | |
|---|---|---|---|---|---|---|---|---|---|---|
| AVAEIV_RS02610 | 529368 | 529380 | -38 | + | 10.4286 | 9.80E-05 | 0.0831 | GGAATGGCCAGCA | - | OprD family porin |
| AVAEIV_RS03730 | 750943 | 750955 | 72 | + | 10.4286 | 9.80E-05 | 0.0782 | GGGCTGGCCAGCA | - | ATP phosphoribosyltransferase regulatory subunit |
| AVAEIV_RS04690 | 956017 | 956029 | 364 | - | 10.4286 | 9.80E-05 | 0.0782 | CAGTTCGCCAGTC | - | PepSY domain-containing protein |
| AVAEIV_RS07395 | 1534242 | 1534254 | 314 | - | 10.4286 | 9.80E-05 | 0.0782 | CGCTGCGCCAGCC | - | feruloyl-CoA synthase |
| AVAEIV_RS08030 | 1668752 | 1668764 | 343 | + | 10.4286 | 9.80E-05 | 0.0831 | CGGCTCGCCTGCA | - | sensor histidine kinase |
| AVAEIV_RS11935 | 2535412 | 2535424 | 50 | + | 10.4286 | 9.80E-05 | 0.0831 | CACGTCGCCATGG | *lpxH* | UDP-2,3-diacylglucosamine diphosphatase |
| AVAEIV_RS20970 | 4461046 | 4461058 | 310 | - | 10.4286 | 9.80E-05 | 0.0831 | CCGATCGCCGCCC | - | efflux RND transporter permease subunit |
| AVAEIV_RS22110 | 4718243 | 4718255 | 219 | - | 10.4286 | 9.80E-05 | 0.0782 | GGAATGGCCAGCA | *lptG* | LPS export ABC transporter permease LptG |
| AVAEIV_RS22365 | 4776831 | 4776843 | 152 | + | 10.4286 | 9.80E-05 | 0.0831 | CCGATCGCCGCCC | - | branched-chain amino acid transaminase |
| AVAEIV_RS23565 | 5022571 | 5022583 | 255 | + | 10.4286 | 9.80E-05 | 0.0782 | TGGAGCGCCATCG | - | peptidylprolyl isomerase |
| AVAEIV_RS25025 | 5327268 | 5327280 | 196 | + | 10.4286 | 9.80E-05 | 0.0782 | CAGATGGCGACCG | - | sugar ABC transporter substrate-binding protein |
| AVAEIV_RS00135 | 28833 | 28845 | -35 | - | 10.4143 | 9.89E-05 | 0.0782 | CGCCTCGGCAGCA | - | copper resistance protein B |
| AVAEIV_RS02210 | 441763 | 441775 | 351 | + | 10.4143 | 9.89E-05 | 0.0782 | CGCTTCGCCGCCG | - | FAD/NAD(P)-binding protein |
| AVAEIV_RS02775 | 567437 | 567449 | 356 | + | 10.4143 | 9.89E-05 | 0.0782 | TCGCTGGCCAGCA | *tkt* | transketolase |
| AVAEIV_RS02850 | 585570 | 585582 | 77 | - | 10.4143 | 9.89E-05 | 0.0782 | CCGCTTGCCGGCG | *ald* | alanine dehydrogenase |
| AVAEIV_RS04080 | 828412 | 828424 | -34 | + | 10.4143 | 9.89E-05 | 0.0782 | CAGCTGGCGATCG | *rsfS* | ribosome silencing factor |
| AVAEIV_RS04085 | 828412 | 828424 | 327 | + | 10.4143 | 9.89E-05 | 0.0782 | CAGCTGGCGATCG | *rlmH* | methyltransferase RlmH |
| AVAEIV_RS04585 | 938599 | 938611 | 353 | - | 10.4143 | 9.89E-05 | 0.0833 | CGCTTCGCCGCCG | - | urease subunit gamma |
| AVAEIV_RS05680 | 1164518 | 1164530 | 274 | + | 10.4143 | 9.89E-05 | 0.0782 | CAGGTGGCTATCG | - | hypothetical protein |
| AVAEIV_RS06385 | 1302492 | 1302504 | 249 | - | 10.4143 | 9.89E-05 | 0.0833 | CCGAATGGCAGCA | - | TatD family hydrolase |
| AVAEIV_RS07160 | 1475042 | 1475054 | 63 | + | 10.4143 | 9.89E-05 | 0.0782 | CAGGGGGCCAGCC | *era* | GTPase Era |
| AVAEIV_RS07445 | 1548793 | 1548805 | 375 | - | 10.4143 | 9.89E-05 | 0.0782 | TCGCTGGCCAGCA | - | efflux RND transporter permease subunit |
| AVAEIV_RS07580 | 1581006 | 1581018 | 46 | - | 10.4143 | 9.89E-05 | 0.0782 | CCGAGGGACATCA | *nqrF* | subunit F |
| AVAEIV_RS12805 | 2722230 | 2722242 | 365 | + | 10.4143 | 9.89E-05 | 0.0833 | CGCTTCGCCGCCG | - | IS4 family transposase |
| AVAEIV_RS13495 | 2918050 | 2918062 | -31 | + | 10.4143 | 9.89E-05 | 0.0782 | CGCATGGCCATGG | - | 3-hydroxyacyl-CoA dehydrogenase |
| AVAEIV_RS13800 | 2988246 | 2988258 | 103 | + | 10.4143 | 9.89E-05 | 0.0833 | CGCTTCGCCGCCG | - | type VI secretion system Vgr family protein |
| AVAEIV_RS14190 | 3090038 | 3090050 | 199 | + | 10.4143 | 9.89E-05 | 0.0782 | CGCTTCGCCGCCG | - | DUF2970 domain-containing protein |
| AVAEIV_RS14925 | 3255087 | 3255099 | 73 | - | 10.4143 | 9.89E-05 | 0.0833 | CGGCAGGCCAGCC | - | GNAT family N-acetyltransferase |
| AVAEIV_RS14930 | 3255087 | 3255099 | 50 | - | 10.4143 | 9.89E-05 | 0.0782 | CGGCAGGCCAGCC | *xthA* | exodeoxyribonuclease III |
| AVAEIV_RS15275 | 3335306 | 3335318 | 64 | + | 10.4143 | 9.89E-05 | 0.0782 | CGGTTCGCCTTCG | - | extracellular solute-binding protein |
| AVAEIV_RS16765 | 3629381 | 3629393 | 74 | - | 10.4143 | 9.89E-05 | 0.0833 | CGCTTCGCCGCCG | - | 3-deoxy-7-phosphoheptulonate synthase |
| AVAEIV_RS17210 | 3732379 | 3732391 | 124 | - | 10.4143 | 9.89E-05 | 0.0833 | CGGCAGGCCAGCC | *leuB* | 3-isopropylmalate dehydrogenase |
| AVAEIV_RS18075 | 3906641 | 3906653 | 161 | + | 10.4143 | 9.89E-05 | 0.0833 | CAGTTGGCCGTGA | *queC* | 7-cyano-7-deazaguanine synthase QueC |
| AVAEIV_RS18445 | 3972138 | 3972150 | 368 | + | 10.4143 | 9.89E-05 | 0.0782 | CAGCAGGCGATCA | - | hypothetical protein |
| AVAEIV_RS19655 | 4190630 | 4190642 | 51 | - | 10.4143 | 9.89E-05 | 0.0782 | TCGCTGGCCAGCA | - | putative RNA methyltransferase |
| AVAEIV_RS20160 | 4306314 | 4306326 | 266 | - | 10.4143 | 9.89E-05 | 0.0833 | CGCTTCGCCGCCG | *ispG* | diphosphate synthase |
| AVAEIV_RS20230 | 4317065 | 4317077 | 322 | + | 10.4143 | 9.89E-05 | 0.0833 | CATCACGCCATCC | *trmJ* | methyltransferase TrmJ |
| AVAEIV_RS20450 | 4345909 | 4345921 | 195 | + | 10.4143 | 9.89E-05 | 0.0782 | CGCTTCGCCGCCG | - | DUF4105 domain-containing protein |
| AVAEIV_RS21975 | 4687468 | 4687480 | 324 | - | 10.4143 | 9.89E-05 | 0.0782 | TCGCTGGCCAGCA | *fabI* | enoyl-ACP reductase FabI |
| AVAEIV_RS22925 | 4911236 | 4911248 | 130 | - | 10.4143 | 9.89E-05 | 0.0782 | GGGTTGGCCATCG | - | anhydro-N-acetylmuramic acid kinase |
| AVAEIV_RS23070 | 4935021 | 4935033 | 162 | + | 10.4143 | 9.89E-05 | 0.0782 | GCCTTCGCCATCG | - | phage virion morphogenesis protein |
| AVAEIV_RS24025 | 5114066 | 5114078 | 82 | - | 10.4143 | 9.89E-05 | 0.0782 | CCGCTCGCCGTCC | *dapF* | diaminopimelate epimerase |
| AVAEIV_RS24155 | 5136576 | 5136588 | 366 | - | 10.4143 | 9.89E-05 | 0.0833 | AAGGTCGCCATCA | - | RidA family protein |
| AVAEIV_RS24260 | 5158862 | 5158874 | 305 | + | 10.4143 | 9.89E-05 | 0.0833 | CGCTTCGCCGCCG | - | sulfite exporter TauE/SafE family protein |
| AVAEIV_RS24765 | 5262733 | 5262745 | 354 | + | 10.4143 | 9.89E-05 | 0.0782 | CACACCGGCAGCA | - | OprD family porin |
| AVAEIV_RS25675 | 5490399 | 5490411 | 254 | + | 10.4143 | 9.89E-05 | 0.0782 | CAGCAGGCGATCA | - | efflux RND transporter permease subunit |
| AVAEIV_RS00350 | 75496 | 75508 | 219 | + | 10.4 | 9.97E-05 | 0.0836 | CCCTCCGCCACCG | - | SMR family transporter |
| AVAEIV_RS01500 | 289114 | 289126 | 344 | - | 10.4 | 9.97E-05 | 0.0836 | CACCGCGCCGGCG | - | YggS family pyridoxal phosphate-dependent enzyme |
| AVAEIV_RS03125 | 642624 | 642636 | 22 | + | 10.4 | 9.97E-05 | 0.0784 | GATTTCGCCAGCA | *rplB* | 50S ribosomal protein L2 |
| AVAEIV_RS03385 | 681152 | 681164 | 74 | + | 10.4 | 9.97E-05 | 0.0836 | CAGGTCGCGATCG | - | termination factor Rho |
| AVAEIV_RS03390 | 681152 | 681164 | 44 | + | 10.4 | 9.97E-05 | 0.0784 | CAGGTCGCGATCG | - | esterase/lipase family protein |
| AVAEIV_RS03830 | 772441 | 772453 | 286 | - | 10.4 | 9.97E-05 | 0.0836 | CACCGCGCCGGCG | *trmA* | tRNA (uridine(54)-C5)-methyltransferase TrmA |

| | | | | | | | | | | |
|---|---|---|---|---|---|---|---|---|---|---|
| AVAEIV_RS04050 | 821340 | 821352 | 193 | + | 10.4 | 9.97E-05 | 0.0784 | CAGGAGGCCGCCA | - | LrgB family protein |
| AVAEIV_RS05845 | 1196893 | 1196905 | -11 | - | 10.4 | 9.97E-05 | 0.0784 | AGGATGGCCATCA | - | family protein |
| AVAEIV_RS08870 | 1842323 | 1842335 | 170 | - | 10.4 | 9.97E-05 | 0.0784 | TCGGTCGCCAGCA | *cas7e* | type I-E CRISPR-associated protein Cas7/Cse4/CasC |
| AVAEIV_RS09915 | 2078508 | 2078520 | 136 | + | 10.4 | 9.97E-05 | 0.0836 | CCTTCCGCCAGCA | - | penicillin acylase family protein |
| AVAEIV_RS13145 | 2851740 | 2851752 | 37 | + | 10.4 | 9.97E-05 | 0.0836 | CAGCTCGCCGCCC | *hutG* | N-formylglutamate deformylase |
| AVAEIV_RS14170 | 3083977 | 3083989 | 162 | + | 10.4 | 9.97E-05 | 0.0836 | CCAAGCGCCGCCA | - | DUF2149 domain-containing protein |
| AVAEIV_RS15515 | 3393603 | 3393615 | -40 | + | 10.4 | 9.97E-05 | 0.0784 | CACAGCGGCAGCG | - | polysaccharide biosynthesis/export family protein |
| AVAEIV_RS18650 | 4015213 | 4015225 | 242 | + | 10.4 | 9.97E-05 | 0.0836 | CAGCTCGCCGCCC | - | TetR/AcrR family transcriptional regulator |
| AVAEIV_RS19805 | 4221918 | 4221930 | 291 | + | 10.4 | 9.97E-05 | 0.0836 | CAGGTGGCCGCCG | - | homoserine dehydrogenase |
| AVAEIV_RS20030 | 4270916 | 4270928 | 257 | + | 10.4 | 9.97E-05 | 0.0784 | CGCGATGCCACCA | - | putative quinol monooxygenase |
| AVAEIV_RS21555 | 4585386 | 4585398 | 118 | + | 10.4 | 9.97E-05 | 0.0784 | CCCTTCGCCACCT | - | (Fe-S)-binding protein |
| AVAEIV_RS23255 | 4959761 | 4959773 | 211 | + | 10.4 | 9.97E-05 | 0.0784 | CGGGACGCCAGCC | - | DedA family protein |
| AVAEIV_RS23665 | 5044343 | 5044355 | 260 | + | 10.4 | 9.97E-05 | 0.0784 | CGCTTCGGCATCG | - | Crp/Fnr family transcriptional regulator |

**S4 Table. Genes differentially expressed in the *amrZ* mutant when compared to the wt strain AEIV, based on the RNAseq data**

| | | | | | | | | | | CORRESPONDANCE TO DJ STRAIN | | | |
|---|---|---|---|---|---|---|---|---|---|---|---|---|---|
| locus_tag | baseMean | log2FoldChange | lfcSE | stat | pvalue | padj | GENE | PRODUCT | protein_ID | locus_tag DJ | GENE DJ | PRODUCT DJ | IDENTITY |
| AVAEIV_RS11870 | 4067.373106 | -4.255542246 | 0.254156482 | -16.74378797 | 6.29E-63 | 2.31E-60 | *nirB* | nitrite reductase large subunit NirB | WP_061288356.1 | Avin_23370 | *nasA* | Assimilatory Nitrite reductase,large subunit; NasA | 99.88 |
| AVAEIV_RS02310 | 2301.014793 | -3.954703106 | 0.355275555 | -11.13136846 | 8.83E-29 | 1.59E-27 | - | hypothetical protein | WP_012699141.1 | Avin_04580 | | hypothetical protein | 100 |
| AVAEIV_RS21140 | 7954.753969 | -3.825567656 | 0.413808349 | -9.244781221 | 2.36E-20 | 1.75E-19 | - | IS630 family transposase | WP_302229235.1 | | | | |
| AVAEIV_RS16845 | 2756.561539 | -3.806804816 | 0.41228466 | -9.233437931 | 2.62E-20 | 1.93E-19 | - | hypothetical protein | WP_012701876.1 | Avin_33410 | | conserved hypothetical protein | 100 |
| AVAEIV_RS05080 | 3100.557093 | -3.795755619 | 0.212427321 | -17.86849071 | 2.08E-71 | 1.07E-68 | - | nucleotide sugar dehydrogenase | WP_012699749.1 | Avin_10970 | *algD* | GDP-mannose 6-dehydrogenase | 100 |
| AVAEIV_RS12030 | 1905.194267 | -3.739095336 | 0.486502794 | -7.685660564 | 1.52E-14 | 6.08E-14 | *phaR* | polyhydroxyalkanoate synthesis repressor PhaR | WP_012700954.1 | Avin_23680 | *phbF* | polyhydroxyalkanoate synthesis regulator | 100 |
| AVAEIV_RS02930 | 3636.702576 | -3.637080007 | 0.538402544 | -6.755317279 | 1.43E-11 | 4.22E-11 | - | RNA-guided endonuclease InsQ/TnpB family protein | WP_302228103.1 | Avin_05900 | | transposase IS891/IS1136/IS1341 | 99.48 |
| AVAEIV_RS15235 | 10046.80096 | -3.60048672 | 0.346873146 | -10.37983701 | 3.06E-25 | 3.94E-24 | - | type I polyketide synthase | WP_061288661.1 | Avin_29560 | *arsA* | Type I fatty acid synthase ArsA | 99.84 |
| AVAEIV_RS20540 | 2457.424517 | -3.589563609 | 0.508343882 | -7.061290079 | 1.65E-12 | 5.39E-12 | - | FKBP-type peptidyl-prolyl cis-trans isomerase | WP_012702617.1 | Avin_41110 | | Peptidylprolyl isomerase, FKBP-type | 100 |
| AVAEIV_RS16385 | 5040.053448 | -3.561830265 | 0.546723207 | -6.514869357 | 7.28E-11 | 2.03E-10 | - | hypothetical protein | WP_302226003.1 | Avin_32050 | | Conserved hypothetical protein | 99.23 |
| AVAEIV_RS13180 | 1216.122827 | -3.556710251 | 0.229620945 | -15.48948526 | 4.09E-54 | 7.00E-52 | - | hypothetical protein | WP_231876346.1 | Avin_26230 | | hypothetical protein | 100 |
| AVAEIV_RS15210 | 1310.063468 | -3.535328399 | 0.223247021 | -15.83594882 | 1.76E-56 | 3.61E-54 | - | ABC transporter permease | WP_012701504.1 | Avin_29500 | | conserved hypothetical protein | 100 |
| AVAEIV_RS18560 | 2100.010594 | -3.452314463 | 0.559293422 | -6.172635561 | 6.72E-10 | 1.72E-09 | - | hypothetical protein | WP_041807600.1 | | | | |
| AVAEIV_RS12310 | 345.2745823 | -3.424890065 | 0.269049054 | -12.72961198 | 4.05E-37 | 1.72E-35 | *fliT* | flagellar protein FliT | WP_012701009.1 | Avin_24270 | *fliT* | flagellar protein | 100 |
| AVAEIV_RS19665 | 10412.95213 | -3.384060966 | 0.682998723 | -4.95471053 | 7.24E-07 | 1.41E-06 | - | cold-shock protein | WP_012702433.1 | Avin_39180 | *cspA* | Cold-shock-like protein | 100 |
| AVAEIV_RS09215 | 14357.14841 | -3.38355425 | 0.726843578 | -4.655133998 | 3.24E-06 | 5.96E-06 | - | cold-shock protein | WP_012700418.1 | Avin_17950 | | Cold-shock DNA-binding domain-containing protein | 100 |
| AVAEIV_RS11865 | 297.544364 | -3.37146651 | 0.163877794 | -20.57305284 | 4.79E-94 | 2.46E-90 | *nirD* | nitrite reductase small subunit NirD | WP_012700922.1 | Avin_23360 | | assimilatory nitrite reductase, small subunit | 100 |
| AVAEIV_RS15205 | 1331.726398 | -3.353578442 | 0.173265851 | -19.35510331 | 1.85E-83 | 1.90E-80 | - | outer membrane lipoprotein-sorting protein | WP_012701503.1 | Avin_29490 | | hypothetical protein | 100 |
| AVAEIV_RS07695 | 216.4333835 | -3.312015411 | 0.449541058 | -7.367548195 | 1.74E-13 | 6.31E-13 | *rpmF* | 50S ribosomal protein L32 | WP_007160601.1 | Avin_14890 | *rpmF* | 50S ribosomal protein L32 | 100 |
| AVAEIV_RS20645 | 1026.397691 | -3.303144347 | 0.427495412 | -7.726736376 | 1.10E-14 | 4.47E-14 | - | hypothetical protein | WP_012702637.1 | Avin_41310 | | conserved hypothetical protein | 100 |
| AVAEIV_RS17290 | 920.7165075 | -3.22291239 | 0.339727115 | -9.486768189 | 2.38E-21 | 2.02E-20 | - | Arc family DNA-binding protein | WP_041807311.1 | Avin_34410 | *algZ* | AlgZ-like DNA binding protein | 100 |
| AVAEIV_RS00275 | 856.2938141 | -3.163429069 | 0.304420191 | -10.39165327 | 2.71E-25 | 3.49E-24 | - | hypothetical protein | WP_012698752.1 | Avin_00570 | | hypothetical protein | 100 |
| AVAEIV_RS07115 | 2376.323755 | -3.146462211 | 0.426135254 | -7.383717219 | 1.54E-13 | 5.60E-13 | *rpoE* | RNA polymerase sigma factor RpoE | WP_061289897.1 | | | | |
| AVAEIV_RS09915 | 6529.57865 | -3.110765325 | 0.389945763 | -7.977430753 | 1.49E-15 | 6.59E-15 | - | penicillin acylase family protein | WP_302228733.1 | Avin_19340 | | Peptidase S45, penicillin amidase family | 99.4 |
| AVAEIV_RS07120 | 1768.643419 | -3.081146112 | 0.334050434 | -9.223595599 | 2.87E-20 | 2.09E-19 | - | sigma-E factor negative regulatory protein | WP_012700008.1 | Avin_13700 | *mucA* | sigma factor algU negative regulatory protein MucA | 100 |
| AVAEIV_RS05405 | 8016.550148 | -3.061899401 | 0.704883799 | -4.343835688 | 1.40E-05 | 2.44E-05 | - | cold-shock protein | WP_012699820.1 | Avin_11700 | | Cold shock domain family protein | 100 |
| AVAEIV_RS08170 | 8719.269141 | -3.050867932 | 0.55449784 | -5.502037539 | 3.75E-08 | 8.18E-08 | *rpsA* | 30S ribosomal protein S1 | WP_012700217.1 | Avin_15870 | *rpsA* | 30S ribosomal protein S1 | 100 |
| AVAEIV_RS03215 | 993.7187205 | -3.012772838 | 0.667597569 | -4.512857711 | 6.40E-06 | 1.15E-05 | *rpmJ* | 50S ribosomal protein L36 | WP_012699323.1 | Avin_06460 | *rpmJ* | 50S ribosomal protein L36 | 100 |
| AVAEIV_RS15230 | 1180.548376 | -3.011821505 | 0.161285955 | -18.67379898 | 8.09E-78 | 5.20E-75 | - | type III polyketide synthase | WP_012701508.1 | Avin_29550 | *arsB* | Type III polyketide synthase | 100 |
| AVAEIV_RS20195 | 1678.305431 | -3.009915856 | 0.154580914 | -19.47145858 | 1.92E-84 | 2.46E-81 | *hscA* | Fe-S protein assembly chaperone HscA | WP_302226983.1 | Avin_40360 | *hscA* | Fe-S protein assembly chaperone HscA | 99.68 |
| AVAEIV_RS15225 | 1028.052798 | -2.968346058 | 0.191375908 | -15.51055249 | 2.94E-54 | 5.22E-52 | - | 3-oxoacyl-[acyl-carrier-protein] synthase III C-terminal domain-containing protein | WP_061288656.1 | Avin_29530 | *arsC* | Type III polyketide synthase | 99.76 |
| AVAEIV_RS03050 | 4608.010719 | -2.962772128 | 0.584684786 | -5.067298136 | 4.04E-07 | 8.05E-07 | *nusG* | transcription termination/antitermination protein NusG | WP_039806181.1 | Avin_06110 | *nusG* | transcription termination/antitermination factor NusG | 100 |
| AVAEIV_RS06825 | 2134.987265 | -2.947224327 | 0.503229046 | -5.856626026 | 4.72E-09 | 1.12E-08 | *rpsI* | 30S ribosomal protein S9 | WP_012699951.1 | Avin_13050 | *rpsI* | 30S ribosomal protein S9 | 100 |
| AVAEIV_RS08510 | 1300.828802 | -2.938340085 | 0.286296406 | -10.26327969 | 1.03E-24 | 1.27E-23 | - | TPM domain-containing protein | WP_302228572.1 | | | | |
| AVAEIV_RS22530 | 1189.704199 | -2.913311165 | 0.38455297 | -7.575838426 | 3.57E-14 | 1.37E-13 | *aroK* | shikimate kinase AroK | WP_012703006.1 | Avin_45250 | *aroK* | shikimate kinase | 100 |
| AVAEIV_RS02255 | 561.864639 | -2.908894217 | 0.175321904 | -16.59173297 | 8.00E-62 | 2.42E-59 | - | NAD(P)/FAD-dependent oxidoreductase | WP_302228012.1 | Avin_04480 | | FAD-dependent pyridine nucleotide-disulphide oxidoreductase | 99.76 |
| AVAEIV_RS01115 | 5845.042107 | -2.895282927 | 0.555823232 | -5.208999483 | 1.90E-07 | 3.92E-07 | - | hypothetical protein | WP_302227906.1 | Avin_02260 | | hypothetical protein | 98.88 |
| AVAEIV_RS12195 | 836.9940843 | -2.884186332 | 0.261715258 | -11.02032167 | 3.05E-28 | 5.14E-27 | - | flagellar basal body rod protein FlgF | WP_012700986.1 | Avin_24040 | *flgF* | flagellar basal-body rod protein | 100 |
| AVAEIV_RS18795 | 1734.364492 | -2.876362803 | 0.311586191 | -9.2313552 | 2.67E-20 | 1.96E-19 | - | sensor domain-containing diguanylate cyclase | WP_012702359.1 | Avin_38420 | | GGDEF Response Regulator | 100 |
| AVAEIV_RS03380 | 527.5701006 | -2.870564335 | 0.183718367 | -15.6248087 | 4.93E-55 | 9.76E-53 | - | cobalamin-binding protein | WP_302228141.1 | Avin_06810 | | Periplasmic binding protein | 99.29 |
| AVAEIV_RS11705 | 1774.726827 | -2.865807729 | 0.58036971 | -4.937900239 | 7.90E-07 | 1.54E-06 | - | TatD family hydrolase | WP_012700886.1 | Avin_23000 | | TatD-related deoxyribonuclease protein | 100 |
| AVAEIV_RS05145 | 2485.546849 | -2.863096274 | 0.284836464 | -10.05171963 | 9.03E-24 | 1.00E-22 | - | sigma-54-dependent transcriptional regulator | WP_012699764.1 | Avin_11120 | *algB* | Response regulator, sigma54-dependent regulatory protein; AlgB | 100 |
| AVAEIV_RS19660 | 339.7711958 | -2.862342344 | 0.158026381 | -18.11306643 | 2.51E-73 | 1.44E-70 | - | hypothetical protein | WP_061288074.1 | Avin_39170 | | conserved hypothetical protein | 99.19 |
| AVAEIV_RS20190 | 236.3851565 | -2.826902999 | 0.159770475 | -17.69352563 | 4.70E-70 | 2.20E-67 | *fdx* | ISC system 2Fe-2S type ferredoxin | WP_041807608.1 | Avin_40350 | *fdx* | Isc ferredoxin | 100 |
| AVAEIV_RS20215 | 1949.242982 | -2.826631484 | 0.320807974 | -8.810976379 | 1.24E-18 | 7.48E-18 | - | IscS subfamily cysteine desulfurase | WP_012702551.1 | Avin_40400 | *iscS* | Cysteine desulfurase IscS | 100 |
| AVAEIV_RS17785 | 1339.63271 | -2.816096401 | 0.264179094 | -10.65980035 | 1.57E-26 | 2.25E-25 | - | hypothetical protein | WP_061288891.1 | Avin_35450 | | hypothetical protein | 100 |
| AVAEIV_RS22085 | 1951.864219 | -2.815798136 | 0.286170795 | -9.839571978 | 7.60E-23 | 7.52E-22 | - | fatty acyl-AMP ligase | WP_197471057.1 | | | | |
| AVAEIV_RS00595 | 1913.781583 | -2.799943249 | 0.857059574 | -3.266917884 | 0.001087252 | 0.001573195 | - | GlsB/YeaQ/YmgE family stress response membrane protein | WP_012698816.1 | Avin_01210 | | conserved hypothetical protein | 100 |
| AVAEIV_RS02720 | 229.3993749 | -2.777242944 | 0.216662931 | -12.81826537 | 1.30E-37 | 5.69E-36 | - | hypothetical protein | pseudo=true | | | | |
| AVAEIV_RS03045 | 6629.172389 | -2.77538392 | 0.598266771 | -4.639040729 | 3.50E-06 | 6.43E-06 | *secE* | preprotein translocase subunit SecE | WP_041806885.1 | Avin_06100 | *secE* | preprotein translocase, SecE subunit | 98.8 |
| AVAEIV_RS22070 | 969.3733161 | -2.769789378 | 0.219486526 | -12.61940505 | 1.65E-36 | 6.79E-35 | *spt* | serine palmitoyltransferase | WP_302227354.1 | Avin_44240 | | 8-amino-7-oxononanoate synthase-like protein | 99.49 |
| AVAEIV_RS08685 | 640.7769204 | -2.767036254 | 0.307451571 | -8.999909291 | 2.26E-19 | 1.48E-18 | - | hypothetical protein | WP_156483737.1 | | | | |
| AVAEIV_RS11855 | 523.8250434 | -2.748453417 | 0.178375454 | -15.40824904 | 1.44E-53 | 2.31E-51 | *cobA* | uroporphyrinogen-III C-methyltransferase | WP_012700920.1 | Avin_23340 | *nasH* | Uroporphyrin-III C-methyltransferase, NasH | 100 |
| AVAEIV_RS19835 | 1422.23229 | -2.745496191 | 0.389677388 | -7.04556198 | 1.85E-12 | 5.99E-12 | *rimM* | ribosome maturation factor RimM | WP_012702469.1 | Avin_39550 | *rimM* | 16S rRNA processing protein RimM | 100 |
| AVAEIV_RS18815 | 707.1345279 | -2.74414538 | 0.223970587 | -12.25225786 | 1.63E-34 | 5.28E-33 | *fpr* | ferredoxin-NADP reductase | WP_061290376.1 | Avin_38470 | *fpr1* | NADPH: ferredoxin reductase | 99.61 |
| AVAEIV_RS17275 | 1349.933473 | -2.728474464 | 0.285674357 | -9.550995369 | 1.28E-21 | 1.12E-20 | - | TIGR04282 family arsenosugar biosynthesis glycosyltransferase | WP_012701968.1 | Avin_34390 | | conserved hypothetical protein | 100 |
| AVAEIV_RS20650 | 598.7011978 | -2.723473787 | 0.183533576 | -14.83910381 | 8.18E-50 | 9.71E-48 | - | YajG family lipoprotein | WP_012702638.1 | Avin_41320 | | conserved hypothetical protein | 100 |
| AVAEIV_RS09315 | 1315.156661 | -2.717175132 | 0.276856943 | -9.814365134 | 9.77E-23 | 9.53E-22 | - | DUF3597 domain-containing protein | WP_082765919.1 | Avin_18200 | | conserved hypothetical protein | 97.65 |
| AVAEIV_RS07920 | 1049.80064 | -2.715784738 | 0.163147499 | -16.64619292 | 3.22E-62 | 1.04E-59 | - | DEAD/DEAH box helicase | WP_302228531.1 | Avin_15370 | | ATP-dependent RNA helicase, DEAD/DEAH box family | 99.82 |
| AVAEIV_RS22080 | 708.1228453 | -2.713518331 | 0.202607929 | -13.39295228 | 6.65E-41 | 3.97E-39 | - | NAD-dependent epimerase/dehydratase family protein | WP_302227357.1 | Avin_44260 | | NAD dependent epimerase/dehydratase | 98.02 |
| AVAEIV_RS19025 | 3237.377449 | -2.709962178 | 0.317933017 | -8.523689056 | 1.55E-17 | 8.26E-17 | *bamA* | outer membrane protein assembly factor BamA | WP_012702406.1 | Avin_38910 | | bacterial surface antigen (D15) | 100 |
| AVAEIV_RS11560 | 71.04506184 | -2.708718367 | 0.297634616 | -9.100817647 | 8.97E-20 | 6.15E-19 | - | FeoA family protein | WP_012700861.1 | Avin_22720 | *feoA* | ferrous iron transport protein A | 100 |
| AVAEIV_RS14380 | 636.6050015 | -2.70843862 | 0.223583712 | -12.1137564 | 8.93E-34 | 2.67E-32 | - | RNA polymerase sigma factor FliA | WP_012701347.1 | Avin_27870 | *fliA* | motility sigma factor 28 | 100 |
| AVAEIV_RS11700 | 2906.186473 | -2.708283854 | 0.709813209 | -3.815488105 | 0.000135914 | 0.000215061 | - | DUF1826 domain-containing protein | WP_012700885.1 | Avin_22990 | | conserved hypothetical protein | 100 |
| AVAEIV_RS04350 | 1285.718606 | -2.704650566 | 0.34519367 | -7.835168493 | 4.68E-15 | 1.97E-14 | - | hypothetical protein | WP_012699565.1 | Avin_09010 | | conserved hypothetical protein | 100 |
| AVAEIV_RS10165 | 60.16669757 | -2.698243492 | 0.370098341 | -7.290612228 | 3.09E-13 | 1.09E-12 | *cydX* | cytochrome bd-I oxidase subunit CydX | WP_012700604.1 | Avin_19870 | | hypothetical transmembrane protein | 100 |
| AVAEIV_RS09720 | 476.5706469 | -2.690132393 | 0.221403643 | -12.15035287 | 5.71E-34 | 1.77E-32 | - | glutaredoxin family protein | WP_012700521.1 | Avin_19000 | | conserved hypothetical protein | 100 |
| AVAEIV_RS05150 | 1533.494556 | -2.685865934 | 0.353576776 | -7.596273613 | 3.05E-14 | 1.18E-13 | - | general stress protein | WP_012699765.1 | Avin_11130 | | hypothetical protein | 100 |
| AVAEIV_RS05515 | 845.4331961 | -2.679760806 | 0.199867597 | -13.40768011 | 5.45E-41 | 3.30E-39 | *rluD* | 23S rRNA pseudouridine(1911/1915/1917) synthase RluD | WP_041807510.1 | Avin_11940 | *rluD* | Pseudouridine synthase | 100 |
| AVAEIV_RS14590 | 2098.50581 | -2.663521841 | 0.528732908 | -5.037556393 | 4.72E-07 | 9.35E-07 | *clpS* | ATP-dependent Clp protease adapter ClpS | WP_012701387.1 | Avin_28280 | *clpS* | ATP-dependent clp protease adaptor protein, ClpS | 100 |

| | | | | | | | | | | | | | |
|---|---|---|---|---|---|---|---|---|---|---|---|---|---|
| AVAEIV_RS09210 | 146.6327515 | -2.655755184 | 0.32632975 | -8.138256419 | 4.01E-16 | 1.87E-15 | - | hypothetical protein | WP_012700417.1 | Avin_17940 | | hypothetical protein | 100 |
| AVAEIV_RS12215 | 559.4398342 | -2.640822371 | 0.197286872 | -13.38569743 | 7.33E-41 | 4.33E-39 | *flgJ* | flagellar assembly peptidoglycan hydrolase FlgJ | WP_012700990.1 | Avin_24080 | *flgJ* | flagellar rod assembly protein/muramidase | 100 |
| AVAEIV_RS12465 | 822.3811192 | -2.639744468 | 0.243232016 | -10.85278373 | 1.93E-27 | 2.99E-26 | - | GNAT family N-acetyltransferase | WP_302229127.1 | Avin_24790 | | N-acetyltransferase | 98.94 |
| AVAEIV_RS12295 | 266.8385685 | -2.63624232 | 0.202743163 | -13.00286669 | 1.18E-38 | 5.66E-37 | *fliE* | flagellar hook-basal body complex protein FliE | WP_012701006.1 | Avin_24240 | *fliE* | flagellar hook-basal body complex protein | 100 |
| AVAEIV_RS06820 | 1385.78507 | -2.63161271 | 0.404182883 | -6.510945465 | 7.47E-11 | 2.08E-10 | *rplM* | 50S ribosomal protein L13 | WP_012699950.1 | Avin_13040 | *rplM* | 50 S ribosomal protein L13 | 100 |
| AVAEIV_RS08625 | 1732.377112 | -2.629797665 | 0.253704079 | -10.3656105 | 3.55E-25 | 4.56E-24 | - | acetyl-CoA hydrolase/transferase family protein | WP_012700293.1 | Avin_16700 | | acetyl-CoA hydrolase/transferase family protein | 100 |
| AVAEIV_RS12025 | 22123.65419 | -2.627241637 | 0.741245939 | -3.544358894 | 0.000393569 | 0.000595976 | - | phasin family protein | WP_012700953.1 | Avin_23670 | *phbP* | phasin protein | 100 |
| AVAEIV_RS06710 | 1320.144269 | -2.62680317 | 0.326349346 | -8.049052965 | 8.34E-16 | 3.78E-15 | *mlaE* | lipid asymmetry maintenance ABC transporter permease subunit MlaE | WP_012699932.1 | Avin_12860 | | organic solvent tolerance ABC efflux transporter, permease component | 100 |
| AVAEIV_RS05045 | 1458.982285 | -2.618446722 | 0.283680548 | -9.230265307 | 2.70E-20 | 1.97E-19 | - | mannuronate-specific alginate lyase | WP_012699742.1 | Avin_10900 | *algL* | alginate lyase; Poly(beta-D-mannuronate) lyase | 100 |
| AVAEIV_RS20210 | 426.8383326 | -2.608859964 | 0.173100682 | -15.07134424 | 2.50E-51 | 3.30E-49 | *iscU* | Fe-S cluster assembly scaffold IscU | WP_012702550.1 | Avin_40390 | *iscU* | Iron-sulfur cluster assembly scaffold protein | 100 |
| AVAEIV_RS07070 | 5251.759488 | -2.606837303 | 0.546841222 | -4.767082652 | 1.87E-06 | 3.53E-06 | - | enoyl-CoA hydratase | WP_302229298.1 | Avin_13550 | | Enoyl-CoA hydratase/isomerase | 99.27 |
| AVAEIV_RS20205 | 290.2432483 | -2.596582676 | 0.16198094 | -16.03017411 | 7.87E-58 | 1.76E-55 | *iscA* | iron-sulfur cluster assembly protein IscA | WP_012702549.1 | Avin_40380 | *iscA* | Iron-sulfur biogenesis protein | 100 |
| AVAEIV_RS26000 | 631.7961189 | -2.596502374 | 0.171773404 | -15.11585791 | 1.27E-51 | 1.82E-49 | *rsmG* | 16S rRNA (guanine(527)-N(7))-methyltransferase RsmG | WP_061290846.1 | Avin_52260 | *gidB* | Glucose inhibited division protein B, gidB | 99.54 |
| AVAEIV_RS03355 | 272.9227768 | -2.590084178 | 0.250900342 | -10.32315923 | 5.54E-25 | 6.91E-24 | *nusB* | transcription antitermination factor NusB | WP_050764334.1 | Avin_06760 | *nusB* | transcription antitermination factor NusB | 100 |
| AVAEIV_RS11795 | 925.3930133 | -2.589018289 | 0.27721551 | -9.339370247 | 9.69E-21 | 7.45E-20 | - | GntR family transcriptional regulator | WP_012700906.1 | Avin_23200 | *gntR* | transcriptional regulator, GntR | 100 |
| AVAEIV_RS14875 | 2115.077896 | -2.588605631 | 0.446058547 | -5.803286693 | 6.50E-09 | 1.52E-08 | *nfuA* | Fe-S biogenesis protein NfuA | WP_012701434.1 | Avin_28760 | *nfuA* | NfuA protein | 100 |
| AVAEIV_RS05420 | 2630.089854 | -2.587705549 | 0.400005274 | -6.469178582 | 9.85E-11 | 2.71E-10 | - | hypothetical protein | WP_231876192.1 | Avin_11740 | | hypothetical protein | 100 |
| AVAEIV_RS15220 | 1358.118064 | -2.587258064 | 0.26760325 | -9.668261002 | 4.11E-22 | 3.80E-21 | - | 4'-phosphopantetheinyl transferase superfamily protein | WP_012701506.1 | Avin_29520 | *arsD* | Type I fatty acid synthase ArsD | 100 |
| AVAEIV_RS10355 | 378.9445651 | -2.585625293 | 0.254625987 | -10.15460095 | 3.16E-24 | 3.69E-23 | - | DUF2934 domain-containing protein | WP_012700640.1 | Avin_20270 | | hypothetical protein | 100 |
| AVAEIV_RS19785 | 534.2840793 | -2.580978526 | 0.215853797 | -11.957068 | 5.96E-33 | 1.70E-31 | - | YaeQ family protein | WP_012702459.1 | Avin_39450 | *yaeQ* | YaeQ family protein | 100 |
| AVAEIV_RS18825 | 669.9593372 | -2.580034421 | 0.228047777 | -11.31356968 | 1.12E-29 | 2.21E-28 | *erdR* | response regulator transcription factor ErdR | WP_012702365.1 | Avin_38490 | | Response regulator, LuxR family | 100 |
| AVAEIV_RS20405 | 850.084493 | -2.577589776 | 0.589613125 | -4.371662819 | 1.23E-05 | 2.16E-05 | *rpmA* | 50S ribosomal protein L27 | WP_012702586.1 | Avin_40780 | *rpmA* | ribosomal protein L27 | 100 |
| AVAEIV_RS20200 | 335.2393264 | -2.576969358 | 0.12797245 | -20.13690722 | 3.51E-90 | 9.01E-87 | *hscB* | co-chaperone HscB | WP_012702548.1 | Avin_40370 | *hscB* | Co-chaperone Hsc20 | 100 |
| AVAEIV_RS16585 | 3229.206738 | -2.572829006 | 0.676624699 | -3.802446185 | 0.000143274 | 0.00022622 | - | PA1414 family protein | WP_012701825.1 | Avin_32840 | | conserved hypothetical protein | 100 |
| AVAEIV_RS09725 | 2138.848962 | -2.565743419 | 0.392171467 | -6.542402071 | 6.05E-11 | 1.69E-10 | - | thermonuclease family protein | WP_012700522.1 | Avin_19010 | | Staphylococcus nuclease (SNase-like) protein | 100 |
| AVAEIV_RS25325 | 1102.140455 | -2.565021939 | 0.316007281 | -8.116971024 | 4.78E-16 | 2.21E-15 | *tssK* | type VI secretion system baseplate subunit TssK | WP_012703518.1 | Avin_50780 | | conserved hypothetical protein | 100 |
| AVAEIV_RS18555 | 551.2719744 | -2.562984366 | 0.521810759 | -4.911712383 | 9.03E-07 | 1.75E-06 | - | hypothetical protein | WP_061289358.1 | Avin_37900 | | hypothetical protein | 92.59 |
| AVAEIV_RS03190 | 294.7747847 | -2.55411148 | 0.21317121 | -11.98150296 | 4.44E-33 | 1.28E-31 | *rplR* | 50S ribosomal protein L18 | WP_012699318.1 | Avin_06410 | *rplR* | 50S ribosomal protein L18 | 100 |
| AVAEIV_RS03815 | 762.4744096 | -2.548921128 | 0.288136548 | -8.846226353 | 9.05E-19 | 5.55E-18 | - | neutral zinc metallopeptidase | WP_302228191.1 | Avin_07770 | | zinc metallopeptidase | 98.24 |
| AVAEIV_RS09085 | 1627.164181 | -2.5479997 | 0.377888515 | -6.742728604 | 1.55E-11 | 4.60E-11 | - | carbohydrate kinase family protein | WP_012700390.1 | Avin_17670 | | PfkB-family carbohydrate kinase | 100 |
| AVAEIV_RS11995 | 2629.18645 | -2.54554128 | 0.579967769 | -4.389108184 | 1.14E-05 | 2.00E-05 | *hupB* | nucleoid-associated protein HU-beta | WP_012700946.1 | Avin_23600 | | histone-like bacterial DNA-binding protein | 100 |
| AVAEIV_RS06655 | 208.3849766 | -2.53746421 | 0.231745471 | -10.94935837 | 6.69E-28 | 1.09E-26 | - | HPr family phosphocarrier protein | WP_012699921.1 | Avin_12750 | *ptsO* | Phosphocarrier NPr protein | 100 |
| AVAEIV_RS23985 | 1434.164482 | -2.536712427 | 0.414448607 | -6.12069237 | 9.32E-10 | 2.35E-09 | - | LytR/AlgR family response regulator transcription factor | WP_012703228.1 | Avin_47610 | *algR* | encystment and alginate biosynthesis response regulator; AlgR | 100 |
| AVAEIV_RS03220 | 857.6641273 | -2.534135459 | 0.324844839 | -7.801064247 | 6.14E-15 | 2.56E-14 | *rpsM* | 30S ribosomal protein S13 | WP_012699324.1 | Avin_06470 | | 30S ribosomal protein S13 | 100 |
| AVAEIV_RS08690 | 816.2079608 | -2.523843926 | 0.162340702 | -15.54658753 | 1.68E-54 | 3.08E-52 | - | substrate-binding periplasmic protein | WP_012700307.1 | Avin_16838 | | amino acid ABC transporter periplasmic protein | 100 |
| AVAEIV_RS12315 | 482.4271487 | -2.51823524 | 0.134801441 | -18.68107063 | 7.06E-78 | 5.18E-75 | - | hypothetical protein | WP_302229103.1 | Avin_24280 | | conserved hypothetical protein | 99.45 |
| AVAEIV_RS18445 | 450.8664398 | -2.517520627 | 0.278217261 | -9.048757851 | 1.45E-19 | 9.71E-19 | - | hypothetical protein | WP_231876250.1 | Avin_37670 | | hypothetical protein | 100 |
| AVAEIV_RS03240 | 792.9402456 | -2.515428713 | 0.355887974 | -7.068035157 | 1.57E-12 | 5.16E-12 | *rplQ* | 50S ribosomal protein L17 | WP_012699327.1 | Avin_06510 | *rplQ* | 50S ribosomal protein L17 | 100 |
| AVAEIV_RS24060 | 1094.197787 | -2.514350341 | 0.275197232 | -9.136539357 | 6.45E-20 | 4.51E-19 | - | ammonium transporter | WP_012703244.1 | Avin_47780 | *amtB* | Ammonium transporter | 100 |
| AVAEIV_RS00805 | 390.2117728 | -2.511668405 | 0.283425691 | -8.86182335 | 7.87E-19 | 4.87E-18 | *nifW* | nitrogenase-stabilizing/protective protein NifW | WP_012698858.1 | Avin_01670 | *nifW* | Nitrogen fixation protein NifW | 100 |
| AVAEIV_RS17580 | 591.8717294 | -2.509468896 | 0.460909486 | -5.444602403 | 5.19E-08 | 1.12E-07 | - | DUF485 domain-containing protein | WP_012702033.1 | Avin_35040 | | conserved hypothetical protein | 100 |
| AVAEIV_RS18870 | 738.0549516 | -2.508987432 | 0.194880376 | -12.87450019 | 6.26E-38 | 2.85E-36 | - | hypothetical protein | WP_012702374.1 | Avin_38580 | | conserved hypothetical protein | 100 |
| AVAEIV_RS01830 | 388.0050508 | -2.508177335 | 0.213423078 | -11.75213738 | 6.89E-32 | 1.74E-30 | *fdhD* | formate dehydrogenase accessory sulfurtransferase FdhD | WP_302229228.1 | Avin_03800 | *fdhD* | formate dehydrogenase family accessory protein FdhD | 98.91 |
| AVAEIV_RS02945 | 556.1750442 | -2.508014783 | 0.260400435 | -9.631377068 | 5.89E-22 | 5.35E-21 | - | glycine cleavage system protein R | WP_012699274.1 | Avin_05930 | | ACT domain-containing regulatory protein | 100 |
| AVAEIV_RS03065 | 992.6865695 | -2.507051234 | 0.40721849 | -6.156526026 | 7.44E-10 | 1.90E-09 | *rplJ* | 50S ribosomal protein L10 | WP_012699294.1 | Avin_06150 | | 50S ribosomal protein L10 | 100 |
| AVAEIV_RS21450 | 2149.71618 | -2.494627678 | 0.387277999 | -6.441439182 | 1.18E-10 | 3.23E-10 | *secG* | preprotein translocase subunit SecG | WP_012702776.1 | Avin_42850 | *secG* | Preprotein translocase SecG subunit | 100 |
| AVAEIV_RS14450 | 435.1920212 | -2.493725145 | 0.16017077 | -15.56916494 | 1.18E-54 | 2.25E-52 | - | transposase | pseudo=true | | | | |
| AVAEIV_RS18800 | 5643.135856 | -2.490305921 | 0.514664749 | -4.838695335 | 1.31E-06 | 2.50E-06 | - | fumarate hydratase | WP_041807368.1 | Avin_38440 | *fumB* | fumarate hydratase, class I | 99.8 |
| AVAEIV_RS00825 | 4112.821676 | -2.48082786 | 0.61177129 | -4.055155743 | 5.01E-05 | 8.30E-05 | - | flavodoxin | WP_012698862.1 | Avin_01710 | *nifF* | Flavodoxin, nifF | 100 |
| AVAEIV_RS25395 | 488.2860521 | -2.472314326 | 0.185221621 | -13.34787111 | 1.22E-40 | 7.12E-39 | *nifX* | nitrogen fixation protein NifX | WP_012703531.1 | Avin_50910 | *nafY* | Nitrogen fixation-related protein, gamma subunit | 100 |
| AVAEIV_RS19775 | 2331.496173 | -2.468810134 | 0.351849663 | -7.016661919 | 2.27E-12 | 7.28E-12 | - | NADH:flavin oxidoreductase/NADH oxidase | WP_061288056.1 | Avin_39430 | | NADH:flavin oxidoreductase/NADH oxidase | 99.73 |
| AVAEIV_RS12945 | 677.1567522 | -2.465029371 | 0.213328751 | -11.55507339 | 6.96E-31 | 1.52E-29 | - | FAD/NAD(P)-binding protein | WP_012701147.1 | Avin_25760 | | Oxidoreductase FAD/NAD(P)-binding:Oxidoreductase FAD-binding region | 100 |
| AVAEIV_RS16285 | 184.0626681 | -2.455721766 | 0.237253368 | -10.35062974 | 4.16E-25 | 5.28E-24 | - | hypothetical protein | WP_367281702.1 | | | | |
| AVAEIV_RS14065 | 2231.877157 | -2.450946185 | 0.409015246 | -5.992310096 | 2.07E-09 | 5.05E-09 | *eno* | phosphopyruvate hydratase | WP_302225444.1 | Avin_22210 | *eno-2* | Enolase | 98.86 |
| AVAEIV_RS18810 | 313.2635322 | -2.448728225 | 0.150645657 | -16.25488758 | 2.06E-59 | 5.58E-57 | *tsaA* | tRNA (N6-threonylcarbamoyladenosine(37)-N6)-methyltransferase TrmO | WP_175555870.1 | Avin_38460 | | Conserved Hypothetical Protein | 100 |
| AVAEIV_RS22715 | 155.356682 | -2.448656636 | 0.227269082 | -10.77426201 | 4.55E-27 | 6.79E-26 | - | hypothetical protein | WP_012703042.1 | Avin_45630 | | Conserved hypothetical protein | 100 |
| AVAEIV_RS15905 | 965.978881 | -2.445448557 | 0.373214477 | -6.552394687 | 5.66E-11 | 1.59E-10 | - | hypothetical protein | WP_012701643.1 | Avin_30940 | | hypothetical protein | 100 |
| AVAEIV_RS22630 | 263.5463039 | -2.443061035 | 0.240157249 | -10.17275576 | 2.62E-24 | 3.11E-23 | - | phosphoribosyl-ATP diphosphatase | WP_012703026.1 | Avin_45450 | *hisE* | Phosphoribosyl-ATP pyrophosphohydrolase | 100 |
| AVAEIV_RS24605 | 997.8597366 | -2.43594268 | 0.164427414 | -14.8146992 | 1.18E-49 | 1.34E-47 | - | acyl-CoA desaturase | WP_302229439.1 | Avin_48940 | | Fatty acid desaturase | 98.99 |
| AVAEIV_RS15185 | 361.9233814 | -2.434511841 | 0.149597165 | -16.27378327 | 1.52E-59 | 4.33E-57 | - | DUF2147 domain-containing protein | WP_302225718.1 | Avin_29440 | | conserved hypothetical protein | 98.66 |
| AVAEIV_RS21480 | 899.2487179 | -2.431292892 | 0.366612757 | -6.631773841 | 3.32E-11 | 9.55E-11 | *greA* | transcription elongation factor GreA | WP_012702782.1 | Avin_42920 | *greA* | Prokaryotic transcription elongation factor GreA | 100 |
| AVAEIV_RS24205 | 520.9376053 | -2.430042638 | 0.121317612 | -20.03041925 | 2.99E-89 | 5.13E-86 | *ftsX* | permease-like cell division protein FtsX | WP_012703274.1 | Avin_48080 | *ftsX* | Cell division protein FtsX | 100 |
| AVAEIV_RS09660 | 1379.714534 | -2.427643714 | 0.36003592 | -6.742781974 | 1.55E-11 | 4.60E-11 | - | hypothetical protein | WP_012700508.1 | Avin_18870 | | hypothetical protein | 100 |
| AVAEIV_RS00755 | 367.8820841 | -2.42413013 | 0.249986585 | -9.697040843 | 3.10E-22 | 2.89E-21 | - | hypothetical protein | WP_231876279.1 | Avin_01560 | | conserved hypothetical protein | 99.26 |
| AVAEIV_RS23275 | 96.23802185 | -2.417263153 | 0.279443583 | -8.650272554 | 5.14E-18 | 2.87E-17 | - | hypothetical protein | pseudo=true | | | | |
| AVAEIV_RS02940 | 1465.216264 | -2.415194807 | 0.301926307 | -7.999285746 | 1.25E-15 | 5.57E-15 | - | malate synthase G | WP_302228106.1 | Avin_05920 | *glcB* | malate synthase G | 99.31 |
| AVAEIV_RS21335 | 858.8462793 | -2.411369369 | 0.359191255 | -6.713329844 | 1.90E-11 | 5.59E-11 | *dksA* | RNA polymerase-binding protein DksA | WP_012702755.1 | Avin_42640 | *dksA* | RNA polymerase binding protein DksA | 100 |
| AVAEIV_RS00215 | 1467.018583 | -2.408177343 | 0.351097144 | -6.859005796 | 6.93E-12 | 2.11E-11 | - | hypothetical protein | WP_012698739.1 | Avin_00440 | | Transcriptional regulatory protein TetR family | 100 |
| AVAEIV_RS23955 | 2201.401967 | -2.405776907 | 0.504219732 | -4.771286708 | 1.83E-06 | 3.46E-06 | - | Rsd/AlgQ family anti-sigma factor | WP_012703222.1 | Avin_47550 | *algQ* | Regulator of RNA polymerase sigma(70) subunit, Rsd/AlgQ | 100 |
| AVAEIV_RS18425 | 612.8532596 | -2.405060654 | 0.561543071 | -4.28294957 | 1.84E-05 | 3.18E-05 | - | hypothetical protein | WP_012702233.1 | Avin_37120 | | hypothetical protein | 100 |
| AVAEIV_RS17360 | 2885.285176 | -2.399162448 | 0.311879836 | -7.692585972 | 1.44E-14 | 5.77E-14 | *acs* | acetate--CoA ligase | WP_302226295.1 | Avin_34530 | *acsA1* | acetate-CoA ligase | 99.69 |
| AVAEIV_RS20220 | 653.0836619 | -2.39781245 | 0.230065939 | -10.42228354 | 1.96E-25 | 2.57E-24 | *iscR* | Fe-S cluster assembly transcriptional regulator IscR | WP_012702552.1 | Avin_40410 | *iscR* | Iron-sulphur cluster assembly transcription factor IscR | 100 |

| | | | | | | | | | | | | |
|---|---|---|---|---|---|---|---|---|---|---|---|---|
| AVAEIV_RS10160 | 474.7159756 | -2.39566007 | 0.42691353 | -5.611581513 | 2.00E-08 | 4.48E-08 | - | cyd operon YbgE family protein | WP_012700603.1 | Avin_19860 | | hypothetical protein | 100 |
| AVAEIV_RS11925 | 366.7812447 | -2.393448986 | 0.203710051 | -11.74929259 | 7.12E-32 | 1.79E-30 | - | DUF1289 domain-containing protein | WP_012700934.1 | Avin_23480 | | conserved hypothetical protein | 100 |
| AVAEIV_RS03350 | 502.2838151 | -2.390715671 | 0.411160005 | -5.814562804 | 6.08E-09 | 1.42E-08 | *ribE* | 6,2C7-dimethyl-8-ribityllumazine synthase | WP_012699351.1 | Avin_06750 | *ribH* | 6,7-dimethyl-8-ribityllumazine synthase | 100 |
| AVAEIV_RS00090 | 353.6783117 | -2.386587882 | 0.258378965 | -9.236773133 | 2.54E-20 | 1.87E-19 | *def* | peptide deformylase | WP_012698713.1 | Avin_00170 | *def* | Formylmethionine deformylase protein | 100 |
| AVAEIV_RS07550 | 2613.35193 | -2.38073043 | 0.285215156 | -8.347138565 | 6.99E-17 | 3.53E-16 | - | glyceraldehyde-3-phosphate dehydrogenase | WP_041807044.1 | Avin_14580 | *gapB* | glyceraldehyde-3-phosphate dehydrogenase | 100 |
| AVAEIV_RS00790 | 1088.027919 | -2.380601959 | 0.239017063 | -9.959966581 | 2.28E-23 | 2.42E-22 | *nifV* | homocitrate synthase | WP_012698855.1 | Avin_01640 | *nifV* | nitrogen fixation homocitrate synthase | 100 |
| AVAEIV_RS09220 | 607.2775359 | -2.374584573 | 0.565764738 | -4.197123669 | 2.70E-05 | 4.59E-05 | - | DUF2789 domain-containing protein | WP_012700419.1 | Avin_17960 | | conserved hypothetical protein | 100 |
| AVAEIV_RS19725 | 336.2120186 | -2.364681267 | 0.248220561 | -9.526532606 | 1.63E-21 | 1.40E-20 | - | O-acetyl-ADP-ribose deacetylase | WP_012702447.1 | Avin_39320 | | Appr-1-p processing protein | 100 |
| AVAEIV_RS07710 | 1014.178328 | -2.361890799 | 0.292661471 | -8.070385185 | 7.01E-16 | 3.22E-15 | *fabG* | 3-oxoacyl-ACP reductase FabG | WP_012700125.1 | Avin_14920 | *fabG* | 3-oxoacyl (acyl carrier protein) reductase | 100 |
| AVAEIV_RS19840 | 1086.002078 | -2.355708491 | 0.423762298 | -5.55903274 | 2.71E-08 | 5.99E-08 | *rpsP* | 30S ribosomal protein S16 | WP_012702470.1 | Avin_39560 | *rpsP* | ribosomal protein S16 | 100 |
| AVAEIV_RS14375 | 857.1145806 | -2.352016162 | 0.125365175 | -18.76131999 | 1.56E-78 | 1.34E-75 | - | HD domain-containing phosphohydrolase | WP_061290720.1 | Avin_27860 | | Metal-dependent phosphohydrolase, HD domain protein | 99.65 |
| AVAEIV_RS09090 | 1628.09499 | -2.350991279 | 0.322789358 | -7.283360552 | 3.26E-13 | 1.15E-12 | - | aldose 1-epimerase family protein | WP_061291863.1 | Avin_17680 | | conserved hypothetical protein | 99.26 |
| AVAEIV_RS03375 | 831.3195606 | -2.346183002 | 0.33637141 | -6.97497746 | 3.06E-12 | 9.67E-12 | *ribA* | GTP cyclohydrolase II | WP_061290418.1 | Avin_06800 | | GTP cyclohydrolase II | 99.52 |
| AVAEIV_RS08820 | 1386.240786 | -2.341654318 | 0.390074117 | -6.003100993 | 1.94E-09 | 4.75E-09 | - | ogr/Delta-like zinc finger family protein | WP_302228600.1 | Avin_17050 | | ParB protein | 87.81 |
| AVAEIV_RS24025 | 677.967176 | -2.339613279 | 0.283567411 | -8.250642301 | 1.58E-16 | 7.70E-16 | *dapF* | diaminopimelate epimerase | WP_012703237.1 | Avin_47710 | *dapF* | diaminopimelate epimerase | 100 |
| AVAEIV_RS15105 | 428.8616602 | -2.336189537 | 0.595831831 | -3.920887433 | 8.82E-05 | 0.000142763 | - | hypothetical protein | WP_231876200.1 | Avin_29230 | | conserved hypothetical protein | 100 |
| AVAEIV_RS06315 | 1723.074886 | -2.33590748 | 0.187785438 | -12.43923654 | 1.60E-35 | 5.67E-34 | - | NAD(P)/FAD-dependent oxidoreductase | WP_012699846.1 | Avin_12000 | *ndh* | uncoupled NADH:quinone oxidoreductase | 100 |
| AVAEIV_RS06415 | 981.8801129 | -2.333099021 | 0.273159773 | -8.541151562 | 1.33E-17 | 7.15E-17 | - | PTS fructose-like transporter subunit IIB | WP_302228466.1 | Avin_12210 | *fruA* | Fructose phosphotransferase system IIBC component | 99.48 |
| AVAEIV_RS25400 | 207.5107739 | -2.332889065 | 0.283406728 | -8.23159381 | 1.85E-16 | 8.93E-16 | - | RnfH family protein | WP_012703532.1 | Avin_50920 | *rnfH* | Electron transport complex, RnfABCDGE type, H subunit | 100 |
| AVAEIV_RS25540 | 3049.492622 | -2.332650077 | 0.313742886 | -7.434909873 | 1.05E-13 | 3.86E-13 | - | hypothetical protein | WP_041807467.1 | | | | |
| AVAEIV_RS12175 | 291.0427674 | -2.330164409 | 0.152832658 | -15.24650848 | 1.74E-52 | 2.71E-50 | *flgB* | flagellar basal body rod protein FlgB | WP_012700982.1 | Avin_24000 | *flgB* | flagellar basal-body rod protein | 100 |
| AVAEIV_RS14090 | 1100.149767 | -2.329611056 | 0.240109856 | -9.702271679 | 2.95E-22 | 2.75E-21 | - | MurR/RpiR family transcriptional regulator | WP_012701291.1 | Avin_27270 | *hexR-1* | transcriptional regulatory protein, RpiR family | 100 |
| AVAEIV_RS11750 | 3665.214431 | -2.325568863 | 0.583067946 | -3.98850405 | 6.65E-05 | 0.000108587 | - | Lpp/OprI family alanine-zipper lipoprotein | WP_012700896.1 | Avin_23100 | *oprI* | outer membrane lipoprotein OprI | 100 |
| AVAEIV_RS02260 | 859.042477 | -2.315313325 | 0.19074603 | -12.13819931 | 6.63E-34 | 2.04E-32 | *cooS* | anaerobic carbon-monoxide dehydrogenase catalytic subunit | WP_012699133.1 | Avin_04490 | *cooS* | carbon-monoxide dehydrogenase, catalytic subunit | 100 |
| AVAEIV_RS19885 | 3094.597568 | -2.315130669 | 0.235166612 | -9.844640133 | 7.23E-23 | 7.19E-22 | *purL* | phosphoribosylformylglycinamidine synthase | WP_012702480.1 | Avin_39660 | *purL* | phosphoribosylformylglycinamidine synthase | 100 |
| AVAEIV_RS03085 | 756.2037733 | -2.314077212 | 0.473943268 | -4.882603825 | 1.05E-06 | 2.02E-06 | *rpsL* | 30S ribosomal protein S12 | WP_012699298.1 | Avin_06190 | *rpsL* | 30S ribosomal protein S12 | 100 |
| AVAEIV_RS12355 | 1129.29418 | -2.31397757 | 0.207361209 | -11.15916316 | 6.46E-29 | 1.18E-27 | - | methyl-accepting chemotaxis protein | WP_302229107.1 | Avin_24360 | | chemotaxis sensory transducer | 99.82 |
| AVAEIV_RS25675 | 7527.616316 | -2.312460663 | 0.510282039 | -4.531730466 | 5.85E-06 | 1.05E-05 | - | efflux RND transporter permease subunit | WP_012703586.1 | Avin_51560 | | multidrug efflux pump RND-family transporter protein | 100 |
| AVAEIV_RS09390 | 1688.208032 | -2.309815112 | 0.500614726 | -4.613957582 | 3.95E-06 | 7.23E-06 | - | hypothetical protein | WP_012700457.1 | Avin_18360 | | hypothetical protein | 100 |
| AVAEIV_RS06715 | 1326.318573 | -2.308407767 | 0.409156127 | -5.6418751 | 1.68E-08 | 3.78E-08 | *mlaD* | outer membrane lipid asymmetry maintenance protein MlaD | WP_012699933.1 | Avin_12870 | | organic solvent tolerance ABC efflux transporter, substrate binding protein | 100 |
| AVAEIV_RS25520 | 4666.812455 | -2.308241478 | 0.283906515 | -8.130287103 | 4.28E-16 | 1.99E-15 | - | glycosyl hydrolase family 28-related protein | WP_415860943.1 | Avin_51190 | *algE1* | Secreted mannuronan C-5 epimerase | 98.65 |
| AVAEIV_RS11800 | 769.0198616 | -2.307348469 | 0.167121462 | -13.80641628 | 2.33E-43 | 1.60E-41 | *prpB* | methylisocitrate lyase | WP_302229046.1 | Avin_23210 | *prpB* | methylisocitrate lyase | 99.66 |
| AVAEIV_RS00670 | 13018.44491 | -2.303338187 | 0.684192363 | -3.366506719 | 0.000761267 | 0.001120434 | *nifH* | nitrogenase iron protein | WP_012698831.1 | Avin_01380 | *nifH* | Nitrogenase iron protein | 100 |
| AVAEIV_RS14095 | 4376.284463 | -2.301063369 | 0.5226517 | -4.402670786 | 1.07E-05 | 1.88E-05 | *edd* | phosphogluconate dehydratase | WP_061291994.1 | Avin_27280 | *edd-2* | 6-phosphogluconate dehydratase | 99.34 |
| AVAEIV_RS16660 | 412.8315926 | -2.29805475 | 0.44793584 | -5.13032123 | 2.89E-07 | 5.88E-07 | - | hypothetical protein | WP_139239932.1 | | | | |
| AVAEIV_RS02235 | 710.2493726 | -2.296368802 | 0.188575848 | -12.17742798 | 4.10E-34 | 1.29E-32 | - | MBL fold metallo-hydrolase | WP_302228008.1 | Avin_04440 | | metallo hydrolase | 99.44 |
| AVAEIV_RS20535 | 171.3227646 | -2.293736861 | 0.156898536 | -14.61923686 | 2.12E-48 | 2.13E-46 | - | glutathione peroxidase | WP_012702616.1 | Avin_41100 | | Glutathione peroxidase | 100 |
| AVAEIV_RS07690 | 591.8289006 | -2.291304272 | 0.238291058 | -9.615569658 | 6.87E-22 | 6.19E-21 | - | YceD family protein | WP_012700122.1 | Avin_14880 | | Conserved hypothetical protein, DUF177 | 100 |
| AVAEIV_RS12280 | 224.2110275 | -2.290481821 | 0.13757417 | -16.64906881 | 3.07E-62 | 1.04E-59 | *fliH* | flagellar assembly protein FliH | WP_012701003.1 | Avin_24210 | *fliH* | Flagellar assembly protein | 100 |
| AVAEIV_RS06410 | 707.696214 | -2.288068522 | 0.304647075 | -7.510554703 | 5.89E-14 | 2.22E-13 | *pfkB* | 1-phosphofructokinase | WP_012699867.1 | Avin_12200 | *fruK* | 1-phosphofructokinase | 100 |
| AVAEIV_RS18090 | 701.7482382 | -2.285079464 | 0.298447531 | -7.656553408 | 1.91E-14 | 7.54E-14 | *pal* | peptidoglycan-associated lipoprotein Pal | WP_012702185.1 | Avin_36630 | *oprL* | peptidoglycan-associated outer membrane lipoprotein | 100 |
| AVAEIV_RS19830 | 1681.757663 | -2.284541 | 0.353798073 | -6.45718893 | 1.07E-10 | 2.93E-10 | *trmD* | tRNA (guanosine(37)-N1)-methyltransferase TrmD | WP_302226904.1 | Avin_39540 | *trmD* | tRNA (guanine-N(1)-)-methyltransferase | 99.6 |
| AVAEIV_RS06400 | 924.1099867 | -2.28169122 | 0.260868412 | -8.746521673 | 2.20E-18 | 1.28E-17 | *cra* | catabolite repressor/activator | WP_012699865.1 | Avin_12180 | *fruR* | Transcriptional regulator protein, LacI family | 100 |
| AVAEIV_RS18085 | 1306.429511 | -2.280708403 | 0.375546323 | -6.073041495 | 1.26E-09 | 3.13E-09 | *ybgF* | tol-pal system protein YbgF | WP_061290568.1 | Avin_36620 | | tol-pal system YbgF-like protein | 99.63 |
| AVAEIV_RS15350 | 607.4963232 | -2.276905079 | 0.273253859 | -8.332563305 | 7.91E-17 | 3.97E-16 | *sdhD* | succinate dehydrogenase,2C hydrophobic membrane anchor protein | WP_012701533.1 | Avin_29800 | *sdhD* | succinate dehydrogenase cytochrome b556 small membrane subunit | 100 |
| AVAEIV_RS11880 | 507.365334 | -2.273597511 | 0.156673088 | -14.51172974 | 1.02E-47 | 9.72E-46 | - | nitrate/nitrite transporter | WP_012700925.1 | Avin_23390 | *narK* | Nitrate/nitrite transporter | 100 |
| AVAEIV_RS10440 | 582.1648612 | -2.270224131 | 0.444484843 | -5.107540042 | 3.26E-07 | 6.59E-07 | *rplT* | 50S ribosomal protein L20 | WP_012700657.1 | Avin_20450 | *rplT* | 50S ribosomal protein L20 | 100 |
| AVAEIV_RS13915 | 1222.985608 | -2.269631276 | 0.470168236 | -4.827274801 | 1.38E-06 | 2.64E-06 | - | FadR/GntR family transcriptional regulator | WP_012701251.1 | Avin_26860 | | Bacterial regulatory protein, GntR family | 100 |
| AVAEIV_RS10035 | 176.8056706 | -2.260181815 | 0.156988764 | -14.39709289 | 5.40E-47 | 4.70E-45 | - | ABC transporter ATP-binding protein | WP_061288587.1 | Avin_19610 | | high affinity branched chain amino acid ABC transporter, ATP binding component | 99.14 |
| AVAEIV_RS11860 | 1529.879204 | -2.257748478 | 0.191454503 | -11.79261098 | 4.26E-32 | 1.10E-30 | - | nitrate reductase | WP_302229051.1 | Avin_23350 | *nasB* | assimilatory nitrate reductase, NasB | 99.45 |
| AVAEIV_RS20745 | 1013.863601 | -2.256028577 | 0.303078579 | -7.44370844 | 9.79E-14 | 3.62E-13 | *ispE* | 4-(cytidine 5'-diphospho)-2-C-methyl-D-erythritol kinase | WP_012702656.1 | Avin_41510 | *ispE* | 4-diphosphocytidyl-2C-methyl-D-erythritol kinase | 100 |
| AVAEIV_RS01145 | 851.2948994 | -2.255683113 | 0.156326794 | -14.42928022 | 3.39E-47 | 3.11E-45 | - | hypothetical protein | WP_050764409.1 | Avin_02320 | | hypothetical protein | 100 |
| AVAEIV_RS12100 | 1959.129244 | -2.255209827 | 0.523643451 | -4.306766035 | 1.66E-05 | 2.86E-05 | - | ribonuclease E inhibitor RraB | WP_012700968.1 | Avin_23830 | | conserved hypothetical protein | 100 |
| AVAEIV_RS06705 | 1225.463653 | -2.247439565 | 0.377056947 | -5.960477804 | 2.52E-09 | 6.10E-09 | - | ATP-binding cassette domain-containing protein | WP_012699931.1 | Avin_12850 | | organic solvent tolerance ABC efflux transporter, ATP binding component | 100 |
| AVAEIV_RS06725 | 677.3458655 | -2.245468859 | 0.254111266 | -8.836557694 | 9.87E-19 | 6.02E-18 | - | lipid asymmetry maintenance protein MlaB | WP_012699935.1 | Avin_12890 | | STAS family protein | 100 |
| AVAEIV_RS16610 | 320.0792089 | -2.244970972 | 0.140463645 | -15.98257668 | 1.69E-57 | 3.62E-55 | - | hypothetical protein | WP_012701830.1 | Avin_32890 | | conserved hypothetical protein | 100 |
| AVAEIV_RS21455 | 1552.154338 | -2.238879721 | 0.223421138 | -10.02089479 | 1.23E-23 | 1.34E-22 | *glmM* | phosphoglucosamine mutase | WP_012702777.1 | Avin_42860 | *glmM* | phosphoglucosamine mutase | 100 |
| AVAEIV_RS12035 | 498.8396432 | -2.238159869 | 0.195774188 | -11.43235426 | 2.88E-30 | 5.97E-29 | *folE* | GTP cyclohydrolase I FolE | WP_012700955.1 | Avin_23690 | *folE* | GTP cyclohydrolase I | 100 |
| AVAEIV_RS21495 | 849.0644154 | -2.237167565 | 0.178428277 | -12.5381896 | 4.61E-36 | 1.74E-34 | *dapB* | 4-hydroxy-tetrahydrodipicolinate reductase | WP_012702785.1 | Avin_42950 | *dapB* | dihydrodipicolinate reductase | 100 |
| AVAEIV_RS07065 | 2352.840886 | -2.235301646 | 0.307135358 | -7.277903997 | 3.39E-13 | 1.19E-12 | - | IS630 family transposase | pseudo=true | | | | |
| AVAEIV_RS24595 | 1966.811722 | -2.231065357 | 0.194031864 | -11.49844828 | 1.34E-30 | 2.84E-29 | - | OprD family porin | WP_012703354.1 | Avin_48920 | | Bacterial outer membrane porin | 100 |
| AVAEIV_RS04660 | 1787.094534 | -2.230433934 | 0.509060814 | -4.381468529 | 1.18E-05 | 2.07E-05 | *bfr* | bacterioferritin | WP_012699660.1 | Avin_10050 | | bacterioferritin protein | 100 |
| AVAEIV_RS00740 | 324.0238207 | -2.227505626 | 0.248514857 | -8.963269444 | 3.15E-19 | 2.03E-18 | - | hypothetical protein | WP_012698845.1 | Avin_01530 | | conserved hypothetical protein | 100 |
| AVAEIV_RS19905 | 2024.188333 | -2.225200275 | 0.522076229 | -4.262213355 | 2.02E-05 | 3.47E-05 | - | 2Fe-2S iron-sulfur cluster binding domain-containing protein | WP_302226922.1 | Avin_39700 | | Ferredoxin | 98.36 |
| AVAEIV_RS21955 | 432.5369109 | -2.224678077 | 0.231481699 | -9.610600245 | 7.21E-22 | 6.47E-21 | - | DUF5064 family protein | WP_012702888.1 | Avin_44000 | | conserved hypothetical protein | 100 |
| AVAEIV_RS20690 | 1774.703501 | -2.219715987 | 0.310518304 | -7.148422363 | 8.78E-13 | 2.95E-12 | - | NAD(P)/FAD-dependent oxidoreductase | WP_012702645.1 | Avin_41390 | | FAD dependent oxidoreductase | 100 |
| AVAEIV_RS03070 | 1932.785345 | -2.218358091 | 0.479386625 | -4.627492664 | 3.70E-06 | 6.79E-06 | *rplL* | 50S ribosomal protein L7/L12 | WP_012699295.1 | Avin_06160 | *rplL* | 50S ribosomal protein L7/L12 | 100 |
| AVAEIV_RS24215 | 784.2446353 | -2.217374408 | 0.197767189 | -11.21204393 | 3.56E-29 | 6.70E-28 | *ftsY* | signal recognition particle-docking protein FtsY | WP_012703276.1 | Avin_48100 | *ftsY* | signal recognition particle-docking protein FtsY | 100 |

| | | | | | | | | | | | | | |
|---|---|---|---|---|---|---|---|---|---|---|---|---|---|
| AVAEIV_RS03765 | 265.6268115 | -2.212079434 | 0.552662293 | -4.002587951 | 6.27E-05 | 0.000102711 | *rpsR* | 30S ribosomal protein S18 | WP_012699436.1 | Avin_07650 | *rpsR* | 30S ribosomal protein S18 | 100 |
| AVAEIV_RS23950 | 1498.262623 | -2.211804835 | 0.352705645 | -6.270965221 | 3.59E-10 | 9.37E-10 | - | AlgP family protein | WP_012703221.1 | Avin_47540 | | regulatory protein | 100 |
| AVAEIV_RS24200 | 3013.247792 | -2.209876752 | 0.553352299 | -3.993616282 | 6.51E-05 | 0.000106406 | *rpoH* | RNA polymerase sigma factor RpoH | WP_012703273.1 | Avin_48070 | *rpoH* | RNA polymerase sigma-32 factor | 100 |
| AVAEIV_RS14595 | 2070.689164 | -2.208356265 | 0.480058804 | -4.600178656 | 4.22E-06 | 7.69E-06 | *cspD* | cold shock domain-containing protein CspD | WP_012701388.1 | Avin_28290 | *cspD* | cold shock protein, CspD | 100 |
| AVAEIV_RS19005 | 629.3199967 | -2.207115039 | 0.230647517 | -9.569212211 | 1.08E-21 | 9.48E-21 | *lpxA* | acyl-ACP--UDP-N-acetylglucosamine O-acyltransferase | WP_012702402.1 | Avin_38870 | *lpxA* | acyl-(acyl-carrier-protein)--UDP-N-acetylglucosamine O-acyltransferase | 100 |
| AVAEIV_RS22625 | 281.5123831 | -2.204673567 | 0.235060866 | -9.379160407 | 6.65E-21 | 5.26E-20 | *hisI* | phosphoribosyl-AMP cyclohydrolase | WP_302227462.1 | Avin_45440 | *hisI* | Phosphoribosyl-AMP cyclohydrolase | 99.25 |
| AVAEIV_RS09070 | 1291.582428 | -2.203974823 | 0.36057967 | -6.112310284 | 9.82E-10 | 2.47E-09 | *pgl* | 6-phosphogluconolactonase | WP_061292207.1 | Avin_17640 | *pgl-2* | 6-phosphogluconolactonase | 99.58 |
| AVAEIV_RS15640 | 41.08563366 | -2.200678776 | 0.252678187 | -8.709413341 | 3.05E-18 | 1.75E-17 | *ccmD* | heme exporter protein CcmD | WP_012701593.1 | Avin_30420 | *ccmD* | Cytochrome c-type biogenesis protein | 100 |
| AVAEIV_RS18915 | 6351.219334 | -2.196173259 | 0.593969104 | -3.697453697 | 0.000217773 | 0.000337933 | - | peptidoglycan DD-metalloendopeptidase family protein | WP_302226640.1 | Avin_38690 | *nlpD* | lipoprotein NlpD | 99.65 |
| AVAEIV_RS11685 | 485.3721811 | -2.193523605 | 0.164522932 | -13.33263135 | 1.50E-40 | 8.64E-39 | - | hypothetical protein | WP_302229030.1 | Avin_22950 | | conserved hypothetical protein | 99.17 |
| AVAEIV_RS12185 | 604.3266598 | -2.193043256 | 0.175126965 | -12.52259044 | 5.62E-36 | 2.09E-34 | *flgD* | flagellar hook assembly protein FlgD | WP_012700984.1 | Avin_24020 | *flgD* | Flagellar hook capping protein | 100 |
| AVAEIV_RS03710 | 1721.108407 | -2.191978902 | 0.320770802 | -6.833473897 | 8.29E-12 | 2.50E-11 | *hflX* | ribosome rescue GTPase HflX | WP_012699426.1 | Avin_07550 | *hflX* | GTP-binding protein HflX | 100 |
| AVAEIV_RS17665 | 1766.819182 | -2.190601941 | 0.312351033 | -7.013269406 | 2.33E-12 | 7.46E-12 | - | FKBP-type peptidyl-prolyl cis-trans isomerase | WP_041807315.1 | Avin_35210 | | Peptidyl-prolyl cis-trans isomerase | 100 |
| AVAEIV_RS06885 | 1245.15569 | -2.188073418 | 0.377720675 | -5.792834655 | 6.92E-09 | 1.61E-08 | *mraZ* | division/cell wall cluster transcriptional repressor MraZ | WP_012699962.1 | Avin_13160 | *mraZ* | MraZ-family protein | 100 |
| AVAEIV_RS21460 | 1854.949888 | -2.183841572 | 0.311630094 | -7.007800638 | 2.42E-12 | 7.74E-12 | *folP* | dihydropteroate synthase | WP_175555939.1 | Avin_42870 | *folP* | dihydropteroate synthase | 100 |
| AVAEIV_RS14075 | 1239.487536 | -2.181863056 | 0.226929161 | -9.614731941 | 6.93E-22 | 6.23E-21 | *pyk* | pyruvate kinase | WP_302225457.1 | Avin_27240 | *pykA-1* | Pyruvate kinase | 99.58 |
| AVAEIV_RS25670 | 2379.106 | -2.180462902 | 0.347969251 | -6.266251668 | 3.70E-10 | 9.64E-10 | - | efflux RND transporter periplasmic adaptor subunit | WP_012703585.1 | Avin_51550 | | multidrug efflux pump membrane fusion protein | 100 |
| AVAEIV_RS02730 | 1209.051618 | -2.180459261 | 0.238240023 | -9.152363383 | 5.57E-20 | 3.92E-19 | - | phosphoenolpyruvate carboxykinase | WP_061288744.1 | Avin_05450 | *pckA* | phosphoenolpyruvate carboxykinase | 99.81 |
| AVAEIV_RS21655 | 171.1078968 | -2.180281129 | 0.50506562 | -4.316827444 | 1.58E-05 | 2.74E-05 | - | hypothetical protein | WP_175555891.1 | | | | |
| AVAEIV_RS06405 | 2193.027276 | -2.177274374 | 0.392700706 | -5.54436074 | 2.95E-08 | 6.50E-08 | *ptsP* | phosphoenolpyruvate--protein phosphotransferase | WP_012699866.1 | Avin_12190 | *fruB* | fructose-specific multiphosphoryl transfer protein | 100 |
| AVAEIV_RS08335 | 862.413068 | -2.174259007 | 0.172227586 | -12.62433654 | 1.55E-36 | 6.43E-35 | - | HlyD family type I secretion periplasmic adaptor subunit | WP_012700255.1 | Avin_16250 | *eexE* | Type I secretion membrane fusion protein | 100 |
| AVAEIV_RS21475 | 330.6646634 | -2.172558774 | 0.230626587 | -9.420244212 | 4.50E-21 | 3.67E-20 | *yhbY* | ribosome assembly RNA-binding protein YhbY | WP_012702781.1 | Avin_42910 | | conserved hypothetical protein | 100 |
| AVAEIV_RS00435 | 740.6149021 | -2.172347887 | 0.190551621 | -11.40031175 | 4.17E-30 | 8.57E-29 | - | zinc-dependent alcohol dehydrogenase family protein | WP_061291434.1 | Avin_00880 | | Zinc-containing alcohol dehydrogenase superfamily | 99.69 |
| AVAEIV_RS12940 | 443.6362388 | -2.170388911 | 0.267379763 | -8.117251968 | 4.77E-16 | 2.20E-15 | - | BCAM0308 family protein | WP_061289705.1 | Avin_25750 | | conserved hypothetical protein | 98.11 |
| AVAEIV_RS03855 | 54.42904457 | -2.169768897 | 0.234167574 | -9.265881107 | 1.93E-20 | 1.45E-19 | - | phosphonate metabolism protein PhnP | pseudo=true | | | | |
| AVAEIV_RS25685 | 558.7546336 | -2.168244655 | 0.300426942 | -7.217211084 | 5.31E-13 | 1.82E-12 | - | 3-hydroxyacyl-CoA dehydrogenase family protein | WP_302227749.1 | Avin_51580 | *hbdH* | hydroxybutyryl-CoA dehydrogenase | 99.52 |
| AVAEIV_RS00510 | 602.2578644 | -2.167704765 | 0.229560989 | -9.442827252 | 3.63E-21 | 3.00E-20 | - | hypothetical protein | WP_012698799.1 | Avin_01040 | | conserved hypothetical protein | 100 |
| AVAEIV_RS12210 | 497.3152641 | -2.167323636 | 0.173015471 | -12.52676207 | 5.33E-36 | 2.00E-34 | - | flagellar basal body P-ring protein FlgI | WP_175555965.1 | Avin_24070 | *flgI* | Flagellar P-ring protein | 100 |
| AVAEIV_RS06905 | 1549.921565 | -2.165397657 | 0.296293468 | -7.308286863 | 2.71E-13 | 9.64E-13 | - | UDP-N-acetylmuramoyl-L-alanyl-D-glutamate--2%2C6-diaminopimelate ligase | WP_012699966.1 | Avin_13200 | *murE* | UDP-N-acetylmuramyl-tripeptide synthetase | 100 |
| AVAEIV_RS06900 | 1710.236889 | -2.162999935 | 0.294348895 | -7.348422133 | 2.01E-13 | 7.22E-13 | - | peptidoglycan D%2CD-transpeptidase FtsI family protein | WP_012699965.1 | Avin_13190 | *ftsI* | Penicilin-binding Protein 3 | 100 |
| AVAEIV_RS23335 | 886.0109776 | -2.162352691 | 0.283779998 | -7.619820662 | 2.54E-14 | 9.90E-14 | - | alpha/beta fold hydrolase | WP_302227541.1 | Avin_46290 | | poly(3-hydroxybutyrate) synthase subunit | 99.14 |
| AVAEIV_RS16690 | 135.4945163 | -2.15981144 | 0.242405671 | -8.90990476 | 5.11E-19 | 3.22E-18 | *cobU* | bifunctional adenosylcobinamide kinase/adenosylcobinamide-phosphate guanylyltransferase | WP_302226063.1 | Avin_33060 | *cobP* | Cobinamide kinase protein | 97.11 |
| AVAEIV_RS02250 | 209.4149258 | -2.155439944 | 0.143995949 | -14.96875402 | 1.18E-50 | 1.47E-48 | - | AAA family ATPase | WP_012699131.1 | Avin_04470 | *cooC* | Carbon monoxide dehydrogenae assesory protein, CooC | 100 |
| AVAEIV_RS02195 | 1507.692617 | -2.15510508 | 0.450480598 | -4.784013095 | 1.72E-06 | 3.25E-06 | - | DUF3617 domain-containing protein | WP_302227999.1 | Avin_04350 | | conserved hypothetical protein | 99.43 |
| AVAEIV_RS00685 | 284.4933982 | -2.154854759 | 0.248912431 | -8.657079712 | 4.84E-18 | 2.72E-17 | *nifT* | putative nitrogen fixation protein NifT | WP_012698834.1 | Avin_01410 | *nifT* | Nitrogen fixation protein | 100 |
| AVAEIV_RS06465 | 386.6789005 | -2.154026867 | 0.186005652 | -11.58043769 | 5.18E-31 | 1.14E-29 | - | Fe2+-dependent dioxygenase | WP_012699881.1 | Avin_12340 | | 2-oxoglutarate and Fe(II)-dependent dioxygenase superfamily | 100 |
| AVAEIV_RS11710 | 1193.075098 | -2.153401858 | 0.275286855 | -7.822392607 | 5.18E-15 | 2.17E-14 | - | transglycosylase SLT domain-containing protein | WP_302229035.1 | Avin_23020 | | Lytic transglycosylase protein | 99.58 |
| AVAEIV_RS03755 | 698.8800151 | -2.146706938 | 0.263629684 | -8.142887814 | 3.86E-16 | 1.80E-15 | *rlmB* | 23S rRNA (guanosine(2251)-2'-O)-methyltransferase RlmB | WP_012699434.1 | Avin_07630 | | RNA methyltransferase TrmH, group 3 | 100 |
| AVAEIV_RS00795 | 970.1132221 | -2.143696734 | 0.21750026 | -9.856065162 | 6.45E-23 | 6.48E-22 | *cysE* | serine O-acetyltransferase | WP_012698856.1 | Avin_01650 | *cysE1nif* | nitrogen fixation serine O-acetyltransferase CysE1 | 100 |
| AVAEIV_RS03090 | 610.3367937 | -2.13807328 | 0.253395958 | -8.437677144 | 3.24E-17 | 1.67E-16 | *rpsG* | 30S ribosomal protein S7 | WP_012699299.1 | Avin_06200 | *rpsG* | 30S ribosomal protein S7 | 100 |
| AVAEIV_RS12020 | 3291.518723 | -2.137452271 | 0.478084975 | -4.470862678 | 7.79E-06 | 1.38E-05 | - | AraC family transcriptional regulator | WP_231876181.1 | Avin_23660 | *phbR* | Transcription activator in PHB biosynthesis, AraC family | 100 |
| AVAEIV_RS20430 | 455.2408495 | -2.133798434 | 0.222373684 | -9.595552852 | 8.35E-22 | 7.42E-21 | - | hypothetical protein | WP_012702592.1 | Avin_40840 | | conserved hypothetical protein | 100 |
| AVAEIV_RS17770 | 2418.553564 | -2.133173998 | 0.421932087 | -5.055728311 | 4.29E-07 | 8.53E-07 | - | polyhydroxyalkanoate granule-associated phasin | WP_231876213.1 | Avin_35400 | | hypothetical protein | 100 |
| AVAEIV_RS24140 | 2640.0731 | -2.131506419 | 0.528316192 | -4.034527901 | 5.47E-05 | 9.03E-05 | - | c-type cytochrome | WP_373860944.1 | Avin_47940 | *cycB* | cytochrome c5 protein | 100 |
| AVAEIV_RS16520 | 1054.704878 | -2.129411801 | 0.403619794 | -5.275786351 | 1.32E-07 | 2.76E-07 | *rssC* | anti-sigma factor antagonist RssC | WP_012701812.1 | Avin_32710 | | antisigma-factor antagonist STAS domain protein | 100 |
| AVAEIV_RS12335 | 378.8691409 | -2.128177197 | 0.152259756 | -13.97727972 | 2.15E-44 | 1.62E-42 | *motA* | flagellar motor stator protein MotA | WP_012701014.1 | Avin_24320 | *motA* | Fagellar motor protein | 100 |
| AVAEIV_RS12305 | 476.4105498 | -2.127844432 | 0.408898261 | -5.20384808 | 1.95E-07 | 4.03E-07 | *fliS* | flagellar export chaperone FliS | WP_231876321.1 | Avin_24260 | *fliS* | flagellar protein FliS | 100 |
| AVAEIV_RS08260 | 844.8562818 | -2.127747172 | 0.61379948 | -3.466518371 | 0.000527246 | 0.000787727 | - | glycosyltransferase family 2 protein | WP_302228549.1 | | | | |
| AVAEIV_RS12180 | 267.191039 | -2.124682675 | 0.168483962 | -12.610593 | 1.85E-36 | 7.47E-35 | *flgC* | flagellar basal body rod protein FlgC | WP_012700983.1 | Avin_24010 | *flgC* | flagellar basal-body rod protein | 100 |
| AVAEIV_RS09680 | 20914.97369 | -2.123446269 | 0.718608829 | -2.954940412 | 0.003127294 | 0.004333537 | *rmf* | ribosome modulation factor | WP_041807109.1 | Avin_18910 | | Ribosome modulation factor | 98.41 |
| AVAEIV_RS04635 | 448.5266784 | -2.123181623 | 0.162448419 | -13.0698817 | 4.89E-39 | 2.54E-37 | - | SDR family oxidoreductase | WP_012699655.1 | Avin_10000 | | Short-chain dehydrogenase/reductase SDR | 100 |
| AVAEIV_RS10900 | 81.16133327 | -2.120950358 | 0.302107678 | -7.02051127 | 2.21E-12 | 7.10E-12 | - | hypothetical protein | WP_012700712.1 | Avin_21080 | | hypothetical protein | 100 |
| AVAEIV_RS16380 | 1319.746699 | -2.11908573 | 0.173288262 | -12.22867437 | 2.18E-34 | 7.02E-33 | - | phage antirepressor N-terminal domain-containing protein | WP_302226001.1 | | | | |
| AVAEIV_RS01120 | 641.791074 | -2.117123605 | 0.212788774 | -9.949413972 | 2.54E-23 | 2.68E-22 | - | hypothetical protein | WP_302227907.1 | Avin_02270 | | hypothetical protein | 98.88 |
| AVAEIV_RS03105 | 1164.855033 | -2.116950681 | 0.710336195 | -2.980209504 | 0.002880513 | 0.004004521 | *rpsJ* | 30S ribosomal protein S10 | WP_003243886.1 | Avin_06240 | *rpsJ* | 30S ribosomal protein S10 | 100 |
| AVAEIV_RS17575 | 1545.124595 | -2.116002818 | 0.266041309 | -7.953662621 | 1.81E-15 | 7.87E-15 | - | cation acetate symporter | WP_012702032.1 | Avin_35030 | *actP* | cation/acetate symporter ActP | 100 |
| AVAEIV_RS09935 | 668.6395842 | -2.114021509 | 0.207944727 | -10.16626651 | 2.80E-24 | 3.31E-23 | - | hypothetical protein | WP_231876195.1 | Avin_19380 | | non-conserved hypothetical protein | 100 |
| AVAEIV_RS19825 | 927.8737384 | -2.113842664 | 0.537502337 | -3.93271344 | 8.40E-05 | 0.000136044 | *rplS* | 50S ribosomal protein L19 | WP_012702467.1 | Avin_39530 | *rplS* | Ribosomal protein L19 | 100 |
| AVAEIV_RS11845 | 1173.062151 | -2.113764592 | 0.430496715 | -4.910059746 | 9.10E-07 | 1.76E-06 | *sigX* | RNA polymerase sigma factor SigX | WP_041807537.1 | Avin_23320 | | RNA polymerase sigma-70 factor | 99.47 |
| AVAEIV_RS03360 | 509.8906836 | -2.108531021 | 0.194071184 | -10.86473002 | 1.70E-27 | 2.64E-26 | *thiL* | thiamine-phosphate kinase | WP_302228138.1 | Avin_06770 | *thiL* | thiamine-monophosphate kinase | 99.38 |
| AVAEIV_RS03125 | 989.3457944 | -2.108361133 | 0.304102822 | -6.933053499 | 4.12E-12 | 1.28E-11 | *rplB* | 50S ribosomal protein L2 | WP_012699305.1 | Avin_06280 | *rplB* | 50S ribosomal protein L2 | 100 |
| AVAEIV_RS06115 | 3229.773167 | -2.107826337 | 0.550406389 | -3.829581885 | 0.000128361 | 0.000203486 | - | Rpn family recombination-promoting nuclease/putative transposase | WP_072432726.1 | | | | |
| AVAEIV_RS00600 | 2835.589249 | -2.106667533 | 0.442578454 | -4.759986654 | 1.94E-06 | 3.65E-06 | *pbpG* | D-alanyl-D-alanine endopeptidase | WP_012698817.1 | Avin_01220 | | penicillin-binding protein 7 precursor | 100 |
| AVAEIV_RS12010 | 6701.198727 | -2.106515342 | 0.641367619 | -3.284411746 | 0.001021955 | 0.001485823 | - | acetyl-CoA C-acetyltransferase | WP_012700950.1 | Avin_23640 | *phbA* | polyhydroxybutyrate biosynthetic beta-ketothiolase | 100 |
| AVAEIV_RS11555 | 1786.105008 | -2.105588107 | 0.319998459 | -6.579994511 | 4.70E-11 | 1.33E-10 | - | heavy metal translocating P-type ATPase | WP_015589524.1 | Avin_22710 | | heavy metal translocating P-type ATPase protein | 99.86 |
| AVAEIV_RS06915 | 993.6666705 | -2.105167726 | 0.301040337 | -6.992975593 | 2.69E-12 | 8.56E-12 | *mraY* | phospho-N-acetylmuramoyl-pentapeptide-transferase | WP_012699968.1 | Avin_13220 | *mraY* | phospho-N-acetylmuramoyl-pentapeptide-transferase | 100 |
| AVAEIV_RS14190 | 186.2852785 | -2.104677145 | 0.276267827 | -7.618249162 | 2.57E-14 | 1.00E-13 | - | DUF2970 domain-containing protein | WP_175556026.1 | Avin_27500 | | conserved hypothetical protein | 100 |
| AVAEIV_RS01395 | 666.7088821 | -2.103101365 | 0.23485063 | -8.955059492 | 3.40E-19 | 2.18E-18 | - | exodeoxyribonuclease III | WP_012698975.1 | Avin_02870 | *crc* | catabolite repression control protein; Crc | 100 |
| AVAEIV_RS00725 | 550.0491829 | -2.101258668 | 0.272641043 | -7.707051886 | 1.29E-14 | 5.18E-14 | - | CCE_0567 family metalloprotein | WP_012698842.1 | Avin_01500 | | conserved hypothetical protein | 100 |
| AVAEIV_RS22980 | 652.6022976 | -2.100884317 | 0.131045965 | -16.03165976 | 7.68E-58 | 1.76E-55 | *trpD* | anthranilate phosphoribosyltransferase | WP_012703084.1 | Avin_46120 | *trpD* | Anthranilate phosphoribosyl transferase | 100 |
| AVAEIV_RS19380 | 2829.899717 | -2.099858933 | 0.538743157 | -3.8976995 | 9.71E-05 | 0.000156602 | - | helix-turn-helix domain-containing protein | WP_302226760.1 | | | | |
| AVAEIV_RS20055 | 372.5419576 | -2.099442119 | 0.2807883 | -7.476957253 | 7.61E-14 | 2.84E-13 | - | hypothetical protein | WP_012702516.1 | Avin_40030 | | hypothetical protein | 100 |

| | | | | | | | | | | | | | |
|---|---|---|---|---|---|---|---|---|---|---|---|---|---|
| AVAEIV_RS04665 | 241.8017958 | -2.09772775 | 0.17522725 | -11.97146991 | 5.01E-33 | 1.44E-31 | *cueR* | Cu(I)-responsive transcriptional regulator | WP_302228282.1 | Avin_10060 | *cueR* | Cu(I)-responsive transcriptional regulator, MerR family | 99.28 |
| AVAEIV_RS24190 | 940.5769344 | -2.089893931 | 0.575283824 | -3.632804961 | 0.000280357 | 0.000430629 | *rpmB* | 50S ribosomal protein L28 | WP_012703271.1 | Avin_48050 | *rpmB* | ribosomal protein L28 | 100 |
| AVAEIV_RS00580 | 884.2783075 | -2.08546189 | 0.204370939 | -10.20429762 | 1.90E-24 | 2.26E-23 | - | asparaginase | WP_302227857.1 | Avin_01180 | | L-Asparaginase, type I | 99.12 |
| AVAEIV_RS15200 | 1369.302349 | -2.084393445 | 0.160123943 | -13.01737522 | 9.75E-39 | 4.82E-37 | - | hypothetical protein | WP_302225723.1 | Avin_29470 | | hypothetical protein | 99.51 |
| AVAEIV_RS11985 | 4453.52367 | -2.083699349 | 0.527031485 | -3.953652499 | 7.70E-05 | 0.000125297 | *clpX* | ATP-dependent Clp protease ATP-binding subunit ClpX | WP_012700944.1 | Avin_23580 | *clpX* | ATP-dependent Clp protease, ATP binding subunit ClpX | 100 |
| AVAEIV_RS21425 | 3085.313271 | -2.083697126 | 0.27520643 | -7.571396965 | 3.69E-14 | 1.41E-13 | *infB* | translation initiation factor IF-2 | WP_012702773.1 | Avin_42820 | *infB* | translation initiation factor IF-2 | 100 |
| AVAEIV_RS19010 | 328.1087012 | -2.07561134 | 0.205994941 | -10.07603064 | 7.05E-24 | 7.95E-23 | *fabZ* | 3-hydroxyacyl-ACP dehydratase FabZ | WP_012702403.1 | Avin_38880 | *fabZ* | beta-hydroxyacyl-(acyl-carrier-protein) dehydratase FabZ | 100 |
| AVAEIV_RS09230 | 1671.650955 | -2.07431911 | 0.499785416 | -4.150419447 | 3.32E-05 | 5.60E-05 | - | hypothetical protein | WP_041807526.1 | Avin_17980 | | conserved hypothetical protein | 100 |
| AVAEIV_RS13155 | 1277.441511 | -2.074317987 | 0.207876253 | -9.978619264 | 1.89E-23 | 2.03E-22 | *hutU* | urocanate hydratase | WP_302225187.1 | Avin_26180 | *hutU* | urocanate hydratase | 99.28 |
| AVAEIV_RS25320 | 599.9815233 | -2.072298915 | 0.258301688 | -8.022785014 | 1.03E-15 | 4.63E-15 | - | DotU family type IV/VI secretion system protein | WP_012703517.1 | Avin_50770 | | conserved hypothetical protein | 100 |
| AVAEIV_RS20890 | 689.4948705 | -2.071875776 | 0.128255911 | -16.15423229 | 1.06E-58 | 2.60E-56 | - | sigma-54-dependent Fis family transcriptional regulator | WP_050764397.1 | Avin_41690 | *acoR* | Sigma54-dependent transcriptional activator protein, AcoR | 99.84 |
| AVAEIV_RS10610 | 1150.643202 | -2.069872918 | 0.268024097 | -7.722712026 | 1.14E-14 | 4.61E-14 | *hpnI* | bacteriohopanetetrol glucosamine biosynthesis glycosyltransferase HpnI | WP_012700674.1 | Avin_20650 | | Glycosyl transferase, family 2 | 100 |
| AVAEIV_RS03195 | 432.885425 | -2.069867415 | 0.223666033 | -9.254276939 | 2.16E-20 | 1.61E-19 | *rpsE* | 30S ribosomal protein S5 | WP_012699319.1 | Avin_06420 | *rpsE* | 30S ribosomal protein S5 | 100 |
| AVAEIV_RS09065 | 2074.895913 | -2.068269876 | 0.417017264 | -4.959674465 | 7.06E-07 | 1.38E-06 | *zwf* | glucose-6-phosphate dehydrogenase | WP_302228639.1 | Avin_17630 | *zwf-2* | Glucose-6-phosphate dehydrogenase | 98.98 |
| AVAEIV_RS16985 | 217.8706874 | -2.066378994 | 0.205831113 | -10.03919653 | 1.03E-23 | 1.13E-22 | - | transposase | pseudo=true | | | | |
| AVAEIV_RS08080 | 454.630179 | -2.066375437 | 0.236670201 | -8.731033414 | 2.52E-18 | 1.46E-17 | - | glucokinase | WP_012700200.1 | Avin_15690 | *glk-1* | glucokinase | 100 |
| AVAEIV_RS15345 | 1789.594759 | -2.065395263 | 0.347464127 | -5.944197119 | 2.78E-09 | 6.70E-09 | *sdhA* | succinate dehydrogenase flavoprotein subunit | WP_041807252.1 | Avin_29790 | *sdhA* | succinate dehydrogenase flavoprotein subunit | 99.77 |
| AVAEIV_RS08055 | 597.9509506 | -2.065027432 | 0.243378283 | -8.484846734 | 2.16E-17 | 1.13E-16 | - | PHP domain-containing protein | WP_012700195.1 | Avin_15640 | | Phosphoesterase (PHP family) | 100 |
| AVAEIV_RS07715 | 873.2434916 | -2.058278406 | 0.666739181 | -3.087081824 | 0.002021319 | 0.002846235 | *acpP* | acyl carrier protein | WP_003283670.1 | Avin_14930 | *acpP* | acyl carrier protein | 100 |
| AVAEIV_RS10455 | 2398.477427 | -2.055641259 | 0.522655486 | -3.933071233 | 8.39E-05 | 0.000135885 | *ihfA* | integration host factor subunit alpha | WP_011913482.1 | Avin_20480 | *ihfA* | integration host factor, alpha subunit | 100 |
| AVAEIV_RS11975 | 1660.186298 | -2.051761563 | 0.343931059 | -5.965618713 | 2.44E-09 | 5.93E-09 | *tig* | trigger factor | WP_061288365.1 | Avin_23550 | *tig* | trigger factor | 99.77 |
| AVAEIV_RS25680 | 3052.589167 | -2.049105671 | 0.419187467 | -4.888279905 | 1.02E-06 | 1.96E-06 | - | efflux transporter outer membrane subunit | WP_302227748.1 | Avin_51570 | | multidrug efflux pump RND-family outer membrane protein | 99.8 |
| AVAEIV_RS24055 | 274.0536947 | -2.048762492 | 0.221420177 | -9.252826547 | 2.19E-20 | 1.63E-19 | - | secondary thiamine-phosphate synthase enzyme YjbQ | WP_012703243.1 | Avin_47770 | | Conserved hypothetical protein | 100 |
| AVAEIV_RS16780 | 353.1312779 | -2.044506639 | 0.232591019 | -8.790135781 | 1.49E-18 | 8.89E-18 | - | SprT family zinc-dependent metalloprotease | WP_012701863.1 | Avin_33250 | | conserved hypothetical protein | 100 |
| AVAEIV_RS24380 | 1899.091561 | -2.042813008 | 0.518633231 | -3.938839411 | 8.19E-05 | 0.000132869 | - | HU family DNA-binding protein | WP_012703311.1 | Avin_48480 | *hupA* | histone-like bacterial DNA-binding protein | 100 |
| AVAEIV_RS11610 | 602.0834598 | -2.041903768 | 0.184730419 | -11.05342466 | 2.11E-28 | 3.64E-27 | - | sensor domain-containing diguanylate cyclase | WP_302229013.1 | Avin_22780 | | GGDEF protein | 99.37 |
| AVAEIV_RS21615 | 162.5283756 | -2.039767349 | 0.4598441 | -4.435780195 | 9.17E-06 | 1.62E-05 | - | hypothetical protein | WP_012700810.1 | | | | |
| AVAEIV_RS21610 | 1026.309007 | -2.038788385 | 0.175878869 | -11.59200305 | 4.52E-31 | 1.02E-29 | *pyk* | pyruvate kinase | WP_302227246.1 | | | | |
| AVAEIV_RS18420 | 410.4801977 | -2.034337534 | 0.442090745 | -4.601628869 | 4.19E-06 | 7.65E-06 | - | FxLYD domain-containing protein | WP_231876249.1 | Avin_37110 | | hypothetical protein | 100 |
| AVAEIV_RS22515 | 3079.899165 | -2.033443405 | 0.233005267 | -8.727027652 | 2.61E-18 | 1.51E-17 | *gltB* | glutamate synthase large subunit | WP_012703003.1 | Avin_45210 | *gltB* | Glutamate synthase large subunit protein | 100 |
| AVAEIV_RS08140 | 1953.29101 | -2.03342731 | 0.183087243 | -11.10632984 | 1.17E-28 | 2.06E-27 | *gyrA* | DNA gyrase subunit A | WP_012700211.1 | Avin_15810 | *gyrA* | DNA gyrase, A subunit | 100 |
| AVAEIV_RS11980 | 792.843233 | -2.03262501 | 0.498256101 | -4.079478415 | 4.51E-05 | 7.50E-05 | *clpP* | ATP-dependent Clp endopeptidase proteolytic subunit ClpP | WP_041807539.1 | Avin_23570 | *clpP* | Peptidase S14, ClpP | 100 |
| AVAEIV_RS07905 | 319.1909588 | -2.030355021 | 0.258168909 | -7.864444373 | 3.71E-15 | 1.57E-14 | - | DUF2790 domain-containing protein | WP_231876277.1 | Avin_15330 | | conserved hypothetical protein | 100 |
| AVAEIV_RS14085 | 2333.69846 | -2.029929552 | 0.425451536 | -4.77123569 | 1.83E-06 | 3.46E-06 | *zwf* | glucose-6-phosphate dehydrogenase | WP_012701290.1 | Avin_27260 | *zwf-1* | Glucose-6-phosphate dehydrogenase | 100 |
| AVAEIV_RS12365 | 873.9937818 | -2.028071228 | 0.220854933 | -9.182820589 | 4.20E-20 | 2.99E-19 | - | PAS domain-containing methyl-accepting chemotaxis protein | WP_012701020.1 | Avin_24380 | | aerotaxis receptor Aer | 100 |
| AVAEIV_RS02825 | 334.8509351 | -2.027859783 | 0.26045374 | -7.785873154 | 6.92E-15 | 2.87E-14 | - | GreA/GreB family elongation factor | WP_302228093.1 | Avin_05670 | | conserved hypothetical protein | 99.38 |
| AVAEIV_RS13470 | 185.4505179 | -2.026210419 | 0.263106726 | -7.701096988 | 1.35E-14 | 5.41E-14 | - | hypothetical protein | pseudo=true | | | | |
| AVAEIV_RS16710 | 105.0809732 | -2.025677401 | 0.197971768 | -10.23215293 | 1.42E-24 | 1.73E-23 | *bluB* | 5,2C6-dimethylbenzimidazole synthase | WP_302229379.1 | Avin_33110 | | cob(II)yrinic acid a,c-diamide reductase protein | 99.54 |
| AVAEIV_RS08755 | 304.2218142 | -2.023137488 | 0.479363637 | -4.220465072 | 2.44E-05 | 4.15E-05 | - | helix-turn-helix domain-containing protein | WP_302228596.1 | | | | |
| AVAEIV_RS17640 | 1605.184585 | -2.022657875 | 0.354934625 | -5.698677255 | 1.21E-08 | 2.75E-08 | - | zinc-binding dehydrogenase | WP_012702045.1 | Avin_35160 | | Zinc-containing alcohol dehydrogenase-like protein | 100 |
| AVAEIV_RS09980 | 1063.359706 | -2.020521614 | 0.397494809 | -5.083139617 | 3.71E-07 | 7.45E-07 | *pdxH* | pyridoxamine 5'-phosphate oxidase | WP_012700568.1 | Avin_19500 | *pdxH* | pyridoxamine 5-phosphate oxidase | 100 |
| AVAEIV_RS22525 | 800.8857793 | -2.019583283 | 0.154745691 | -13.05098234 | 6.27E-39 | 3.19E-37 | *aroB* | 3-dehydroquinate synthase | WP_012703005.1 | Avin_45240 | *aroB* | 3-dehydroquinate synthase | 100 |
| AVAEIV_RS14100 | 1113.063192 | -2.014461429 | 0.29001904 | -6.94596267 | 3.76E-12 | 1.18E-11 | - | LacI family DNA-binding transcriptional regulator | WP_302225465.1 | Avin_27300 | | Transcriptional regulator, LacI family | 99.71 |
| AVAEIV_RS03825 | 377.8587083 | -2.013169017 | 0.227841443 | -8.835833321 | 9.94E-19 | 6.05E-18 | - | DUF4345 family protein | WP_302228193.1 | Avin_07790 | | conserved hypothetical protein | 99.4 |
| AVAEIV_RS22635 | 234.4377131 | -2.011721756 | 0.465517062 | -4.321478032 | 1.55E-05 | 2.69E-05 | - | twin-arginine translocase TatA/TatE family subunit | WP_012703027.1 | Avin_45460 | *tatA* | Twin-arginine translocation protein | 100 |
| AVAEIV_RS25390 | 292.5045729 | -2.011536504 | 0.137568154 | -14.62210872 | 2.03E-48 | 2.09E-46 | - | NifB/NifX family molybdenum-iron cluster-binding protein | WP_012703530.1 | Avin_50900 | | nitrogen fixation-related protein | 100 |
| AVAEIV_RS20160 | 725.0200202 | -2.009392423 | 0.198561724 | -10.11973699 | 4.52E-24 | 5.16E-23 | *ispG* | flavodoxin-dependent (E)-4-hydroxy-3-methylbut-2-enyl-diphosphate synthase | WP_012702540.1 | Avin_40290 | *ispG* | 4-hydroxy-3-methylbut-2-en-1-yl diphosphate synthase | 100 |
| AVAEIV_RS07090 | 575.1498836 | -2.009301456 | 0.203121889 | -9.892097156 | 4.50E-23 | 4.61E-22 | *ung* | uracil-DNA glycosylase | WP_012700002.1 | Avin_13610 | *ung* | uracil-DNA glycosylase | 100 |
| AVAEIV_RS04080 | 157.1761858 | -2.008864199 | 0.227679205 | -8.823222131 | 1.11E-18 | 6.73E-18 | *rsfS* | ribosome silencing factor | WP_061290624.1 | Avin_08370 | | iojap-related protein | 99.15 |
| AVAEIV_RS11535 | 452.9067109 | -2.00746525 | 0.432384961 | -4.642773064 | 3.44E-06 | 6.32E-06 | - | ferritin-like domain-containing protein | WP_012700856.1 | Avin_22670 | | hypothetical protein | 100 |
| AVAEIV_RS07125 | 1959.611776 | -2.006907378 | 0.396811849 | -5.057579261 | 4.25E-07 | 8.45E-07 | - | MucB/RseB C-terminal domain-containing protein | WP_302228496.1 | Avin_13710 | *mucB* | sigma factor algU negative regulatory protein MucB | 99.69 |
| AVAEIV_RS08535 | 352.9778408 | -2.006191746 | 0.266550064 | -7.526510099 | 5.21E-14 | 1.97E-13 | - | DUF411 domain-containing protein | WP_415860969.1 | | | | |
| AVAEIV_RS16980 | 1121.473433 | -2.005884529 | 0.216338875 | -9.271955992 | 1.83E-20 | 1.38E-19 | - | YMGG-like glycine zipper-containing protein | WP_231876234.1 | | | | |
| AVAEIV_RS25330 | 301.9915229 | -2.000440616 | 0.246358676 | -8.120033164 | 4.66E-16 | 2.16E-15 | - | hypothetical protein | WP_012703519.1 | Avin_50790 | | conserved hypothetical protein | 100 |
| AVAEIV_RS12360 | 1197.822566 | -2.000323803 | 0.263899082 | -7.579881637 | 3.46E-14 | 1.33E-13 | - | methyl-accepting chemotaxis protein | WP_302229109.1 | Avin_24370 | | methyl accepting chemotaxis sensory transducer | 99.48 |
| AVAEIV_RS13770 | 114.2125003 | 2.000413532 | 0.260744291 | 7.671936061 | 1.69E-14 | 6.74E-14 | - | dienelactone hydrolase family protein | WP_302225363.1 | Avin_26540 | | Conserved hypothetical protein | 99.64 |
| AVAEIV_RS02895 | 193.1972956 | 2.001890882 | 0.196169403 | 10.20490885 | 1.88E-24 | 2.25E-23 | - | MFS transporter | WP_012699265.1 | Avin_05840 | | tartrate transporter | 100 |
| AVAEIV_RS18275 | 26.78529917 | 2.006129965 | 0.379267406 | 5.289486873 | 1.23E-07 | 2.57E-07 | - | hypothetical protein | WP_302226506.1 | | | | |
| AVAEIV_RS16165 | 103.1212892 | 2.006470728 | 0.218618809 | 9.177941923 | 4.39E-20 | 3.12E-19 | - | MetQ/NlpA family ABC transporter substrate-binding protein | WP_302225971.1 | Avin_31510 | | NLPA lipoprotein | 99.63 |
| AVAEIV_RS10540 | 164.5139603 | 2.007136829 | 0.251526027 | 7.979837532 | 1.47E-15 | 6.47E-15 | - | AAA family ATPase | WP_302228820.1 | | | | |
| AVAEIV_RS21085 | 123.44475 | 2.009204708 | 0.244277248 | 8.225099639 | 1.95E-16 | 9.38E-16 | - | NAD(P)H-dependent flavin oxidoreductase | WP_012702711.1 | Avin_42160 | | 2-nitropropane dioxygenase-like dioxygenase | 100 |
| AVAEIV_RS15545 | 143.3460891 | 2.009215791 | 0.297351237 | 6.757045337 | 1.41E-11 | 4.17E-11 | - | Gfo/Idh/MocA family protein | WP_302225801.1 | Avin_30210 | | oxidoreductase protein | 98.86 |
| AVAEIV_RS15450 | 82.33482831 | 2.00978535 | 0.269142799 | 7.467356947 | 8.18E-14 | 3.05E-13 | - | glycosyltransferase | WP_061291789.1 | Avin_29990 | | Glycosyl transferase, group 1 family protein | 98.68 |
| AVAEIV_RS16175 | 73.38191398 | 2.011627384 | 0.262723774 | 7.656815193 | 1.91E-14 | 7.54E-14 | - | methionine ABC transporter permease | WP_012701701.1 | Avin_31530 | | transmembrane ABC transporter protein | 100 |
| AVAEIV_RS04260 | 62.89541122 | 2.013624122 | 0.292109628 | 6.893384988 | 5.45E-12 | 1.68E-11 | - | 2-hydroxymuconate tautomerase | WP_012699545.1 | Avin_08790 | *dmpI* | 4-oxalocrotonate tautomerase | 100 |
| AVAEIV_RS04065 | 200.8349332 | 2.01478202 | 0.225004641 | 8.954402043 | 3.42E-19 | 2.19E-18 | - | bifunctional DedA family/phosphatase PAP2 family protein | WP_012699504.1 | Avin_08340 | | Acid phosphatase/vanadium-dependent haloperoxidase superfamily | 100 |
| AVAEIV_RS18290 | 24.92140327 | 2.016128684 | 0.393230012 | 5.127097679 | 2.94E-07 | 5.98E-07 | - | hypothetical protein | WP_302226512.1 | | | | |
| AVAEIV_RS08845 | 182.2299733 | 2.016411932 | 0.255985531 | 7.877054302 | 3.35E-15 | 1.42E-14 | - | CitMHS family transporter | WP_012700335.1 | Avin_17120 | | Citrate transporter | 100 |
| AVAEIV_RS16085 | 121.8135574 | 2.016823877 | 0.22849671 | 8.82648979 | 1.08E-18 | 6.56E-18 | - | LLM class flavin-dependent oxidoreductase | WP_012701681.1 | Avin_31330 | | alkanesulfonate monooxygenase | 100 |
| AVAEIV_RS07795 | 77.8018961 | 2.018091142 | 0.281329069 | 7.173418483 | 7.31E-13 | 2.48E-12 | *mhpD* | 2-keto-4-pentenoate hydratase | WP_302228527.1 | Avin_15110 | *mhpD* | 4-oxalocrotonate decarboxylase | 99.62 |

| | | | | | | | | | | | | | |
|---|---|---|---|---|---|---|---|---|---|---|---|---|---|
| AVAEIV_RS15435 | 167.5107598 | 2.019067456 | 0.205105709 | 9.844033409 | 7.27E-23 | 7.20E-22 | - | UDP-glucose dehydrogenase family protein | WP_302225772.1 | Avin_29960 | | UDP-glucose/GDP-mannose dehydrogenase protein | 99.77 |
| AVAEIV_RS23750 | 137.2729394 | 2.01949319 | 0.226785216 | 8.904871417 | 5.34E-19 | 3.36E-18 | - | FecR domain-containing protein | WP_302227584.1 | Avin_47180 | | Anti-sigma factor, FecR family | 99.04 |
| AVAEIV_RS21790 | 77.58190692 | 2.021397733 | 0.256071308 | 7.893886078 | 2.93E-15 | 1.25E-14 | - | sugar ABC transporter substrate-binding protein | WP_302227281.1 | Avin_43620 | | ribose ABC transporter | 98.81 |
| AVAEIV_RS21750 | 81.68162662 | 2.021757826 | 0.24008086 | 8.421153709 | 3.73E-17 | 1.91E-16 | - | AMP-binding protein | WP_302227271.1 | Avin_43550 | | long-chain-fatty-acid--CoA ligase-related protein | 99.22 |
| AVAEIV_RS13230 | 197.5542761 | 2.022192392 | 0.265584427 | 7.614122608 | 2.65E-14 | 1.03E-13 | - | tyrosine-type recombinase/integrase | pseudo=true | | | | |
| AVAEIV_RS21745 | 65.79584322 | 2.024127405 | 0.306261483 | 6.609147794 | 3.87E-11 | 1.10E-10 | - | ABC transporter ATP-binding protein | WP_012702842.1 | Avin_43540 | | ABC transporter, ATP binding component | 100 |
| AVAEIV_RS23035 | 168.2028341 | 2.026835474 | 0.254320774 | 7.969602485 | 1.59E-15 | 6.99E-15 | - | phage portal protein | WP_231876270.1 | | | | |
| AVAEIV_RS12700 | 67.8320601 | 2.029330273 | 0.335524306 | 6.048236255 | 1.46E-09 | 3.63E-09 | - | DUF485 domain-containing protein | WP_012701100.1 | Avin_25240 | | DUF485 family protein | 100 |
| AVAEIV_RS15505 | 99.43253237 | 2.029582133 | 0.305124376 | 6.651655177 | 2.90E-11 | 8.40E-11 | - | glycosyltransferase family 2 protein | WP_012701563.1 | Avin_30100 | | Glycosyl transferase, family 2 protein | 100 |
| AVAEIV_RS13720 | 129.2672471 | 2.029799349 | 0.200649255 | 10.11615692 | 4.68E-24 | 5.33E-23 | - | FecCD family ABC transporter permease | WP_012701211.1 | Avin_26440 | | iron(III) ABC transporter-permease protein | 100 |
| AVAEIV_RS16170 | 135.8035067 | 2.030314797 | 0.253997075 | 7.993457402 | 1.31E-15 | 5.82E-15 | - | methionine ABC transporter ATP-binding protein | WP_012701700.1 | Avin_31520 | | ABC transporter, ATP-binding component | 100 |
| AVAEIV_RS23390 | 62.13911729 | 2.031486918 | 0.322802816 | 6.293275089 | 3.11E-10 | 8.17E-10 | - | universal stress protein | WP_012703110.1 | Avin_46410 | | universal stress protein family | 100 |
| AVAEIV_RS05275 | 268.7839156 | 2.032320178 | 0.14667906 | 13.85555771 | 1.18E-43 | 8.29E-42 | - | sigma-54-dependent Fis family transcriptional regulator | WP_302228345.1 | Avin_11420 | | sigma54-dependent activator protein | 99.85 |
| AVAEIV_RS08285 | 202.5471529 | 2.032577171 | 0.258325227 | 7.868287554 | 3.60E-15 | 1.52E-14 | *istA* | IS21 family transposase | pseudo=true | | | | |
| AVAEIV_RS11260 | 160.0204135 | 2.034762982 | 0.243081655 | 8.370697407 | 5.73E-17 | 2.90E-16 | - | MDR/zinc-dependent alcohol dehydrogenase-like family protein | WP_302228961.1 | Avin_22030 | | Zinc-containing alcohol dehydrogenase superfamily | 99.73 |
| AVAEIV_RS02150 | 100.1393472 | 2.035536368 | 0.261393303 | 7.787255237 | 6.85E-15 | 2.84E-14 | *trfA* | plasmid replication initiator TrfA | WP_367281695.1 | | | | |
| AVAEIV_RS16935 | 291.6752014 | 2.035662706 | 0.253089915 | 8.043239137 | 8.75E-16 | 3.95E-15 | - | cytochrome c | WP_012701894.1 | Avin_33640 | | dehydrogenase cytochrome c subunit | 100 |
| AVAEIV_RS11105 | 118.9003955 | 2.036233189 | 0.264406466 | 7.701147468 | 1.35E-14 | 5.41E-14 | - | FecR domain-containing protein | WP_061291569.1 | Avin_21540 | | Anti-sigma factor, FecR family | 99.69 |
| AVAEIV_RS15695 | 74.8621335 | 2.036491929 | 0.28971012 | 7.029412456 | 2.07E-12 | 6.69E-12 | *ssuC* | aliphatic sulfonate ABC transporter permease SsuC | WP_012701600.1 | Avin_30490 | *ssuC* | aliphatic sulfonate ABC transporter permease protein | 100 |
| AVAEIV_RS00235 | 95.97618918 | 2.036867276 | 0.253377464 | 8.038865178 | 9.07E-16 | 4.09E-15 | - | ATP-binding cassette domain-containing protein | WP_302227831.1 | Avin_00480 | | ABC transporter, ATP binding component | 99.69 |
| AVAEIV_RS10120 | 113.8716832 | 2.041299413 | 0.26409896 | 7.729297436 | 1.08E-14 | 4.39E-14 | - | phosphonate ABC transporter ATP-binding protein | WP_012700596.1 | Avin_19780 | | Phosphonate ABC transporter, ATP binding component | 100 |
| AVAEIV_RS21805 | 113.639066 | 2.041589793 | 0.218609904 | 9.33896294 | 9.73E-21 | 7.46E-20 | - | SfnB family sulfur acquisition oxidoreductase | WP_302227285.1 | Avin_43650 | | Acyl-CoA dehydrogenase | 99.28 |
| AVAEIV_RS18295 | 70.77028624 | 2.0445621 | 0.278531349 | 7.340509806 | 2.13E-13 | 7.63E-13 | - | hypothetical protein | WP_302226514.1 | | | | |
| AVAEIV_RS25695 | 106.2496448 | 2.046812196 | 0.222521224 | 9.198278517 | 3.64E-20 | 2.61E-19 | - | CoA transferase subunit B | WP_012703590.1 | Avin_51600 | | 3-oxoacid CoA-transferase | 100 |
| AVAEIV_RS23370 | 124.9240549 | 2.046978248 | 0.198949504 | 10.28893367 | 7.90E-25 | 9.75E-24 | - | LysR family transcriptional regulator | WP_012703106.1 | Avin_46370 | | Transcriptional regulator, LysR-family | 100 |
| AVAEIV_RS06065 | 200.2738829 | 2.047688452 | 0.29833943 | 6.863619905 | 6.71E-12 | 2.05E-11 | - | hypothetical protein | WP_072432735.1 | Avin_11030 | | hypothetical protein | 100 |
| AVAEIV_RS16100 | 70.36134578 | 2.04819022 | 0.258215103 | 7.932108518 | 2.15E-15 | 9.32E-15 | - | ABC transporter substrate-binding protein | WP_012701684.1 | Avin_31360 | | ABC transporter periplasmic aliphatic sulfonate-binding protein | 100 |
| AVAEIV_RS20360 | 66.18697493 | 2.048231792 | 0.296307258 | 6.912526568 | 4.76E-12 | 1.48E-11 | - | hypothetical protein | WP_012702576.1 | Avin_40680 | | hypothetical protein | 100 |
| AVAEIV_RS01790 | 102.4996705 | 2.052817304 | 0.225507178 | 9.103112908 | 8.78E-20 | 6.03E-19 | - | LLM class flavin-dependent oxidoreductase | WP_012699059.1 | Avin_03710 | | Alkanesulfonate monooxygenase | 100 |
| AVAEIV_RS00240 | 77.67292906 | 2.053108491 | 0.282100095 | 7.277943269 | 3.39E-13 | 1.19E-12 | - | sulfotransferase family 2 domain-containing protein | WP_302227832.1 | Avin_00490 | | hypothetical protein | 98.33 |
| AVAEIV_RS01355 | 82.44775219 | 2.056230184 | 0.266603346 | 7.712694587 | 1.23E-14 | 4.97E-14 | - | NifU family protein | WP_012698967.1 | Avin_02790 | *vnfU* | NifU C-terminal domain-containing protein, VnfU | 100 |
| AVAEIV_RS08935 | 177.6423421 | 2.057646934 | 0.244203056 | 8.42596719 | 3.58E-17 | 1.84E-16 | - | PAS domain-containing sensor histidine kinase | WP_012700357.1 | Avin_17340 | | intracellular sensory histidine protein kinase, two component | 100 |
| AVAEIV_RS11650 | 146.8003964 | 2.059871052 | 0.291272214 | 7.071979239 | 1.53E-12 | 5.02E-12 | - | general secretion pathway protein GspB | WP_012700876.1 | Avin_22870 | | hypothetical protein | 100 |
| AVAEIV_RS17890 | 87.32407444 | 2.060300292 | 0.273047735 | 7.545568155 | 4.50E-14 | 1.70E-13 | - | integrating conjugative element protein | WP_415860934.1 | | | | |
| AVAEIV_RS17035 | 134.7418819 | 2.060898464 | 0.234642039 | 8.783159557 | 1.59E-18 | 9.41E-18 | - | efflux RND transporter periplasmic adaptor subunit | WP_302226174.1 | Avin_33860 | *mexE* | multidrug efflux RND membrane protein, HlyD family | 99.76 |
| AVAEIV_RS19440 | 128.7021798 | 2.06208143 | 0.238184583 | 8.657493277 | 4.82E-18 | 2.71E-17 | - | helix-turn-helix domain-containing protein | WP_302226778.1 | | | | |
| AVAEIV_RS05410 | 42.06910186 | 2.062791461 | 0.327050312 | 6.307260328 | 2.84E-10 | 7.50E-10 | - | hypothetical protein | WP_012699821.1 | Avin_11710 | | hypothetical protein | 100 |
| AVAEIV_RS17925 | 145.4608679 | 2.064449 | 0.207344893 | 9.956594403 | 2.36E-23 | 2.50E-22 | - | hypothetical protein | WP_072432881.1 | | | | |
| AVAEIV_RS22045 | 164.8635444 | 2.067773134 | 0.2527445 | 8.18127845 | 2.81E-16 | 1.33E-15 | - | DUF2955 domain-containing protein | WP_302227345.1 | Avin_44190 | | membrane protein | 99.42 |
| AVAEIV_RS25040 | 141.6200131 | 2.068134424 | 0.191346565 | 10.80831746 | 3.14E-27 | 4.78E-26 | *hppD* | 4-hydroxyphenylpyruvate dioxygenase | WP_302227661.1 | Avin_50170 | *hppD* | 4-hydroxyphenylpyruvate dioxygenase | 99.13 |
| AVAEIV_RS06550 | 244.0680978 | 2.069298192 | 0.173572086 | 11.92183742 | 9.11E-33 | 2.49E-31 | - | ABC transporter substrate-binding protein | WP_012699899.1 | Avin_12520 | | dipeptide ABC transporter, periplasmic substrate-binding component | 100 |
| AVAEIV_RS17900 | 298.3489967 | 2.070803265 | 0.239353362 | 8.651657298 | 5.08E-18 | 2.84E-17 | - | TIGR03747 family integrating conjugative element membrane protein | WP_072432876.1 | | | | |
| AVAEIV_RS25940 | 328.8179265 | 2.071736751 | 0.18531435 | 11.17958081 | 5.13E-29 | 9.56E-28 | - | di-heme oxidoredictase family protein | WP_061290850.1 | Avin_52140 | | thiol oxidoreductase | 98.67 |
| AVAEIV_RS03995 | 159.9247837 | 2.072070639 | 0.230325868 | 8.996256716 | 2.34E-19 | 1.53E-18 | - | ABC transporter substrate-binding protein | WP_302228208.1 | Avin_08170 | | sulfate ester transport system substrate-binding protein | 99.43 |
| AVAEIV_RS13090 | 182.99066 | 2.073511772 | 0.252330157 | 8.217455243 | 2.08E-16 | 9.95E-16 | - | LysR family transcriptional regulator | WP_012701170.1 | Avin_26010 | | Transcriptional regulator, LysR family | 100 |
| AVAEIV_RS16050 | 141.137484 | 2.075422151 | 0.202973192 | 10.22510474 | 1.53E-24 | 1.86E-23 | - | gallate dioxygenase | WP_012701674.1 | Avin_31250 | | Extradiol ring-cleavage dioxygenase, class III protein | 100 |
| AVAEIV_RS16045 | 181.8446815 | 2.077404139 | 0.237089566 | 8.762106961 | 1.92E-18 | 1.12E-17 | - | MFS transporter | WP_061291631.1 | Avin_31240 | | Major facilitator superfamily protein | 99.78 |
| AVAEIV_RS17085 | 174.0367818 | 2.078285965 | 0.264288385 | 7.863705261 | 3.73E-15 | 1.58E-14 | - | polyprenyl synthetase family protein | WP_231876231.1 | Avin_33990 | | geranylgeranyl pyrophosphate synthase protein | 99.38 |
| AVAEIV_RS08365 | 194.5004811 | 2.078987483 | 0.254582522 | 8.166261629 | 3.18E-16 | 1.50E-15 | - | MFS transporter | WP_156483631.1 | Avin_16310 | | major facilitator superfamily transporter | 99.75 |
| AVAEIV_RS02080 | 165.4535763 | 2.079571113 | 0.316623253 | 6.56796711 | 5.10E-11 | 1.44E-10 | - | hypothetical protein | WP_302227989.1 | | | | |
| AVAEIV_RS24630 | 272.975046 | 2.081197465 | 0.190409902 | 10.93009051 | 8.28E-28 | 1.34E-26 | *anfD* | nitrogenase iron-iron protein,2C alpha chain | WP_012703361.1 | Avin_48990 | *anfD* | nitrogenase iron-iron protein, alpha chain | 100 |
| AVAEIV_RS17030 | 125.0597393 | 2.081346305 | 0.225870125 | 9.214792369 | 3.12E-20 | 2.26E-19 | - | LysR family transcriptional regulator | WP_012701915.1 | Avin_33850 | *mexT* | Transcriptional regulator MexT | 100 |
| AVAEIV_RS08805 | 17.07625018 | 2.08996326 | 0.469494108 | 4.451521811 | 8.53E-06 | 1.51E-05 | - | helix-turn-helix transcriptional regulator | WP_012700326.1 | Avin_17020 | | Prophage CP4-57 regulatory, AlpA-like protein | 100 |
| AVAEIV_RS24740 | 219.2401534 | 2.091097825 | 0.2061363 | 10.14424835 | 3.51E-24 | 4.04E-23 | - | IlvD/Edd family dehydratase | WP_302227636.1 | Avin_49250 | | L-arabonate dehydratase | 98.96 |
| AVAEIV_RS21730 | 110.435684 | 2.091896846 | 0.22180078 | 9.431422403 | 4.05E-21 | 3.32E-20 | - | branched-chain amino acid ABC transporter permease | WP_012702840.1 | Avin_43520 | | amino acid ABC transporter-like protein | 100 |
| AVAEIV_RS01810 | 116.1659539 | 2.092374857 | 0.218098654 | 9.593708248 | 8.50E-22 | 7.54E-21 | - | ABC transporter substrate-binding protein | WP_012699064.1 | Avin_03760 | | ABC transporter, substrate-binding protein, aliphatic sulphonate | 100 |
| AVAEIV_RS17790 | 77.31242696 | 2.092705225 | 0.259484816 | 8.064846553 | 7.33E-16 | 3.35E-15 | - | peptide-methionine (S)-S-oxide reductase | WP_081434900.1 | | | | |
| AVAEIV_RS23135 | 94.44828006 | 2.093766117 | 0.247771891 | 8.450377907 | 2.90E-17 | 1.50E-16 | - | baseplate J/gp47 family protein | WP_302227516.1 | | | | |
| AVAEIV_RS25050 | 187.5740962 | 2.094348321 | 0.23891988 | 8.765902262 | 1.85E-18 | 1.09E-17 | *fahA* | fumarylacetoacetase | WP_302227662.1 | Avin_50190 | *fahA* | fumarylacetoacetase | 99.78 |
| AVAEIV_RS21755 | 101.114872 | 2.094730224 | 0.288441387 | 7.262238779 | 3.81E-13 | 1.33E-12 | *sfnG* | dimethylsulfone monooxygenase SfnG | WP_012702844.1 | Avin_43560 | *sfnG* | dimethyl sulfone monooxygenase; SfnG | 100 |
| AVAEIV_RS07430 | 173.205162 | 2.095008111 | 0.212399771 | 9.863513963 | 5.99E-23 | 6.05E-22 | - | ATP-binding protein | WP_012700067.1 | Avin_14320 | | periplasmic sensory histidine protein kinase | 100 |
| AVAEIV_RS25055 | 94.58655 | 2.101816131 | 0.278626281 | 7.543495616 | 4.58E-14 | 1.73E-13 | *maiA* | maleylacetoacetate isomerase | WP_302227663.1 | Avin_50200 | *maiA* | maleylacetoacetate isomerase | 98.17 |
| AVAEIV_RS18300 | 51.21997004 | 2.104074978 | 0.319445674 | 6.586644138 | 4.50E-11 | 1.28E-10 | - | hypothetical protein | WP_302226516.1 | | | | |
| AVAEIV_RS13945 | 319.7451688 | 2.105107589 | 0.190956226 | 11.02403224 | 2.93E-28 | 4.97E-27 | - | OprD family porin | WP_302225411.1 | Avin_26920 | | OprD family outer membrane porin | 99.06 |
| AVAEIV_RS11035 | 93.00220415 | 2.105903869 | 0.26452309 | 7.961134392 | 1.70E-15 | 7.45E-15 | - | ABC transporter substrate-binding protein | WP_302228921.1 | Avin_21350 | | ABC-type Fe3+ transport system, substrate binding component | 99.71 |
| AVAEIV_RS15460 | 111.335747 | 2.107025772 | 0.235668713 | 8.940625794 | 3.87E-19 | 2.46E-18 | - | glycosyltransferase family 4 protein | WP_012701554.1 | Avin_30010 | | Conserved hypothetical protein | 100 |
| AVAEIV_RS13300 | 335.0669031 | 2.1070378 | 0.200926236 | 10.48662354 | 9.95E-26 | 1.33E-24 | - | MFS transporter | WP_302225220.1 | | | | |

| | | | | | | | | | | | | | |
|---|---|---|---|---|---|---|---|---|---|---|---|---|---|
| AVAEIV_RS25810 | 242.5901429 | 2.110117494 | 0.258605188 | 8.159610049 | 3.36E-16 | 1.58E-15 | - | glutathione S-transferase family protein | WP_302227764.1 | Avin_51860 | | Glutathione S-transferase, N-terminal:Glutathione S-transferase, C-terminal | 97.26 |
| AVAEIV_RS09105 | 96.71697907 | 2.110689505 | 0.235209971 | 8.973639564 | 2.87E-19 | 1.86E-18 | - | sigma-70 family RNA polymerase sigma factor | WP_012700394.1 | Avin_17710 | | sigma factor, FecI family | 100 |
| AVAEIV_RS11135 | 97.0245999 | 2.113989998 | 0.265253091 | 7.969709181 | 1.59E-15 | 6.99E-15 | - | ABC transporter ATP-binding protein | WP_012700759.1 | Avin_21610 | | aliphatic sulfonates ABC transporter, ATP binding component | 100 |
| AVAEIV_RS10345 | 135.9793065 | 2.119784092 | 0.21802906 | 9.722484194 | 2.42E-22 | 2.29E-21 | - | FAD-dependent oxidoreductase | WP_012700638.1 | Avin_20240 | | succinate dehydrogenase/fumarate reductase flavoprotein subunit | 100 |
| AVAEIV_RS01840 | 91.26521592 | 2.120262446 | 0.273437153 | 7.754112494 | 8.90E-15 | 3.64E-14 | *fdxH* | formate dehydrogenase subunit beta | WP_302229230.1 | Avin_03820 | *fdhH* | formate dehydrogenase, beta subunit | 98.98 |
| AVAEIV_RS05720 | 206.6640541 | 2.120349404 | 0.220464643 | 9.617639246 | 6.74E-22 | 6.09E-21 | - | hypothetical protein | WP_139240022.1 | | | | |
| AVAEIV_RS16595 | 163.0822772 | 2.120627848 | 0.237947196 | 8.912178345 | 5.00E-19 | 3.16E-18 | - | AAA family ATPase | WP_012701827.1 | Avin_32860 | | ATPase, AAA family protein | 100 |
| AVAEIV_RS08420 | 232.6962858 | 2.120641423 | 0.214147805 | 9.902699809 | 4.05E-23 | 4.17E-22 | - | electron transfer flavoprotein-ubiquinone oxidoreductase | WP_302228560.1 | Avin_17400 | | Electron-transferring-flavoprotein ubiquinone oxidoreductase | 99.64 |
| AVAEIV_RS11070 | 46.19596432 | 2.121333814 | 0.325358278 | 6.519993376 | 7.03E-11 | 1.96E-10 | - | amino acid ABC transporter permease | WP_061291058.1 | Avin_21450 | | polar amino acid ABC transporter, inner membrane permease component | 99.59 |
| AVAEIV_RS15165 | 118.7907632 | 2.122481295 | 0.245736409 | 8.637227605 | 5.76E-18 | 3.19E-17 | - | ABC transporter substrate-binding protein | WP_302225712.1 | Avin_29390 | | ABC-type nitrate/sulfonate/bicarbonate transport system,substrate binding component | 99.16 |
| AVAEIV_RS06220 | 114.989123 | 2.123716503 | 0.260946564 | 8.138511081 | 4.00E-16 | 1.86E-15 | - | DUF3158 family protein | WP_072432707.1 | | | | |
| AVAEIV_RS21940 | 96.26232816 | 2.124572367 | 0.24547117 | 8.655078981 | 4.93E-18 | 2.76E-17 | - | class II aldolase/adducin family protein | WP_012702885.1 | Avin_43970 | | class II aldolase protein | 100 |
| AVAEIV_RS21115 | 13.79116695 | 2.12480807 | 0.480194036 | 4.424894749 | 9.65E-06 | 1.70E-05 | - | extradiol ring-cleavage dioxygenase | WP_012702717.1 | Avin_42220 | | Extradiol ring-cleavage dioxygenase, class III enzyme, subunit A | 100 |
| AVAEIV_RS18225 | 105.0118014 | 2.125590896 | 0.232581412 | 9.139126277 | 6.30E-20 | 4.41E-19 | - | hypothetical protein | WP_302226486.1 | | | | |
| AVAEIV_RS12725 | 71.17366619 | 2.132944538 | 0.257739186 | 8.275592736 | 1.28E-16 | 6.31E-16 | - | DUF3325 domain-containing protein | WP_061289399.1 | Avin_25310 | | membrane protein | 99.07 |
| AVAEIV_RS10690 | 212.1740535 | 2.136015031 | 0.200258888 | 10.66626833 | 1.46E-26 | 2.11E-25 | - | SGNH/GDSL hydrolase family protein | WP_175556007.1 | Avin_20820 | | Lipolytic enzyme, G-D-S-L domain protein | 100 |
| AVAEIV_RS17415 | 283.6357732 | 2.136363547 | 0.17608885 | 12.13230451 | 7.12E-34 | 2.17E-32 | - | MFS transporter | WP_012701996.1 | Avin_34670 | | Transmembrane transport protein | 100 |
| AVAEIV_RS08735 | 37.15066438 | 2.136414972 | 0.364261382 | 5.865060303 | 4.49E-09 | 1.06E-08 | - | hypothetical protein | WP_302228593.1 | | | | |
| AVAEIV_RS21025 | 162.0675242 | 2.136477912 | 0.612748507 | 3.486712552 | 0.000488997 | 0.000732071 | - | HigA family addiction module antitoxin | WP_012702698.1 | Avin_42030 | | antitoxin protein | 100 |
| AVAEIV_RS03965 | 77.56724028 | 2.136894062 | 0.301225772 | 7.093994801 | 1.30E-12 | 4.31E-12 | - | ABC transporter ATP-binding protein | WP_302228205.1 | Avin_08090 | | ABC transporter, ATP-binding component | 99.26 |
| AVAEIV_RS06225 | 279.2440937 | 2.146494208 | 0.260321129 | 8.245562754 | 1.64E-16 | 8.03E-16 | - | PFL_4669 family integrating conjugative element protein | WP_072432706.1 | | | | |
| AVAEIV_RS01640 | 148.6893656 | 2.152542309 | 0.208663247 | 10.31586702 | 5.97E-25 | 7.42E-24 | - | methyl-accepting chemotaxis protein | WP_302227949.1 | Avin_03380 | | methyl accepting chemotaxis sensory transducer | 99.81 |
| AVAEIV_RS14460 | 290.7174547 | 2.153624475 | 0.185887702 | 11.58562104 | 4.87E-31 | 1.08E-29 | - | NAD(P)/FAD-dependent oxidoreductase | WP_415860990.1 | Avin_28030 | | HI0933-like protein | 100 |
| AVAEIV_RS26040 | 245.3328261 | 2.154631062 | 0.218811771 | 9.846961376 | 7.06E-23 | 7.05E-22 | - | phage integrase N-terminal domain-containing protein | WP_302227789.1 | Avin_52320 | | integrase | 86.36 |
| AVAEIV_RS17800 | 281.9751085 | 2.156561293 | 0.413572671 | 5.214467604 | 1.84E-07 | 3.81E-07 | - | NAD(P)-dependent alcohol dehydrogenase | WP_012702076.1 | Avin_35470 | | Zinc-containing alcohol dehydrogenase | 100 |
| AVAEIV_RS17300 | 377.5829228 | 2.156600169 | 0.245830354 | 8.77271719 | 1.74E-18 | 1.03E-17 | - | DUF4124 domain-containing protein | WP_072432481.1 | Avin_34430 | | conserved hypothetical protein | 100 |
| AVAEIV_RS19415 | 78.54939725 | 2.157438 | 0.283981908 | 7.597096648 | 3.03E-14 | 1.17E-13 | - | recombination protein NinB | WP_302226770.1 | | | | |
| AVAEIV_RS01805 | 83.31502611 | 2.161247749 | 0.268029268 | 8.063476672 | 7.42E-16 | 3.39E-15 | - | Bug family tripartite tricarboxylate transporter substrate binding protein | WP_302227956.1 | Avin_03750 | | extra-cytoplasmic solute receptor, Bug family | 99.4 |
| AVAEIV_RS02615 | 265.1923358 | 2.161526372 | 0.187478645 | 11.52945377 | 9.37E-31 | 2.03E-29 | - | aldehyde dehydrogenase (NADP(+)) | WP_302228069.1 | Avin_05180 | | NAD-dependent aldehyde dehydrogenase | 99.43 |
| AVAEIV_RS11310 | 149.0624371 | 2.163219224 | 0.257282801 | 8.407943373 | 4.17E-17 | 2.13E-16 | *tal* | transaldolase | WP_012700804.1 | | | | |
| AVAEIV_RS06060 | 390.2154528 | 2.164356528 | 0.393757541 | 5.496673209 | 3.87E-08 | 8.42E-08 | - | DUF6094 domain-containing protein | WP_072432736.1 | | | | |
| AVAEIV_RS10735 | 91.99166762 | 2.169153303 | 0.279083836 | 7.772407508 | 7.70E-15 | 3.18E-14 | - | acetone carboxylase subunit gamma | WP_012700699.1 | Avin_20930 | *acxC* | Acetone carboxylase gamma subunit; AcxC | 100 |
| AVAEIV_RS10830 | 85.250911 | 2.172387399 | 0.248852286 | 8.729626047 | 2.56E-18 | 1.48E-17 | - | hypothetical protein | WP_302228881.1 | | | | |
| AVAEIV_RS04295 | 18.34271551 | 2.172801955 | 0.498910289 | 4.355095496 | 1.33E-05 | 2.32E-05 | - | phenol hydroxylase subunit | WP_302228241.1 | Avin_08860 | *dmpK* | Phenol hydroxylase subunit DmpK | 98.82 |
| AVAEIV_RS12695 | 340.3604392 | 2.173545919 | 0.209693239 | 10.36536002 | 3.56E-25 | 4.56E-24 | - | cation acetate symporter | WP_231876261.1 | Avin_25230 | | Na+/solute symporter | 99.82 |
| AVAEIV_RS12730 | 318.1335841 | 2.175234743 | 0.168235799 | 12.92967821 | 3.06E-38 | 1.44E-36 | - | PepSY-associated TM helix domain-containing protein | WP_302229173.1 | Avin_25320 | | membrane protein | 99.81 |
| AVAEIV_RS25645 | 194.8870162 | 2.175748901 | 0.238544827 | 9.120922589 | 7.45E-20 | 5.18E-19 | - | LpxA family transferase | WP_012703579.1 | Avin_51490 | | LpxA family transferase | 100 |
| AVAEIV_RS06390 | 158.0870582 | 2.180304669 | 0.213540822 | 10.2102476 | 1.78E-24 | 2.14E-23 | - | UDP-glucose dehydrogenase family protein | WP_061288525.1 | Avin_12150 | | UDP-glucose 6-dehydrogenase | 99.78 |
| AVAEIV_RS21885 | 174.3074392 | 2.182733824 | 0.208550477 | 10.46621352 | 1.23E-25 | 1.64E-24 | - | trans-sulfuration enzyme family protein | WP_302227308.1 | Avin_43850 | *metB* | Cystathionine gamma-synthase | 99.75 |
| AVAEIV_RS25130 | 110.9439468 | 2.183156609 | 0.276735275 | 7.888971182 | 3.05E-15 | 1.30E-14 | - | condensation domain-containing protein | WP_302227669.1 | Avin_50370 | | Aspartate racemase | 99.33 |
| AVAEIV_RS26140 | 72.59437767 | 2.183736815 | 0.364465081 | 5.9916215 | 2.08E-09 | 5.07E-09 | - | hypothetical protein | WP_082765933.1 | Avin_52490 | | hypothetical protein | 94.87 |
| AVAEIV_RS11520 | 80.25288764 | 2.188467736 | 0.258145126 | 8.477664368 | 2.30E-17 | 1.20E-16 | - | FecR family protein | WP_012700853.1 | Avin_22640 | | Anti-sigma factor protein, FecR family | 100 |
| AVAEIV_RS20470 | 139.4305566 | 2.1943763 | 0.285263412 | 7.692456185 | 1.44E-14 | 5.77E-14 | - | FecR domain-containing protein | WP_302227007.1 | Avin_40930 | | Anti-sigma factor protein, FecR family | 98.07 |
| AVAEIV_RS24465 | 144.9271526 | 2.198442754 | 0.200355279 | 10.97272189 | 5.17E-28 | 8.55E-27 | - | OmpA family protein | WP_302227624.1 | Avin_48650 | | OmpA/MotB family protein | 99.62 |
| AVAEIV_RS05940 | 68.4217482 | 2.200522027 | 0.322286384 | 6.8278467 | 8.62E-12 | 2.60E-11 | - | TIGR03745 family integrating conjugative element membrane protein | WP_302228432.1 | Avin_36220 | | conserved hypothetical protein | 99.12 |
| AVAEIV_RS22540 | 94.57737485 | 2.20052406 | 0.237485811 | 9.265918022 | 1.93E-20 | 1.45E-19 | - | pilus assembly protein PilP | WP_012703008.1 | Avin_45270 | *pilP* | Pilus assembly protein | 100 |
| AVAEIV_RS19430 | 68.22574934 | 2.201189918 | 0.286763471 | 7.67597739 | 1.64E-14 | 6.53E-14 | - | hypothetical protein | WP_175556070.1 | | | | |
| AVAEIV_RS15835 | 26.11359244 | 2.20592345 | 0.385858521 | 5.716922996 | 1.08E-08 | 2.48E-08 | - | phenol hydroxylase subunit | WP_302225857.1 | Avin_30760 | *lapK* | Multi-component phenol hydoxylase, assembly subunit; LapK | 97.7 |
| AVAEIV_RS17055 | 141.1063213 | 2.20616832 | 0.24108366 | 9.15104873 | 5.64E-20 | 3.96E-19 | - | aldo/keto reductase | WP_302226180.1 | Avin_33900 | | Aldo/Keto reductase with a Twin-arginine translocation pathway signal | 99.47 |
| AVAEIV_RS21100 | 148.6087304 | 2.20868358 | 0.243108606 | 9.085172335 | 1.04E-19 | 7.05E-19 | - | SRPBCC family protein | WP_302227147.1 | Avin_42190 | | aromatic ring hydroxylating dioxygenase, alpha subunit | 99.78 |
| AVAEIV_RS24460 | 80.66039761 | 2.20878613 | 0.292486946 | 7.551742605 | 4.29E-14 | 1.63E-13 | - | DUF4398 domain-containing protein | WP_012703326.1 | Avin_48640 | | conserved hypothetical protein | 100 |
| AVAEIV_RS01260 | 115.993908 | 2.209369009 | 0.240157712 | 9.199658802 | 3.59E-20 | 2.58E-19 | - | phosphonate ABC transporter ATP-binding protein | WP_231876173.1 | Avin_02560 | | ABC transporter | 99.6 |
| AVAEIV_RS03980 | 106.3889806 | 2.21038394 | 0.269765128 | 8.193734892 | 2.53E-16 | 1.21E-15 | - | LysR family transcriptional regulator | WP_231876423.1 | Avin_08140 | | Transcriptional regulator, LysR family | 99.67 |
| AVAEIV_RS00880 | 96.93138188 | 2.211712337 | 0.286000829 | 7.7332375 | 1.05E-14 | 4.26E-14 | - | DUF2388 domain-containing protein | WP_061289820.1 | Avin_01820 | | conserved hypothetical protein | 99.04 |
| AVAEIV_RS04475 | 241.6015602 | 2.211808186 | 0.161752097 | 13.67406193 | 1.45E-42 | 9.56E-41 | - | IucA/IucC family protein | WP_302228258.1 | Avin_09310 | | siderophore biosynthesis protein, IucA/IucC family | 99.32 |
| AVAEIV_RS05485 | 25.27351032 | 2.215589076 | 0.405614564 | 5.462301592 | 4.70E-08 | 1.02E-07 | - | hypothetical protein | WP_367281696.1 | | | | |
| AVAEIV_RS25485 | 159.4586259 | 2.218729095 | 0.318732567 | 6.961099449 | 3.38E-12 | 1.06E-11 | *hmpA* | NO-inducible flavohemoprotein | WP_302227720.1 | Avin_51090 | *hmp* | Flavohemoglobin Hmp | 99.75 |
| AVAEIV_RS02800 | 141.9880743 | 2.219812908 | 0.220369824 | 10.07312555 | 7.26E-24 | 8.17E-23 | - | oxidoreductase | WP_231876204.1 | Avin_05620 | | Oxidoreductase, NAD-binding Rossmann fold, GFO_IDH_MocA family | 99.7 |
| AVAEIV_RS05910 | 124.0728625 | 2.224344341 | 0.627750081 | 3.543359703 | 0.000395063 | 0.000598063 | - | hypothetical protein | WP_072432992.1 | | | | |
| AVAEIV_RS02685 | 171.4188636 | 2.228604624 | 0.237802214 | 9.371673164 | 7.14E-21 | 5.63E-20 | - | O-antigen ligase family protein | WP_012699216.1 | Avin_05350 | | O-antigen polymerase, Wzy protein | 100 |
| AVAEIV_RS02300 | 36.04040884 | 2.22943997 | 0.332990087 | 6.695214226 | 2.15E-11 | 6.28E-11 | - | transposase | pseudo=true | | | | |
| AVAEIV_RS09450 | 292.1088946 | 2.229797864 | 0.24284952 | 9.181808804 | 4.24E-20 | 3.01E-19 | - | aldehyde dehydrogenase family protein | WP_012700465.1 | Avin_18440 | | Aldehyde dehydrogenase family protein | 100 |
| AVAEIV_RS01845 | 106.2010701 | 2.232290114 | 0.239010175 | 9.339728394 | 9.66E-21 | 7.44E-20 | - | formate dehydrogenase subunit gamma | WP_012699071.1 | Avin_03830 | *fdhI* | formate dehydrogenase, gamma subunit | 100 |
| AVAEIV_RS19525 | 278.7841782 | 2.232546862 | 0.184035143 | 12.1310899 | 7.23E-34 | 2.17E-32 | - | hypothetical protein | WP_302226814.1 | | | | |
| AVAEIV_RS01200 | 119.3327752 | 2.233333536 | 0.233550716 | 9.562520611 | 1.15E-21 | 1.01E-20 | - | ABC transporter substrate-binding protein | WP_012698932.1 | Avin_02430 | | hypothetical protein | 100 |
| AVAEIV_RS16010 | 41.8729905 | 2.23358654 | 0.382927201 | 5.832927334 | 5.45E-09 | 1.28E-08 | - | SH3 domain-containing protein | WP_302225917.1 | Avin_31170 | | hypothetical protein | 99.04 |
| AVAEIV_RS04585 | 60.92867842 | 2.235129828 | 0.328687321 | 6.800170503 | 1.04E-11 | 3.13E-11 | - | urease subunit gamma | WP_012699645.1 | Avin_09900 | *ureA* | Urease, gamma subunit | 100 |
| AVAEIV_RS17080 | 136.1456715 | 2.236327174 | 0.237496837 | 9.416239819 | 4.68E-21 | 3.81E-20 | - | styrene monooxygenase/indole monooxygenase family protein | WP_302226185.1 | Avin_33980 | | oxygenase subunit protein | 97.28 |

| | | | | | | | | | | | | | |
|---|---|---|---|---|---|---|---|---|---|---|---|---|---|
| AVAEIV_RS15860 | 162.3536852 | 2.23764674 | 0.210601602 | 10.62502239 | 2.28E-26 | 3.23E-25 | - | LysR substrate-binding domain-containing protein | WP_012701634.1 | Avin_30850 | | LysR family transcriptional regulator | 100 |
| AVAEIV_RS15400 | 143.2724033 | 2.239857661 | 0.234257043 | 9.561538173 | 1.16E-21 | 1.02E-20 | - | low molecular weight protein-tyrosine-phosphatase | WP_012701542.1 | Avin_29890 | | Low molecular weight phosphotyrosine protein phosphatase | 100 |
| AVAEIV_RS24530 | 169.2434638 | 2.24333448 | 0.205058991 | 10.93994694 | 7.42E-28 | 1.21E-26 | - | IS110-like element ISAzvi12 family transposase | pseudo=true | | | | |
| AVAEIV_RS22195 | 82.64306856 | 2.245753462 | 0.287697537 | 7.805953028 | 5.91E-15 | 2.46E-14 | - | transporter substrate-binding domain-containing protein | WP_012702937.1 | Avin_44520 | | Bacterial extracellular solute-binding protein, family 3 | 100 |
| AVAEIV_RS22730 | 62.39266689 | 2.250462074 | 0.275404338 | 8.171483745 | 3.05E-16 | 1.44E-15 | - | YkgJ family cysteine cluster protein | WP_061290405.1 | Avin_45660 | | conserved hypothetical protein | 99.25 |
| AVAEIV_RS07395 | 152.4532664 | 2.251201633 | 0.209737985 | 10.73339976 | 7.09E-27 | 1.04E-25 | - | feruloyl-CoA synthase | WP_302228504.1 | Avin_14250 | *fcs* | feruoyl-CoA synthetase | 99.2 |
| AVAEIV_RS12900 | 1255.42028 | 2.255626414 | 0.182840359 | 12.33658932 | 5.75E-35 | 1.93E-33 | - | non-ribosomal peptide synthase/polyketide synthase | WP_302229202.1 | Avin_25650 | | Non-ribosomal peptide synthase:Amino acid adenylation | 98.8 |
| AVAEIV_RS06245 | 64.60814427 | 2.258692357 | 0.336881793 | 6.704702964 | 2.02E-11 | 5.90E-11 | - | Arc family DNA-binding protein | WP_072432698.1 | | | | |
| AVAEIV_RS20020 | 94.75994919 | 2.259027876 | 0.256770945 | 8.797832931 | 1.39E-18 | 8.37E-18 | - | hypothetical protein | WP_302226955.1 | Avin_39940 | | hypothetical protein | 99.74 |
| AVAEIV_RS19555 | 101.5897028 | 2.262676322 | 0.233006917 | 9.710768893 | 2.71E-22 | 2.55E-21 | - | hypothetical protein | WP_302226824.1 | | | | |
| AVAEIV_RS15045 | 59.7707433 | 2.264090331 | 0.297232798 | 7.617229134 | 2.59E-14 | 1.01E-13 | - | hypothetical protein | WP_302225630.1 | Avin_29110 | | hypothetical protein | 98.86 |
| AVAEIV_RS23540 | 58.9564466 | 2.264573652 | 0.284322267 | 7.964812861 | 1.65E-15 | 7.25E-15 | - | SDR family NAD(P)-dependent oxidoreductase | WP_012703143.1 | Avin_46750 | | Short-chain dehydrogenase/reductase SDR | 100 |
| AVAEIV_RS04010 | 194.735839 | 2.268123873 | 0.257921507 | 8.793853214 | 1.45E-18 | 8.62E-18 | - | aryl-sulfate sulfotransferase | WP_302228209.1 | Avin_08210 | | conserved hypothetical protein | 99.78 |
| AVAEIV_RS14525 | 61.92258796 | 2.268492763 | 0.31773856 | 7.139494699 | 9.37E-13 | 3.14E-12 | - | TusE/DsrC/DsvC family sulfur relay protein | WP_012701375.1 | Avin_28150 | | dissimilatory sulfite reductase, DsrC-like protein | 100 |
| AVAEIV_RS13705 | 376.6594978 | 2.270997484 | 0.182533825 | 12.44151592 | 1.56E-35 | 5.59E-34 | - | TonB-dependent receptor | WP_231876344.1 | Avin_26410 | | TonB-dependent siderophore receptor | 100 |
| AVAEIV_RS18285 | 32.73298424 | 2.27173514 | 0.36586316 | 6.209248131 | 5.32E-10 | 1.37E-09 | - | hypothetical protein | WP_302226510.1 | | | | |
| AVAEIV_RS04415 | 148.1652586 | 2.272862505 | 0.260270506 | 8.732693299 | 2.49E-18 | 1.44E-17 | - | amidohydrolase family protein | WP_012699582.1 | Avin_09180 | | Amidohydrolase 2 | 100 |
| AVAEIV_RS18830 | 103.2741409 | 2.274159747 | 0.251771987 | 9.032616276 | 1.68E-19 | 1.12E-18 | - | tRNA-uridine aminocarboxypropyltransferase | WP_302229400.1 | Avin_38500 | | Conserved hypothetical protein | 99.14 |
| AVAEIV_RS16155 | 114.0127612 | 2.274180966 | 0.268167138 | 8.480461054 | 2.24E-17 | 1.17E-16 | - | SfnB family sulfur acquisition oxidoreductase | WP_012701697.1 | Avin_31490 | | Acyl-CoA dehydrogenase family protein | 100 |
| AVAEIV_RS16035 | 125.4548193 | 2.275669226 | 0.270488882 | 8.413171025 | 3.99E-17 | 2.04E-16 | - | LysR substrate-binding domain-containing protein | WP_061291629.1 | Avin_31220 | | Transcriptional regulator, LysR family | 99.68 |
| AVAEIV_RS25090 | 144.4160367 | 2.276105508 | 0.267109489 | 8.521245404 | 1.58E-17 | 8.43E-17 | - | maltoporin | WP_012703470.1 | Avin_50270 | | LamB type porin | 100 |
| AVAEIV_RS01665 | 98.97601042 | 2.279844248 | 0.237534758 | 9.597939547 | 8.16E-22 | 7.27E-21 | - | DUF1615 domain-containing protein | WP_197471041.1 | Avin_03430 | | DUF1615 family hypothetical protein | 99.73 |
| AVAEIV_RS16040 | 134.2592225 | 2.27997503 | 0.306749286 | 7.432698724 | 1.06E-13 | 3.92E-13 | - | hypothetical protein | WP_302225928.1 | Avin_31230 | | hypothetical protein | 98.9 |
| AVAEIV_RS20285 | 88.21242786 | 2.282068526 | 0.288352745 | 7.914155716 | 2.49E-15 | 1.07E-14 | - | IS110-like element ISAzvi4 family transposase | pseudo=true | | | | |
| AVAEIV_RS09570 | 80.93848124 | 2.2846987 | 0.285195624 | 8.010987935 | 1.14E-15 | 5.08E-15 | - | ABC transporter permease | WP_302228696.1 | Avin_18680 | | ABC transporter, inner membrane permease component | 99.69 |
| AVAEIV_RS11015 | 154.7779811 | 2.287517436 | 0.219222205 | 10.43469767 | 1.72E-25 | 2.26E-24 | - | ABC transporter permease | WP_012700735.1 | Avin_21310 | | ABC transporter, permease component | 100 |
| AVAEIV_RS11475 | 81.37022778 | 2.28760486 | 0.300355684 | 7.616319528 | 2.61E-14 | 1.01E-13 | - | ABC transporter permease | WP_012700842.1 | Avin_22530 | | ABC transporter inner-membrane protein | 100 |
| AVAEIV_RS05125 | 103.2041092 | 2.289257832 | 0.334466171 | 6.844512338 | 7.67E-12 | 2.33E-11 | - | hypothetical protein | WP_302228328.1 | Avin_11080 | | conserved hypothetical protein | 99.46 |
| AVAEIV_RS04430 | 194.1219913 | 2.289414596 | 0.210992991 | 10.85066656 | 1.98E-27 | 3.05E-26 | *rhmD* | L-rhamnonate dehydratase | WP_302228256.1 | Avin_09220 | | Mandelate racemase/muconate lactonizing enzyme | 99.49 |
| AVAEIV_RS03835 | 233.6957669 | 2.290824294 | 0.185350257 | 12.35943411 | 4.33E-35 | 1.47E-33 | - | NCS2 family permease | WP_061289046.1 | Avin_07810 | | Xanthine/uracil/vitamin C permease family | 99.54 |
| AVAEIV_RS07385 | 94.09355113 | 2.293088629 | 0.248411387 | 9.231012536 | 2.68E-20 | 1.96E-19 | - | p-hydroxycinnamoyl CoA hydratase/lyase | WP_012700058.1 | Avin_14230 | *ech* | p-hydroxycinnamoyl CoA hydratase/lyase | 100 |
| AVAEIV_RS24755 | 277.649885 | 2.293486943 | 0.19440005 | 11.79776928 | 4.01E-32 | 1.04E-30 | - | aldehyde dehydrogenase (NADP(+)) | WP_302227638.1 | Avin_49290 | | Aldehyde dehydrogenase | 97.72 |
| AVAEIV_RS20040 | 95.79591861 | 2.296952509 | 0.24823699 | 9.253063019 | 2.18E-20 | 1.63E-19 | - | glycerol dehydrogenase | WP_302226956.1 | Avin_39990 | | glycerol dehydrogenase | 99.47 |
| AVAEIV_RS13835 | 41.24515931 | 2.299810992 | 0.335080089 | 6.863466582 | 6.72E-12 | 2.05E-11 | *tssB* | type VI secretion system contractile sheath small subunit | WP_012701234.1 | Avin_26670 | | conserved hypothetical protein (UCP028301) | 100 |
| AVAEIV_RS01555 | 154.1424363 | 2.299838145 | 0.222692468 | 10.32741776 | 5.30E-25 | 6.66E-24 | - | 16S rRNA (uracil(1498)-N(3))-methyltransferase | WP_302227941.1 | Avin_03190 | | conserved hypothetical protein | 99.16 |
| AVAEIV_RS18655 | 231.808055 | 2.300142773 | 0.225101276 | 10.21825737 | 1.64E-24 | 1.98E-23 | *quiC* | 3-dehydroshikimate dehydratase QuiC | WP_302226586.1 | Avin_38100 | *hppD* | 4-hydroxyphenylpyruvate dioxygenase | 99.84 |
| AVAEIV_RS05745 | 361.5458938 | 2.300495877 | 0.237107269 | 9.702342246 | 2.95E-22 | 2.75E-21 | *fadB* | fatty acid oxidation complex subunit alpha FadB | WP_302229286.1 | | | | |
| AVAEIV_RS01265 | 49.80398134 | 2.301159088 | 0.356570679 | 6.453584721 | 1.09E-10 | 3.00E-10 | - | NifB/NifX family molybdenum-iron cluster-binding protein | WP_012698946.1 | Avin_02570 | *vnfY* | nitrogen fixation-related protein, vnfY | 100 |
| AVAEIV_RS04535 | 115.2341165 | 2.30442633 | 0.248915625 | 9.257861286 | 2.09E-20 | 1.56E-19 | - | alpha/beta fold hydrolase | WP_302228264.1 | Avin_09470 | | Alpha/beta hydrolase fold protein | 99.66 |
| AVAEIV_RS04005 | 348.7812004 | 2.306074316 | 0.211096043 | 10.92428965 | 8.82E-28 | 1.42E-26 | - | TonB-dependent receptor | WP_415860955.1 | Avin_08200 | | TonB-dependent siderophore receptor | 99.87 |
| AVAEIV_RS11350 | 108.3437768 | 2.306261902 | 0.237911001 | 9.693801021 | 3.20E-22 | 2.97E-21 | - | acetyl-CoA acetyltransferase | WP_012700814.1 | Avin_22240 | | Thiolase protein | 100 |
| AVAEIV_RS01970 | 64.11376878 | 2.308555162 | 0.286086158 | 8.069440252 | 7.06E-16 | 3.24E-15 | - | hypothetical protein | WP_302227977.1 | Avin_04110 | | hypothetical protein | 89.69 |
| AVAEIV_RS14130 | 181.6881683 | 2.311163529 | 0.219084139 | 10.54920517 | 5.12E-26 | 7.02E-25 | - | mannitol dehydrogenase family protein | WP_302225477.1 | Avin_27360 | *mtlD* | Mannitol dehydrogenase | 99.39 |
| AVAEIV_RS11210 | 15.90920645 | 2.314154752 | 0.49783714 | 4.648417258 | 3.34E-06 | 6.16E-06 | - | ferredoxin family protein | WP_061291639.1 | Avin_21920 | *asfB* | 4Fe-4S ferredoxin protein | 98.46 |
| AVAEIV_RS25705 | 83.70539456 | 2.315897183 | 0.250961322 | 9.228104029 | 2.75E-20 | 2.00E-19 | - | LysR family transcriptional regulator | WP_012703592.1 | Avin_51620 | | LysR family transcriptional regulator protein | 100 |
| AVAEIV_RS21000 | 316.4137391 | 2.321859976 | 0.264018091 | 8.794321492 | 1.44E-18 | 8.59E-18 | - | SulP family inorganic anion transporter | WP_302227122.1 | Avin_41950 | | sulphate transporter | 99.58 |
| AVAEIV_RS04425 | 240.4766109 | 2.324177769 | 0.23373558 | 9.94361992 | 2.69E-23 | 2.83E-22 | - | MFS transporter | WP_302228254.1 | Avin_09210 | | MFS transporter | 98.84 |
| AVAEIV_RS11420 | 118.383822 | 2.32597076 | 0.250312961 | 9.292250601 | 1.51E-20 | 1.15E-19 | - | TauD/TfdA dioxygenase family protein | WP_012700828.1 | Avin_22390 | | Taurine dioxygenase protein | 100 |
| AVAEIV_RS23145 | 57.93838073 | 2.326747082 | 0.282318212 | 8.241576276 | 1.70E-16 | 8.28E-16 | - | phage tail protein | pseudo=true | | | | |
| AVAEIV_RS05945 | 59.45096932 | 2.328731732 | 0.303899913 | 7.662824606 | 1.82E-14 | 7.21E-14 | - | TIGR03758 family integrating conjugative element protein | WP_072432885.1 | Avin_36210 | | hypothetical conserved | 98.73 |
| AVAEIV_RS13865 | 108.8329485 | 2.328803749 | 0.259951453 | 8.958610258 | 3.29E-19 | 2.11E-18 | *tssK* | type VI secretion system baseplate subunit TssK | WP_302225391.1 | Avin_26730 | | conserved hypothetical protein | 99.77 |
| AVAEIV_RS18705 | 102.1740509 | 2.33152767 | 0.258713356 | 9.012011221 | 2.02E-19 | 1.34E-18 | - | CoA transferase subunit A | WP_012702339.1 | Avin_38220 | *pcaI* | 3-oxoadipate:succinyl-CoA transferase subunit A | 100 |
| AVAEIV_RS24845 | 129.2386609 | 2.331680576 | 0.200484469 | 11.63023045 | 2.89E-31 | 6.79E-30 | - | tripartite tricarboxylate transporter permease | WP_302227645.1 | Avin_49480 | *tctA* | tripartite tricarboxylate transporter large transmembrane protein | 99.8 |
| AVAEIV_RS00635 | 113.3905197 | 2.332747814 | 0.248667954 | 9.380974818 | 6.54E-21 | 5.18E-20 | - | bacteriohemerythrin | WP_012698824.1 | Avin_01310 | | hemerythrin-like, metal binding protein | 100 |
| AVAEIV_RS04200 | 114.6336996 | 2.333166463 | 0.283258332 | 8.236885544 | 1.77E-16 | 8.58E-16 | *dmpE* | 2-oxopent-4-enoate hydratase | WP_012699533.1 | Avin_08670 | *xylJ* | 4-oxo-4-pentenoate hydratase, XylJ | 100 |
| AVAEIV_RS06070 | 174.9769738 | 2.335701903 | 0.520250395 | 4.489572573 | 7.14E-06 | 1.27E-05 | - | hypothetical protein | WP_072432734.1 | | | | |
| AVAEIV_RS21080 | 67.57834227 | 2.337923923 | 0.330152928 | 7.081336325 | 1.43E-12 | 4.71E-12 | - | 2-keto-4-pentenoate hydratase | WP_012702710.1 | Avin_42150 | | 2-hydroxypent-2,4-dienoate hydratase | 100 |
| AVAEIV_RS23530 | 39.64336603 | 2.338776317 | 0.332868689 | 7.026122885 | 2.12E-12 | 6.83E-12 | - | GlcG/HbpS family heme-binding protein | WP_012703140.1 | Avin_46720 | | Conserved hypothetical protein | 100 |
| AVAEIV_RS23225 | 101.2873724 | 2.338826192 | 0.232907792 | 10.04185462 | 9.98E-24 | 1.10E-22 | - | hypothetical protein | WP_302227529.1 | | | | |
| AVAEIV_RS23525 | 182.1906099 | 2.338940001 | 0.18370771 | 12.73185542 | 3.93E-37 | 1.69E-35 | - | MFS transporter | WP_012703139.1 | Avin_46710 | | Major facilitator superfamily transporter | 100 |
| AVAEIV_RS25650 | 151.7905716 | 2.341755534 | 0.215716313 | 10.85571833 | 1.87E-27 | 2.91E-26 | - | hypothetical protein | WP_175555998.1 | | | | |
| AVAEIV_RS12925 | 145.9941195 | 2.342300887 | 0.2152963 | 10.87942936 | 1.44E-27 | 2.29E-26 | - | site-specific integrase | WP_302229210.1 | Avin_25710 | | cointegrate resolution protein S | 99.39 |
| AVAEIV_RS02505 | 297.8146 | 2.346969106 | 0.155811722 | 15.06285326 | 2.84E-51 | 3.65E-49 | - | TonB-dependent receptor | WP_302228055.1 | Avin_04960 | | TonB-dependent receptor | 99.41 |
| AVAEIV_RS19600 | 130.7877476 | 2.351741434 | 0.261127076 | 9.006118669 | 2.13E-19 | 1.40E-18 | - | hypothetical protein | WP_302226842.1 | | | | |
| AVAEIV_RS15825 | 30.79656585 | 2.353335982 | 0.402275369 | 5.850062334 | 4.91E-09 | 1.16E-08 | - | MmoB/DmpM family protein | WP_012701625.1 | Avin_30740 | *lapM* | Multi-component phenol hydoxylase, activator subunit; LapM | 100 |
| AVAEIV_RS00325 | 85.94318327 | 2.353681545 | 0.300295683 | 7.837880046 | 4.58E-15 | 1.93E-14 | - | DUF3175 domain-containing protein | WP_012698762.1 | Avin_00670 | | hypothetical protein | 100 |
| AVAEIV_RS09670 | 55.92087918 | 2.354620944 | 0.310279201 | 7.588716675 | 3.23E-14 | 1.25E-13 | - | hypothetical protein | WP_082765920.1 | Avin_18890 | | hypothetical protein | 98.36 |
| AVAEIV_RS11640 | 59.67557102 | 2.355070273 | 0.289151715 | 8.144756375 | 3.80E-16 | 1.78E-15 | - | YgiW/YdeI family stress tolerance OB fold protein | WP_012700874.1 | Avin_22850 | | Conserved hypothetical protein 156 | 100 |
| AVAEIV_RS16115 | 74.48506582 | 2.358961904 | 0.286259375 | 8.240645074 | 1.71E-16 | 8.34E-16 | - | ABC transporter substrate-binding protein | WP_012701687.1 | Avin_31390 | | ABC nitrate/sulfonate/bicarbonate family transporter, periplasmic ligand binding protein | 100 |
| AVAEIV_RS09520 | 145.9115566 | 2.362046255 | 0.224841875 | 10.5053663 | 8.16E-26 | 1.10E-24 | - | ParA family protein | WP_302228688.1 | | | | |

| | | | | | | | | | | | | | |
|---|---|---|---|---|---|---|---|---|---|---|---|---|---|
| AVAEIV_RS12895 | 169.057782 | 2.363200409 | 0.234809074 | 10.06434874 | 7.94E-24 | 8.88E-23 | - | aspartate aminotransferase family protein | WP_012701135.1 | Avin_25640 | | Diaminobutyrate-2-oxoglutarate aminotransferase | 100 |
| AVAEIV_RS01820 | 42.46067555 | 2.367513135 | 0.33262584 | 7.117646456 | 1.10E-12 | 3.66E-12 | - | hypothetical protein | WP_012699066.1 | Avin_03780 | | hypothetical protein | 100 |
| AVAEIV_RS16425 | 290.954417 | 2.370467449 | 0.245458401 | 9.657308273 | 4.58E-22 | 4.21E-21 | - | Bcr/CflA family multidrug efflux MFS transporter | WP_231876385.1 | Avin_32370 | | Drug resistance transporter Bcr/CflA subfamily protein | 100 |
| AVAEIV_RS04000 | 175.8216806 | 2.370855028 | 0.261103261 | 9.080143318 | 1.08E-19 | 7.37E-19 | - | LysR family transcriptional regulator | WP_012699488.1 | Avin_24730 | | LysR family transcriptional regulator protein | 99.68 |
| AVAEIV_RS02500 | 91.80175948 | 2.371274019 | 0.244065064 | 9.715745392 | 2.58E-22 | 2.44E-21 | - | GGDEF domain-containing protein | WP_012699177.1 | Avin_04950 | | GGDEF domain protein | 100 |
| AVAEIV_RS13685 | 173.5506307 | 2.375653956 | 0.238764677 | 9.949771402 | 2.53E-23 | 2.67E-22 | - | aldo/keto reductase | WP_302225334.1 | Avin_26350 | | Aldo/keto reductase protein | 99.67 |
| AVAEIV_RS07195 | 110.0413528 | 2.375960614 | 0.252381445 | 9.414165192 | 4.77E-21 | 3.87E-20 | - | protease inhibitor I42 family protein | WP_012700023.1 | Avin_13870 | | conserved hypothetical protein | 100 |
| AVAEIV_RS21890 | 140.7545392 | 2.37627621 | 0.23485379 | 10.11810886 | 4.59E-24 | 5.23E-23 | - | amidohydrolase family protein | WP_012702874.1 | Avin_43860 | | Amidohydrolase | 100 |
| AVAEIV_RS02155 | 37.23139304 | 2.376310093 | 0.447571636 | 5.309340229 | 1.10E-07 | 2.31E-07 | - | helix-turn-helix transcriptional regulator | WP_367281688.1 | | | | |
| AVAEIV_RS05900 | 42.45553951 | 2.378243207 | 0.378536734 | 6.282727647 | 3.33E-10 | 8.72E-10 | - | TIGR03752 family integrating conjugative element protein | pseudo=true | | | | |
| AVAEIV_RS21900 | 253.2314106 | 2.379987526 | 0.199462328 | 11.93201516 | 8.06E-33 | 2.24E-31 | - | acyl-CoA dehydrogenase family protein | WP_012702877.1 | Avin_43890 | | Acyl-CoA dehydrogenase-related protein | 100 |
| AVAEIV_RS19955 | 39.16402885 | 2.381067465 | 0.359643035 | 6.620641125 | 3.58E-11 | 1.02E-10 | - | DUF1778 domain-containing protein | WP_061288032.1 | Avin_39810 | | conserved hypothetical protein | 98.97 |
| AVAEIV_RS25725 | 227.2766639 | 2.382346851 | 0.170902794 | 13.93977705 | 3.63E-44 | 2.71E-42 | - | bifunctional diguanylate cyclase/phosphodiesterase | WP_012703596.1 | Avin_51660 | | EAL/GGDEF domain-containing protein | 100 |
| AVAEIV_RS13750 | 304.6735788 | 2.383203241 | 0.239740026 | 9.940781618 | 2.77E-23 | 2.91E-22 | - | hypothetical protein | WP_302225356.1 | Avin_26500 | | hypothetical protein | 99.88 |
| AVAEIV_RS06365 | 77.72956728 | 2.390562638 | 0.312475422 | 7.650402138 | 2.00E-14 | 7.89E-14 | - | pilin | WP_302228462.1 | Avin_12104 | *pilA* | fimbrial protein | 99.39 |
| AVAEIV_RS20090 | 199.7280056 | 2.393207536 | 0.200672774 | 11.92592041 | 8.67E-33 | 2.40E-31 | *leuA* | 2-isopropylmalate synthase | WP_012702523.1 | Avin_40120 | | 2-isopropylmalate synthase | 100 |
| AVAEIV_RS16120 | 84.7292064 | 2.394339797 | 0.265859623 | 9.006030199 | 2.14E-19 | 1.40E-18 | - | ABC transporter substrate-binding protein | WP_302225951.1 | Avin_31400 | | ABC nitrate/sulfonate/bicarbonate family transporter, periplasmic ligand binding protein | 99.68 |
| AVAEIV_RS11125 | 80.33360358 | 2.405075042 | 0.301846878 | 7.967864551 | 1.61E-15 | 7.09E-15 | - | class II aldolase/adducin family protein | WP_012700757.1 | Avin_21590 | | Class II aldolase/adducin domain protein | 100 |
| AVAEIV_RS05735 | 162.7730857 | 2.405263031 | 0.26173377 | 9.189731333 | 3.94E-20 | 2.81E-19 | - | electron transfer flavoprotein subunit alpha/FixB family protein | WP_302228411.1 | | | | |
| AVAEIV_RS24735 | 186.1929051 | 2.407469154 | 0.20776241 | 11.58760698 | 4.76E-31 | 1.06E-29 | - | FadR/GntR family transcriptional regulator | WP_012703383.1 | Avin_49240 | | Bacterial regulatory protein, GntR family | 100 |
| AVAEIV_RS11120 | 99.27943247 | 2.407583838 | 0.303350783 | 7.936633016 | 2.08E-15 | 9.00E-15 | - | D-isomer specific 2-hydroxyacid dehydrogenase family protein | WP_061291565.1 | Avin_21580 | | D-isomer specific 2-hydroxyacid dehydrogenase | 99.68 |
| AVAEIV_RS00375 | 124.8899943 | 2.411705062 | 0.299169696 | 8.061328057 | 7.55E-16 | 3.44E-15 | - | hypothetical protein | WP_231876391.1 | Avin_00770 | | transmembrane protein | 100 |
| AVAEIV_RS24690 | 36.09236868 | 2.412041458 | 0.380333414 | 6.341913089 | 2.27E-10 | 6.02E-10 | - | hypothetical protein | WP_012703374.1 | Avin_49130 | | hypothetical protein | 100 |
| AVAEIV_RS02875 | 229.6825013 | 2.414951317 | 0.20529528 | 11.7633066 | 6.03E-32 | 1.54E-30 | *mdtD* | multidrug transporter subunit MdtD | WP_012699261.1 | Avin_05800 | | Drug resistance transporter EmrB/QacA subfamily | 100 |
| AVAEIV_RS04290 | 244.5971681 | 2.417874783 | 0.255127332 | 9.477129587 | 2.61E-21 | 2.20E-20 | - | aromatic/alkene monooxygenase hydroxylase subunit beta | WP_302228240.1 | Avin_08850 | *dpmL* | Phenol hydroxylase subunit P1 | 99.7 |
| AVAEIV_RS25060 | 126.65731 | 2.419415163 | 0.297050299 | 8.144799615 | 3.80E-16 | 1.78E-15 | - | maltoporin | WP_012703464.1 | Avin_50210 | | LamB type porin | 100 |
| AVAEIV_RS25135 | 26.7220409 | 2.421618785 | 0.384505985 | 6.298000236 | 3.02E-10 | 7.93E-10 | - | MbtH family protein | WP_012703479.1 | Avin_50380 | | MbtH-like protein | 100 |
| AVAEIV_RS24980 | 88.98917649 | 2.423187507 | 0.27920839 | 8.678777549 | 4.00E-18 | 2.27E-17 | *iolE* | myo-inosose-2 dehydratase | WP_012703448.1 | Avin_50050 | *iolE* | 2-keto-myo-inositol dehydratase | 100 |
| AVAEIV_RS02460 | 292.1084937 | 2.424446467 | 0.205765464 | 11.78257236 | 4.80E-32 | 1.23E-30 | - | efflux RND transporter periplasmic adaptor subunit | WP_302228044.1 | | | | |
| AVAEIV_RS13950 | 263.7194296 | 2.425359514 | 0.273790178 | 8.858460624 | 8.11E-19 | 5.01E-18 | - | maltoporin | WP_012701259.1 | Avin_16650 | | LamB type porin | 94.34 |
| AVAEIV_RS11470 | 77.58006298 | 2.425933793 | 0.294662453 | 8.232924717 | 1.83E-16 | 8.84E-16 | - | ABC transporter permease | WP_302228995.1 | Avin_22520 | | ABC transporter inner-membrane protein | 99.64 |
| AVAEIV_RS12905 | 70.16940056 | 2.42615532 | 0.270646026 | 8.964311648 | 3.12E-19 | 2.01E-18 | - | RNA polymerase factor sigma-70 | WP_085930851.1 | Avin_25660 | | RNA polymerase sigma factor, ECF subfamily, PvdS-like protein | 100 |
| AVAEIV_RS13550 | 22.4983279 | 2.427552618 | 0.465386169 | 5.216211348 | 1.83E-07 | 3.78E-07 | - | hypothetical protein | WP_415860884.1 | | | | |
| AVAEIV_RS05855 | 31.73417483 | 2.42939153 | 0.422923482 | 5.744281488 | 9.23E-09 | 2.12E-08 | - | hypothetical protein | WP_367281697.1 | | | | |
| AVAEIV_RS15970 | 205.3593791 | 2.434037036 | 0.227164514 | 10.7148647 | 8.67E-27 | 1.27E-25 | - | type II toxin-antitoxin system HipA family toxin | WP_302225903.1 | Avin_31080 | | conserved hypothetical protein | 96.33 |
| AVAEIV_RS10725 | 515.59996 | 2.435802767 | 0.190207377 | 12.80603731 | 1.52E-37 | 6.61E-36 | - | hydantoinase/oxoprolinase family protein | WP_302228854.1 | Avin_20910 | *acxA* | Acetone carboxylase beta subunit; AcxA | 99.72 |
| AVAEIV_RS01910 | 31.35592529 | 2.43706448 | 0.395499111 | 6.161997369 | 7.18E-10 | 1.83E-09 | - | DUF1656 domain-containing protein | WP_012699085.1 | Avin_03990 | | conserved hypothetical protein | 100 |
| AVAEIV_RS11245 | 107.0282749 | 2.443064001 | 0.24502678 | 9.970599968 | 2.05E-23 | 2.20E-22 | - | ABC transporter permease | WP_012700789.1 | Avin_21990 | | ABC transporter, inner membrane permease component | 100 |
| AVAEIV_RS00865 | 227.3972548 | 2.445049294 | 0.2320686 | 10.53589023 | 5.90E-26 | 8.05E-25 | - | AEC family transporter | WP_012698869.1 | Avin_01790 | | conserved hypothetical protein | 100 |
| AVAEIV_RS15000 | 107.5202437 | 2.445812762 | 0.268281559 | 9.116589185 | 7.75E-20 | 5.36E-19 | - | sensor histidine kinase | WP_302225618.1 | Avin_29010 | | periplasmic sensory histidine protein kinase, two-component | 99.05 |
| AVAEIV_RS00230 | 64.96286914 | 2.447350998 | 0.327738806 | 7.467382411 | 8.18E-14 | 3.05E-13 | - | GTP-binding protein | WP_012698742.1 | Avin_00470 | | nickel incorporation protein | 100 |
| AVAEIV_RS10375 | 146.2548557 | 2.447662508 | 0.212010672 | 11.54499673 | 7.82E-31 | 1.70E-29 | - | helix-turn-helix domain-containing protein | WP_302228776.1 | Avin_20310 | | transcriptional regulator, AraC family | 97.97 |
| AVAEIV_RS08950 | 219.9433203 | 2.447674565 | 0.218547616 | 11.19973126 | 4.09E-29 | 7.67E-28 | - | long-chain-fatty-acid--CoA ligase | WP_012700360.1 | Avin_17370 | | long-chain-fatty-acid--CoA ligase | 100 |
| AVAEIV_RS23140 | 107.597014 | 2.451458177 | 0.332692418 | 7.368542355 | 1.73E-13 | 6.26E-13 | - | phage tail protein | pseudo=true | | | | |
| AVAEIV_RS16160 | 210.2407222 | 2.452790013 | 0.200791057 | 12.21563378 | 2.56E-34 | 8.19E-33 | - | LLM class flavin-dependent oxidoreductase | WP_302225970.1 | Avin_31500 | | Flavin-dependent oxidoreductase | 99.14 |
| AVAEIV_RS08965 | 186.217573 | 2.453941308 | 0.274949217 | 8.925071087 | 4.45E-19 | 2.82E-18 | - | electron transfer flavoprotein-ubiquinone oxidoreductase | WP_302228621.1 | Avin_16430 | | Electron transfer flavoprotein-ubiquinone oxidoreductase | 99.09 |
| AVAEIV_RS20045 | 221.2173203 | 2.457429633 | 0.178309043 | 13.78185641 | 3.28E-43 | 2.22E-41 | - | OprD family porin | WP_012702513.1 | Avin_40000 | | OprD family outer membrane porin | 100 |
| AVAEIV_RS13810 | 97.78520317 | 2.461866133 | 0.25779479 | 9.549712527 | 1.30E-21 | 1.13E-20 | *tssG* | type VI secretion system baseplate subunit TssG | WP_302225377.1 | Avin_26620 | | conserved hypothetical protein | 99.7 |
| AVAEIV_RS18385 | 201.449598 | 2.46622097 | 0.232827112 | 10.59249907 | 3.23E-26 | 4.52E-25 | - | sensor histidine kinase | WP_012702225.1 | Avin_37040 | | sensory histidine protein kinase, two component | 100 |
| AVAEIV_RS01255 | 121.6851463 | 2.468877441 | 0.236977131 | 10.41820968 | 2.05E-25 | 2.67E-24 | *phnD* | phosphate/phosphite/phosphonate ABC transporter substrate-binding protein | WP_012698944.1 | Avin_02550 | | phosphonate transport system substrate-binding protein | 100 |
| AVAEIV_RS03970 | 195.3108493 | 2.469276923 | 0.235969912 | 10.46437192 | 1.26E-25 | 1.66E-24 | - | ABC transporter permease | WP_012699480.1 | Avin_08100 | | ABC transporter, permease component | 100 |
| AVAEIV_RS13935 | 225.6120764 | 2.470928758 | 0.254396017 | 9.712922337 | 2.66E-22 | 2.50E-21 | - | MFS transporter | WP_061290242.1 | Avin_26900 | *kguT* | 2-ketogluconate transporter | 99.54 |
| AVAEIV_RS23095 | 29.48397699 | 2.473770322 | 0.386856706 | 6.394539074 | 1.61E-10 | 4.35E-10 | - | DUF1353 domain-containing protein | WP_302227512.1 | | | | |
| AVAEIV_RS09500 | 149.6312825 | 2.479178118 | 0.270364833 | 9.169750705 | 4.74E-20 | 3.36E-19 | - | TolC family protein | WP_061291401.1 | Avin_18540 | | outer membrane efflux protein | 99.77 |
| AVAEIV_RS16030 | 144.1317502 | 2.482792704 | 0.227482491 | 10.91421453 | 9.86E-28 | 1.58E-26 | - | dicarboxylate/amino acid:cation symporter | WP_061291627.1 | Avin_31210 | *dctA-3* | C4-dicarboxylate transport protein | 99.77 |
| AVAEIV_RS24970 | 122.1405141 | 2.483084721 | 0.24005503 | 10.34381458 | 4.46E-25 | 5.65E-24 | - | MurR/RpiR family transcriptional regulator | WP_302227651.1 | Avin_50020 | | Sugar isomerase (SIS) | 99.68 |
| AVAEIV_RS10700 | 246.07252 | 2.483694286 | 0.247366934 | 10.04052661 | 1.01E-23 | 1.12E-22 | - | MFS transporter | WP_302228846.1 | Avin_20840 | | multidrug/chloramphenicol efflux transporter, major facilitator superfamily MFS_1 | 99.75 |
| AVAEIV_RS01855 | 101.4393972 | 2.488671235 | 0.296732104 | 8.386929494 | 4.99E-17 | 2.53E-16 | *moaA* | GTP 3',2C8-cyclase MoaA | WP_302227960.1 | Avin_03850 | *moaA* | molybdenum cofactor biosynthesis protein A | 99.7 |
| AVAEIV_RS25625 | 157.9564783 | 2.491599845 | 0.241680032 | 10.30949815 | 6.38E-25 | 7.91E-24 | - | SDR family NAD(P)-dependent oxidoreductase | WP_302227744.1 | Avin_51440 | | Short-chain dehydrogenase/reductase (SDR) family protein | 99.62 |
| AVAEIV_RS10390 | 42.1501573 | 2.493644359 | 0.379077303 | 6.578194833 | 4.76E-11 | 1.35E-10 | - | NIPSNAP family protein | WP_012700648.1 | Avin_20350 | | conserved hypothetical protein | 100 |
| AVAEIV_RS08940 | 291.2170171 | 2.49390315 | 0.244137176 | 10.21517163 | 1.70E-24 | 2.04E-23 | - | 3-hydroxyacyl-CoA dehydrogenase NAD-binding domain-containing protein | WP_302229318.1 | Avin_17350 | | 3-hydroxyacyl-CoA dehydrogenase | 99.71 |
| AVAEIV_RS18255 | 109.3486148 | 2.49422404 | 0.241539269 | 10.32637075 | 5.35E-25 | 6.71E-24 | - | hypothetical protein | WP_302226498.1 | | | | |
| AVAEIV_RS15895 | 134.8740699 | 2.498403526 | 0.245760451 | 10.1660113 | 2.81E-24 | 3.31E-23 | - | ABC transporter substrate-binding protein | WP_012701641.1 | Avin_30920 | | ABC transporter, periplasmic binding protein | 100 |
| AVAEIV_RS25110 | 118.8382357 | 2.502049924 | 0.235755779 | 10.61288905 | 2.60E-26 | 3.66E-25 | - | sugar ABC transporter substrate-binding protein | WP_012703474.1 | Avin_50320 | | ABC transporter substrate binding protein | 100 |
| AVAEIV_RS15700 | 105.9415664 | 2.511054525 | 0.241465921 | 10.39920876 | 2.50E-25 | 3.24E-24 | *ssuD* | FMNH2-dependent alkanesulfonate monooxygenase | WP_012701601.1 | Avin_30500 | *ssuD* | alkanesulfonate monooxygenase | 100 |
| AVAEIV_RS11430 | 114.7240807 | 2.513645038 | 0.278539996 | 9.024359412 | 1.81E-19 | 1.20E-18 | - | ABC transporter substrate-binding protein | WP_012700831.1 | Avin_22420 | | ABC nitrate/sulfonate/bicarbonate family transporter protein | 100 |
| AVAEIV_RS11200 | 81.69003838 | 2.514989813 | 0.288564187 | 8.715529959 | 2.89E-18 | 1.66E-17 | - | alkene reductase | WP_302228950.1 | Avin_21900 | *nemA* | NADH:flavin oxidoreductase/NADH oxidase | 99.16 |

| | | | | | | | | | | | | | |
|---|---|---|---|---|---|---|---|---|---|---|---|---|---|
| AVAEIV_RS01160 | 94.51949029 | 2.516450929 | 0.268359667 | 9.377157752 | 6.78E-21 | 5.35E-20 | - | MerR family transcriptional regulator | WP_012698924.1 | Avin_02350 | | transcriptional regulator, MerR family | 100 |
| AVAEIV_RS23205 | 127.3064164 | 2.51849573 | 0.24640767 | 10.22084959 | 1.60E-24 | 1.93E-23 | - | S24 family peptidase | WP_307189437.1 | | | | |
| AVAEIV_RS25080 | 195.1925056 | 2.52431775 | 0.201289468 | 12.54073439 | 4.47E-36 | 1.70E-34 | - | TIM barrel protein | WP_012703468.1 | Avin_50250 | | conserved hypothetical protein | 100 |
| AVAEIV_RS05740 | 152.3781253 | 2.525353733 | 0.237587078 | 10.62917124 | 2.18E-26 | 3.10E-25 | - | electron transfer flavoprotein subunit beta/FixA family protein | WP_302228412.1 | Avin_16410 | *etfB3* | electron transfer flavoprotein beta subunit | 97.99 |
| AVAEIV_RS16065 | 78.99890853 | 2.530996448 | 0.319346117 | 7.925558867 | 2.27E-15 | 9.78E-15 | - | 4-carboxy-4-hydroxy-2-oxoadipate aldolase/oxaloacetate decarboxylase | WP_061291633.1 | Avin_31280 | | 4-carboxy-4-hydroxy-2-oxoadipate aldolase protein | 99.58 |
| AVAEIV_RS04230 | 153.4762178 | 2.535373638 | 0.249726595 | 10.15259764 | 3.23E-24 | 3.74E-23 | - | 2-hydroxymuconic semialdehyde dehydrogenase | WP_012699539.1 | Avin_08730 | *xylG* | 2-hydroxymuconic semi-aldehyde dehydrogenase | 100 |
| AVAEIV_RS08810 | 61.04673275 | 2.535506764 | 0.299210829 | 8.473980615 | 2.37E-17 | 1.24E-16 | - | ogr/Delta-like zinc finger family protein | WP_231876374.1 | Avin_17030 | | hypothetical protein | 100 |
| AVAEIV_RS23050 | 127.3923589 | 2.539662788 | 0.238425893 | 10.65179104 | 1.71E-26 | 2.45E-25 | - | phage major capsid protein%2C P2 family | WP_302227504.1 | | | | |
| AVAEIV_RS25630 | 127.4358781 | 2.540580694 | 0.235647533 | 10.78127431 | 4.22E-27 | 6.34E-26 | - | SMP-30/gluconolactonase/LRE family protein | WP_302227745.1 | Avin_51450 | | gluconolactonase | 99.3 |
| AVAEIV_RS21860 | 74.11718637 | 2.543044494 | 0.293862962 | 8.653844901 | 4.98E-18 | 2.79E-17 | - | ABC transporter ATP-binding protein | WP_302227300.1 | Avin_43780 | | ABC transporter, LivG family | 99.62 |
| AVAEIV_RS00625 | 87.22264516 | 2.545961119 | 0.272360203 | 9.347772139 | 8.95E-21 | 6.96E-20 | *modB* | molybdate ABC transporter permease subunit | WP_012698822.1 | Avin_01290 | | ABC transporter, membrane spanning protein | 100 |
| AVAEIV_RS19425 | 342.8663199 | 2.551920397 | 0.277544985 | 9.194619019 | 3.76E-20 | 2.70E-19 | - | hypothetical protein | WP_302226774.1 | | | | |
| AVAEIV_RS10760 | 128.1798688 | 2.554276924 | 0.226388524 | 11.2827138 | 1.60E-29 | 3.10E-28 | - | CRISPR-associated endonuclease Cas1 | WP_302228863.1 | | | | |
| AVAEIV_RS10840 | 261.7664431 | 2.556082214 | 0.207484746 | 12.31937416 | 7.12E-35 | 2.36E-33 | - | RAMP superfamily CRISPR-associated protein | WP_302228886.1 | | | | |
| AVAEIV_RS09485 | 48.40680405 | 2.559639898 | 0.328203845 | 7.798933303 | 6.24E-15 | 2.59E-14 | - | MerR family transcriptional regulator | WP_415860988.1 | Avin_18510 | | hypothetical protein | 98.04 |
| AVAEIV_RS11220 | 282.2866816 | 2.562299875 | 0.194136927 | 13.19841578 | 8.96E-40 | 5.01E-38 | - | molybdopterin guanine dinucleotide-containing S/N-oxide reductase | WP_302228954.1 | Avin_21940 | *torG* | trimethylamine-N-oxide reductase (cytochrome c) | 99.87 |
| AVAEIV_RS20910 | 210.9633548 | 2.563908336 | 0.247476155 | 10.36022374 | 3.76E-25 | 4.80E-24 | - | alpha-ketoacid dehydrogenase subunit beta | WP_012702676.1 | Avin_41740 | *acoB* | acetoin:2,6-dichlorophenolindophenol oxidoreductase beta subunit AcoB | 100 |
| AVAEIV_RS13740 | 101.1681192 | 2.564187576 | 0.352672857 | 7.270725621 | 3.58E-13 | 1.25E-12 | - | alpha/beta fold hydrolase | WP_302225352.1 | Avin_26480 | | epoxide hydrolase protein | 99.66 |
| AVAEIV_RS05115 | 392.2513438 | 2.568727777 | 0.239089869 | 10.74377509 | 6.34E-27 | 9.37E-26 | - | cytochrome ubiquinol oxidase subunit I | WP_302228325.1 | Avin_11050 | | Cytochrome bd ubiquinol oxidase, subunit I | 99.79 |
| AVAEIV_RS15875 | 91.30764867 | 2.569820584 | 0.319861134 | 8.034175818 | 9.42E-16 | 4.24E-15 | - | ABC transporter substrate-binding protein | WP_302225873.1 | Avin_30880 | | ABC transporter, periplasmic substrate-binding protein | 99.12 |
| AVAEIV_RS23045 | 186.3648548 | 2.577440879 | 0.253641888 | 10.16173195 | 2.94E-24 | 3.44E-23 | - | GPO family capsid scaffolding protein | WP_302227503.1 | | | | |
| AVAEIV_RS23515 | 49.00297025 | 2.577714269 | 0.340128315 | 7.578652388 | 3.49E-14 | 1.34E-13 | - | SDR family oxidoreductase | WP_012703137.1 | Avin_46690 | | hypothetical protein | 100 |
| AVAEIV_RS09475 | 49.79928112 | 2.579045655 | 0.390049161 | 6.612104096 | 3.79E-11 | 1.08E-10 | - | hypothetical protein | WP_041816954.1 | | | | |
| AVAEIV_RS17060 | 42.13354187 | 2.580504315 | 0.369482872 | 6.984097257 | 2.87E-12 | 9.08E-12 | - | putative quinol monooxygenase | WP_302226182.1 | Avin_33910 | | Antibiotic biosynthesis monooxygenase protein | 98.98 |
| AVAEIV_RS11025 | 93.2218018 | 2.583066116 | 0.274465247 | 9.411268431 | 4.90E-21 | 3.96E-20 | - | TauD/TfdA dioxygenase family protein | WP_302228920.1 | Avin_21330 | | Taurine catabolism dioxygenase, TauD/TfdA family | 99.7 |
| AVAEIV_RS25700 | 89.54783253 | 2.586746193 | 0.245889941 | 10.51993499 | 6.99E-26 | 9.46E-25 | - | CoA transferase subunit A | WP_012703591.1 | Avin_51610 | | 3-oxoacid CoA-transferase | 100 |
| AVAEIV_RS06010 | 137.0109903 | 2.587564237 | 0.301758906 | 8.574939071 | 9.91E-18 | 5.41E-17 | - | lytic transglycosylase | WP_072432873.1 | | | | |
| AVAEIV_RS13255 | 77.51307861 | 2.589423266 | 0.262163316 | 9.877138046 | 5.23E-23 | 5.30E-22 | - | aminotransferase class V-fold PLP-dependent enzyme | WP_302225207.1 | | | | |
| AVAEIV_RS05890 | 913.3856109 | 2.59180814 | 0.35953723 | 7.208733686 | 5.65E-13 | 1.93E-12 | - | conjugative transfer ATPase | WP_302228425.1 | Avin_36280 | | lipoprotein | 96.24 |
| AVAEIV_RS21075 | 153.1554699 | 2.593052652 | 0.23955365 | 10.8245174 | 2.63E-27 | 4.03E-26 | - | feruloyl-CoA synthase | WP_302227141.1 | Avin_42140 | | AMP-dependent synthetase and ligase | 99.84 |
| AVAEIV_RS16060 | 95.16222206 | 2.595200767 | 0.305966535 | 8.481975861 | 2.21E-17 | 1.16E-16 | *galB* | 4-oxalmesaconate hydratase | WP_012701676.1 | Avin_31270 | | LmbE-like protein | 100 |
| AVAEIV_RS15880 | 136.983194 | 2.600197157 | 0.258003703 | 10.07813891 | 6.90E-24 | 7.80E-23 | - | SDR family NAD(P)-dependent oxidoreductase | WP_012701638.1 | Avin_30890 | | Short-chain dehydrogenase/reductase SDR protein | 100 |
| AVAEIV_RS21780 | 84.68097276 | 2.601532266 | 0.291039309 | 8.938765956 | 3.94E-19 | 2.50E-18 | - | ABC transporter permease | WP_012702848.1 | Avin_43600 | | Bacterial inner-membrane translocator, araH family | 100 |
| AVAEIV_RS21740 | 28.33843909 | 2.604821822 | 0.431257264 | 6.040064801 | 1.54E-09 | 3.80E-09 | - | hypothetical protein | WP_302227268.1 | | | | |
| AVAEIV_RS04480 | 100.2577328 | 2.605502795 | 0.268420427 | 9.706797743 | 2.82E-22 | 2.64E-21 | - | MFS transporter | WP_302228259.1 | Avin_09320 | | transporter, MFS superfamily | 99.47 |
| AVAEIV_RS11225 | 101.7319393 | 2.607403235 | 0.305889214 | 8.524011698 | 1.54E-17 | 8.24E-17 | - | alpha/beta fold hydrolase | WP_302228955.1 | Avin_21950 | | Alpha/beta hydrolase fold protein | 98.17 |
| AVAEIV_RS05895 | 64.82188735 | 2.614325919 | 0.293826758 | 8.897507961 | 5.71E-19 | 3.59E-18 | - | TIGR03751 family conjugal transfer lipoprotein | WP_302228426.1 | Avin_36270 | | conserved hypothetical protein | 96.62 |
| AVAEIV_RS06485 | 54.56612005 | 2.617893007 | 0.309799989 | 8.450268231 | 2.91E-17 | 1.50E-16 | - | DUF2271 domain-containing protein | WP_175555904.1 | Avin_12390 | | conserved hypothetical protein | 100 |
| AVAEIV_RS25935 | 144.36728 | 2.623286619 | 0.277102818 | 9.466834871 | 2.88E-21 | 2.42E-20 | - | ABC transporter substrate-binding protein | WP_082765947.1 | Avin_52130 | | periplasmic binding protein | 99.49 |
| AVAEIV_RS04420 | 104.3634814 | 2.623532102 | 0.274179796 | 9.568655826 | 1.08E-21 | 9.52E-21 | - | maltoporin | WP_302228253.1 | Avin_09190 | | LamB type porin | 99.76 |
| AVAEIV_RS13050 | 47.22738879 | 2.626528589 | 0.39590506 | 6.634238495 | 3.26E-11 | 9.39E-11 | - | arsenate reductase ArsC | WP_302225172.1 | Avin_25910 | | Low molecular weight phosphotyrosine protein phosphatase | 99.36 |
| AVAEIV_RS17965 | 161.4639584 | 2.627123029 | 0.300452082 | 8.743900234 | 2.25E-18 | 1.31E-17 | - | TIGR03756 family integrating conjugative element protein | WP_302226419.1 | | | | |
| AVAEIV_RS25560 | 225.4479093 | 2.630089869 | 0.236495418 | 11.12110287 | 9.90E-29 | 1.76E-27 | - | MFS transporter | WP_012703563.1 | Avin_51310 | *dgoT* | D-galactonate transporter | 100 |
| AVAEIV_RS07775 | 168.154528 | 2.631193879 | 0.236467787 | 11.1270711 | 9.26E-29 | 1.65E-27 | - | acyl-CoA dehydrogenase family protein | WP_012700137.1 | Avin_15060 | | oxidoreductase | 100 |
| AVAEIV_RS05655 | 179.1121824 | 2.635081945 | 0.492345786 | 5.35209607 | 8.69E-08 | 1.84E-07 | - | TIGR03756 family integrating conjugative element protein | WP_302228397.1 | Avin_36350 | | conserved hypothetical protein | 99.68 |
| AVAEIV_RS23040 | 449.3933972 | 2.63552736 | 0.429873184 | 6.130941532 | 8.74E-10 | 2.21E-09 | - | terminase large subunit domain-containing protein | WP_302227501.1 | | | | |
| AVAEIV_RS19925 | 101.9498757 | 2.645374016 | 0.250149032 | 10.57519191 | 3.88E-26 | 5.40E-25 | - | D-2-hydroxyacid dehydrogenase family protein | WP_302226929.1 | Avin_39750 | | D-isomer specific 2-hydroxyacid dehydrogenase | 99.69 |
| AVAEIV_RS17885 | 109.8115179 | 2.646949648 | 0.277188903 | 9.549262699 | 1.31E-21 | 1.13E-20 | - | lytic transglycosylase domain-containing protein | pseudo=true | | | | |
| AVAEIV_RS02020 | 81.48348217 | 2.647292744 | 0.287367904 | 9.212207427 | 3.19E-20 | 2.31E-19 | - | hypothetical protein | WP_012699107.1 | Avin_04220 | | hypothetical protein | 100 |
| AVAEIV_RS17065 | 153.8568058 | 2.651783975 | 0.289455093 | 9.161296645 | 5.13E-20 | 3.62E-19 | - | LacI family DNA-binding transcriptional regulator | WP_231876233.1 | Avin_33920 | | LacI family transcriptional regulator protein | 99.72 |
| AVAEIV_RS21855 | 117.5755009 | 2.653323195 | 0.240733077 | 11.02184724 | 3.00E-28 | 5.07E-27 | - | branched-chain amino acid ABC transporter permease | WP_012702865.1 | Avin_43770 | *livH* | Bacterial inner-membrane translocator | 100 |
| AVAEIV_RS16940 | 644.9333719 | 2.656074012 | 0.446653675 | 5.946607314 | 2.74E-09 | 6.61E-09 | - | GMC family oxidoreductase | WP_302226144.1 | Avin_33650 | | dehydrogenase flavoprotein subunit | 99.66 |
| AVAEIV_RS16195 | 189.5987013 | 2.661145747 | 0.457477863 | 5.816993487 | 5.99E-09 | 1.40E-08 | - | aliphatic sulfonate ABC transporter substrate-binding protein | WP_012701715.1 | Avin_31680 | | ABC transporter, aliphatic sulfonate substrate-binding protein | 100 |
| AVAEIV_RS23440 | 24.8629102 | 2.662838978 | 0.432102366 | 6.162518852 | 7.16E-10 | 1.83E-09 | - | EF-hand domain-containing protein | WP_061290265.1 | Avin_46530 | | hypothetical protein | 98.02 |
| AVAEIV_RS23100 | 51.69020977 | 2.667750929 | 0.345793038 | 7.714877494 | 1.21E-14 | 4.89E-14 | - | hypothetical protein | WP_061289593.1 | | | | |
| AVAEIV_RS05885 | 55.31123348 | 2.668171835 | 0.327103066 | 8.156975921 | 3.44E-16 | 1.61E-15 | - | hypothetical protein | WP_302228424.1 | Avin_36290 | | conserved hypothetical protein | 93.94 |
| AVAEIV_RS15975 | 180.5291942 | 2.674311058 | 0.324796577 | 8.233803088 | 1.81E-16 | 8.79E-16 | - | acyl-CoA dehydrogenase family protein | WP_302225905.1 | Avin_31100 | | Acyl-CoA dehydrogenase family protein | 99.5 |
| AVAEIV_RS16465 | 111.7803989 | 2.679517549 | 0.611087571 | 4.384833987 | 1.16E-05 | 2.04E-05 | - | hypothetical protein | WP_156483698.1 | Avin_32540 | | hypothetical protein | 100 |
| AVAEIV_RS11050 | 64.21418313 | 2.681577657 | 0.292591209 | 9.164929013 | 4.96E-20 | 3.50E-19 | - | transporter substrate-binding domain-containing protein | WP_061291066.1 | Avin_21410 | | ABC transporter, substrate binding protein, family 3 | 99.32 |
| AVAEIV_RS13760 | 32.46394003 | 2.686784549 | 0.413546511 | 6.496934398 | 8.20E-11 | 2.27E-10 | - | ACT domain-containing protein | WP_302225359.1 | Avin_26520 | | conserved hypothetical protein | 99.22 |
| AVAEIV_RS00650 | 477.7609919 | 2.687872503 | 0.226653418 | 11.85895418 | 1.93E-32 | 5.18E-31 | *aas* | bifunctional acyl-ACP--phospholipid O-acyltransferase/long-chain-fatty-acid--ACP ligase | WP_302227866.1 | Avin_01340 | | AAS bifunctional protein | 99.72 |
| AVAEIV_RS24990 | 174.4447848 | 2.691362322 | 0.209616808 | 12.8394395 | 9.86E-38 | 4.41E-36 | - | sugar porter family MFS transporter | WP_012703450.1 | Avin_50070 | *iolT* | myo-inositol transporter | 100 |
| AVAEIV_RS25125 | 189.8824258 | 2.694440452 | 0.254424006 | 10.59035466 | 3.30E-26 | 4.61E-25 | *fhuB* | Fe(3+)-hydroxamate ABC transporter permease FhuB | WP_302227668.1 | Avin_50350 | *fhuB* | iron(III)-siderophore ABC transporter permease | 99.7 |
| AVAEIV_RS14935 | 186.6037457 | 2.6969283 | 0.223435869 | 12.07025672 | 1.52E-33 | 4.51E-32 | - | substrate-binding domain-containing protein | WP_231876208.1 | Avin_28890 | | periplasmic phosphate binding protein | 100 |
| AVAEIV_RS17045 | 173.5767615 | 2.701558207 | 0.225027954 | 12.00543382 | 3.33E-33 | 9.72E-32 | - | cupin | WP_302226177.1 | Avin_33880 | | Twin-arginine translocation pathway signal and cupin region containing protein | 99.49 |
| AVAEIV_RS24985 | 97.07611142 | 2.702058746 | 0.278792523 | 9.692005799 | 3.26E-22 | 3.02E-21 | *iolB* | 5-deoxy-glucuronate isomerase | WP_302227652.1 | Avin_50060 | *iolB* | Myo-inositol catabolism protein IolB | 99.27 |
| AVAEIV_RS01280 | 50.23777475 | 2.70416922 | 0.314392294 | 8.601257949 | 7.88E-18 | 4.34E-17 | *vnfG* | V-containing nitrogenase subunit delta | WP_012698949.1 | Avin_02600 | *vnfG* | vanadium nitrogenase, delta subunit, VnfG | 100 |
| AVAEIV_RS02900 | 193.0521445 | 2.704808214 | 0.216548849 | 12.49052221 | 8.41E-36 | 3.09E-34 | - | Bug family tripartite tricarboxylate transporter substrate binding protein | WP_302228101.1 | Avin_05850 | | conserved hypothetical protein | 99.39 |
| AVAEIV_RS10785 | 268.3891857 | 2.704838359 | 0.233142805 | 11.60163773 | 4.04E-31 | 9.12E-30 | *cas6* | CRISPR system precrRNA processing endoribonuclease RAMP protein Cas6 | WP_302228870.1 | | | | |

| | | | | | | | | | | | | | |
|---|---|---|---|---|---|---|---|---|---|---|---|---|---|
| AVAEIV_RS02890 | 10.67126227 | 2.706778452 | 0.602376747 | 4.493497574 | 7.01E-06 | 1.25E-05 | - | hypothetical protein | WP_175555935.1 | | | | |
| AVAEIV_RS15885 | 122.4404641 | 2.706780768 | 0.291924121 | 9.272206622 | 1.82E-20 | 1.38E-19 | - | ABC transporter permease | WP_012701639.1 | Avin_30900 | | ABC transporter, permease protein | 100 |
| AVAEIV_RS21735 | 81.21932467 | 2.709430315 | 0.278271917 | 9.736628649 | 2.10E-22 | 2.00E-21 | - | branched-chain amino acid ABC transporter permease | WP_302227264.1 | Avin_43530 | | amino acid ABC transporter-like protein | 99.66 |
| AVAEIV_RS13845 | 147.862982 | 2.713406355 | 0.221209764 | 12.26621424 | 1.38E-34 | 4.50E-33 | - | serine/threonine protein kinase | WP_012701236.1 | Avin_26690 | | Protein kinase | 100 |
| AVAEIV_RS25120 | 168.7257225 | 2.714087889 | 0.229095119 | 11.84699133 | 2.23E-32 | 5.88E-31 | - | ABC transporter substrate-binding protein | WP_012703476.1 | Avin_50340 | *fhuD* | iron(III)-siderophore-binding periplasmic protein | 100 |
| AVAEIV_RS01850 | 99.80927972 | 2.718790397 | 0.337777081 | 8.049067117 | 8.34E-16 | 3.78E-15 | *fdhE* | formate dehydrogenase accessory protein FdhE | WP_012699072.1 | Avin_03840 | *fdhE* | formate dehydrogenase accessory protein FdhE | 100 |
| AVAEIV_RS21050 | 208.2711285 | 2.723724158 | 0.228008483 | 11.94571412 | 6.84E-33 | 1.93E-31 | - | OprD family porin | WP_302227134.1 | Avin_42090 | | Bacterial outer membrane porin, OprD family | 98.83 |
| AVAEIV_RS25025 | 177.1577097 | 2.72579134 | 0.196914345 | 13.84252296 | 1.41E-43 | 9.81E-42 | - | sugar ABC transporter substrate-binding protein | WP_302227656.1 | Avin_50140 | | Periplasmic substrate-binding protein of sugar ABC transporter | 99.03 |
| AVAEIV_RS21785 | 110.8983763 | 2.726605924 | 0.283180912 | 9.628494763 | 6.06E-22 | 5.50E-21 | - | sugar ABC transporter ATP-binding protein | WP_012702849.1 | Avin_43610 | | ABC transporter | 100 |
| AVAEIV_RS13055 | 32.51441628 | 2.727596395 | 0.433215562 | 6.296164396 | 3.05E-10 | 8.02E-10 | - | metalloregulator ArsR/SmtB family transcription factor | WP_012701162.1 | Avin_25920 | | regulatory protein, ArsR family | 100 |
| AVAEIV_RS01275 | 188.7130376 | 2.728205751 | 0.315736339 | 8.640772093 | 5.58E-18 | 3.10E-17 | *vnfK* | V-containing nitrogenase subunit beta | WP_061288188.1 | Avin_02590 | *vnfK* | vanadium nitrogenase beta subunit, vnfK | 99.79 |
| AVAEIV_RS18360 | 58.82290558 | 2.733549092 | 0.339126953 | 8.060548033 | 7.60E-16 | 3.46E-15 | - | hypothetical protein | WP_061290574.1 | Avin_36980 | | hypothetical protein | 98.59 |
| AVAEIV_RS14040 | 72.31188497 | 2.741478899 | 0.363664449 | 7.53848474 | 4.75E-14 | 1.80E-13 | - | FagA protein | WP_236616141.1 | Avin_27170 | *fagA* | fur associated gene A protein | 100 |
| AVAEIV_RS01270 | 135.4524255 | 2.745688961 | 0.217954922 | 12.59750839 | 2.18E-36 | 8.75E-35 | - | hypothetical protein | WP_302227920.1 | Avin_02580 | | hypothetical protein | 98.67 |
| AVAEIV_RS18260 | 334.9052061 | 2.745861094 | 0.211083247 | 13.00842739 | 1.10E-38 | 5.31E-37 | - | toprim domain-containing protein | WP_302226500.1 | | | | |
| AVAEIV_RS04955 | 178.1294297 | 2.7459041 | 0.209174737 | 13.12732185 | 2.30E-39 | 1.22E-37 | *mmsB* | 3-hydroxyisobutyrate dehydrogenase | WP_302228313.1 | Avin_10700 | | 3-hydroxyisobutyrate dehydrogenase | 99.33 |
| AVAEIV_RS05925 | 181.0397079 | 2.748872333 | 0.270729309 | 10.15358233 | 3.19E-24 | 3.72E-23 | - | TIGR03749 family integrating conjugative element protein | WP_302228429.1 | Avin_36250 | | conserved hypothetical protein | 96.09 |
| AVAEIV_RS13000 | 117.2382862 | 2.750272379 | 0.322096798 | 8.538651712 | 1.36E-17 | 7.30E-17 | - | LysR family transcriptional regulator | WP_012701157.1 | Avin_25870 | | LysR family transcriptional regulator protein | 100 |
| AVAEIV_RS02480 | 164.0739804 | 2.752041906 | 0.194514031 | 14.14829505 | 1.91E-45 | 1.51E-43 | - | hypothetical protein | WP_302228051.1 | | | | |
| AVAEIV_RS23080 | 42.02429315 | 2.75784885 | 0.360437078 | 7.651401641 | 1.99E-14 | 7.84E-14 | - | DUF2597 family protein | WP_061289605.1 | | | | |
| AVAEIV_RS04500 | 74.02178934 | 2.759045774 | 0.326388577 | 8.453254703 | 2.83E-17 | 1.47E-16 | - | HpcH/HpaI aldolase family protein | WP_061290162.1 | Avin_09360 | | 2,4-dihydroxyhept-2-ene-1,7-dioic acid aldolase, HpcH/HpaI family | 99.61 |
| AVAEIV_RS13875 | 25.57105414 | 2.761672414 | 0.477612171 | 5.782248827 | 7.37E-09 | 1.71E-08 | - | TssQ family T6SS-associated lipoprotein | WP_197471077.1 | Avin_26760 | | hypothetical protein | 99.06 |
| AVAEIV_RS00630 | 116.080278 | 2.762413508 | 0.251110731 | 11.00077839 | 3.79E-28 | 6.36E-27 | *modA* | molybdate ABC transporter substrate-binding protein | WP_302227862.1 | Avin_01300 | *modA2* | periplasmic molybdate-binding protein, ModA2 | 99.6 |
| AVAEIV_RS07380 | 40.19306071 | 2.768174022 | 0.403785591 | 6.855554241 | 7.10E-12 | 2.16E-11 | - | acyl-CoA thioesterase | WP_061291849.1 | Avin_14220 | | 4-hydroxybenzoyl CoA thioesterase | 99.31 |
| AVAEIV_RS23150 | 165.190056 | 2.769438195 | 0.210270881 | 13.17081174 | 1.29E-39 | 7.14E-38 | - | hypothetical protein | WP_302227517.1 | | | | |
| AVAEIV_RS09575 | 89.24505747 | 2.776463985 | 0.315704482 | 8.794502912 | 1.44E-18 | 8.59E-18 | - | ABC transporter permease | WP_302228697.1 | Avin_18690 | | ABC transporter, inner membrane permease component | 99.67 |
| AVAEIV_RS23090 | 34.76428826 | 2.780360991 | 0.415256649 | 6.695524317 | 2.15E-11 | 6.27E-11 | - | D-Ala-D-Ala carboxypeptidase family metallohydrolase | WP_061289599.1 | | | | |
| AVAEIV_RS20475 | 167.1077787 | 2.781909055 | 0.239645723 | 11.60842357 | 3.73E-31 | 8.53E-30 | - | sigma-70 family RNA polymerase sigma factor | WP_012702602.1 | Avin_40940 | | RNA polymerase sigma factor, FecI family | 100 |
| AVAEIV_RS25300 | 84.06457941 | 2.790941592 | 0.364510588 | 7.656681812 | 1.91E-14 | 7.54E-14 | - | molybdate ABC transporter permease subunit | WP_012703513.1 | Avin_50720 | | ABC transporter, permease protein | 100 |
| AVAEIV_RS02515 | 80.75060761 | 2.79177544 | 0.326768277 | 8.543593856 | 1.30E-17 | 7.02E-17 | *tolQ* | protein TolQ | WP_012699180.1 | Avin_04980 | | TolQ-like protein | 100 |
| AVAEIV_RS20355 | 57.54119931 | 2.792379735 | 0.399677386 | 6.986584262 | 2.82E-12 | 8.93E-12 | - | toprim domain-containing protein | WP_367281683.1 | Avin_40670 | | DnaG-like primase protein | 99.22 |
| AVAEIV_RS19420 | 46.05729535 | 2.797616298 | 0.386515638 | 7.238041677 | 4.55E-13 | 1.57E-12 | - | hypothetical protein | WP_302226772.1 | | | | |
| AVAEIV_RS15865 | 597.4540231 | 2.802476342 | 0.247798587 | 11.30949282 | 1.18E-29 | 2.31E-28 | - | TonB-dependent siderophore receptor | WP_302225869.1 | Avin_30860 | | TonB-dependent siderophore receptor | 99.86 |
| AVAEIV_RS02175 | 628.7777879 | 2.821357195 | 0.268601211 | 10.50388857 | 8.29E-26 | 1.11E-24 | - | methyltransferase | WP_302227998.1 | Avin_04300 | | Conserved Hypothetical Protein | 99.76 |
| AVAEIV_RS15490 | 351.9396748 | 2.831046755 | 0.429435065 | 6.592490885 | 4.33E-11 | 1.23E-10 | - | DegT/DnrJ/EryC1/StrS family aminotransferase | WP_012701560.1 | Avin_30070 | | DegT/DnrJ/EryC1/StrS aminotransferase family protein | 100 |
| AVAEIV_RS04190 | 191.5233557 | 2.832992999 | 0.315084541 | 8.991215461 | 2.45E-19 | 1.60E-18 | - | MFS transporter | WP_302228230.1 | Avin_08630 | | benzoate transporter | 99.77 |
| AVAEIV_RS04175 | 75.55829754 | 2.83506445 | 0.277041576 | 10.23335373 | 1.41E-24 | 1.72E-23 | *benB* | benzoate 1,2C2-dioxygenase small subunit | WP_061290612.1 | Avin_08600 | *benB* | benzoate-1,2-dioxygenase oxygenase component, beta subunit | 98.77 |
| AVAEIV_RS16150 | 191.479103 | 2.843460114 | 0.229011722 | 12.41622084 | 2.13E-35 | 7.41E-34 | - | SfnB family sulfur acquisition oxidoreductase | WP_302225964.1 | Avin_31480 | | Acyl-CoA dehydrogenase family protein | 99.53 |
| AVAEIV_RS05265 | 70.92330362 | 2.853339866 | 0.286376657 | 9.963590956 | 2.20E-23 | 2.35E-22 | - | acetone carboxylase subunit gamma | WP_175555958.1 | Avin_11400 | *apc2* | acetophenone carboxylase gamma subunit | 100 |
| AVAEIV_RS15445 | 127.5918536 | 2.85464355 | 0.288218773 | 9.904433083 | 3.98E-23 | 4.10E-22 | - | glycosyltransferase family 2 protein | WP_012701551.1 | Avin_29980 | | Glycosyl transferase, family 2 protein | 100 |
| AVAEIV_RS16110 | 110.1430525 | 2.862205022 | 0.308401547 | 9.280773884 | 1.68E-20 | 1.28E-19 | - | ABC transporter substrate-binding protein | WP_302225949.1 | Avin_31380 | | ABC nitrate/sulfonate/bicarbonate family transporter, periplasmic ligand binding protein | 99.69 |
| AVAEIV_RS25480 | 252.7935125 | 2.865370753 | 0.26658103 | 10.74859209 | 6.02E-27 | 8.91E-26 | *norR* | nitric oxide reductase transcriptional regulator NorR | pseudo=true | | | | |
| AVAEIV_RS10395 | 212.9431556 | 2.866681952 | 0.243774913 | 11.75954457 | 6.31E-32 | 1.60E-30 | - | MFS transporter | WP_415860977.1 | Avin_20360 | | major facilitator superfamily (MFS) permease | 96.63 |
| AVAEIV_RS07390 | 121.298594 | 2.867328676 | 0.300099754 | 9.554585236 | 1.24E-21 | 1.08E-20 | - | aldehyde dehydrogenase | WP_302228503.1 | Avin_14240 | *vdh* | vanillin:NAD+ oxidoreductase | 99.79 |
| AVAEIV_RS03325 | 76.71207995 | 2.87620615 | 0.284754236 | 10.1006615 | 5.49E-24 | 6.21E-23 | - | hypothetical protein | WP_012699345.1 | Avin_06690 | | conserved hypothetical protein | 100 |
| AVAEIV_RS25030 | 244.0435111 | 2.879988592 | 0.236399523 | 12.18271742 | 3.84E-34 | 1.22E-32 | - | Gfo/Idh/MocA family protein | WP_302227658.1 | Avin_50150 | | oxidoreductase family, NAD-binding Rossmann fold domain protein | 99.42 |
| AVAEIV_RS05270 | 173.6430619 | 2.891723126 | 0.232497664 | 12.4376438 | 1.63E-35 | 5.75E-34 | - | hydantoinase/oxoprolinase family protein | WP_302228343.1 | Avin_11410 | *apc1* | acetophenone carboxylase | 98.92 |
| AVAEIV_RS02465 | 265.036224 | 2.898396534 | 0.261201943 | 11.09829621 | 1.28E-28 | 2.25E-27 | - | TolC family protein | WP_367281689.1 | | | | |
| AVAEIV_RS00100 | 74.75901592 | 2.902353842 | 0.325118027 | 8.927077562 | 4.37E-19 | 2.78E-18 | *dprA* | DNA-processing protein DprA | WP_061290474.1 | Avin_00190 | *dprA* | DNA processing protein, DprA (SMF family) | 99.45 |
| AVAEIV_RS21905 | 145.7085818 | 2.909944864 | 0.250567218 | 11.61343006 | 3.52E-31 | 8.08E-30 | - | sigma-54 interaction domain-containing protein | WP_012702878.1 | Avin_43900 | *sfnR* | Sigma54-dependent transcriptional activator protein | 100 |
| AVAEIV_RS10955 | 51.05915072 | 2.91219293 | 0.369359136 | 7.884448077 | 3.16E-15 | 1.34E-14 | - | isochorismatase | WP_175556018.1 | Avin_21200 | *entB* | Isochorismatase (2,3 dihydro-2,3 dihydroxybenzoate synthase) | 100 |
| AVAEIV_RS16005 | 34.36015345 | 2.919844802 | 0.417488348 | 6.993835433 | 2.67E-12 | 8.52E-12 | - | hypothetical protein | WP_302225915.1 | Avin_31160 | | hypothetical protein | 96.43 |
| AVAEIV_RS21795 | 142.6800077 | 2.921023613 | 0.244299289 | 11.95674217 | 5.99E-33 | 1.70E-31 | - | sugar ABC transporter substrate-binding protein | WP_302227282.1 | Avin_43630 | | periplasmic sugar binding protein | 99.09 |
| AVAEIV_RS15840 | 128.4542504 | 2.923395058 | 0.2921236 | 10.00739091 | 1.41E-23 | 1.54E-22 | - | catechol 2%2C3-dioxygenase | WP_302225858.1 | Avin_30770 | *lapB* | catechol 2,3-dioxygenase, LapB | 99.35 |
| AVAEIV_RS18305 | 108.5812502 | 2.923418537 | 0.287655308 | 10.16292228 | 2.90E-24 | 3.41E-23 | - | ORF6N domain-containing protein | WP_302226517.1 | | | | |
| AVAEIV_RS15980 | 134.7097001 | 2.926389066 | 0.321109716 | 9.113361944 | 7.99E-20 | 5.52E-19 | - | ABC transporter substrate-binding protein | WP_302225907.1 | Avin_31110 | | ABC transporter substrate binding protein | 98.06 |
| AVAEIV_RS11235 | 186.3551092 | 2.930519508 | 0.25818816 | 11.35032492 | 7.39E-30 | 1.48E-28 | - | ABC transporter substrate-binding protein | WP_302228957.1 | Avin_21970 | | ABC transporter, taurine periplasmic binding protein | 99.39 |
| AVAEIV_RS13205 | 144.1293551 | 2.932056581 | 0.279779459 | 10.47988509 | 1.07E-25 | 1.42E-24 | - | RES family NAD+ phosphorylase | WP_012701195.1 | Avin_26280 | | conserved hypothetical protein | 100 |
| AVAEIV_RS05660 | 281.8721165 | 2.932819408 | 0.64378042 | 4.5556207 | 5.22E-06 | 9.42E-06 | - | TIGR03757 family integrating conjugative element protein | WP_302229280.1 | Avin_36340 | | conserved hypothetical protein | 98.35 |
| AVAEIV_RS10330 | 57.74254543 | 2.936730615 | 0.347718423 | 8.445714765 | 3.02E-17 | 1.56E-16 | - | shikimate dehydrogenase family protein | WP_175555915.1 | Avin_20210 | | shikimate 5-dehydrogenase | 100 |
| AVAEIV_RS10985 | 106.8811133 | 2.937537109 | 0.313651727 | 9.365601576 | 7.56E-21 | 5.93E-20 | - | MmgE/PrpD family protein | WP_302228917.1 | Avin_21260 | | MmgE/PrpD family protein | 99.12 |
| AVAEIV_RS20900 | 145.0292792 | 2.943852865 | 0.237153621 | 12.41327396 | 2.21E-35 | 7.64E-34 | - | ATP-NAD kinase family protein | WP_012702674.1 | Avin_41720 | | ATP-NAD AcoX kinase | 100 |
| AVAEIV_RS11045 | 121.8794253 | 2.945785382 | 0.252117381 | 11.68418206 | 1.54E-31 | 3.78E-30 | - | LysR substrate-binding domain-containing protein | WP_201765945.1 | Avin_21400 | | Transcriptional regulator, LysR family | 100 |
| AVAEIV_RS07815 | 195.5218269 | 2.945974251 | 0.254198572 | 11.58926357 | 4.67E-31 | 1.04E-29 | - | OprD family porin | WP_302228530.1 | | | | |
| AVAEIV_RS02905 | 117.5620279 | 2.949574352 | 0.271389127 | 10.86843231 | 1.63E-27 | 2.55E-26 | - | acyltransferase | WP_302229243.1 | Avin_05860 | | Phospholipid/glycerol acyltransferase | 100 |
| AVAEIV_RS20015 | 90.87231429 | 2.950101396 | 0.305564522 | 9.654593969 | 4.70E-22 | 4.31E-21 | - | SDR family oxidoreductase | WP_302226954.1 | Avin_39930 | | conserved hypothetical protein | 100 |
| AVAEIV_RS25930 | 311.391024 | 2.959563534 | 0.205982366 | 14.36804323 | 8.21E-47 | 6.92E-45 | - | TonB-dependent receptor plug domain-containing protein | WP_302227783.1 | Avin_52120 | | TonB-dependent receptor | 99.85 |
| AVAEIV_RS12825 | 1200.536814 | 2.962196241 | 0.415882585 | 7.122674404 | 1.06E-12 | 3.53E-12 | - | TonB-dependent siderophore receptor | WP_302229188.1 | Avin_25550 | | TonB-dependent siderophore receptor/ferripyoverdine receptor | 91.95 |

| | | | | | | | | | | | | | |
|---|---|---|---|---|---|---|---|---|---|---|---|---|---|
| AVAEIV_RS16095 | 107.3474203 | 2.963941565 | 0.317346385 | 9.339767863 | 9.65E-21 | 7.44E-20 | - | acyl-CoA dehydrogenase family protein | WP_302225945.1 | Avin_31350 | | DszC-like desulfurization enzyme | 99.75 |
| AVAEIV_RS05915 | 27.52247827 | 2.986144752 | 0.467170578 | 6.391979482 | 1.64E-10 | 4.42E-10 | - | hypothetical protein | WP_254796286.1 | | | | |
| AVAEIV_RS10775 | 98.47227274 | 2.989485669 | 0.310555747 | 9.626244891 | 6.20E-22 | 5.61E-21 | *cas2* | CRISPR-associated endonuclease Cas2 | WP_302228868.1 | | | | |
| AVAEIV_RS23060 | 35.01591829 | 2.989906361 | 0.413604819 | 7.228896346 | 4.87E-13 | 1.68E-12 | - | head completion/stabilization protein | WP_302227507.1 | | | | |
| AVAEIV_RS13765 | 109.6536232 | 2.995569827 | 0.309208306 | 9.687869854 | 3.40E-22 | 3.14E-21 | - | 2OG-Fe dioxygenase family protein | WP_302225361.1 | Avin_26530 | | Conserved hypothetical protein | 99.66 |
| AVAEIV_RS23125 | 320.1969516 | 3.002685574 | 0.210476986 | 14.26609923 | 3.56E-46 | 2.90E-44 | - | phage tail tape measure protein | WP_302227515.1 | | | | |
| AVAEIV_RS24765 | 194.3515149 | 3.008873511 | 0.206882485 | 14.54387745 | 6.39E-48 | 6.31E-46 | - | OprD family porin | WP_302227640.1 | Avin_49310 | | Bacterial outer membrane porin | 99.52 |
| AVAEIV_RS07400 | 79.47644787 | 3.010719925 | 0.291236904 | 10.33770062 | 4.76E-25 | 6.00E-24 | - | thiolase family protein | WP_302228505.1 | Avin_14260 | *aat1* | beta-ketothiolase | 99.52 |
| AVAEIV_RS13850 | 39.35709368 | 3.018701001 | 0.389641367 | 7.747383253 | 9.38E-15 | 3.84E-14 | *tagF* | type VI secretion system-associated protein TagF | WP_061290225.1 | Avin_26700 | | conserved hypothetical protein | 99.57 |
| AVAEIV_RS11240 | 187.4088003 | 3.019806984 | 0.647906243 | 4.660870327 | 3.15E-06 | 5.81E-06 | - | ABC transporter ATP-binding protein | WP_302228960.1 | Avin_21980 | | ABC transporter, ATP binding component | 99.63 |
| AVAEIV_RS11425 | 154.1542133 | 3.033273715 | 0.312812179 | 9.696789056 | 3.11E-22 | 2.89E-21 | - | LLM class flavin-dependent oxidoreductase | WP_012700829.1 | Avin_22400 | *dszA* | xenobiotic compound monooxygenase, DszA family, A subunit protein | 100 |
| AVAEIV_RS11055 | 75.5438656 | 3.035319224 | 0.374280722 | 8.109739674 | 5.07E-16 | 2.34E-15 | - | Ldh family oxidoreductase | WP_012700742.1 | Avin_21420 | *vbiC* | Malate/L-lactate dehydrogenase | 100 |
| AVAEIV_RS10770 | 45.37573728 | 3.039887208 | 0.364185787 | 8.347078104 | 7.00E-17 | 3.53E-16 | - | hypothetical protein | WP_302228866.1 | | | | |
| AVAEIV_RS11020 | 151.5023027 | 3.059053185 | 0.290327741 | 10.53655146 | 5.86E-26 | 8.01E-25 | - | FAD-dependent oxidoreductase | WP_012700736.1 | Avin_21320 | | fumarate reductase/succinate dehyfrogenase, flavoprotein subunit | 100 |
| AVAEIV_RS10980 | 114.9461746 | 3.071047482 | 0.273403772 | 11.2326449 | 2.82E-29 | 5.39E-28 | *purB* | adenylosuccinate lyase | WP_061291346.1 | Avin_21250 | | adenylosuccinate lyase | 99.78 |
| AVAEIV_RS11065 | 102.4519529 | 3.083575689 | 0.333110905 | 9.256904048 | 2.10E-20 | 1.57E-19 | - | aminotransferase class I/II-fold pyridoxal phosphate-dependent enzyme | WP_061291060.1 | Avin_21440 | | aspartate aminotransferase | 99.01 |
| AVAEIV_RS20935 | 140.8345031 | 3.083810072 | 0.326136784 | 9.455572696 | 3.21E-21 | 2.68E-20 | - | STAS domain-containing protein | pseudo=true | | | | |
| AVAEIV_RS04490 | 86.46186371 | 3.085782287 | 0.309741621 | 9.962439902 | 2.23E-23 | 2.37E-22 | - | siderophore biosynthesis protein PvsA | WP_012699598.1 | Avin_09340 | | conserved hypothetical protein | 100 |
| AVAEIV_RS15475 | 103.6009184 | 3.087501375 | 0.275154855 | 11.22095911 | 3.22E-29 | 6.08E-28 | - | glycosyltransferase family 4 protein | WP_012701557.1 | Avin_30040 | | Glycosyl transferase, group 1 family protein | 100 |
| AVAEIV_RS20340 | 24.66338931 | 3.097192091 | 0.501290349 | 6.178439495 | 6.47E-10 | 1.66E-09 | - | hypothetical protein | WP_012702570.1 | Avin_40620 | | hypothetical protein | 100 |
| AVAEIV_RS23375 | 159.7536535 | 3.113567777 | 0.288283396 | 10.80037152 | 3.43E-27 | 5.20E-26 | - | MFS transporter | WP_015589523.1 | Avin_46380 | | major facilitator superfamily protein | 99.61 |
| AVAEIV_RS21840 | 138.7656803 | 3.116451277 | 0.237369427 | 13.12911827 | 2.24E-39 | 1.20E-37 | - | LLM class flavin-dependent oxidoreductase | WP_061291802.1 | Avin_43740 | | monooxygenase, NtaA/SnaA/SoxA/DszA family | 99.12 |
| AVAEIV_RS10765 | 167.6293283 | 3.119468934 | 0.239652637 | 13.01662679 | 9.84E-39 | 4.82E-37 | *cas1* | CRISPR-associated endonuclease Cas1 | WP_302228864.1 | | | | |
| AVAEIV_RS17795 | 36.20951995 | 3.136737194 | 0.457868648 | 6.850735919 | 7.35E-12 | 2.23E-11 | - | hypothetical protein | WP_156483618.1 | Avin_35460 | | hypothetical protein | 100 |
| AVAEIV_RS11060 | 117.3927381 | 3.141278348 | 0.30330948 | 10.35667713 | 3.90E-25 | 4.97E-24 | - | FAD-binding oxidoreductase | WP_012700743.1 | Avin_21430 | | FAD linked oxidoreductase | 100 |
| AVAEIV_RS15890 | 94.93127368 | 3.155358213 | 0.336095431 | 9.388280587 | 6.10E-21 | 4.85E-20 | - | ABC transporter ATP-binding protein | WP_302225880.1 | Avin_30910 | | ABC transporter, ATP-binding protein | 99.64 |
| AVAEIV_RS04255 | 118.6938318 | 3.16032895 | 0.538040243 | 5.873778011 | 4.26E-09 | 1.01E-08 | - | transporter | WP_415860985.1 | Avin_08780 | | Conserved hypothetical protein | 99.63 |
| AVAEIV_RS13065 | 238.6271346 | 3.169914816 | 0.297125458 | 10.66860725 | 1.43E-26 | 2.06E-25 | - | sigma-54-dependent transcriptional regulator | WP_012701164.1 | Avin_25940 | | sigma54-dependent activator protein | 100 |
| AVAEIV_RS23740 | 163.558703 | 3.170114949 | 0.279967843 | 11.32313953 | 1.01E-29 | 1.99E-28 | - | ABC transporter ATP-binding protein | WP_302227582.1 | Avin_47160 | | ABC transporter | 99.62 |
| AVAEIV_RS14820 | 67.71741087 | 3.173997523 | 0.357738667 | 8.87239155 | 7.16E-19 | 4.43E-18 | *gspG* | type II secretion system major pseudopilin GspG | WP_012701421.1 | Avin_28620 | *gspG1* | General secretion pathway protein G | 100 |
| AVAEIV_RS07375 | 201.6061277 | 3.185056793 | 0.246099777 | 12.94213602 | 2.60E-38 | 1.24E-36 | - | OprD family porin | WP_302228502.1 | Avin_14210 | | Bacterial outer membrane porin | 98.82 |
| AVAEIV_RS11075 | 63.61385222 | 3.185647019 | 0.410817155 | 7.75441575 | 8.88E-15 | 3.64E-14 | - | amino acid ABC transporter permease | WP_012700746.1 | Avin_21460 | | polar amino acid ABC transporter, inner membrane permease component | 100 |
| AVAEIV_RS09020 | 139.3461066 | 3.193241612 | 0.253962819 | 12.57365792 | 2.95E-36 | 1.14E-34 | - | PhnD/SsuA/transferrin family substrate-binding protein | WP_302228632.1 | Avin_17530 | | Phosphonate ABC transporter, periplasmic phosphate-binding protein | 99.64 |
| AVAEIV_RS13200 | 87.31188562 | 3.197652985 | 0.289734737 | 11.03648467 | 2.55E-28 | 4.36E-27 | - | MbcA/ParS/Xre antitoxin family protein | WP_302225194.1 | Avin_26270 | | conserved hypothetical protein | 99.12 |
| AVAEIV_RS10385 | 343.0677173 | 3.259620039 | 0.456560336 | 7.139516471 | 9.37E-13 | 3.14E-12 | - | FAD-dependent oxidoreductase | WP_012700647.1 | Avin_20340 | | Fumarate reductase/succinate dehydrogenase related flavoprotein | 100 |
| AVAEIV_RS13870 | 50.68358449 | 3.276871615 | 0.369129168 | 8.877303392 | 6.85E-19 | 4.25E-18 | *tssJ* | type VI secretion system lipoprotein TssJ | WP_061290232.1 | Avin_26740 | | conserved hypothetical protein | 98.96 |
| AVAEIV_RS09110 | 50.65222405 | 3.279592488 | 0.411993483 | 7.960301854 | 1.72E-15 | 7.50E-15 | - | DUF2790 domain-containing protein | WP_231876284.1 | Avin_17720 | | conserved hypothetical protein | 100 |
| AVAEIV_RS11230 | 151.1286538 | 3.281444252 | 0.311768362 | 10.52526378 | 6.61E-26 | 8.96E-25 | - | TauD/TfdA dioxygenase family protein | WP_012700786.1 | Avin_21960 | | Taurine catabolism dioxygenase TauD/TfdA family | 100 |
| AVAEIV_RS08745 | 69.45322233 | 3.298849999 | 0.364372416 | 9.053511894 | 1.38E-19 | 9.32E-19 | - | helix-turn-helix transcriptional regulator | WP_062832573.1 | | | | |
| AVAEIV_RS03960 | 203.2615534 | 3.317824547 | 0.302289613 | 10.97564853 | 5.00E-28 | 8.30E-27 | - | energy transducer TonB | WP_302228203.1 | Avin_08080 | *tonB* | TonB protein | 98.55 |
| AVAEIV_RS21875 | 149.1829589 | 3.332248555 | 0.244742022 | 13.61535111 | 3.25E-42 | 2.09E-40 | - | LLM class flavin-dependent oxidoreductase | WP_302227305.1 | Avin_43810 | *seuA* | monoxygenase, NtaA/DszA family | 99.79 |
| AVAEIV_RS23055 | 93.08704269 | 3.333717442 | 0.308928447 | 10.79122842 | 3.79E-27 | 5.73E-26 | *gpM* | phage terminase small subunit | WP_302227505.1 | | | | |
| AVAEIV_RS13860 | 102.8411293 | 3.354717351 | 0.339203446 | 9.889986054 | 4.60E-23 | 4.70E-22 | - | DotU family type VI secretion system protein | WP_302225389.1 | Avin_26720 | | OmpA/MotB domain protein | 99.77 |
| AVAEIV_RS23380 | 86.53397278 | 3.356451589 | 0.30921201 | 10.8548552 | 1.89E-27 | 2.93E-26 | - | HlyD family secretion protein | WP_061290280.1 | Avin_46390 | | secretion protein | 99.44 |
| AVAEIV_RS05310 | 59.4333061 | 3.386927674 | 0.425116217 | 7.967062987 | 1.62E-15 | 7.13E-15 | - | LysR family transcriptional regulator | WP_302228354.1 | Avin_11490 | | Transcriptional regulator, LysR family | 99.01 |
| AVAEIV_RS04275 | 69.73475922 | 3.393619078 | 0.35747199 | 9.493384576 | 2.24E-21 | 1.90E-20 | - | phenol hydroxylase subunit P4 | WP_012699548.1 | Avin_08820 | *dmpO* | Phenol hydroxylase subunit P4 | 100 |
| AVAEIV_RS10755 | 55.45755826 | 3.394666261 | 0.387769891 | 8.754331722 | 2.05E-18 | 1.20E-17 | *cas2* | CRISPR-associated endonuclease Cas2 | WP_302228861.1 | | | | |
| AVAEIV_RS20350 | 60.24546166 | 3.425252819 | 0.363808382 | 9.414991474 | 4.73E-21 | 3.84E-20 | - | hypothetical protein | WP_302226997.1 | Avin_40650 | | hypothetical protein | 99.15 |
| AVAEIV_RS06490 | 83.88709329 | 3.426799042 | 0.337756434 | 10.14576984 | 3.46E-24 | 3.99E-23 | - | PepSY-associated TM helix domain-containing protein | WP_302228472.1 | Avin_12400 | | PepSY-associated TM helix | 97.49 |
| AVAEIV_RS23130 | 22.0429716 | 3.432548403 | 0.536585287 | 6.397022969 | 1.58E-10 | 4.28E-10 | - | DUF2590 family protein | WP_061289582.1 | | | | |
| AVAEIV_RS11180 | 65.65555181 | 3.446412631 | 0.384607892 | 8.960847408 | 3.22E-19 | 2.07E-18 | - | hypothetical protein | WP_012700776.1 | Avin_21860 | | hypothetical protein | 100 |
| AVAEIV_RS17960 | 505.3382301 | 3.458077213 | 0.814192667 | 4.247246819 | 2.16E-05 | 3.70E-05 | - | TIGR03757 family integrating conjugative element protein | WP_302229394.1 | | | | |
| AVAEIV_RS19930 | 66.71830892 | 3.458227795 | 0.34161655 | 10.12312721 | 4.36E-24 | 5.00E-23 | - | DMT family transporter | WP_302226931.1 | Avin_39760 | | Conserved Hypothetical Protein | 99.66 |
| AVAEIV_RS11170 | 172.3565037 | 3.477088429 | 0.318866518 | 10.90452659 | 1.10E-27 | 1.75E-26 | - | energy transducer TonB | WP_302228945.1 | Avin_21840 | | TonB C-terminal domain protein, ferric siderophore transporter | 95.8 |
| AVAEIV_RS08415 | 103.0827542 | 3.486429455 | 0.308766442 | 11.29147789 | 1.45E-29 | 2.82E-28 | - | electron transfer flavoprotein subunit alpha/FixB family protein | WP_012700272.1 | Avin_16420 | *etfA3* | Electron transfer flavoprotein, alpha subunit | 100 |
| AVAEIV_RS24090 | 217.2712204 | 3.508139454 | 0.269066164 | 13.03820372 | 7.42E-39 | 3.74E-37 | - | DoxX family protein | WP_012703250.1 | Avin_47840 | | DoxX-like protein | 100 |
| AVAEIV_RS11080 | 57.76819152 | 3.515903546 | 0.414636312 | 8.479487791 | 2.26E-17 | 1.18E-16 | - | amino acid ABC transporter ATP-binding protein | WP_302228929.1 | Avin_21470 | | amino acid ABC transporter, ATP binding component | 99.63 |
| AVAEIV_RS23070 | 115.7807611 | 3.545561053 | 0.284165136 | 12.47711491 | 9.95E-36 | 3.63E-34 | - | phage virion morphogenesis protein | WP_302227508.1 | | | | |
| AVAEIV_RS18265 | 21.81731028 | 3.551929427 | 0.597095794 | 5.948676011 | 2.70E-09 | 6.53E-09 | - | hypothetical protein | WP_302226502.1 | | | | |
| AVAEIV_RS16090 | 214.8535769 | 3.565370663 | 0.240285478 | 14.83806134 | 8.31E-50 | 9.71E-48 | - | LLM class flavin-dependent oxidoreductase | WP_012701682.1 | Avin_31340 | *dszA* | dibenzothiophene sulfone monooxygenase | 100 |
| AVAEIV_RS24745 | 306.5653811 | 3.571148461 | 0.278896212 | 12.80457857 | 1.55E-37 | 6.68E-36 | - | MFS transporter | WP_302227637.1 | Avin_49260 | | major facilitator family transporter | 99.77 |
| AVAEIV_RS25115 | 59.73765472 | 3.599571715 | 0.39512047 | 9.110061339 | 8.23E-20 | 5.68E-19 | - | ABC transporter ATP-binding protein | WP_012703475.1 | Avin_50330 | *fhuC* | iron(III)-siderophore ABC transporter ATP-binding protein | 100 |
| AVAEIV_RS11195 | 160.556411 | 3.639838298 | 0.303606556 | 11.98866833 | 4.07E-33 | 1.18E-31 | - | LLM class flavin-dependent oxidoreductase | WP_302228949.1 | Avin_21890 | | monooxygenase, NtaA/SnaA/SoxA family | 99.55 |
| AVAEIV_RS20335 | 161.2215024 | 3.646702082 | 0.293129166 | 12.4405979 | 1.57E-35 | 5.62E-34 | - | hypothetical protein | WP_012702569.1 | Avin_40610 | | hypothetical protein | 100 |
| AVAEIV_RS04485 | 188.9124818 | 3.693026505 | 0.314755559 | 11.7329985 | 8.63E-32 | 2.15E-30 | - | IucA/IucC family protein | WP_302228260.1 | Avin_09330 | | siderophore biosynthesis protein, IucA/IucC family | 98.47 |
| AVAEIV_RS15830 | 193.9333413 | 3.721192055 | 0.275887806 | 13.4880628 | 1.84E-41 | 1.14E-39 | - | aromatic/alkene monooxygenase hydroxylase subunit beta | WP_012701626.1 | Avin_30750 | *lapL* | Multi-component phenol hydoxylase, beta subunit; LapL | 100 |
| AVAEIV_RS20895 | 107.7274442 | 3.739737244 | 0.588512792 | 6.354555579 | 2.09E-10 | 5.57E-10 | - | SDR family NAD(P)-dependent oxidoreductase | WP_012702673.1 | Avin_41710 | | Short-chain dehydrogenase/reductase SDR | 100 |
| AVAEIV_RS24760 | 253.7411194 | 3.760872334 | 0.249431942 | 15.0777495 | 2.27E-51 | 3.07E-49 | - | OprD family porin | WP_302227639.1 | Avin_49300 | | OprD family outer membrane porin | 97.67 |
| AVAEIV_RS15985 | 119.1569155 | 3.78006639 | 0.352788573 | 10.71482094 | 8.67E-27 | 1.27E-25 | - | ABC transporter permease | WP_302225909.1 | Avin_31120 | | ABC transporter permease protein | 99.4 |

| | | | | | | | | | | | | | |
|---|---|---|---|---|---|---|---|---|---|---|---|---|---|
| AVAEIV_RS24840 | 61.51588609 | 3.949873485 | 0.655291273 | 6.027660751 | 1.66E-09 | 4.10E-09 | - | tripartite tricarboxylate transporter TctB family protein | WP_012703404.1 | Avin_49470 | *tctB* | tripartite tricarboxylate transporter small transmembrane protein | 100 |
| AVAEIV_RS23745 | 57.5119068 | 4.034709019 | 0.409822237 | 9.845022182 | 7.20E-23 | 7.18E-22 | - | sigma-70 family RNA polymerase sigma factor | WP_061288955.1 | Avin_47170 | | RNA polymerase sigma factor, FecI family | 99.38 |
| AVAEIV_RS21865 | 62.89525784 | 4.069553097 | 0.432713273 | 9.404733694 | 5.22E-21 | 4.19E-20 | - | AMP-binding protein | WP_012702867.1 | Avin_43790 | | conserved hypothetical protein | 100 |
| AVAEIV_RS23085 | 11.02418553 | 4.697025646 | 1.078215367 | 4.356296329 | 1.32E-05 | 2.31E-05 | - | TraR/DksA family transcriptional regulator | WP_302227511.1 | | | | |
| AVAEIV_RS12890 | 224.5923002 | 4.751081223 | 0.724454766 | 6.558147519 | 5.45E-11 | 1.53E-10 | - | MbtH family protein | WP_012701134.1 | Avin_25630 | | MbtH-like protein | 100 |
